# Epithelial cells maintain memory of prior infection with *Streptococcus pneumoniae* through di-methylation of histone H3

**DOI:** 10.1101/2023.05.19.541441

**Authors:** Christine Chevalier, Claudia Chica, Justine Matheau, Michael G. Connor, Adrien Pain, Melanie A. Hamon

## Abstract

Epithelial cells are the first point of contact for bacteria entering the respiratory tract. *Streptococcus pneumoniae* is an obligate human pathobiont of the nasal mucosa, carried asymptomatically but also the cause of severe pneumonia. The role of the epithelium in maintaining homeostatic interactions or mounting an inflammatory response to invasive *S. pneumoniae* is currently poorly understood. However, studies have shown that chromatin modifications, at the histone level, induced by bacterial pathogens interfere with the host transcriptional program and promote infection. In this study, we demonstrate that *S. pneumoniae* actively induces di-methylation of lysine 4 on histone H3 (H3K4me2), which persists for at least 9 days upon clearance of bacteria with antibiotics. We show that infection establishes a unique epigenetic program affecting the transcriptional response of epithelial cells, rendering them more permissive upon secondary infection. Our results establish H3K4me2 as a unique modification induced by infection, distinct from H3K4me3, which localizes to enhancer regions genome-wide. Therefore, this study reveals evidence that bacterial infection leaves a memory in epithelial cells after bacterial clearance, in an epigenomic mark, thereby altering cellular responses for subsequent infection.

## Introduction

The external and internal surfaces of the body are covered with epithelia, which form a barrier to the environment. As such, epithelial barriers and epithelial cells are the primary responders to environmental assault and constitute the first line of defense against invading pathogens. They play an important role in controlling the initial steps in the inflammatory response upon contact with virulent pathogens, while maintaining tissue homeostasis under resting state conditions (1, 2). Therefore, a tight regulation of epithelial cell responses is paramount to ensure homoeostatic maintenance even under contact with commensal bacteria.

Pathogenic bacteria are sensed by the epithelia, mainly through pattern recognition receptors, but also have evolved mechanisms to subvert host responses to promote infection (3). One of these strategies involves reprogramming of host transcription, effectively downregulating host inflammatory responses or promoting cell survival (4). There are multiple levels at which bacteria can modulate host transcription; at the level of signal transduction, transcription factor regulation, or even at the level of chromatin organization. Indeed, eukaryotic DNA is organized in a complex structure combining DNA and associated proteins into chromatin. The primary unit of chromatin, the nucleosome, is composed of an octamer of 2 copies of each core histone H2A, H2B, H3 and H4, around which DNA is wrapped (5). The position of nucleosomes as well as the extent to which DNA is wrapped plays an essential role in accessibility of genetic regions to transcription factor binding and transcription machinery, and thereby can regulate transcription. The chromatin structure is dynamically regulated at the nucleosome level by post-translational modifications of histone proteins as well as remodeling enzymes that reposition nucleosomes along the genome. Interestingly, bacterial pathogens have been shown to actively induce chromatin modifications as a potent transcriptional reprograming strategy (6).

*Streptococcus pneumoniae* is one example of a bacterium that actively drives chromatin remodeling (7, 8). *S. pneumoniae* (or *pneumococcus*) is a natural colonizer of the human upper respiratory tract, which is carried asymptomatically in a large proportion of the population (9–11). However, it is the cause of the largest number of community acquired bacterial pneumonia (12), making it a deadly pathogen. Considering that vaccine evasion is prominent and that antibiotic resistance is continuously increasing, the *pneumococcus* is on the WHO list of priority pathogens (13). This bacterium has long been defined by its capsule, which has been used to classify *S. pneumoniae* into different serotypes, and against which the vaccine is directed. However, genome-wide analysis of over 2 000 strains has revealed that, although serotype is a potent virulence determinant, genomic variation beyond serotype largely contributes to virulence (14). Strikingly, *pneumococcus* has been shown to actively modify host chromatin to drive infection, but also to maintain homeostatic colonizing conditions, depending on the bacterial strain (7, 8). Histone modifications induced by *pneumococcus* have been shown to be mediated by bacterial proteins rather than capsule (7, 8).

Chromatin modifications in an inflammatory setting have been documented for many years. For instance, stimulation of macrophages with the bacterial outer membrane component lipopolysaccharide (LPS) causes chromatin remodeling at numerous inflammatory genes (15). Although many of these modifications are transient and return to basal state once the stimulus has been removed, in some instances stimuli induce lasting modifications which alter the transcriptional response of the cell (16). This is best exemplified in immune memory (also termed “trained immunity”), in which innate immune cells exposed to a particular stimulus (infectious agent, inflammatory signals, etc.) mount a greater and faster response against secondary challenge, whether it is homologous or heterologous (17). Although still the subject of many investigations, chromatin remodeling is attributed for maintaining such memory responses. Indeed, immune memory has been associated with histones modifications at enhancer regions, transcription of long non-coding RNAs (lncRNAs), DNA methylation and reprograming of cellular metabolism (18). Most of the studies of immune memory have concentrated on cells of the immune system, such as monocytes, macrophages or NK cells, but little has been done on epithelial cells. Only recent studies have emerged documenting memory responses in these cells important for barrier functions (19, 20). Immune memory in different settings is therefore an important host response for protection against infection. However, how bacteria impact immune memory, or whether bacteria-induced histone modifications impose their own memory has not been explored.

In this study, we demonstrate that *S. pneumoniae* infection induces a persistent histone modification, di-methylation of histone H3 on lysine 4 (H3K4me2), which is maintained at least 9 days upon clearance of bacteria with antibiotics. We show that infection establishes a unique epigenetic program affecting the transcriptional response of epithelial cells, rendering them more permissive upon secondary infection. Our results establish H3K4me2 as a unique modification induced by infection, distinct from H3K4me3, which localizes to enhancer regions genome-wide. Importantly, we show that epithelial cells are changed following pneumococcal infection and respond differently upon a secondary infection many days after. Mainly, their metabolism and cell compartmentalization is altered, and they are more permissive to infection. Therefore, this study reveals evidence that bacterial infection leaves a memory in epithelial cells after bacterial clearance, via an epigenomic mark, thereby altering cellular responses for subsequent infections.

## Results

### Cells respond differently in primary and secondary infections

We hypothesized that epithelial cells could retain memory of a first infection, and this would result in a different response upon secondary infection. To test this, we set up an infection model in which alveolar epithelial lung A549 cells were infected with *Streptococcus pneumoniae* TIGR4 strain (Figure 1A). The control cells were uninfected (UI) and were treated in the same way as the infected cells. The cells were infected once (primary infection, 1°) or were maintained in culture for several cell divisions following a first infection (PI) or were infected twice (secondary infection, 2°). To ensure that phenotypes observed were inherited and thus due to memory, the cells in PI and 2° conditions were passaged twice following a first infection, ensuring that cells exposed to bacteria a second time were mostly descendants of those exposed the first time. Bacteria were grown to mid-log phase (∼1×10^8^ CFU/mL) and the cells were infected with a multiplicity of infection (MOI) of 20 or 35 according to experiment. After 1h or 3h of infection, cells were washed, and cocktail antibiotics (Ab) were added to the medium. Importantly, no more CFUs were recovered 24 hours post infection. We controlled that cell proliferation did not change after 1° infection by staining cells with the CellTrace™ CFSE dye (Figure S1).

**Figure 1:**
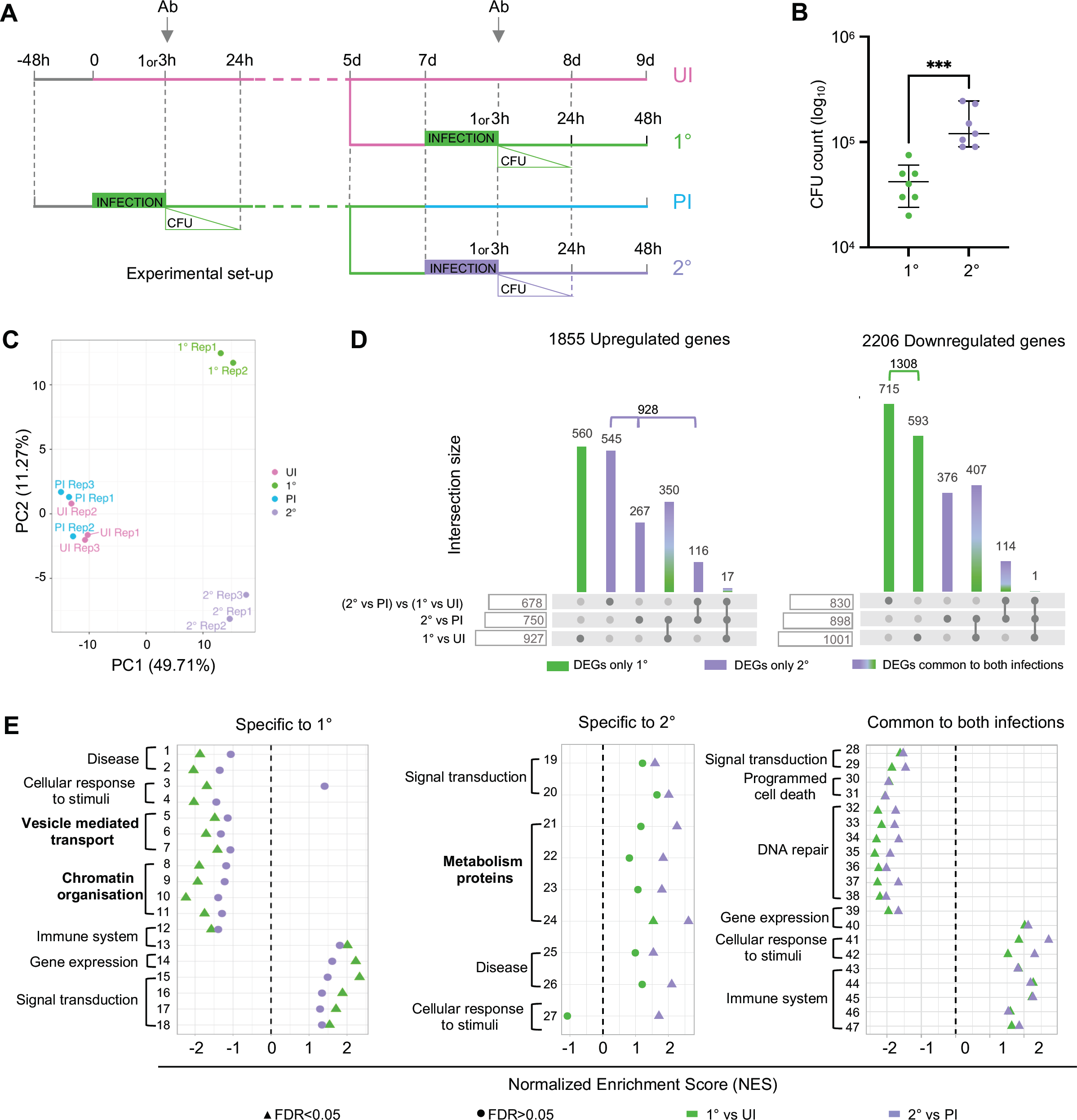
Differential infection efficiency and host cell transcriptome between primary and secondary infections. **(A)** Experimental set-up showing pink for uninfected cells (UI), green for primary infection with *S. pneumoniae* (*Spn*) (1°), blue for cells maintained in culture post infection with *Spn* (PI), purple for secondary infection with *Spn* (2°), hour (h), d (day). Cell washes followed by addition of antibiotics cocktail (Ab) after 1h or 3h of infection. Number of CFU after 24h of infection. **(B)** Cells were collected at 1h post primary (1°) and secondary (2°) infection with MOI 35 for CFU counts, 7 biological replicates, lines are the mean ± SEM and statistical significance was calculated by Mann-Whitney test, ***p =0,0006. **(C)** Principal component (PC) analysis of gene expression samples of the UI, PI, 1° and 2° infection. Principal components calculated on count data after removal of replicate batch effect with ComBat. Samples coloured according to time point. **(D)** Differentially expression genes analysis of transcriptome. Number of genes up or downregulated at 3h post 1° or 2° infection in comparison to UI or PI (at 7 d) and between infections ((2° vs PI) vs (1° vs UI)), adjusted p-value < 0.05. Column hight indicate the number of genes defined as differentially expressed (DEG) in one or more of the above mentioned comparisons. Column colour specifies whether the subset of genes is differentially expressed only in the 1° (green) or 2° (purple) infection or in both (green/purple gradient). **(E)** Functional scoring analysis. Reactome categories significantly enriched in the 1° in comparison to UI (1°vsUI) and 2° infection in comparison to PI (2°vsPI). Pathways are separated depending on whether they are enriched only in the 1° (left) or 2° (middle) infection or in both (right). Normalized enrichment score (NES) calculated for each significatively enriched pathway with a false discovery rate (FDR). See complete list of pathways corresponding to each category in supplementary table S1.

Using this protocol, we evaluated the infection efficiency during the primary (1°) and secondary (2°) infection, by comparing the number of bacteria recovered. Following 1° infection, 4×10^4^ bacteria are recovered on average. Interestingly, approximately 10-fold more bacteria are recovered following a 2° infection. Therefore, cells are more permissive to infection upon on a second exposure (Figure 1B).

To better understand the differences between 1° and 2° infection, we performed transcriptomic analysis of the four conditions, UI, 1°, PI and 2°. Differentially expressed genes (DEGs, adjusted p-value < 0.05) were determined for the 1° and 2° infections (1° vs UI and 2° vs PI) and between infections ((2° vs PI) vs (1° vs UI)). As shown in Figure 1C, the primary source of variability of the time course (PC1 ∼50%) coincides with the differences between UI/PI time points and the 1° and 2° infections. These results indicate that cells return to a basal transcriptional state following 1° infection, which is indistinguishable from UI condition. However, the variation between the 1° and 2° infection (PC2 ∼11%), reflects the differential response between the two infections. In total, when comparing the DEGs, there are 1855 upregulated and 2206 downregulated genes (Figure 1D). Of which, 560 upregulated and 1308 downregulated are significantly different only in the 1° infection in comparison to uninfected cells (green bars), and 376 downregulated and 928 upregulated are specific of the 2° infection (purple bars). These results show that the 1° infection is dominated by downregulated DEGs, while upregulated DEGs prevail during the 2° infection, and highlight the observation that cells following 1° infection respond differently to a 2° infection.

By performing Gene Set Enrichment Analysis (GSEA), we obtained a functional description of the transcriptome using the gene sets reported in the Reactome database (Figure 1E and Table S1). A normalized enrichment score (NES) is calculated for each significatively enriched pathway (FDR>0.05) and indicates expression levels. Interestingly, some of functional categories identified are different between the two infections and include Chromatin organization and Vesicle mediated transport as specific to 1° infection, whereas metabolism of proteins is specific to 2° infection. It should be noted that although some functional categories are shared between the different infections, they do not include the same pathways in each infection (Table S1). Overall, our results reveal that the transcriptional status of cells during a 1° infection is different than that during a 2° infection, suggesting that cells become altered following 1° infection.

### *S. pneumoniae* actively modifies cells following a primary infection

Gene expression analysis and GSEA uncovered different cellular functions during 1° and 2° infection, one of which being related to metabolism. Thus, we monitored the metabolic activity of cells using alamarBlue, an oxidation-reduction indicator which becomes fluorescent in the reducing environment of metabolically active cells (Figure 2A). Interestingly, alamarBlue fluorescence increased at a significantly faster rate following 1° infection compared to 2° infection, demonstrating a loss of oxidation-reduction rate during 2° infection. Strikingly, if the 1° infection is performed with inactivated bacteria, the loss in metabolic activity during 2° infection is not observed. These results therefore show *Streptococcus pneumoniae* (*Spn*) is actively modifying host cells during a 1° infection, which results in an altered metabolic activity upon 2° infection.

**Figure 2:**
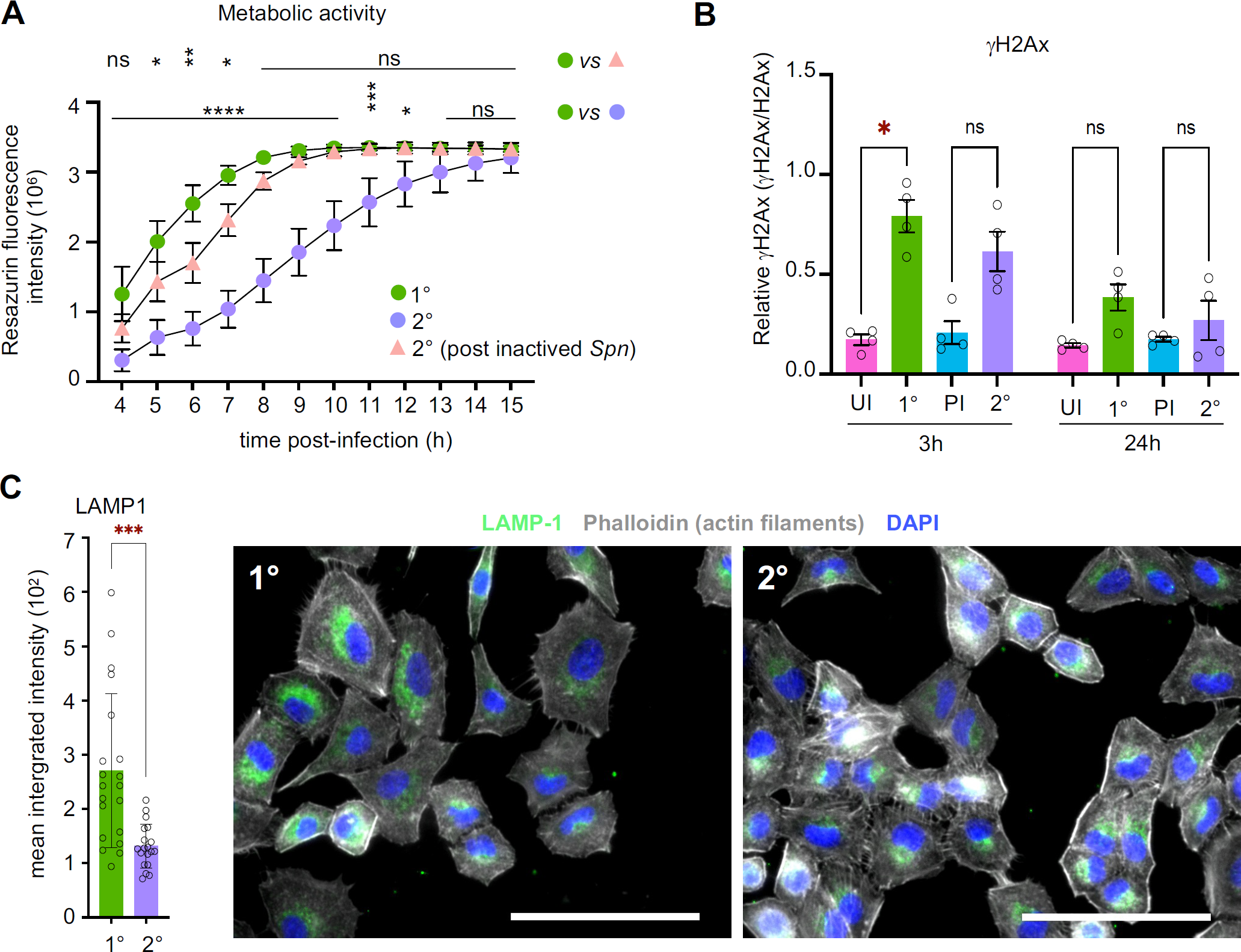
*S. pneumoniae* actively modifies cells following primary infection. **(A)** Cells metabolic activity measured by alamarBlue assay. A549 cells treated at 3h post infection (MOI 20), 1 measure of fluorescence/h for 12h. Results are expressed as resazurin fluorescence intensity. Plot shows mean ± SEM from 4 biological replicates. Statistical significance was determined by two-way ANOVA with FDR Benjamini-Hochberg correction for multiple comparisons for primary (1°) versus secondary (2°) infection, and primary (1°) versus secondary^§^ (2°) infection (^§^*Spn* inactived at 1° infection then *Spn* live at 2°), ns = not significant, *p < 0.05, **p <0.01, ***p <0.001, ****p <0.0001). **(B)** Immunoblot detection of γH2Ax from infected and uninfected A549 cells at 3h and 24h post infection (MOI 20), at primary infection (1°), PI (7d or 8d) and secondary (2°) infection. Histogram show actin-normalized ratio of γH2Ax to total H2Ax from 4 biological replicates. Error bars are the standard error (SEM) of the mean of γH2Ax levels. Statistical significance was determined by two-way ANOVA comparing all means with Tukey’s multiple comparisons test (ns = not significant, *p =0.0333). **(C)** Quantification of LAMP1 normalized to the segmented to the cytoplasm and filtered to nuclei at 1h (MOI 35) in 1° and 2° conditions. Plot shows quantify of 1029 cells, mean integrated intensity of 39 Regions of Interest (1029 cells total) ± SD, statistical significance was determined by Mann-Whitney test (***p <0.0001). Representative images of immunofluorescence microscopy of A549 cells stained LAMP1 (GFP; green), nucleus (DAPI; blue) and actin (Phalloidin; grey) at 1° and 2° infection. Crop of images microscopy taken at 20x magnification. Scale bar = 100 μm.

DNA damage was another category of genes enriched by GSEA. We therefore measured *Spn* induced DNA damage response after 1° and 2° infection by monitoring the level of γH2Ax, which is essential to the efficient recognition and/or repair of DNA double strand breaks (Figure 2B and S2). Infection clearly induces a measurable level of DNA damage, as the levels of γH2Ax increases after 1° infection, though slightly less after 2° infection. DNA damage is no longer observed 24 hours after infection, indicating that cells have recovered following both 1° and 2° infection.

Similarly, GSEA pointed to a differential expression of vesicle mediated transport proteins specifically in the 1° infection. We monitored cell transport by immunofluorescence marking of lysosomes with the LAMP1 marker (Figure 2C). Interestingly, in a very heterogeneous manner, more LAMP1-positive lysosomal compartments are detected in the 1° infection compared to the 2°, even though more bacteria are present. Therefore, cell compartments are altered following 1° infection. In addition, we noticed stronger staining of actin stress fibers during the 2° infection, which further supports a physiological modification of host cells. Thus, our results clearly show that following infection with 1° infection, cells have both biological phenotypes and transcriptional activity altered and inherited/persist in daugther cells that were not exposed to bacteria.

### Cells maintain an epigenetic mark after infection

Our data demonstrate that 1° infection changes the cells in a lasting manner, suggesting a mechanism to maintain a memory of infection through multiple cell divisions. This raised the question of whether an epigenetic mechanism could contribute to these changes, with a particular interest in histone modifications. To identify potential modifications, we performed a multiplex ELISA, which analyzed twenty-one modified histone H3 patterns simultaneously in histone extracts from cells at 3 hours (1°) and 7 days (PI) after infection compared to uninfected cells (UI) (Figure 3A and S3A). Interestingly only one histone modification was significantly detected at 7 days following bacteria challenge, di-methylation of histone H3 on lysine 4 (H3K4me2). Surprisingly, this modification is not detected directly after 1° infection, and appears later. We further measured H3K4me2 levels in whole cell extracts by immunoblotting (Figure 3B and S3B). By comparing the level of modified H3, we confirmed the significant increase at 7 days (PI) compared to 3 hours (1°) and uninfected cells (UI). Given that H3K4me2 is often associated with H3K4me3, as the methyltransferase is the same, we also measured levels of tri-methylation under the same conditions. However, the level of H3K4me3 does not change either upon 3h of infection (1°) or at 7 days (PI) after infection (Figure 3C and S3C). With this data, we concluded that infection leaves one specific epigenomic mark after bacterial clearance and that it persists for at least 7 days.

**Figure 3:**
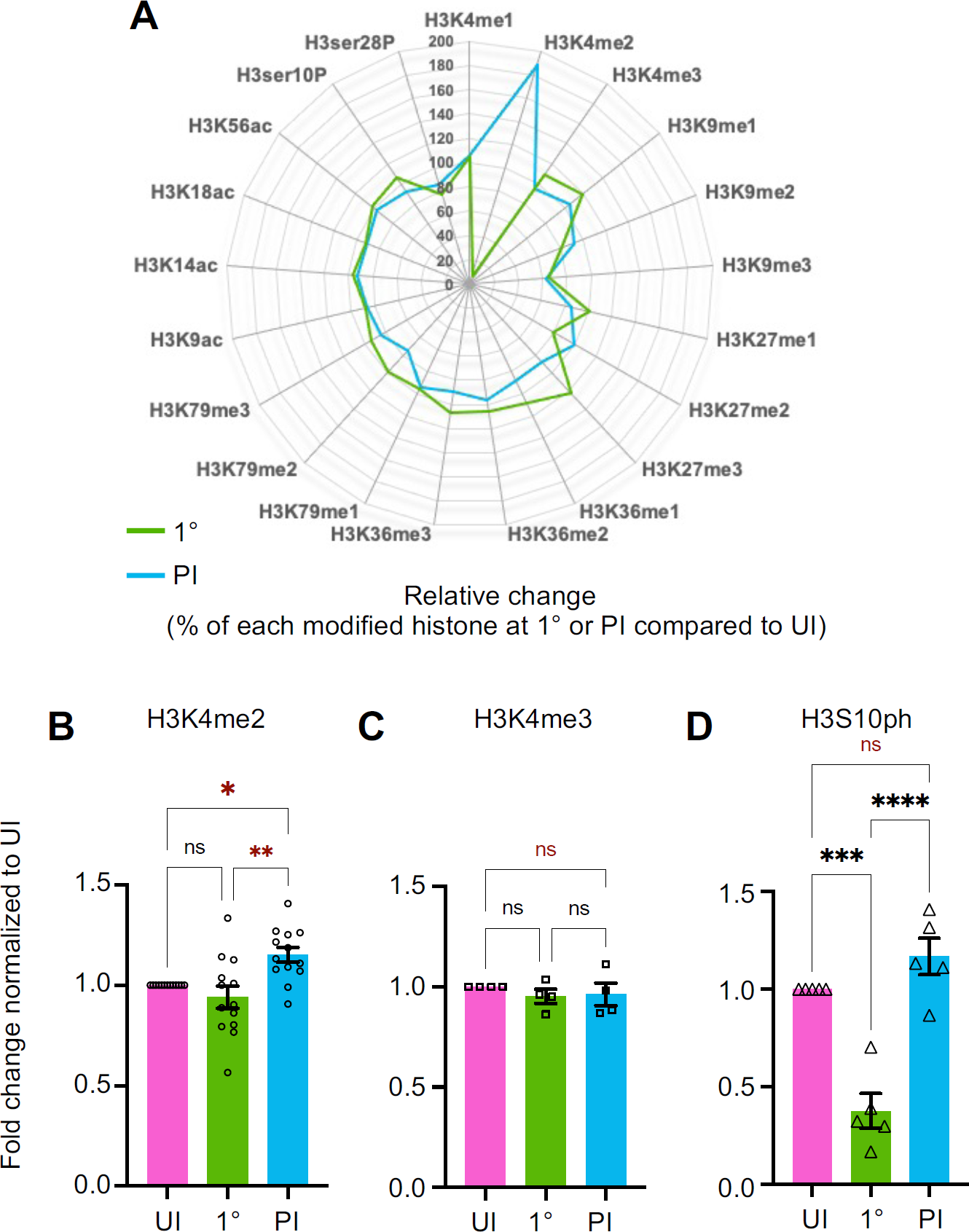
Cells maintain an epigenetic mark after primary infection. **(A)** Histone H3 modifications by Multiplex ELISA assay. Quantitation of twenty-one modified histone H3 patterns simultaneously. Radar plots representing total H3-normalized ratio of the relative change (%) of each histone H3 modification between 1° infection (3h) (MOI 20) or PI (7d) and uninfected cells. See table heatmaps with values in supplementary figure S3.A. **(B)** Immunoblot detection of H3K4me2 from uninfected (UI) and infected (MOI 20) whole cells A549 lysates at 1° infection (3h) and PI (7d). Histogram show mean ± SEM of values expressed as normalized band intensity relative to actin followed by fold change of infected cells at 1° infection (3h) or PI (7d) to uninfected cells (UI) for 14 biological replicates. Statistical significance was determined by one-way ANOVA method with Tukey’s multiple comparisons test (ns = not significant, *p =0.0209 **p =0.0011). See representative image of immunoblot in supplementary figure S3.B. **(C)** Immunoblot detection of H3K4me3 from uninfected (UI) and infected (MOI 20) whole A549 cells lysates at 3h (1°) and 7d (PI) post infection. Data points represent mean ± SEM from 4 biological replicates. Histogram show the values expressed as normalized band intensity relative to actin followed by fold change of infected cells at 1° infection (3h) or PI (7d) to uninfected (UI). Statistical significance was determined by one-way ANOVA method with Tukey’s multiple comparisons test (ns = not significant). See representative image of immunoblot in supplementary figure S3.C. **(D)** Immunoblot detection of H3S10ph from uninfected (UI) and infected (MOI 20) whole cells lysates of A549 cells at 1° infection (3h) or PI (7d). Data points represent mean ± SEM from 5 biological replicates. Histogram show values expressed as normalized band intensity relative to actin followed by fold change of infected cells at 1° infection (3h) or PI (7d) to uninfected (UI). Statistical significance was determined by one-way ANOVA method with Tukey’s multiple comparisons test (ns = not significant, **p =0.0002, ***p <0.0001). See representative image of immunoblot in supplementary figure S3.D.

To further assess the specificity of this lasting modification, we measured the levels of H3 phosphorylation, as we had previously shown that *Spn* infection induces dephosphorylation on serine 10 (7). Although a significant decrease in phosphorylation levels is observed 3h post infection (1°), the levels returned to uninfected levels by 7 days (PI) (Figure 3D and S3D). Therefore, H3S10 dephosphorylation is transient in comparison the H3K4me2, which is maintained.

To evaluate how the increase in di-methylation levels of H3K4 is distributed from cell to cell, we performed immunofluorescence analysis to quantify the level of histone modification at individual cell resolution. Images show a slight variability of H3K4me2 levels from one cell to another, with some cells staining more intensely than others under all conditions (Figure 4A). Greater magnification reveals di-methylation of H3K4 is localized to euchromatin under all conditions, the form required for transcriptional activity (Figure S4A). Images were quantified to evaluate the levels of H3K4me2 in Figure 4B. The global levels significantly increased at 48 hours following infection (1°) and were still elevated 9 days (PI). Images show that both the number and the intensity of fluorescence increase in both conditions. This increased level of H3K4 di-methylation 9 days after primary infection (PI, live) is not enhanced after a secondary infection (2°, live) (Figure S4B), suggesting that a second bacterial challenge does not induce a change in the persistence of this epigenetic mark. In contrast, tri-methylation levels on the same histone H3 residue (H3K4me3) exhibit a different pattern of methylation, with a small increase at 24 hours following infection (1°) but returning to basal levels 8 days (PI) (Figure 4C). Thus, our data demonstrate that H3K4me2 is persistent at least 9 days following infection with *Spn* and that it is specifically maintained in the di-methylated form.

**Figure 4:**
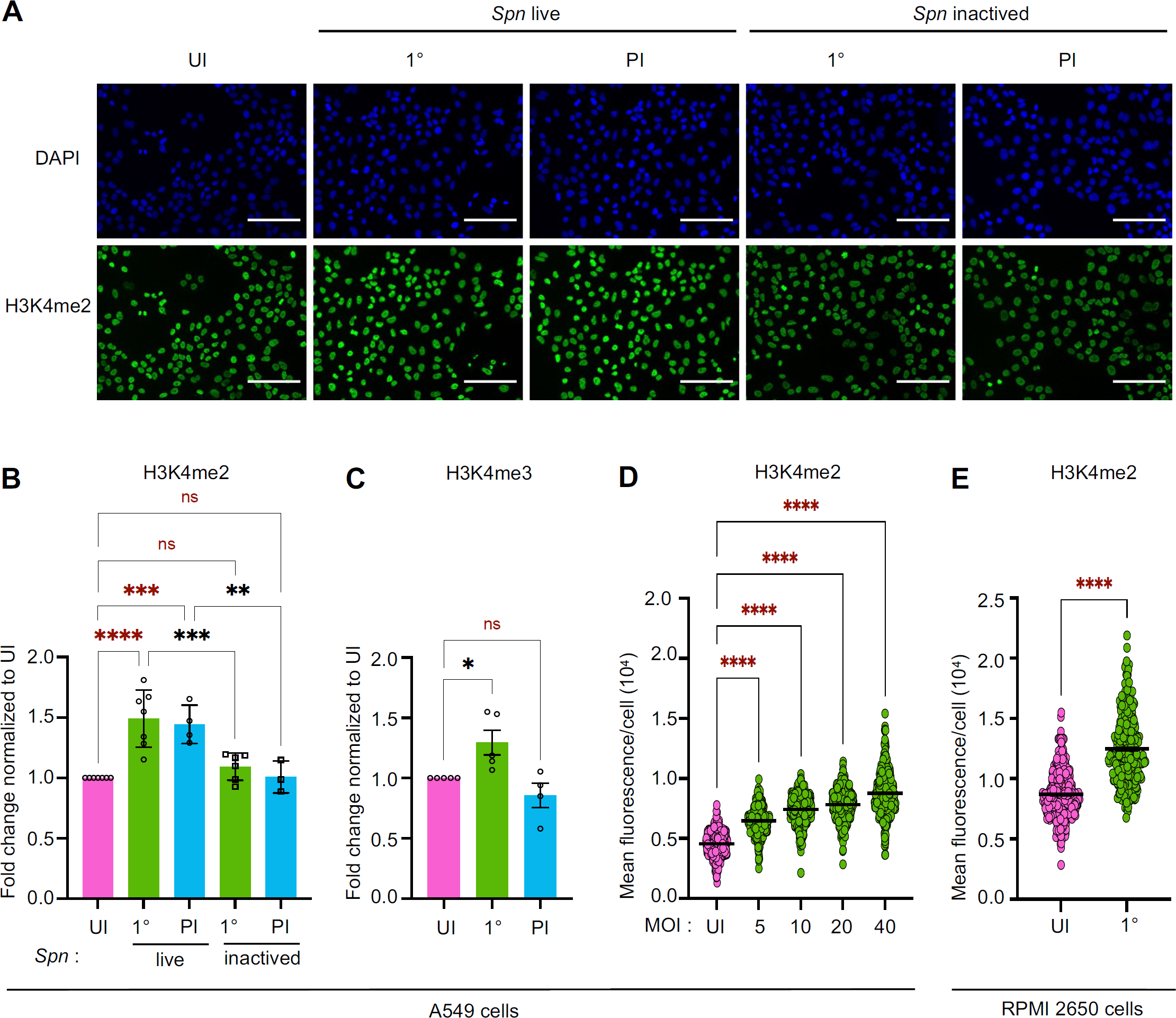
The increase in H3K4me2 levels is actively induced by live bacteria. **(A)** Representative images of immunofluorescence detection of nuclear H3K4me2 in A549 cells uninfected (UI) or infected (MOI 20) at 1° infection (48h) and PI (9d). Paraformaldehyde fixed cells stained for H3K4me2 (GFP; green) and nuclei (DAPI; blue). Images microscopy taken at 20x magnification. Scale bar is 100μm. **(B)** Quantification of H3K4me2 normalized to the segmented nuclei using DAPI signal. Data points expressed as fold change of mean fluorescence intensity of infected cells at 1° infection (48h) and PI (9d) to uninfected (UI) with *Spn* live and *Spn* inactived. Graphs display quantification from 3 to 7 biological replicates with the mean values of each condition. Statistical significance was determined by one-way ANOVA with Fisher’s LSD test (ns = not significant, **p = 0.0012, ***p = 0.0001, ****p <0.0001). **(C)** Quantification of H3K4me3 normalized to the segmented nuclei using DAPI signal. Data points expressed as fold change of mean fluorescence intensity of infected cells at 1° infection (24h) and PI (8d) to uninfected (UI). Graphs display quantification from 5 biological replicates. Statistical significance was determined by one-way ANOVA with Fisher’s LSD test (ns = not significant, *p = 0,02). **(D)** Quantification of H3K4me2 according to Multiplicities of infection (MOI) normalized to the segmented nuclei using DAPI signal at 1° infection (48h). Data points represent the mean of fluorescence intensity of H3K4me2 within individual nuclei. Histogram shows the mean and error bars represent SEM from 435 to 480 nuclei at uninfected cells (UI) and each MOI. Statistical significance was determined by one-way ANOVA with Fisher’s LSD test (****p <0.0001). **(E)** Quantification of nuclear H3K4me2 from RPMI 2650 cells normalized to the segmented nuclei using DAPI signal. Data points represent the mean fluorescence intensity of H3K4me2 within individual nuclei. Histogram shows the mean and error bars represent SEM from 340 to 480 nuclei at 1° infection (MOI 10) (48h) and uninfected cells (UI). Statistical significance was determined by two-tailed unpaired t test (****p <0.0001).

To demonstrate that H3K4 di-methylation is actively induced during infection, we determined whether the increase in H3K4 di-methylation is proportional to the bacterial load. We performed infections with increasing numbers of bacteria and measured H3K4me2 levels by immunofluorescence (Figure 4D). These results clearly show that increasing the bacterial load increases the levels of H3K4me2. We also monitored H3K4me2 levels upon incubation of cells with inactivated bacteria (Figure 4B). Indeed, our alamarBlue data indicated that metabolic activity was altered in cells only if they were previously exposed to live bacteria. Interestingly, and in agreement with what we observe with alamarBlue, H3K4me2 levels do not increase upon contact with inactivated bacteria (Figure 4D). These results allow us to determine that the H3K4me2 epigenetic modification of cells is a feature of a live infection and is not due to the passive presence of bacterial factors.

A549 cells are terminally differentiated type II pneumocytes with little potential to transmit epigenetic changes in a physiological setting. Therefore, we performed infections of undifferentiated nasal cells, RPMI 2650, and measured levels of H3K4me2 by immunofluorescence. Similarly to A549 cells, we observe a strong increase in the level of H3K4me2 after 48 h of infection (1°) (MOI 10) compared to uninfected cells (UI) (Figure 4D). In these cells, we also confirmed the specificity of our antibody by performing the same immunofluorescence experiments with another H3K4me2 antibody cataloged in the Histone Specificity Database (http://www.histoneantibodies.com). Similarly, we were observed a strong increase in the level of H3K4me2 after 48 h of infection (1°) compared to uninfected cells (UI) (Figure S4C), independently of the antibody used. Together these results show that H3K4 is di-methylated by infection in undifferentiated cells and is not restricted to alveolar epithelial A549 cells.

### Methylome dynamics during 1° infection

To study the distribution of H3K4me2 along the host cell genome, we combined chromatin immunoprecipitation with sequencing (ChIP-seq) to profile the methylome upon infection. The analysis recovered approximately ∼28 000 peaks 7 days post infection (PI) but only ∼12 600 directly 3 hours after infection (1°) (Figure 5A), which is consistent with immunoblotting and immunofluorescence data showing a steady increase in H3K4me2 levels following primary infection until 7 and 9 days. Interestingly most of the peaks recovered following infection (1°) are shared peaks with both uninfected (UI) and PI conditions, and only ∼1 700 (∼13%) are unique to that 1° condition, supporting the idea that this time point is transitional. In contrast, more than ∼30% of peaks unique to the PI condition (8 221) are unique and distinct from either UI or 1° conditions. These results are consistent with a principal component analysis along the infection time course (Figure 5B) which reveals that the first source of variability for H3K4me2 coincides with the difference between UI and PI conditions (PC1 ∼44%). Therefore, genome wide epigenomic changes take place upon 1° infection. The second source of variability (PC2 ∼27%) corresponds with the distance between the two biological replicates of the 1° infection time point, possibly underlying the heterogenous response of the epigenome after the 1° infection. Taken together, these data show that upon infection, a large number of peaks differ in comparison to uninfected (UI), and this difference continuously increases up to 7 days (PI).

**Figure 5:**
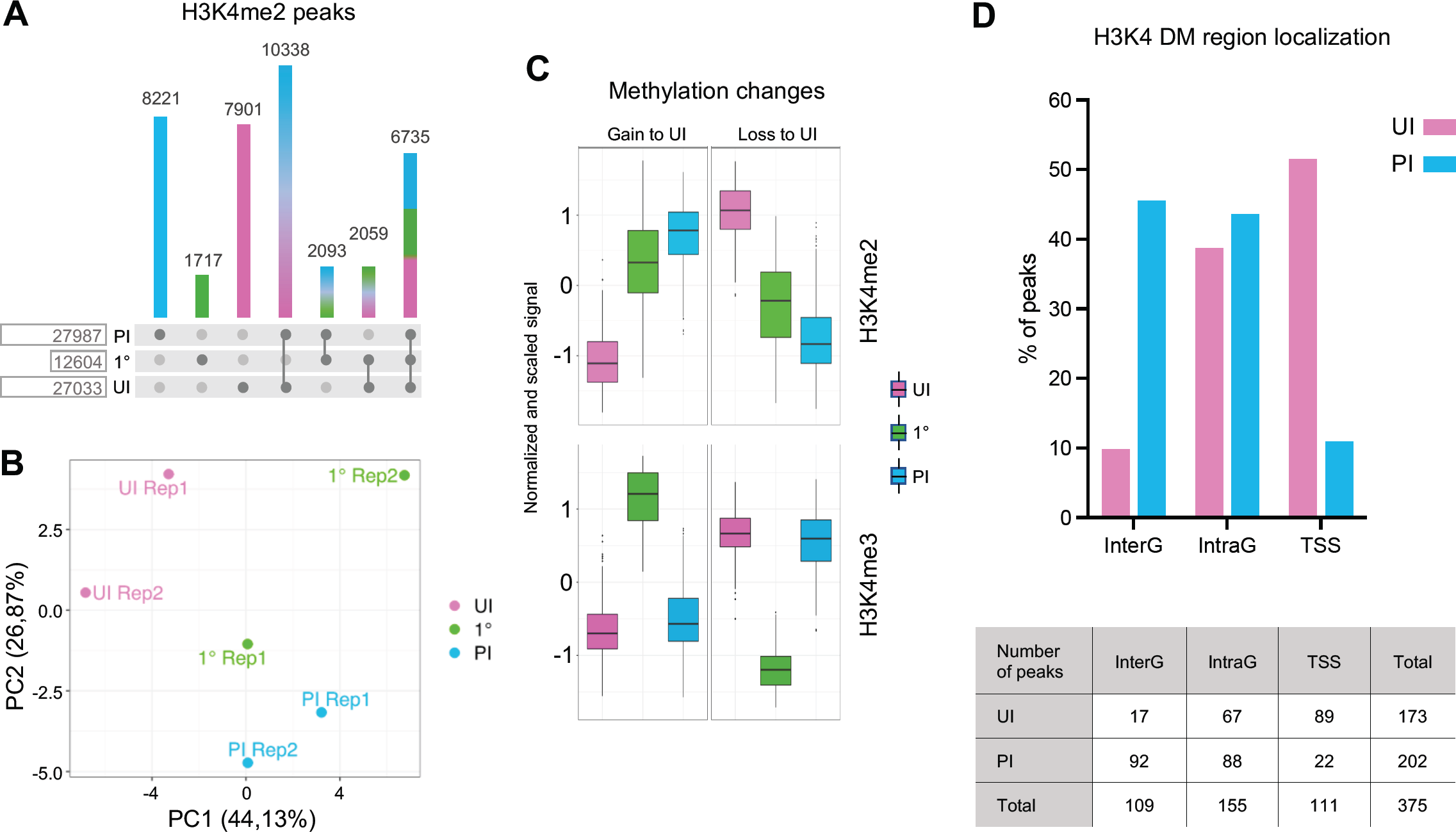
Methylome dynamics during primary infection. **(A)** Number of reproducible peaks between replicates per time point are compared among the three times points UI, 1° and PI by simple overlap of at least 50% of the peak length. **(B)** Principal component (PC) analysis of the H3K4me2 peaks by sample UI, 1° and PI. Principal components are calculated on peak enrichment, i.e. read counts, after removal of replicate batch effect. **(C)** Methylation changes for H3K4me2 and H3K4me3. Count distribution is plotted per time point (UI, 1°, PI) for the most dynamic regions (DMRs), i.e. those showing the biggest changes in methylation. Read counts per peak are normalised by sequencing depth, peak length and the replicate batch effect is corrected; they are subsequently centred and scaled by their standard deviation. Peaks are clustered and two global profiles are shown: gain and loss of methylation with respect to UI. **(D)** H3K4me2 DMRs localization. Percentage and number of UI peaks (loss of methylation) and PI peaks (gain of methylation) according to the localization: transcription start site (TSS) for peaks overlapping the 2Kb interval centred around the transcription start site; intragenic (IntraG) for peaks located within the gene annotations and outside the TSS interval; intergenic (InterG) for all other peaks.

We further clustered the enrichment of H3K4me2 for the most dynamic peaks, i.e. those showing the biggest changes in methylation (Figure 5C), and separated them into two global profiles, those that gain and those that lose in signal intensity compared to UI. These profiles reveal that the gain or loss observed following infection (1°) are maintained at 7 days (PI). Interestingly, we compared this H3K4me2 methylation profile to that of ChIP-seq we performed on H3K4me3, and observed a very different peak pattern. Indeed, tri-methylation peaks were detected rapidly after infection (1°), but disappeared, leaving no differential peaks at 7 days (PI) compared to UI. These results further support that pneumococcal infection specifically induces di-methylation, a modification that is maintained genome-wide at least 7 days (PI).

We proceeded to map the localization of the most significant H3K4me2 peaks, which we name “differentially methylated” (DM), that were detected 7 days post-infection (PI) compared to those detected in uninfected condition (UI) (Figure 5D). Under these stringent conditions, we mapped 375 peaks, 173 from UI and 202 for PI conditions. Interestingly, although H3K4me2 in uninfected cells is lowly abundant in intergenic regions (InterG), and most abundant in intragenic regions (IntraG) and at transcription start sites (TSS), distribution is altered upon infection. In PI condition, most peaks are present at InterG and IntraG regions, and not at TSS. By incorporating ENCODE data to our data of these 375 DM peaks, we performed unbiased clustering according to chromatin state, profile, and localization (Figure S5A). Strikingly, peaks from UI condition clustered together according to their localization to TSS and regions known to accumulate H3K4me3. These peaks are also shared with our ChIP-seq data on H3K4me3. In contrast, PI peaks are unique and cluster in regions known to have marks of enhancer regions (H3K27ac and H3K4me1). Therefore, altogether, our ChIP-seq data show that after 7 days (PI), infection has remodeled the host methylome as cells have differential peaks to uninfected condition (UI), and this process occurs mainly in enhancer regions of the genome.

### Identification of the regulatory modes underlying primary and secondary infections

Even though our ChIP-seq data identified a significant number of di-methylated peaks 7 days post infection (PI), at this same time point, there are no gene expression changes between UI and PI conditions. However, upon 2° infection, gene expression is altered in a significant manner compared to 1° infection, raising the question of a possible link between H3K4me2 peaks and transcriptional changes. To address this point, we integrated our transcriptome data with our ChIP-seq data in order to find links between the two analyses (Figure S5B). A direct intersection of the differentially methylated peaks (DMs) with the differentially regulated genes (DEs) only yielded 29 direct associations (dark line continuous in Figure S5B). Amongst these, only 9 were linked with a gain in methylation, which is we decided was not sufficient to further our analysis. Therefore, we extended our study to include regions of methylation and genes that displayed a upwards trend even though they were not statistically significant (black dotted lines in Figure S5B). Using this method, we predicted 3845 regulatory links between H3K4me2 regions and genes. Together with the above 29, we analyzed 3874 associations from a global perspective, using a Multiple Factor Analysis (MFA) (21). This factorial method is aimed at uncovering the main sources of variability that characterize a dataset among multiple sets of quantitative and qualitative variables. To perform this method, we used both transcriptome and methylome, more precisely: the complete transcriptome dataset of the UI cells, post 1° infection, PI and post 2° infection (TOME); the H3K4me2 log fold change of PI vs UI and the average H3K4me2 level of the 1° infection time course (METH). We added the epigenome profile of the UI cells (EPIG) described by six key histone marks symptomatic of the main active and repressive chromatin states (CS). We retrieved ten ENCODE samples H3K4me3, H3K4me2, H3K4me1, H3K27ac, H3K79me2 and H3K27me3 from the ENCODE database for A549 cells (22). Finally, we considered two additional variables that describe the regulatory links between genes and H3K4me2 peaks, as estimated by the T-Gene tool of the MEME suite (23): the distance in bp separating each gene-peak pair (DIST) and the correlation between chromatin and gene expression changes in an independent tissue panel (CORR) (Figures S6A).

Factors resulting from the MFA summarize the structure of the joint groups of variables, as shown by the cumulative percentage of variance associated to the first 5 factors/dimension (Dim) and the correlation coefficients of groups of variables with each factor (Figure S6B). Such factors unveil correlations among the different groups, one of which is very interesting for our study, namely the correlation between DMRs and DEGs in Dim1 and that between TOME and METH in Dim3. (Figure S6B and Figure 6A). Indeed, earlier in the differential expression analysis, we highlighted that after 1° infection, cells respond by downregulated DEGs while upregulated DEGs prevail in the 2° infection. Furthermore, in the methylome analysis, an increase in dynamic H3K4me2 peaks at 7 days (PI) was demonstrated. This correlation suggests a strong link between an increase in gene expression levels and the increase in H3K4 di-methylation at 7 days (PI). Additionally, Dim2 unveils a correlation between the EPIG, CS and DMR, further supporting the H3K4me2 epigenome profile at enhancer region (Figure S5A).

**Figure 6:**
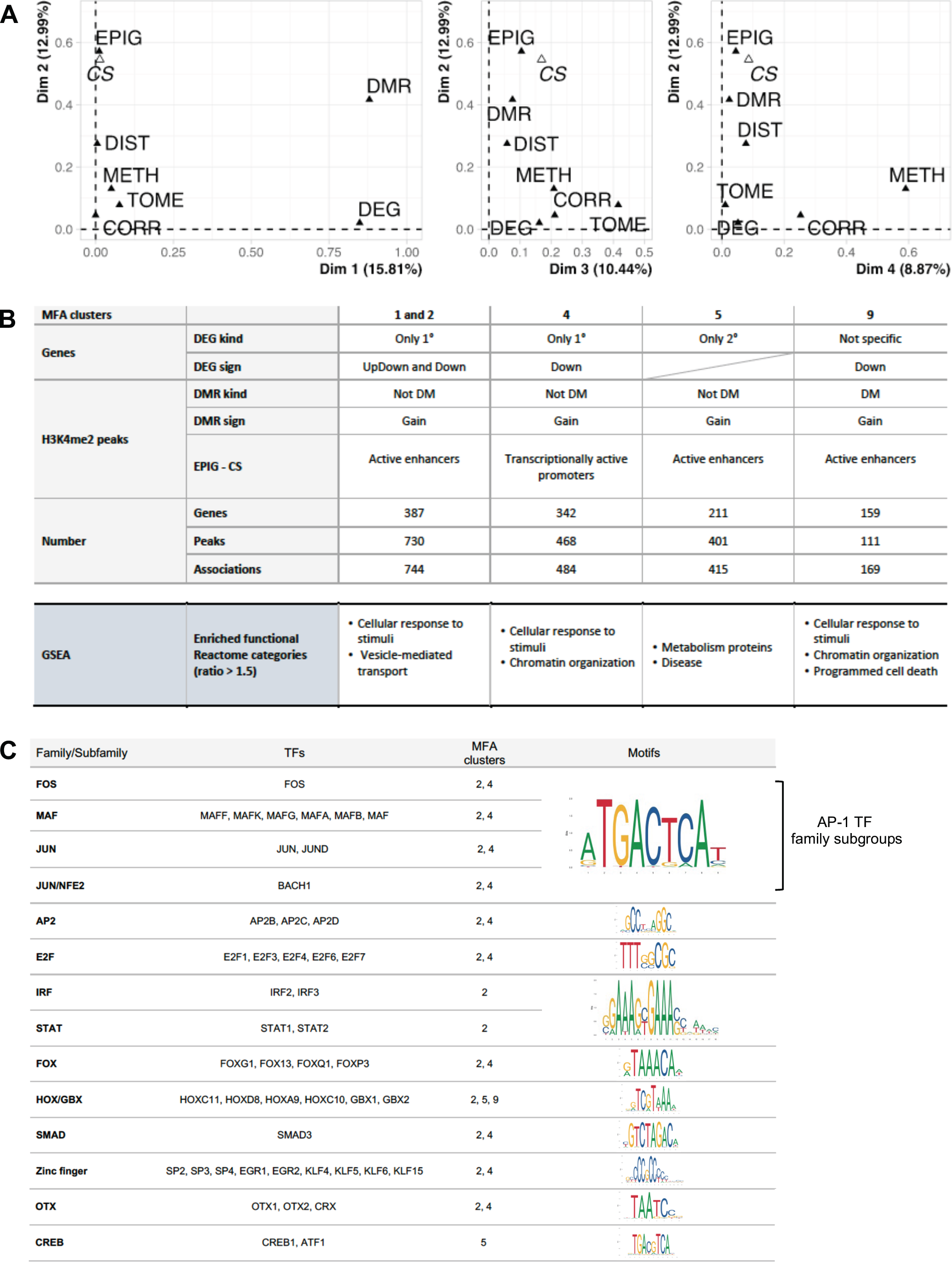
Identification of the regulatory modes underlying the primary and secondary infections. **(A)** Contribution of groups of variables to the definition of factors/dimensions. Groups of variables constitute the Multiple Factor Analysis (MFA) input matrix and describe the genes (TOME = Transcriptome, DEG = Differentially expressed gene), the H3K4me2 peaks (Methylome = METH, DMR = Differentially methylated region, EPIG = Chromatin profile, CS = Chromatin state) and their association (DIST = distance, CORR = correlation). **(B)** Description of the four regulatory modes associated to an increase of H3K4me2. Clusters of gene-peak associations are characterised in terms of the categorical variables used in the MFA (DMR, DEG), the chromatin state of the peaks (EPIG, CS). Representation of enriched functional Reactome categories from GSEA for each MFA cluster: the observed number of genes belonging to a given Gene Set and the expected number per cluster (with ratio >1.5). **(C)** Transcription factors (TFs) associated to the regulatory modes. Table lists the TFs with binding motifs enriched in H3K4me2 peaks from the selected MFA clusters.

Using the top factors obtained by the MFA, we clustered the regulatory links and characterize them using the variables enriched among dynamically marked H3K4me2 region, differentially expressed genes and the functional categories specifically enriched. This analysis allows a subclassification of the data into smaller clusters unified by common features (Figure 6B, and S6C). We focused our attention on clusters 1, 2, 4, 5 and 9 that contain most of the peaks gaining H3K4me2 methylation. MFA cluster 9 is the only cluster including H3K4me2 peaks that are differentially methylated in a significant manner between UI and PI conditions, while all other clusters include all H3K4me2 peaks. Such an analysis reveals that most H3K4me2 peaks gained in active enhancers are associated with downregulated genes, either specific to 1° infection (MFA clusters 1 and 2), or in a nonspecific manner (MFA cluster 9). At the 1° infection, we also find peaks associate with transcriptionally active promoters, but at downregulated genes, further supporting a role of H3K4me2 in transcriptional repression (MFA cluster 4). Thus, H3K4me2 seems to play a role in gene downregulation, and GSEA analysis of these, attributes their function to chromatin organization and cell response to external stimuli. Furthermore, it is interesting to note that the functional analysis of MFA clusters 1 and 2 (only 1° infection) reveal a role in vesicle mediated transport and supports our LAMP1 data showing more LAMP1-positive lysosomal compartments detected in the 1° infection. Peaks associate with active enhancers exclusively mark genes specific to 2° infection regardless of up or down regulation (MFA cluster 5). Interestingly, functional analysis of this cluster displays a role in metabolism and support our alamarBlue data exhibiting distinct metabolism dynamics upon 1° and 2° infection. Therefore, although straightforward global links between transcriptome and epigenome yielded a very small number of genes, multiple factor analysis generated meaningful associations by clustering our data into smaller clusters revealing that infection induced H3K4me2 probably has different roles depending on the genomic localization and the transcriptional context in which it is placed.

We reasoned that each MFA clusters could be regulated by different transcription factors. Therefore, we performed a transcription factor motif enrichment analysis on the H3K4me2 peaks of clusters gaining marking upon infection, i.e. MFA clusters 1, 2, 4, 5 and 9 (Figure 6C). Interestingly, we obtained the enrichment of TFs known to be involved in immune regulation, such as SMAD3, and IRF and STAT family members, all shown to have a role during infection by viral and bacterial pathogens. Other TFs families, such as Fox, Hox, Zinc finger, OTX and CREB that have more extended cellular functions. Remarkably, this analysis reveals significant motif enrichment for the Activator Protein-1 (AP-1) family members. This ubiquitous dimeric protein complex composed of different Jun, Fos and MAF subfamilies, has been shown to play a role mediating inflammation and in transcriptional memory responses (24–26). Therefore, *S. pneumoniae* infection modifies host chromatin in an impactful way by impacting major transcription factor regulatory processes.

## Discussion

In our study, we demonstrate that infection by *S. pneumoniae* results in a long-lasting modification of epithelial cells characterized by changes in transcription, phenotype and a specific histone modification, H3K4me2. This modification is detectable shortly after infection, specific to di-methylation of lysine 4 and persists for at least 9 days after bacteria are cleared with antibiotics. Genome-wide analysis of H3K4me2 peaks shows that they are mainly localized to enhancer regions in which pioneer transcription factors are predicted to bind, and when correlated with transcriptional regulation of nearest genes these are mostly repressed. Interestingly, the observed epigenetic modifications arise upon interaction of epithelial cells with live bacteria, indicating an active pathogenic mechanism, resulting in a higher bacterial load upon secondary infection.

Bacterial presence or production of secondary metabolites are types of stimuli that are sensed and to which eukaryotic cells respond through integration of signal transduction resulting in chromatin rearrangements and transcriptional modulation. However, certain bacteria impose their own infection-favoring histone modifications to reprogram host transcription (27–29). To target chromatin, bacteria have been shown to either hijack and repurpose host enzymes or inject factors termed nucleomodulins which will directly modify histones (30). We had previously shown that *pneumococcus* dephosphorylates histone H3S10 by relocalizing the host PP1 phosphatase to promote infection, and that the pneumolysin toxin PLY and pyruvate oxidase SpxB were the main bacterial factors mediating this modification (7). However, we had not assessed the lasting potential of this mark. In our current work, we show that H3S10 dephosphorylation is a transient modification that rapidly returns to pre-infection levels shortly after clearing of bacteria. In contrast H3K4me2 is acquired during infection and persists multiple cell passages post antibiotic treatment. This is the first characterization of a mark that is maintained and thereby affects cell physiology over time. Although we do not know the mechanism by which *S. pneumoniae* induces this modification, we show that it is an active mechanism that only occurs if bacteria are alive. Indeed, when we performed experiments with paraformaldehyde fixed bacteria, which retains surface proteins and capsule intact, H3K4 is not methylated and phenotypic changes to host cells do not occur. Such a finding would suggest that H3K4me2 is induced by a bacterial factor secreted during infection that is not present on the surface previously to interaction with host cells. Whether this factor indirectly induces H3 methylation through activation of host machinery or directly through secretion of methyltransferase mimic remains to be determined. Nevertheless, in combination with our finding that the impact on a secondary infection is to increase bacterial numbers approximately 10-fold, our results indicate that this process is a mechanism driven by pneumococcus for its own benefit, and not a host stress response.

Bacteria-induced histone modifications have often been associated to virulence mechanisms as mostly pathogenic bacteria have been studied. However, we have recently shown that an asymptomatic pneumococcal colonizing strain activates a histone demethylase, KDM6B, to drive a homeostatic host response (8). Disruption of this process leads to host cell damage *in vitro* and *in vivo* dissemination of the asymptomatic strain to the circulation. Although the effect of KDM6B is not on H3K4, as it is not a direct substrate, it is interesting to think that similarly to the histone modification induced by KDM6B, di-methylation could be important to better colonize respiratory epithelial cells. Whether H3K4me2 methylation is a virulence mechanism driving infection, or a means to enhance asymptomatic colonization remains to be determined.

The respiratory epithelium is composed of three major cell types: ciliated, secretory and basal cells. These play critical roles in providing a barrier and defense system against inhaled pathogens (31). The cells in contact with invading bacteria are thought to be terminally differentiated and constantly renewed. In our study, we show that H3K4me2 is induced in terminally differentiated type II pneumocytes (A549 cells), but also in undifferentiated nasal cells (RPMI cells). These results indicate that pneumococcus induced epigenetic modifications occur in precursor cells and therefore have the potential to be transmitted to cells of the epithelium over multiple generations. Furthermore, we show that H3K4me2 is maintained through cell division. We hypothesize that H3K4me2 occurs initially in cells in contact with *pneumococcus* and this mark is transmitted through cell division. Therefore 7 days later, cells display H3K4me2 even if they were not initially in contact with bacteria. Epigenetic transmission is more likely than the possibility that continuous cellular signals perpetuate the mark over time as we no longer detect any live bacteria after 24 hours, and no more bacterial products after 3 days. Furthermore, cell divisions over time ensure that after 7 days cells are in majority renewed. Therefore, we show that *S. pneumoniae* epigenetically modifies epithelial cells during infection and the epigenomic mark is inherited by daughter cells that were not in contact with bacteria, illustrating the potential to modify large numbers of cells.

H3K4 has the potential to be mono-, di- or tri-methylated and the number of methyl groups could define different chromatin states. H3K4me1 is a well-established feature of enhancers, often associated with active and poised enhancers, and H3K4me3 marks the promoters of active genes (32, 33). H3K4me2 is less well defined, and often characterized along with H3K4me3 as studies studying function do so by inactivating a methyltransferase, most often shared between di- and tri-methylation (34, 35). In yeast, H3K4me2 has no clear association with transcriptional status, in animals H3K4me2 was suggested to be involved in transcriptional activation, and in plants the association is with repression (36). As the genomic localization and association with other histone modifications and chromatin features are important for defining the role of a particular mark, it may be that a generalized role for H3K4me2 cannot be made. Alternatively, heterogeneity in the cell population could make bulk analysis difficult to extract overarching features. This is what we find in our analysis, where direct genome-wide association of H3K4me2 peaks with transcriptional activity was not possible on the global level. Instead, unbiased multiple factor analysis allowed us to divide our findings into smaller clusters where a combination of chromatin states, genomic localization and transcriptional regulation allowed us to draw some common features. This type of analysis has not previously been used to link transcriptome and epigenome data, but could be a good solution as the integration of ChIP-seq and RNA-seq data remains difficult and is slowly being addressed (37). Intriguingly, if H3K4me2 was a “true” epigenetic mark, it’s function would be in maintaining chromatin states and not directly on transcriptional regulation.

Epigenetic transcriptional memory has been documented in multiple organisms and thought to play a role in adaptation to changes in environmental conditions. In plants upon drought stress, a subset of genes display an increased transcription rate upon re-exposure to the dehydration, along with a maintenance of high levels of H3K4me3 and phosphorylation of polymerase II at serine 5 (38). Similarly, hyperosmotic priming of *Arabidobsis* seedlings led to enhance drought tolerance in plants correlated with alterations in H3K27me3 profiles (39). In yeast, localization within the nucleus at the periphery, as well as association with the histone variant H2A.Z, allows for robust reactivation (40). In mammalian cells, one report shows that in HeLa cells, certain Interferon-γ regulated genes accumulate persistently high levels of H3K4me2, correlating with facilitated promoter accessibility for multiple cell cycles (41). In all these studies, the histone mark is shown to be transmitted through the cell cycle and maintained through multiple cell divisions. Our results fit perfectly with already published data on epigenetic transcriptional memory. Infection specifically induces H3K4me2, which we identified by screening 21 different histone H3 marks, and is maintained through multiple cell divisions affecting transcription upon secondary infection.

Innate immune memory (also known as trained immunity) is the process by which innate immune cells such as macrophages, monocytes, and natural killer cells, mount a faster and increased response to homologous or heterologous secondary challenge and display associated histone modifications (17). For instance, functional reprogramming of monocytes by *Candida albicans* leading to stronger response and protection from infection, is associated with maintenance of H3K4me3 (42). Natural killer cells remember previous encounters with multiple pathogens (43). Memory to the bacterial factor LPS (44), as well as to infection with *S. pneumoniae* correlates with H3K4me1 at the promoter of genes of interest (45). The best characterized innate immune mechanism is in response to LPS, which depending on the dose either leads to lasting unresponsiveness or hyperresponsiveness (46). Prolonged exposure to LPS induces tolerance as displayed by suppression of inflammatory cytokines secretion, and is associated with H3K9 and DNA methylation (47). At lower doses, LPS induces de *novo* enhancers marked with H3K4me1, which is retained upon signal removal and leads to an augmented secondary response to stimulation (48). Thus, innate immune memory and epigenetic transcriptional memory probably operate along similar mechanistic features. Here we show that bacteria have their own mechanism of inducing transcriptional memory, and therefore pathogen manipulation of innate immune memory needs to be further studied to understand the impact on infectious processes.

By analysis of ENCODE data, one report has found that 90% transcription factor binding sites overlap specifically with H3K4me2 enriched regions, suggesting a tight regulation of H3K4me2 by transcription factor binding (49). In our study, we performed a transcription factor motif search and found enrichment of certain binding sites in MFA cluster dependent manner. This result is interesting as it suggests there are specific genomic regions targeted by H3K4me2, which by Reactome analysis have biological significance for infection. Whether these transcription factors are necessary to initiate memory or maintain it remains to be determined. In yeast, transcriptional memory of the INO1 gene has been shown to depend on H3K4me2 and initiated by a particular transcription factor Sfl1, which binds to a subset of genes upon repression of their transcription (50, 51). Interestingly, our MFA analysis links most gain of methylation peaks with genes downregulated upon primary infection. This could suggest that the transcription factors that we identify could act similarly to Sfl1 in binding to repressed genes thereby initiating memory at these loci. Alternatively, a subset of the identified transcription factors could be maintained after bacterial clearance, thereby maintaining an open chromatin state and transcriptional memory. This mechanism would be similar to what has been reported in a skin inflammation model, where a combination of the transcription factors STAT3 and FOS are transiently induced, but only JUN remains bound to memory domains (24).

In this study, we provide a demonstration that bacteria modify host chromatin in a lasting manner, and for their benefit. Characterizing this process could have important consequences for our understanding of bacterial infection and/or colonization, particularly important for *S. pneumoniae*. Particularly, as epigenetic mechanisms are reversible, targeting the processes exploited by bacteria could be a prolific and novel strategy to develop host directed therapies.

## Materials and methods

### Cell culture condition

The human alveolar epithelial cell line A549 (ATCC CCL-185) was cultured in F-12K culture medium (Gibco™ 21127-030) supplemented with 10% fetal calf serum (FCS) and 1% L-glutamine (Gibco™ 25030-024). The human nasal septum epithelial cell line RPMI 2650 (ATCC CCL-30) cells was cultured in Advanced MEM culture medium (Gibco™ 12492-013) supplemented with 2.5% FCS and 4 mM L-glutamine. The cells were incubated at 37 °C in a humidified atmosphere with 5% CO_2_. For assays (Figure 1A), 1×10^5^ A549 cells/mL or 2×10^5^ RMI 2650 cells/mL were seeded in 6-well plates 2 days before infection (-48h). When A549 cells were grown to semi-confluence, they were serum-starved (0.25% FCS) for 24h before use in experiments and for RPMI 2650 cells, culture medium (2.5 % FCS) has only been renewed.

### *Streptococcus pneumoniae* culture and cell infection

Experimental starter stocks of *Streptococcus pneumoniae* TIGR4 strain were prepared on 5% Columbia blood agar plates (Biomerieux 43041) from frozen permanent stocks. Bacteria are grown Todd–Hewitt (BD 249240) broth supplemented with 50 mM HEPES (Sigma H3375) (TH+H) at 37°C with 5% CO_2_ to mid-log phase 0.6 (OD_600_). Aliquots were made in TH+H media supplemented with Luria–Bertani (BD 244620) and 15% glycerol final concentration then frozen at −80 °C.

All experiments were performed with frozen experimental starters of *S. pneumoniae* (*Spn*) less than 1 month old. For experiments, starters were grown to mid-log phase (0.6 OD_600_) in TH+H broth at 37°C with 5% CO_2_, washed twice in PBS and concentrated in 1 ml PBS. Bacteria were diluted in the serum-low cell culture medium (0.25% FCS) for A549 cells or in the cell culture medium (2.5% FCS) for RPMI 2650 to get a multiplicity of infection of 20:1 (MOI 20), 10:1 (MOI 10) or 35:1 (MOI 35) according to experiment.

The first week, after 1h or 3h of *Spn* infection, cells were washed twice with PBS to remove the unattached bacteria and cultured in medium containing an antibiotics cocktail (penicillin-streptomycin (10 μg /ml) and gentamicin (200 μg /ml)) for 5 days (5d) (Figure 1A). The control cells uninfected were treated in the same way as the infected cells. 2 days (2d) after infection or no-infection, the cells were recovered with PBS supplemented with 10 mM EDTA, then 1×10^6^ cells were seeded in flask (Corning™ 430641U) with medium including the antibiotics cocktail.

For inactivated *Spn*, the pellet of bacteria was incubated in 4% PFA for 10 min at room temperature, then washed twice in PBS before infection.

The second week at 5 days (5d) for the experimental conditions, uninfected cells (UI), 1° infection (1°), 2° infection (2°) and cells maintained post 1° infection (PI), 1×10^5^ A549 cells/mL or 2×10^5^ RMI 2650 cells/mL were seeded in each well of 6-well plates. 24h before infection or no-infection, A549 cells were serum-starved (0.25% FCS) and for RPMI 2650 cells, the culture medium (2.5%) has been renewed. After 1h or 3h of *Spn* infection, cells were washed twice with PBS to remove the unattached bacteria and then either collected for analysis, or either cultured in medium containing the antibiotics cocktail. Samples were collected for analysis at different times.

To determine the amount of *Spn* that have colonized cells, PBS washed cells were lysed using sterile ddH2O. Lysates and serial dilutions were plated on 5% Columbia blood agar plates overnight at 37°C with 5% CO_2_, and the CFUs were counted.

### Cell proliferation with CellTrace™ CFSE by Cytometry

24 h after infection or no-infection, cells were incubated at 37°C with 5% CO_2_ in serum-low cell culture medium with penicillin-streptomycin (10 μg /ml) and gentamicin (200 μg /ml) containing 2.5 mM CellTrace^TM^ CFSE (Invitrogen™ C34554). After 6h, 3 days (3d) and 5 days (d), the cells were recovered with PBS supplemented with 10 mM EDTA. Cells were washed in PBS in preparation for viability staining using fixable viability dye (eFluor780, ebioscience) for 5 minutes at 4°C. Cell were washed in PBS and fixed using commercial fixation buffer (Biolegend). After washes, cells were resuspended in PBS, and the acquisition of the different samples was performed on MACSQuant (Miltenyi Biotec) flow-cytometer and analysis was completed using FlowJo Software.

### Cell viability alamarBlue assay

3 h after infection, A549 cells were incubated at 37°C with 5% CO_2_ in serum-low cell culture medium with antibiotics cocktail (penicillin-streptomycin (10 μg /ml) and gentamicin (200 μg /ml)) containing 10% alamarBlue reagent (Molecular Probes™ DAL1025) for 12 h. One measure of fluorescence (Ex/Em 560/590 nm) was then read each hour (h) using a Cytation 5 (BioTek) driven by the Gene 5.1.11 Software. Redox of alamarBlue (Resazurin) produces fluorescence whose intensity is measured. Fluorescence readings were blank corrected to wells containing only culture medium and results are expressed as fluorescence intensity.

### Immunoblotting and quantification

Cells were lysed using laemmli buffer (4% SDS, 20% glycerol, 200mM DTT, 0.01% bromophenol blue and 0.1 M Tris HCl, pH 6.8). Samples were sonicated for 4 sec, boiled for 10 min. Proteins were separated on 15% SDS-PAGE and were transferred to PVDF membrane under 2.5A/25V condition for 7 min using a semidry transfer system (Trans-Blot Turbo, BioRad). Transferred membranes were first blocked by TBS-Tween20 (0.1%) with 5% milk or 5% BSA, and then incubated overnight at 4°C with primary antibodies for H3K4me2 (1:2000; Abcam ab32356), H3K4me3 (1:1000; Diagenode C15410003), H3S10ph clone MC463 (1:2000; Millipore 04-817), anti-γH2A.X (S139) (2OE3) (1:1000; CST 9718S), anti-H2A.X (1:1000; CST 2595S) or actin AC-15 monoclonal (1:10.000; Sigma A5441). Membranes were washed with TBS-Tween20 (0.1%) and incubated with secondary-HRP conjugated antibodies (1:10.000; Anti Rabbit IgG HRPO, Millipore BI2407; Anti Mouse IgG HRPO, Millipore BI2413C) for 1h at room temperature. After washing with TBS-Tween20 (0.1%), the immunoreactive bands were visualized using Clarity ECL substrate (BioRad). Immunoblot signal was acquired on a ChemiDoc Imaging Systems (BioRad) and analyzed using Image Lab software (BioRad).

### Histone H3 modification Multiplex assay

For cells infected and uninfected, the histone extraction kit (OP-0006) and the EpiQuik^TM^ Histone H3 modification Multiplex assay kit (P-3100) were purchased from Epigentek and protocol followed according to manufacturers’ instructions. 100 ng of total histone proteins extracted from A549 cells was used. We analyzed twenty-one modified histone H3 patterns simultaneously in histone extracts from cells at 3 h (1°) and 7 days (PI) after infection compared to uninfected cells (UI).

### Immunofluorescence microscopy and Fiji or CellProfiler analysis

Cells were grown on acid-washed and ultraviolet-treated coverslips in 6-well plates (TPP 92006). After infection, cells were washed 2 times with PBS and fixed in 2.5% paraformaldehyde for 10 min at room temperature. After 3 washes, cells were blocked in 5% BSA for 6h at +4°C, then permeabilized overnight in 5% BSA 0.5% Tween20 at +4°C. Immunostaining was performed overnight at +4°C with primary antibodies (H3K4me2 1:1000; Abcam ab32356) (H3K4me3 1:200; Diagenode C15410003) (H3K4me2 1:1000; EpiGentek A4032) (LAMP1 1:1000; Abcam ab32356) diluted in 5% BSA 0.5% Tween20, then washed three times in 0.5% Tween20 and three times in PBS. Cells were incubated for 1h at room temperature with Alexa Fluor 488 secondary antibody (1:1000; Invitrogen^TM^ A11034) and Alexa Fluor 647 phalloidin (1:1000; Invitrogen^TM^ A22287) in 5% BSA 0.5% Tween20, then washed three times in 0.5% Tween20 and three times in PBS. Cells were incubated for 5 min at room temperature with 4,6-diamidino-2-phenylindole (DAPI) (300 nM; Invitrogen^TM^ D1306) diluted in PBS, then washed three times in 0.5% Tween20 and three times in PBS. The coverslips were rinsed briefly in distilled water and were mounted on slides using ProLong^TM^ Gold antifade reagent (Invitrogen^TM^ P36930). For quantification, all images were acquired using Cytation 5 (BioTek) driven by the Gene 5.1.11 Software and processing was done with ImageJ or CellProfiler software.

### Statistical analysis

Statistical tests are reported in the figure legends. Appropriate parametric or nonparametric tests were used. Data plots and statistics were generated using Prism (version 9, GraphPad Software Inc.).

### RNA samples and sequencing

RNA was extracted from cells using RNeasy Mini Kit (Qiagen, 74104) and protocol followed according to manufacturers’ instructions. Chloroform extraction/Na-Acetate 3M (pH 5.5), isopropanol precipitated and washed 3 times in 70% ethanol prior to being suspended in molecular grade water. Extracted RNA quality was assessed and quantified with RNA 6000 Nano kit (Agilent, 5067-1511) using 2100 Bioanalyzer instrument (Agilent). The Affymetrix Human Gene array 2.1 microarrays was performed by Eurofins Genomics (France).

### Transcriptome analysis

Expression profile of the complete time course, i.e. UI, 1°, 2° infections and PI in triplicates, was measured using the Affymetrix Human Gene array 2.1 microarray. Data were deposited into the Gene Expression Omnibus (GEO) repository of the National Center for Biotechnology Information under accession number <u>GSE230142</u>.

Expression signal was normalized using the Robust Multichip Average (RMA) method, from the affy R package (doi: <u>10.1093/bioinformatics/btg405</u>). Control and lowly express probes were filtered out and the replicate effect was removed using the ComBat approach implemented in the sva R package (52). Replicate 3 from 1° infection was filter out after principal component analysis. Differential expression analysis was performed using the limma method (53) and the following comparison evaluated: 1° infection vs UI cells, 2° infections vs PI and (2° vs PI) vs (1° vs UI), i.e. 2° infection vs 1° infection. Genes with an adjusted p-value < 0.05 were defined as differentially expressed (DEGs).

For the functional characterization, we performed a Gene Set Enrichment Analysis (GSEA) for each comparison, by ranking the genes with the t-statistic, which is the log2 fold change divided by its standard error. We used the Reactome database gene sets and the GSEA implementation in https://gitlab.pasteur.fr/hvaret/fgsa_scripts.

### ChIP sequencing assays

After infection or no-infection, the cells were washed with PBS in each well of 6-well plates (TPP 92006). Cells were cross-linked with 1% formaldehyde 8 min at room temperature, followed by quenching with 0.125 M glycine for 5 min. After two washes in PBS, cells were collected by scraping, then pelleted and lysed on ice for 5 min in 0.25% Triton X-100, 10 mM Tris-HCl (pH 8), 10 mM EDTA, 0.5 mM EGTA and proteases inhibitors. The soluble fraction was eliminated by centrifugation and chromatin was extracted with 250 mM NaCl, 50 mM Tris-HCl (pH 8), 1 mM EDTA, 0.5 mM EGTA and proteases inhibitors cocktail for 30 min on ice. Chromatin was resuspended in 1% SDS, 10 mM Tris-HCl (pH 8), 1 mM EDTA, 0.5 mM EGTA and protease inhibitor cocktail, then fragmented by sonication (10 cycles of 30 sec ‘on’ and 30 sec ‘off’) using Bioruptor Pico (Diagenode). Sheared chromatin was cleared by centrifugation, and tested for shearing efficiency analysis with High Sensitivity DNA kit (Agilent, 5067-4626) using 2100 Bioanalyzer instrument (Agilent). 2 μg of antibodies ChIP-grade (H3K4me2, Abcam ab32356; H3K4me3, Diagenode C15410003; H3, Abcam ab1791; Normal Rabbit IgG, Abcam ab37415) were used per ChIP and were bound to DiaMag protein G-coated magnetic beads (Diagenode, C03010021) overnight at 4°C with gentle rotation. 6-7 μg/IP experimental chromatin were diluted 10 times in 0.6% Triton X-100, 0.06% sodium deoxycholate (NaDOC), 150 mM NaCl, 12 mM Tris-HCl, 1 mM EDTA, 0.5 mM EGTA and proteases inhibitors cocktail. 2% of ChIP sample volume was reserved to serve as input. Diluted chromatin was then added to antibody bound DiaMag beads and incubated at 4°C overnight with gentle rotation. ChIP samples were washed sequentially 5 min with buffer 1 (1% Triton X-100, 0.1% NaDOC, 150 mM NaCl, 10 mM Tris-HCl (pH 8)), buffer 2 (0.5% NP-40, 0.5% Triton X-100, 0.5 NaDOC, 150 mM NaCl, 10 mM Tris-HCl (pH 8)), buffer 3 (0.7% Triton X-100, 0.1% NaDOC, 250 mM NaCl, 10 mM Tris-HCl (pH 8)), buffer 4 (0.5% NP-40, 0.5% NaDOC, 250 mM LiCl, 20 mM Tris-HCl (pH 8), 1 mM EDTA), buffer 5 (0,1% NP-40, 150 mM NaCl, 20 mM Tris-HCl (pH 8), 1 mM EDTA) and buffer 6 (10 mM Tris-HCl (pH 8), 1 mM EDTA). ChIP and input samples were treated with IPure Kit (Diagenode, C03010015) following manufacturers’ instructions. ChIP and input DNA were quantified with Qubit 4 fluorometer (ThermoFisher). Biomics Platform, C2RT, Institut Pasteur generated the libraries with 10 ng of each sample and sequencing was performed with Illumina Hiseq2500.

### Methylome analysis

H3K4me3 and H3K4me2 were profiled using ChIPseq for the UI cells, 1° infection (3h) and PI (7d) in duplicates. Complete analysis from the raw sequencing data to the differential marking analysis was done following the ePeak approach (54). Default parameters were used for the filtering, mapping of reads and for narrow peak calling. For H3K4me2, reproducible peaks were determined using the intersection approach, requiring at least 40% length overlap between replicates. For H3K4me3, reproducible peaks were determined using the IDR method. Data were deposited into the Gene Expression Omnibus (GEO) repository of the National Center for Biotechnology Information under accession number <u>GSE230142</u>.

Differential analysis was performed using the limma method (53) with a model considering the biological factor of interest, i.e. time (UI, 1° infection and PI) and the batch effect, i.e. replicate. Prior to the statistical test, the systematic differences between samples due to technical variation such as sequencing depth were normalized using the cyclic lowness method (i.e. locally fitting a smooth curve).

Time course profiles were calculated using the read counts distribution over the dynamically marked regions (DMRs): adjusted p-value < 0.1 and log fold change > abs (0) for H3K4me3, adjusted p-value < 0.3 and log fold change > abs (0) for H3K4me2. Read counts over DMRs per time point and replicate were normalized to account for systematic technical variation (sequencing depth and replicate), differences in region length and they were subsequently scaled. Finally, DMRs were clustered using the Euclidean distance and the Ward agglomerating method to define the two main profiles: Gain and Loss of marking.

DMRs were classified according to their localization with respect to the transcriptional starting site (TSS) of the closest gene in: TSS if they overlap a window of 2Kbp centered on the TSS; intragenic if they overlap the gene annotation but not the TSS window; intergenic if they overlap noncoding regions between gene annotations and don’t overlap any TSS window.

An epigenomic characterization of H3K4me2 DMRs was done for the UI condition using publicly available information for the cell line in the ENCODE portal (https://www.encodeproject.org/, (55)). We downloaded the call sets with the following identifiers: ENCFF628ANV, ENCFF510LTC, ENCFF556OVF, ENCFF250OYQ, ENCFF189JCB, ENCFF421ZNH (H3K4me3); ENCFF876DTJ, ENCFF975YWP, ENCFF419LFZ (H3K4me2); ENCFF137KNW, ENCFF103BLQ, ENCFF663ILW (H3K27ac); ENCFF165ZPD, ENCFF569ZJQ, ENCFF613NHX (H3K4me1); ENCFF892LYD, ENCFF154HXA, ENCFF931LYX (H3K79me2); ENCFF819QSK, ENCFF134YLO, ENCFF336AWS (H3K27me3). Coverage over peaks was normalized by length and scaled. Peaks with similar epigenomic profiles were clustered using the Euclidean distance and the Ward agglomerating method.

### Integrative analysis

Joint analysis of the complete dataset was performed by Multiple factor analysis (MFA) (21).

Firstly, we predicted the regulatory links between DEs and DMs using T-gene (23) and selected the most significant links (distance < 500Kb and correlation p-value < 0.05). The joint dataset contains the 3874 links holding either a differentially expressed gene (DEs) and/or a dynamically marked region (DMs) and either a not DMs and/or a not DEs. For each link we defined five groups of continuous variables:

- Transcriptome (TOME) consisting of the gene expression over the four time points (UI, 1°, 2° infections and PI) in triplicates;
- H3K4me2 methylome (METH) measured by the average methylation over le three time points (UI, 1°, PI) and the log fold change between PI and UI cells;
- Epigenome of the UI condition (EPIG) including the coverage of the above-mentioned histone modifications over the H3K4me2 regions;
- Distance (DIST) and correlation (CORR) of the regulatory link between genes and regions;

and two groups of categorical variables:

- DEG kind and sing, describing whether genes are specific of the 1° or 2° infection, or Not specific; and whether genes are going Up, Down, UpDown or DownUp along the time course;
- DMR kind and sing, indicating whether regions are DM or NotDM; and whether regions belong to the Gain or Loss of H3K4me2 profile along the 1° infection time course.

The chromatin state (CS) of each H3K4me2 region, calculated from the EPIG group as described in the previous section, was added as a supplementary variable to facilitate the interpretation of the MFA dimensions/factors. To control for the within group variability, all continuous variables were normalized and scaled. MFA was performed and a clustering of the 3874 was obtained using the top 10 dimensions/factors. Each cluster was then defined as a combination of the continuous and categorical variables more significantly associated to the genes and regions of the corresponding links (v.test statistic > 5).

### Transcription factor motif enrichment analysis

Transcription factor motif enrichment analysis was performed on the previously defined MFA clusters 1, 2, 4, 5 and 9, separating the peaks corresponding to a gain or loss of methylation. For each peak, 500bp were extracted around the summit and then used as input for MEME-chip V5.1.1 (56). For the analysis, the HOCOMOCO v11 was used as target database for the motifs and the parameter ccut (maximum size of a sequence before it is cut down to a centered section) was set to 0. All the other parameters were the default parameters of the MEME-chip web version.

**Figure S1:**
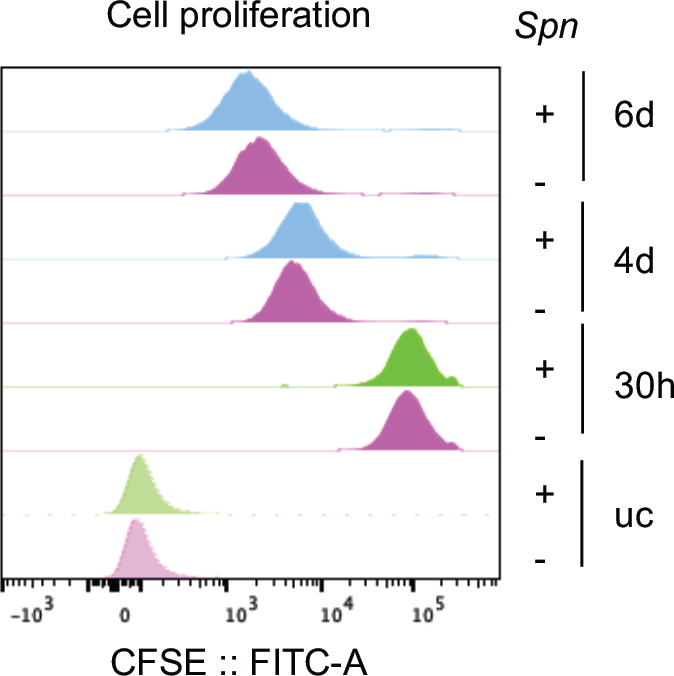
Analysis of A549 cell proliferation. Every generation of cells appears as a different peak on flow cytometry histogram for uninfected (-) and infected with *Spn* (+) cells (MOI 20) at 30 hours (30h), 4 days (4d) and 6 days (6d) post infection. Unstrained control (uc) for uninfected (-) and infected (+) cells.

**Table S1:**
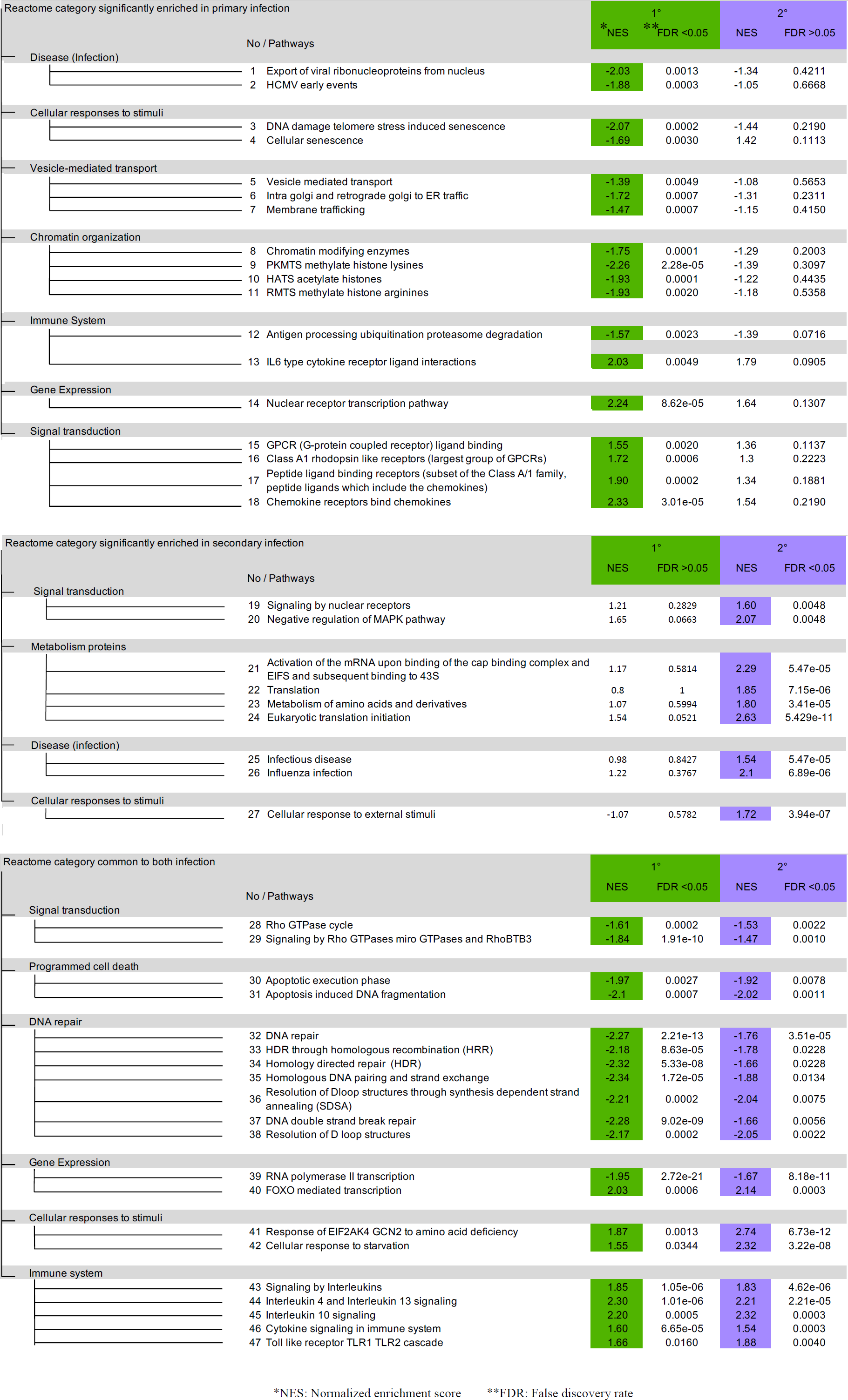
Reactome pathways and corresponding categories Figure S2: Immunoblot of γH2Ax levels. Representative image of immunoblot detection of γH2Ax from infected and uninfected A549 cells at 3h and 24h post infection, at primary infection (1°), PI (7d or 8d) and secondary (2°) infection.

**Figure S2:**
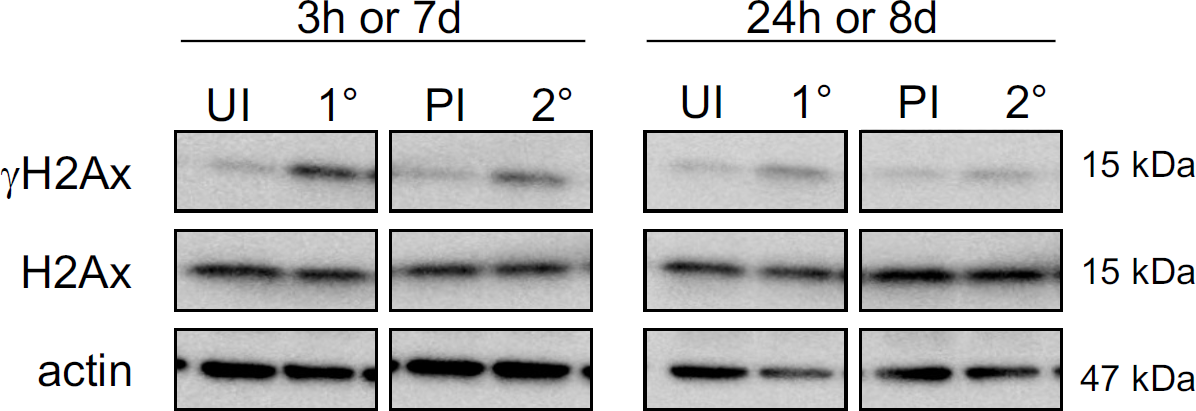
Immunoblot of γ levels. Representative image of immunoblot detection of γ from infected and uninfected A549 cells at 3h and 24h post infection, at primary infection (1°), PI (7d or 8d) and secondary (2°) infection.

**Figure S3:**
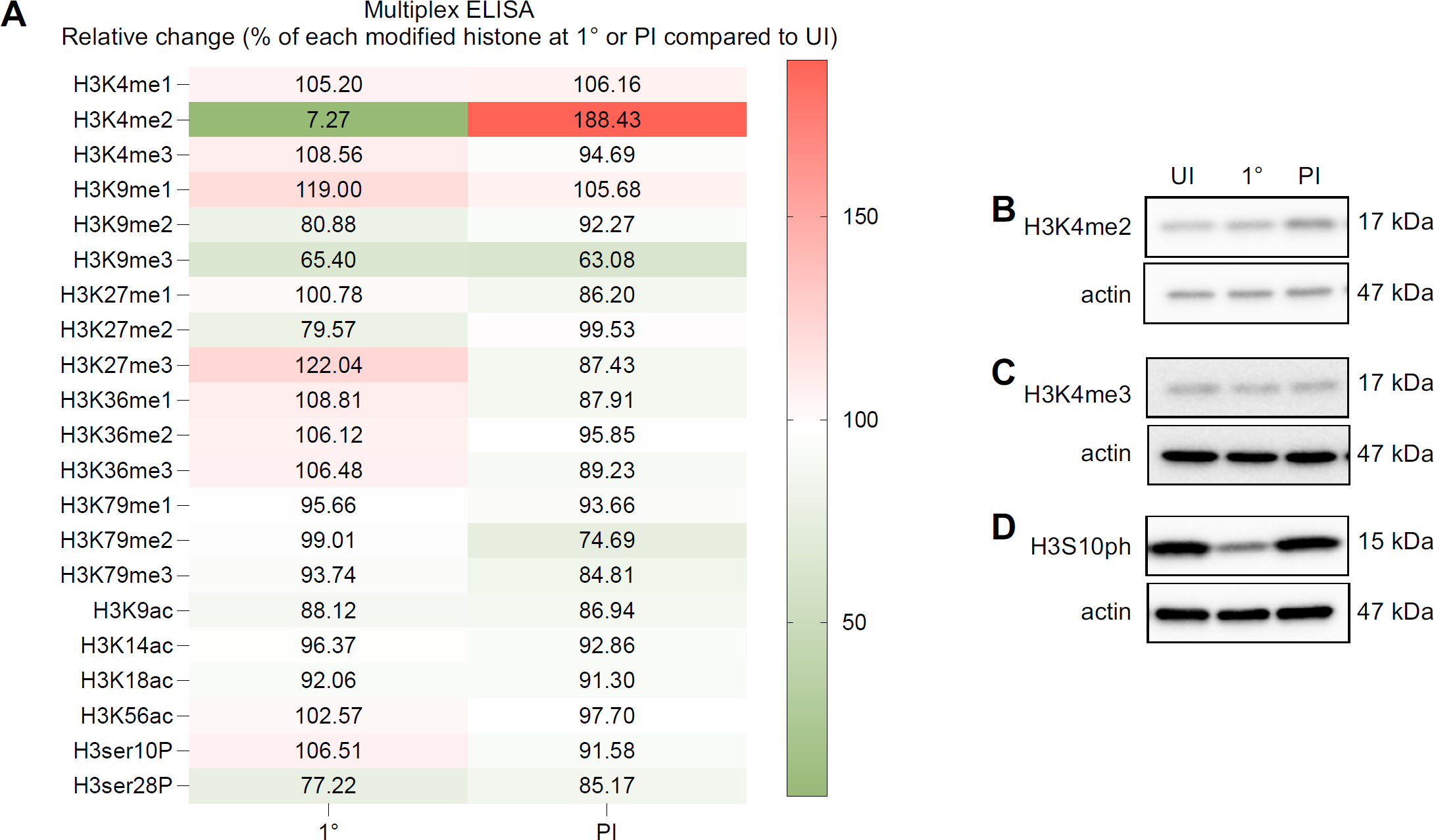
H3K4me2 is a specific mark lasting beyond primary infection. **(A)** Histone H3 modifications by Multiplex ELISA assay from infected cells at uninfected, 1° infection (3h) and PI (7d). 100 ng of total histone proteins extracted A549 cells were used. Quantitation of twenty-one modified histone H3 patterns simultaneously. Histone H3 modifications by Multiplex ELISA assay. Quantitation of twenty-one modified histone H3 patterns simultaneously. Table heatmaps representing total H3-normalized ratio of the relative. change (%) of each histone H3 modification between 1° (3h) or PI (7d) and uninfected cells. **(B)** Representative image of Immunoblot detection of H3K4me2 and actin from uninfected (UI) and infected whole cells lysates of A549 cells at 1° infection (3h) and PI (7d) and actin. **(C)** Representative image of Immunoblot detection of H3K4me3 and actin from uninfected (UI) and infected whole cells lysates of A549 cells at 1° infection (3h) and PI (7d). **(D)** Representative image of Immunoblot detection of H3S10ph and actin from uninfected (UI) and infected whole A549 cells lysates at 1° infection (3h) and PI (7d).

**Figure S4:**
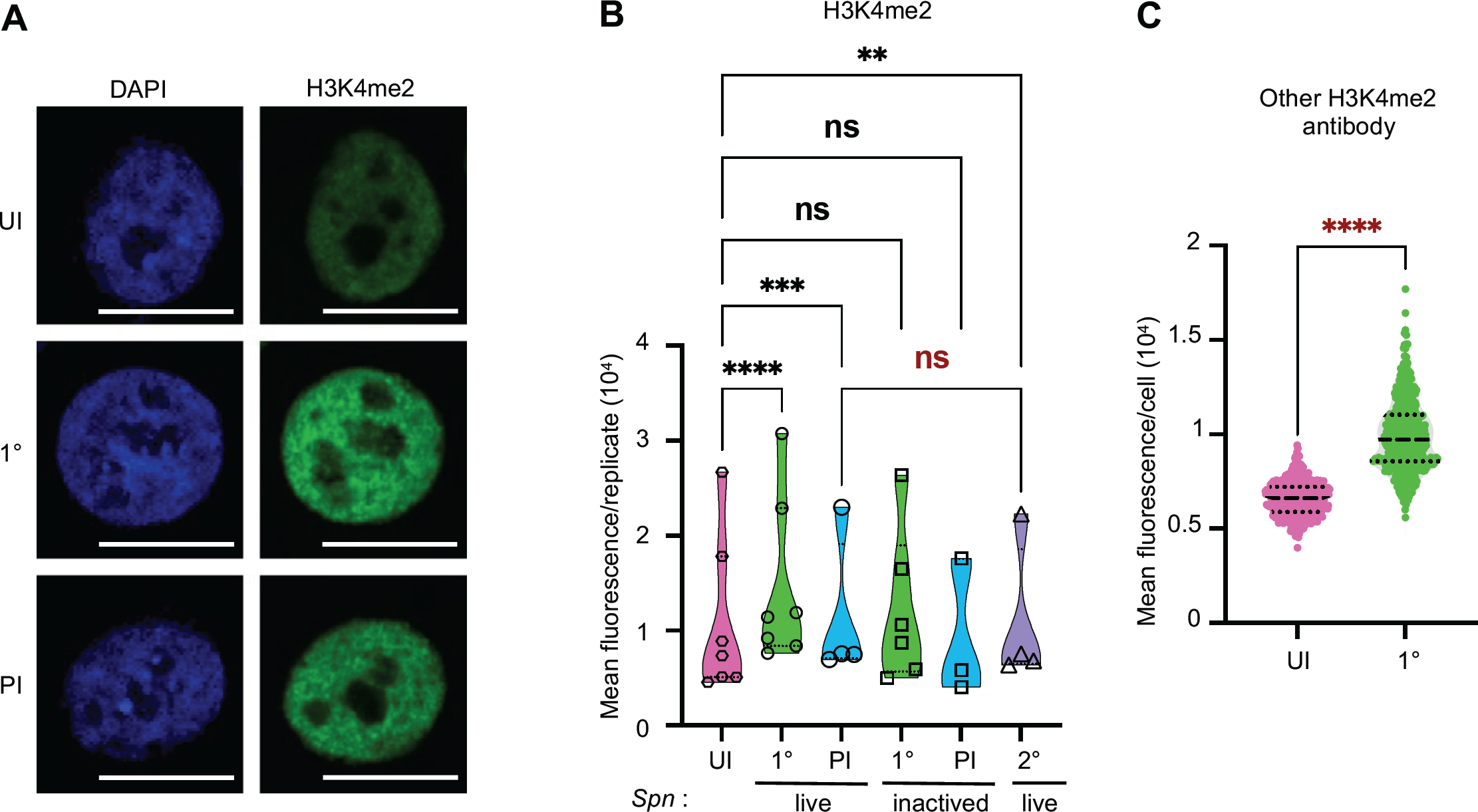
The increase in H3K4me2 levels is actively induced by live bacteria. **(A)** Representative images of immunofluorescence confocal microscopy detection of H3K4me2 in nuclear of paraformaldehyde fixed A549 cells uninfected and infected at 1° infection (48h) and PI (9d). Cells stained for H3K4me2 (GFP; green) and nuclei (DAPI; blue). Crop of images taken at 63x magnification. Scale bar is 10μm. **(B)** Quantification of H3K4me2 normalized to the segmented nuclei using DAPI signal. Data points expressed mean fluorescence intensity for each biological replicates for uninfected cells (UI), for infected cells (MOI 20) at 1° infection (48h) and PI (9d) with *Spn* live and *Spn* inactived, and at 2° infection (48h) with *Spn* live. Violin plot (truncated) show all points, statistical significance was determined by ANOVA with matching across each biological replicate and Tukey’s multiple comparisons test with a single pooled variance (ns = not significant, **p = 0.005, ***p <0.0009, ****p <0.0001). **(C)** Quantification of nuclear H3K4me2 from RPMI 2650 cells normalized to the segmented nuclei using DAPI signal Data points represent the mean fluorescence intensity (MFI) of H3K4me2 (Epigentek antibody) within individual nuclei. Violin plot (truncated) show 500 nuclei at 1° infection (MOI 10, 48h) and uninfected cells (UI), statistical significance was determined by Mann-Whitney test (****p <0.0001).

**Figure S5:**
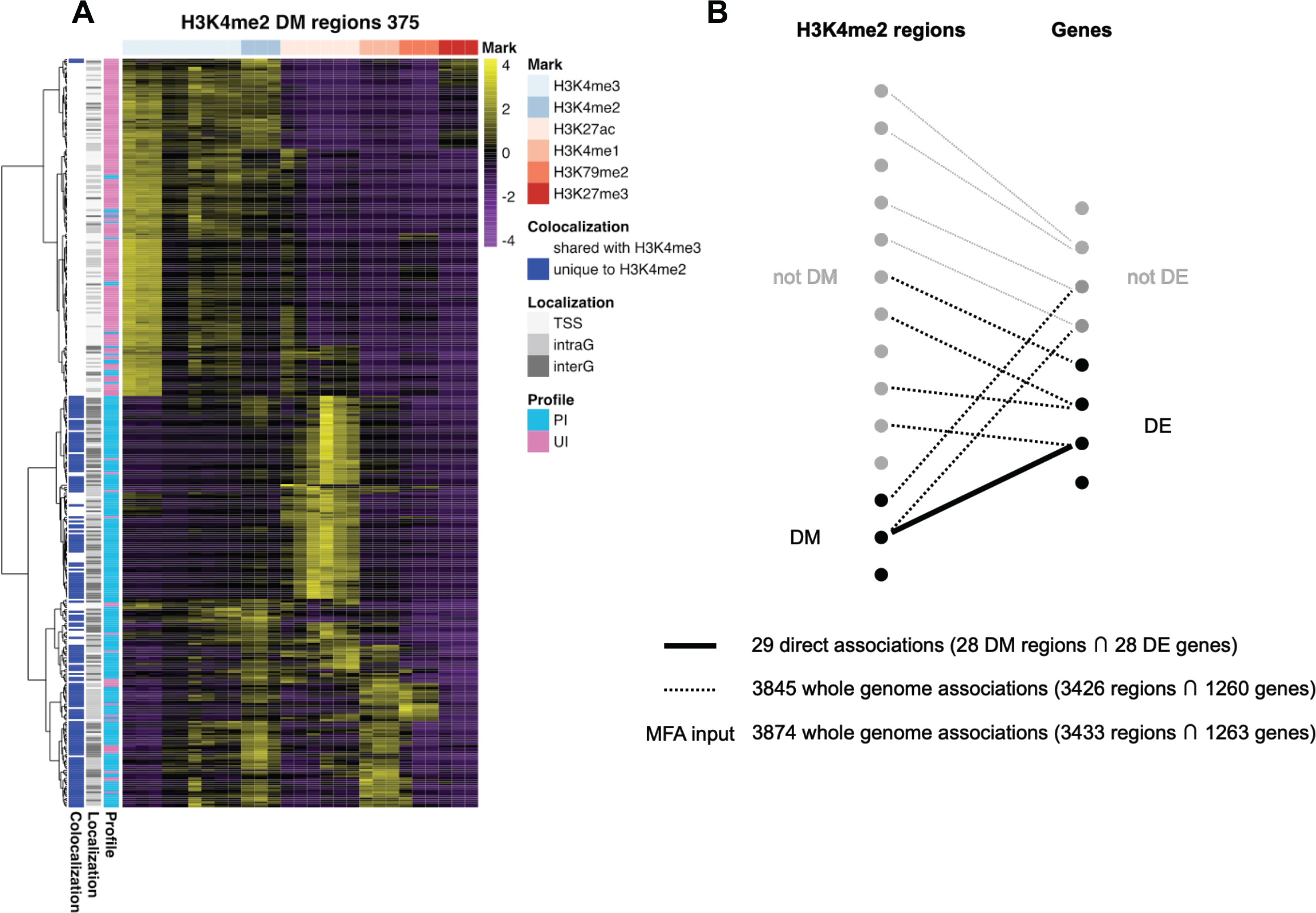
Chromatin profiling over H3K4me2 differentially methylated peaks and regulatory association to genes. **(A)** Chromatin profiling over the H3K4me2 dynamic peaks. Coverage for key histone modifications recovered from the ENCODE portal for A549 cells is shown for the 375 H3K4me2 dynamic peaks (DM). Histone coverage is normalised by peak length, centred and scaled among samples. Peaks are clustered according to their chromatin state and annotated according to their profile: PI (gain of methylation) or UI (loss of methylation); their localization with respect to the nearest gene (TSS = overlapping the 2 Kb interval centred around the transcription start site; IntraG = located within the gene annotations and outside the TSS interval; InterG = all other peaks); the colocalization with H3K4me3 dynamic peaks (shared or unique). **(B)** Association between methylome and transcriptome. Dots represent H3K4me2 regions or genes and lines are regulatory links predicted by the T-Gene tool of the MEME suite. Black lines show regulatory links where either a DMs and/or a DEs (continuous lines) or either a not DMs and/or a not DEs (dotted lines) are involved. These account for the 3874 (29 + 3845) associations used as input for the multiple factor analysis (MFA). Grey lines represent regulatory links between not DMs and not DEs and are therefore not included in the downstream integrative analysis.

**Figure S6:**
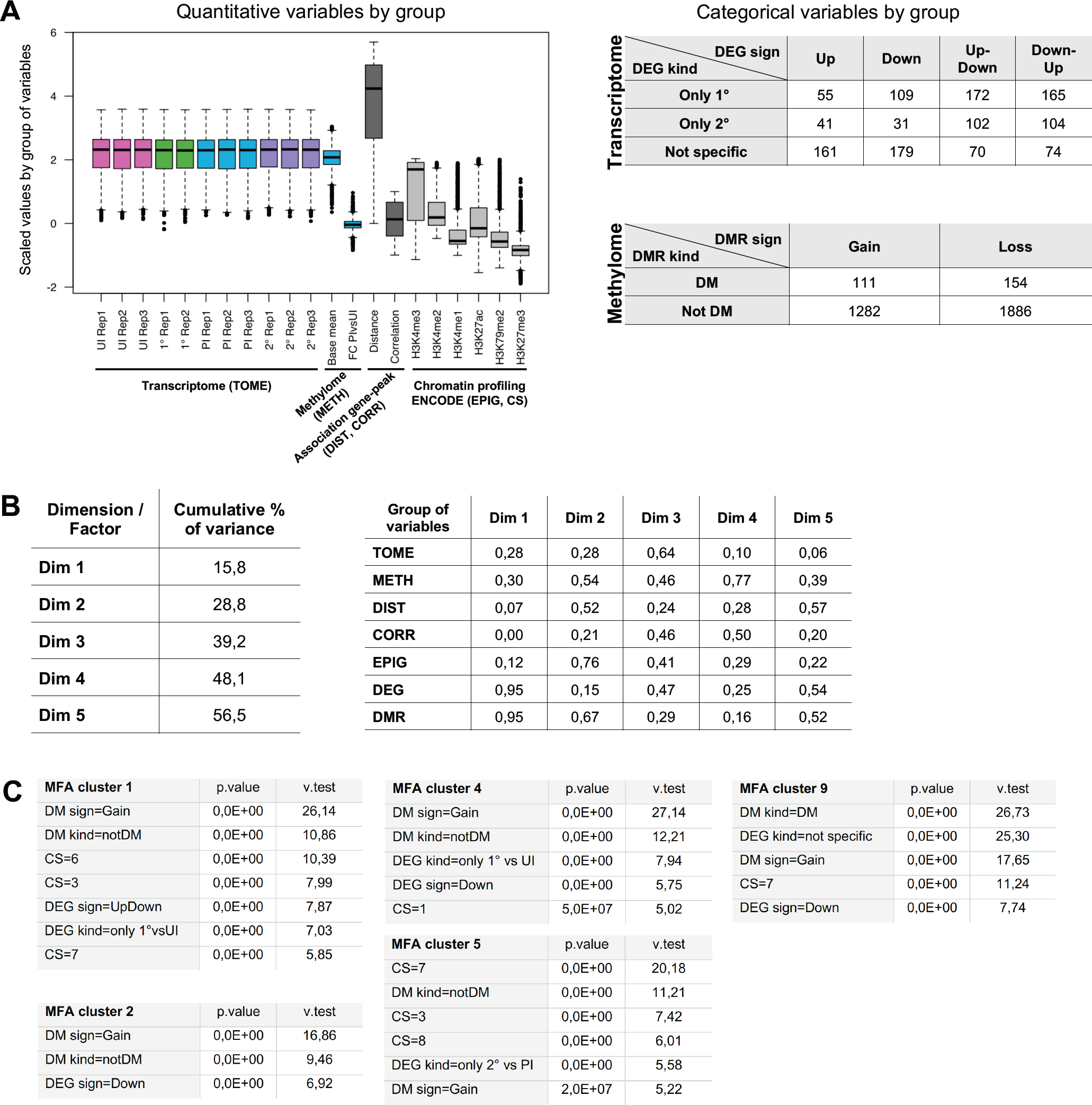
Multiple Factor Analysis (MFA) input data and output metrics. **(A)** Groups of variables used for MFA. Groups of variables constitute the MFA input matrix and describe the genes (TOME = Transcriptome, DEG = Differentially expressed gene), the H3K4me2 peaks (Methylome = METH, DMR = Differentially methylated region, EPIG = Chromatin profile, CS = Chromatin state) and their association (DIST = distance, CORR = correlation). Box plots of quantitative variables after normalization and scaling (left). Tables with number of genes/peaks classified according to the differential expression/methylation analysis (right). **(B)** Quantitative description of factors/dimensions. Cumulative percentage of input dataset variance explained by first five factors (left). Correlation coefficients of groups with the first five factors (right). **(C)** Description of clusters in terms of significant categorical variables.

## Author contributions

C.Chevalier coordinated the study. C.Chevalier and M.A.H conceived and designed all experiments; C.Chevalier performed experiments and analyzed data; C.Chica supervised all the bioinformatic analysis and performed the transcriptome and integrative analysis; J.M. and M.G.C. contributed some experiments and analysis; A.P. performed the bioinformatic analysis for ChIP-seq and MEME-chip. C.Chevalier, C.Chica and M.A.H. conceived and write the manuscript. M.A.H. supervised the work and secured the fundings. All authors approved the final manuscript.

## Acknowledgements

We would like to thank Tiphaine Marie-Noelle Camarasa for her help with the acquisition of the samples performed on MACSQuant flow-cytometer. We thank the Biomics platform, C2RT, Institut Pasteur in particular Juliana Pipoli Da Fonseca for processing the libraries and the sequencing of ChIP-seq. We were thankful to all members of the Chromatin and Infection Unit and the GORE expertise group of the Hub for their precious advice throughout the project and for critical reading of this manuscript. We thank the UTechS PBI platform, C2RT, Institut Pasteur, for giving us access to the LSM780 confocal microscopy driven by Zen software (Zeiss) to acquire some images. We would like to thank Thomas Kohler for providing of *Streptococcus pneumoniae* TIGR4 strain used in this study.

Work in the Chromatin and Infection Unit (headed by Melanie Hamon) is supported by the Institut Pasteur, the Agence Nationale de la Recherche (ANR 17 CE12 0007 01 EPIBACTIN), the Fondation pour la Recherche Médicale (FRM608 EQU202003010152), the Fondation iXCore-iXLife, the Don Prix CANETTI 2020, the EMBO Young Investigator Program. M.A.H. is a member of the Laboratoire d’Excellence “Integrative Biology of Emerging Infectious Diseases” Agence Nationale de la Recherche (ANR-10-LABX-62-IBEID). M.G.C. is supported by a Springboard to Independence grant (AirwayStasis) from the French Government’s Investissement d’Avenir program, the Laboratoire d’Excellence ‘‘Integrative Biology of Emerging Infectious Diseases” (ANR-10-LABX-62-605IBEID) and Pasteur-Weizmann research fund. Bioinformatics and Biostatistics Hub is supported by Institut Pasteur.

