## Supplemental Table 2 for "Epithelial cells maintain memory of prior infection with *Streptococcus pneumoniae* through di-methylation of histone H3"

MFA_SuppTable

| peakIDgene_association | peakID | SYMBOL | ENTREZID | Distance | Correlation | CS | DEG_kind | DEG_sign | DM_kind | DM_sign | MFAclust |
|---|---|---|---|---|---|---|---|---|---|---|---|
| **1_10002631_10003225_116362** | 1_10002631_10003225 | RBP7 | 116362 | 4.73279569828933 | -0.67411576 | 1 | only_DIvsD7 | UpDown | notDM | Gain | 2 |
| **1_10093388_10095341_116362** | 1_10093388_10095341 | RBP7 | 116362 | 4.55769964435121 | -0.48739825 | 1 | only_DIvsD7 | UpDown | notDM | Gain | 4 |
| **1_10531319_10532459_116362** | 1_10531319_10532459 | RBP7 | 116362 | 5.67583056382001 | -0.91684049 | 1 | only_DIvsD7 | UpDown | notDM | Loss | 3 |
| **1_109126041_109129440_113802** | 1_109126041_109129440 | HENMT1 | 113802 | 4.87067891038296 | 0.77888146 | 3 | only_H3vsUI | DownUp | notDM | Gain | 5 |
| **1_109233541_109234680_113802** | 1_109233541_109234680 | HENMT1 | 113802 | 4.47420169019139 | 0.60201219 | 2 | only_H3vsUI | DownUp | notDM | Loss | 6 |
| **1_109235717_109238358_113802** | 1_109235717_109238358 | HENMT1 | 113802 | 4.50481055315069 | 0.60201219 | 1 | only_H3vsUI | DownUp | notDM | Gain | 4 |
| **1_109261889_109263630_22911** | 1_109261889_109263630 | WDR47 | 22911 | 5.50659702273885 | 0.80620105 | 3 | only_H3vsUI | DownUp | notDM | Gain | 1 |
| **1_109372459_109374149_22911** | 1_109372459_109374149 | WDR47 | 22911 | 5.32335318308465 | 0.76230724 | 3 | only_H3vsUI | DownUp | notDM | Loss | 5 |
| **1_109582724_109583699_22911** | 1_109582724_109583699 | WDR47 | 22911 | 2.99956548822598 | 0.84507643 | 1 | only_H3vsUI | DownUp | notDM | Loss | 3 |
| **1_109654733_109655421_113802** | 1_109654733_109655421 | HENMT1 | 113802 | 5.65416787516103 | 0.97100718 | 4 | only_H3vsUI | DownUp | notDM | Loss | 5 |
| **1_109654733_109655421_57535** | 1_109654733_109655421 | ELAPOR1 | 57535 | 3.11693964655076 | -0.04306847 | 4 | only_H3vsUI | UpDown | notDM | Loss | 3 |
| **1_109722412_109725460_22911** | 1_109722412_109725460 | WDR47 | 22911 | 5.13889293738991 | -0.66757784 | 7 | only_H3vsUI | DownUp | notDM | Gain | 5 |
| **1_109722412_109725460_57535** | 1_109722412_109725460 | ELAPOR1 | 57535 | 3.56960796754682 | 0 | 7 | only_H3vsUI | UpDown | notDM | Gain | 1 |
| **1_109739143_109741537_22911** | 1_109739143_109741537 | WDR47 | 22911 | 5.18878228884603 | 0.67239992 | 7 | only_H3vsUI | DownUp | notDM | Gain | 5 |
| **1_109739143_109741537_57535** | 1_109739143_109741537 | ELAPOR1 | 57535 | 3.10619089726342 | 0 | 7 | only_H3vsUI | UpDown | notDM | Gain | 1 |
| **1_109965966_109968692_2773** | 1_109965966_109968692 | GNAI3 | 2773 | 5.08828496414573 | -0.67943072 | 1 | only_DIvsD7 | DownUp | notDM | Gain | 2 |
| **1_109969365_109970761_2773** | 1_109969365_109970761 | GNAI3 | 2773 | 5.08088972521044 | -0.67943072 | 1 | only_DIvsD7 | DownUp | notDM | Gain | 2 |
| **1_110075161_110075627_2773** | 1_110075161_110075627 | GNAI3 | 2773 | 4.19331943037448 | 0.31856486 | 4 | only_DIvsD7 | DownUp | notDM | Loss | 7 |
| **1_110089164_110090986_2773** | 1_110089164_110090986 | GNAI3 | 2773 | 2.39445168082622 | 0.27355356 | 2 | only_DIvsD7 | DownUp | notDM | Loss | 8 |
| **1_110091672_110093017_2773** | 1_110091672_110093017 | GNAI3 | 2773 | 2.64443858946784 | 0.27355356 | 2 | only_DIvsD7 | DownUp | notDM | Loss | 7 |
| **1_110105091_110106869_2773** | 1_110105091_110106869 | GNAI3 | 2773 | 4.14176323027579 | 0.31377998 | 8 | only_DIvsD7 | DownUp | notDM | Gain | 5 |
| **1_110414761_110416611_1435** | 1_110414761_110416611 | CSF1 | 1435 | 4.5668144635045 | -0.53450226 | 3 | only_H3vsUI | Down | notDM | Loss | 2 |
| **1_110419208_110420954_1435** | 1_110419208_110420954 | CSF1 | 1435 | 4.51240420191506 | -0.51047513 | 7 | only_H3vsUI | Down | notDM | Gain | 2 |
| **1_110452319_110453167_1435** | 1_110452319_110453167 | CSF1 | 1435 | 2.51321760006794 | 0.77710839 | 3 | only_H3vsUI | Down | notDM | Gain | 1 |
| **1_110453864_110457726_1435** | 1_110453864_110457726 | CSF1 | 1435 | 2.57287160220048 | 0.79335769 | 1 | only_H3vsUI | Down | notDM | Loss | 8 |
| **1_110576229_110577054_1435** | 1_110576229_110577054 | CSF1 | 1435 | 5.08898258060466 | -0.65229381 | 2 | only_H3vsUI | Down | notDM | Gain | 2 |
| **1_110878499_110880999_440600** | 1_110878499_110880999 | RBM15-AS1 | 440600 | 2.90036712865647 | 0.5646474 | 1 | only_H3vsUI | UpDown | notDM | Gain | 4 |
| **1_110882982_110884025_440600** | 1_110882982_110884025 | RBM15-AS1 | 440600 | 3.07591176148278 | 0.5646474 | 2 | only_H3vsUI | UpDown | notDM | Loss | 3 |
| **1_110884247_110887571_440600** | 1_110884247_110887571 | RBM15-AS1 | 440600 | 3.39022836246913 | 0.79085319 | 8 | only_H3vsUI | UpDown | notDM | Loss | 6 |
| **1_110930916_110938027_440600** | 1_110930916_110938027 | RBM15-AS1 | 440600 | 4.69130256338348 | -0.57338331 | 1 | only_H3vsUI | UpDown | notDM | Loss | 6 |
| **1_111989545_111991429_55924** | 1_111989545_111991429 | INKA2 | 55924 | 5.46323134277515 | -0.77252303 | 1 | only_H3vsUI | DownUp | notDM | Loss | 7 |
| **1_112163153_112164607_55924** | 1_112163153_112164607 | INKA2 | 55924 | 5.069816042572 | -0.78336948 | 2 | only_H3vsUI | DownUp | notDM | Gain | 2 |

| | | | | | | | | | | | |
|---|---|---|---|---|---|---|---|---|---|---|---|
| **1_112297186_112298043_11218** | 1_112297186_112298043 | DDX20 | 11218 | 2.68033551341456 | -0.31336933 | 1 | only_DIvsD7 | UpDown | notDM | Gain | 4 |
| **1_112297186_112298043_55924** | 1_112297186_112298043 | INKA2 | 55924 | 1.94939000664491 | -0.01009482 | 1 | only_H3vsUI | DownUp | notDM | Gain | 4 |
| **1_112299014_112300353_11218** | 1_112299014_112300353 | DDX20 | 11218 | 2.69460519893357 | -0.31336933 | 1 | only_DIvsD7 | UpDown | notDM | Loss | 3 |
| **1_112299014_112300353_55924** | 1_112299014_112300353 | INKA2 | 55924 | 4.22968184231768 | -0.68368212 | 1 | only_H3vsUI | DownUp | notDM | Loss | 7 |
| **1_11321466_11322301_93611** | 1_11321466_11322301 | FBXO44 | 93611 | 5.59396578180077 | -0.86988731 | 2 | only_DIvsD7 | DownUp | notDM | Loss | 7 |
| **1_11332426_11332864_93611** | 1_11332426_11332864 | FBXO44 | 93611 | 5.58217135465617 | 0.97507113 | 4 | only_DIvsD7 | DownUp | notDM | Gain | 2 |
| **1_113614688_113615365_10099625** | 1_113614688_113615365 | LRIG2-DT | 100996251 | 2.55990662503611 | 0.24876458 | 4 | only_H3vsUI | DownUp | notDM | Gain | 1 |
| **1_113616315_113617284_10099625** | 1_113616315_113617284 | LRIG2-DT | 100996251 | 2.77085201164214 | 0.25209049 | 1 | only_H3vsUI | DownUp | notDM | Gain | 4 |
| **1_1165566_1167165_57801** | 1_1165566_1167165 | HES4 | 57801 | 5.36175804675684 | 0.73939337 | 2 | only_H3vsUI | UpDown | notDM | Loss | 6 |
| **1_11740236_11740923_93611** | 1_11740236_11740923 | FBXO44 | 93611 | 4.40353230484943 | -0.76423793 | 2 | only_DIvsD7 | DownUp | notDM | Loss | 7 |
| **1_1207115_1208691_57801** | 1_1207115_1208691 | HES4 | 57801 | 5.43397133820086 | 0.84615119 | 2 | only_H3vsUI | UpDown | notDM | Loss | 6 |
| **1_12079797_12080834_1185** | 1_12079797_12080834 | CLCN6 | 1185 | 5.32945686617456 | 0.73602466 | 2 | notSpecific | Up | DM | Loss | 10 |
| **1_12079797_12080834_60672** | 1_12079797_12080834 | MIIP | 60672 | 2.32014628611105 | -0.13822925 | 2 | notSpecific | UpDown | DM | Loss | 10 |
| **1_12079797_12080834_7133** | 1_12079797_12080834 | TNFRSF1B | 7133 | 5.16502757007997 | 0.76067348 | 2 | notSpecific | Down | DM | Loss | 10 |
| **1_1308860_1310204_57801** | 1_1308860_1310204 | HES4 | 57801 | 5.57206962369713 | 0.86942802 | 1 | only_H3vsUI | UpDown | notDM | Loss | 6 |
| **1_1311071_1311975_57801** | 1_1311071_1311975 | HES4 | 57801 | 5.57463422876789 | 0.86942802 | 2 | only_H3vsUI | UpDown | notDM | Gain | 1 |
| **1_145655642_145657684_5174** | 1_145655642_145657684 | PDZK1 | 5174 | 4.84535853856165 | 0.76698057 | 6 | only_H3vsUI | UpDown | notDM | Loss | 5 |
| **1_145713247_145714831_5174** | 1_145713247_145714831 | PDZK1 | 5174 | 4.11038766587509 | 0.38324939 | 1 | only_H3vsUI | UpDown | notDM | Gain | 1 |
| **1_145725611_145729923_5174** | 1_145725611_145729923 | PDZK1 | 5174 | 0 | 0.79788453 | 7 | only_H3vsUI | UpDown | notDM | Loss | 3 |
| **1_145757955_145759558_5174** | 1_145757955_145759558 | PDZK1 | 5174 | 4.48048124406175 | 0.69563586 | 6 | only_H3vsUI | UpDown | notDM | Loss | 3 |
| **1_145824554_145826723_653519** | 1_145824554_145826723 | GPR89A | 653519 | 2.58092497567562 | -0.12804517 | 2 | only_H3vsUI | DownUp | notDM | Loss | 8 |
| **1_147050942_147052420_51205** | 1_147050942_147052420 | ACP6 | 51205 | 4.95498981771614 | 0.5928842 | 6 | only_H3vsUI | Down | notDM | Gain | 1 |
| **1_147072313_147073500_51205** | 1_147072313_147073500 | ACP6 | 51205 | 4.83959744622976 | 0.70996314 | 2 | only_H3vsUI | Down | notDM | Gain | 1 |
| **1_147082634_147085454_51205** | 1_147082634_147085454 | ACP6 | 51205 | 4.75713020782785 | 0.53334139 | 6 | only_H3vsUI | Down | notDM | Gain | 1 |
| **1_147140409_147142621_51205** | 1_147140409_147142621 | ACP6 | 51205 | 0 | 0.89796892 | 2 | only_H3vsUI | Down | notDM | Loss | 8 |
| **1_147142679_147143352_51205** | 1_147142679_147143352 | ACP6 | 51205 | 2.50105926221775 | 0.77748077 | 2 | only_H3vsUI | Down | notDM | Gain | 1 |
| **1_149781254_149783882_653604** | 1_149781254_149783882 | H3C13 | 653604 | 3.13193929521042 | 0.00257128 | 2 | only_DIvsD7 | Up | notDM | Gain | 4 |
| **1_150121171_150122016_54507** | 1_150121171_150122016 | ADAMTSL4 | 54507 | 5.60191773658965 | -0.91058366 | 4 | only_DIvsD7 | DownUp | notDM | Loss | 7 |
| **1_150122665_150123693_54507** | 1_150122665_150123693 | ADAMTSL4 | 54507 | 5.60009253045092 | -0.90841206 | 1 | only_DIvsD7 | DownUp | notDM | Loss | 7 |
| **1_150459962_150461357_54507** | 1_150459962_150461357 | ADAMTSL4 | 54507 | 4.78205676374009 | 0.74099448 | 2 | only_DIvsD7 | DownUp | notDM | Gain | 1 |
| **1_150480240_150480961_54507** | 1_150480240_150480961 | ADAMTSL4 | 54507 | 4.61211601294338 | 0.42112202 | 6 | only_DIvsD7 | DownUp | notDM | Gain | 1 |
| **1_150520547_150521690_54507** | 1_150520547_150521690 | ADAMTSL4 | 54507 | 2.29003461136252 | 0.5979398 | 4 | only_DIvsD7 | DownUp | notDM | Gain | 4 |
| **1_150532716_150537979_54507** | 1_150532716_150537979 | ADAMTSL4 | 54507 | 4.03426739703803 | -0.20418712 | 5 | only_DIvsD7 | DownUp | notDM | Loss | 6 |
| **1_150539633_150542496_54507** | 1_150539633_150542496 | ADAMTSL4 | 54507 | 4.24922282399667 | -0.74401446 | 7 | only_DIvsD7 | DownUp | notDM | Gain | 1 |

| | | | | | | | | | | | |
|---|---|---|---|---|---|---|---|---|---|---|---|
| **1_150953420_150955489_54507** | 1_150953420_150955489 | ADAMTSL4 | 54507 | 5.635019044463 | 0.84322854 | 3 | only_DIvsD7 | DownUp | notDM | Gain | 1 |
| **1_151652184_151653810_51686** | 1_151652184_151653810 | OAZ3 | 51686 | 4.91188171700819 | 0.7285131 | 3 | only_H3vsUI | UpDown | notDM | Loss | 5 |
| **1_151689308_151690333_11189** | 1_151689308_151690333 | CELF3 | 11189 | 1.30102999566398 | 0.00800082 | 4 | only_DIvsD7 | Up | notDM | Loss | 3 |
| **1_151693974_151694690_11189** | 1_151693974_151694690 | CELF3 | 11189 | 3.67080228426094 | 0.03310871 | 2 | only_DIvsD7 | Up | notDM | Loss | 3 |
| **1_151734604_151735634_51686** | 1_151734604_151735634 | OAZ3 | 51686 | 0 | 0.41414152 | 1 | only_H3vsUI | UpDown | notDM | Loss | 3 |
| **1_151736072_151737733_51686** | 1_151736072_151737733 | OAZ3 | 51686 | 2.79865064544527 | 0.41414152 | 1 | only_H3vsUI | UpDown | notDM | Loss | 6 |
| **1_152473185_152476296_353133** | 1_152473185_152476296 | LCE1C | 353133 | 5.48117308162896 | 0.96336184 | 7 | only_DIvsD7 | DownUp | notDM | Loss | 5 |
| **1_152717625_152718405_353133** | 1_152717625_152718405 | LCE1C | 353133 | 4.78321015485203 | 0.96496757 | 4 | only_DIvsD7 | DownUp | notDM | Gain | 1 |
| **1_154296746_154297893_89872** | 1_154296746_154297893 | AQP10 | 89872 | 3.50270017531056 | 0.09072872 | 1 | only_H3vsUI | DownUp | notDM | Gain | 4 |
| **1_154298233_154299842_89872** | 1_154298233_154299842 | AQP10 | 89872 | 3.66642437251876 | 0.07200588 | 1 | only_H3vsUI | DownUp | notDM | Loss | 6 |
| **1_154539690_154541404_80308** | 1_154539690_154541404 | FLAD1 | 80308 | 5.6175287258864 | 0.83265859 | 4 | only_DIvsD7 | DownUp | notDM | Gain | 1 |
| **1_154907807_154911074_80308** | 1_154907807_154911074 | FLAD1 | 80308 | 4.65160748748126 | -0.85764596 | 2 | only_DIvsD7 | DownUp | notDM | Loss | 7 |
| **1_154940623_154946839_80308** | 1_154940623_154946839 | FLAD1 | 80308 | 3.95755940189748 | -0.40192407 | 5 | only_DIvsD7 | DownUp | notDM | Loss | 7 |
| **1_154947525_154950491_80308** | 1_154947525_154950491 | FLAD1 | 80308 | 3.7337588355872 | -0.40192407 | 1 | only_DIvsD7 | DownUp | notDM | Loss | 7 |
| **1_154955788_154957961_80308** | 1_154955788_154957961 | FLAD1 | 80308 | 0 | 0.23495364 | 1 | only_DIvsD7 | DownUp | notDM | Loss | 8 |
| **1_154971352_154971949_55891** | 1_154971352_154971949 | LENEP | 55891 | 3.72345567203519 | 0 | 4 | only_DIvsD7 | Up | notDM | Gain | 1 |
| **1_154971352_154971949_80308** | 1_154971352_154971949 | FLAD1 | 80308 | 4.1888441465469 | -0.3422186 | 4 | only_DIvsD7 | DownUp | notDM | Gain | 2 |
| **1_154972133_154973050_80308** | 1_154972133_154973050 | FLAD1 | 80308 | 4.21026499903229 | -0.77853146 | 5 | only_DIvsD7 | DownUp | notDM | Loss | 7 |
| **1_154973216_154974573_80308** | 1_154973216_154974573 | FLAD1 | 80308 | 4.23832215637555 | -0.77853146 | 1 | only_DIvsD7 | DownUp | notDM | Loss | 7 |
| **1_154976588_154981577_80308** | 1_154976588_154981577 | FLAD1 | 80308 | 4.31561353195342 | -0.46549477 | 5 | only_DIvsD7 | DownUp | notDM | Loss | 7 |
| **1_154988952_154989836_80308** | 1_154988952_154989836 | FLAD1 | 80308 | 4.51913204044551 | -0.55987646 | 7 | only_DIvsD7 | DownUp | notDM | Loss | 7 |
| **1_154990097_154992273_80308** | 1_154990097_154992273 | FLAD1 | 80308 | 4.53392450481999 | -0.65264829 | 3 | only_DIvsD7 | DownUp | notDM | Loss | 7 |
| **1_154993579_154996577_80308** | 1_154993579_154996577 | FLAD1 | 80308 | 4.57711248826435 | 0.48539881 | 6 | only_DIvsD7 | DownUp | notDM | Gain | 5 |
| **1_154996797_154999950_80308** | 1_154996797_154999950 | FLAD1 | 80308 | 4.61225390609644 | 0.66842745 | 6 | only_DIvsD7 | DownUp | notDM | Gain | 5 |
| **1_155023844_155025552_80308** | 1_155023844_155025552 | FLAD1 | 80308 | 4.83211915016312 | -0.68486952 | 1 | only_DIvsD7 | DownUp | notDM | Loss | 7 |
| **1_155035262_155036492_80308** | 1_155035262_155036492 | FLAD1 | 80308 | 4.89993535101983 | -0.60481173 | 2 | only_DIvsD7 | DownUp | notDM | Gain | 2 |
| **1_155056291_155059319_80308** | 1_155056291_155059319 | FLAD1 | 80308 | 5.00167314959487 | -0.75732588 | 2 | only_DIvsD7 | DownUp | notDM | Loss | 7 |
| **1_155105700_155108133_80128** | 1_155105700_155108133 | TRIM46 | 80128 | 4.5921545422762 | -0.48774113 | 1 | only_DIvsD7 | UpDown | notDM | Loss | 6 |
| **1_155105700_155108133_80308** | 1_155105700_155108133 | FLAD1 | 80308 | 5.17549731731105 | -0.84568361 | 1 | only_DIvsD7 | DownUp | notDM | Loss | 7 |
| **1_155108384_155109854_80128** | 1_155108384_155109854 | TRIM46 | 80128 | 4.5726044405492 | -0.40129653 | 1 | only_DIvsD7 | UpDown | notDM | Loss | 3 |
| **1_155108384_155109854_80308** | 1_155108384_155109854 | FLAD1 | 80308 | 5.18337520071103 | -0.86338387 | 1 | only_DIvsD7 | DownUp | notDM | Loss | 7 |
| **1_155111131_155113501_80128** | 1_155111131_155113501 | TRIM46 | 80128 | 4.5280163411892 | -0.4537276 | 1 | only_DIvsD7 | UpDown | notDM | Loss | 7 |
| **1_155111131_155113501_80308** | 1_155111131_155113501 | FLAD1 | 80308 | 5.19096446634751 | -0.67927357 | 1 | only_DIvsD7 | DownUp | notDM | Loss | 7 |
| **1_155140061_155140923_80128** | 1_155140061_155140923 | TRIM46 | 80128 | 3.78922805726733 | 0.25092389 | 4 | only_DIvsD7 | UpDown | notDM | Gain | 1 |

| | | | | | | | | | | | |
|---|---|---|---|---|---|---|---|---|---|---|---|
| **1_155144210_155145548_80128** | 1_155144210_155145548 | TRIM46 | 80128 | 3.22608411597582 | -0.56350094 | 1 | only_DIvsD7 | UpDown | notDM | Loss | 3 |
| **1_155144210_155145548_80308** | 1_155144210_155145548 | FLAD1 | 80308 | 5.27486185184662 | -0.76590789 | 1 | only_DIvsD7 | DownUp | notDM | Loss | 7 |
| **1_155145906_155147175_80128** | 1_155145906_155147175 | TRIM46 | 80128 | 1.7481880270062 | -0.56350094 | 1 | only_DIvsD7 | UpDown | notDM | Loss | 3 |
| **1_155145906_155147175_80308** | 1_155145906_155147175 | FLAD1 | 80308 | 5.27875588670724 | -0.76590789 | 1 | only_DIvsD7 | DownUp | notDM | Loss | 7 |
| **1_155148586_155149167_80128** | 1_155148586_155149167 | TRIM46 | 80128 | 3.13289976994448 | -0.10311545 | 6 | only_DIvsD7 | UpDown | notDM | Gain | 1 |
| **1_155162831_155164891_80308** | 1_155162831_155164891 | FLAD1 | 80308 | 5.31601020640611 | 0.85667388 | 1 | only_DIvsD7 | DownUp | notDM | Loss | 6 |
| **1_155165104_155166526_80308** | 1_155165104_155166526 | FLAD1 | 80308 | 5.32075262813679 | 0.74367746 | 1 | only_DIvsD7 | DownUp | notDM | Gain | 5 |
| **1_155209919_155212172_80308** | 1_155209919_155212172 | FLAD1 | 80308 | 5.4048576534514 | -0.87209882 | 2 | only_DIvsD7 | DownUp | notDM | Gain | 2 |
| **1_155212322_155214441_80308** | 1_155212322_155214441 | FLAD1 | 80308 | 5.40910430067654 | 0.83997708 | 1 | only_DIvsD7 | DownUp | notDM | Loss | 6 |
| **1_155230495_155232023_80308** | 1_155230495_155232023 | FLAD1 | 80308 | 5.43878592837204 | -0.76240381 | 2 | only_DIvsD7 | DownUp | notDM | Loss | 7 |
| **1_155232250_155232864_80308** | 1_155232250_155232864 | FLAD1 | 80308 | 5.44155217975391 | -0.76240381 | 3 | only_DIvsD7 | DownUp | notDM | Gain | 2 |
| **1_156162691_156163595_11243** | 1_156162691_156163595 | PMF1 | 11243 | 4.28282595915353 | 0.41106796 | 1 | only_H3vsUI | UpDown | notDM | Gain | 4 |
| **1_156164027_156165580_11243** | 1_156164027_156165580 | PMF1 | 11243 | 4.23537692240224 | 0.36946483 | 2 | only_H3vsUI | UpDown | notDM | Loss | 6 |
| **1_156181888_156182656_11243** | 1_156181888_156182656 | PMF1 | 11243 | 2.11058971029925 | -0.27534236 | 4 | only_H3vsUI | UpDown | notDM | Gain | 4 |
| **1_156183117_156184523_11243** | 1_156183117_156184523 | PMF1 | 11243 | 2.48000694295715 | -0.27986982 | 1 | only_H3vsUI | UpDown | notDM | Gain | 4 |
| **1_156194474_156195321_11243** | 1_156194474_156195321 | PMF1 | 11243 | 4.06829720503145 | -0.10435558 | 6 | only_H3vsUI | UpDown | notDM | Gain | 1 |
| **1_156679382_156680737_11243** | 1_156679382_156680737 | PMF1 | 11243 | 5.69597785457797 | 0.86788264 | 6 | only_H3vsUI | UpDown | notDM | Gain | 1 |
| **1_157012227_157015324_9826** | 1_157012227_157015324 | ARHGEF11 | 9826 | 0 | -0.04291899 | 2 | only_H3vsUI | DownUp | notDM | Gain | 8 |
| **1_157016124_157016704_9826** | 1_157016124_157016704 | ARHGEF11 | 9826 | 2.98407703390283 | -0.01046307 | 1 | only_H3vsUI | DownUp | notDM | Gain | 2 |
| **1_160175368_160177396_729867** | 1_160175368_160177396 | LOC729867 | 729867 | 3.10174707394637 | 0 | 1 | only_DIvsD7 | Up | notDM | Loss | 6 |
| **1_161013855_161015624_9507** | 1_161013855_161015624 | ADAMTS4 | 9507 | 5.18532396127145 | -0.86013956 | 1 | only_H3vsUI | UpDown | notDM | Loss | 3 |
| **1_161015974_161016815_9507** | 1_161015974_161016815 | ADAMTS4 | 9507 | 5.18193500873911 | -0.86013956 | 1 | only_H3vsUI | UpDown | notDM | Loss | 3 |
| **1_161084044_161087703_27005** | 1_161084044_161087703 | USP21 | 27005 | 4.61918727840529 | 0.58318151 | 1 | only_DIvsD7 | UpDown | notDM | Gain | 5 |
| **1_161100725_161102177_9507** | 1_161100725_161102177 | ADAMTS4 | 9507 | 4.82393045512556 | -0.59382049 | 1 | only_H3vsUI | UpDown | notDM | Loss | 3 |
| **1_161102804_161104908_9507** | 1_161102804_161104908 | ADAMTS4 | 9507 | 4.80576583966357 | -0.59382049 | 1 | only_H3vsUI | UpDown | notDM | Gain | 4 |
| **1_161122773_161123486_27005** | 1_161122773_161123486 | USP21 | 27005 | 3.76004532796581 | 0.39916554 | 2 | only_DIvsD7 | UpDown | notDM | Gain | 4 |
| **1_161122773_161123486_9507** | 1_161122773_161123486 | ADAMTS4 | 9507 | 4.65668262017265 | -0.80328663 | 2 | only_H3vsUI | UpDown | notDM | Gain | 2 |
| **1_161123979_161124974_27005** | 1_161123979_161124974 | USP21 | 27005 | 3.63012264285931 | 0.39916554 | 1 | only_DIvsD7 | UpDown | notDM | Gain | 4 |
| **1_161123979_161124974_9507** | 1_161123979_161124974 | ADAMTS4 | 9507 | 4.6421973321232 | -0.80328663 | 1 | only_H3vsUI | UpDown | notDM | Gain | 4 |
| **1_161128391_161129194_27005** | 1_161128391_161129194 | USP21 | 27005 | 1.67209785793572 | 0.06917781 | 1 | only_DIvsD7 | UpDown | notDM | Loss | 3 |
| **1_161128391_161129194_9507** | 1_161128391_161129194 | ADAMTS4 | 9507 | 4.59824319165362 | -0.8678183 | 1 | only_H3vsUI | UpDown | notDM | Loss | 3 |
| **1_161129484_161130710_27005** | 1_161129484_161130710 | USP21 | 27005 | 2.33645973384853 | 0.06983586 | 1 | only_DIvsD7 | UpDown | notDM | Loss | 8 |
| **1_161129484_161130710_9507** | 1_161129484_161130710 | ADAMTS4 | 9507 | 4.58131236223185 | -0.8678183 | 1 | only_H3vsUI | UpDown | notDM | Loss | 3 |
| **1_161134830_161136039_27005** | 1_161134830_161136039 | USP21 | 27005 | 3.06557971472845 | 0.00134534 | 1 | only_DIvsD7 | UpDown | notDM | Loss | 7 |

| | | | | | | | | | | | |
|---|---|---|---|---|---|---|---|---|---|---|---|
| **1_161134830_161136039_9507** | 1_161134830_161136039 | ADAMTS4 | 9507 | 4.51597975627943 | -0.58183012 | 1 | only_H3vsUI | UpDown | notDM | Loss | 3 |
| **1_161136240_161137824_27005** | 1_161136240_161137824 | USP21 | 27005 | 3.84341966520492 | 0.21434787 | 2 | only_DIvsD7 | UpDown | notDM | Gain | 4 |
| **1_161136240_161137824_9507** | 1_161136240_161137824 | ADAMTS4 | 9507 | 4.49168379284727 | -0.58183012 | 2 | only_H3vsUI | UpDown | notDM | Gain | 4 |
| **1_161145563_161147103_27005** | 1_161145563_161147103 | USP21 | 27005 | 4.21285318994711 | 0.42730963 | 1 | only_DIvsD7 | UpDown | notDM | Loss | 6 |
| **1_161145563_161147103_9507** | 1_161145563_161147103 | ADAMTS4 | 9507 | 4.33727951603794 | -0.54896212 | 1 | only_H3vsUI | UpDown | notDM | Loss | 3 |
| **1_161147350_161148146_27005** | 1_161147350_161148146 | USP21 | 27005 | 4.25796640950818 | 0.42730963 | 5 | only_DIvsD7 | UpDown | notDM | Gain | 1 |
| **1_161147350_161148146_9507** | 1_161147350_161148146 | ADAMTS4 | 9507 | 4.31592838261019 | -0.54896212 | 5 | only_H3vsUI | UpDown | notDM | Gain | 1 |
| **1_161170660_161172013_9507** | 1_161170660_161172013 | ADAMTS4 | 9507 | 3.25911584418507 | -0.1344029 | 2 | only_H3vsUI | UpDown | notDM | Gain | 4 |
| **1_161172273_161173583_9507** | 1_161172273_161173583 | ADAMTS4 | 9507 | 3.53516748511494 | -0.1344029 | 1 | only_H3vsUI | UpDown | notDM | Gain | 4 |
| **1_161195840_161197748_27005** | 1_161195840_161197748 | USP21 | 27005 | 4.82348727085081 | 0.54922058 | 1 | only_DIvsD7 | UpDown | notDM | Loss | 6 |
| **1_161282922_161284140_27005** | 1_161282922_161284140 | USP21 | 27005 | 5.18654669695217 | 0.76087202 | 2 | only_DIvsD7 | UpDown | notDM | Gain | 1 |
| **1_161282922_161284140_9507** | 1_161282922_161284140 | ADAMTS4 | 9507 | 5.05720189858267 | -0.72055858 | 2 | only_H3vsUI | UpDown | notDM | Gain | 4 |
| **1_161284243_161285706_27005** | 1_161284243_161285706 | USP21 | 27005 | 5.19026444736645 | 0.76087202 | 1 | only_DIvsD7 | UpDown | notDM | Loss | 6 |
| **1_161284243_161285706_9507** | 1_161284243_161285706 | ADAMTS4 | 9507 | 5.06220204541968 | -0.72055858 | 1 | only_H3vsUI | UpDown | notDM | Loss | 3 |
| **1_161366741_161369514_9507** | 1_161366741_161369514 | ADAMTS4 | 9507 | 5.29643921061205 | -0.77254393 | 5 | only_H3vsUI | UpDown | notDM | Gain | 4 |
| **1_161369758_161371091_9507** | 1_161369758_161371091 | ADAMTS4 | 9507 | 5.30301020011761 | -0.75789116 | 1 | only_H3vsUI | UpDown | notDM | Gain | 4 |
| **1_161508899_161509912_27005** | 1_161508899_161509912 | USP21 | 27005 | 5.57929645653911 | -0.90902851 | 1 | only_DIvsD7 | UpDown | notDM | Loss | 7 |
| **1_161580256_161582112_27005** | 1_161580256_161582112 | USP21 | 27005 | 5.65411009262783 | -0.83342821 | 1 | only_DIvsD7 | UpDown | notDM | Loss | 7 |
| **1_161582538_161583587_27005** | 1_161582538_161583587 | USP21 | 27005 | 5.65630235870965 | -0.87379931 | 2 | only_DIvsD7 | UpDown | notDM | Gain | 2 |
| **1_16301305_16302078_149563** | 1_16301305_16302078 | SRARP | 149563 | 4.45718525655367 | 0.62484104 | 2 | only_H3vsUI | UpDown | notDM | Loss | 3 |
| **1_16302672_16303573_149563** | 1_16302672_16303573 | SRARP | 149563 | 4.43391377509016 | 0.49081316 | 4 | only_H3vsUI | UpDown | notDM | Loss | 3 |
| **1_16562342_16563622_149563** | 1_16562342_16563622 | SRARP | 149563 | 5.36476293186808 | -0.76852619 | 2 | only_H3vsUI | UpDown | notDM | Gain | 4 |
| **1_16562342_16563622_26099** | 1_16562342_16563622 | SZRD1 | 26099 | 5.11434739224287 | 0.82547512 | 2 | only_DIvsD7 | DownUp | notDM | Gain | 1 |
| **1_16660929_16662405_149563** | 1_16660929_16662405 | SRARP | 149563 | 5.51877706892677 | 0.99845769 | 6 | only_H3vsUI | UpDown | notDM | Loss | 3 |
| **1_16660929_16662405_26099** | 1_16660929_16662405 | SZRD1 | 26099 | 4.49607127622064 | -0.85799688 | 6 | only_DIvsD7 | DownUp | notDM | Loss | 7 |
| **1_16675178_16678625_26099** | 1_16675178_16678625 | SZRD1 | 26099 | 2.64933485871214 | 0.00577768 | 1 | only_DIvsD7 | DownUp | notDM | Loss | 6 |
| **1_16679148_16679686_26099** | 1_16679148_16679686 | SZRD1 | 26099 | 1.90308998699194 | 0.02178446 | 5 | only_DIvsD7 | DownUp | notDM | Gain | 4 |
| **1_16694436_16696932_26099** | 1_16694436_16696932 | SZRD1 | 26099 | 2.83122969386706 | 0.60283322 | 1 | only_DIvsD7 | DownUp | notDM | Gain | 4 |
| **1_16698112_16703266_26099** | 1_16698112_16703266 | SZRD1 | 26099 | 3.63888842470508 | 0.37525905 | 7 | only_DIvsD7 | DownUp | notDM | Loss | 6 |
| **1_1675718_1677172_339456** | 1_1675718_1677172 | TMEM52 | 339456 | 5.23936455584478 | 0.80663997 | 2 | only_DIvsD7 | DownUp | notDM | Gain | 1 |
| **1_178060839_178062656_10030240** | 1_178060839_178062656 | RASAL2-AS1 | 100302401 | 1.90308998699194 | -0.03475708 | 2 | only_H3vsUI | DownUp | notDM | Loss | 3 |
| **1_178063693_178066943_10030240** | 1_178063693_178066943 | RASAL2-AS1 | 100302401 | 2.76042248342321 | 0.11859647 | 2 | only_H3vsUI | DownUp | notDM | Gain | 4 |
| **1_1789363_1792353_339456** | 1_1789363_1792353 | TMEM52 | 339456 | 4.76611528322141 | -0.60362135 | 8 | only_DIvsD7 | DownUp | notDM | Loss | 7 |
| **1_179711613_179713333_148753** | 1_179711613_179713333 | FAM163A | 148753 | 0 | 0 | 4 | notSpecific | UpDown | DM | Loss | 10 |

| | | | | | | | | | | | |
|---|---|---|---|---|---|---|---|---|---|---|---|
| **1_182572645_182574262_6004** | 1_182572645_182574262 | RGS16 | 6004 | 0 | 0.97159727 | 4 | only_DIvsD7 | Up | notDM | Loss | 3 |
| **1_182583276_182584709_6004** | 1_182583276_182584709 | RGS16 | 6004 | 3.98833595585605 | 0.98507974 | 4 | only_DIvsD7 | Up | notDM | Loss | 3 |
| **1_1839404_1840304_339456** | 1_1839404_1840304 | TMEM52 | 339456 | 4.01678271248684 | 0.16203582 | 2 | only_DIvsD7 | DownUp | notDM | Loss | 3 |
| **1_184005058_184006153_23127** | 1_184005058_184006153 | COLGALT2 | 23127 | 2.85186960072977 | 0.72347078 | 4 | only_DIvsD7 | UpDown | notDM | Loss | 3 |
| **1_184019847_184020745_23127** | 1_184019847_184020745 | COLGALT2 | 23127 | 4.11347539853674 | 0.78504535 | 2 | only_DIvsD7 | UpDown | notDM | Loss | 3 |
| **1_184021237_184022767_23127** | 1_184021237_184022767 | COLGALT2 | 23127 | 4.15763806410092 | 0.78504535 | 2 | only_DIvsD7 | UpDown | notDM | Gain | 1 |
| **1_184985603_184987537_6045** | 1_184985603_184987537 | RNF2 | 6045 | 4.43332171163078 | -0.77648317 | 6 | only_DIvsD7 | UpDown | notDM | Gain | 2 |
| **1_185013638_185014422_6045** | 1_185013638_185014422 | RNF2 | 6045 | 2.3747483460101 | -0.27061966 | 2 | only_DIvsD7 | UpDown | notDM | Loss | 3 |
| **1_185015008_185017426_6045** | 1_185015008_185017426 | RNF2 | 6045 | 2.54654266347813 | 0.50521603 | 2 | only_DIvsD7 | UpDown | notDM | Loss | 6 |
| **1_19274220_19282231_126917** | 1_19274220_19282231 | IFFO2 | 126917 | 2.97772360528885 | 0.74817683 | 1 | notSpecific | Down | DM | Loss | 10 |
| **1_19274220_19282231_23352** | 1_19274220_19282231 | UBR4 | 23352 | 5.40575603977396 | -0.82102953 | 1 | notSpecific | Down | DM | Loss | 10 |
| **1_19274220_19282231_51154** | 1_19274220_19282231 | MRTO4 | 51154 | 5.47100257424789 | -0.96514016 | 1 | notSpecific | Up | DM | Loss | 10 |
| **1_19274220_19282231_8659** | 1_19274220_19282231 | ALDH4A1 | 8659 | 4.65325111601366 | 0.61656 | 1 | notSpecific | Up | DM | Loss | 10 |
| **1_192775926_192777952_5997** | 1_192775926_192777952 | RGS2 | 5997 | 2.3384564936046 | 0.58764883 | 5 | only_DIvsD7 | Up | notDM | Gain | 4 |
| **1_192778211_192780036_5997** | 1_192778211_192780036 | RGS2 | 5997 | 1.61278385671974 | 0.63101094 | 2 | only_DIvsD7 | Up | notDM | Loss | 6 |
| **1_192873169_192874901_5997** | 1_192873169_192874901 | RGS2 | 5997 | 4.97773274823433 | 0.72661486 | 4 | only_DIvsD7 | Up | notDM | Loss | 5 |
| **1_19335793_19337645_23065** | 1_19335793_19337645 | EMC1 | 23065 | 5.38093807641452 | 0.93489293 | 7 | only_H3vsUI | DownUp | notDM | Gain | 5 |
| **1_19335793_19337645_54896** | 1_19335793_19337645 | SLC66A1 | 54896 | 5.47882324499362 | 0.94363693 | 7 | only_DIvsD7 | DownUp | notDM | Gain | 5 |
| **1_19337809_19340741_23065** | 1_19337809_19340741 | EMC1 | 23065 | 5.37530871897566 | 0.93559838 | 7 | only_H3vsUI | DownUp | notDM | Gain | 5 |
| **1_19337809_19340741_54896** | 1_19337809_19340741 | SLC66A1 | 54896 | 5.47433283755217 | 0.94455688 | 7 | only_DIvsD7 | DownUp | notDM | Gain | 5 |
| **1_19534475_19536441_23065** | 1_19534475_19536441 | EMC1 | 23065 | 4.61870688721103 | 0.77431448 | 1 | only_H3vsUI | DownUp | notDM | Loss | 6 |
| **1_19537029_19539562_23065** | 1_19537029_19539562 | EMC1 | 23065 | 4.5848059743755 | 0.77431448 | 1 | only_H3vsUI | DownUp | notDM | Loss | 6 |
| **1_19576683_19577869_23065** | 1_19576683_19577869 | EMC1 | 23065 | 2.13033376849501 | 0.11433042 | 1 | only_H3vsUI | DownUp | notDM | Loss | 8 |
| **1_19578530_19580569_23065** | 1_19578530_19580569 | EMC1 | 23065 | 2.71264970162721 | -0.0187384 | 1 | only_H3vsUI | DownUp | notDM | Loss | 8 |
| **1_19636768_19638429_54896** | 1_19636768_19638429 | SLC66A1 | 54896 | 2.60959440922522 | 0.06586046 | 2 | only_DIvsD7 | DownUp | notDM | Loss | 7 |
| **1_19638925_19640798_54896** | 1_19638925_19640798 | SLC66A1 | 54896 | 1.96378782734556 | 0.04906644 | 1 | only_DIvsD7 | DownUp | notDM | Loss | 8 |
| **1_19766457_19768352_54896** | 1_19766457_19768352 | SLC66A1 | 54896 | 5.10593235218658 | 0.69319184 | 8 | only_DIvsD7 | DownUp | notDM | Loss | 5 |
| **1_202162193_202163038_148713** | 1_202162193_202163038 | PTPRVP | 148713 | 3.79323144705652 | -0.00365573 | 6 | only_H3vsUI | UpDown | notDM | Gain | 1 |
| **1_202427516_202429614_51094** | 1_202427516_202429614 | ADIPOR1 | 51094 | 5.69705838196914 | 0.9152875 | 6 | only_H3vsUI | DownUp | notDM | Gain | 5 |
| **1_202771448_202773565_51094** | 1_202771448_202773565 | ADIPOR1 | 51094 | 5.1871059690316 | 0.81935532 | 3 | only_H3vsUI | DownUp | notDM | Gain | 5 |
| **1_202925552_202927206_51094** | 1_202925552_202927206 | ADIPOR1 | 51094 | 2.31806333496276 | 0.16855801 | 1 | only_H3vsUI | DownUp | notDM | Loss | 8 |
| **1_202927584_202929430_51094** | 1_202927584_202929430 | ADIPOR1 | 51094 | 1.83884909073726 | 0.13014303 | 1 | only_H3vsUI | DownUp | notDM | Gain | 4 |
| **1_202934942_202936297_51094** | 1_202934942_202936297 | ADIPOR1 | 51094 | 3.87662191603427 | 0.20129034 | 1 | only_H3vsUI | DownUp | notDM | Loss | 7 |
| **1_202936685_202937585_51094** | 1_202936685_202937585 | ADIPOR1 | 51094 | 3.9670797341445 | 0.14697426 | 2 | only_H3vsUI | DownUp | notDM | Loss | 7 |

| | | | | | | | | | | | |
|---|---|---|---|---|---|---|---|---|---|---|---|
| **1_204181902_204184057_127845** | 1_204181902_204184057 | GOLT1A | 127845 | 0 | 0.70328267 | 1 | only_DIvsD7 | DownUp | notDM | Loss | 6 |
| **1_204247333_204249061_127845** | 1_204247333_204249061 | GOLT1A | 127845 | 4.80695964660353 | 0.99020665 | 8 | only_DIvsD7 | DownUp | notDM | Loss | 5 |
| **1_204251365_204253800_127845** | 1_204251365_204253800 | GOLT1A | 127845 | 4.83344674187101 | 0.99480837 | 6 | only_DIvsD7 | DownUp | DM | Loss | 10 |
| **1_204299425_204302477_127845** | 1_204299425_204302477 | GOLT1A | 127845 | 5.06523228958451 | 0.99077831 | 7 | only_DIvsD7 | DownUp | notDM | Loss | 5 |
| **1_204474620_204475478_127845** | 1_204474620_204475478 | GOLT1A | 127845 | 5.4644925281683 | -0.86831815 | 4 | only_DIvsD7 | DownUp | notDM | Gain | 2 |
| **1_205179112_205180388_25778** | 1_205179112_205180388 | DSTYK | 25778 | 2.48713837547719 | -0.47576388 | 1 | only_DIvsD7 | Down | notDM | Loss | 8 |
| **1_205180924_205181754_25778** | 1_205180924_205181754 | DSTYK | 25778 | 2.3654879848909 | -0.53705895 | 2 | only_DIvsD7 | Down | notDM | Gain | 4 |
| **1_205197318_205198621_25778** | 1_205197318_205198621 | DSTYK | 25778 | 4.22102280520484 | 0.28530083 | 2 | only_DIvsD7 | Down | notDM | Loss | 7 |
| **1_205780408_205782213_254428** | 1_205780408_205782213 | SLC41A1 | 254428 | 1.96378782734556 | 0.59547814 | 2 | notSpecific | Down | DM | Loss | 10 |
| **1_205780408_205782213_64710** | 1_205780408_205782213 | NUCKS1 | 64710 | 4.78537255040007 | 0.61279334 | 2 | notSpecific | Down | DM | Loss | 10 |
| **1_207063687_207065714_11009** | 1_207063687_207065714 | IL24 | 11009 | 3.70543604658525 | 0.28321991 | 4 | only_H3vsUI | UpDown | notDM | Gain | 1 |
| **1_207068199_207072853_11009** | 1_207068199_207072853 | IL24 | 11009 | 0 | 0.28340261 | 6 | only_H3vsUI | UpDown | notDM | Loss | 3 |
| **1_207082102_207083863_11009** | 1_207082102_207083863 | IL24 | 11009 | 4.05284778340086 | 0.32073953 | 2 | only_H3vsUI | UpDown | notDM | Loss | 3 |
| **1_209584302_209585498_3664** | 1_209584302_209585498 | IRF6 | 3664 | 5.59537716031882 | 0.9853252 | 6 | only_DIvsD7 | UpDown | notDM | Gain | 1 |
| **1_209861645_209863605_3664** | 1_209861645_209863605 | IRF6 | 3664 | 5.06365229998569 | 0.73938861 | 7 | only_DIvsD7 | UpDown | notDM | Loss | 5 |
| **1_209920337_209921112_3664** | 1_209920337_209921112 | IRF6 | 3664 | 4.76550463911944 | 0.98513912 | 4 | only_DIvsD7 | UpDown | notDM | Gain | 1 |
| **1_209956550_209957804_3664** | 1_209956550_209957804 | IRF6 | 3664 | 4.33417217272461 | -0.9912954 | 1 | only_DIvsD7 | UpDown | notDM | Loss | 3 |
| **1_209998824_210001214_3664** | 1_209998824_210001214 | IRF6 | 3664 | 4.28862923466499 | 0.36205823 | 1 | only_DIvsD7 | UpDown | notDM | Gain | 1 |
| **1_210001991_210003431_3664** | 1_210001991_210003431 | IRF6 | 3664 | 4.35272263746202 | 0.48744977 | 2 | only_DIvsD7 | UpDown | notDM | Loss | 3 |
| **1_210052833_210053746_3664** | 1_210052833_210053746 | IRF6 | 3664 | 4.86596814831917 | 0.98641293 | 4 | only_DIvsD7 | UpDown | notDM | Gain | 1 |
| **1_210406427_210408391_3664** | 1_210406427_210408391 | IRF6 | 3664 | 5.63439463550726 | 0.79934401 | 2 | only_DIvsD7 | UpDown | notDM | Loss | 3 |
| **1_212094074_212095365_55248** | 1_212094074_212095365 | PACC1 | 55248 | 5.69273502778714 | -0.83708662 | 7 | only_DIvsD7 | UpDown | notDM | Gain | 2 |
| **1_212117620_212119865_55248** | 1_212117620_212119865 | PACC1 | 55248 | 5.67059741503275 | -0.90480935 | 3 | only_DIvsD7 | UpDown | notDM | Gain | 5 |
| **1_21220816_21223435_1889** | 1_21220816_21223435 | ECE1 | 1889 | 5.59474161376386 | 0.84380398 | 7 | notSpecific | Down | DM | Gain | 9 |
| **1_21220816_21223435_50809** | 1_21220816_21223435 | HP1BP3 | 50809 | 5.02939189526321 | 0.7024543 | 7 | notSpecific | Down | DM | Gain | 9 |
| **1_21220816_21223435_8672** | 1_21220816_21223435 | EIF4G3 | 8672 | 3.47304880508854 | -0.02168277 | 7 | notSpecific | Down | DM | Gain | 9 |
| **1_212586381_212588124_55248** | 1_212586381_212588124 | PACC1 | 55248 | 2.07918124604762 | 0.91312975 | 1 | only_DIvsD7 | UpDown | notDM | Loss | 6 |
| **1_212605236_212606139_55248** | 1_212605236_212606139 | PACC1 | 55248 | 4.23032116891908 | 0.56728137 | 4 | only_DIvsD7 | UpDown | notDM | Gain | 1 |
| **1_212606543_212608026_55248** | 1_212606543_212608026 | PACC1 | 55248 | 4.26249855101477 | 0.55882309 | 1 | only_DIvsD7 | UpDown | notDM | Loss | 6 |
| **1_212782190_212784373_55248** | 1_212782190_212784373 | PACC1 | 55248 | 5.28768754472184 | -0.94985054 | 2 | only_DIvsD7 | UpDown | notDM | Loss | 7 |
| **1_2143147_2143914_11079** | 1_2143147_2143914 | RER1 | 11079 | 5.25386117004511 | 0.71856384 | 7 | notSpecific | DownUp | DM | Gain | 9 |
| **1_2143147_2143914_199990** | 1_2143147_2143914 | FAAP20 | 199990 | 2.39093510710338 | 0.00751231 | 7 | notSpecific | Up | DM | Gain | 9 |
| **1_2143147_2143914_2782** | 1_2143147_2143914 | GNB1 | 2782 | 5.50692289049268 | 0.79251188 | 7 | notSpecific | DownUp | DM | Gain | 9 |
| **1_2143147_2143914_5192** | 1_2143147_2143914 | PEX10 | 5192 | 5.30124057742575 | 0.74211689 | 7 | notSpecific | Up | DM | Gain | 9 |

| | | | | | | | | | | | |
|---|---|---|---|---|---|---|---|---|---|---|---|
| **1_217262425_217264006_2104** | 1_217262425_217264006 | ESRRG | 2104 | 0 | 0.00060687 | 4 | only_DIvsD7 | UpDown | notDM | Loss | 3 |
| **1_217309051_217309383_2104** | 1_217309051_217309383 | ESRRG | 2104 | 3.23401081758718 | 0 | 4 | only_DIvsD7 | UpDown | notDM | Gain | 1 |
| **1_21965461_21966267_84196** | 1_21965461_21966267 | USP48 | 84196 | 5.15670963497883 | -0.77365091 | 7 | only_H3vsUI | Down | notDM | Gain | 2 |
| **1_22107727_22109213_84196** | 1_22107727_22109213 | USP48 | 84196 | 2.4345689040342 | -0.38803463 | 2 | only_H3vsUI | Down | notDM | Loss | 8 |
| **1_226068291_226071923_10637** | 1_226068291_226071923 | LEFTY1 | 10637 | 3.69231804425928 | -0.11936458 | 1 | only_DIvsD7 | DownUp | notDM | Loss | 6 |
| **1_227914874_227916121_79169** | 1_227914874_227916121 | C1orf35 | 79169 | 5.57399652277935 | -0.96831942 | 1 | only_H3vsUI | DownUp | notDM | Gain | 2 |
| **1_227921359_227922902_79169** | 1_227921359_227922902 | C1orf35 | 79169 | 5.56607080939705 | 0.78120341 | 1 | only_H3vsUI | DownUp | notDM | Loss | 6 |
| **1_227923378_227925078_79169** | 1_227923378_227925078 | C1orf35 | 79169 | 5.56349651088075 | 0.78120341 | 2 | only_H3vsUI | DownUp | notDM | Loss | 6 |
| **1_227950342_227954565_79169** | 1_227950342_227954565 | C1orf35 | 79169 | 5.52701862344951 | 0.92206474 | 7 | only_H3vsUI | DownUp | notDM | Loss | 5 |
| **1_228230659_228233040_79169** | 1_228230659_228233040 | C1orf35 | 79169 | 4.76419855665556 | -0.67057057 | 6 | only_H3vsUI | DownUp | notDM | Gain | 1 |
| **1_228268565_228270184_79169** | 1_228268565_228270184 | C1orf35 | 79169 | 4.32149486673959 | -0.36123167 | 1 | only_H3vsUI | DownUp | notDM | Loss | 7 |
| **1_228270923_228276637_375** | 1_228270923_228276637 | ARF1 | 375 | 2.63848925695464 | 0.66629815 | 1 | notSpecific | Up | DM | Loss | 10 |
| **1_228270923_228276637_79169** | 1_228270923_228276637 | C1orf35 | 79169 | 4.16172726971602 | -0.24085791 | 1 | only_H3vsUI | DownUp | DM | Loss | 10 |
| **1_228295676_228296878_79169** | 1_228295676_228296878 | C1orf35 | 79169 | 3.66162340922923 | -0.85272008 | 1 | only_H3vsUI | DownUp | notDM | Loss | 7 |
| **1_228297149_228298398_79169** | 1_228297149_228298398 | C1orf35 | 79169 | 3.78254428401001 | -0.85272008 | 1 | only_H3vsUI | DownUp | notDM | Loss | 7 |
| **1_228326581_228327582_79169** | 1_228326581_228327582 | C1orf35 | 79169 | 4.5501427090794 | 0.68173675 | 1 | only_H3vsUI | DownUp | notDM | Gain | 4 |
| **1_228328236_228329824_79169** | 1_228328236_228329824 | C1orf35 | 79169 | 4.569935436876 | 0.68173675 | 1 | only_H3vsUI | DownUp | notDM | Loss | 6 |
| **1_228352282_228353193_574432** | 1_228352282_228353193 | IBA57-DT | 574432 | 0 | -0.00290959 | 1 | only_DIvsD7 | DownUp | notDM | Gain | 4 |
| **1_228352282_228353193_79169** | 1_228352282_228353193 | C1orf35 | 79169 | 4.78633964071584 | 0.58287007 | 1 | only_H3vsUI | DownUp | notDM | Gain | 4 |
| **1_228354007_228355044_574432** | 1_228354007_228355044 | IBA57-DT | 574432 | 2.90091306773767 | -0.00097999 | 1 | only_DIvsD7 | DownUp | notDM | Gain | 4 |
| **1_228354007_228355044_79169** | 1_228354007_228355044 | C1orf35 | 79169 | 4.79842273640344 | 0.58107957 | 1 | only_H3vsUI | DownUp | notDM | Gain | 4 |
| **1_228633134_228634165_79169** | 1_228633134_228634165 | C1orf35 | 79169 | 5.53408451600575 | 0.85492287 | 4 | only_H3vsUI | DownUp | notDM | Loss | 3 |
| **1_228675703_228676899_79169** | 1_228675703_228676899 | C1orf35 | 79169 | 5.58502621773448 | 0.93326997 | 2 | only_H3vsUI | DownUp | notDM | Gain | 1 |
| **1_231297294_231298967_128061** | 1_231297294_231298967 | C1orf131 | 128061 | 4.89191082383833 | -0.62575166 | 2 | notSpecific | Down | DM | Loss | 10 |
| **1_231297294_231298967_440730** | 1_231297294_231298967 | TRIM67 | 440730 | 0 | -0.0069035 | 2 | notSpecific | Up | DM | Loss | 10 |
| **1_231468767_231473506_149371** | 1_231468767_231473506 | EXOC8 | 149371 | 1.96848294855394 | -0.24637319 | 2 | notSpecific | Down | DM | Loss | 10 |
| **1_231468767_231473506_83932** | 1_231468767_231473506 | SPRTN | 83932 | 2.46089784275655 | -0.29553593 | 2 | notSpecific | Down | DM | Loss | 10 |
| **1_235400289_235401262_51742** | 1_235400289_235401262 | ARID4B | 51742 | 4.95554825349415 | -0.72665022 | 6 | only_DIvsD7 | UpDown | notDM | Gain | 2 |
| **1_235487880_235490528_51742** | 1_235487880_235490528 | ARID4B | 51742 | 3.00216606175651 | 0.52502903 | 1 | only_DIvsD7 | UpDown | notDM | Loss | 6 |
| **1_235492321_235496030_51742** | 1_235492321_235496030 | ARID4B | 51742 | 2.89817648349768 | 0.52502903 | 1 | only_DIvsD7 | UpDown | notDM | Loss | 6 |
| **1_235564147_235567647_51742** | 1_235564147_235567647 | ARID4B | 51742 | 4.86116387790122 | -0.7484256 | 7 | only_DIvsD7 | UpDown | notDM | Gain | 5 |
| **1_235570313_235573115_51742** | 1_235570313_235573115 | ARID4B | 51742 | 4.89643251440964 | -0.75759816 | 7 | only_DIvsD7 | UpDown | notDM | Gain | 5 |
| **1_23901983_23903761_259283** | 1_23901983_23903761 | MDS2 | 259283 | 3.62582671328571 | 0.00450059 | 7 | only_H3vsUI | UpDown | notDM | Loss | 3 |
| **1_23903777_23906016_259283** | 1_23903777_23906016 | MDS2 | 259283 | 3.29446622616159 | 0.00264468 | 7 | only_H3vsUI | UpDown | notDM | Gain | 1 |

| | | | | | | | | | | | |
|---|---|---|---|---|---|---|---|---|---|---|---|
| **1_24070495_24072631_57095** | 1_24070495_24072631 | PITHD1 | 57095 | 4.50873166944314 | -0.66685895 | 2 | only_H3vsUI | Up | notDM | Loss | 7 |
| **1_24103358_24104665_57095** | 1_24103358_24104665 | PITHD1 | 57095 | 2.36361197989214 | 0.03013323 | 2 | only_H3vsUI | Up | notDM | Gain | 4 |
| **1_24105332_24107360_57095** | 1_24105332_24107360 | PITHD1 | 57095 | 2.64246452024212 | -0.24784403 | 2 | only_H3vsUI | Up | notDM | Loss | 8 |
| **1_24115485_24117537_57095** | 1_24115485_24117537 | PITHD1 | 57095 | 4.02497797209563 | -0.51067121 | 3 | only_H3vsUI | Up | notDM | Gain | 4 |
| **1_24117856_24118629_57095** | 1_24117856_24118629 | PITHD1 | 57095 | 4.11270552103089 | -0.32051233 | 1 | only_H3vsUI | Up | notDM | Loss | 7 |
| **1_24125242_24126689_57095** | 1_24125242_24126689 | PITHD1 | 57095 | 4.30854307178468 | -0.44593617 | 1 | only_H3vsUI | Up | notDM | Loss | 7 |
| **1_24127466_24128319_57095** | 1_24127466_24128319 | PITHD1 | 57095 | 4.35358928157948 | -0.33624537 | 2 | only_H3vsUI | Up | notDM | Loss | 7 |
| **1_24150483_24151706_57095** | 1_24150483_24151706 | PITHD1 | 57095 | 4.65886959220196 | -0.70243967 | 2 | only_H3vsUI | Up | notDM | Gain | 2 |
| **1_24341226_24342136_57095** | 1_24341226_24342136 | PITHD1 | 57095 | 5.37352436791786 | -0.84178515 | 6 | only_H3vsUI | Up | notDM | Gain | 2 |
| **1_244524211_244526215_200159** | 1_244524211_244526215 | C1orf100 | 200159 | 3.91708519064057 | 0.01439394 | 4 | notSpecific | Up | DM | Gain | 9 |
| **1_244524211_244526215_51029** | 1_244524211_244526215 | DESI2 | 51029 | 5.46268393907709 | 0.86800233 | 4 | notSpecific | UpDown | DM | Gain | 9 |
| **1_245316158_245319858_55083** | 1_245316158_245319858 | KIF26B | 55083 | 0 | 0.06223824 | 4 | only_H3vsUI | UpDown | notDM | Loss | 3 |
| **1_245320057_245320629_55083** | 1_245320057_245320629 | KIF26B | 55083 | 3.24846371755103 | -0.00710793 | 2 | only_H3vsUI | UpDown | notDM | Loss | 3 |
| **1_25755932_25756502_55219** | 1_25755932_25756502 | MACO1 | 55219 | 2.95904139232109 | -0.20392864 | 2 | only_H3vsUI | Up | notDM | Gain | 2 |
| **1_25758030_25759795_55219** | 1_25758030_25759795 | MACO1 | 55219 | 2.80888586735981 | 0.75795833 | 2 | only_H3vsUI | Up | notDM | Loss | 6 |
| **1_26323076_26324499_79947** | 1_26323076_26324499 | DHDDS | 79947 | 5.63784882460541 | -0.84221529 | 1 | only_DIvsD7 | UpDown | notDM | Gain | 2 |
| **1_26405825_26407236_149420** | 1_26405825_26407236 | PDIK1L | 149420 | 4.49283020567869 | 0.6838235 | 6 | notSpecific | Down | DM | Loss | 10 |
| **1_26405825_26407236_64793** | 1_26405825_26407236 | CEP85 | 64793 | 5.18598953379288 | 0.73673422 | 6 | notSpecific | Up | DM | Loss | 10 |
| **1_26496574_26497783_79947** | 1_26496574_26497783 | DHDDS | 79947 | 5.41667378533392 | -0.80190413 | 1 | only_DIvsD7 | UpDown | notDM | Loss | 7 |
| **1_26560990_26562681_79947** | 1_26560990_26562681 | DHDDS | 79947 | 5.29272999135092 | -0.79834369 | 1 | only_DIvsD7 | UpDown | notDM | Loss | 7 |
| **1_26676254_26679122_79947** | 1_26676254_26679122 | DHDDS | 79947 | 4.90138202708215 | -0.73469386 | 7 | only_DIvsD7 | UpDown | notDM | Loss | 7 |
| **1_26757682_26758575_3151** | 1_26757682_26758575 | HMGN2 | 3151 | 4.60668225702875 | 0.51903817 | 1 | notSpecific | Down | DM | Gain | 9 |
| **1_26757682_26758575_79947** | 1_26757682_26758575 | DHDDS | 79947 | 2.50514997831991 | -0.23440546 | 1 | only_DIvsD7 | UpDown | DM | Gain | 1 |
| **1_26758908_26760317_79947** | 1_26758908_26760317 | DHDDS | 79947 | 1.20411998265592 | -0.23440546 | 1 | only_DIvsD7 | UpDown | notDM | Loss | 8 |
| **1_26797039_26798726_79947** | 1_26797039_26798726 | DHDDS | 79947 | 4.58160836603206 | -0.45552956 | 2 | only_DIvsD7 | UpDown | notDM | Loss | 7 |
| **1_26799707_26801602_79947** | 1_26799707_26801602 | DHDDS | 79947 | 4.61146839047983 | -0.48572612 | 2 | only_DIvsD7 | UpDown | notDM | Loss | 7 |
| **1_26856577_26858657_79947** | 1_26856577_26858657 | DHDDS | 79947 | 4.98982788083891 | 0.79355966 | 1 | only_DIvsD7 | UpDown | notDM | Loss | 6 |
| **1_27019729_27022000_79947** | 1_27019729_27022000 | DHDDS | 79947 | 5.41639084105839 | -0.79831675 | 5 | only_DIvsD7 | UpDown | notDM | Loss | 7 |
| **1_27024439_27032343_79947** | 1_27024439_27032343 | DHDDS | 79947 | 5.42416265997488 | -0.82320895 | 2 | only_DIvsD7 | UpDown | notDM | Loss | 7 |
| **1_27032456_27034756_79947** | 1_27032456_27034756 | DHDDS | 79947 | 5.4370795828399 | -0.91752553 | 7 | only_DIvsD7 | UpDown | notDM | Gain | 5 |
| **1_27225242_27226708_10726** | 1_27225242_27226708 | NUDC | 10726 | 1.34242268082221 | 0.02088907 | 1 | only_H3vsUI | DownUp | notDM | Loss | 8 |
| **1_27227037_27227770_10726** | 1_27227037_27227770 | NUDC | 10726 | 2.49136169383427 | 0.02088907 | 1 | only_H3vsUI | DownUp | notDM | Gain | 4 |
| **1_27248466_27250680_10726** | 1_27248466_27250680 | NUDC | 10726 | 2.39967372148104 | -0.31679876 | 1 | only_H3vsUI | DownUp | notDM | Loss | 8 |
| **1_27317401_27318985_10726** | 1_27317401_27318985 | NUDC | 10726 | 4.84001822252221 | -0.60838297 | 7 | only_H3vsUI | DownUp | notDM | Loss | 7 |

| | | | | | | | | | | | |
|---|---|---|---|---|---|---|---|---|---|---|---|
| **1_31768172_31769280_2170** | 1_31768172_31769280 | FABP3 | 2170 | 4.88446116271517 | -0.87000416 | 2 | only_DIvsD7 | DownUp | notDM | Loss | 3 |
| **1_31770070_31771875_2170** | 1_31770070_31771875 | FABP3 | 2170 | 4.86950160269722 | -0.87000416 | 1 | only_DIvsD7 | DownUp | notDM | Loss | 3 |
| **1_32083298_32083848_2170** | 1_32083298_32083848 | FABP3 | 2170 | 5.37543863687221 | 0.93800252 | 2 | only_DIvsD7 | DownUp | notDM | Loss | 3 |
| **1_32108814_32110224_2170** | 1_32108814_32110224 | FABP3 | 2170 | 5.4197790223869 | -0.84902381 | 1 | only_DIvsD7 | DownUp | notDM | Gain | 2 |
| **1_32110919_32111792_2170** | 1_32110919_32111792 | FABP3 | 2170 | 5.42324751278082 | -0.84902763 | 1 | only_DIvsD7 | DownUp | notDM | Loss | 3 |
| **1_32228590_32229504_2170** | 1_32228590_32229504 | FABP3 | 2170 | 5.5828232814713 | 0.93591384 | 2 | only_DIvsD7 | DownUp | notDM | Loss | 3 |
| **1_32478348_32479053_8668** | 1_32478348_32479053 | EIF3I | 8668 | 5.31997793840026 | 0.8514832 | 5 | only_DIvsD7 | DownUp | notDM | Gain | 2 |
| **1_32480142_32482796_8668** | 1_32480142_32482796 | EIF3I | 8668 | 5.31212655879285 | 0.8514832 | 1 | only_DIvsD7 | DownUp | notDM | Loss | 6 |
| **1_32644290_32645177_8668** | 1_32644290_32645177 | EIF3I | 8668 | 4.63139303071276 | 0.57328045 | 3 | only_DIvsD7 | DownUp | notDM | Gain | 2 |
| **1_32645733_32648050_8668** | 1_32645733_32648050 | EIF3I | 8668 | 4.60121229031088 | 0.57328045 | 1 | only_DIvsD7 | DownUp | notDM | Loss | 6 |
| **1_32670477_32671176_8668** | 1_32670477_32671176 | EIF3I | 8668 | 4.2252058659659 | 0.59890055 | 1 | only_DIvsD7 | DownUp | notDM | Gain | 1 |
| **1_32671311_32672532_8668** | 1_32671311_32672532 | EIF3I | 8668 | 4.18864729599972 | 0.59890055 | 1 | only_DIvsD7 | DownUp | notDM | Loss | 7 |
| **1_32686379_32687636_8668** | 1_32686379_32687636 | EIF3I | 8668 | 2.52633927738984 | 0.73692629 | 2 | only_DIvsD7 | DownUp | notDM | Loss | 6 |
| **1_32688227_32689895_8668** | 1_32688227_32689895 | EIF3I | 8668 | 1.91907809237607 | -0.02097445 | 1 | only_DIvsD7 | DownUp | notDM | Loss | 8 |
| **1_32756538_32757468_8668** | 1_32756538_32757468 | EIF3I | 8668 | 4.83612781582605 | 0.61057308 | 5 | only_DIvsD7 | DownUp | notDM | Loss | 7 |
| **1_32758145_32759073_8668** | 1_32758145_32759073 | EIF3I | 8668 | 4.84618860999743 | 0.55776005 | 1 | only_DIvsD7 | DownUp | notDM | Loss | 7 |
| **1_33168877_33170319_8668** | 1_33168877_33170319 | EIF3I | 8668 | 5.68206200170878 | 0.83114013 | 7 | only_DIvsD7 | DownUp | notDM | Loss | 5 |
| **1_3713288_3714081_1677** | 1_3713288_3714081 | DFFB | 1677 | 4.77774571959004 | 0.61846315 | 2 | only_H3vsUI | Down | notDM | Gain | 1 |
| **1_3772157_3773231_1677** | 1_3772157_3773231 | DFFB | 1677 | 2.90687353472207 | 0.33798089 | 2 | only_H3vsUI | Down | notDM | Loss | 8 |
| **1_39669499_39672599_8761** | 1_39669499_39672599 | PABPC4 | 8761 | 5.56804206215561 | -0.80147527 | 5 | only_DIvsD7 | DownUp | notDM | Loss | 7 |
| **1_40023996_40025164_8761** | 1_40023996_40025164 | PABPC4 | 8761 | 3.69897000433602 | 0.07274589 | 1 | only_DIvsD7 | DownUp | notDM | Loss | 7 |
| **1_40038581_40041488_8761** | 1_40038581_40041488 | PABPC4 | 8761 | 2.96801571399364 | 0.61825159 | 2 | only_DIvsD7 | DownUp | notDM | Loss | 6 |
| **1_40347429_40348865_8761** | 1_40347429_40348865 | PABPC4 | 8761 | 5.48425996785924 | -0.80516937 | 1 | only_DIvsD7 | DownUp | notDM | Gain | 2 |
| **1_40357255_40360129_8761** | 1_40357255_40360129 | PABPC4 | 8761 | 5.49803196450112 | -0.77582388 | 6 | only_DIvsD7 | DownUp | notDM | Gain | 2 |
| **1_42804399_42807436_79717** | 1_42804399_42807436 | PPCS | 79717 | 5.05984730143806 | -0.82945686 | 3 | only_H3vsUI | DownUp | notDM | Gain | 5 |
| **1_42846672_42847391_79717** | 1_42846672_42847391 | PPCS | 79717 | 4.87414538461843 | 0.85377448 | 2 | only_H3vsUI | DownUp | notDM | Loss | 3 |
| **1_42922514_42924036_79717** | 1_42922514_42924036 | PPCS | 79717 | 2.45331834004704 | 0.50098135 | 1 | only_H3vsUI | DownUp | notDM | Loss | 6 |
| **1_42927483_42928694_79717** | 1_42927483_42928694 | PPCS | 79717 | 3.72181061521255 | 0.56248046 | 2 | only_H3vsUI | DownUp | notDM | Gain | 4 |
| **1_42928966_42930138_79717** | 1_42928966_42930138 | PPCS | 79717 | 3.82949674972018 | 0.34522053 | 2 | only_H3vsUI | DownUp | notDM | Loss | 7 |
| **1_43124252_43125632_79717** | 1_43124252_43125632 | PPCS | 79717 | 5.30543521028796 | -0.71332991 | 1 | only_H3vsUI | DownUp | notDM | Loss | 7 |
| **1_43146583_43147735_64175** | 1_43146583_43147735 | P3H1 | 64175 | 4.92921450373739 | -0.80713575 | 1 | only_H3vsUI | DownUp | notDM | Loss | 7 |
| **1_43149038_43153376_64175** | 1_43149038_43153376 | P3H1 | 64175 | 4.89917459730477 | -0.7626651 | 2 | only_H3vsUI | DownUp | notDM | Gain | 2 |
| **1_43233225_43234530_64175** | 1_43233225_43234530 | P3H1 | 64175 | 2.67394199863409 | -0.25342554 | 1 | only_H3vsUI | DownUp | notDM | Gain | 4 |
| **1_43287807_43289744_79717** | 1_43287807_43289744 | PPCS | 79717 | 5.56297886461296 | -0.86337823 | 7 | only_H3vsUI | DownUp | notDM | Gain | 2 |

| | | | | | | | | | | | |
|---|---|---|---|---|---|---|---|---|---|---|---|
| **1_43432149_43433579_64175** | 1_43432149_43433579 | P3H1 | 64175 | 5.29984928260137 | -0.73457729 | 7 | only_H3vsUI | DownUp | notDM | Gain | 2 |
| **1_45241661_45244369_8891** | 1_45241661_45244369 | EIF2B3 | 8891 | 5.3178356882976 | 0.73446113 | 1 | only_H3vsUI | DownUp | notDM | Loss | 6 |
| **1_45389918_45391041_79654** | 1_45389918_45391041 | HECTD3 | 79654 | 4.93430145907575 | 0.88848397 | 4 | only_H3vsUI | DownUp | notDM | Gain | 1 |
| **1_45389918_45391041_8891** | 1_45389918_45391041 | EIF2B3 | 8891 | 3.10957854690439 | -0.04271108 | 4 | only_H3vsUI | DownUp | notDM | Gain | 4 |
| **1_45450875_45452102_79654** | 1_45450875_45452102 | HECTD3 | 79654 | 4.39619934709574 | -0.35945335 | 1 | only_H3vsUI | DownUp | notDM | Loss | 7 |
| **1_45450875_45452102_8891** | 1_45450875_45452102 | EIF2B3 | 8891 | 2.25767857486918 | -0.48167971 | 1 | only_H3vsUI | DownUp | notDM | Loss | 8 |
| **1_45475409_45476828_79654** | 1_45475409_45476828 | HECTD3 | 79654 | 2.2405492482826 | -0.13218161 | 2 | only_H3vsUI | DownUp | notDM | Loss | 8 |
| **1_45477064_45477785_79654** | 1_45477064_45477785 | HECTD3 | 79654 | 1.69019608002851 | -0.13523855 | 4 | only_H3vsUI | DownUp | notDM | Loss | 3 |
| **1_45477876_45479176_79654** | 1_45477876_45479176 | HECTD3 | 79654 | 2.93500315145365 | -0.13523855 | 1 | only_H3vsUI | DownUp | notDM | Loss | 8 |
| **1_45636867_45639250_8891** | 1_45636867_45639250 | EIF2B3 | 8891 | 5.26620111149449 | -0.74653858 | 6 | only_H3vsUI | DownUp | notDM | Gain | 2 |
| **1_45955790_45956786_10420** | 1_45955790_45956786 | TESK2 | 10420 | 1.47712125471966 | -0.02793704 | 2 | only_DIvsD7 | DownUp | notDM | Loss | 3 |
| **1_45957235_45958063_10420** | 1_45957235_45958063 | TESK2 | 10420 | 2.59879050676312 | 0.53828113 | 4 | only_DIvsD7 | DownUp | notDM | Loss | 3 |
| **1_45965949_45968378_10420** | 1_45965949_45968378 | TESK2 | 10420 | 3.9596613702735 | 0.31747513 | 1 | only_DIvsD7 | DownUp | notDM | Gain | 1 |
| **1_45979944_45986330_10420** | 1_45979944_45986330 | TESK2 | 10420 | 4.36376235886938 | -0.61701403 | 1 | only_DIvsD7 | DownUp | notDM | Loss | 7 |
| **1_45987767_45990271_10420** | 1_45987767_45990271 | TESK2 | 10420 | 4.49039396098156 | -0.55024236 | 5 | only_DIvsD7 | DownUp | notDM | Gain | 1 |
| **1_46050550_46051840_4678** | 1_46050550_46051840 | NASP | 4678 | 2.91115760873998 | 0.34154952 | 1 | only_DIvsD7 | DownUp | notDM | Loss | 7 |
| **1_46149984_46151853_10420** | 1_46149984_46151853 | TESK2 | 10420 | 5.28594192805612 | -0.89222038 | 2 | only_DIvsD7 | DownUp | notDM | Loss | 7 |
| **1_46149984_46151853_3652** | 1_46149984_46151853 | IPP | 3652 | 4.80935767021111 | 0.55666553 | 2 | only_H3vsUI | Down | notDM | Loss | 7 |
| **1_46152978_46153831_10420** | 1_46152978_46153831 | TESK2 | 10420 | 5.29262152260145 | -0.89222038 | 1 | only_DIvsD7 | DownUp | notDM | Gain | 2 |
| **1_46152978_46153831_3652** | 1_46152978_46153831 | IPP | 3652 | 4.79582442409235 | 0.55666553 | 1 | only_H3vsUI | Down | notDM | Gain | 1 |
| **1_46154152_46155352_10420** | 1_46154152_46155352 | TESK2 | 10420 | 5.29521292311844 | -0.89222038 | 1 | only_DIvsD7 | DownUp | notDM | Loss | 7 |
| **1_46154152_46155352_3652** | 1_46154152_46155352 | IPP | 3652 | 4.78511619502192 | 0.55666553 | 1 | only_H3vsUI | Down | notDM | Loss | 7 |
| **1_46215449_46216118_3652** | 1_46215449_46216118 | IPP | 3652 | 2.3096301674259 | 0.55634681 | 2 | only_H3vsUI | Down | notDM | Gain | 1 |
| **1_46215449_46216118_4678** | 1_46215449_46216118 | NASP | 4678 | 5.12943207415557 | -0.6661911 | 2 | only_DIvsD7 | DownUp | notDM | Gain | 2 |
| **1_46267821_46269072_3652** | 1_46267821_46269072 | IPP | 3652 | 4.71182409451883 | -0.48839095 | 5 | only_H3vsUI | Down | notDM | Gain | 2 |
| **1_46269862_46271699_3652** | 1_46269862_46271699 | IPP | 3652 | 4.72870270093655 | -0.55309379 | 2 | only_H3vsUI | Down | notDM | Loss | 7 |
| **1_46356494_46359189_4678** | 1_46356494_46359189 | NASP | 4678 | 5.44053914482955 | -0.92806369 | 7 | only_DIvsD7 | DownUp | notDM | Gain | 5 |
| **1_52519133_52520726_104355143** | 1_52519133_52520726 | TXNDC12-AS1 | 104355143 | 3.40568778667278 | 0 | 1 | only_H3vsUI | UpDown | DM | Loss | 10 |
| **1_52519133_52520726_112970** | 1_52519133_52520726 | KTI12 | 112970 | 4.29329624514846 | 0.2902526 | 1 | notSpecific | UpDown | DM | Loss | 10 |
| **1_52519133_52520726_51060** | 1_52519133_52520726 | TXNDC12 | 51060 | 2.0899051114394 | -0.24559252 | 1 | notSpecific | DownUp | DM | Loss | 10 |
| **1_52519133_52520726_5865** | 1_52519133_52520726 | RAB3B | 5865 | 4.79740604967638 | -0.83780048 | 1 | notSpecific | Down | DM | Loss | 10 |
| **1_52519133_52520726_91408** | 1_52519133_52520726 | BTF3L4 | 91408 | 3.10311925354571 | -0.22807523 | 1 | notSpecific | UpDown | DM | Loss | 10 |
| **1_52521145_52521752_104355143** | 1_52521145_52521752 | TXNDC12-AS1 | 104355143 | 3.65867902858245 | 0 | 1 | only_H3vsUI | UpDown | notDM | Gain | 1 |
| **1_59246114_59247280_3725** | 1_59246114_59247280 | JUN | 3725 | 3.39898106665813 | 0.46134193 | 2 | only_DIvsD7 | Up | notDM | Gain | 7 |

| | | | | | | | | | | | |
|---|---|---|---|---|---|---|---|---|---|---|---|
| **1_59250131_59252029_3725** | 1_59250131_59252029 | JUN | 3725 | 2.54157924394658 | 0.62243159 | 1 | only_DIvsD7 | Up | notDM | Loss | 6 |
| **1_59314811_59316713_3725** | 1_59314811_59316713 | JUN | 3725 | 4.81310039706002 | 0.78408198 | 7 | only_DIvsD7 | Up | notDM | Gain | 5 |
| **1_59607586_59610370_3725** | 1_59607586_59610370 | JUN | 3725 | 5.55364397758961 | 0.83600266 | 7 | only_DIvsD7 | Up | notDM | Gain | 5 |
| **1_59761305_59762120_55277** | 1_59761305_59762120 | FGGY | 55277 | 2.54157924394658 | -0.62007073 | 5 | only_H3vsUI | Up | notDM | Loss | 3 |
| **1_60224871_60225236_55277** | 1_60224871_60225236 | FGGY | 55277 | 3.54765165835997 | 0 | 4 | only_H3vsUI | Up | notDM | Gain | 1 |
| **1_6085111_6086159_388591** | 1_6085111_6086159 | RNF207 | 388591 | 5.25534729382412 | 0.836277 | 2 | only_DIvsD7 | Down | notDM | Loss | 3 |
| **1_6257159_6259187_388591** | 1_6257159_6259187 | RNF207 | 388591 | 3.84528412634799 | -0.00843844 | 2 | only_DIvsD7 | Down | notDM | Loss | 8 |
| **1_6264703_6265326_388591** | 1_6264703_6265326 | RNF207 | 388591 | 2.93651374247889 | 0.80758579 | 3 | only_DIvsD7 | Down | notDM | Gain | 1 |
| **1_6266410_6267347_388591** | 1_6266410_6267347 | RNF207 | 388591 | 2.34830486304816 | 0.87471255 | 1 | only_DIvsD7 | Down | notDM | Loss | 3 |
| **1_6293995_6295613_388591** | 1_6293995_6295613 | RNF207 | 388591 | 4.44416975476857 | 0.56831863 | 2 | only_DIvsD7 | Down | notDM | Loss | 7 |
| **1_6639287_6639878_388591** | 1_6639287_6639878 | RNF207 | 388591 | 5.57182524904083 | 0.9680676 | 2 | only_DIvsD7 | Down | notDM | Gain | 2 |
| **1_6640450_6641434_388591** | 1_6640450_6641434 | RNF207 | 388591 | 5.57317689446655 | 0.9680676 | 1 | only_DIvsD7 | Down | notDM | Gain | 1 |
| **1_67394398_67395712_23169** | 1_67394398_67395712 | SLC35D1 | 23169 | 5.09367028261262 | 0.73665919 | 4 | only_DIvsD7 | UpDown | notDM | Gain | 1 |
| **1_67396728_67401823_23169** | 1_67396728_67401823 | SLC35D1 | 23169 | 5.07173476387976 | 0.73665919 | 2 | only_DIvsD7 | UpDown | notDM | Loss | 6 |
| **1_67409502_67410807_23169** | 1_67409502_67410807 | SLC35D1 | 23169 | 5.03733086293853 | 0.86046086 | 7 | only_DIvsD7 | UpDown | notDM | Gain | 1 |
| **1_67517487_67519209_23169** | 1_67517487_67519209 | SLC35D1 | 23169 | 2.75891189239797 | 0.68478154 | 1 | only_DIvsD7 | UpDown | notDM | Loss | 6 |
| **1_67520186_67520537_23169** | 1_67520186_67520537 | SLC35D1 | 23169 | 2.60852603357719 | 0.62216479 | 4 | only_DIvsD7 | UpDown | notDM | Gain | 1 |
| **1_67965962_67966659_26135** | 1_67965962_67966659 | SERBP1 | 26135 | 4.84426587954094 | 0.55187934 | 4 | notSpecific | Down | DM | Gain | 9 |
| **1_70669490_70670702_55631** | 1_70669490_70670702 | LRRC40 | 55631 | 2.77959649125782 | -0.04412115 | 2 | only_H3vsUI | DownUp | notDM | Loss | 8 |
| **1_70671711_70673694_55631** | 1_70671711_70673694 | LRRC40 | 55631 | 2.61278385671974 | -0.08074225 | 1 | only_H3vsUI | DownUp | notDM | Loss | 8 |
| **1_70686176_70686971_55631** | 1_70686176_70686971 | LRRC40 | 55631 | 4.17245697440059 | -0.28209463 | 1 | only_H3vsUI | DownUp | notDM | Gain | 2 |
| **1_70687579_70689703_55631** | 1_70687579_70689703 | LRRC40 | 55631 | 4.21160104414524 | -0.28209463 | 1 | only_H3vsUI | DownUp | notDM | Gain | 2 |
| **1_761929_762512_57801** | 1_761929_762512 | HES4 | 57801 | 5.23814901621621 | 0.82243872 | 1 | only_H3vsUI | UpDown | notDM | Loss | 3 |
| **1_7830504_7831142_9341** | 1_7830504_7831142 | VAMP3 | 9341 | 2.27415784926368 | 0.65739023 | 2 | only_H3vsUI | DownUp | notDM | Loss | 3 |
| **1_7831759_7832958_9341** | 1_7831759_7832958 | VAMP3 | 9341 | 2.63548374681491 | -0.29489252 | 1 | only_H3vsUI | DownUp | notDM | Loss | 8 |
| **1_78440508_78444310_54810** | 1_78440508_78444310 | GIPC2 | 54810 | 2.96236933567002 | -0.05242067 | 2 | only_H3vsUI | UpDown | notDM | Loss | 6 |
| **1_78445864_78449086_54810** | 1_78445864_78449086 | GIPC2 | 54810 | 2.80617997398389 | -0.05242067 | 1 | only_H3vsUI | UpDown | notDM | Gain | 4 |
| **1_78449196_78451744_54810** | 1_78449196_78451744 | GIPC2 | 54810 | 3.59900923982334 | -0.06919896 | 8 | only_H3vsUI | UpDown | notDM | Gain | 1 |
| **1_84969246_84971405_2787** | 1_84969246_84971405 | GNG5 | 2787 | 2.92634244662566 | 0.78563394 | 1 | only_DIvsD7 | Up | notDM | Gain | 4 |
| **1_84972415_84973444_2787** | 1_84972415_84973444 | GNG5 | 2787 | 2.22788670461367 | 0.78563394 | 1 | only_DIvsD7 | Up | notDM | Loss | 6 |
| **1_85038493_85039951_2787** | 1_85038493_85039951 | GNG5 | 2787 | 4.82116621599834 | 0.60901833 | 2 | only_DIvsD7 | Up | notDM | Gain | 1 |
| **1_85040361_85040944_2787** | 1_85040361_85040944 | GNG5 | 2787 | 4.83324276095665 | 0.71070347 | 2 | only_DIvsD7 | Up | notDM | Loss | 3 |
| **1_85098021_85099270_2787** | 1_85098021_85099270 | GNG5 | 2787 | 5.09959432599779 | -0.68279042 | 6 | only_DIvsD7 | Up | notDM | Gain | 2 |
| **1_85156503_85157606_2787** | 1_85156503_85157606 | GNG5 | 2787 | 5.26542399587234 | 0.90494722 | 1 | only_DIvsD7 | Up | notDM | Gain | 1 |

| | | | | | | | | | | | |
|---|---|---|---|---|---|---|---|---|---|---|---|
| **1_85247583_85248325_2787** | 1_85247583_85248325 | GNG5 | 2787 | 5.4398645761449 | -0.76449134 | 2 | only_DIvsD7 | Up | notDM | Loss | 7 |
| **1_85871811_85873897_23576** | 1_85871811_85873897 | DDAH1 | 23576 | 3.21298618473667 | 0.04721374 | 3 | notSpecific | DownUp | DM | Gain | 9 |
| **1_85871811_85873897_3491** | 1_85871811_85873897 | CCN1 | 3491 | 5.23690992975907 | 0.70209593 | 3 | notSpecific | Up | DM | Gain | 9 |
| **1_85871811_85873897_8915** | 1_85871811_85873897 | BCL10 | 8915 | 5.11285290755318 | 0.85529098 | 3 | notSpecific | Up | DM | Gain | 9 |
| **1_8928968_8937819_100505975** | 1_8928968_8937819 | ENO1-AS1 | 100505975 | 3.03181227133037 | -0.0705588 | 5 | only_DIvsD7 | DownUp | notDM | Loss | 6 |
| **1_892946_894419_57801** | 1_892946_894419 | HES4 | 57801 | 4.61420094372898 | 0.78706386 | 2 | only_H3vsUI | UpDown | notDM | Loss | 6 |
| **1_89354838_89356734_2959** | 1_89354838_89356734 | GTF2B | 2959 | 2.67851837904011 | 0.60118015 | 1 | notSpecific | Up | DM | Loss | 10 |
| **1_8949069_8951793_100505975** | 1_8949069_8951793 | ENO1-AS1 | 100505975 | 4.0076197745174 | -0.11620088 | 7 | only_DIvsD7 | DownUp | notDM | Gain | 1 |
| **1_92545341_92545784_284697** | 1_92545341_92545784 | BTBD8 | 284697 | 1.89762709129044 | 0.05829137 | 2 | only_DIvsD7 | Down | notDM | Loss | 3 |
| **1_92546222_92547338_284697** | 1_92546222_92547338 | BTBD8 | 284697 | 2.11058971029925 | 0.09015916 | 2 | only_DIvsD7 | Down | notDM | Gain | 4 |
| **1_948126_948750_57801** | 1_948126_948750 | HES4 | 57801 | 4.09954252869533 | 0.59490461 | 2 | only_H3vsUI | UpDown | notDM | Gain | 1 |
| **1_9649359_9650458_116362** | 1_9649359_9650458 | RBP7 | 116362 | 5.60939909285427 | -0.78776031 | 2 | only_DIvsD7 | UpDown | notDM | Loss | 3 |
| **1_9970829_9971857_116362** | 1_9970829_9971857 | RBP7 | 116362 | 4.93154939852217 | -0.80614992 | 5 | only_DIvsD7 | UpDown | notDM | Loss | 3 |
| **10_101677074_101679304_282991** | 10_101677074_101679304 | BLOC1S2 | 282991 | 5.56482460604751 | 0.8387251 | 5 | only_H3vsUI | Down | notDM | Gain | 5 |
| **10_101943336_101945588_282991** | 10_101943336_101945588 | BLOC1S2 | 282991 | 5.00368451514523 | -0.67043932 | 1 | only_H3vsUI | Down | notDM | Loss | 2 |
| **10_101945983_101948127_282991** | 10_101945983_101948127 | BLOC1S2 | 282991 | 4.99261094870673 | -0.67043932 | 1 | only_H3vsUI | Down | notDM | Loss | 2 |
| **10_102025592_102027280_282991** | 10_102025592_102027280 | BLOC1S2 | 282991 | 4.28239550474253 | -0.45978821 | 2 | only_H3vsUI | Down | notDM | Loss | 7 |
| **10_102027520_102028632_282991** | 10_102027520_102028632 | BLOC1S2 | 282991 | 4.25061514699063 | -0.45853047 | 2 | only_H3vsUI | Down | notDM | Gain | 4 |
| **10_102043721_102046236_282991** | 10_102043721_102046236 | BLOC1S2 | 282991 | 0 | -0.1827012 | 1 | only_H3vsUI | Down | notDM | Gain | 4 |
| **10_102046543_102047243_282991** | 10_102046543_102047243 | BLOC1S2 | 282991 | 2.02530586526477 | -0.09904681 | 5 | only_H3vsUI | Down | notDM | Gain | 4 |
| **10_102059470_102062364_282991** | 10_102059470_102062364 | BLOC1S2 | 282991 | 4.12574143912872 | 0.36840588 | 6 | only_H3vsUI | Down | notDM | Loss | 5 |
| **10_102059470_102062364_90271** | 10_102059470_102062364 | OLMALINC | 90271 | 4.85131339664339 | 0.59238538 | 6 | only_DIvsD7 | DownUp | notDM | Loss | 5 |
| **10_102104711_102106584_282991** | 10_102104711_102106584 | BLOC1S2 | 282991 | 4.76547482962791 | -0.58338944 | 1 | only_H3vsUI | Down | notDM | Loss | 2 |
| **10_102107110_102115112_90271** | 10_102107110_102115112 | OLMALINC | 90271 | 4.26152455647267 | -0.51015884 | 1 | only_DIvsD7 | DownUp | notDM | Loss | 8 |
| **10_102118526_102126104_90271** | 10_102118526_102126104 | OLMALINC | 90271 | 3.86147466885717 | -0.22000927 | 3 | only_DIvsD7 | DownUp | notDM | Loss | 6 |
| **10_102129131_102133132_90271** | 10_102129131_102133132 | OLMALINC | 90271 | 2.38201704257487 | -0.52867867 | 7 | only_DIvsD7 | DownUp | notDM | Gain | 4 |
| **10_102133321_102135748_90271** | 10_102133321_102135748 | OLMALINC | 90271 | 0 | -0.33844233 | 5 | only_DIvsD7 | DownUp | notDM | Loss | 8 |
| **10_102135921_102137057_282991** | 10_102135921_102137057 | BLOC1S2 | 282991 | 4.95331986065931 | -0.63356077 | 6 | only_H3vsUI | Down | notDM | Gain | 2 |
| **10_102135921_102137057_90271** | 10_102135921_102137057 | OLMALINC | 90271 | 3.40671045860979 | -0.08884832 | 6 | only_DIvsD7 | DownUp | notDM | Gain | 1 |
| **10_102151649_102153023_282991** | 10_102151649_102153023 | BLOC1S2 | 282991 | 5.02206527628701 | -0.75615463 | 6 | only_H3vsUI | Down | notDM | Loss | 2 |
| **10_102151649_102153023_90271** | 10_102151649_102153023 | OLMALINC | 90271 | 4.26195243284413 | 0.25222522 | 6 | only_DIvsD7 | DownUp | notDM | Loss | 1 |
| **10_102295818_102297602_90271** | 10_102295818_102297602 | OLMALINC | 90271 | 5.21071436884006 | -0.76314735 | 1 | only_DIvsD7 | DownUp | notDM | Loss | 7 |
| **10_102314009_102315623_55719** | 10_102314009_102315623 | SLF2 | 55719 | 5.55280687624589 | 0.79328191 | 6 | only_H3vsUI | Down | notDM | Loss | 1 |
| **10_102430385_102430877_90271** | 10_102430385_102430877 | OLMALINC | 90271 | 5.47277838282808 | 0.83004728 | 4 | only_DIvsD7 | DownUp | notDM | Gain | 2 |

| | | | | | | | | | | | |
|---|---|---|---|---|---|---|---|---|---|---|---|
| **10_102489047_102492364_55719** | 10_102489047_102492364 | SLF2 | 55719 | 5.25613300278068 | -0.72403484 | 5 | only_H3vsUI | Down | notDM | Loss | 2 |
| **10_102649154_102651446_55719** | 10_102649154_102651446 | SLF2 | 55719 | 4.32786956875663 | -0.25772707 | 3 | only_H3vsUI | Down | notDM | Loss | 2 |
| **10_102671416_102672542_55719** | 10_102671416_102672542 | SLF2 | 55719 | 2.25285303097989 | 0.4577059 | 5 | only_H3vsUI | Down | notDM | Loss | 8 |
| **10_102673111_102675462_55719** | 10_102673111_102675462 | SLF2 | 55719 | 2.57634135020579 | 0.46127107 | 2 | only_H3vsUI | Down | notDM | Loss | 8 |
| **10_102824840_102826843_81621** | 10_102824840_102826843 | KAZALD1 | 81621 | 3.58274496569128 | 0.60050226 | 1 | notSpecific | Up | DM | Loss | 10 |
| **10_102824840_102826843_84545** | 10_102824840_102826843 | MRPL43 | 84545 | 4.89015264586983 | -0.67577711 | 1 | notSpecific | DownUp | DM | Loss | 10 |
| **10_102973176_102974944_27343** | 10_102973176_102974944 | POLL | 27343 | 5.57178916303633 | 0.8070915 | 4 | only_H3vsUI | DownUp | notDM | Loss | 6 |
| **10_102988713_102989654_27343** | 10_102988713_102989654 | POLL | 27343 | 5.55432073981151 | -0.87388496 | 4 | only_H3vsUI | DownUp | notDM | Loss | 7 |
| **10_103329060_103330125_27343** | 10_103329060_103330125 | POLL | 27343 | 4.24941850723599 | -0.77133111 | 1 | only_H3vsUI | DownUp | notDM | Loss | 7 |
| **10_103330206_103331179_27343** | 10_103330206_103331179 | POLL | 27343 | 4.2252575769241 | -0.82353601 | 2 | only_H3vsUI | DownUp | notDM | Loss | 7 |
| **10_103345975_103347826_27343** | 10_103345975_103347826 | POLL | 27343 | 1.76342799356294 | -0.85269188 | 2 | only_H3vsUI | DownUp | notDM | Gain | 4 |
| **10_103348108_103349419_27343** | 10_103348108_103349419 | POLL | 27343 | 2.35602585719312 | -0.90799296 | 2 | only_H3vsUI | DownUp | notDM | Gain | 4 |
| **10_103452052_103454347_27343** | 10_103452052_103454347 | POLL | 27343 | 5.01714607419451 | 0.84441043 | 1 | only_H3vsUI | DownUp | notDM | Gain | 5 |
| **10_103454939_103455778_27343** | 10_103454939_103455778 | POLL | 27343 | 5.02903457821761 | 0.84441043 | 2 | only_H3vsUI | DownUp | notDM | Loss | 3 |
| **10_103541376_103544363_27343** | 10_103541376_103544363 | POLL | 27343 | 5.2866697471303 | -0.79724327 | 2 | only_H3vsUI | DownUp | notDM | Loss | 7 |
| **10_103590086_103591543_30819** | 10_103590086_103591543 | KCNIP2 | 30819 | 0 | 0 | 5 | only_H3vsUI | UpDown | notDM | Loss | 3 |
| **10_103599872_103600953_30819** | 10_103599872_103600953 | KCNIP2 | 30819 | 2.38916608436453 | 0.09596797 | 4 | only_H3vsUI | UpDown | notDM | Loss | 3 |
| **10_103638532_103640621_27343** | 10_103638532_103640621 | POLL | 27343 | 5.4631894892612 | 0.83280302 | 7 | only_H3vsUI | DownUp | notDM | Gain | 5 |
| **10_103697336_103700993_79004** | 10_103697336_103700993 | CUEDC2 | 79004 | 5.6914581300802 | 0.96422589 | 7 | only_H3vsUI | DownUp | notDM | Loss | 5 |
| **10_104000808_104002344_79004** | 10_104000808_104002344 | CUEDC2 | 79004 | 5.27874902940791 | -0.76761899 | 1 | only_H3vsUI | DownUp | notDM | Loss | 7 |
| **10_104004168_104005060_79004** | 10_104004168_104005060 | CUEDC2 | 79004 | 5.27267455975412 | 0.80415134 | 4 | only_H3vsUI | DownUp | notDM | Gain | 1 |
| **10_104152552_104155465_79004** | 10_104152552_104155465 | CUEDC2 | 79004 | 4.5676614544739 | 0.556955 | 1 | only_H3vsUI | DownUp | notDM | Loss | 6 |
| **10_104155656_104157042_79004** | 10_104155656_104157042 | CUEDC2 | 79004 | 4.5475778134058 | -0.51476418 | 1 | only_H3vsUI | DownUp | notDM | Loss | 7 |
| **10_104161074_104162072_79004** | 10_104161074_104162072 | CUEDC2 | 79004 | 4.4821157646751 | 0.37590097 | 3 | only_H3vsUI | DownUp | notDM | Loss | 1 |
| **10_104178970_104180822_79004** | 10_104178970_104180822 | CUEDC2 | 79004 | 4.06145247908719 | -0.65622783 | 1 | only_H3vsUI | DownUp | notDM | Loss | 7 |
| **10_104180939_104183153_79004** | 10_104180939_104183153 | CUEDC2 | 79004 | 2.95856388322197 | 0.01345523 | 2 | only_H3vsUI | DownUp | notDM | Loss | 8 |
| **10_104191017_104192388_79004** | 10_104191017_104192388 | CUEDC2 | 79004 | 0 | -0.12248968 | 1 | only_H3vsUI | DownUp | notDM | Loss | 8 |
| **10_104208802_104210208_79004** | 10_104208802_104210208 | CUEDC2 | 79004 | 4.21693085374102 | -0.42642837 | 1 | only_H3vsUI | DownUp | notDM | Gain | 2 |
| **10_104210645_104211799_79004** | 10_104210645_104211799 | CUEDC2 | 79004 | 4.26297287880537 | -0.56239965 | 1 | only_H3vsUI | DownUp | notDM | Loss | 7 |
| **10_104220570_104222120_79004** | 10_104220570_104222120 | CUEDC2 | 79004 | 4.45072626202188 | -0.66062929 | 1 | only_H3vsUI | DownUp | notDM | Loss | 7 |
| **10_104260315_104261824_79004** | 10_104260315_104261824 | CUEDC2 | 79004 | 4.83186337814383 | -0.78686881 | 2 | only_H3vsUI | DownUp | notDM | Loss | 7 |
| **10_104262667_104263595_79004** | 10_104262667_104263595 | CUEDC2 | 79004 | 4.84722706015004 | -0.62266801 | 4 | only_H3vsUI | DownUp | notDM | Loss | 7 |
| **10_104264021_104265502_79004** | 10_104264021_104265502 | CUEDC2 | 79004 | 4.85550704128735 | -0.5795197 | 1 | only_H3vsUI | DownUp | notDM | Loss | 7 |
| **10_104363386_104365721_79004** | 10_104363386_104365721 | CUEDC2 | 79004 | 5.23311546148003 | 0.72242295 | 7 | only_H3vsUI | DownUp | notDM | Loss | 5 |

| | | | | | | | | | | | |
|---|---|---|---|---|---|---|---|---|---|---|---|
| **10_104401129_104401905_79004** | 10_104401129_104401905 | CUEDC2 | 79004 | 5.31954950058645 | 0.83307393 | 1 | only_H3vsUI | DownUp | notDM | Gain | 1 |
| **10_104403860_104408377_79004** | 10_104403860_104408377 | CUEDC2 | 79004 | 5.32519536598393 | 0.83307393 | 1 | only_H3vsUI | DownUp | notDM | Loss | 6 |
| **10_111674227_111676148_7511** | 10_111674227_111676148 | XPNPEP1 | 7511 | 3.8500945943867 | 0.85675908 | 8 | only_H3vsUI | DownUp | notDM | Loss | 5 |
| **10_111681226_111682941_7511** | 10_111681226_111682941 | XPNPEP1 | 7511 | 2.39619934709574 | 0.86715527 | 2 | only_H3vsUI | DownUp | notDM | Gain | 4 |
| **10_111683448_111684065_7511** | 10_111683448_111684065 | XPNPEP1 | 7511 | 2.40823996531185 | 0.86706089 | 2 | only_H3vsUI | DownUp | notDM | Loss | 3 |
| **10_111724795_111726374_7511** | 10_111724795_111726374 | XPNPEP1 | 7511 | 4.61834101587079 | 0.66925072 | 6 | only_H3vsUI | DownUp | notDM | Gain | 1 |
| **10_111765383_111767517_120** | 10_111765383_111767517 | ADD3 | 120 | 2.69372694892365 | 0.25362809 | 2 | notSpecific | Up | DM | Loss | 10 |
| **10_112082990_112085515_7511** | 10_112082990_112085515 | XPNPEP1 | 7511 | 5.6018460486079 | 0.85888485 | 3 | only_H3vsUI | DownUp | notDM | Loss | 5 |
| **10_112155875_112158448_7511** | 10_112155875_112158448 | XPNPEP1 | 7511 | 5.6745203659497 | 0.93208286 | 7 | only_H3vsUI | DownUp | notDM | Gain | 5 |
| **10_112166731_112170268_7511** | 10_112166731_112170268 | XPNPEP1 | 7511 | 5.68438300475143 | 0.86854167 | 7 | only_H3vsUI | DownUp | notDM | Gain | 5 |
| **10_112170409_112174376_7511** | 10_112170409_112174376 | XPNPEP1 | 7511 | 5.68767429623806 | 0.89747213 | 7 | only_H3vsUI | DownUp | notDM | Gain | 5 |
| **10_112644938_112646916_27250** | 10_112644938_112646916 | PDCD4 | 27250 | 2.78958071216443 | 0.39330621 | 3 | notSpecific | Up | DM | Gain | 9 |
| **10_11601634_11603261_254427** | 10_11601634_11603261 | PROSER2 | 254427 | 5.4184305658546 | -0.89590026 | 6 | only_H3vsUI | Up | notDM | Gain | 2 |
| **10_11644321_11646209_254427** | 10_11644321_11646209 | PROSER2 | 254427 | 5.34070183873168 | -0.73243668 | 8 | only_H3vsUI | Up | notDM | Gain | 2 |
| **10_11651022_11653088_254427** | 10_11651022_11653088 | PROSER2 | 254427 | 5.32684974505714 | -0.88432611 | 2 | only_H3vsUI | Up | notDM | Gain | 2 |
| **10_11653851_11654741_254427** | 10_11653851_11654741 | PROSER2 | 254427 | 5.32345424247583 | -0.81301904 | 2 | only_H3vsUI | Up | notDM | Gain | 2 |
| **10_11865776_11868423_254427** | 10_11865776_11868423 | PROSER2 | 254427 | 2.58092497567562 | 0.64355477 | 2 | only_H3vsUI | Up | notDM | Loss | 6 |
| **10_11872231_11875102_254427** | 10_11872231_11875102 | PROSER2 | 254427 | 0 | 0.20020449 | 6 | only_H3vsUI | Up | notDM | Gain | 4 |
| **10_11916211_11917943_254427** | 10_11916211_11917943 | PROSER2 | 254427 | 4.70600047641098 | 0.64922119 | 7 | only_H3vsUI | Up | notDM | Gain | 5 |
| **10_120327627_120328409_143384** | 10_120327627_120328409 | CACUL1 | 143384 | 5.26986770659952 | 0.92990965 | 4 | only_H3vsUI | DownUp | notDM | Gain | 1 |
| **10_120488346_120489446_143384** | 10_120488346_120489446 | CACUL1 | 143384 | 2.67851837904011 | 0.00093703 | 6 | only_H3vsUI | DownUp | notDM | Loss | 7 |
| **10_120511362_120514216_143384** | 10_120511362_120514216 | CACUL1 | 143384 | 2.53781909507327 | 0.29144542 | 1 | only_H3vsUI | DownUp | notDM | Gain | 4 |
| **10_120514843_120515698_143384** | 10_120514843_120515698 | CACUL1 | 143384 | 2.45484486000851 | 0.29144542 | 1 | only_H3vsUI | DownUp | notDM | Loss | 8 |
| **10_120629710_120631510_143384** | 10_120629710_120631510 | CACUL1 | 143384 | 5.06127148534342 | 0.75486631 | 6 | only_H3vsUI | DownUp | notDM | Gain | 1 |
| **10_120840479_120841753_404636** | 10_120840479_120841753 | DENND10 | 404636 | 4.33937192931759 | -0.27336249 | 1 | only_H3vsUI | DownUp | notDM | Loss | 3 |
| **10_120859763_120861124_404636** | 10_120859763_120861124 | DENND10 | 404636 | 3.39357520326959 | 0.45129449 | 6 | only_H3vsUI | DownUp | notDM | Loss | 3 |
| **10_120864193_120865761_404636** | 10_120864193_120865761 | DENND10 | 404636 | 2.77597433112937 | -0.54813159 | 2 | only_H3vsUI | DownUp | notDM | Gain | 4 |
| **10_120872545_120874247_404636** | 10_120872545_120874247 | DENND10 | 404636 | 0 | 0.03054236 | 3 | only_H3vsUI | DownUp | notDM | Gain | 4 |
| **10_120879913_120880398_404636** | 10_120879913_120880398 | DENND10 | 404636 | 1.32221929473392 | 0 | 7 | only_H3vsUI | DownUp | notDM | Gain | 1 |
| **10_120890472_120891899_404636** | 10_120890472_120891899 | DENND10 | 404636 | 4.42933231282833 | -0.90810068 | 5 | only_H3vsUI | DownUp | notDM | Loss | 3 |
| **10_120923864_120925104_404636** | 10_120923864_120925104 | DENND10 | 404636 | 4.7800867795984 | -0.65543906 | 1 | only_H3vsUI | DownUp | notDM | Loss | 3 |
| **10_120925299_120926437_404636** | 10_120925299_120926437 | DENND10 | 404636 | 4.79030627994527 | -0.65543906 | 1 | only_H3vsUI | DownUp | notDM | Loss | 3 |
| **10_123885466_123888501_5654** | 10_123885466_123888501 | HTRA1 | 5654 | 5.52184781040692 | 0.91355663 | 8 | only_H3vsUI | DownUp | notDM | Loss | 5 |
| **10_123897264_123902922_5654** | 10_123897264_123902922 | HTRA1 | 5654 | 5.50259370415364 | 0.83598565 | 7 | only_H3vsUI | DownUp | notDM | Loss | 5 |

| | | | | | | | | | | | |
|---|---|---|---|---|---|---|---|---|---|---|---|
| **10_124132564_124133959_5654** | 10_124132564_124133959 | HTRA1 | 5654 | 4.93993338202504 | 0.67034123 | 1 | only_H3vsUI | DownUp | notDM | Loss | 6 |
| **10_124134831_124137155_5654** | 10_124134831_124137155 | HTRA1 | 5654 | 4.92370501741396 | 0.61048717 | 1 | only_H3vsUI | DownUp | notDM | Gain | 5 |
| **10_124220310_124220847_5654** | 10_124220310_124220847 | HTRA1 | 5654 | 2.29003461136252 | 0.63460547 | 2 | only_H3vsUI | DownUp | notDM | Loss | 3 |
| **10_124221463_124222817_5654** | 10_124221463_124222817 | HTRA1 | 5654 | 2.62736585659273 | 0.95195244 | 1 | only_H3vsUI | DownUp | notDM | Loss | 6 |
| **10_124227215_124229586_5654** | 10_124227215_124229586 | HTRA1 | 5654 | 3.79070728732768 | 0.95532393 | 3 | only_H3vsUI | DownUp | notDM | Gain | 5 |
| **10_124263670_124264908_5654** | 10_124263670_124264908 | HTRA1 | 5654 | 3.11394335230684 | 0.00423828 | 6 | only_H3vsUI | DownUp | notDM | Gain | 1 |
| **10_124609080_124615404_5654** | 10_124609080_124615404 | HTRA1 | 5654 | 5.58887537671176 | 0.90225532 | 6 | only_H3vsUI | DownUp | notDM | Gain | 5 |
| **10_124620777_124627663_5654** | 10_124620777_124627663 | HTRA1 | 5654 | 5.60177543524002 | 0.93248246 | 8 | only_H3vsUI | DownUp | notDM | Gain | 5 |
| **10_131906712_131908801_387723** | 10_131906712_131908801 | C10orf143 | 387723 | 2.44870631990508 | -0.00308538 | 2 | only_DIvsD7 | DownUp | notDM | Loss | 6 |
| **10_131909601_131911457_387723** | 10_131909601_131911457 | C10orf143 | 387723 | 2.71767050300226 | -0.07611582 | 1 | only_DIvsD7 | DownUp | notDM | Gain | 4 |
| **10_134775678_134776749_1019275** | 10_134775678_134776749 | LINC01166 | 101927590 | 3.31069331234336 | 0 | 2 | only_H3vsUI | UpDown | notDM | Loss | 3 |
| **10_16933431_16934006_7431** | 10_16933431_16934006 | VIM | 7431 | 5.52969080161011 | 0.86479391 | 6 | only_DIvsD7 | Up | notDM | Loss | 5 |
| **10_17070946_17072217_7431** | 10_17070946_17072217 | VIM | 7431 | 5.30188037974408 | 0.80139607 | 4 | only_DIvsD7 | Up | notDM | Gain | 1 |
| **10_17156901_17158939_7431** | 10_17156901_17158939 | VIM | 7431 | 5.05564585999466 | 0.67845581 | 2 | only_DIvsD7 | Up | notDM | Gain | 5 |
| **10_17241167_17243385_7431** | 10_17241167_17243385 | VIM | 7431 | 4.44569758989516 | 0.79678142 | 1 | only_DIvsD7 | Up | notDM | Loss | 6 |
| **10_17243744_17244278_7431** | 10_17243744_17244278 | VIM | 7431 | 4.43157281858565 | 0.76105523 | 3 | only_DIvsD7 | Up | notDM | Loss | 5 |
| **10_17254485_17259199_7431** | 10_17254485_17259199 | VIM | 7431 | 4.08249813850578 | 0.70596399 | 7 | only_DIvsD7 | Up | notDM | Loss | 5 |
| **10_17268617_17270511_7431** | 10_17268617_17270511 | VIM | 7431 | 2.89209460269048 | 0.78868817 | 5 | only_DIvsD7 | Up | notDM | Gain | 5 |
| **10_17272620_17276370_7431** | 10_17272620_17276370 | VIM | 7431 | 3.12450422483428 | 0.78722268 | 1 | only_DIvsD7 | Up | notDM | Gain | 4 |
| **10_17552751_17555739_7431** | 10_17552751_17555739 | VIM | 7431 | 5.44942131232427 | 0.75757807 | 7 | only_DIvsD7 | Up | notDM | Gain | 5 |
| **10_17712661_17713901_7431** | 10_17712661_17713901 | VIM | 7431 | 5.64480576265061 | -0.83315085 | 8 | only_DIvsD7 | Up | notDM | Loss | 7 |
| **10_21336455_21337895_8028** | 10_21336455_21337895 | MLLT10 | 8028 | 5.68608008782795 | -0.87266784 | 6 | only_H3vsUI | Down | notDM | Loss | 2 |
| **10_21782651_21784738_8028** | 10_21782651_21784738 | MLLT10 | 8028 | 4.58726184969253 | 0.5902103 | 1 | only_H3vsUI | Down | notDM | Loss | 4 |
| **10_21803878_21805652_8028** | 10_21803878_21805652 | MLLT10 | 8028 | 4.24603055119058 | -0.35146774 | 1 | only_H3vsUI | Down | notDM | Loss | 8 |
| **10_21806389_21809372_8028** | 10_21806389_21809372 | MLLT10 | 8028 | 4.14693383456276 | 0.42497136 | 2 | only_H3vsUI | Down | notDM | Gain | 4 |
| **10_21813010_21814883_8028** | 10_21813010_21814883 | MLLT10 | 8028 | 3.9238654751855 | 0.48074253 | 1 | only_H3vsUI | Down | notDM | Gain | 4 |
| **10_21815567_21816094_8028** | 10_21815567_21816094 | MLLT10 | 8028 | 3.85618492672717 | 0.48074253 | 5 | only_H3vsUI | Down | notDM | Loss | 7 |
| **10_21816263_21822838_8028** | 10_21816263_21822838 | MLLT10 | 8028 | 2.63848925695464 | 0.78392443 | 5 | only_H3vsUI | Down | notDM | Gain | 4 |
| **10_21824194_21825659_8028** | 10_21824194_21825659 | MLLT10 | 8028 | 2.89652621748956 | -0.1242453 | 1 | only_H3vsUI | Down | notDM | Loss | 8 |
| **10_22012808_22014627_8028** | 10_22012808_22014627 | MLLT10 | 8028 | 5.27769171096085 | 0.76236363 | 6 | only_H3vsUI | Down | notDM | Loss | 5 |
| **10_27387979_27389147_22852** | 10_27387979_27389147 | ANKRD26 | 22852 | 2.2405492482826 | -0.05618074 | 1 | only_DIvsD7 | Down | notDM | Loss | 3 |
| **10_27389518_27390418_22852** | 10_27389518_27390418 | ANKRD26 | 22852 | 1.99563519459755 | -0.16000841 | 2 | only_DIvsD7 | Down | notDM | Loss | 3 |
| **10_27529956_27531012_22852** | 10_27529956_27531012 | ANKRD26 | 22852 | 5.14779067854977 | 0.65918751 | 5 | only_DIvsD7 | Down | notDM | Loss | 3 |
| **10_27531215_27533303_22852** | 10_27531215_27533303 | ANKRD26 | 22852 | 5.15166397977208 | 0.74915307 | 1 | only_DIvsD7 | Down | notDM | Loss | 6 |

| | | | | | | | | | | | |
|---|---|---|---|---|---|---|---|---|---|---|---|
| **10_28031102_28035245_283078** | 10_28031102_28035245 | MKX | 283078 | 0 | -0.00621248 | 2 | notSpecific | Up | DM | Loss | 10 |
| **10_32200302_32204053_80314** | 10_32200302_32204053 | EPC1 | 80314 | 5.63554506647507 | -0.93255525 | 8 | only_DIvsD7 | Up | notDM | Loss | 7 |
| **10_32632796_32635201_80314** | 10_32632796_32635201 | EPC1 | 80314 | 2.97589113640179 | -0.8804649 | 2 | only_DIvsD7 | Up | notDM | Gain | 4 |
| **10_32636395_32640544_80314** | 10_32636395_32640544 | EPC1 | 80314 | 2.39967372148104 | -0.88966681 | 5 | only_DIvsD7 | Up | notDM | Gain | 4 |
| **10_32665003_32669321_80314** | 10_32665003_32669321 | EPC1 | 80314 | 0 | 0.18834213 | 1 | only_DIvsD7 | Up | notDM | Gain | 8 |
| **10_33614337_33616977_8829** | 10_33614337_33616977 | NRP1 | 8829 | 3.81117269506655 | 0.84370756 | 8 | notSpecific | Down | DM | Gain | 9 |
| **10_35929928_35933445_8325** | 10_35929928_35933445 | FZD8 | 8325 | 0 | 0.00457997 | 2 | notSpecific | Up | DM | Loss | 10 |
| **10_38144226_38146489_7587** | 10_38144226_38146489 | ZNF37A | 7587 | 5.37433767948755 | 0.75970633 | 1 | only_DIvsD7 | UpDown | notDM | Loss | 6 |
| **10_38146718_38148113_7587** | 10_38146718_38148113 | ZNF37A | 7587 | 5.37134867683192 | 0.75970509 | 1 | only_DIvsD7 | UpDown | notDM | Gain | 1 |
| **10_38265736_38266480_7587** | 10_38265736_38266480 | ZNF37A | 7587 | 5.06738706507458 | 0.74291581 | 1 | only_DIvsD7 | UpDown | notDM | Gain | 1 |
| **10_38298565_38299354_7587** | 10_38298565_38299354 | ZNF37A | 7587 | 4.92389652468197 | 0.77669473 | 2 | only_DIvsD7 | UpDown | notDM | Gain | 1 |
| **10_38300086_38301423_7587** | 10_38300086_38301423 | ZNF37A | 7587 | 4.91305582402144 | 0.77669473 | 1 | only_DIvsD7 | UpDown | notDM | Loss | 6 |
| **10_38382541_38383156_7587** | 10_38382541_38383156 | ZNF37A | 7587 | 2.03742649794062 | 0.66811419 | 4 | only_DIvsD7 | UpDown | notDM | Loss | 3 |
| **10_38383361_38384659_7587** | 10_38383361_38384659 | ZNF37A | 7587 | 1.99563519459755 | 0.6035396 | 1 | only_DIvsD7 | UpDown | notDM | Loss | 6 |
| **10_43571137_43572336_5979** | 10_43571137_43572336 | RET | 5979 | 2.14612803567824 | -0.23988527 | 2 | only_H3vsUI | UpDown | notDM | Loss | 3 |
| **10_45453290_45454827_220980** | 10_45453290_45454827 | TMEM72-AS1 | 220980 | 2.49276038902684 | 0.01945205 | 1 | only_H3vsUI | UpDown | notDM | Loss | 3 |
| **10_45455238_45456025_220980** | 10_45455238_45456025 | TMEM72-AS1 | 220980 | 2.01283722470517 | 0.03085082 | 2 | only_H3vsUI | UpDown | notDM | Gain | 1 |
| **10_4766768_4769705_100216001** | 10_4766768_4769705 | MANCR | 100216001 | 4.66674255755647 | 0.86657419 | 7 | notSpecific | DownUp | DM | Gain | 9 |
| **10_49812084_49813928_58504** | 10_49812084_49813928 | ARHGAP22 | 58504 | 0 | 0.7885004 | 2 | notSpecific | Down | DM | Loss | 10 |
| **10_53457575_53459308_23283** | 10_53457575_53459308 | CSTF2T | 23283 | 1.68124123737559 | -0.02065621 | 1 | only_H3vsUI | Down | notDM | Loss | 8 |
| **10_53459446_53460504_23283** | 10_53459446_53460504 | CSTF2T | 23283 | 1.96848294855394 | -0.02065621 | 1 | only_H3vsUI | Down | notDM | Gain | 4 |
| **10_6621068_6622057_5588** | 10_6621068_6622057 | PRKCQ | 5588 | 2.26007138798507 | 0.21874816 | 1 | only_H3vsUI | UpDown | notDM | Loss | 3 |
| **10_6622553_6624163_5588** | 10_6622553_6624163 | PRKCQ | 5588 | 2.50105926221775 | -0.20493737 | 1 | only_H3vsUI | UpDown | notDM | Loss | 8 |
| **10_71227902_71228783_1305** | 10_71227902_71228783 | COL13A1 | 1305 | 5.52232162256252 | 0.89150031 | 7 | only_H3vsUI | Up | notDM | Loss | 5 |
| **10_71233810_71235663_1305** | 10_71233810_71235663 | COL13A1 | 1305 | 5.51325223567 | 0.90181614 | 7 | only_H3vsUI | Up | notDM | Loss | 5 |
| **10_71528060_71530299_1305** | 10_71528060_71530299 | COL13A1 | 1305 | 4.49618212925727 | 0.97048191 | 6 | only_H3vsUI | Up | notDM | Loss | 5 |
| **10_71560417_71562887_1305** | 10_71560417_71562887 | COL13A1 | 1305 | 0 | 0.63822899 | 2 | only_H3vsUI | Up | notDM | Loss | 3 |
| **10_71566907_71567660_1305** | 10_71566907_71567660 | COL13A1 | 1305 | 3.71775369321072 | 0.71339313 | 6 | only_H3vsUI | Up | notDM | Gain | 1 |
| **10_71622969_71625541_1305** | 10_71622969_71625541 | COL13A1 | 1305 | 4.78734001725697 | 0.62210854 | 6 | only_H3vsUI | Up | notDM | Loss | 5 |
| **10_71628263_71630016_1305** | 10_71628263_71630016 | COL13A1 | 1305 | 4.82332422168878 | 0.6356445 | 6 | only_H3vsUI | Up | notDM | Loss | 5 |
| **10_72043230_72044209_64106** | 10_72043230_72044209 | NPFFR1 | 64106 | 0 | 0 | 4 | notSpecific | DownUp | DM | Loss | 10 |
| **10_76584039_76585006_23522** | 10_76584039_76585006 | KAT6B | 23522 | 2.52504480703685 | 0.05679737 | 1 | only_H3vsUI | Down | notDM | Gain | 4 |
| **10_76586915_76588273_23522** | 10_76586915_76588273 | KAT6B | 23522 | 2.71349054309394 | -0.04998997 | 2 | only_H3vsUI | Down | notDM | Gain | 4 |
| **10_77159087_77159747_84858** | 10_77159087_77159747 | ZNF503 | 84858 | 3.24723654950676 | 0.78321604 | 1 | only_DIvsD7 | Up | notDM | Loss | 3 |

| | | | | | | | | | | | |
|---|---|---|---|---|---|---|---|---|---|---|---|
| **10_77159978_77161056_84858** | 10_77159978_77161056 | ZNF503 | 84858 | 2.66086547800387 | 0.78335658 | 1 | only_DIvsD7 | Up | notDM | Loss | 6 |
| **10_77162242_77164396_84858** | 10_77162242_77164396 | ZNF503 | 84858 | 2.86391737695786 | 0.7833655 | 1 | only_DIvsD7 | Up | notDM | Loss | 6 |
| **10_77168142_77169323_84858** | 10_77168142_77169323 | ZNF503 | 84858 | 3.82157902791201 | 0.77864462 | 2 | only_DIvsD7 | Up | notDM | Loss | 6 |
| **10_77169492_77169919_84858** | 10_77169492_77169919 | ZNF503 | 84858 | 3.90205731080847 | 0.77893463 | 1 | only_DIvsD7 | Up | notDM | Loss | 3 |
| **10_77187568_77191228_84858** | 10_77187568_77191228 | ZNF503 | 84858 | 4.41592441297116 | 0.60071724 | 2 | only_DIvsD7 | Up | notDM | Loss | 6 |
| **10_77523824_77525806_84858** | 10_77523824_77525806 | ZNF503 | 84858 | 5.55899160949107 | 0.86711498 | 7 | only_DIvsD7 | Up | notDM | Gain | 5 |
| **10_80827168_80828534_283050** | 10_80827168_80828534 | ZMIZ1-AS1 | 283050 | 0 | 0.01147302 | 1 | notSpecific | UpDown | DM | Loss | 10 |
| **10_80827168_80828534_57178** | 10_80827168_80828534 | ZMIZ1 | 57178 | 2.41329976408125 | 0.78076683 | 1 | notSpecific | Down | DM | Loss | 10 |
| **10_88124626_88126236_2894** | 10_88124626_88126236 | GRID1 | 2894 | 0 | -0.00612638 | 4 | notSpecific | DownUp | DM | Loss | 10 |
| **10_91294180_91295337_387700** | 10_91294180_91295337 | SLC16A12 | 387700 | 0 | -0.01669127 | 4 | only_DIvsD7 | UpDown | notDM | Loss | 3 |
| **10_91312189_91314932_387700** | 10_91312189_91314932 | SLC16A12 | 387700 | 3.16643011384328 | 0 | 6 | only_DIvsD7 | UpDown | notDM | Gain | 1 |
| **10_94403523_94405216_3087** | 10_94403523_94405216 | HHEX | 3087 | 4.6483307317143 | 0.59921161 | 7 | only_H3vsUI | Up | notDM | Loss | 5 |
| **10_94448105_94449432_3087** | 10_94448105_94449432 | HHEX | 3087 | 2.44870631990508 | 0.88409211 | 1 | only_H3vsUI | Up | notDM | Loss | 6 |
| **10_94450358_94455748_3087** | 10_94450358_94455748 | HHEX | 3087 | 2.81157500587059 | 0.88507474 | 1 | only_H3vsUI | Up | notDM | Gain | 4 |
| **10_94455961_94460787_3087** | 10_94455961_94460787 | HHEX | 3087 | 3.7959494989028 | 0.91890034 | 7 | only_H3vsUI | Up | notDM | Gain | 5 |
| **10_94626780_94628500_3087** | 10_94626780_94628500 | HHEX | 3087 | 5.24346468849127 | -0.72003772 | 6 | only_H3vsUI | Up | notDM | Gain | 2 |
| **10_94651087_94655368_3087** | 10_94651087_94655368 | HHEX | 3087 | 5.3040098666968 | 0.8797905 | 3 | only_H3vsUI | Up | notDM | Gain | 5 |
| **10_99255285_99258077_80019** | 10_99255285_99258077 | UBTD1 | 80019 | 2.73957234445009 | -0.12606078 | 1 | only_H3vsUI | Up | notDM | Loss | 8 |
| **10_99259420_99260664_80019** | 10_99259420_99260664 | UBTD1 | 80019 | 2.90145832139611 | 0.16394254 | 2 | only_H3vsUI | Up | notDM | Gain | 4 |
| **11_104838480_104839820_837** | 11_104838480_104839820 | CASP4 | 837 | 0 | 0.94528949 | 2 | only_DIvsD7 | UpDown | notDM | Gain | 4 |
| **11_1071468_1074724_4583** | 11_1071468_1074724 | MUC2 | 4583 | 2.18184358794477 | 0 | 3 | only_H3vsUI | Up | notDM | Loss | 6 |
| **11_107326799_107328391_143884** | 11_107326799_107328391 | CWF19L2 | 143884 | 2.26007138798507 | -0.34042343 | 1 | only_H3vsUI | DownUp | notDM | Loss | 8 |
| **11_107431828_107436110_91801** | 11_107431828_107436110 | ALKBH8 | 91801 | 2.54406804435028 | 0.10468764 | 1 | only_H3vsUI | Down | notDM | Gain | 4 |
| **11_107880403_107882070_38** | 11_107880403_107882070 | ACAT1 | 38 | 5.04228010749836 | 0.7099234 | 2 | only_H3vsUI | DownUp | notDM | Loss | 6 |
| **11_107963564_107965226_38** | 11_107963564_107965226 | ACAT1 | 38 | 4.43165319734658 | -0.44808766 | 3 | only_H3vsUI | DownUp | notDM | Gain | 1 |
| **11_107992490_107993611_38** | 11_107992490_107993611 | ACAT1 | 38 | 2.29666519026153 | 0.45346452 | 2 | only_H3vsUI | DownUp | notDM | Gain | 4 |
| **11_111173025_111175244_5450** | 11_111173025_111175244 | POU2AF1 | 5450 | 4.73947740631835 | 0.997668 | 3 | only_H3vsUI | UpDown | notDM | Gain | 1 |
| **11_111178313_111179624_5450** | 11_111178313_111179624 | POU2AF1 | 5450 | 4.70336017179565 | 0.99865329 | 3 | only_H3vsUI | UpDown | notDM | Gain | 1 |
| **11_11605079_11607399_27122** | 11_11605079_11607399 | DKK3 | 27122 | 5.62638759304879 | 0.94555039 | 3 | only_H3vsUI | UpDown | notDM | Gain | 1 |
| **11_116880765_116883689_6876** | 11_116880765_116883689 | TAGLN | 6876 | 5.2704692064532 | 0.76399798 | 8 | only_H3vsUI | UpDown | notDM | Loss | 5 |
| **11_117049928_117051160_6876** | 11_117049928_117051160 | TAGLN | 6876 | 4.27595598175374 | -0.3885956 | 2 | only_H3vsUI | UpDown | notDM | Loss | 3 |
| **11_117100970_117102691_6876** | 11_117100970_117102691 | TAGLN | 6876 | 4.48959289982189 | -0.66232916 | 1 | only_H3vsUI | UpDown | notDM | Loss | 8 |
| **11_11996522_12000047_27122** | 11_11996522_12000047 | DKK3 | 27122 | 4.48955069775057 | 0.71952341 | 7 | only_H3vsUI | UpDown | notDM | Loss | 5 |
| **11_12008504_12011252_27122** | 11_12008504_12011252 | DKK3 | 27122 | 4.29371603482611 | 0.7482916 | 7 | only_H3vsUI | UpDown | notDM | Gain | 1 |

| | | | | | | | | | | | |
|---|---|---|---|---|---|---|---|---|---|---|---|
| **11_12012896_12016656_27122** | 11_12012896_12016656 | DKK3 | 27122 | 4.12952877385878 | 0.96032909 | 7 | only_H3vsUI | UpDown | notDM | Gain | 1 |
| **11_12018783_12021055_27122** | 11_12018783_12021055 | DKK3 | 27122 | 3.98913822421579 | 0.7733572 | 4 | only_H3vsUI | UpDown | notDM | Loss | 6 |
| **11_12026125_12027436_27122** | 11_12026125_12027436 | DKK3 | 27122 | 3.54182876678131 | 0.93294237 | 6 | only_H3vsUI | UpDown | notDM | Gain | 1 |
| **11_12030347_12030730_27122** | 11_12030347_12030730 | DKK3 | 27122 | 0 | 0.97223416 | 2 | only_H3vsUI | UpDown | notDM | Loss | 3 |
| **11_12031016_12031822_27122** | 11_12031016_12031822 | DKK3 | 27122 | 2.58994960132571 | 0.99161784 | 4 | only_H3vsUI | UpDown | notDM | Loss | 3 |
| **11_12086934_12090321_27122** | 11_12086934_12090321 | DKK3 | 27122 | 4.75198695044155 | 0.60229067 | 7 | only_H3vsUI | UpDown | notDM | Gain | 1 |
| **11_12092421_12094871_27122** | 11_12092421_12094871 | DKK3 | 27122 | 4.78969350127866 | 0.96662889 | 7 | only_H3vsUI | UpDown | notDM | Loss | 5 |
| **11_12100482_12102348_27122** | 11_12100482_12102348 | DKK3 | 27122 | 4.84308944349941 | 0.85807462 | 7 | only_H3vsUI | UpDown | notDM | Gain | 1 |
| **11_12106490_12112758_27122** | 11_12106490_12112758 | DKK3 | 27122 | 4.87900981526319 | 0.73046688 | 7 | only_H3vsUI | UpDown | notDM | Gain | 1 |
| **11_12130670_12131984_27122** | 11_12130670_12131984 | DKK3 | 27122 | 4.99941330634205 | 0.77002578 | 4 | only_H3vsUI | UpDown | notDM | Loss | 6 |
| **11_12132391_12134038_27122** | 11_12132391_12134038 | DKK3 | 27122 | 5.00683386009661 | 0.8651759 | 1 | only_H3vsUI | UpDown | notDM | Loss | 6 |
| **11_12180615_12181981_27122** | 11_12180615_12181981 | DKK3 | 27122 | 5.17554080401677 | 0.748235 | 6 | only_H3vsUI | UpDown | notDM | Loss | 5 |
| **11_12306472_12310805_27122** | 11_12306472_12310805 | DKK3 | 27122 | 5.44021144792458 | 0.77990329 | 7 | only_H3vsUI | UpDown | notDM | Gain | 1 |
| **11_12450374_12453144_27122** | 11_12450374_12453144 | DKK3 | 27122 | 5.62285307450874 | 0.82526389 | 6 | only_H3vsUI | UpDown | notDM | Loss | 5 |
| **11_125755809_125757296_83480** | 11_125755809_125757296 | PUS3 | 83480 | 4.1992339305369 | -0.65879699 | 5 | only_DIvsD7 | UpDown | notDM | Gain | 4 |
| **11_125758082_125764293_83480** | 11_125758082_125764293 | PUS3 | 83480 | 3.28623185402855 | 0.00530861 | 5 | only_DIvsD7 | UpDown | notDM | Loss | 6 |
| **11_125772392_125773966_83480** | 11_125772392_125773966 | PUS3 | 83480 | 0 | -0.53342405 | 2 | only_DIvsD7 | UpDown | notDM | Loss | 3 |
| **11_125949886_125950950_84881** | 11_125949886_125950950 | RPUSD4 | 84881 | 5.11606952295632 | 0.67993405 | 6 | only_H3vsUI | Up | notDM | Gain | 1 |
| **11_126080127_126081356_84881** | 11_126080127_126081356 | RPUSD4 | 84881 | 2.26951294421792 | -0.55658958 | 1 | only_H3vsUI | Up | notDM | Loss | 8 |
| **11_126081919_126083072_84881** | 11_126081919_126083072 | RPUSD4 | 84881 | 2.57978359661681 | -0.55658958 | 1 | only_H3vsUI | Up | notDM | Loss | 8 |
| **11_129870255_129871727_56980** | 11_129870255_129871727 | PRDM10 | 56980 | 2.9795483747041 | -0.25043716 | 1 | only_H3vsUI | UpDown | notDM | Gain | 4 |
| **11_129873327_129874767_56980** | 11_129873327_129874767 | PRDM10 | 56980 | 2.79934054945358 | -0.24912938 | 1 | only_H3vsUI | UpDown | notDM | Gain | 4 |
| **11_1408865_1410766_9024** | 11_1408865_1410766 | BRSK2 | 9024 | 2.8750612633917 | -0.01993565 | 1 | notSpecific | Up | DM | Loss | 10 |
| **11_17411135_17411746_10083** | 11_17411135_17411746 | USH1C | 10083 | 5.18809000655107 | 0.98014827 | 4 | only_H3vsUI | DownUp | notDM | Loss | 3 |
| **11_18812509_18814123_84867** | 11_18812509_18814123 | PTPN5 | 84867 | 0 | 0.09637133 | 4 | only_H3vsUI | Up | notDM | Loss | 3 |
| **11_2422291_2423836_5002** | 11_2422291_2423836 | SLC22A18 | 5002 | 5.69869022859221 | -0.92261339 | 1 | only_DIvsD7 | DownUp | notDM | Loss | 7 |
| **11_2906391_2907079_5002** | 11_2906391_2907079 | SLC22A18 | 5002 | 4.21576970841764 | 0.31534278 | 2 | only_DIvsD7 | DownUp | notDM | Loss | 3 |
| **11_2922407_2923332_5002** | 11_2922407_2923332 | SLC22A18 | 5002 | 2.26007138798507 | 0.65721203 | 1 | only_DIvsD7 | DownUp | notDM | Loss | 3 |
| **11_2923530_2926402_5002** | 11_2923530_2926402 | SLC22A18 | 5002 | 1.27875360095283 | 0.63884926 | 1 | only_DIvsD7 | DownUp | notDM | Loss | 6 |
| **11_2932441_2935311_5002** | 11_2932441_2935311 | SLC22A18 | 5002 | 3.95085145888855 | 0.6995064 | 3 | only_DIvsD7 | DownUp | notDM | Loss | 5 |
| **11_2948162_2950587_5002** | 11_2948162_2950587 | SLC22A18 | 5002 | 4.43477642166315 | 0.39001071 | 1 | only_DIvsD7 | DownUp | notDM | Loss | 6 |
| **11_2950827_2952213_5002** | 11_2950827_2952213 | SLC22A18 | 5002 | 4.47535152292802 | 0.38958513 | 3 | only_DIvsD7 | DownUp | notDM | Loss | 7 |
| **11_2953747_2955190_5002** | 11_2953747_2955190 | SLC22A18 | 5002 | 4.51584736153344 | 0.4294104 | 6 | only_DIvsD7 | DownUp | notDM | Gain | 1 |
| **11_2962007_2962914_5002** | 11_2962007_2962914 | SLC22A18 | 5002 | 4.58541560565975 | 0.61625687 | 7 | only_DIvsD7 | DownUp | notDM | Loss | 5 |

| | | | | | | | | | | | |
|---|---|---|---|---|---|---|---|---|---|---|---|
| **11_3009159_3013001_5002** | 11_3009159_3013001 | SLC22A18 | 5002 | 4.93271722603936 | 0.71766726 | 1 | only_DIvsD7 | DownUp | notDM | Loss | 6 |
| **11_3013832_3014935_5002** | 11_3013832_3014935 | SLC22A18 | 5002 | 4.95578873728801 | 0.71067888 | 1 | only_DIvsD7 | DownUp | notDM | Loss | 6 |
| **11_34705906_34706951_26298** | 11_34705906_34706951 | EHF | 26298 | 4.80089570699307 | 0.77187609 | 7 | notSpecific | Down | DM | Gain | 9 |
| **11_34705906_34706951_960** | 11_34705906_34706951 | CD44 | 960 | 5.69449639234358 | 0.90834946 | 7 | notSpecific | UpDown | DM | Gain | 9 |
| **11_36382408_36383832_79899** | 11_36382408_36383832 | PRR5L | 79899 | 4.14983469671578 | 0.77664681 | 6 | only_H3vsUI | Down | notDM | Gain | 1 |
| **11_36396811_36400350_79899** | 11_36396811_36400350 | PRR5L | 79899 | 0 | 0.77338785 | 2 | only_H3vsUI | Down | notDM | Loss | 8 |
| **11_36441610_36443861_79899** | 11_36441610_36443861 | PRR5L | 79899 | 4.50023647482564 | 0.80589349 | 3 | only_H3vsUI | Down | notDM | Gain | 1 |
| **11_36452270_36454414_79899** | 11_36452270_36454414 | PRR5L | 79899 | 4.32401479803629 | -0.24619303 | 7 | only_H3vsUI | Down | notDM | Loss | 7 |
| **11_36474143_36476292_79899** | 11_36474143_36476292 | PRR5L | 79899 | 0 | -0.0747733 | 3 | only_H3vsUI | Down | notDM | Loss | 4 |
| **11_43958435_43959958_387763** | 11_43958435_43959958 | C11orf96 | 387763 | 3.61257195406518 | 0.03574686 | 6 | only_H3vsUI | Up | notDM | Loss | 3 |
| **11_43964438_43966488_387763** | 11_43964438_43966488 | C11orf96 | 387763 | 2.5854607295085 | 0.00865032 | 4 | only_H3vsUI | Up | notDM | Loss | 3 |
| **11_44997861_45003722_221122** | 11_44997861_45003722 | LINC02685 | 221122 | 3.38201704257487 | 0 | 7 | only_DIvsD7 | DownUp | notDM | Loss | 5 |
| **11_45112262_45114675_56981** | 11_45112262_45114675 | PRDM11 | 56981 | 2.94939000664491 | 0 | 6 | only_H3vsUI | UpDown | notDM | Loss | 3 |
| **11_45167218_45168217_56981** | 11_45167218_45168217 | PRDM11 | 56981 | 1 | 0.05980212 | 1 | only_H3vsUI | UpDown | notDM | Loss | 3 |
| **11_45168939_45170521_56981** | 11_45168939_45170521 | PRDM11 | 56981 | 1.67209785793572 | -0.00215314 | 1 | only_H3vsUI | UpDown | notDM | Loss | 3 |
| **11_45200426_45202085_56981** | 11_45200426_45202085 | PRDM11 | 56981 | 3.12839926871781 | 7.795e-05 | 1 | only_H3vsUI | UpDown | notDM | Loss | 6 |
| **11_45202514_45203999_56981** | 11_45202514_45203999 | PRDM11 | 56981 | 0 | 0.00198129 | 1 | only_H3vsUI | UpDown | notDM | Loss | 3 |
| **11_4627300_4629384_55128** | 11_4627300_4629384 | TRIM68 | 55128 | 1.89209460269048 | 0.34488922 | 1 | only_H3vsUI | Down | notDM | Gain | 4 |
| **11_46951073_46952651_8567** | 11_46951073_46952651 | MADD | 8567 | 5.53027271503066 | 0.92362231 | 7 | only_H3vsUI | DownUp | notDM | Gain | 5 |
| **11_47235202_47236393_53** | 11_47235202_47236393 | ACP2 | 53 | 4.53119781201017 | 0.41672874 | 5 | only_DIvsD7 | Down | notDM | Loss | 7 |
| **11_47236521_47240806_53** | 11_47236521_47240806 | ACP2 | 53 | 4.47077788333512 | 0.43233765 | 1 | only_DIvsD7 | Down | notDM | Loss | 4 |
| **11_47236521_47240806_8567** | 11_47236521_47240806 | MADD | 8567 | 4.70276646769512 | 0.53550286 | 1 | only_H3vsUI | DownUp | notDM | Loss | 6 |
| **11_47268998_47270333_53** | 11_47268998_47270333 | ACP2 | 53 | 1.32221929473392 | -0.01308694 | 1 | only_DIvsD7 | Down | notDM | Loss | 8 |
| **11_47268998_47270333_8567** | 11_47268998_47270333 | MADD | 8567 | 4.32974355001123 | 0.39591576 | 1 | only_H3vsUI | DownUp | notDM | Loss | 7 |
| **11_47270556_47272007_53** | 11_47270556_47272007 | ACP2 | 53 | 2.22271647114758 | -0.01468261 | 1 | only_DIvsD7 | Down | notDM | Loss | 8 |
| **11_47270556_47272007_8567** | 11_47270556_47272007 | MADD | 8567 | 4.29431188090201 | 0.39591576 | 1 | only_H3vsUI | DownUp | notDM | Loss | 7 |
| **11_47278340_47281112_8567** | 11_47278340_47281112 | MADD | 8567 | 4.02481393262931 | 0.41743054 | 3 | only_H3vsUI | DownUp | notDM | Loss | 5 |
| **11_47289862_47291109_53** | 11_47289862_47291109 | ACP2 | 53 | 4.29021274691953 | 0.20752786 | 1 | only_DIvsD7 | Down | notDM | Loss | 7 |
| **11_47289862_47291109_8567** | 11_47289862_47291109 | MADD | 8567 | 2.77158748088126 | 0.5471277 | 1 | only_H3vsUI | DownUp | notDM | Loss | 6 |
| **11_47292152_47293730_8567** | 11_47292152_47293730 | MADD | 8567 | 2.65801139665711 | 0.65314658 | 2 | only_H3vsUI | DownUp | notDM | Loss | 6 |
| **11_5300923_5302744_3046** | 11_5300923_5302744 | HBE1 | 3046 | 3.97941178263444 | 0.16916607 | 7 | notSpecific | Down | DM | Gain | 9 |
| **11_56699328_56701740_85456** | 11_56699328_56701740 | TNKS1BP1 | 85456 | 5.58891007062446 | 0.9860942 | 7 | notSpecific | DownUp | DM | Gain | 9 |
| **11_57016683_57017272_26519** | 11_57016683_57017272 | TIMM10 | 26519 | 5.44860739459344 | -0.76943404 | 7 | only_DIvsD7 | DownUp | notDM | Gain | 2 |
| **11_57092671_57094105_26519** | 11_57092671_57094105 | TIMM10 | 26519 | 5.30999618300418 | -0.90198621 | 1 | only_DIvsD7 | DownUp | notDM | Loss | 7 |

| | | | | | | | | | | | |
|---|---|---|---|---|---|---|---|---|---|---|---|
| **11_57208345_57209138_26519** | 11_57208345_57209138 | TIMM10 | 26519 | 4.95006775767345 | -0.61410607 | 7 | only_DIvsD7 | DownUp | notDM | Gain | 2 |
| **11_57224979_57227810_26519** | 11_57224979_57227810 | TIMM10 | 26519 | 4.84798578262003 | -0.62141225 | 5 | only_DIvsD7 | DownUp | notDM | Loss | 7 |
| **11_57249926_57251442_26519** | 11_57249926_57251442 | TIMM10 | 26519 | 4.66993022532399 | 0.52304229 | 2 | only_DIvsD7 | DownUp | notDM | Gain | 5 |
| **11_57281869_57284035_26519** | 11_57281869_57284035 | TIMM10 | 26519 | 4.15146178712433 | 0.46024615 | 2 | only_DIvsD7 | DownUp | notDM | Gain | 5 |
| **11_57296533_57298116_26519** | 11_57296533_57298116 | TIMM10 | 26519 | 2.20682587603185 | -0.14186275 | 1 | only_DIvsD7 | DownUp | notDM | Loss | 8 |
| **11_57298444_57299791_26519** | 11_57298444_57299791 | TIMM10 | 26519 | 2.23044892137827 | 0.11461306 | 2 | only_DIvsD7 | DownUp | notDM | Loss | 7 |
| **11_57405018_57405706_26519** | 11_57405018_57405706 | TIMM10 | 26519 | 5.02834347301674 | -0.61601891 | 3 | only_DIvsD7 | DownUp | notDM | Gain | 2 |
| **11_57415836_57417256_26519** | 11_57415836_57417256 | TIMM10 | 26519 | 5.07052178904262 | 0.68216919 | 4 | only_DIvsD7 | DownUp | notDM | Loss | 7 |
| **11_57539532_57541168_26519** | 11_57539532_57541168 | TIMM10 | 26519 | 5.3824817232521 | -0.76942463 | 8 | only_DIvsD7 | DownUp | notDM | Gain | 2 |
| **11_58328025_58330655_9404** | 11_58328025_58330655 | LPXN | 9404 | 4.10311925354571 | 0.62793056 | 3 | only_H3vsUI | DownUp | notDM | Gain | 5 |
| **11_58339835_58342150_9404** | 11_58339835_58342150 | LPXN | 9404 | 3.07371835034612 | 0.55928345 | 7 | only_H3vsUI | DownUp | notDM | Gain | 1 |
| **11_58342998_58345557_9404** | 11_58342998_58345557 | LPXN | 9404 | 0 | 0.63064455 | 1 | only_H3vsUI | DownUp | notDM | Loss | 8 |
| **11_58347253_58351255_9404** | 11_58347253_58351255 | LPXN | 9404 | 3.59339684230021 | 0.63064455 | 1 | only_H3vsUI | DownUp | notDM | Gain | 4 |
| **11_58940042_58942982_23220** | 11_58940042_58942982 | DTX4 | 23220 | 1.89762709129044 | 0.02065146 | 2 | only_H3vsUI | Down | notDM | Loss | 8 |
| **11_61195532_61196862_5866** | 11_61195532_61196862 | RAB3IL1 | 5866 | 5.69097533793287 | -0.80896058 | 2 | only_H3vsUI | DownUp | notDM | Loss | 7 |
| **11_61197703_61199472_5866** | 11_61197703_61199472 | RAB3IL1 | 5866 | 5.68866004143607 | -0.80896058 | 1 | only_H3vsUI | DownUp | notDM | Gain | 2 |
| **11_615047_616650_6343** | 11_615047_616650 | SCT | 6343 | 4.02094105986232 | -0.05363236 | 2 | only_H3vsUI | UpDown | notDM | Loss | 3 |
| **11_61519040_61519904_2237** | 11_61519040_61519904 | FEN1 | 2237 | 4.60766943668824 | -0.43017091 | 2 | only_H3vsUI | DownUp | notDM | Loss | 7 |
| **11_61558583_61560334_2237** | 11_61558583_61560334 | FEN1 | 2237 | 0 | 0.18431929 | 1 | only_H3vsUI | DownUp | notDM | Loss | 8 |
| **11_61558583_61560334_5866** | 11_61558583_61560334 | RAB3IL1 | 5866 | 5.105189880947 | 0.69558486 | 1 | only_H3vsUI | DownUp | notDM | Loss | 6 |
| **11_61560425_61562624_2237** | 11_61560425_61562624 | FEN1 | 2237 | 0.602059991327962 | 0.22359214 | 2 | only_H3vsUI | DownUp | notDM | Gain | 8 |
| **11_61560425_61562624_5866** | 11_61560425_61562624 | RAB3IL1 | 5866 | 5.09731285139915 | 0.66132127 | 2 | only_H3vsUI | DownUp | notDM | Gain | 5 |
| **11_61581501_61582622_2237** | 11_61581501_61582622 | FEN1 | 2237 | 4.33029199146816 | 0.43677707 | 1 | only_H3vsUI | DownUp | notDM | Gain | 1 |
| **11_61584700_61589045_2237** | 11_61584700_61589045 | FEN1 | 2237 | 4.38523079511221 | -0.48911733 | 5 | only_H3vsUI | DownUp | notDM | Loss | 7 |
| **11_61595623_61603423_2237** | 11_61595623_61603423 | FEN1 | 2237 | 4.54656733859998 | 0.72229498 | 1 | only_H3vsUI | DownUp | notDM | Loss | 6 |
| **11_61595623_61603423_5866** | 11_61595623_61603423 | RAB3IL1 | 5866 | 4.92592544728667 | 0.59413381 | 1 | only_H3vsUI | DownUp | notDM | Loss | 6 |
| **11_61657243_61658886_5866** | 11_61657243_61658886 | RAB3IL1 | 5866 | 4.46020602745745 | 0.50570474 | 1 | only_H3vsUI | DownUp | notDM | Loss | 6 |
| **11_61684514_61684855_2237** | 11_61684514_61684855 | FEN1 | 2237 | 5.09374728393938 | 0.68964703 | 2 | only_H3vsUI | DownUp | notDM | Loss | 7 |
| **11_61684514_61684855_5866** | 11_61684514_61684855 | RAB3IL1 | 5866 | 2.15533603746506 | 0.35972441 | 2 | only_H3vsUI | DownUp | notDM | Loss | 3 |
| **11_61685157_61685942_5866** | 11_61685157_61685942 | RAB3IL1 | 5866 | 3.25527250510331 | 0.82051555 | 2 | only_H3vsUI | DownUp | notDM | Loss | 3 |
| **11_61687343_61688191_5866** | 11_61687343_61688191 | RAB3IL1 | 5866 | 0 | 0.96156038 | 6 | only_H3vsUI | DownUp | notDM | Loss | 4 |
| **11_61730741_61733579_2237** | 11_61730741_61733579 | FEN1 | 2237 | 5.23126564843027 | -0.86163276 | 1 | only_H3vsUI | DownUp | notDM | Loss | 7 |
| **11_61735339_61739221_2237** | 11_61735339_61739221 | FEN1 | 2237 | 5.24283450300892 | -0.89016161 | 5 | only_H3vsUI | DownUp | notDM | Loss | 7 |
| **11_61739709_61743332_2237** | 11_61739709_61743332 | FEN1 | 2237 | 5.25355122258959 | -0.69524201 | 7 | only_H3vsUI | DownUp | notDM | Gain | 5 |

| | | | | | | | | | | | |
|---|---|---|---|---|---|---|---|---|---|---|---|
| **11_62160287_62161458_5866** | 11_62160287_62161458 | RAB3IL1 | 5866 | 5.67444776750189 | 0.99956783 | 6 | only_H3vsUI | DownUp | notDM | Gain | 1 |
| **11_62165703_62166698_5866** | 11_62165703_62166698 | RAB3IL1 | 5866 | 5.67939700427413 | 0.99954457 | 3 | only_H3vsUI | DownUp | notDM | Loss | 5 |
| **11_62306948_62313275_1937** | 11_62306948_62313275 | EEF1G | 1937 | 4.45154082857195 | -0.72208377 | 1 | notSpecific | Up | DM | Loss | 10 |
| **11_62306948_62313275_51035** | 11_62306948_62313275 | UBXN1 | 51035 | 5.12298545415801 | -0.77585596 | 1 | notSpecific | Up | DM | Loss | 10 |
| **11_62306948_62313275_64852** | 11_62306948_62313275 | TUT1 | 64852 | 4.66096029177608 | -0.54879552 | 1 | notSpecific | Down | DM | Loss | 10 |
| **11_62306948_62313275_79026** | 11_62306948_62313275 | AHNAK | 79026 | 2.99738638439731 | 0.46997788 | 1 | notSpecific | Down | DM | Loss | 10 |
| **11_63296290_63300663_5920** | 11_63296290_63300663 | PLAAT4 | 5920 | 3.5585885831082 | -0.1791468 | 7 | only_H3vsUI | UpDown | notDM | Gain | 1 |
| **11_63303592_63304964_5920** | 11_63303592_63304964 | PLAAT4 | 5920 | 0 | 0.14566873 | 6 | only_H3vsUI | UpDown | notDM | Loss | 3 |
| **11_63533954_63535752_65998** | 11_63533954_63535752 | C11orf95 | 65998 | 2.55870857053317 | -0.0003328 | 2 | only_H3vsUI | UpDown | notDM | Loss | 3 |
| **11_63606908_63610032_2011** | 11_63606908_63610032 | MARK2 | 2011 | 2.70757017609794 | -0.46619458 | 1 | notSpecific | Up | DM | Loss | 10 |
| **11_63606908_63610032_55611** | 11_63606908_63610032 | OTUB1 | 55611 | 5.15809097318556 | 0.66489538 | 1 | notSpecific | DownUp | DM | Loss | 10 |
| **11_63606908_63610032_79829** | 11_63606908_63610032 | NAA40 | 79829 | 4.98427071127802 | 0.67061591 | 1 | notSpecific | Down | DM | Loss | 10 |
| **11_63683802_63684422_56834** | 11_63683802_63684422 | GPR137 | 56834 | 5.5662111522465 | -0.79052354 | 1 | only_H3vsUI | DownUp | notDM | Loss | 7 |
| **11_63684808_63686700_56834** | 11_63684808_63686700 | GPR137 | 56834 | 5.56351668184391 | -0.81720292 | 4 | only_H3vsUI | DownUp | notDM | Loss | 7 |
| **11_64035276_64037399_56834** | 11_64035276_64037399 | GPR137 | 56834 | 2.13353890837022 | 0.00369621 | 1 | only_H3vsUI | DownUp | notDM | Loss | 8 |
| **11_64037695_64038494_56834** | 11_64037695_64038494 | GPR137 | 56834 | 2.21218760440396 | 0.00382345 | 1 | only_H3vsUI | DownUp | notDM | Loss | 8 |
| **11_64038703_64041239_56834** | 11_64038703_64041239 | GPR137 | 56834 | 3.06855689507236 | 0.00367701 | 1 | only_H3vsUI | DownUp | notDM | Loss | 8 |
| **11_64050639_64052823_56834** | 11_64050639_64052823 | GPR137 | 56834 | 0 | 0.49512596 | 1 | only_H3vsUI | DownUp | notDM | Loss | 8 |
| **11_64053402_64054520_56834** | 11_64053402_64054520 | GPR137 | 56834 | 2.82930377283102 | 0.54657076 | 1 | only_H3vsUI | DownUp | notDM | Loss | 6 |
| **11_64071216_64072490_56834** | 11_64071216_64072490 | GPR137 | 56834 | 4.2756798301027 | -0.36332048 | 1 | only_H3vsUI | DownUp | notDM | Loss | 7 |
| **11_64073363_64075419_56834** | 11_64073363_64075419 | GPR137 | 56834 | 4.31462551920127 | -0.4366211 | 1 | only_H3vsUI | DownUp | notDM | Loss | 7 |
| **11_64122353_64123928_56834** | 11_64122353_64123928 | GPR137 | 56834 | 4.84277144575971 | 0.54097442 | 5 | only_H3vsUI | DownUp | notDM | Gain | 1 |
| **11_64691048_64693131_84447** | 11_64691048_64693131 | SYVN1 | 84447 | 5.31988230446639 | 0.76704474 | 1 | only_H3vsUI | DownUp | notDM | Loss | 6 |
| **11_64765385_64767563_84447** | 11_64765385_64767563 | SYVN1 | 84447 | 5.12836372232786 | 0.79902331 | 3 | only_H3vsUI | DownUp | notDM | Gain | 5 |
| **11_64780323_64781639_84447** | 11_64780323_64781639 | SYVN1 | 84447 | 5.08031255604503 | 0.64824244 | 1 | only_H3vsUI | DownUp | notDM | Loss | 6 |
| **11_64849445_64851346_84447** | 11_64849445_64851346 | SYVN1 | 84447 | 4.70420201115623 | 0.71210509 | 1 | only_H3vsUI | DownUp | notDM | Loss | 6 |
| **11_64851949_64853372_84447** | 11_64851949_64853372 | SYVN1 | 84447 | 4.68645751046911 | 0.71210509 | 2 | only_H3vsUI | DownUp | notDM | Loss | 6 |
| **11_64863954_64865288_84447** | 11_64863954_64865288 | SYVN1 | 84447 | 4.56423984423673 | 0.63016194 | 1 | only_H3vsUI | DownUp | notDM | Loss | 6 |
| **11_64877992_64879103_84447** | 11_64877992_64879103 | SYVN1 | 84447 | 4.35985444675036 | 0.49136236 | 6 | only_H3vsUI | DownUp | notDM | Loss | 1 |
| **11_64879275_64880943_84447** | 11_64879275_64880943 | SYVN1 | 84447 | 4.32347898813016 | 0.38663709 | 2 | only_H3vsUI | DownUp | notDM | Loss | 7 |
| **11_64883468_64885057_84447** | 11_64883468_64885057 | SYVN1 | 84447 | 4.22775819611091 | 0.89434608 | 1 | only_H3vsUI | DownUp | notDM | Loss | 6 |
| **11_64887984_64889529_84447** | 11_64887984_64889529 | SYVN1 | 84447 | 4.09607536608511 | 0.82715249 | 1 | only_H3vsUI | DownUp | notDM | Loss | 6 |
| **11_64889894_64891285_84447** | 11_64889894_64891285 | SYVN1 | 84447 | 4.03019478535675 | 0.82715249 | 2 | only_H3vsUI | DownUp | notDM | Gain | 4 |
| **11_64899939_64901874_84447** | 11_64899939_64901874 | SYVN1 | 84447 | 1.99122607569249 | 0.81327362 | 2 | only_H3vsUI | DownUp | notDM | Loss | 6 |

| | | | | | | | | | | | |
|---|---|---|---|---|---|---|---|---|---|---|---|
| **11_64902634_64903975_84447** | 11_64902634_64903975 | SYVN1 | 84447 | 2.83569057149243 | 0.9044506 | 2 | only_H3vsUI | DownUp | notDM | Gain | 4 |
| **11_64917077_64919553_84447** | 11_64917077_64919553 | SYVN1 | 84447 | 4.17828612877757 | -0.14388638 | 7 | only_H3vsUI | DownUp | notDM | Gain | 5 |
| **11_65121374_65122071_84447** | 11_65121374_65122071 | SYVN1 | 84447 | 5.34128610702863 | 0.75858813 | 4 | only_H3vsUI | DownUp | notDM | Loss | 3 |
| **11_65122348_65123281_84447** | 11_65122348_65123281 | SYVN1 | 84447 | 5.34320961969215 | 0.77908651 | 2 | only_H3vsUI | DownUp | notDM | Loss | 3 |
| **11_65242620_65244987_84447** | 11_65242620_65244987 | SYVN1 | 84447 | 5.5323351653377 | 0.78782284 | 5 | only_H3vsUI | DownUp | notDM | Gain | 5 |
| **11_65263191_65264815_84447** | 11_65263191_65264815 | SYVN1 | 84447 | 5.55779823806405 | 0.87171987 | 7 | only_H3vsUI | DownUp | notDM | Gain | 5 |
| **11_65265230_65266445_84447** | 11_65265230_65266445 | SYVN1 | 84447 | 5.56024268437266 | 0.87171987 | 5 | only_H3vsUI | DownUp | notDM | Gain | 5 |
| **11_65269628_65275039_84447** | 11_65269628_65275039 | SYVN1 | 84447 | 5.56546882583143 | 0.82858988 | 1 | only_H3vsUI | DownUp | notDM | Loss | 6 |
| **11_65326021_65329791_80198** | 11_65326021_65329791 | MUS81 | 80198 | 5.47434157930158 | -0.78700745 | 7 | only_DIvsD7 | UpDown | notDM | Loss | 7 |
| **11_65584381_65585495_1521** | 11_65584381_65585495 | CTSW | 1521 | 4.79089008008012 | -0.59140939 | 4 | only_DIvsD7 | Up | notDM | Gain | 4 |
| **11_65584381_65585495_80198** | 11_65584381_65585495 | MUS81 | 80198 | 4.62718144714753 | -0.57152471 | 4 | only_DIvsD7 | UpDown | notDM | Gain | 2 |
| **11_65621831_65624746_1521** | 11_65621831_65624746 | CTSW | 1521 | 4.35380086513886 | 0.37884644 | 1 | only_DIvsD7 | Up | notDM | Loss | 6 |
| **11_65621831_65624746_80198** | 11_65621831_65624746 | MUS81 | 80198 | 0 | 0.00112893 | 1 | only_DIvsD7 | UpDown | notDM | Loss | 8 |
| **11_65625880_65628065_1521** | 11_65625880_65628065 | CTSW | 1521 | 4.28366299005883 | 0.40346549 | 5 | only_DIvsD7 | Up | notDM | Loss | 6 |
| **11_65625880_65628065_80198** | 11_65625880_65628065 | MUS81 | 80198 | 0 | 0.05153188 | 5 | only_DIvsD7 | UpDown | notDM | Loss | 8 |
| **11_65628293_65630073_1521** | 11_65628293_65630073 | CTSW | 1521 | 4.23573039737941 | 0.40346549 | 1 | only_DIvsD7 | Up | notDM | Loss | 6 |
| **11_65628293_65630073_80198** | 11_65628293_65630073 | MUS81 | 80198 | 2.4456042032736 | 0.05153188 | 1 | only_DIvsD7 | UpDown | notDM | Loss | 6 |
| **11_65640455_65641760_1521** | 11_65640455_65641760 | CTSW | 1521 | 3.74530905994083 | 0.00086199 | 2 | only_DIvsD7 | Up | notDM | Loss | 3 |
| **11_65640455_65641760_80198** | 11_65640455_65641760 | MUS81 | 80198 | 4.09964611712323 | -0.58780356 | 2 | only_DIvsD7 | UpDown | notDM | Loss | 7 |
| **11_65653783_65655916_1521** | 11_65653783_65655916 | CTSW | 1521 | 3.80996352171401 | 0.30231864 | 1 | only_DIvsD7 | Up | notDM | Loss | 6 |
| **11_65656110_65657989_1521** | 11_65656110_65657989 | CTSW | 1521 | 3.94398887507377 | 0.29193375 | 1 | only_DIvsD7 | Up | notDM | Loss | 6 |
| **11_65658114_65660769_1521** | 11_65658114_65660769 | CTSW | 1521 | 4.03318241372919 | 0.30945245 | 1 | only_DIvsD7 | Up | notDM | Loss | 6 |
| **11_65668091_65671695_1521** | 11_65668091_65671695 | CTSW | 1521 | 4.31731102020681 | -0.45369554 | 7 | only_DIvsD7 | Up | notDM | Loss | 5 |
| **11_65668091_65671695_80198** | 11_65668091_65671695 | MUS81 | 80198 | 4.60440967113295 | -0.51819349 | 7 | only_DIvsD7 | UpDown | notDM | Loss | 7 |
| **11_65678816_65681750_80198** | 11_65678816_65681750 | MUS81 | 80198 | 4.7058808101553 | 0.68496503 | 7 | only_DIvsD7 | UpDown | notDM | Loss | 5 |
| **11_65682103_65686239_80198** | 11_65682103_65686239 | MUS81 | 80198 | 4.73423159598098 | -0.5569912 | 1 | only_DIvsD7 | UpDown | notDM | Loss | 6 |
| **11_65687203_65688724_64837** | 11_65687203_65688724 | KLC2 | 64837 | 5.52692182345106 | 0.79613897 | 1 | only_H3vsUI | UpDown | notDM | Loss | 6 |
| **11_65873820_65875046_80198** | 11_65873820_65875046 | MUS81 | 80198 | 5.39059247958822 | -0.75433207 | 7 | only_DIvsD7 | UpDown | notDM | Gain | 2 |
| **11_65891095_65893176_64837** | 11_65891095_65893176 | KLC2 | 64837 | 5.12057064108368 | 0.70832865 | 7 | only_H3vsUI | UpDown | notDM | Gain | 5 |
| **11_66023630_66025087_64837** | 11_66023630_66025087 | KLC2 | 64837 | 2.01703333929878 | 0.32331159 | 1 | only_H3vsUI | UpDown | notDM | Gain | 4 |
| **11_66025318_66027170_1521** | 11_66025318_66027170 | CTSW | 1521 | 5.57753775449384 | 0.93717047 | 1 | only_DIvsD7 | Up | notDM | Loss | 3 |
| **11_66025318_66027170_64837** | 11_66025318_66027170 | KLC2 | 64837 | 2.74429298312268 | 0.48065337 | 1 | only_H3vsUI | UpDown | notDM | Loss | 6 |
| **11_66035196_66035908_64837** | 11_66035196_66035908 | KLC2 | 64837 | 3.99646789026172 | -0.81541576 | 1 | only_H3vsUI | UpDown | notDM | Loss | 7 |
| **11_66036357_66037962_64837** | 11_66036357_66037962 | KLC2 | 64837 | 4.04453976039241 | -0.81541576 | 1 | only_H3vsUI | UpDown | notDM | Loss | 7 |

| | | | | | | | | | | | |
|---|---|---|---|---|---|---|---|---|---|---|---|
| **11_66055170_66056350_64837** | 11_66055170_66056350 | KLC2 | 64837 | 4.47556950212748 | -0.70782231 | 1 | only_H3vsUI | UpDown | notDM | Loss | 7 |
| **11_66056874_66058483_64837** | 11_66056874_66058483 | KLC2 | 64837 | 4.49964585017235 | -0.70782231 | 1 | only_H3vsUI | UpDown | notDM | Loss | 7 |
| **11_66059372_66060235_64837** | 11_66059372_66060235 | KLC2 | 64837 | 4.53269069478065 | 0.37168503 | 2 | only_H3vsUI | UpDown | notDM | Loss | 3 |
| **11_66102460_66105259_64837** | 11_66102460_66105259 | KLC2 | 64837 | 4.88811206910148 | 0.79203999 | 1 | only_H3vsUI | UpDown | notDM | Loss | 6 |
| **11_66153339_66154568_64837** | 11_66153339_66154568 | KLC2 | 64837 | 5.10777621892082 | -0.69772602 | 2 | only_H3vsUI | UpDown | notDM | Gain | 2 |
| **11_67805404_67809331_10312** | 11_67805404_67809331 | TCIRG1 | 10312 | 0 | -0.04038527 | 2 | notSpecific | DownUp | DM | Loss | 10 |
| **11_67805404_67809331_221** | 11_67805404_67809331 | ALDH3B1 | 221 | 4.44116077403512 | 0.50174405 | 2 | notSpecific | Up | DM | Loss | 10 |
| **11_67805404_67809331_4728** | 11_67805404_67809331 | NDUFS8 | 4728 | 3.84341966520492 | 0.59109219 | 2 | notSpecific | DownUp | DM | Loss | 10 |
| **11_67805404_67809331_81622** | 11_67805404_67809331 | UNC93B1 | 81622 | 4.52908370450271 | 0.6643799 | 2 | notSpecific | Up | DM | Loss | 10 |
| **11_69086525_69089531_26579** | 11_69086525_69089531 | MYEOV | 26579 | 3.34791518650169 | 0 | 7 | notSpecific | Down | DM | Loss | 10 |
| **11_71496562_71497839_55199** | 11_71496562_71497839 | FAM86C1P | 55199 | 2.8561244442423 | -0.10296545 | 1 | only_H3vsUI | Down | notDM | Gain | 4 |
| **11_71498741_71500300_55199** | 11_71498741_71500300 | FAM86C1P | 55199 | 2.25285303097989 | 0.02626121 | 1 | only_H3vsUI | Down | notDM | Loss | 8 |
| **11_71523529_71524909_285407** | 11_71523529_71524909 | ALG1L9P | 285407 | 0 | 0.01035271 | 1 | only_H3vsUI | DownUp | notDM | Loss | 3 |
| **11_71523529_71524909_55199** | 11_71523529_71524909 | FAM86C1P | 55199 | 4.39736636126529 | -0.35806637 | 1 | only_H3vsUI | Down | notDM | Loss | 7 |
| **11_71748753_71754523_220074** | 11_71748753_71754523 | LRTOMT | 220074 | 4.5721277878771 | -0.54903796 | 5 | only_H3vsUI | UpDown | notDM | Loss | 6 |
| **11_71780279_71783956_220074** | 11_71780279_71783956 | LRTOMT | 220074 | 3.89779198193922 | 0.16842015 | 8 | only_H3vsUI | UpDown | notDM | Loss | 5 |
| **11_71788891_71791298_220074** | 11_71788891_71791298 | LRTOMT | 220074 | 2.78390357927274 | 0.15891673 | 1 | only_H3vsUI | UpDown | notDM | Loss | 6 |
| **11_71791963_71793709_220074** | 11_71791963_71793709 | LRTOMT | 220074 | 1.77815125038364 | 0.15891673 | 1 | only_H3vsUI | UpDown | notDM | Gain | 4 |
| **11_71811681_71814206_220074** | 11_71811681_71814206 | LRTOMT | 220074 | 4.29618237255544 | 0.23661462 | 1 | only_H3vsUI | UpDown | notDM | Loss | 6 |
| **11_71822201_71823355_220074** | 11_71822201_71823355 | LRTOMT | 220074 | 3.39654803798713 | 0 | 1 | only_H3vsUI | UpDown | notDM | Gain | 4 |
| **11_71823871_71825156_220074** | 11_71823871_71825156 | LRTOMT | 220074 | 3.61930207587561 | 0 | 3 | only_H3vsUI | UpDown | notDM | Gain | 1 |
| **11_71933226_71934297_2350** | 11_71933226_71934297 | FOLR2 | 2350 | 3.56181666431896 | 0 | 2 | only_H3vsUI | UpDown | notDM | Gain | 4 |
| **11_71933226_71934297_81570** | 11_71933226_71934297 | CLPB | 81570 | 5.32486249925932 | 0.75759054 | 2 | only_DIvsD7 | UpDown | notDM | Gain | 1 |
| **11_72144655_72145176_81570** | 11_72144655_72145176 | CLPB | 81570 | 1 | 0.09964966 | 2 | only_DIvsD7 | UpDown | notDM | Loss | 3 |
| **11_72300590_72301747_81570** | 11_72300590_72301747 | CLPB | 81570 | 5.19037092299724 | 0.77122795 | 1 | only_DIvsD7 | UpDown | notDM | Loss | 3 |
| **11_72352787_72353352_81570** | 11_72352787_72353352 | CLPB | 81570 | 5.31641280663544 | 0.76714773 | 4 | only_DIvsD7 | UpDown | notDM | Loss | 3 |
| **11_7266907_7268517_8495** | 11_7266907_7268517 | PPFIBP2 | 8495 | 5.54394518367714 | 0.99476891 | 6 | only_H3vsUI | Down | notDM | Loss | 1 |
| **11_72999482_73001211_9828** | 11_72999482_73001211 | ARHGEF17 | 9828 | 4.25825405350715 | 0.26703643 | 7 | only_DIvsD7 | UpDown | notDM | Loss | 5 |
| **11_73017382_73018524_9828** | 11_73017382_73018524 | ARHGEF17 | 9828 | 2.90902085421116 | 0.66618468 | 1 | only_DIvsD7 | UpDown | notDM | Loss | 6 |
| **11_73018878_73021441_9828** | 11_73018878_73021441 | ARHGEF17 | 9828 | 0 | 0.58857178 | 1 | only_DIvsD7 | UpDown | notDM | Loss | 6 |
| **11_73044598_73047233_9828** | 11_73044598_73047233 | ARHGEF17 | 9828 | 4.40253649180391 | 0.75674003 | 3 | only_DIvsD7 | UpDown | notDM | Loss | 5 |
| **11_74383038_74385471_9789** | 11_74383038_74385471 | SPCS2 | 9789 | 5.43905149584081 | 0.82378982 | 7 | only_DIvsD7 | UpDown | notDM | Gain | 5 |
| **11_74658882_74660047_9789** | 11_74658882_74660047 | SPCS2 | 9789 | 2.41497334797082 | -0.09510164 | 1 | only_DIvsD7 | UpDown | notDM | Gain | 4 |
| **11_74660553_74662138_9789** | 11_74660553_74662138 | SPCS2 | 9789 | 2.36361197989214 | -0.34941431 | 1 | only_DIvsD7 | UpDown | notDM | Loss | 8 |

| | | | | | | | | | | | |
|---|---|---|---|---|---|---|---|---|---|---|---|
| **11_75131419_75134778_283212** | 11_75131419_75134778 | KLHL35 | 283212 | 3.81016528454315 | -0.0948823 | 7 | only_H3vsUI | Up | notDM | Loss | 5 |
| **11_75131419_75134778_84649** | 11_75131419_75134778 | DGAT2 | 84649 | 5.53781909507327 | -0.78270715 | 7 | only_DIvsD7 | UpDown | notDM | Loss | 7 |
| **11_7534033_7534868_8495** | 11_7534033_7534868 | PPFIBP2 | 8495 | 2.12710479836481 | 0.89654386 | 2 | only_H3vsUI | Down | notDM | Loss | 3 |
| **11_7535170_7536063_8495** | 11_7535170_7536063 | PPFIBP2 | 8495 | 0 | 0.91484654 | 2 | only_H3vsUI | Down | notDM | Loss | 3 |
| **11_75478691_75480491_84649** | 11_75478691_75480491 | DGAT2 | 84649 | 0 | 0.3473706 | 2 | only_DIvsD7 | UpDown | notDM | Gain | 4 |
| **11_75917168_75921909_7481** | 11_75917168_75921909 | WNT11 | 7481 | 0 | 0.01706689 | 2 | only_H3vsUI | DownUp | notDM | Loss | 3 |
| **11_7592462_7599625_8495** | 11_7592462_7599625 | PPFIBP2 | 8495 | 0 | 0.60606532 | 7 | only_H3vsUI | Down | notDM | Gain | 4 |
| **11_7621140_7622690_8495** | 11_7621140_7622690 | PPFIBP2 | 8495 | 3.43536650661266 | 0.33775681 | 7 | only_H3vsUI | Down | notDM | Gain | 1 |
| **11_7624783_7631163_8495** | 11_7624783_7631163 | PPFIBP2 | 8495 | 0 | 0.26666039 | 5 | only_H3vsUI | Down | notDM | Loss | 8 |
| **11_7632003_7635918_8495** | 11_7632003_7635918 | PPFIBP2 | 8495 | 4.56666135837174 | 0.81076019 | 6 | only_H3vsUI | Down | notDM | Loss | 5 |
| **11_77529311_77531406_51773** | 11_77529311_77531406 | RSF1 | 51773 | 2.53147891704225 | 0.13514569 | 1 | only_DIvsD7 | UpDown | notDM | Loss | 6 |
| **11_77532218_77533444_51773** | 11_77532218_77533444 | RSF1 | 51773 | 2.19589965240923 | -0.09822143 | 1 | only_DIvsD7 | UpDown | notDM | Loss | 8 |
| **11_77533577_77534892_51773** | 11_77533577_77534892 | RSF1 | 51773 | 3.263399331334 | 0.13611758 | 4 | only_DIvsD7 | UpDown | notDM | Gain | 4 |
| **11_77704115_77705632_51773** | 11_77704115_77705632 | RSF1 | 51773 | 5.23566477377609 | 0.77276605 | 1 | only_DIvsD7 | UpDown | notDM | Gain | 5 |
| **11_77748053_77748838_51773** | 11_77748053_77748838 | RSF1 | 51773 | 5.33507659731441 | 0.78699862 | 6 | only_DIvsD7 | UpDown | notDM | Gain | 1 |
| **11_78135104_78136203_101928865** | 11_78135104_78136203 | LINC02728 | 101928865 | 3.67024585307412 | 0 | 3 | only_H3vsUI | DownUp | notDM | Gain | 1 |
| **11_79129756_79130937_26011** | 11_79129756_79130937 | TENM4 | 26011 | 4.32337587193386 | 0.24400744 | 6 | only_H3vsUI | UpDown | notDM | Loss | 3 |
| **11_79148083_79149484_26011** | 11_79148083_79149484 | TENM4 | 26011 | 3.39950066131461 | 0.24386607 | 4 | only_H3vsUI | UpDown | notDM | Loss | 3 |
| **11_79149606_79152496_26011** | 11_79149606_79152496 | TENM4 | 26011 | 0 | 0.24401236 | 4 | only_H3vsUI | UpDown | notDM | Loss | 3 |
| **11_87835273_87839108_23682** | 11_87835273_87839108 | RAB38 | 23682 | 4.84194730992282 | 0.89400182 | 3 | notSpecific | UpDown | DM | Gain | 9 |
| **11_8956130_8957210_56673** | 11_8956130_8957210 | C11orf16 | 56673 | 3.20303288701471 | 0 | 6 | only_DIvsD7 | UpDown | notDM | Loss | 3 |
| **11_9925788_9929897_23075** | 11_9925788_9929897 | SWAP70 | 23075 | 5.38051152489229 | 0.83038958 | 7 | notSpecific | Down | DM | Gain | 9 |
| **12_104322319_104323509_728568** | 12_104322319_104323509 | C12orf73 | 728568 | 4.43909574130175 | 0.48542495 | 1 | only_DIvsD7 | UpDown | notDM | Loss | 6 |
| **12_104324362_104327833_728568** | 12_104324362_104327833 | C12orf73 | 728568 | 4.36475730657121 | 0.48542495 | 1 | only_DIvsD7 | UpDown | notDM | Loss | 6 |
| **12_104348944_104350745_728568** | 12_104348944_104350745 | C12orf73 | 728568 | 2.39619934709574 | 0.65707454 | 1 | only_DIvsD7 | UpDown | notDM | Loss | 6 |
| **12_104351151_104351653_728568** | 12_104351151_104351653 | C12orf73 | 728568 | 2.20411998265592 | 0.65707454 | 1 | only_DIvsD7 | UpDown | notDM | Gain | 1 |
| **12_104351828_104353416_728568** | 12_104351828_104353416 | C12orf73 | 728568 | 2.92272545799326 | 0.55633465 | 6 | only_DIvsD7 | UpDown | notDM | Loss | 5 |
| **12_104358061_104359462_728568** | 12_104358061_104359462 | C12orf73 | 728568 | 3.8494194137969 | 0.63745081 | 1 | only_DIvsD7 | UpDown | notDM | Loss | 6 |
| **12_104359994_104361993_728568** | 12_104359994_104361993 | C12orf73 | 728568 | 3.95438725014451 | 0.63745081 | 1 | only_DIvsD7 | UpDown | notDM | Loss | 6 |
| **12_110839565_110841149_10094** | 12_110839565_110841149 | ARPC3 | 10094 | 4.67156158910706 | 0.67340583 | 1 | only_H3vsUI | DownUp | notDM | Loss | 6 |
| **12_110841540_110842661_10094** | 12_110841540_110842661 | ARPC3 | 10094 | 4.65734273681463 | 0.67340583 | 5 | only_H3vsUI | DownUp | notDM | Gain | 1 |
| **12_110886681_110887795_10094** | 12_110886681_110887795 | ARPC3 | 10094 | 2.47129171105894 | 0.60193131 | 1 | only_H3vsUI | DownUp | notDM | Gain | 4 |
| **12_110888272_110888627_10094** | 12_110888272_110888627 | ARPC3 | 10094 | 2.10380372095596 | 0.58011448 | 4 | only_H3vsUI | DownUp | notDM | Gain | 2 |
| **12_110904286_110906006_10094** | 12_110904286_110906006 | ARPC3 | 10094 | 4.20946139462999 | 0.55927932 | 1 | only_H3vsUI | DownUp | notDM | Loss | 6 |

| | | | | | | | | | | | |
|---|---|---|---|---|---|---|---|---|---|---|---|
| **12_111051877_111053375_10094** | 12_111051877_111053375 | ARPC3 | 10094 | 5.21428473136997 | 0.68761193 | 2 | only_H3vsUI | DownUp | notDM | Loss | 6 |
| **12_112121606_112123487_80724** | 12_112121606_112123487 | ACAD10 | 80724 | 2.56937390961505 | -0.13724808 | 2 | only_DIvsD7 | Down | notDM | Loss | 8 |
| **12_112123910_112125671_80724** | 12_112123910_112125671 | ACAD10 | 80724 | 1 | -0.13480081 | 1 | only_DIvsD7 | Down | notDM | Gain | 4 |
| **12_117316265_117318896_26259** | 12_117316265_117318896 | FBXW8 | 26259 | 4.47530791395619 | -0.44991825 | 4 | notSpecific | Up | DM | Loss | 10 |
| **12_117316265_117318896_8739** | 12_117316265_117318896 | HRK | 8739 | 2.52762990087134 | 0.03457821 | 4 | notSpecific | Up | DM | Loss | 10 |
| **12_117588613_117590181_23014** | 12_117588613_117590181 | FBXO21 | 23014 | 4.5808793780608 | -0.62497206 | 7 | notSpecific | Down | DM | Gain | 9 |
| **12_118488687_118492451_51347** | 12_118488687_118492451 | TAOK3 | 51347 | 5.48355306551291 | -0.86525909 | 7 | only_DIvsD7 | UpDown | notDM | Gain | 5 |
| **12_118493124_118495821_51347** | 12_118493124_118495821 | TAOK3 | 51347 | 5.49814231826636 | -0.78023624 | 7 | only_DIvsD7 | UpDown | notDM | Loss | 7 |
| **12_118539289_118541732_51347** | 12_118539289_118541732 | TAOK3 | 51347 | 5.40694873595035 | -0.92116541 | 1 | only_DIvsD7 | UpDown | notDM | Loss | 7 |
| **12_118541969_118542976_51347** | 12_118541969_118542976 | TAOK3 | 51347 | 5.4048268772829 | -0.9230217 | 1 | only_DIvsD7 | UpDown | notDM | Loss | 7 |
| **12_118568605_118570619_51347** | 12_118568605_118570619 | TAOK3 | 51347 | 5.38045003801369 | -0.86677861 | 6 | only_DIvsD7 | UpDown | notDM | Loss | 7 |
| **12_118572347_118573744_51347** | 12_118572347_118573744 | TAOK3 | 51347 | 5.37476117308385 | -0.79407579 | 1 | only_DIvsD7 | UpDown | notDM | Loss | 7 |
| **12_118573980_118575801_51347** | 12_118573980_118575801 | TAOK3 | 51347 | 5.37097544935897 | -0.78076224 | 1 | only_DIvsD7 | UpDown | notDM | Loss | 7 |
| **12_118744566_118746246_51347** | 12_118744566_118746246 | TAOK3 | 51347 | 4.80959337956935 | 0.66813162 | 4 | only_DIvsD7 | UpDown | notDM | Gain | 1 |
| **12_118772846_118777373_51347** | 12_118772846_118777373 | TAOK3 | 51347 | 4.29088010526716 | 0.95065115 | 7 | only_DIvsD7 | UpDown | notDM | Gain | 5 |
| **12_118793152_118797637_51347** | 12_118793152_118797637 | TAOK3 | 51347 | 0 | 0.76639505 | 3 | only_DIvsD7 | UpDown | notDM | Gain | 4 |
| **12_118798068_118799595_51347** | 12_118798068_118799595 | TAOK3 | 51347 | 3.06445798922692 | 0.79727162 | 6 | only_DIvsD7 | UpDown | notDM | Gain | 1 |
| **12_118808059_118810441_51347** | 12_118808059_118810441 | TAOK3 | 51347 | 2.40483371661994 | 0.05858095 | 1 | only_DIvsD7 | UpDown | notDM | Loss | 6 |
| **12_118810986_118812491_51347** | 12_118810986_118812491 | TAOK3 | 51347 | 2.46239799789896 | 0.05786225 | 1 | only_DIvsD7 | UpDown | DM | Gain | 1 |
| **12_118810986_118812491_64426** | 12_118810986_118812491 | SUDS3 | 64426 | 3.27137687189407 | -0.91330736 | 1 | notSpecific | Down | DM | Gain | 9 |
| **12_118813329_118814068_51347** | 12_118813329_118814068 | TAOK3 | 51347 | 3.42045085910607 | -0.53816517 | 2 | only_DIvsD7 | UpDown | notDM | Gain | 2 |
| **12_118814789_118816284_51347** | 12_118814789_118816284 | TAOK3 | 51347 | 3.61246596395314 | 0.10480128 | 2 | only_DIvsD7 | UpDown | notDM | Gain | 4 |
| **12_119615708_119619326_26353** | 12_119615708_119619326 | HSPB8 | 26353 | 0 | 0.10680365 | 2 | only_DIvsD7 | Up | DM | Loss | 10 |
| **12_120876100_120878203_84274** | 12_120876100_120878203 | COQ5 | 84274 | 4.94817325515851 | -0.59719803 | 2 | only_H3vsUI | DownUp | notDM | Loss | 7 |
| **12_120907684_120909391_84274** | 12_120907684_120909391 | COQ5 | 84274 | 4.76014341989789 | 0.4992345 | 2 | only_H3vsUI | DownUp | notDM | Loss | 7 |
| **12_120911641_120913864_84274** | 12_120911641_120913864 | COQ5 | 84274 | 4.72501272534116 | -0.75643884 | 6 | only_H3vsUI | DownUp | notDM | Loss | 7 |
| **12_120965775_120966809_84274** | 12_120965775_120966809 | COQ5 | 84274 | 2.16136800223497 | 0.21112068 | 2 | only_H3vsUI | DownUp | notDM | Loss | 8 |
| **12_120967007_120968416_84274** | 12_120967007_120968416 | COQ5 | 84274 | 1.65321251377534 | 0.21004267 | 1 | only_H3vsUI | DownUp | notDM | Loss | 6 |
| **12_120972616_120978304_84274** | 12_120972616_120978304 | COQ5 | 84274 | 3.75319991419942 | 0.01571625 | 1 | only_H3vsUI | DownUp | notDM | Loss | 8 |
| **12_120987020_120989058_84274** | 12_120987020_120989058 | COQ5 | 84274 | 4.30252573296074 | 0.2990307 | 8 | only_H3vsUI | DownUp | notDM | Gain | 1 |
| **12_121266258_121268017_84274** | 12_121266258_121268017 | COQ5 | 84274 | 5.47610091266533 | -0.77952312 | 6 | only_H3vsUI | DownUp | notDM | Gain | 2 |
| **12_121266258_121268017_8638** | 12_121266258_121268017 | OASL | 8638 | 5.31959319563817 | 0.76819687 | 6 | only_H3vsUI | UpDown | notDM | Gain | 1 |
| **12_121287398_121288604_8638** | 12_121287398_121288604 | OASL | 8638 | 5.27449729789461 | 0.72498653 | 6 | only_H3vsUI | UpDown | notDM | Gain | 1 |
| **12_121452632_121454073_8638** | 12_121452632_121454073 | OASL | 8638 | 4.3556047509588 | 0.30272764 | 2 | only_H3vsUI | UpDown | notDM | Gain | 4 |

| | | | | | | | | | | | |
|---|---|---|---|---|---|---|---|---|---|---|---|
| **12_121476014_121477812_8638** | 12_121476014_121477812 | OASL | 8638 | 0 | 0.96258852 | 1 | only_H3vsUI | UpDown | notDM | Loss | 3 |
| **12_121497775_121499019_8638** | 12_121497775_121499019 | OASL | 8638 | 4.32277731470321 | 0.5864279 | 4 | only_H3vsUI | UpDown | notDM | Loss | 3 |
| **12_122109677_122111236_283385** | 12_122109677_122111236 | MORN3 | 283385 | 3.32613095671079 | -0.046015 | 2 | only_H3vsUI | DownUp | notDM | Gain | 4 |
| **12_122277649_122279000_65082** | 12_122277649_122279000 | VPS33A | 65082 | 5.67400548195852 | 0.92317839 | 2 | only_DIvsD7 | Down | notDM | Loss | 7 |
| **12_122433755_122435267_65082** | 12_122433755_122435267 | VPS33A | 65082 | 5.4994148761036 | -0.89530162 | 6 | only_DIvsD7 | Down | notDM | Gain | 2 |
| **12_122458640_122459602_605** | 12_122458640_122459602 | BCL7A | 605 | 2.28103336724773 | 0.07117362 | 2 | only_DIvsD7 | UpDown | notDM | Loss | 3 |
| **12_122460686_122462844_605** | 12_122460686_122462844 | BCL7A | 605 | 2.95230800966213 | -0.11217558 | 2 | only_DIvsD7 | UpDown | notDM | Gain | 4 |
| **12_122478018_122480348_65082** | 12_122478018_122480348 | VPS33A | 65082 | 5.43252194557613 | -0.93476268 | 7 | only_DIvsD7 | Down | notDM | Gain | 2 |
| **12_122500017_122503153_65082** | 12_122500017_122503153 | VPS33A | 65082 | 5.39412058022113 | 0.81421555 | 7 | only_DIvsD7 | Down | notDM | Loss | 5 |
| **12_122516999_122520475_65082** | 12_122516999_122520475 | VPS33A | 65082 | 5.362650203681 | -0.72876813 | 1 | only_DIvsD7 | Down | notDM | Loss | 7 |
| **12_122610684_122611934_56616** | 12_122610684_122611934 | DIABLO | 56616 | 4.99416752190719 | 0.64450243 | 8 | only_DIvsD7 | DownUp | notDM | Loss | 5 |
| **12_122687434_122688114_56616** | 12_122687434_122688114 | DIABLO | 56616 | 4.35210530333973 | 0.71825879 | 1 | only_DIvsD7 | DownUp | notDM | Gain | 1 |
| **12_122708035_122710335_56616** | 12_122708035_122710335 | DIABLO | 56616 | 2.34635297445064 | 0.28821102 | 1 | only_DIvsD7 | DownUp | notDM | Loss | 8 |
| **12_122708035_122710335_65082** | 12_122708035_122710335 | VPS33A | 65082 | 4.60883613318473 | 0.45177459 | 1 | only_DIvsD7 | Down | notDM | Loss | 7 |
| **12_122711055_122711843_56616** | 12_122711055_122711843 | DIABLO | 56616 | 2.65127801399814 | 0.31127198 | 2 | only_DIvsD7 | DownUp | notDM | Loss | 3 |
| **12_122711055_122711843_65082** | 12_122711055_122711843 | VPS33A | 65082 | 4.59357402399756 | 0.41465372 | 2 | only_DIvsD7 | Down | notDM | Loss | 2 |
| **12_122749449_122750842_65082** | 12_122749449_122750842 | VPS33A | 65082 | 2.35602585719312 | -0.57052461 | 1 | only_DIvsD7 | Down | notDM | Loss | 8 |
| **12_122751156_122752525_65082** | 12_122751156_122752525 | VPS33A | 65082 | 1.95424250943932 | -0.57052461 | 1 | only_DIvsD7 | Down | notDM | Gain | 4 |
| **12_122897479_122898800_55596** | 12_122897479_122898800 | ZCCHC8 | 55596 | 4.93811426115711 | -0.58921829 | 8 | only_H3vsUI | DownUp | notDM | Gain | 2 |
| **12_122948180_122949879_55596** | 12_122948180_122949879 | ZCCHC8 | 55596 | 4.55193769536484 | -0.49762631 | 3 | only_H3vsUI | DownUp | notDM | Loss | 7 |
| **12_122948180_122949879_56616** | 12_122948180_122949879 | DIABLO | 56616 | 5.37589395751608 | -0.86317725 | 3 | only_DIvsD7 | DownUp | notDM | Loss | 7 |
| **12_122955639_122956535_55596** | 12_122955639_122956535 | ZCCHC8 | 55596 | 4.46215832102648 | -0.78481143 | 3 | only_H3vsUI | DownUp | notDM | Gain | 2 |
| **12_122982203_122985083_55596** | 12_122982203_122985083 | ZCCHC8 | 55596 | 2.63948648926859 | 0.28149926 | 1 | only_H3vsUI | DownUp | notDM | Loss | 6 |
| **12_123006442_123010327_55596** | 12_123006442_123010327 | ZCCHC8 | 55596 | 4.32068622098333 | 0.25243609 | 2 | only_H3vsUI | DownUp | notDM | Gain | 4 |
| **12_123011865_123013543_55596** | 12_123011865_123013543 | ZCCHC8 | 55596 | 4.42076413747179 | 0.45721282 | 1 | only_H3vsUI | DownUp | notDM | Loss | 6 |
| **12_123162969_123164846_55596** | 12_123162969_123164846 | ZCCHC8 | 55596 | 5.24908334587822 | -0.71460513 | 7 | only_H3vsUI | DownUp | notDM | Loss | 7 |
| **12_123162969_123164846_65082** | 12_123162969_123164846 | VPS33A | 65082 | 5.61490564885382 | -0.839362 | 7 | only_DIvsD7 | Down | notDM | Loss | 2 |
| **12_123370603_123374151_55596** | 12_123370603_123374151 | ZCCHC8 | 55596 | 5.58555885769424 | -0.88535943 | 6 | only_H3vsUI | DownUp | notDM | Loss | 7 |
| **12_129306451_129308066_144423** | 12_129306451_129308066 | GLT1D1 | 144423 | 4.47674470297268 | 0.71369566 | 1 | only_H3vsUI | UpDown | notDM | Loss | 3 |
| **12_129308681_129309755_144423** | 12_129308681_129309755 | GLT1D1 | 144423 | 4.45155618307643 | 0.70172896 | 1 | only_H3vsUI | UpDown | notDM | Loss | 3 |
| **12_12955202_12958805_2012** | 12_12955202_12958805 | EMP1 | 2012 | 5.59208011334447 | 0.93706331 | 7 | only_H3vsUI | Up | notDM | Gain | 5 |
| **12_12971639_12973833_2012** | 12_12971639_12973833 | EMP1 | 2012 | 5.5750549951812 | 0.86433246 | 7 | only_H3vsUI | Up | notDM | Gain | 5 |
| **12_12989230_12993836_2012** | 12_12989230_12993836 | EMP1 | 2012 | 5.55130602256871 | 0.81134497 | 7 | only_H3vsUI | Up | notDM | Loss | 5 |
| **12_13023561_13027992_2012** | 12_13023561_13027992 | EMP1 | 2012 | 5.50748615944177 | 0.90373684 | 7 | only_H3vsUI | Up | notDM | Gain | 5 |

| | | | | | | | | | | | |
|---|---|---|---|---|---|---|---|---|---|---|---|
| **12_13033767_13036366_2012** | 12_13033767_13036366 | EMP1 | 2012 | 5.49603247097214 | 0.92717761 | 7 | only_H3vsUI | Up | notDM | Loss | 5 |
| **12_13041840_13044359_2012** | 12_13041840_13044359 | EMP1 | 2012 | 5.48481072470109 | 0.98258872 | 1 | only_H3vsUI | Up | notDM | Loss | 6 |
| **12_13044465_13057486_2012** | 12_13044465_13057486 | EMP1 | 2012 | 5.46572777033372 | 0.99818253 | 1 | only_H3vsUI | Up | notDM | Loss | 6 |
| **12_13153389_13154258_2012** | 12_13153389_13154258 | EMP1 | 2012 | 5.29105789443428 | 0.93101026 | 1 | only_H3vsUI | Up | notDM | Loss | 3 |
| **12_13249854_13251078_2012** | 12_13249854_13251078 | EMP1 | 2012 | 4.99405306358768 | 0.95130685 | 3 | only_H3vsUI | Up | notDM | Gain | 1 |
| **12_13251358_13254092_2012** | 12_13251358_13254092 | EMP1 | 2012 | 4.98057598977957 | 0.84944514 | 7 | only_H3vsUI | Up | notDM | Loss | 5 |
| **12_13254272_13256385_2012** | 12_13254272_13256385 | EMP1 | 2012 | 4.97003522556807 | 0.82444543 | 5 | only_H3vsUI | Up | notDM | Gain | 5 |
| **12_132626922_132628438_317781** | 12_132626922_132628438 | DDX51 | 317781 | 2.64640372622307 | 0.07608901 | 1 | only_DIvsD7 | DownUp | notDM | Loss | 7 |
| **12_132629229_132630688_317781** | 12_132629229_132630688 | DDX51 | 317781 | 2.54530711646582 | -0.07302461 | 1 | only_DIvsD7 | DownUp | notDM | Loss | 7 |
| **12_133003085_133003733_5827** | 12_133003085_133003733 | PXMP2 | 5827 | 5.41580105895506 | -0.79647932 | 6 | notSpecific | UpDown | DM | Loss | 10 |
| **12_13339378_13341327_2012** | 12_13339378_13341327 | EMP1 | 2012 | 3.92085349612126 | 0.61720824 | 6 | only_H3vsUI | Up | notDM | Loss | 5 |
| **12_13347539_13351768_2012** | 12_13347539_13351768 | EMP1 | 2012 | 0 | 0.96968339 | 1 | only_H3vsUI | Up | notDM | Loss | 8 |
| **12_13364312_13365146_2012** | 12_13364312_13365146 | EMP1 | 2012 | 4.16426360802266 | 0.98105559 | 6 | only_H3vsUI | Up | notDM | Loss | 5 |
| **12_13374789_13377677_2012** | 12_13374789_13377677 | EMP1 | 2012 | 4.39922362152727 | 0.67134562 | 6 | only_H3vsUI | Up | notDM | Loss | 5 |
| **12_13394902_13396164_2012** | 12_13394902_13396164 | EMP1 | 2012 | 4.65501350914057 | 0.72243943 | 6 | only_H3vsUI | Up | notDM | Gain | 1 |
| **12_1798843_1799989_79602** | 12_1798843_1799989 | ADIPOR2 | 79602 | 2.31175386105575 | 0.40715526 | 5 | only_DIvsD7 | Up | notDM | Loss | 6 |
| **12_1800620_1802263_79602** | 12_1800620_1802263 | ADIPOR2 | 79602 | 2.63245729218472 | 0.40578779 | 1 | only_DIvsD7 | Up | notDM | Gain | 4 |
| **12_27088618_27090784_26127** | 12_27088618_27090784 | FGFR1OP2 | 26127 | 2.72672720902657 | -0.31445198 | 2 | only_H3vsUI | UpDown | notDM | Gain | 4 |
| **12_27091709_27094362_26127** | 12_27091709_27094362 | FGFR1OP2 | 26127 | 2.59659709562646 | -0.26015492 | 1 | only_H3vsUI | UpDown | notDM | Loss | 6 |
| **12_2799841_2801560_2288** | 12_2799841_2801560 | FKBP4 | 2288 | 5.01097801217474 | 0.79997618 | 4 | notSpecific | Up | DM | Loss | 10 |
| **12_2799841_2801560_2305** | 12_2799841_2801560 | FOXM1 | 2305 | 5.26634225596734 | 0.74878816 | 4 | notSpecific | Down | DM | Loss | 10 |
| **12_31467694_31476988_79857** | 12_31467694_31476988 | FLJ13224 | 79857 | 2.41995574848976 | -0.0005872 | 2 | only_H3vsUI | UpDown | notDM | Loss | 6 |
| **12_31479321_31480338_79857** | 12_31479321_31480338 | FLJ13224 | 79857 | 3.31659930209386 | -0.00108489 | 2 | only_H3vsUI | UpDown | notDM | Loss | 3 |
| **12_45444384_45445835_51054** | 12_45444384_45445835 | PLEKHA8P1 | 51054 | 5.21472466552687 | -0.68035661 | 4 | only_DIvsD7 | UpDown | notDM | Loss | 7 |
| **12_45584194_45587363_51054** | 12_45584194_45587363 | PLEKHA8P1 | 51054 | 4.35077118305317 | -0.66545869 | 6 | only_DIvsD7 | UpDown | notDM | Loss | 7 |
| **12_45607749_45609739_51054** | 12_45607749_45609739 | PLEKHA8P1 | 51054 | 1.70757017609794 | -0.75302088 | 1 | only_DIvsD7 | UpDown | notDM | Loss | 8 |
| **12_45610465_45613774_51054** | 12_45610465_45613774 | PLEKHA8P1 | 51054 | 2.83122969386706 | -0.70185058 | 1 | only_DIvsD7 | UpDown | notDM | Loss | 8 |
| **12_45624557_45630375_51054** | 12_45624557_45630375 | PLEKHA8P1 | 51054 | 4.16938049531195 | -0.64902843 | 7 | only_DIvsD7 | UpDown | notDM | Gain | 5 |
| **12_47474080_47482926_347902** | 12_47474080_47482926 | AMIGO2 | 347902 | 2.72015930340596 | 0.65521853 | 5 | notSpecific | Down | DM | Gain | 9 |
| **12_48576060_48577273_387856** | 12_48576060_48577273 | CCDC184 | 387856 | 1.9731278535997 | 0.08667136 | 1 | only_DIvsD7 | UpDown | notDM | Loss | 6 |
| **12_48579002_48580338_387856** | 12_48579002_48580338 | CCDC184 | 387856 | 3.2143138974244 | -0.05408374 | 6 | only_DIvsD7 | UpDown | notDM | Loss | 3 |
| **12_50419461_50420080_91012** | 12_50419461_50420080 | CERS5 | 91012 | 5.14925607231515 | -0.80338169 | 4 | only_H3vsUI | Down | notDM | Gain | 2 |
| **12_50469169_50469856_91012** | 12_50469169_50469856 | CERS5 | 91012 | 4.9605706585355 | -0.73901636 | 6 | only_H3vsUI | Down | notDM | Loss | 7 |
| **12_50475785_50478843_91012** | 12_50475785_50478843 | CERS5 | 91012 | 4.91513062641384 | 0.91566181 | 1 | only_H3vsUI | Down | notDM | Gain | 4 |

| | | | | | | | | | | | |
|---|---|---|---|---|---|---|---|---|---|---|---|
| **12_50559514_50560899_91012** | 12_50559514_50560899 | CERS5 | 91012 | 2.31175386105575 | -0.40391899 | 1 | only_H3vsUI | Down | notDM | Loss | 8 |
| **12_50614103_50617013_91012** | 12_50614103_50617013 | CERS5 | 91012 | 4.72369368884136 | 0.58027183 | 4 | only_H3vsUI | Down | notDM | Loss | 4 |
| **12_50633926_50637239_91012** | 12_50633926_50637239 | CERS5 | 91012 | 4.86239974763749 | -0.64619015 | 6 | only_H3vsUI | Down | notDM | Loss | 2 |
| **12_50642342_50643010_91012** | 12_50642342_50643010 | CERS5 | 91012 | 4.90983940484344 | -0.94028728 | 6 | only_H3vsUI | Down | notDM | Gain | 2 |
| **12_50676153_50677318_91012** | 12_50676153_50677318 | CERS5 | 91012 | 5.06080356874801 | -0.69629033 | 1 | only_H3vsUI | Down | notDM | Loss | 7 |
| **12_50677461_50678765_91012** | 12_50677461_50678765 | CERS5 | 91012 | 5.06571412729979 | -0.79462716 | 1 | only_H3vsUI | Down | notDM | Loss | 7 |
| **12_50790082_50791377_91012** | 12_50790082_50791377 | CERS5 | 91012 | 5.3596609708971 | -0.74066702 | 6 | only_H3vsUI | Down | notDM | Gain | 2 |
| **12_51415927_51420005_25840** | 12_51415927_51420005 | METTL7A | 25840 | 4.9873042019775 | 0.79882609 | 2 | only_H3vsUI | UpDown | notDM | Loss | 6 |
| **12_51415927_51420005_7024** | 12_51415927_51420005 | TFCP2 | 7024 | 5.16708683161889 | -0.74104036 | 2 | only_H3vsUI | Down | notDM | Loss | 2 |
| **12_51420299_51421301_25840** | 12_51420299_51421301 | METTL7A | 25840 | 5.00642753167152 | 0.79882609 | 1 | only_H3vsUI | UpDown | notDM | Loss | 3 |
| **12_51420299_51421301_7024** | 12_51420299_51421301 | TFCP2 | 7024 | 5.16323892063956 | -0.74104036 | 1 | only_H3vsUI | Down | notDM | Loss | 7 |
| **12_51421506_51423307_25840** | 12_51421506_51423307 | METTL7A | 25840 | 5.01156198597876 | 0.71380718 | 4 | only_H3vsUI | UpDown | notDM | Gain | 1 |
| **12_51475145_51477208_7024** | 12_51475145_51477208 | TFCP2 | 7024 | 4.95288442433021 | -0.85329566 | 1 | only_H3vsUI | Down | notDM | Gain | 2 |
| **12_51564482_51566550_7024** | 12_51564482_51566550 | TFCP2 | 7024 | 2.57634135020579 | -0.00989934 | 1 | only_H3vsUI | Down | notDM | Loss | 8 |
| **12_51567036_51567626_7024** | 12_51567036_51567626 | TFCP2 | 7024 | 2.04921802267018 | -0.00989934 | 5 | only_H3vsUI | Down | notDM | Gain | 4 |
| **12_51631622_51632475_25840** | 12_51631622_51632475 | METTL7A | 25840 | 5.4952861816854 | 0.83901708 | 1 | only_H3vsUI | UpDown | notDM | Gain | 1 |
| **12_51631622_51632475_7024** | 12_51631622_51632475 | TFCP2 | 7024 | 4.81089085559314 | -0.64321344 | 1 | only_H3vsUI | Down | notDM | Gain | 2 |
| **12_51633049_51634299_25840** | 12_51633049_51634299 | METTL7A | 25840 | 5.49726284815176 | 0.84647544 | 1 | only_H3vsUI | UpDown | notDM | Loss | 6 |
| **12_51633049_51634299_7024** | 12_51633049_51634299 | TFCP2 | 7024 | 4.82036568504324 | -0.70293807 | 1 | only_H3vsUI | Down | notDM | Loss | 7 |
| **12_52670137_52672905_3852** | 12_52670137_52672905 | KRT5 | 3852 | 5.38262750888013 | 0.74440243 | 7 | only_H3vsUI | UpDown | notDM | Gain | 1 |
| **12_52674399_52677020_144501** | 12_52674399_52677020 | KRT80 | 144501 | 4.94751704353469 | 0.71484356 | 7 | notSpecific | Up | DM | Gain | 9 |
| **12_52674399_52677020_3852** | 12_52674399_52677020 | KRT5 | 3852 | 5.37515862457543 | 0.74320544 | 7 | only_H3vsUI | UpDown | DM | Gain | 9 |
| **12_52674399_52677020_3853** | 12_52674399_52677020 | KRT6A | 3853 | 5.32255419274733 | 0.74847977 | 7 | notSpecific | Down | DM | Gain | 9 |
| **12_52674399_52677020_3854** | 12_52674399_52677020 | KRT6B | 3854 | 5.22760650715254 | 0.75430256 | 7 | notSpecific | UpDown | DM | Gain | 9 |
| **12_52674399_52677020_3855** | 12_52674399_52677020 | KRT7 | 3855 | 4.67554005571353 | 0.72840805 | 7 | notSpecific | DownUp | DM | Gain | 9 |
| **12_52674399_52677020_9119** | 12_52674399_52677020 | KRT75 | 9119 | 5.17923859559814 | 0.74810685 | 7 | notSpecific | UpDown | DM | Gain | 9 |
| **12_52738408_52740845_3852** | 12_52738408_52740845 | KRT5 | 3852 | 5.23904658855118 | 0.99926211 | 6 | only_H3vsUI | UpDown | notDM | Gain | 1 |
| **12_52888679_52890467_3852** | 12_52888679_52890467 | KRT5 | 3852 | 4.37615705778325 | 0.90082996 | 6 | only_H3vsUI | UpDown | notDM | Gain | 1 |
| **12_52932016_52934095_3852** | 12_52932016_52934095 | KRT5 | 3852 | 4.2498096094018 | -0.27941167 | 6 | only_H3vsUI | UpDown | notDM | Loss | 3 |
| **12_53214813_53217273_3856** | 12_53214813_53217273 | KRT8 | 3856 | 4.91166886928687 | -0.69096582 | 6 | only_H3vsUI | Down | notDM | Loss | 2 |
| **12_53220880_53221825_3856** | 12_53220880_53221825 | KRT8 | 3856 | 4.88673882256963 | -0.65064063 | 6 | only_H3vsUI | Down | notDM | Gain | 2 |
| **12_53272228_53273442_3856** | 12_53272228_53273442 | KRT8 | 3856 | 4.40529512304229 | 0.44782814 | 7 | only_H3vsUI | Down | notDM | Loss | 5 |
| **12_53288187_53297441_3856** | 12_53288187_53297441 | KRT8 | 3856 | 3.15472820744016 | 0.67074447 | 1 | only_H3vsUI | Down | notDM | Loss | 4 |
| **12_53299022_53304221_3856** | 12_53299022_53304221 | KRT8 | 3856 | 2.19312459835446 | 0.81568234 | 7 | only_H3vsUI | Down | notDM | Loss | 4 |

| | | | | | | | | | | | |
|---|---|---|---|---|---|---|---|---|---|---|---|
| **12_53316089_53317329_3856** | 12_53316089_53317329 | KRT8 | 3856 | 4.23610880158728 | 0.49915913 | 7 | only_H3vsUI | Down | notDM | Gain | 1 |
| **12_53317820_53321134_3856** | 12_53317820_53321134 | KRT8 | 3856 | 4.27770087628879 | 0.52834296 | 7 | only_H3vsUI | Down | notDM | Gain | 5 |
| **12_53337935_53342718_3856** | 12_53337935_53342718 | KRT8 | 3856 | 4.59183229527802 | 0.75649387 | 7 | only_H3vsUI | Down | notDM | Loss | 5 |
| **12_53343471_53347564_3856** | 12_53343471_53347564 | KRT8 | 3856 | 4.6493835437054 | 0.75689522 | 1 | only_H3vsUI | Down | notDM | Loss | 4 |
| **12_53367844_53371597_3856** | 12_53367844_53371597 | KRT8 | 3856 | 4.83871059795315 | 0.59440228 | 7 | only_H3vsUI | Down | notDM | Loss | 5 |
| **12_53385859_53388040_3856** | 12_53385859_53388040 | KRT8 | 3856 | 4.9394843079786 | 0.69600262 | 7 | only_H3vsUI | Down | notDM | Gain | 5 |
| **12_53397829_53399858_3856** | 12_53397829_53399858 | KRT8 | 3856 | 4.99547285217985 | 0.9564313 | 2 | only_H3vsUI | Down | notDM | Gain | 4 |
| **12_53400814_53405785_3856** | 12_53400814_53405785 | KRT8 | 3856 | 5.00837871027616 | 0.94998758 | 1 | only_H3vsUI | Down | notDM | Gain | 5 |
| **12_53437577_53440882_3856** | 12_53437577_53440882 | KRT8 | 3856 | 5.14211090267296 | 0.8601943 | 1 | only_H3vsUI | Down | notDM | Loss | 4 |
| **12_53446647_53448216_3856** | 12_53446647_53448216 | KRT8 | 3856 | 5.16961860100345 | 0.76197624 | 5 | only_H3vsUI | Down | notDM | Gain | 4 |
| **12_53737942_53738556_3856** | 12_53737942_53738556 | KRT8 | 3856 | 5.64253969911208 | -0.89699542 | 2 | only_H3vsUI | Down | notDM | Loss | 7 |
| **12_54190642_54193381_23583** | 12_54190642_54193381 | SMUG1 | 23583 | 5.59033348637268 | -0.78410368 | 6 | only_H3vsUI | Down | notDM | Gain | 2 |
| **12_54221731_54223545_3227** | 12_54221731_54223545 | HOXC11 | 3227 | 5.15645222682501 | -0.68482273 | 6 | only_H3vsUI | DownUp | notDM | Loss | 3 |
| **12_54331324_54332312_3227** | 12_54331324_54332312 | HOXC11 | 3227 | 4.53908865047503 | 0.95586311 | 2 | only_H3vsUI | DownUp | notDM | Loss | 3 |
| **12_54332850_54335920_3227** | 12_54332850_54335920 | HOXC11 | 3227 | 4.49126361626467 | 0.88276337 | 2 | only_H3vsUI | DownUp | notDM | Loss | 6 |
| **12_54346789_54347811_3227** | 12_54346789_54347811 | HOXC11 | 3227 | 4.28107884072894 | 0.67166565 | 4 | only_H3vsUI | DownUp | notDM | Loss | 3 |
| **12_54372337_54373610_3227** | 12_54372337_54373610 | HOXC11 | 3227 | 3.73455982157948 | 0.95429905 | 6 | only_H3vsUI | DownUp | notDM | Loss | 3 |
| **12_54378923_54379618_3227** | 12_54378923_54379618 | HOXC11 | 3227 | 4.07965147707384 | 0.9538155 | 2 | only_H3vsUI | DownUp | notDM | Gain | 1 |
| **12_54382326_54384571_3227** | 12_54382326_54384571 | HOXC11 | 3227 | 4.1879717016474 | 0.89994903 | 2 | only_H3vsUI | DownUp | notDM | Loss | 6 |
| **12_54406350_54407727_3227** | 12_54406350_54407727 | HOXC11 | 3227 | 4.59593690626917 | -0.62454502 | 3 | only_H3vsUI | DownUp | notDM | Loss | 3 |
| **12_54409473_54410351_23583** | 12_54409473_54410351 | SMUG1 | 23583 | 5.23647175970172 | 0.71086063 | 3 | only_H3vsUI | Down | notDM | Loss | 1 |
| **12_54409473_54410351_3227** | 12_54409473_54410351 | HOXC11 | 3227 | 4.62903223107103 | 0.58508581 | 3 | only_H3vsUI | DownUp | notDM | Loss | 3 |
| **12_54410532_54412751_23583** | 12_54410532_54412751 | SMUG1 | 23583 | 5.23038249478947 | 0.89400813 | 1 | only_H3vsUI | Down | notDM | Loss | 4 |
| **12_54410532_54412751_3227** | 12_54410532_54412751 | HOXC11 | 3227 | 4.63970557341718 | 0.46796555 | 1 | only_H3vsUI | DownUp | notDM | Loss | 6 |
| **12_54581211_54582559_23583** | 12_54581211_54582559 | SMUG1 | 23583 | 2.23552844690755 | 0.18770021 | 1 | only_H3vsUI | Down | notDM | Gain | 4 |
| **12_54584830_54585461_23583** | 12_54584830_54585461 | SMUG1 | 23583 | 3.32387060654051 | 0.47503602 | 6 | only_H3vsUI | Down | notDM | Gain | 1 |
| **12_54585580_54586143_23583** | 12_54585580_54586143 | SMUG1 | 23583 | 3.44839710345777 | -0.64805143 | 6 | only_H3vsUI | Down | notDM | Loss | 2 |
| **12_54608384_54609356_23583** | 12_54608384_54609356 | SMUG1 | 23583 | 4.40874860618424 | 0.3990531 | 4 | only_H3vsUI | Down | notDM | Gain | 1 |
| **12_54609723_54611870_23583** | 12_54609723_54611870 | SMUG1 | 23583 | 4.43137984884194 | 0.80084531 | 2 | only_H3vsUI | Down | notDM | Loss | 4 |
| **12_54651378_54655487_23583** | 12_54651378_54655487 | SMUG1 | 23583 | 4.83633677716889 | 0.85772361 | 5 | only_H3vsUI | Down | notDM | Loss | 4 |
| **12_54761945_54763823_23583** | 12_54761945_54763823 | SMUG1 | 23583 | 5.25339374267052 | 0.69737478 | 4 | only_H3vsUI | Down | notDM | Loss | 1 |
| **12_57442505_57443349_4640** | 12_57442505_57443349 | MYO1A | 4640 | 2.7427251313047 | -0.09878391 | 6 | only_H3vsUI | Up | notDM | Loss | 3 |
| **12_57482952_57484911_1649** | 12_57482952_57484911 | DDIT3 | 1649 | 5.63285192599825 | 0.83816734 | 1 | only_DIvsD7 | Up | notDM | Loss | 6 |
| **12_57623022_57625298_1649** | 12_57623022_57625298 | DDIT3 | 1649 | 5.46090235098037 | 0.76482112 | 1 | only_DIvsD7 | Up | notDM | Loss | 6 |

| | | | | | | | | | | | |
|---|---|---|---|---|---|---|---|---|---|---|---|
| **12_57764379_57767409_1649** | 12_57764379_57767409 | DDIT3 | 1649 | 5.16699814398315 | -0.84122173 | 7 | only_DIvsD7 | Up | notDM | Gain | 5 |
| **12_57822576_57824229_1649** | 12_57822576_57824229 | DDIT3 | 1649 | 4.95458980612469 | 0.68531615 | 2 | only_DIvsD7 | Up | notDM | Gain | 5 |
| **12_57824936_57825800_1649** | 12_57824936_57825800 | DDIT3 | 1649 | 4.94694817795238 | 0.68531615 | 5 | only_DIvsD7 | Up | notDM | Gain | 1 |
| **12_57882031_57883707_1649** | 12_57882031_57883707 | DDIT3 | 1649 | 4.48563626235112 | 0.34351935 | 1 | only_DIvsD7 | Up | notDM | Loss | 6 |
| **12_57910678_57914300_1649** | 12_57910678_57914300 | DDIT3 | 1649 | 0 | 0.01477761 | 1 | only_DIvsD7 | Up | notDM | Loss | 8 |
| **12_57914688_57916364_1649** | 12_57914688_57916364 | DDIT3 | 1649 | 2.5910646070265 | -0.07587374 | 1 | only_DIvsD7 | Up | notDM | Gain | 4 |
| **12_57916975_57918677_1649** | 12_57916975_57918677 | DDIT3 | 1649 | 3.42764837118693 | 0.46602667 | 1 | only_DIvsD7 | Up | notDM | Loss | 6 |
| **12_57938659_57940670_1649** | 12_57938659_57940670 | DDIT3 | 1649 | 4.38669511177562 | 0.52182741 | 1 | only_DIvsD7 | Up | notDM | Loss | 6 |
| **12_58014711_58016073_65012** | 12_58014711_58016073 | SLC26A10 | 65012 | 3.00860017176192 | 0 | 2 | only_DIvsD7 | DownUp | notDM | Loss | 3 |
| **12_58024721_58026438_1649** | 12_58024721_58026438 | DDIT3 | 1649 | 5.04305954198676 | 0.64795712 | 1 | only_DIvsD7 | Up | notDM | Loss | 6 |
| **12_58024721_58026438_65012** | 12_58024721_58026438 | SLC26A10 | 65012 | 3.8079407212155 | 0.00342535 | 1 | only_DIvsD7 | DownUp | notDM | Loss | 3 |
| **12_58026590_58027968_1649** | 12_58026590_58027968 | DDIT3 | 1649 | 5.05034881699508 | 0.67379113 | 1 | only_DIvsD7 | Up | notDM | Gain | 1 |
| **12_58144143_58146025_1649** | 12_58144143_58146025 | DDIT3 | 1649 | 5.36143506063787 | 0.74867283 | 1 | only_DIvsD7 | Up | notDM | Gain | 5 |
| **12_58164385_58168096_1649** | 12_58164385_58168096 | DDIT3 | 1649 | 5.39809111686044 | 0.88041318 | 1 | only_DIvsD7 | Up | notDM | Loss | 6 |
| **12_58237281_58240244_1649** | 12_58237281_58240244 | DDIT3 | 1649 | 5.50917966412525 | 0.83998186 | 1 | only_DIvsD7 | Up | notDM | Loss | 6 |
| **12_58335467_58336964_1649** | 12_58335467_58336964 | DDIT3 | 1649 | 5.62445639759322 | 0.80007108 | 1 | only_DIvsD7 | Up | notDM | Loss | 6 |
| **12_64953943_64956013_2799** | 12_64953943_64956013 | GNS | 2799 | 5.29493994391899 | 0.74905981 | 6 | only_DIvsD7 | DownUp | notDM | Gain | 2 |
| **12_65127398_65128506_2799** | 12_65127398_65128506 | GNS | 2799 | 4.39308360208946 | 0.76445875 | 6 | only_DIvsD7 | DownUp | notDM | Gain | 2 |
| **12_65151360_65153121_2799** | 12_65151360_65153121 | GNS | 2799 | 1.81954393554187 | -0.59210538 | 2 | only_DIvsD7 | DownUp | notDM | Loss | 8 |
| **12_65153374_65154558_2799** | 12_65153374_65154558 | GNS | 2799 | 2.17318626841227 | -0.66241693 | 2 | only_DIvsD7 | DownUp | notDM | Loss | 8 |
| **12_66270316_66271685_84298** | 12_66270316_66271685 | LLPH | 84298 | 5.40277363372269 | -0.81274063 | 6 | only_H3vsUI | DownUp | notDM | Gain | 2 |
| **12_66273319_66274984_84298** | 12_66273319_66274984 | LLPH | 84298 | 5.39706880929667 | -0.76339499 | 6 | only_H3vsUI | DownUp | notDM | Gain | 2 |
| **12_66319393_66321639_84298** | 12_66319393_66321639 | LLPH | 84298 | 5.3071621660683 | -0.77582589 | 6 | only_H3vsUI | DownUp | notDM | Loss | 7 |
| **12_66442291_66443566_84298** | 12_66442291_66443566 | LLPH | 84298 | 4.90839386108778 | -0.57762937 | 6 | only_H3vsUI | DownUp | notDM | Gain | 1 |
| **12_66522664_66524286_84298** | 12_66522664_66524286 | LLPH | 84298 | 2.29446622616159 | 0.41471039 | 2 | only_H3vsUI | DownUp | notDM | Gain | 4 |
| **12_66524806_66526712_84298** | 12_66524806_66526712 | LLPH | 84298 | 2.41497334797082 | 0.45929244 | 1 | only_H3vsUI | DownUp | notDM | Gain | 4 |
| **12_66561853_66563259_84298** | 12_66561853_66563259 | LLPH | 84298 | 4.57255796087217 | 0.43138412 | 5 | only_H3vsUI | DownUp | notDM | Gain | 5 |
| **12_66629247_66630362_51643** | 12_66629247_66630362 | TMBIM4 | 51643 | 4.81555782610211 | 0.69650657 | 2 | notSpecific | Down | DM | Gain | 9 |
| **12_6795677_6798259_171017** | 12_6795677_6798259 | ZNF384 | 171017 | 2.62117628177504 | 0.72024979 | 1 | notSpecific | Up | DM | Loss | 10 |
| **12_6795677_6798259_196500** | 12_6795677_6798259 | PIANP | 196500 | 4.03730695089709 | -0.06621903 | 1 | notSpecific | DownUp | DM | Loss | 10 |
| **12_6795677_6798259_50813** | 12_6795677_6798259 | COPS7A | 50813 | 4.54648096953927 | -0.68655396 | 1 | notSpecific | Up | DM | Loss | 10 |
| **12_6795677_6798259_51147** | 12_6795677_6798259 | ING4 | 51147 | 4.36867729541279 | 0.53884719 | 1 | notSpecific | UpDown | DM | Loss | 10 |
| **12_6795677_6798259_51258** | 12_6795677_6798259 | MRPL51 | 51258 | 5.28482085958242 | 0.79385626 | 1 | notSpecific | DownUp | DM | Loss | 10 |
| **12_6795677_6798259_8079** | 12_6795677_6798259 | MLF2 | 8079 | 4.80602387135308 | -0.73783528 | 1 | notSpecific | Down | DM | Loss | 10 |

| | | | | | | | | | | | |
|---|---|---|---|---|---|---|---|---|---|---|---|
| **12_6795677_6798259_83461** | 12_6795677_6798259 | CDCA3 | 83461 | 5.20993034455343 | 0.82052924 | 1 | notSpecific | Down | DM | Loss | 10 |
| **12_6795677_6798259_9918** | 12_6795677_6798259 | NCAPD2 | 9918 | 5.28422989671813 | 0.72291522 | 1 | notSpecific | Down | DM | Loss | 10 |
| **12_687107_688833_4815** | 12_687107_688833 | NINJ2 | 4815 | 4.48787336598845 | 0.45281459 | 6 | only_H3vsUI | UpDown | notDM | Loss | 5 |
| **12_7122560_7125718_10162** | 12_7122560_7125718 | LPCAT3 | 10162 | 1.78532983501077 | 0.09117871 | 1 | notSpecific | Up | DM | Loss | 10 |
| **12_7122560_7125718_10436** | 12_7122560_7125718 | EMG1 | 10436 | 4.62845023788495 | 0.54004646 | 1 | notSpecific | Up | DM | Loss | 10 |
| **12_7122560_7125718_2026** | 12_7122560_7125718 | ENO2 | 2026 | 4.99477452700426 | 0.66687882 | 1 | notSpecific | Up | DM | Loss | 10 |
| **12_7122560_7125718_716** | 12_7122560_7125718 | C1S | 716 | 4.62596032180296 | 0.5051234 | 1 | notSpecific | UpDown | DM | Loss | 10 |
| **12_7122560_7125718_83461** | 12_7122560_7125718 | CDCA3 | 83461 | 5.20991427465055 | 0.72588133 | 1 | notSpecific | Down | DM | Loss | 10 |
| **12_718964_720314_4815** | 12_718964_720314 | NINJ2 | 4815 | 0 | 0.30227139 | 3 | only_H3vsUI | UpDown | notDM | Gain | 4 |
| **12_72665344_72666361_29953** | 12_72665344_72666361 | TRHDE | 29953 | 2.22788670461367 | 0.00463039 | 4 | only_H3vsUI | Down | notDM | Loss | 3 |
| **12_732281_733236_4815** | 12_732281_733236 | NINJ2 | 4815 | 4.10376952319367 | 0.2305493 | 6 | only_H3vsUI | UpDown | notDM | Loss | 5 |
| **12_739669_744139_4815** | 12_739669_744139 | NINJ2 | 4815 | 3.93771844361726 | -0.43015313 | 7 | only_H3vsUI | UpDown | notDM | Gain | 4 |
| **12_77158322_77160172_1466** | 12_77158322_77160172 | CSRP2 | 1466 | 5.05164637216553 | -0.67494407 | 1 | only_DIvsD7 | Up | notDM | Loss | 7 |
| **12_77271784_77272690_1466** | 12_77271784_77272690 | CSRP2 | 1466 | 2.04139268515822 | 0.88258976 | 1 | only_DIvsD7 | Up | notDM | Gain | 4 |
| **12_77273013_77274262_1466** | 12_77273013_77274262 | CSRP2 | 1466 | 2.33445375115093 | 0.87167556 | 1 | only_DIvsD7 | Up | notDM | Loss | 6 |
| **12_8332975_8334534_440078** | 12_8332975_8334534 | FAM66C | 440078 | 2.04921802267018 | 0.02105391 | 2 | only_DIvsD7 | UpDown | notDM | Loss | 3 |
| **12_86158957_86159709_4922** | 12_86158957_86159709 | NTS | 4922 | 5.03489705065408 | 0.67354638 | 6 | only_H3vsUI | DownUp | notDM | Gain | 1 |
| **12_96194808_96199782_59277** | 12_96194808_96199782 | NTN4 | 59277 | 4.01173956138832 | 0.59925811 | 7 | notSpecific | Down | DM | Gain | 9 |
| **12_96335734_96336386_120935** | 12_96335734_96336386 | CCDC38 | 120935 | 1.63346845557959 | 0.0018778 | 5 | only_H3vsUI | Up | notDM | Gain | 1 |
| **12_96337222_96338114_120935** | 12_96337222_96338114 | CCDC38 | 120935 | 2.90091306773767 | 0.00224662 | 2 | only_H3vsUI | Up | notDM | Gain | 4 |
| **13_108867969_108868672_3981** | 13_108867969_108868672 | LIG4 | 3981 | 1.91907809237607 | 0.11264105 | 2 | only_DIvsD7 | DownUp | notDM | Gain | 4 |
| **13_111358811_111360086_3621** | 13_111358811_111360086 | ING1 | 3621 | 3.74350976472843 | 0.02636331 | 1 | only_DIvsD7 | Up | notDM | Loss | 3 |
| **13_111363495_111364659_3621** | 13_111363495_111364659 | ING1 | 3621 | 2.985426474083 | 0.04216919 | 2 | only_DIvsD7 | Up | notDM | Loss | 6 |
| **13_111368352_111370689_3621** | 13_111368352_111370689 | ING1 | 3621 | 3.01072386539177 | -0.01356574 | 1 | only_DIvsD7 | Up | notDM | Loss | 6 |
| **13_113840302_113841742_7027** | 13_113840302_113841742 | TFDP1 | 7027 | 5.5991371476082 | 0.84624506 | 5 | only_H3vsUI | Down | notDM | Gain | 1 |
| **13_114103790_114105163_113622** | 13_114103790_114105163 | ADPRHL1 | 113622 | 0 | 0.00966842 | 1 | notSpecific | Up | DM | Loss | 10 |
| **13_114239797_114240845_7027** | 13_114239797_114240845 | TFDP1 | 7027 | 2.86981820797933 | 0.75799059 | 2 | only_H3vsUI | Down | notDM | Loss | 8 |
| **13_114534734_114535368_2621** | 13_114534734_114535368 | GAS6 | 2621 | 3.12936759572299 | 0 | 3 | only_DIvsD7 | DownUp | notDM | Gain | 1 |
| **13_114540064_114543172_2621** | 13_114540064_114543172 | GAS6 | 2621 | 3.02077548819356 | -0.00041373 | 5 | only_DIvsD7 | DownUp | notDM | Loss | 6 |
| **13_115078593_115079663_283489** | 13_115078593_115079663 | CHAMP1 | 283489 | 2.51321760006794 | 0.22487206 | 1 | notSpecific | Down | DM | Gain | 9 |
| **13_115078593_115079663_65110** | 13_115078593_115079663 | UPF3A | 65110 | 4.49792434257526 | -0.44948191 | 1 | notSpecific | UpDown | DM | Gain | 9 |
| **13_115078593_115079663_8881** | 13_115078593_115079663 | CDC16 | 8881 | 4.89338998445112 | -0.64509872 | 1 | notSpecific | Down | DM | Gain | 9 |
| **13_41704836_41706683_89890** | 13_41704836_41706683 | KBTBD6 | 89890 | 2.30102999566398 | -0.62646507 | 1 | only_H3vsUI | Down | notDM | Gain | 4 |
| **13_41753133_41754443_89890** | 13_41753133_41754443 | KBTBD6 | 89890 | 4.66513990661433 | -0.49889715 | 3 | only_H3vsUI | Down | notDM | Loss | 2 |

| | | | | | | | | | | | |
|---|---|---|---|---|---|---|---|---|---|---|---|
| **13_44451687_44453300_144811** | 13_44451687_44453300 | LACC1 | 144811 | 2.59659709562646 | 0.0667094 | 2 | only_DIvsD7 | DownUp | notDM | Loss | 7 |
| **13_44454464_44455461_144811** | 13_44454464_44455461 | LACC1 | 144811 | 2.88761730033574 | 0.11060375 | 1 | only_DIvsD7 | DownUp | notDM | Gain | 2 |
| **13_45561658_45563496_26747** | 13_45561658_45563496 | NUFIP1 | 26747 | 2.0899051114394 | -0.45422454 | 1 | only_H3vsUI | DownUp | notDM | Loss | 8 |
| **13_45563993_45565229_26747** | 13_45563993_45565229 | NUFIP1 | 26747 | 2.57634135020579 | -0.50385053 | 1 | only_H3vsUI | DownUp | notDM | Loss | 7 |
| **13_47127949_47130492_2098** | 13_47127949_47130492 | ESD | 2098 | 5.38161499898881 | 0.87064217 | 1 | only_DIvsD7 | DownUp | notDM | Loss | 6 |
| **13_47371415_47372347_2098** | 13_47371415_47372347 | ESD | 2098 | 2.17318626841227 | 0.73158037 | 5 | only_DIvsD7 | DownUp | notDM | Loss | 3 |
| **13_50016974_50017999_83852** | 13_50016974_50017999 | SETDB2 | 83852 | 2.70926996097583 | 0.20830995 | 1 | only_H3vsUI | Down | notDM | Gain | 4 |
| **13_50018722_50019990_83852** | 13_50018722_50019990 | SETDB2 | 83852 | 2.33041377334919 | 0.20830995 | 1 | only_H3vsUI | Down | notDM | Loss | 3 |
| **13_52432261_52433757_83446** | 13_52432261_52433757 | CCDC70 | 83446 | 3.37309598707873 | 0 | 4 | only_H3vsUI | UpDown | notDM | Loss | 3 |
| **13_53001806_53003463_51028** | 13_53001806_53003463 | VPS36 | 51028 | 1.60205999132796 | 0.01378747 | 6 | notSpecific | Down | DM | Gain | 9 |
| **13_53001806_53003463_55901** | 13_53001806_53003463 | THSD1 | 55901 | 4.33334649842439 | 0.76466065 | 6 | notSpecific | Down | DM | Gain | 9 |
| **13_67767446_67769227_5101** | 13_67767446_67769227 | PCDH9 | 5101 | 4.54706054685457 | 0.64656737 | 3 | notSpecific | Down | DM | Gain | 9 |
| **13_73357126_73358211_10464** | 13_73357126_73358211 | PIBF1 | 10464 | 2.9532763366673 | 0.19103109 | 2 | notSpecific | Up | DM | Gain | 9 |
| **13_73357126_73358211_22894** | 13_73357126_73358211 | DIS3 | 22894 | 2.95133751879592 | -0.29614586 | 2 | notSpecific | Up | DM | Gain | 9 |
| **13_73357126_73358211_440145** | 13_73357126_73358211 | MZT1 | 440145 | 4.74274869094496 | -0.52073662 | 2 | notSpecific | Down | DM | Gain | 9 |
| **13_73357126_73358211_79866** | 13_73357126_73358211 | BORA | 79866 | 4.74085198208798 | -0.4972293 | 2 | notSpecific | Down | DM | Gain | 9 |
| **13_76110156_76111747_170622** | 13_76110156_76111747 | COMMD6 | 170622 | 2.38916608436453 | -0.36797317 | 2 | only_H3vsUI | Up | notDM | Loss | 8 |
| **13_76112411_76112877_170622** | 13_76112411_76112877 | COMMD6 | 170622 | 2.62531245096167 | -0.36797317 | 5 | only_H3vsUI | Up | notDM | Gain | 2 |
| **13_76124094_76125347_170622** | 13_76124094_76125347 | COMMD6 | 170622 | 2.72591163229505 | 0.0438148 | 1 | only_H3vsUI | Up | notDM | Loss | 3 |
| **13_76340853_76348179_170622** | 13_76340853_76348179 | COMMD6 | 170622 | 5.35963251129465 | 0.79663843 | 7 | only_H3vsUI | Up | notDM | Gain | 5 |
| **13_79975457_79979734_64062** | 13_79975457_79979734 | RBM26 | 64062 | 2.79028516403324 | -0.46829133 | 1 | only_H3vsUI | Down | notDM | Loss | 8 |
| **13_79980627_79981973_64062** | 13_79980627_79981973 | RBM26 | 64062 | 2.4456042032736 | -0.48317725 | 1 | only_H3vsUI | Down | notDM | Gain | 4 |
| **13_95246034_95248156_160897** | 13_95246034_95248156 | GPR180 | 160897 | 3.77829599108883 | -0.24125231 | 2 | notSpecific | Down | DM | Loss | 10 |
| **13_95246034_95248156_23483** | 13_95246034_95248156 | TGDS | 23483 | 2.55144999797288 | -0.25999867 | 2 | notSpecific | Down | DM | Loss | 10 |
| **13_96326744_96328867_100289274** | 13_96326744_96328867 | DNAJC3-DT | 100289274 | 2.49554433754645 | -0.00868236 | 1 | only_H3vsUI | UpDown | notDM | Gain | 4 |
| **13_96329889_96331437_100289274** | 13_96329889_96331437 | DNAJC3-DT | 100289274 | 2.85247999363686 | -0.02817749 | 1 | only_H3vsUI | UpDown | notDM | Gain | 4 |
| **13_98087048_98088184_5911** | 13_98087048_98088184 | RAP2A | 5911 | 2.75891189239797 | 0.23708267 | 2 | only_H3vsUI | Up | notDM | Gain | 4 |
| **14_100034649_100046734_84193** | 14_100034649_100046734 | SETD3 | 84193 | 4.94168531437513 | 0.65864495 | 7 | notSpecific | UpDown | DM | Loss | 9 |
| **14_102029805_102032179_1735** | 14_102029805_102032179 | DIO3 | 1735 | 3.32613095671079 | 0 | 2 | notSpecific | UpDown | DM | Loss | 10 |
| **14_102974430_102975715_122416** | 14_102974430_102975715 | ANKRD9 | 122416 | 2.6170003411209 | 0.1268599 | 1 | only_DIvsD7 | DownUp | notDM | Loss | 7 |
| **14_102976417_102977368_122416** | 14_102976417_102977368 | ANKRD9 | 122416 | 2.46389298898591 | 0.0279324 | 2 | only_DIvsD7 | DownUp | notDM | Gain | 4 |
| **14_102982727_102983270_122416** | 14_102982727_102983270 | ANKRD9 | 122416 | 3.81960973275158 | -0.13783732 | 1 | only_DIvsD7 | DownUp | notDM | Gain | 2 |
| **14_102983482_102983855_122416** | 14_102983482_102983855 | ANKRD9 | 122416 | 3.86664172056604 | -0.08899512 | 2 | only_DIvsD7 | DownUp | notDM | Loss | 7 |
| **14_102990082_102990927_122416** | 14_102990082_102990927 | ANKRD9 | 122416 | 4.14476096077607 | -0.23060539 | 2 | only_DIvsD7 | DownUp | notDM | Loss | 7 |

| | | | | | | | | | | | |
|---|---|---|---|---|---|---|---|---|---|---|---|
| **14_103056578_103058260_23186** | 14_103056578_103058260 | RCOR1 | 23186 | 2.98855895687862 | 0.00252729 | 2 | only_DIvsD7 | UpDown | notDM | Loss | 6 |
| **14_103059687_103061877_23186** | 14_103059687_103061877 | RCOR1 | 23186 | 2.65896484266443 | 0.0044713 | 1 | only_DIvsD7 | UpDown | notDM | Gain | 4 |
| **14_21428749_21429656_64745** | 14_21428749_21429656 | METTL17 | 64745 | 4.45460097709776 | -0.9287066 | 4 | only_H3vsUI | DownUp | notDM | Gain | 2 |
| **14_21438674_21439657_64745** | 14_21438674_21439657 | METTL17 | 64745 | 4.26677246351307 | -0.21805323 | 1 | only_H3vsUI | DownUp | notDM | Gain | 2 |
| **14_21454095_21456221_64745** | 14_21454095_21456221 | METTL17 | 64745 | 3.23704079137919 | 0.13935409 | 6 | only_H3vsUI | DownUp | notDM | Gain | 1 |
| **14_21456772_21457812_64745** | 14_21456772_21457812 | METTL17 | 64745 | 2.51587384371168 | 0.34837429 | 5 | only_H3vsUI | DownUp | notDM | Loss | 6 |
| **14_21458329_21460232_64745** | 14_21458329_21460232 | METTL17 | 64745 | 2.28330122870355 | 0.17268173 | 1 | only_H3vsUI | DownUp | notDM | Loss | 8 |
| **14_21522566_21525075_64745** | 14_21522566_21525075 | METTL17 | 64745 | 4.80908139073092 | 0.65640994 | 6 | only_H3vsUI | DownUp | notDM | Gain | 1 |
| **14_21530544_21532496_64745** | 14_21530544_21532496 | METTL17 | 64745 | 4.85978055396536 | 0.65698027 | 6 | only_H3vsUI | DownUp | notDM | Gain | 1 |
| **14_21536781_21538309_64745** | 14_21536781_21538309 | METTL17 | 64745 | 4.89566559452078 | 0.92373607 | 2 | only_H3vsUI | DownUp | notDM | Loss | 6 |
| **14_23283755_23285569_122704** | 14_23283755_23285569 | MRPL52 | 122704 | 4.13257984765974 | 0.5271601 | 5 | only_H3vsUI | UpDown | notDM | Gain | 1 |
| **14_23286611_23291194_122704** | 14_23286611_23291194 | MRPL52 | 122704 | 3.9000939015434 | 0.35122826 | 7 | only_H3vsUI | UpDown | notDM | Loss | 5 |
| **14_23293368_23294527_122704** | 14_23293368_23294527 | MRPL52 | 122704 | 3.66388929862266 | 0.69285143 | 3 | only_H3vsUI | UpDown | notDM | Gain | 1 |
| **14_23299274_23300673_122704** | 14_23299274_23300673 | MRPL52 | 122704 | 2.19589965240923 | 0.04810059 | 1 | only_H3vsUI | UpDown | notDM | Gain | 4 |
| **14_23321793_23323203_122704** | 14_23321793_23323203 | MRPL52 | 122704 | 4.35608324914729 | 0.33459601 | 2 | only_H3vsUI | UpDown | notDM | Gain | 4 |
| **14_23386470_23388110_122704** | 14_23386470_23388110 | MRPL52 | 122704 | 4.94135736493269 | 0.76838648 | 1 | only_H3vsUI | UpDown | notDM | Gain | 1 |
| **14_23388669_23389686_122704** | 14_23388669_23389686 | MRPL52 | 122704 | 4.95207528956265 | 0.76838929 | 4 | only_H3vsUI | UpDown | notDM | Gain | 1 |
| **14_24616954_24618491_10379** | 14_24616954_24618491 | IRF9 | 10379 | 4.13719581194055 | 0.14821741 | 2 | only_H3vsUI | UpDown | notDM | Loss | 6 |
| **14_24629459_24631743_10379** | 14_24629459_24631743 | IRF9 | 10379 | 2.66558099101795 | 0.16063348 | 1 | only_H3vsUI | UpDown | notDM | Loss | 6 |
| **14_24739689_24740651_23351** | 14_24739689_24740651 | KHNYN | 23351 | 5.20000460559912 | 0.69963116 | 1 | only_H3vsUI | DownUp | notDM | Loss | 7 |
| **14_24740875_24741756_23351** | 14_24740875_24741756 | KHNYN | 23351 | 5.19696609782507 | 0.69963116 | 2 | only_H3vsUI | DownUp | notDM | Gain | 1 |
| **14_24899500_24901641_23351** | 14_24899500_24901641 | KHNYN | 23351 | 2.55750720190566 | 0.73044947 | 1 | only_H3vsUI | DownUp | notDM | Loss | 6 |
| **14_24899500_24901641_643866** | 14_24899500_24901641 | CBLN3 | 643866 | 2.88705437805096 | 0.01777003 | 1 | only_H3vsUI | DownUp | notDM | Loss | 8 |
| **14_24910192_24911965_23351** | 14_24910192_24911965 | KHNYN | 23351 | 4.04348017002255 | 0.74689629 | 1 | only_H3vsUI | DownUp | notDM | Loss | 6 |
| **14_24912288_24913115_23351** | 14_24912288_24913115 | KHNYN | 23351 | 4.11889272537362 | 0.74689629 | 1 | only_H3vsUI | DownUp | notDM | Gain | 4 |
| **14_36987966_36990505_7080** | 14_36987966_36990505 | NKX2-1 | 7080 | 0 | 0.00083288 | 4 | notSpecific | DownUp | DM | Loss | 10 |
| **14_37585034_37587471_145282** | 14_37585034_37587471 | MIPOL1 | 145282 | 4.90170346238146 | 0.62002785 | 3 | only_DIvsD7 | UpDown | notDM | Gain | 5 |
| **14_37591221_37592481_145282** | 14_37591221_37592481 | MIPOL1 | 145282 | 4.87352403885685 | 0.64883509 | 6 | only_DIvsD7 | UpDown | notDM | Gain | 1 |
| **14_37640502_37643196_145282** | 14_37640502_37643196 | MIPOL1 | 145282 | 4.37948681371727 | 0.86015045 | 2 | only_DIvsD7 | UpDown | notDM | Gain | 5 |
| **14_37665953_37666998_145282** | 14_37665953_37666998 | MIPOL1 | 145282 | 2.19865708695442 | 0.89201041 | 5 | only_DIvsD7 | UpDown | notDM | Gain | 1 |
| **14_37667374_37668616_145282** | 14_37667374_37668616 | MIPOL1 | 145282 | 2.34439227368511 | 0.89156577 | 2 | only_DIvsD7 | UpDown | notDM | Gain | 4 |
| **14_42075708_42078096_145581** | 14_42075708_42078096 | LRFN5 | 145581 | 0 | 0 | 4 | notSpecific | Up | DM | Loss | 10 |
| **14_50696032_50697901_79944** | 14_50696032_50697901 | L2HGDH | 79944 | 4.90840994914717 | -0.58957658 | 1 | only_H3vsUI | Down | notDM | Loss | 8 |
| **14_50698697_50699626_79944** | 14_50698697_50699626 | L2HGDH | 79944 | 4.89905954732876 | -0.61251382 | 2 | only_H3vsUI | Down | notDM | Loss | 7 |

| | | | | | | | | | | | |
|---|---|---|---|---|---|---|---|---|---|---|---|
| **14_50777153_50778694_79944** | 14_50777153_50778694 | L2HGDH | 79944 | 2.40483371661994 | 0.25580639 | 2 | only_H3vsUI | Down | notDM | Gain | 4 |
| **14_50779423_50780635_79944** | 14_50779423_50780635 | L2HGDH | 79944 | 2.67942789661212 | 0.25580639 | 1 | only_H3vsUI | Down | notDM | Gain | 4 |
| **14_50787937_50788949_79944** | 14_50787937_50788949 | L2HGDH | 79944 | 3.95385629822499 | 0.25104416 | 4 | only_H3vsUI | Down | notDM | Gain | 1 |
| **14_50855956_50857713_79944** | 14_50855956_50857713 | L2HGDH | 79944 | 4.88689662897036 | -0.67397449 | 6 | only_H3vsUI | Down | notDM | Loss | 2 |
| **14_53017888_53019170_57544** | 14_53017888_53019170 | TXNDC16 | 57544 | 2.12057393120585 | -0.09577143 | 1 | only_DIvsD7 | DownUp | notDM | Loss | 8 |
| **14_53020014_53021151_57544** | 14_53020014_53021151 | TXNDC16 | 57544 | 2.85430604180108 | 0.13255157 | 2 | only_DIvsD7 | DownUp | notDM | Gain | 4 |
| **14_61091289_61092886_6495** | 14_61091289_61092886 | SIX1 | 6495 | 4.36726271478424 | 0.56294619 | 4 | only_H3vsUI | Down | notDM | Gain | 1 |
| **14_61113287_61116174_6495** | 14_61113287_61116174 | SIX1 | 6495 | 0.845098040014257 | 0.78455472 | 1 | only_H3vsUI | Down | notDM | Loss | 8 |
| **14_61116338_61117294_6495** | 14_61116338_61117294 | SIX1 | 6495 | 2.20411998265592 | 0.68572297 | 2 | only_H3vsUI | Down | notDM | Loss | 3 |
| **14_61123463_61124884_6495** | 14_61123463_61124884 | SIX1 | 6495 | 3.86242955610601 | -0.26103295 | 2 | only_H3vsUI | Down | notDM | Gain | 4 |
| **14_61125418_61126227_6495** | 14_61125418_61126227 | SIX1 | 6495 | 3.96567197122011 | 0.41276072 | 6 | only_H3vsUI | Down | notDM | Gain | 1 |
| **14_61257060_61258456_6495** | 14_61257060_61258456 | SIX1 | 6495 | 5.14885550836807 | 0.68081974 | 6 | only_H3vsUI | Down | notDM | Gain | 1 |
| **14_61736139_61737067_161291** | 14_61736139_61737067 | TMEM30B | 161291 | 4.03714750363255 | 0.06656594 | 2 | only_H3vsUI | Down | notDM | Loss | 3 |
| **14_61751628_61753647_161291** | 14_61751628_61753647 | TMEM30B | 161291 | 3.56478438450399 | -0.33769578 | 6 | only_H3vsUI | Down | notDM | Gain | 1 |
| **14_65115017_65116724_3306** | 14_65115017_65116724 | HSPA2 | 3306 | 5.03165079355126 | 0.6776416 | 7 | notSpecific | Up | DM | Gain | 9 |
| **14_65115017_65116724_9495** | 14_65115017_65116724 | AKAP5 | 9495 | 5.26198094295258 | 0.69886184 | 7 | notSpecific | UpDown | DM | Gain | 9 |
| **14_70076714_70078031_9766** | 14_70076714_70078031 | SUSD6 | 9766 | 2.44715803134222 | 0.27056709 | 1 | only_H3vsUI | DownUp | notDM | Loss | 3 |
| **14_70078855_70080569_9766** | 14_70078855_70080569 | SUSD6 | 9766 | 2.73798732633343 | 0.27167248 | 1 | only_H3vsUI | DownUp | notDM | Gain | 4 |
| **14_77227272_77227878_22846** | 14_77227272_77227878 | VASH1 | 22846 | 2.55388302664387 | 0.01540371 | 2 | only_DIvsD7 | Down | notDM | Loss | 3 |
| **14_77228031_77229326_22846** | 14_77228031_77229326 | VASH1 | 22846 | 0 | 0.01160515 | 2 | only_DIvsD7 | Down | notDM | Loss | 3 |
| **14_77605654_77608514_283576** | 14_77605654_77608514 | ZDHHC22 | 283576 | 0 | 0.9983321 | 4 | notSpecific | UpDown | DM | Loss | 10 |
| **14_77605654_77608514_85457** | 14_77605654_77608514 | CIPC | 85457 | 4.61360929057012 | 0.63348397 | 4 | notSpecific | Down | DM | Loss | 10 |
| **14_77964271_77965841_10598** | 14_77964271_77965841 | AHSA1 | 10598 | 4.60095112599417 | -0.88541982 | 2 | notSpecific | Up | DM | Loss | 10 |
| **14_77964271_77965841_145501** | 14_77964271_77965841 | ISM2 | 145501 | 0 | 0.99687894 | 2 | notSpecific | UpDown | DM | Loss | 10 |
| **14_77964271_77965841_22938** | 14_77964271_77965841 | SNW1 | 22938 | 5.41773069794064 | -0.9548991 | 2 | notSpecific | Up | DM | Loss | 10 |
| **14_80682260_80684208_1734** | 14_80682260_80684208 | DIO2 | 1734 | 0 | 0 | 7 | notSpecific | Up | DM | Gain | 9 |
| **14_89259480_89260085_161436** | 14_89259480_89260085 | EML5 | 161436 | 2.58658730467176 | 0.00570382 | 2 | only_DIvsD7 | UpDown | notDM | Gain | 1 |
| **14_93671385_93673208_55148** | 14_93671385_93673208 | UBR7 | 55148 | 2.28780172993023 | 0 | 1 | only_H3vsUI | Down | notDM | Gain | 4 |
| **14_93673741_93676100_55148** | 14_93673741_93676100 | UBR7 | 55148 | 2.53402610605613 | 0 | 1 | only_H3vsUI | Down | notDM | Gain | 4 |
| **14_95949230_95950647_161176** | 14_95949230_95950647 | SYNE3 | 161176 | 3.84874318189568 | -0.00494753 | 6 | only_H3vsUI | DownUp | notDM | Loss | 3 |
| **14_99988192_99990305_84193** | 14_99988192_99990305 | SETD3 | 84193 | 4.61255075811203 | 0.50211505 | 3 | notSpecific | UpDown | DM | Loss | 10 |
| **15_32967202_32972260_6447** | 15_32967202_32972260 | SCG5 | 6447 | 0 | 0.09080674 | 3 | notSpecific | Down | DM | Gain | 9 |
| **15_32967202_32972260_9824** | 15_32967202_32972260 | ARHGAP11A | 9824 | 4.77402034534064 | -0.68743057 | 3 | notSpecific | Down | DM | Gain | 9 |
| **15_34329589_34330689_51234** | 15_34329589_34330689 | EMC4 | 51234 | 5.27079990941709 | -0.75540644 | 1 | only_H3vsUI | UpDown | notDM | Gain | 2 |

| | | | | | | | | | | | |
|---|---|---|---|---|---|---|---|---|---|---|---|
| **15_34499698_34501815_51234** | 15_34499698_34501815 | EMC4 | 51234 | 4.18712572810173 | -0.16016682 | 1 | only_H3vsUI | UpDown | notDM | Loss | 8 |
| **15_34515462_34517093_51234** | 15_34515462_34517093 | EMC4 | 51234 | 2.17026171539496 | -0.275083 | 1 | only_H3vsUI | UpDown | notDM | Gain | 4 |
| **15_34517367_34519378_51234** | 15_34517367_34519378 | EMC4 | 51234 | 2.11058971029925 | -0.275083 | 2 | only_H3vsUI | UpDown | notDM | Gain | 4 |
| **15_34628871_34629914_51234** | 15_34628871_34629914 | EMC4 | 51234 | 5.04779259602973 | -0.69557957 | 1 | only_H3vsUI | UpDown | notDM | Gain | 2 |
| **15_38542898_38543747_10125** | 15_38542898_38543747 | RASGRP1 | 10125 | 5.49590632996283 | -0.99090457 | 1 | only_H3vsUI | DownUp | notDM | Loss | 7 |
| **15_38545389_38548951_10125** | 15_38545389_38548951 | RASGRP1 | 10125 | 5.48863108174437 | -0.77995995 | 2 | only_H3vsUI | DownUp | notDM | Gain | 2 |
| **15_38746757_38748241_10125** | 15_38746757_38748241 | RASGRP1 | 10125 | 5.03649715003302 | -0.72777513 | 1 | only_H3vsUI | DownUp | notDM | Loss | 7 |
| **15_38746757_38748241_283742** | 15_38746757_38748241 | FAM98B | 283742 | 2.63447727016073 | 0.00465438 | 1 | only_DIvsD7 | UpDown | notDM | Loss | 8 |
| **15_41198064_41199936_54567** | 15_41198064_41199936 | DLL4 | 54567 | 4.33469495886716 | 0.33818002 | 4 | only_DIvsD7 | Up | notDM | Loss | 3 |
| **15_41220739_41223575_54567** | 15_41220739_41223575 | DLL4 | 54567 | 0 | 0.8423441 | 4 | only_DIvsD7 | Up | notDM | Loss | 3 |
| **15_41233118_41233804_54567** | 15_41233118_41233804 | DLL4 | 54567 | 4.06344595312303 | 0.50719972 | 1 | only_DIvsD7 | Up | notDM | Loss | 3 |
| **15_41542088_41543074_11261** | 15_41542088_41543074 | CHP1 | 11261 | 4.27074869022875 | 0.91192418 | 7 | notSpecific | Up | DM | Gain | 9 |
| **15_41542088_41543074_51203** | 15_41542088_41543074 | NUSAP1 | 51203 | 4.91379266677031 | -0.84968646 | 7 | notSpecific | Up | DM | Gain | 9 |
| **15_41542088_41543074_55192** | 15_41542088_41543074 | DNAJC17 | 55192 | 5.64583376978391 | -0.83913492 | 7 | notSpecific | Up | DM | Gain | 9 |
| **15_44037784_44038405_440278** | 15_44037784_44038405 | CATSPER2P1 | 440278 | 1.89762709129044 | 0.11508595 | 1 | only_H3vsUI | UpDown | notDM | Loss | 3 |
| **15_44039346_44040840_440278** | 15_44039346_44040840 | CATSPER2P1 | 440278 | 2.93701610746481 | 0.11508595 | 2 | only_H3vsUI | UpDown | notDM | Gain | 4 |
| **15_45878050_45879256_26258** | 15_45878050_45879256 | BLOC1S6 | 26258 | 2.20951501454263 | 0.00487983 | 2 | only_H3vsUI | UpDown | notDM | Gain | 4 |
| **15_45879850_45881870_26258** | 15_45879850_45881870 | BLOC1S6 | 26258 | 2.63848925695464 | 0.00487983 | 1 | only_H3vsUI | UpDown | notDM | Loss | 8 |
| **15_55580737_55582951_9488** | 15_55580737_55582951 | PIGB | 9488 | 4.45037229495525 | 0.68616468 | 1 | only_H3vsUI | DownUp | notDM | Gain | 5 |
| **15_55609753_55611027_9488** | 15_55609753_55611027 | PIGB | 9488 | 2.02938377768521 | 0.62449067 | 1 | only_H3vsUI | DownUp | notDM | Gain | 4 |
| **15_55611466_55612647_9488** | 15_55611466_55612647 | PIGB | 9488 | 2.52504480703685 | 0.62449067 | 1 | only_H3vsUI | DownUp | notDM | Gain | 4 |
| **15_55699268_55700259_9488** | 15_55699268_55700259 | PIGB | 9488 | 4.94503505913721 | 0.75644701 | 1 | only_H3vsUI | DownUp | notDM | Gain | 5 |
| **15_55700723_55702077_9488** | 15_55700723_55702077 | PIGB | 9488 | 4.95214802799067 | 0.61794325 | 1 | only_H3vsUI | DownUp | notDM | Loss | 6 |
| **15_55790688_55791709_9488** | 15_55790688_55791709 | PIGB | 9488 | 5.2542023406908 | 0.70614327 | 1 | only_H3vsUI | DownUp | notDM | Gain | 1 |
| **15_60882966_60884492_6095** | 15_60882966_60884492 | RORA | 6095 | 2.33445375115093 | 0.49870708 | 4 | notSpecific | UpDown | DM | Loss | 10 |
| **15_61519164_61521066_6095** | 15_61519164_61521066 | RORA | 6095 | 2.65609820201283 | 0.00011278 | 2 | notSpecific | UpDown | DM | Loss | 10 |
| **15_65202933_65205370_348094** | 15_65202933_65205370 | ANKDD1A | 348094 | 0 | 0.41776689 | 1 | only_H3vsUI | UpDown | notDM | Loss | 3 |
| **15_65577499_65579152_54956** | 15_65577499_65579152 | PARP16 | 54956 | 0 | -0.07760751 | 1 | only_H3vsUI | DownUp | notDM | Loss | 8 |
| **15_65579416_65580384_54956** | 15_65579416_65580384 | PARP16 | 54956 | 2.76937732607614 | -0.06113548 | 1 | only_H3vsUI | DownUp | notDM | Gain | 4 |
| **15_65595492_65596807_54956** | 15_65595492_65596807 | PARP16 | 54956 | 4.21176109370736 | 0.59165855 | 5 | only_H3vsUI | DownUp | notDM | Gain | 1 |
| **15_65669021_65670155_54956** | 15_65669021_65670155 | PARP16 | 54956 | 4.95333920325574 | 0.70818208 | 2 | only_H3vsUI | DownUp | notDM | Loss | 6 |
| **15_67358448_67376269_4088** | 15_67358448_67376269 | SMAD3 | 4088 | 2.42651126136458 | 0.11770822 | 1 | notSpecific | UpDown | DM | Loss | 10 |
| **15_68125437_68127546_10116** | 15_68125437_68127546 | FEM1B | 10116 | 5.64600748463467 | 0.82424507 | 4 | only_DIvsD7 | UpDown | notDM | Loss | 6 |
| **15_68519709_68521733_10116** | 15_68519709_68521733 | FEM1B | 10116 | 4.68492611137529 | -0.54786106 | 1 | only_DIvsD7 | UpDown | notDM | Loss | 7 |

| | | | | | | | | | | | |
|---|---|---|---|---|---|---|---|---|---|---|---|
| **15_68522200_68523439_10116** | 15_68522200_68523439 | FEM1B | 10116 | 4.66934477867316 | -0.54786106 | 1 | only_DIvsD7 | UpDown | notDM | Loss | 7 |
| **15_68549673_68550198_10116** | 15_68549673_68550198 | FEM1B | 10116 | 4.29981226549572 | 0.85955269 | 2 | only_DIvsD7 | UpDown | notDM | Gain | 1 |
| **15_68563028_68566606_10116** | 15_68563028_68566606 | FEM1B | 10116 | 3.54851225634104 | 0.80115971 | 7 | only_DIvsD7 | UpDown | notDM | Gain | 5 |
| **15_68568383_68569998_10116** | 15_68568383_68569998 | FEM1B | 10116 | 2.15836249209525 | 0.19040685 | 1 | only_DIvsD7 | UpDown | notDM | Gain | 4 |
| **15_68570630_68574830_10116** | 15_68570630_68574830 | FEM1B | 10116 | 2.69108149212297 | 0.17674862 | 1 | only_DIvsD7 | UpDown | notDM | Loss | 6 |
| **15_68598728_68599117_10116** | 15_68598728_68599117 | FEM1B | 10116 | 4.45619896465924 | 0.84900414 | 4 | only_DIvsD7 | UpDown | notDM | Gain | 1 |
| **15_68935711_68937180_10116** | 15_68935711_68937180 | FEM1B | 10116 | 5.5629729247227 | 0.80283989 | 6 | only_DIvsD7 | UpDown | notDM | Gain | 1 |
| **15_69106537_69110552_8125** | 15_69106537_69110552 | ANP32A | 8125 | 0 | 0.17918269 | 1 | only_DIvsD7 | Down | notDM | Gain | 8 |
| **15_69113729_69115714_8125** | 15_69113729_69115714 | ANP32A | 8125 | 2.71432975974523 | 0.10377638 | 2 | only_DIvsD7 | Down | notDM | Gain | 4 |
| **15_72475731_72477708_196996** | 15_72475731_72477708 | GRAMD2A | 196996 | 4.09443618801796 | 0.37792401 | 1 | only_H3vsUI | UpDown | notDM | Loss | 6 |
| **15_74471311_74473674_8482** | 15_74471311_74473674 | SEMA7A | 8482 | 5.40247804564705 | -0.84246889 | 7 | only_DIvsD7 | DownUp | notDM | Loss | 7 |
| **15_75073359_75074306_192683** | 15_75073359_75074306 | SCAMP5 | 192683 | 5.3296703721267 | -0.91656805 | 2 | only_H3vsUI | DownUp | notDM | Gain | 2 |
| **15_75074927_75077108_192683** | 15_75074927_75077108 | SCAMP5 | 192683 | 5.32393652859314 | -0.75232819 | 1 | only_H3vsUI | DownUp | notDM | Loss | 7 |
| **15_75074927_75077108_57184** | 15_75074927_75077108 | FAM219B | 57184 | 5.08661250230788 | -0.68256653 | 1 | only_DIvsD7 | DownUp | notDM | Loss | 7 |
| **15_75163594_75165408_57184** | 15_75163594_75165408 | FAM219B | 57184 | 4.5285439208087 | -0.52966949 | 1 | only_DIvsD7 | DownUp | notDM | Loss | 7 |
| **15_75163594_75165408_8482** | 15_75163594_75165408 | SEMA7A | 8482 | 5.64077649795333 | 0.81125463 | 1 | only_DIvsD7 | DownUp | notDM | Loss | 6 |
| **15_75165842_75166865_57184** | 15_75165842_75166865 | FAM219B | 57184 | 4.50939072064236 | -0.52166615 | 4 | only_DIvsD7 | DownUp | notDM | Loss | 7 |
| **15_75181681_75184253_57184** | 15_75181681_75184253 | FAM219B | 57184 | 4.17394343728438 | -0.69990505 | 1 | only_DIvsD7 | DownUp | notDM | Loss | 7 |
| **15_75197066_75199000_57184** | 15_75197066_75199000 | FAM219B | 57184 | 2.25285303097989 | -0.41781728 | 1 | only_DIvsD7 | DownUp | notDM | Loss | 8 |
| **15_75199264_75200168_57184** | 15_75199264_75200168 | FAM219B | 57184 | 1.94448267215017 | -0.53649916 | 1 | only_DIvsD7 | DownUp | notDM | Loss | 7 |
| **15_75227007_75230138_57184** | 15_75227007_75230138 | FAM219B | 57184 | 4.44452881131714 | -0.8753536 | 1 | only_DIvsD7 | DownUp | notDM | Gain | 2 |
| **15_75287326_75287742_192683** | 15_75287326_75287742 | SCAMP5 | 192683 | 2.29666519026153 | -0.35369051 | 4 | only_H3vsUI | DownUp | notDM | Loss | 3 |
| **15_75287326_75287742_57184** | 15_75287326_75287742 | FAM219B | 57184 | 4.94522231663534 | -0.67242231 | 4 | only_DIvsD7 | DownUp | notDM | Loss | 7 |
| **15_75287988_75289095_192683** | 15_75287988_75289095 | SCAMP5 | 192683 | 1.70757017609794 | 0.83466439 | 1 | only_H3vsUI | DownUp | notDM | Loss | 3 |
| **15_75316136_75317846_192683** | 15_75316136_75317846 | SCAMP5 | 192683 | 4.45023370753275 | -0.86056398 | 1 | only_H3vsUI | DownUp | notDM | Loss | 7 |
| **15_75321507_75323143_192683** | 15_75321507_75323143 | SCAMP5 | 192683 | 4.52595134124801 | -0.76401667 | 8 | only_H3vsUI | DownUp | notDM | Loss | 7 |
| **15_75492648_75494103_192683** | 15_75492648_75494103 | SCAMP5 | 192683 | 5.31114117979464 | -0.86601649 | 2 | only_H3vsUI | DownUp | notDM | Loss | 7 |
| **15_75494598_75498264_192683** | 15_75494598_75498264 | SCAMP5 | 192683 | 5.31525852654083 | -0.90728571 | 1 | only_H3vsUI | DownUp | notDM | Loss | 7 |
| **15_76010355_76011314_161753** | 15_76010355_76011314 | ODF3L1 | 161753 | 3.69940408181534 | 0.00651282 | 6 | only_H3vsUI | DownUp | notDM | Loss | 3 |
| **15_76012899_76013886_161753** | 15_76012899_76013886 | ODF3L1 | 161753 | 3.38614210893082 | 0.00723079 | 6 | only_H3vsUI | DownUp | notDM | Gain | 1 |
| **15_81615254_81616547_80765** | 15_81615254_81616547 | STARD5 | 80765 | 0 | 0.43361334 | 1 | only_DIvsD7 | Up | notDM | Loss | 3 |
| **15_81616689_81617687_80765** | 15_81616689_81617687 | STARD5 | 80765 | 2.32014628611105 | 0.30235859 | 1 | only_DIvsD7 | Up | notDM | Loss | 3 |
| **15_82336504_82338349_84206** | 15_82336504_82338349 | MEX3B | 84206 | 2.12710479836481 | 0.09057832 | 1 | notSpecific | DownUp | DM | Loss | 10 |
| **15_88797286_88799384_4916** | 15_88797286_88799384 | NTRK3 | 4916 | 0.845098040014257 | -0.00254783 | 2 | notSpecific | Up | DM | Loss | 10 |

| | | | | | | | | | | | |
|---|---|---|---|---|---|---|---|---|---|---|---|
| **15_90691696_90693622_10519** | 15_90691696_90693622 | CIB1 | 10519 | 4.92250747775183 | 0.76037645 | 4 | only_DIvsD7 | DownUp | notDM | Loss | 5 |
| **15_90732933_90736919_10519** | 15_90732933_90736919 | CIB1 | 10519 | 4.60596191794896 | 0.44057678 | 7 | only_DIvsD7 | DownUp | notDM | Loss | 5 |
| **15_90743302_90747556_10519** | 15_90743302_90747556 | CIB1 | 10519 | 4.47310725263346 | -0.49125249 | 1 | only_DIvsD7 | DownUp | notDM | Loss | 7 |
| **15_90752805_90757357_10519** | 15_90752805_90757357 | CIB1 | 10519 | 4.29935473495851 | -0.27658404 | 7 | only_DIvsD7 | DownUp | notDM | Loss | 5 |
| **15_90764019_90764566_10519** | 15_90764019_90764566 | CIB1 | 10519 | 4.10428220709444 | -0.11309094 | 7 | only_DIvsD7 | DownUp | notDM | Loss | 7 |
| **15_90774273_90777139_10519** | 15_90774273_90777139 | CIB1 | 10519 | 2.14921911265538 | 0.47248096 | 1 | only_DIvsD7 | DownUp | notDM | Loss | 6 |
| **15_90777521_90778743_10519** | 15_90777521_90778743 | CIB1 | 10519 | 2.38738982633873 | 0.43714324 | 2 | only_DIvsD7 | DownUp | notDM | Loss | 6 |
| **15_90791993_90794040_10519** | 15_90791993_90794040 | CIB1 | 10519 | 4.16778977915909 | 0.54388498 | 1 | only_DIvsD7 | DownUp | notDM | Gain | 5 |
| **15_90961809_90965763_10519** | 15_90961809_90965763 | CIB1 | 10519 | 5.26607168875179 | -0.80315279 | 8 | only_DIvsD7 | DownUp | notDM | Loss | 7 |
| **15_91135089_91136462_91433** | 15_91135089_91136462 | RCCD1 | 91433 | 5.55828246504375 | 0.87044778 | 3 | only_DIvsD7 | UpDown | notDM | Loss | 5 |
| **15_91471864_91475762_91433** | 15_91471864_91475762 | RCCD1 | 91433 | 4.34918035899638 | 0.34186073 | 1 | only_DIvsD7 | UpDown | notDM | Loss | 6 |
| **15_91478520_91480244_91433** | 15_91478520_91480244 | RCCD1 | 91433 | 4.2519543982274 | 0.27411008 | 1 | only_DIvsD7 | UpDown | notDM | Gain | 4 |
| **15_91492415_91493288_91433** | 15_91492415_91493288 | RCCD1 | 91433 | 3.68295692630121 | 0.0277944 | 7 | only_DIvsD7 | UpDown | notDM | Loss | 1 |
| **15_91497502_91500598_91433** | 15_91497502_91500598 | RCCD1 | 91433 | 0 | 0.05836244 | 1 | only_DIvsD7 | UpDown | notDM | Loss | 8 |
| **15_91642861_91644391_374659** | 15_91642861_91644391 | HDDC3 | 374659 | 5.22294266145597 | -0.74121646 | 4 | notSpecific | Down | DM | Loss | 10 |
| **15_91642861_91644391_9899** | 15_91642861_91644391 | SV2B | 9899 | 0 | 0.00449517 | 4 | notSpecific | DownUp | DM | Loss | 10 |
| **15_92936488_92937076_8128** | 15_92936488_92937076 | ST8SIA2 | 8128 | 0 | 0.04456607 | 4 | only_H3vsUI | DownUp | notDM | Gain | 1 |
| **15_92937170_92938477_8128** | 15_92937170_92938477 | ST8SIA2 | 8128 | 2.05690485133647 | 0.01425169 | 4 | only_H3vsUI | DownUp | notDM | Loss | 3 |
| **15_96874711_96875288_7026** | 15_96874711_96875288 | NR2F2 | 7026 | 2.88479536394898 | 0.68574844 | 1 | notSpecific | Down | DM | Loss | 10 |
| **16_10480118_10481609_80063** | 16_10480118_10481609 | ATF7IP2 | 80063 | 2.31806333496276 | 0.78353736 | 1 | only_H3vsUI | DownUp | notDM | Gain | 4 |
| **16_10739812_10741382_80063** | 16_10739812_10741382 | ATF7IP2 | 80063 | 5.41479959542523 | 0.74210127 | 6 | only_H3vsUI | DownUp | notDM | Loss | 1 |
| **16_10741582_10742236_80063** | 16_10741582_10742236 | ATF7IP2 | 80063 | 5.41775725379743 | 0.75774535 | 6 | only_H3vsUI | DownUp | notDM | Loss | 3 |
| **16_11033462_11036049_28955** | 16_11033462_11036049 | DEXI | 28955 | 2.42975228000241 | 0.32786457 | 1 | only_H3vsUI | UpDown | notDM | Loss | 6 |
| **16_11036498_11038242_28955** | 16_11036498_11038242 | DEXI | 28955 | 2.26245108973043 | -0.2183148 | 1 | only_H3vsUI | UpDown | notDM | Loss | 8 |
| **16_11110909_11111648_28955** | 16_11110909_11111648 | DEXI | 28955 | 4.87270389621682 | 0.57775911 | 6 | only_H3vsUI | UpDown | notDM | Gain | 1 |
| **16_1127895_1129283_6755** | 16_1127895_1129283 | SSTR5 | 6755 | 0 | 0.10416494 | 6 | only_H3vsUI | UpDown | notDM | Gain | 4 |
| **16_1138261_1141658_6755** | 16_1138261_1141658 | SSTR5 | 6755 | 3.97689995098294 | 0.10049624 | 5 | only_H3vsUI | UpDown | notDM | Loss | 6 |
| **16_11589873_11592812_400499** | 16_11589873_11592812 | LOC400499 | 400499 | 3.62981719601852 | 0 | 7 | only_H3vsUI | UpDown | notDM | Gain | 1 |
| **16_12005654_12008964_2935** | 16_12005654_12008964 | GSPT1 | 2935 | 2.93550726582471 | 0.31181417 | 1 | notSpecific | DownUp | DM | Loss | 10 |
| **16_12996127_12997447_729993** | 16_12996127_12997447 | SHISA9 | 729993 | 2.82865989653532 | 0.38631275 | 4 | only_H3vsUI | UpDown | notDM | Loss | 3 |
| **16_19117944_19120269_162073** | 16_19117944_19120269 | ITPRIPL2 | 162073 | 3.69775227416775 | 0.20008951 | 3 | only_DIvsD7 | UpDown | notDM | Gain | 1 |
| **16_19121781_19124685_162073** | 16_19121781_19124685 | ITPRIPL2 | 162073 | 2.75587485567249 | 0.2464187 | 4 | only_DIvsD7 | UpDown | notDM | Gain | 4 |
| **16_19125632_19131820_162073** | 16_19125632_19131820 | ITPRIPL2 | 162073 | 2.57978359661681 | 0.08502228 | 5 | only_DIvsD7 | UpDown | notDM | Loss | 4 |
| **16_19528151_19529820_9738** | 16_19528151_19529820 | CCP110 | 9738 | 3.73150818359603 | -0.19295366 | 8 | only_DIvsD7 | UpDown | notDM | Gain | 1 |

| | | | | | | | | | | | |
|---|---|---|---|---|---|---|---|---|---|---|---|
| **16_19530790_19533186_9738** | 16_19530790_19533186 | CCP110 | 9738 | 3.3059958827708 | 0.20260548 | 1 | only_DIvsD7 | UpDown | notDM | Loss | 6 |
| **16_19533496_19534273_9738** | 16_19533496_19534273 | CCP110 | 9738 | 2.97127584873811 | 0.20260548 | 4 | only_DIvsD7 | UpDown | notDM | Gain | 4 |
| **16_19535382_19537472_9738** | 16_19535382_19537472 | CCP110 | 9738 | 2.02118929906994 | -0.1993675 | 1 | only_DIvsD7 | UpDown | notDM | Loss | 8 |
| **16_1978620_1979742_254528** | 16_1978620_1979742 | MEIOB | 254528 | 4.75224825417777 | 0.99738674 | 1 | notSpecific | Down | DM | Loss | 10 |
| **16_1978620_1979742_4716** | 16_1978620_1979742 | NDUFB10 | 4716 | 4.47389552558097 | 0.37836303 | 1 | notSpecific | DownUp | DM | Loss | 10 |
| **16_1978620_1979742_51734** | 16_1978620_1979742 | MSRB1 | 51734 | 4.12862540487595 | 0.524882 | 1 | notSpecific | DownUp | DM | Loss | 10 |
| **16_1978620_1979742_7249** | 16_1978620_1979742 | TSC2 | 7249 | 5.07276442105075 | -0.6805437 | 1 | notSpecific | Down | DM | Loss | 10 |
| **16_1978620_1979742_81889** | 16_1978620_1979742 | FAHD1 | 81889 | 5.00602510585807 | -0.72793744 | 1 | notSpecific | Up | DM | Loss | 10 |
| **16_1978620_1979742_84231** | 16_1978620_1979742 | TRAF7 | 84231 | 5.35415647791543 | -0.79329625 | 1 | notSpecific | Up | DM | Loss | 10 |
| **16_2009663_2011541_9143** | 16_2009663_2011541 | SYNGR3 | 9143 | 4.45423489574827 | -0.39282332 | 2 | only_H3vsUI | UpDown | notDM | Loss | 8 |
| **16_2011767_2014680_9143** | 16_2011767_2014680 | SYNGR3 | 9143 | 4.40255368035325 | -0.33470792 | 1 | only_H3vsUI | UpDown | notDM | Loss | 8 |
| **16_2015183_2016717_9143** | 16_2015183_2016717 | SYNGR3 | 9143 | 4.36604920980024 | -0.44295949 | 1 | only_H3vsUI | UpDown | notDM | Loss | 7 |
| **16_2020592_2021917_9143** | 16_2020592_2021917 | SYNGR3 | 9143 | 4.2559957267224 | -0.28826758 | 1 | only_H3vsUI | UpDown | notDM | Loss | 7 |
| **16_2022242_2023808_9143** | 16_2022242_2023808 | SYNGR3 | 9143 | 4.20787662159197 | -0.28826758 | 1 | only_H3vsUI | UpDown | notDM | Loss | 7 |
| **16_2031176_2033946_9143** | 16_2031176_2033946 | SYNGR3 | 9143 | 3.78211414747907 | 0.27870921 | 1 | only_H3vsUI | UpDown | notDM | Loss | 6 |
| **16_2034558_2036338_9143** | 16_2034558_2036338 | SYNGR3 | 9143 | 3.55738688205951 | 0.03927737 | 1 | only_H3vsUI | UpDown | notDM | Loss | 6 |
| **16_2037867_2039843_9143** | 16_2037867_2039843 | SYNGR3 | 9143 | 2.01703333929878 | 0.06729653 | 4 | only_H3vsUI | UpDown | notDM | Loss | 6 |
| **16_2040349_2041261_9143** | 16_2040349_2041261 | SYNGR3 | 9143 | 2.54530711646582 | 0.38001672 | 2 | only_H3vsUI | UpDown | notDM | Loss | 3 |
| **16_2045168_2046666_9143** | 16_2045168_2046666 | SYNGR3 | 9143 | 3.71800316826702 | 0.36263777 | 7 | only_H3vsUI | UpDown | notDM | Gain | 1 |
| **16_2054108_2055155_9143** | 16_2054108_2055155 | SYNGR3 | 9143 | 4.14952701375435 | -0.23711727 | 8 | only_H3vsUI | UpDown | notDM | Loss | 3 |
| **16_2056852_2059452_9143** | 16_2056852_2059452 | SYNGR3 | 9143 | 4.22809223915698 | -0.21800777 | 1 | only_H3vsUI | UpDown | notDM | Loss | 8 |
| **16_2059923_2061289_9143** | 16_2059923_2061289 | SYNGR3 | 9143 | 4.30057374688543 | -0.30962073 | 1 | only_H3vsUI | UpDown | notDM | Loss | 7 |
| **16_21168270_21168565_55567** | 16_21168270_21168565 | DNAH3 | 55567 | 3.34202768808747 | 0 | 6 | only_H3vsUI | UpDown | notDM | Gain | 1 |
| **16_21168844_21171168_55567** | 16_21168844_21171168 | DNAH3 | 55567 | 0 | 0 | 2 | only_H3vsUI | UpDown | notDM | Gain | 4 |
| **16_21516409_21531356_100271836** | 16_21516409_21531356 | SMG1P3 | 100271836 | 2.16136800223497 | 0.05529653 | 1 | notSpecific | Up | DM | Loss | 10 |
| **16_23765656_23766644_63928** | 16_23765656_23766644 | CHP2 | 63928 | 0 | 0.00505583 | 4 | notSpecific | Up | DM | Loss | 10 |
| **16_2517994_2518745_4917** | 16_2517994_2518745 | NTN3 | 4917 | 3.44027921323559 | 0.00311195 | 4 | only_H3vsUI | UpDown | notDM | Loss | 3 |
| **16_2523024_2523858_4917** | 16_2523024_2523858 | NTN3 | 4917 | 3.18355453361886 | 0.00331834 | 2 | only_H3vsUI | UpDown | notDM | Loss | 3 |
| **16_2524097_2524944_4917** | 16_2524097_2524944 | NTN3 | 4917 | 3.41480627950101 | 0.00106863 | 1 | only_H3vsUI | UpDown | notDM | Loss | 3 |
| **16_2525326_2526551_4917** | 16_2525326_2526551 | NTN3 | 4917 | 3.58297192910481 | 0.00153239 | 2 | only_H3vsUI | UpDown | notDM | Loss | 3 |
| **16_2564595_2567510_51005** | 16_2564595_2567510 | AMDHD2 | 51005 | 3.45591024038274 | -0.29876611 | 1 | only_H3vsUI | Down | notDM | Loss | 8 |
| **16_2570720_2571726_51005** | 16_2570720_2571726 | AMDHD2 | 51005 | 2.55144999797288 | -0.32869986 | 2 | only_H3vsUI | Down | notDM | Loss | 8 |
| **16_2581431_2581975_51005** | 16_2581431_2581975 | AMDHD2 | 51005 | 4.04402990994647 | 0.33004849 | 4 | only_H3vsUI | Down | notDM | Loss | 2 |
| **16_2582180_2583738_51005** | 16_2582180_2583738 | AMDHD2 | 51005 | 4.07258073264895 | 0.43315294 | 2 | only_H3vsUI | Down | notDM | Gain | 1 |

| | | | | | | | | | | | |
|---|---|---|---|---|---|---|---|---|---|---|---|
| **16_2586285_2587578_9088** | 16_2586285_2587578 | PKMYT1 | 9088 | 5.64633313537874 | -0.82563394 | 2 | only_H3vsUI | DownUp | notDM | Loss | 7 |
| **16_2588413_2589815_51005** | 16_2588413_2589815 | AMDHD2 | 51005 | 4.25645314493904 | -0.47154487 | 2 | only_H3vsUI | Down | notDM | Gain | 4 |
| **16_27324143_27325026_3566** | 16_27324143_27325026 | IL4R | 3566 | 2.36172783601759 | 0.6624238 | 2 | only_DIvsD7 | UpDown | notDM | Loss | 3 |
| **16_27325447_27328226_3566** | 16_27325447_27328226 | IL4R | 3566 | 2.30535136944662 | 0.61621556 | 1 | only_DIvsD7 | UpDown | notDM | Loss | 6 |
| **16_27333951_27335027_3566** | 16_27333951_27335027 | IL4R | 3566 | 3.93981866282138 | 0.16871 | 8 | only_DIvsD7 | UpDown | notDM | Gain | 1 |
| **16_27337399_27339973_3566** | 16_27337399_27339973 | IL4R | 3566 | 4.08443327678654 | 0.79417064 | 7 | only_DIvsD7 | UpDown | notDM | Gain | 5 |
| **16_27374721_27377672_3566** | 16_27374721_27377672 | IL4R | 3566 | 4.69432435211631 | 0.60138369 | 7 | only_DIvsD7 | UpDown | notDM | Gain | 5 |
| **16_27558394_27560846_23247** | 16_27558394_27560846 | KATNIP | 23247 | 2.79448804665917 | -0.63885244 | 1 | only_H3vsUI | DownUp | notDM | Gain | 4 |
| **16_27561628_27562882_23247** | 16_27561628_27562882 | KATNIP | 23247 | 2.20951501454263 | -0.79783125 | 1 | only_H3vsUI | DownUp | notDM | Gain | 4 |
| **16_27577916_27580815_23247** | 16_27577916_27580815 | KATNIP | 23247 | 4.21616590228599 | 0.30942555 | 7 | only_H3vsUI | DownUp | notDM | Gain | 5 |
| **16_27577916_27580815_3566** | 16_27577916_27580815 | IL4R | 3566 | 5.40255539917077 | -0.79983923 | 7 | only_DIvsD7 | UpDown | notDM | Gain | 5 |
| **16_27589257_27591436_23247** | 16_27589257_27591436 | KATNIP | 23247 | 4.44390417422135 | -0.45701043 | 3 | only_H3vsUI | DownUp | notDM | Loss | 7 |
| **16_2769603_2771010_9088** | 16_2769603_2771010 | PKMYT1 | 9088 | 5.41398336535747 | 0.7826605 | 1 | only_H3vsUI | DownUp | notDM | Loss | 3 |
| **16_2771543_2772044_9088** | 16_2771543_2772044 | PKMYT1 | 9088 | 5.41224880876327 | 0.7430643 | 5 | only_H3vsUI | DownUp | notDM | Gain | 2 |
| **16_279474_280146_3039** | 16_279474_280146 | HBA1 | 3039 | 4.72260924600993 | -0.77624319 | 2 | notSpecific | Down | DM | Gain | 9 |
| **16_279474_280146_3049** | 16_279474_280146 | HBQ1 | 3049 | 4.69040874361649 | -0.74417185 | 2 | notSpecific | Up | DM | Gain | 9 |
| **16_279474_280146_55692** | 16_279474_280146 | LUC7L | 55692 | 1.14612803567824 | -0.30368703 | 2 | notSpecific | Down | DM | Gain | 9 |
| **16_279474_280146_79622** | 16_279474_280146 | SNRNP25 | 79622 | 5.24464073656714 | -0.74080716 | 2 | notSpecific | Up | DM | Gain | 9 |
| **16_279474_280146_83986** | 16_279474_280146 | FAM234A | 83986 | 3.66950283410434 | -0.21571205 | 2 | notSpecific | Down | DM | Gain | 9 |
| **16_279474_280146_9727** | 16_279474_280146 | RAB11FIP3 | 9727 | 5.29108900005696 | 0.7519736 | 2 | notSpecific | DownUp | DM | Gain | 9 |
| **16_285006_286419_65990** | 16_285006_286419 | ANTKMT | 65990 | 5.68550885851482 | 0.80850966 | 1 | only_H3vsUI | Down | notDM | Loss | 7 |
| **16_28503716_28504137_1201** | 16_28503716_28504137 | CLN3 | 1201 | 2.57287160220048 | 0.61249619 | 3 | notSpecific | UpDown | DM | Gain | 9 |
| **16_28503716_28504137_55911** | 16_28503716_28504137 | APOBR | 55911 | 3.26881190373978 | -0.57286975 | 3 | notSpecific | DownUp | DM | Gain | 9 |
| **16_28962624_28964123_11273** | 16_28962624_28964123 | ATXN2L | 11273 | 5.10792867486992 | 0.67455609 | 1 | notSpecific | DownUp | DM | Loss | 10 |
| **16_28962624_28964123_6799** | 16_28962624_28964123 | SULT1A2 | 6799 | 5.54931966644702 | 0.84638923 | 1 | notSpecific | Up | DM | Loss | 10 |
| **16_28962624_28964123_83985** | 16_28962624_28964123 | SPNS1 | 83985 | 4.34190912017977 | 0.318935 | 1 | notSpecific | Up | DM | Loss | 10 |
| **16_28962624_28964123_84901** | 16_28962624_28964123 | NFATC2IP | 84901 | 2.49276038902684 | 0.41265624 | 1 | notSpecific | Down | DM | Loss | 10 |
| **16_2917364_2918273_9088** | 16_2917364_2918273 | PKMYT1 | 9088 | 5.04977991527238 | 0.84350102 | 1 | only_H3vsUI | DownUp | notDM | Gain | 1 |
| **16_2940109_2941403_9088** | 16_2940109_2941403 | PKMYT1 | 9088 | 4.94946319617663 | 0.79239738 | 7 | only_H3vsUI | DownUp | notDM | Gain | 1 |
| **16_2945593_2946445_9088** | 16_2945593_2946445 | PKMYT1 | 9088 | 4.92459455212075 | 0.76265135 | 3 | only_H3vsUI | DownUp | notDM | Loss | 5 |
| **16_2954428_2955351_9088** | 16_2954428_2955351 | PKMYT1 | 9088 | 4.87595787913683 | 0.81491868 | 1 | only_H3vsUI | DownUp | notDM | Loss | 3 |
| **16_2961073_2961756_9088** | 16_2961073_2961756 | PKMYT1 | 9088 | 4.83671644948754 | 0.63862707 | 1 | only_H3vsUI | DownUp | notDM | Loss | 3 |
| **16_29872464_29874887_440356** | 16_29872464_29874887 | CDIPTOSP | 440356 | 2.42975228000241 | 0 | 5 | only_H3vsUI | UpDown | notDM | Loss | 6 |
| **16_29872464_29874887_79153** | 16_29872464_29874887 | GDPD3 | 79153 | 5.39789136496597 | 0.93161528 | 5 | only_H3vsUI | DownUp | notDM | Loss | 6 |

| | | | | | | | | | | | |
|---|---|---|---|---|---|---|---|---|---|---|---|
| **16_29909671_29911577_79153** | 16_29909671_29911577 | GDPD3 | 79153 | 5.32895420457959 | 0.88188545 | 2 | only_H3vsUI | DownUp | notDM | Gain | 1 |
| **16_29911791_29913556_79153** | 16_29911791_29913556 | GDPD3 | 79153 | 5.32490566304473 | 0.87846453 | 1 | only_H3vsUI | DownUp | notDM | Loss | 6 |
| **16_29955592_29957698_79153** | 16_29955592_29957698 | GDPD3 | 79153 | 5.2231349605404 | 0.74014325 | 7 | only_H3vsUI | DownUp | notDM | Gain | 5 |
| **16_29972357_29973219_79153** | 16_29972357_29973219 | GDPD3 | 79153 | 5.1808137757544 | 0.76914879 | 1 | only_H3vsUI | DownUp | notDM | Gain | 1 |
| **16_29973511_29974621_79153** | 16_29973511_29974621 | GDPD3 | 79153 | 5.17677979354061 | 0.76914879 | 1 | only_H3vsUI | DownUp | notDM | Loss | 6 |
| **16_30086499_30087248_79153** | 16_30086499_30087248 | GDPD3 | 79153 | 4.57531488058295 | -0.69974907 | 1 | only_H3vsUI | DownUp | notDM | Gain | 2 |
| **16_30087685_30089585_79153** | 16_30087685_30089585 | GDPD3 | 79153 | 4.54745471057273 | -0.7000476 | 2 | only_H3vsUI | DownUp | notDM | Loss | 7 |
| **16_30103391_30104303_79153** | 16_30103391_30104303 | GDPD3 | 79153 | 4.31293860901313 | 0.24531237 | 1 | only_H3vsUI | DownUp | notDM | Loss | 3 |
| **16_30106664_30107289_79153** | 16_30106664_30107289 | GDPD3 | 79153 | 4.2447717614953 | 0.20202924 | 2 | only_H3vsUI | DownUp | notDM | Loss | 3 |
| **16_3013054_3015089_9088** | 16_3013054_3015089 | PKMYT1 | 9088 | 4.18551382421854 | 0.60385329 | 1 | only_H3vsUI | DownUp | notDM | Loss | 6 |
| **16_30133511_30134356_79153** | 16_30133511_30134356 | GDPD3 | 79153 | 3.93726707221141 | 0.05341182 | 1 | only_H3vsUI | DownUp | notDM | Loss | 3 |
| **16_3018395_3019175_9088** | 16_3018395_3019175 | PKMYT1 | 9088 | 4.05088221066403 | 0.37877409 | 2 | only_H3vsUI | DownUp | notDM | Loss | 3 |
| **16_3019583_3021423_9088** | 16_3019583_3021423 | PKMYT1 | 9088 | 3.95822931418838 | -0.34338629 | 1 | only_H3vsUI | DownUp | notDM | Loss | 7 |
| **16_3028931_3030317_9088** | 16_3028931_3030317 | PKMYT1 | 9088 | 2.27646180417324 | -0.38423216 | 1 | only_H3vsUI | DownUp | notDM | Loss | 8 |
| **16_3030634_3031821_9088** | 16_3030634_3031821 | PKMYT1 | 9088 | 2.11727129565576 | -0.38423216 | 1 | only_H3vsUI | DownUp | notDM | Loss | 8 |
| **16_30362995_30366372_79153** | 16_30362995_30366372 | GDPD3 | 79153 | 5.37683052559301 | 0.74475247 | 1 | only_H3vsUI | DownUp | notDM | Loss | 6 |
| **16_30366727_30368767_79153** | 16_30366727_30368767 | GDPD3 | 79153 | 5.38358380017263 | 0.74475247 | 1 | only_H3vsUI | DownUp | notDM | Loss | 6 |
| **16_30417939_30418860_51333** | 16_30417939_30418860 | ZNF771 | 51333 | 0 | 0 | 1 | only_H3vsUI | UpDown | notDM | Loss | 3 |
| **16_30419395_30420898_51333** | 16_30419395_30420898 | ZNF771 | 51333 | 2.8208579894397 | 0 | 1 | only_H3vsUI | UpDown | notDM | Loss | 3 |
| **16_3054866_3055704_9088** | 16_3054866_3055704 | PKMYT1 | 9088 | 4.38673076520983 | 0.38686566 | 4 | only_H3vsUI | DownUp | notDM | Gain | 4 |
| **16_30750782_30753922_5261** | 16_30750782_30753922 | PHKG2 | 5261 | 3.75358305889291 | -0.01524902 | 7 | only_DIvsD7 | Up | notDM | Loss | 6 |
| **16_30758108_30759483_5261** | 16_30758108_30759483 | PHKG2 | 5261 | 2.03742649794062 | 0.04618201 | 5 | only_DIvsD7 | Up | notDM | Gain | 4 |
| **16_30904267_30905225_1489** | 16_30904267_30905225 | CTF1 | 1489 | 3.4320066872696 | 0.13391277 | 1 | only_H3vsUI | UpDown | notDM | Loss | 3 |
| **16_30905639_30907206_1489** | 16_30905639_30907206 | CTF1 | 1489 | 2.85913829729453 | 0.23776128 | 4 | only_H3vsUI | UpDown | notDM | Loss | 3 |
| **16_30935445_30936617_29108** | 16_30935445_30936617 | PYCARD | 29108 | 5.44309862872004 | -0.79610389 | 1 | notSpecific | Down | DM | Loss | 10 |
| **16_30935445_30936617_54620** | 16_30935445_30936617 | FBXL19 | 54620 | 1.46239799789896 | -0.04514348 | 1 | notSpecific | Up | DM | Loss | 10 |
| **16_30935445_30936617_84148** | 16_30935445_30936617 | KAT8 | 84148 | 5.28413057213467 | 0.82334888 | 1 | notSpecific | Up | DM | Loss | 10 |
| **16_30935445_30936617_93129** | 16_30935445_30936617 | ORAI3 | 93129 | 4.37604745204147 | 0.29555178 | 1 | notSpecific | DownUp | DM | Loss | 10 |
| **16_3095015_3095628_9088** | 16_3095015_3095628 | PKMYT1 | 9088 | 4.80964050609339 | 0.61657787 | 2 | only_H3vsUI | DownUp | notDM | Loss | 3 |
| **16_3095743_3096808_9088** | 16_3095743_3096808 | PKMYT1 | 9088 | 4.81451395236824 | 0.59283614 | 2 | only_H3vsUI | DownUp | notDM | Loss | 3 |
| **16_31226382_31227924_493829** | 16_31226382_31227924 | TRIM72 | 493829 | 3.01786771896351 | 0.05025415 | 2 | only_DIvsD7 | DownUp | notDM | Loss | 3 |
| **16_31498661_31499825_6524** | 16_31498661_31499825 | SLC5A2 | 6524 | 3.62572390952576 | 0 | 1 | only_H3vsUI | Up | notDM | Loss | 3 |
| **16_3178819_3180286_9088** | 16_3178819_3180286 | PKMYT1 | 9088 | 5.17144560683597 | 0.71550661 | 4 | only_H3vsUI | DownUp | notDM | Loss | 3 |
| **16_3183001_3184610_9088** | 16_3183001_3184610 | PKMYT1 | 9088 | 5.18326414798012 | 0.85913234 | 1 | only_H3vsUI | DownUp | notDM | Gain | 4 |

| | | | | | | | | | | | |
|---|---|---|---|---|---|---|---|---|---|---|---|
| **16_3185296_3186560_9088** | 16_3185296_3186560 | PKMYT1 | 9088 | 5.18975131726347 | 0.83281922 | 2 | only_H3vsUI | DownUp | notDM | Loss | 6 |
| **16_3201429_3202891_9088** | 16_3201429_3202891 | PKMYT1 | 9088 | 5.23288688810502 | 0.7423879 | 1 | only_H3vsUI | DownUp | notDM | Gain | 1 |
| **16_3222896_3226212_9088** | 16_3222896_3226212 | PKMYT1 | 9088 | 5.28438786629956 | 0.70218895 | 2 | only_H3vsUI | DownUp | notDM | Loss | 6 |
| **16_3283503_3284951_7752** | 16_3283503_3284951 | ZNF200 | 7752 | 2.35218251811136 | 0.17413829 | 2 | only_H3vsUI | Down | notDM | Loss | 8 |
| **16_3660267_3661475_84464** | 16_3660267_3661475 | SLX4 | 84464 | 2.09691001300806 | -0.32122321 | 1 | notSpecific | Down | DM | Loss | 10 |
| **16_430303_431636_729480** | 16_430303_431636 | RPL23AP5 | 729480 | 2.66464197555613 | 0.04913755 | 1 | only_DIvsD7 | DownUp | notDM | Loss | 3 |
| **16_432189_433027_729480** | 16_432189_433027 | RPL23AP5 | 729480 | 1.61278385671974 | -0.09749373 | 4 | only_DIvsD7 | DownUp | notDM | Loss | 3 |
| **16_445798_447042_65990** | 16_445798_447042 | ANTKMT | 65990 | 5.51070181046007 | -0.77146167 | 5 | only_H3vsUI | Down | notDM | Loss | 2 |
| **16_447894_449072_65990** | 16_447894_449072 | ANTKMT | 65990 | 5.50797319628135 | -0.93329212 | 5 | only_H3vsUI | Down | notDM | Loss | 2 |
| **16_50058212_50058882_55027** | 16_50058212_50058882 | HEATR3 | 55027 | 4.61299565603235 | -0.70389659 | 2 | only_H3vsUI | Down | notDM | Gain | 2 |
| **16_50059477_50060967_55027** | 16_50059477_50060967 | HEATR3 | 55027 | 4.59029555935402 | -0.41589002 | 2 | only_H3vsUI | Down | notDM | Gain | 4 |
| **16_5008297_5009022_5493** | 16_5008297_5009022 | PPL | 5493 | 3.23578087032756 | 0 | 2 | notSpecific | Down | DM | Loss | 10 |
| **16_5008297_5009022_57407** | 16_5008297_5009022 | NMRAL1 | 57407 | 5.68463623978711 | 0.8109545 | 2 | notSpecific | DownUp | DM | Loss | 10 |
| **16_5008297_5009022_9717** | 16_5008297_5009022 | SEC14L5 | 9717 | 0 | 0.03611191 | 2 | notSpecific | UpDown | DM | Loss | 10 |
| **16_500966_502122_65990** | 16_500966_502122 | ANTKMT | 65990 | 5.4298120115668 | -0.76494027 | 3 | only_H3vsUI | Down | notDM | Gain | 2 |
| **16_50099034_50099740_55027** | 16_50099034_50099740 | HEATR3 | 55027 | 2.19865708695442 | -0.08496591 | 2 | only_H3vsUI | Down | notDM | Gain | 4 |
| **16_50100386_50102360_55027** | 16_50100386_50102360 | HEATR3 | 55027 | 2.69108149212297 | -0.03592111 | 1 | only_H3vsUI | Down | notDM | Gain | 4 |
| **16_56442048_56444301_4493** | 16_56442048_56444301 | MT1E | 4493 | 5.33328198946365 | -0.73965883 | 7 | only_DIvsD7 | UpDown | notDM | Gain | 1 |
| **16_56554076_56554932_9688** | 16_56554076_56554932 | NUP93 | 9688 | 5.32036649446046 | -0.72053694 | 1 | only_H3vsUI | DownUp | notDM | Loss | 7 |
| **16_56634127_56634823_4493** | 16_56634127_56634823 | MT1E | 4493 | 4.39612957530959 | -0.6537865 | 4 | only_DIvsD7 | UpDown | notDM | Gain | 1 |
| **16_56639629_56642355_4493** | 16_56639629_56642355 | MT1E | 4493 | 4.23964977716666 | 0.38161434 | 7 | only_DIvsD7 | UpDown | notDM | Gain | 1 |
| **16_56642547_56646114_4493** | 16_56642547_56646114 | MT1E | 4493 | 4.13369854611578 | 0.322717 | 1 | only_DIvsD7 | UpDown | notDM | Loss | 6 |
| **16_56660734_56661711_4493** | 16_56660734_56661711 | MT1E | 4493 | 3.00774777800074 | 0.76646731 | 3 | only_DIvsD7 | UpDown | notDM | Loss | 3 |
| **16_56676559_56678370_4493** | 16_56676559_56678370 | MT1E | 4493 | 4.2264194486486 | -0.45647232 | 4 | only_DIvsD7 | UpDown | notDM | Loss | 3 |
| **16_56762661_56763868_9688** | 16_56762661_56763868 | NUP93 | 9688 | 2.23044892137827 | -0.31034611 | 2 | only_H3vsUI | DownUp | notDM | Loss | 8 |
| **16_56764349_56765766_9688** | 16_56764349_56765766 | NUP93 | 9688 | 2.49692964807321 | -0.14804942 | 1 | only_H3vsUI | DownUp | notDM | Loss | 8 |
| **16_56844446_56850900_51090** | 16_56844446_56850900 | PLLP | 51090 | 5.6699673699085 | 0.91241031 | 7 | only_DIvsD7 | DownUp | notDM | Gain | 5 |
| **16_56844446_56850900_9688** | 16_56844446_56850900 | NUP93 | 9688 | 4.9053154630834 | -0.67101028 | 7 | only_H3vsUI | DownUp | notDM | Gain | 5 |
| **16_57022661_57024495_4493** | 16_57022661_57024495 | MT1E | 4493 | 5.55984081785637 | 0.86006734 | 2 | only_DIvsD7 | UpDown | notDM | Loss | 3 |
| **16_57293973_57295211_51090** | 16_57293973_57295211 | PLLP | 51090 | 4.36901165388415 | -0.48330111 | 6 | only_DIvsD7 | DownUp | notDM | Loss | 7 |
| **16_57310102_57312619_51090** | 16_57310102_57312619 | PLLP | 51090 | 3.77677380241211 | 0.85535273 | 6 | only_DIvsD7 | DownUp | notDM | Loss | 5 |
| **16_57332466_57333748_51090** | 16_57332466_57333748 | PLLP | 51090 | 4.14204514815774 | 0.12673735 | 1 | only_DIvsD7 | DownUp | notDM | Loss | 7 |
| **16_57334022_57334362_51090** | 16_57334022_57334362 | PLLP | 51090 | 4.18822517270528 | 0.12674449 | 1 | only_DIvsD7 | DownUp | notDM | Gain | 2 |
| **16_57373526_57373947_51090** | 16_57373526_57373947 | PLLP | 51090 | 4.73980169262349 | -0.55738446 | 6 | only_DIvsD7 | DownUp | notDM | Loss | 7 |

| | | | | | | | | | | | |
|---|---|---|---|---|---|---|---|---|---|---|---|
| **16_57505705_57508716_51090** | 16_57505705_57508716 | PLLP | 51090 | 5.27209235661588 | 0.82721597 | 7 | only_DIvsD7 | DownUp | notDM | Loss | 5 |
| **16_58035657_58036767_4324** | 16_58035657_58036767 | MMP15 | 4324 | 4.3561023781134 | 0.34026166 | 2 | only_H3vsUI | DownUp | notDM | Loss | 7 |
| **16_58058626_58062336_4324** | 16_58058626_58062336 | MMP15 | 4324 | 0 | 0.14181922 | 2 | only_H3vsUI | DownUp | notDM | Loss | 8 |
| **16_67192377_67193771_3299** | 16_67192377_67193771 | HSF4 | 3299 | 3.63083451782805 | 0.07363618 | 1 | only_H3vsUI | DownUp | notDM | Gain | 4 |
| **16_67193910_67195776_3299** | 16_67193910_67195776 | HSF4 | 3299 | 3.35583449588494 | 0.07557821 | 1 | only_H3vsUI | DownUp | notDM | Loss | 6 |
| **16_67203030_67204340_3299** | 16_67203030_67204340 | HSF4 | 3299 | 1.74036268949424 | 0.00745173 | 5 | only_H3vsUI | DownUp | notDM | Loss | 6 |
| **16_67204691_67206025_3299** | 16_67204691_67206025 | HSF4 | 3299 | 3.23451728351269 | 0.00741076 | 5 | only_H3vsUI | DownUp | notDM | Loss | 6 |
| **16_67206821_67209499_3299** | 16_67206821_67209499 | HSF4 | 3299 | 3.58500927990246 | -0.00826122 | 1 | only_H3vsUI | DownUp | notDM | Loss | 6 |
| **16_676126_678296_65990** | 16_676126_678296 | ANTKMT | 65990 | 4.9678427097243 | 0.71569286 | 3 | only_H3vsUI | Down | notDM | Loss | 5 |
| **16_68002669_68003418_123904** | 16_68002669_68003418 | NRN1L | 123904 | 4.92371019439656 | -0.83092823 | 2 | notSpecific | Up | DM | Loss | 10 |
| **16_68002669_68003418_6560** | 16_68002669_68003418 | SLC12A4 | 6560 | 2.16435285578444 | 0.39554607 | 2 | notSpecific | Up | DM | Loss | 10 |
| **16_69138931_69140103_54921** | 16_69138931_69140103 | CHTF8 | 54921 | 4.42117600185209 | 0.58991797 | 1 | only_H3vsUI | DownUp | notDM | Loss | 7 |
| **16_69140350_69142911_1001** | 16_69140350_69142911 | CDH3 | 1001 | 5.66384032944165 | 0.98471294 | 1 | notSpecific | Down | DM | Loss | 10 |
| **16_69140350_69142911_3038** | 16_69140350_69142911 | HAS3 | 3038 | 2.28330122870355 | 0.99134342 | 1 | notSpecific | Up | DM | Loss | 10 |
| **16_69140350_69142911_54921** | 16_69140350_69142911 | CHTF8 | 54921 | 4.36443842063993 | 0.59133499 | 1 | only_H3vsUI | DownUp | DM | Loss | 10 |
| **16_69140350_69142911_84916** | 16_69140350_69142911 | UTP4 | 84916 | 4.37333504995457 | 0.35256274 | 1 | notSpecific | Up | DM | Loss | 10 |
| **16_69140350_69142911_999** | 16_69140350_69142911 | CDH1 | 999 | 5.56714051528848 | 0.97999522 | 1 | notSpecific | UpDown | DM | Loss | 10 |
| **16_69163986_69165320_54921** | 16_69163986_69165320 | CHTF8 | 54921 | 2.86628733908419 | -0.50205306 | 2 | only_H3vsUI | DownUp | notDM | Gain | 2 |
| **16_69166832_69169095_54921** | 16_69166832_69169095 | CHTF8 | 54921 | 2.89209460269048 | -0.50205306 | 2 | only_H3vsUI | DownUp | notDM | Loss | 8 |
| **16_69206903_69208340_54921** | 16_69206903_69208340 | CHTF8 | 54921 | 4.61125584484748 | 0.50669237 | 3 | only_H3vsUI | DownUp | notDM | Gain | 5 |
| **16_69213015_69214001_54921** | 16_69213015_69214001 | CHTF8 | 54921 | 4.66773305253327 | 0.45669668 | 3 | only_H3vsUI | DownUp | notDM | Loss | 5 |
| **16_69219985_69220925_54921** | 16_69219985_69220925 | CHTF8 | 54921 | 4.7318545784044 | -0.62061884 | 2 | only_H3vsUI | DownUp | notDM | Loss | 7 |
| **16_69221763_69225315_54921** | 16_69221763_69225315 | CHTF8 | 54921 | 4.7459409540183 | -0.66367364 | 1 | only_H3vsUI | DownUp | notDM | Loss | 7 |
| **16_69249073_69253906_54921** | 16_69249073_69253906 | CHTF8 | 54921 | 4.91921411512699 | 0.7332459 | 6 | only_H3vsUI | DownUp | notDM | Gain | 5 |
| **16_69344336_69344984_54921** | 16_69344336_69344984 | CHTF8 | 54921 | 5.25111236940359 | -0.86199663 | 1 | only_H3vsUI | DownUp | notDM | Loss | 7 |
| **16_69373888_69376181_146456** | 16_69373888_69376181 | TMED6 | 146456 | 3.97918403367442 | 0.25710206 | 2 | notSpecific | Up | DM | Loss | 10 |
| **16_69373888_69376181_51388** | 16_69373888_69376181 | NIP7 | 51388 | 2.27875360095283 | -0.27315699 | 2 | notSpecific | Up | DM | Loss | 10 |
| **16_69373888_69376181_80777** | 16_69373888_69376181 | CYB5B | 80777 | 4.91512534614147 | 0.83007009 | 2 | notSpecific | DownUp | DM | Loss | 10 |
| **16_69418067_69419384_54921** | 16_69418067_69419384 | CHTF8 | 54921 | 5.40142639087422 | -0.82314816 | 1 | only_H3vsUI | DownUp | notDM | Loss | 7 |
| **16_69420034_69420824_54921** | 16_69420034_69420824 | CHTF8 | 54921 | 5.40480293875506 | -0.87991418 | 1 | only_H3vsUI | DownUp | notDM | Gain | 2 |
| **16_69458910_69461484_54921** | 16_69458910_69461484 | CHTF8 | 54921 | 5.46665709214544 | -0.7947919 | 1 | only_H3vsUI | DownUp | notDM | Gain | 2 |
| **16_69659889_69661140_7014** | 16_69659889_69661140 | TERF2 | 7014 | 5.38026914378247 | -0.79010348 | 8 | notSpecific | UpDown | DM | Gain | 9 |
| **16_70720232_70721772_92154** | 16_70720232_70721772 | MTSS2 | 92154 | 2.44715803134222 | 0.62281845 | 3 | notSpecific | DownUp | DM | Loss | 10 |
| **16_724745_725647_65990** | 16_724745_725647 | ANTKMT | 65990 | 4.65812592075971 | 0.5005284 | 6 | only_H3vsUI | Down | notDM | Loss | 2 |

| | | | | | | | | | | | |
|---|---|---|---|---|---|---|---|---|---|---|---|
| **16_73078333_73081906_463** | 16_73078333_73081906 | ZFHX3 | 463 | 2.56702636615906 | 0 | 1 | only_DIvsD7 | Down | notDM | Gain | 4 |
| **16_73082329_73091640_463** | 16_73082329_73091640 | ZFHX3 | 463 | 2.95182303531591 | 0.00428077 | 5 | only_DIvsD7 | Down | notDM | Loss | 4 |
| **16_73092911_73095817_463** | 16_73092911_73095817 | ZFHX3 | 463 | 2.57863920996807 | 0.00591474 | 1 | only_DIvsD7 | Down | notDM | Loss | 8 |
| **16_732758_734095_65990** | 16_732758_734095 | ANTKMT | 65990 | 4.5689522871951 | 0.6611088 | 1 | only_H3vsUI | Down | notDM | Loss | 7 |
| **16_734530_734815_65990** | 16_734530_734815 | ANTKMT | 65990 | 4.56043272380657 | 0.65626248 | 1 | only_H3vsUI | Down | notDM | Gain | 2 |
| **16_744280_745260_65990** | 16_744280_745260 | ANTKMT | 65990 | 4.41328299563082 | 0.62579771 | 4 | only_H3vsUI | Down | notDM | Loss | 2 |
| **16_761797_762836_65990** | 16_761797_762836 | ANTKMT | 65990 | 3.92027989463295 | 0.57070626 | 7 | only_H3vsUI | Down | notDM | Gain | 1 |
| **16_763703_764891_65990** | 16_763703_764891 | ANTKMT | 65990 | 3.79712898779655 | 0.11761425 | 5 | only_H3vsUI | Down | notDM | Loss | 2 |
| **16_765330_767318_65990** | 16_765330_767318 | ANTKMT | 65990 | 3.58444430716518 | -0.70988293 | 2 | only_H3vsUI | Down | notDM | Loss | 8 |
| **16_768884_771277_65990** | 16_768884_771277 | ANTKMT | 65990 | 0 | 0.14981739 | 1 | only_H3vsUI | Down | notDM | Loss | 8 |
| **16_771572_772139_65990** | 16_771572_772139 | ANTKMT | 65990 | 2.61909333062674 | -0.02035677 | 2 | only_H3vsUI | Down | notDM | Loss | 8 |
| **16_777200_778058_65990** | 16_777200_778058 | ANTKMT | 65990 | 3.78132445566699 | 0.29362053 | 4 | only_H3vsUI | Down | notDM | Loss | 7 |
| **16_79634332_79636658_4094** | 16_79634332_79636658 | MAF | 4094 | 0 | 0.93136912 | 4 | only_H3vsUI | UpDown | notDM | Loss | 3 |
| **16_80040113_80041255_4094** | 16_80040113_80041255 | MAF | 4094 | 5.60799514256532 | 0.97926891 | 6 | only_H3vsUI | UpDown | notDM | Gain | 1 |
| **16_80074188_80074967_4094** | 16_80074188_80074967 | MAF | 4094 | 5.64303693670422 | 0.95785964 | 6 | only_H3vsUI | UpDown | notDM | Gain | 1 |
| **16_810554_811621_65990** | 16_810554_811621 | ANTKMT | 65990 | 4.59547417586085 | 0.63192428 | 6 | only_H3vsUI | Down | notDM | Loss | 1 |
| **16_84390385_84391284_9914** | 16_84390385_84391284 | ATP2C2 | 9914 | 4.03542973818455 | -0.09030849 | 6 | only_DIvsD7 | UpDown | notDM | Loss | 3 |
| **16_84565104_84567046_23406** | 16_84565104_84567046 | COTL1 | 23406 | 4.92756539241292 | 0.6093168 | 7 | only_DIvsD7 | DownUp | notDM | Loss | 5 |
| **16_84632554_84637576_23406** | 16_84632554_84637576 | COTL1 | 23406 | 4.14946545099545 | 0.27485172 | 7 | only_DIvsD7 | DownUp | notDM | Loss | 5 |
| **16_84638012_84638922_23406** | 16_84638012_84638922 | COTL1 | 23406 | 4.10591874028637 | 0.31141014 | 8 | only_DIvsD7 | DownUp | notDM | Loss | 7 |
| **16_84647909_84651277_23406** | 16_84647909_84651277 | COTL1 | 23406 | 2.60959440922522 | 0.87009344 | 1 | only_DIvsD7 | DownUp | notDM | Loss | 6 |
| **16_84652169_84654478_23406** | 16_84652169_84654478 | COTL1 | 23406 | 2.68841982200271 | 0.61374279 | 7 | only_DIvsD7 | DownUp | notDM | Gain | 5 |
| **16_84681207_84681974_23406** | 16_84681207_84681974 | COTL1 | 23406 | 4.47020461541772 | 0.66104117 | 1 | only_DIvsD7 | DownUp | notDM | Loss | 7 |
| **16_84682550_84683520_23406** | 16_84682550_84683520 | COTL1 | 23406 | 4.48952256075789 | 0.42895753 | 1 | only_DIvsD7 | DownUp | notDM | Loss | 7 |
| **16_84683641_84684918_23406** | 16_84683641_84684918 | COTL1 | 23406 | 4.50460677064195 | 0.42248294 | 6 | only_DIvsD7 | DownUp | notDM | Gain | 2 |
| **16_84758805_84768620_23406** | 16_84758805_84768620 | COTL1 | 23406 | 5.02988678082149 | -0.91739833 | 7 | only_DIvsD7 | DownUp | notDM | Gain | 5 |
| **16_84788199_84788501_23406** | 16_84788199_84788501 | COTL1 | 23406 | 5.1351899172031 | 0.8046279 | 8 | only_DIvsD7 | DownUp | notDM | Gain | 2 |
| **16_856439_857607_65990** | 16_856439_857607 | ANTKMT | 65990 | 4.93086246912773 | 0.74440265 | 1 | only_H3vsUI | Down | notDM | Loss | 7 |
| **16_86542574_86543843_2294** | 16_86542574_86543843 | FOXF1 | 2294 | 2.48000694295715 | -0.00217239 | 4 | notSpecific | UpDown | DM | Loss | 10 |
| **16_86587698_86588612_64779** | 16_86587698_86588612 | MTHFSD | 64779 | 2.36172783601759 | 0.37432196 | 1 | only_DIvsD7 | Down | notDM | Gain | 4 |
| **16_86588957_86589884_64779** | 16_86588957_86589884 | MTHFSD | 64779 | 2.07188200730613 | 0.37432196 | 1 | only_DIvsD7 | Down | notDM | Loss | 8 |
| **16_86597584_86598549_64779** | 16_86597584_86598549 | MTHFSD | 64779 | 3.16435285578444 | 0 | 2 | only_DIvsD7 | Down | notDM | Gain | 4 |
| **16_87887767_87900612_8140** | 16_87887767_87900612 | SLC7A5 | 8140 | 3.40036527334994 | 0.13826974 | 5 | notSpecific | Down | DM | Loss | 10 |
| **16_89006983_89007551_100129697** | 16_89006983_89007551 | LOC100129697 | 100129697 | 2.63948648926859 | 0 | 4 | notSpecific | Up | DM | Loss | 10 |

| | | | | | | | | | | | |
|---|---|---|---|---|---|---|---|---|---|---|---|
| **16_89006983_89007551_51693** | 16_89006983_89007551 | TRAPPC2L | 51693 | 4.92163966277704 | 0.68335349 | 4 | notSpecific | UpDown | DM | Loss | 10 |
| **16_89006983_89007551_863** | 16_89006983_89007551 | CBFA2T3 | 863 | 1.7481880270062 | 0 | 4 | notSpecific | UpDown | DM | Loss | 10 |
| **16_89282324_89284003_197320** | 16_89282324_89284003 | ZNF778 | 197320 | 2.03742649794062 | 0.3415602 | 1 | only_DIvsD7 | Down | notDM | Loss | 6 |
| **16_89284205_89285815_197320** | 16_89284205_89285815 | ZNF778 | 197320 | 1.98227123303957 | 0.05135386 | 1 | only_DIvsD7 | Down | notDM | Loss | 8 |
| **16_89895232_89896419_2175** | 16_89895232_89896419 | FANCA | 2175 | 4.08532626242399 | 0.56459541 | 2 | notSpecific | Down | DM | Loss | 10 |
| **16_89895232_89896419_22980** | 16_89895232_89896419 | TCF25 | 22980 | 4.63900810352324 | -0.54566704 | 2 | notSpecific | Up | DM | Loss | 10 |
| **16_89895232_89896419_84501** | 16_89895232_89896419 | SPIRE2 | 84501 | 2.51454775266029 | 0.87152484 | 2 | notSpecific | Up | DM | Loss | 10 |
| **16_89989969_89992612_172** | 16_89989969_89992612 | AFG3L1P | 172 | 4.66693896721471 | 0.62101473 | 2 | notSpecific | Down | DM | Loss | 10 |
| **16_89989969_89992612_22980** | 16_89989969_89992612 | TCF25 | 22980 | 4.69829197594449 | 0.5133034 | 2 | notSpecific | Up | DM | Loss | 10 |
| **16_89989969_89992612_2622** | 16_89989969_89992612 | GAS8 | 2622 | 4.98443729479608 | 0.76686743 | 2 | notSpecific | Down | DM | Loss | 10 |
| **16_89989969_89992612_54849** | 16_89989969_89992612 | DEF8 | 54849 | 4.35295391171009 | 0.55878462 | 2 | notSpecific | Up | DM | Loss | 10 |
| **17_14199959_14201713_9953** | 17_14199959_14201713 | HS3ST3B1 | 9953 | 3.42942926438179 | 0 | 2 | only_H3vsUI | Up | notDM | Loss | 3 |
| **17_14206115_14206594_9953** | 17_14206115_14206594 | HS3ST3B1 | 9953 | 3.23477029516092 | 0 | 4 | only_H3vsUI | Up | notDM | Loss | 3 |
| **17_16115718_16118512_57336** | 17_16115718_16118512 | ZNF287 | 57336 | 5.54897504423598 | -0.81584191 | 1 | only_H3vsUI | Down | notDM | Loss | 2 |
| **17_16119003_16122410_57336** | 17_16119003_16122410 | ZNF287 | 57336 | 5.54416605975773 | -0.81584191 | 1 | only_H3vsUI | Down | notDM | Gain | 4 |
| **17_1612580_1619073_124997** | 17_1612580_1619073 | WDR81 | 124997 | 2.87215627274829 | 0.05472011 | 1 | only_H3vsUI | DownUp | notDM | Loss | 6 |
| **17_1619938_1620448_124997** | 17_1619938_1620448 | WDR81 | 124997 | 2.04139268515822 | 0.11278102 | 5 | only_H3vsUI | DownUp | notDM | Gain | 4 |
| **17_1620577_1622140_124997** | 17_1620577_1622140 | WDR81 | 124997 | 2.87448181769947 | 0.10239462 | 5 | only_H3vsUI | DownUp | notDM | Gain | 4 |
| **17_1623436_1624511_124997** | 17_1623436_1624511 | WDR81 | 124997 | 3.5572665288699 | 0.12536462 | 4 | only_H3vsUI | DownUp | notDM | Loss | 7 |
| **17_16256947_16257524_57336** | 17_16256947_16257524 | ZNF287 | 57336 | 5.33243239995032 | -0.80872562 | 4 | only_H3vsUI | Down | notDM | Loss | 7 |
| **17_1627034_1627908_124997** | 17_1627034_1627908 | WDR81 | 124997 | 1.88649072517248 | 0.02081101 | 1 | only_H3vsUI | DownUp | notDM | Loss | 8 |
| **17_1628122_1629490_124997** | 17_1628122_1629490 | WDR81 | 124997 | 2.14612803567824 | 0.05851309 | 1 | only_H3vsUI | DownUp | notDM | Loss | 8 |
| **17_16288619_16291970_57336** | 17_16288619_16291970 | ZNF287 | 57336 | 5.25659989815348 | -0.73314015 | 1 | only_H3vsUI | Down | notDM | Loss | 2 |
| **17_16380341_16381075_388341** | 17_16380341_16381075 | LRRC75A | 388341 | 4.1581513253927 | 0.20299381 | 3 | only_DIvsD7 | Up | notDM | Loss | 5 |
| **17_1638826_1640779_124997** | 17_1638826_1640779 | WDR81 | 124997 | 3.7052648623174 | 0.02702885 | 6 | only_H3vsUI | DownUp | notDM | Loss | 7 |
| **17_16394175_16395223_388341** | 17_16394175_16395223 | LRRC75A | 388341 | 2.38916608436453 | 0.05082088 | 4 | only_DIvsD7 | Up | notDM | Loss | 3 |
| **17_16395595_16396342_388341** | 17_16395595_16396342 | LRRC75A | 388341 | 2.11394335230684 | -0.01335347 | 2 | only_DIvsD7 | Up | notDM | Loss | 3 |
| **17_16439240_16440342_57336** | 17_16439240_16440342 | ZNF287 | 57336 | 4.50757254376457 | -0.71646309 | 1 | only_H3vsUI | Down | notDM | Gain | 4 |
| **17_16471506_16472946_57336** | 17_16471506_16472946 | ZNF287 | 57336 | 0 | 0.9948342 | 2 | only_H3vsUI | Down | notDM | Gain | 4 |
| **17_16492000_16493217_57336** | 17_16492000_16493217 | ZNF287 | 57336 | 4.28963353900965 | -0.67689387 | 2 | only_H3vsUI | Down | notDM | Loss | 7 |
| **17_17107415_17110521_10400** | 17_17107415_17110521 | PEMT | 10400 | 5.57429176627511 | 0.79534677 | 1 | notSpecific | Up | DM | Loss | 10 |
| **17_17107415_17110521_201164** | 17_17107415_17110521 | PLD6 | 201164 | 0 | 0.69415932 | 1 | notSpecific | Up | DM | Loss | 10 |
| **17_17107415_17110521_8533** | 17_17107415_17110521 | COPS3 | 8533 | 4.86974200928067 | -0.69607747 | 1 | notSpecific | Up | DM | Loss | 10 |
| **17_17683824_17685952_10743** | 17_17683824_17685952 | RAI1 | 10743 | 0 | 0.01354225 | 2 | notSpecific | Down | DM | Loss | 10 |

| | | | | | | | | | | | |
|---|---|---|---|---|---|---|---|---|---|---|---|
| **17_17683824_17685952_140771** | 17_17683824_17685952 | SMCR5 | 140771 | 2.99255351783214 | 0 | 2 | notSpecific | Up | DM | Loss | 10 |
| **17_17683824_17685952_6720** | 17_17683824_17685952 | SREBF1 | 6720 | 4.61232813812498 | 0.57783345 | 2 | notSpecific | DownUp | DM | Loss | 10 |
| **17_18215775_18218145_140775** | 17_18215775_18218145 | SMCR8 | 140775 | 2.75739602879302 | -0.10683841 | 1 | notSpecific | Up | DM | Loss | 10 |
| **17_18215775_18218145_7156** | 17_18215775_18218145 | TOP3A | 7156 | 2.11394335230684 | 0.38488096 | 1 | notSpecific | Down | DM | Loss | 10 |
| **17_26553904_26556049_23615** | 17_26553904_26556049 | PYY2 | 23615 | 1.81954393554187 | 0 | 1 | only_H3vsUI | Up | notDM | Gain | 4 |
| **17_27071396_27074788_116238** | 17_27071396_27074788 | TLCD1 | 116238 | 4.24174565257064 | -0.55128964 | 1 | notSpecific | Down | DM | Loss | 10 |
| **17_27071396_27074788_284086** | 17_27071396_27074788 | NEK8 | 284086 | 4.19245593851198 | 0.65119537 | 1 | notSpecific | Up | DM | Loss | 10 |
| **17_27071396_27074788_55731** | 17_27071396_27074788 | FAM222B | 55731 | 4.97797039728116 | -0.96561965 | 1 | notSpecific | Down | DM | Loss | 10 |
| **17_27071396_27074788_83871** | 17_27071396_27074788 | RAB34 | 83871 | 4.41438832693107 | -0.74001756 | 1 | notSpecific | Up | DM | Loss | 10 |
| **17_27071396_27074788_9618** | 17_27071396_27074788 | TRAF4 | 9618 | 2.55509444857832 | -0.49214384 | 1 | notSpecific | Up | DM | Loss | 10 |
| **17_28431160_28432467_84081** | 17_28431160_28432467 | NSRP1 | 84081 | 4.05556944006099 | -0.31169049 | 2 | notSpecific | Down | DM | Gain | 9 |
| **17_29886306_29889243_55813** | 17_29886306_29889243 | UTP6 | 55813 | 5.5308257069915 | -0.88152638 | 1 | only_H3vsUI | Down | notDM | Gain | 2 |
| **17_30168964_30170216_55813** | 17_30168964_30170216 | UTP6 | 55813 | 4.76727463123829 | -0.63797954 | 4 | only_H3vsUI | Down | notDM | Gain | 2 |
| **17_30227051_30228582_55813** | 17_30227051_30228582 | UTP6 | 55813 | 2.17609125905568 | -0.02383534 | 2 | only_H3vsUI | Down | notDM | Gain | 4 |
| **17_30250018_30251334_55813** | 17_30250018_30251334 | UTP6 | 55813 | 4.32815526196995 | 0.54202669 | 4 | only_H3vsUI | Down | notDM | Gain | 1 |
| **17_30333433_30334300_55813** | 17_30333433_30334300 | UTP6 | 55813 | 5.01996327331919 | 0.79251235 | 2 | only_H3vsUI | Down | notDM | Loss | 7 |
| **17_30587865_30589932_162494** | 17_30587865_30589932 | RHBDL3 | 162494 | 3.51375015008182 | -0.01735589 | 6 | only_H3vsUI | DownUp | notDM | Loss | 3 |
| **17_30593256_30594418_162494** | 17_30593256_30594418 | RHBDL3 | 162494 | 1.79239168949825 | 0.02379248 | 2 | only_H3vsUI | DownUp | notDM | Loss | 3 |
| **17_30598105_30599353_162494** | 17_30598105_30599353 | RHBDL3 | 162494 | 3.5164031484474 | 0.02850253 | 6 | only_H3vsUI | DownUp | notDM | Gain | 1 |
| **17_30601338_30603054_162494** | 17_30601338_30603054 | RHBDL3 | 162494 | 3.8140477209956 | 0.02863455 | 3 | only_H3vsUI | DownUp | notDM | Gain | 1 |
| **17_33306138_33307455_3980** | 17_33306138_33307455 | LIG3 | 3980 | 1.94448267215017 | -0.65344106 | 2 | only_H3vsUI | DownUp | notDM | Loss | 8 |
| **17_33308033_33309581_3980** | 17_33308033_33309581 | LIG3 | 3980 | 2.69284691927723 | -0.65344106 | 1 | only_H3vsUI | DownUp | notDM | Gain | 4 |
| **17_33405424_33416092_3980** | 17_33405424_33416092 | LIG3 | 3980 | 4.99060964931747 | -0.62796706 | 1 | only_H3vsUI | DownUp | notDM | Loss | 7 |
| **17_33416544_33417502_3980** | 17_33416544_33417502 | LIG3 | 3980 | 5.03735078863406 | -0.65950176 | 3 | only_H3vsUI | DownUp | notDM | Gain | 2 |
| **17_36610375_36613322_3927** | 17_36610375_36613322 | LASP1 | 3927 | 5.61602891153288 | 0.84052286 | 1 | notSpecific | DownUp | DM | Loss | 10 |
| **17_37027070_37042190_3927** | 17_37027070_37042190 | LASP1 | 3927 | 2.87966920563205 | -0.87256392 | 1 | notSpecific | DownUp | DM | Loss | 10 |
| **17_37027070_37042190_9349** | 17_37027070_37042190 | RPL23 | 9349 | 4.23065324717922 | 0.32898549 | 1 | notSpecific | Up | DM | Loss | 10 |
| **17_37308471_37310010_57125** | 17_37308471_37310010 | PLXDC1 | 57125 | 0 | -0.01195955 | 1 | only_DIvsD7 | UpDown | notDM | Loss | 3 |
| **17_37310125_37310726_57125** | 17_37310125_37310726 | PLXDC1 | 57125 | 2.80753502806885 | 0 | 1 | only_DIvsD7 | UpDown | notDM | Loss | 3 |
| **17_37618286_37622245_10948** | 17_37618286_37622245 | STARD3 | 10948 | 5.23322208789954 | -0.8199888 | 1 | only_DIvsD7 | DownUp | notDM | Loss | 7 |
| **17_37729931_37731115_10948** | 17_37729931_37731115 | STARD3 | 10948 | 4.79453684105076 | 0.51580044 | 2 | only_DIvsD7 | DownUp | notDM | Loss | 7 |
| **17_37762219_37763939_2064** | 17_37762219_37763939 | ERBB2 | 2064 | 4.96527697848537 | -0.62255559 | 4 | notSpecific | Down | DM | Loss | 10 |
| **17_37762219_37763939_22806** | 17_37762219_37763939 | IKZF3 | 22806 | 5.40898746176042 | 0.99803032 | 4 | notSpecific | DownUp | DM | Loss | 10 |
| **17_37762219_37763939_4761** | 17_37762219_37763939 | NEUROD2 | 4761 | 2.41161970596323 | 0 | 4 | notSpecific | UpDown | DM | Loss | 10 |

| | | | | | | | | | | | |
|---|---|---|---|---|---|---|---|---|---|---|---|
| **17_37774513_37775776_10948** | 17_37774513_37775776 | STARD3 | 10948 | 4.24447504468901 | 0.31230638 | 1 | only_DIvsD7 | DownUp | notDM | Loss | 6 |
| **17_37778353_37779857_10948** | 17_37778353_37779857 | STARD3 | 10948 | 4.12959322836793 | 0.54319157 | 4 | only_DIvsD7 | DownUp | notDM | Gain | 5 |
| **17_37791656_37793366_10948** | 17_37791656_37793366 | STARD3 | 10948 | 0 | -0.256968 | 1 | only_DIvsD7 | DownUp | notDM | Loss | 8 |
| **17_37793565_37795585_10948** | 17_37793565_37795585 | STARD3 | 10948 | 2.16435285578444 | -0.34508012 | 1 | only_DIvsD7 | DownUp | notDM | Loss | 8 |
| **17_37819917_37821265_10948** | 17_37819917_37821265 | STARD3 | 10948 | 4.42465300041526 | -0.70505363 | 4 | only_DIvsD7 | DownUp | notDM | Loss | 7 |
| **17_37821671_37823958_10948** | 17_37821671_37823958 | STARD3 | 10948 | 4.45239984591144 | -0.74266537 | 6 | only_DIvsD7 | DownUp | notDM | Loss | 7 |
| **17_37824638_37825311_10948** | 17_37824638_37825311 | STARD3 | 10948 | 4.49564145324872 | -0.75170782 | 4 | only_DIvsD7 | DownUp | notDM | Loss | 7 |
| **17_37844269_37845688_10948** | 17_37844269_37845688 | STARD3 | 10948 | 4.70629095725876 | -0.49038484 | 1 | only_DIvsD7 | DownUp | notDM | Gain | 2 |
| **17_38135593_38136748_10948** | 17_38135593_38136748 | STARD3 | 10948 | 5.53424700670854 | -0.7710599 | 2 | only_DIvsD7 | DownUp | notDM | Gain | 2 |
| **17_38485842_38489285_147184** | 17_38485842_38489285 | TMEM99 | 147184 | 5.68670239137177 | -0.92650363 | 7 | only_DIvsD7 | DownUp | notDM | Loss | 7 |
| **17_38600156_38612238_147179** | 17_38600156_38612238 | WIPF2 | 147179 | 5.3513111253286 | -0.7506691 | 1 | notSpecific | UpDown | DM | Loss | 10 |
| **17_38600156_38612238_3487** | 17_38600156_38612238 | IGFBP4 | 3487 | 2.64836001098093 | 0.73332594 | 1 | notSpecific | UpDown | DM | Loss | 10 |
| **17_38600156_38612238_7153** | 17_38600156_38612238 | TOP2A | 7153 | 4.41478956911867 | 0.4260312 | 1 | notSpecific | Down | DM | Loss | 10 |
| **17_38600156_38612238_84951** | 17_38600156_38612238 | TNS4 | 84951 | 4.65907911565073 | -0.76231393 | 1 | notSpecific | UpDown | DM | Loss | 10 |
| **17_38935346_38936489_147184** | 17_38935346_38936489 | TMEM99 | 147184 | 4.58961454063127 | -0.40480351 | 4 | only_DIvsD7 | DownUp | notDM | Gain | 2 |
| **17_38960025_38961033_147184** | 17_38960025_38961033 | TMEM99 | 147184 | 4.15775888604686 | -0.43112454 | 6 | only_DIvsD7 | DownUp | notDM | Gain | 2 |
| **17_38965578_38968187_147184** | 17_38965578_38968187 | TMEM99 | 147184 | 3.85564028089015 | -0.70825692 | 4 | only_DIvsD7 | DownUp | notDM | Gain | 5 |
| **17_38973539_38975007_147184** | 17_38973539_38975007 | TMEM99 | 147184 | 2.54654266347813 | -0.15290325 | 2 | only_DIvsD7 | DownUp | notDM | Gain | 4 |
| **17_38975485_38977532_147184** | 17_38975485_38977532 | TMEM99 | 147184 | 2.11058971029925 | -0.15297538 | 1 | only_DIvsD7 | DownUp | notDM | Loss | 8 |
| **17_39005609_39012714_147184** | 17_39005609_39012714 | TMEM99 | 147184 | 4.47999256210707 | 0.40993376 | 7 | only_DIvsD7 | DownUp | notDM | Gain | 5 |
| **17_39056901_39061176_147184** | 17_39056901_39061176 | TMEM99 | 147184 | 4.9111096471919 | 0.77272674 | 7 | only_DIvsD7 | DownUp | notDM | Loss | 5 |
| **17_39238357_39240890_100132386** | 17_39238357_39240890 | KRTAP4-9 | 100132386 | 0 | 0 | 6 | only_H3vsUI | DownUp | notDM | Gain | 4 |
| **17_39260021_39261692_100132386** | 17_39260021_39261692 | KRTAP4-9 | 100132386 | 0 | 0 | 6 | only_H3vsUI | DownUp | notDM | Gain | 4 |
| **17_39821826_39824348_60681** | 17_39821826_39824348 | FKBP10 | 60681 | 5.16021334191679 | -0.66272941 | 4 | only_DIvsD7 | DownUp | notDM | Loss | 7 |
| **17_39928389_39937126_60681** | 17_39928389_39937126 | FKBP10 | 60681 | 4.50692424213177 | 0.6551377 | 7 | only_DIvsD7 | DownUp | notDM | Loss | 5 |
| **17_39947938_39951629_60681** | 17_39947938_39951629 | FKBP10 | 60681 | 4.23889879222784 | 0.25055656 | 7 | only_DIvsD7 | DownUp | notDM | Loss | 5 |
| **17_39956411_39958192_60681** | 17_39956411_39958192 | FKBP10 | 60681 | 4.03546976348128 | -0.06523503 | 7 | only_DIvsD7 | DownUp | notDM | Loss | 7 |
| **17_39966531_39967866_55175** | 17_39966531_39967866 | KLHL11 | 55175 | 4.73087106285947 | 0.59835993 | 2 | only_H3vsUI | Down | notDM | Loss | 7 |
| **17_39966531_39967866_60681** | 17_39966531_39967866 | FKBP10 | 60681 | 3.11991541025799 | 0.98685347 | 2 | only_DIvsD7 | DownUp | notDM | Loss | 6 |
| **17_39968070_39971505_55175** | 17_39968070_39971505 | KLHL11 | 55175 | 4.70046141359804 | 0.59563205 | 1 | only_H3vsUI | Down | notDM | Loss | 4 |
| **17_39968070_39971505_60681** | 17_39968070_39971505 | FKBP10 | 60681 | 0 | 0.98663052 | 1 | only_DIvsD7 | DownUp | notDM | Loss | 8 |
| **17_39992643_39993359_60681** | 17_39992643_39993359 | FKBP10 | 60681 | 4.37036503045392 | -0.70343176 | 2 | only_DIvsD7 | DownUp | notDM | Gain | 2 |
| **17_40018720_40021424_55175** | 17_40018720_40021424 | KLHL11 | 55175 | 2.40312052117582 | 0.48488378 | 2 | only_H3vsUI | Down | notDM | Loss | 8 |
| **17_40018720_40021424_60681** | 17_40018720_40021424 | FKBP10 | 60681 | 4.69688037168276 | 0.60721973 | 2 | only_DIvsD7 | DownUp | notDM | Loss | 6 |

| | | | | | | | | | | | |
|---|---|---|---|---|---|---|---|---|---|---|---|
| **17_40153902_40157177_60681** | 17_40153902_40157177 | FKBP10 | 60681 | 5.26651627099891 | 0.73049329 | 8 | only_DIvsD7 | DownUp | notDM | Loss | 5 |
| **17_40201198_40202564_60681** | 17_40201198_40202564 | FKBP10 | 60681 | 5.36538314253447 | 0.84750583 | 1 | only_DIvsD7 | DownUp | notDM | Loss | 6 |
| **17_40948706_40950988_124817** | 17_40948706_40950988 | CNTD1 | 124817 | 0 | -0.0867904 | 1 | only_H3vsUI | UpDown | notDM | Loss | 3 |
| **17_40953705_40958096_124817** | 17_40953705_40958096 | CNTD1 | 124817 | 3.45530177165708 | -0.04122781 | 6 | only_H3vsUI | UpDown | notDM | Loss | 6 |
| **17_42422556_42424157_10052** | 17_42422556_42424157 | GJC1 | 10052 | 5.68486599911498 | -0.9571231 | 1 | only_DIvsD7 | UpDown | notDM | Loss | 7 |
| **17_42441331_42441910_10052** | 17_42441331_42441910 | GJC1 | 10052 | 5.66863747367112 | 0.99828099 | 2 | only_DIvsD7 | UpDown | notDM | Gain | 1 |
| **17_42635665_42639714_10052** | 17_42635665_42639714 | GJC1 | 10052 | 5.42888929209379 | 0.998053 | 2 | only_DIvsD7 | UpDown | notDM | Loss | 6 |
| **17_42733015_42734100_10052** | 17_42733015_42734100 | GJC1 | 10052 | 5.24074887801809 | -0.77919752 | 2 | only_DIvsD7 | UpDown | notDM | Gain | 2 |
| **17_42897174_42899046_10052** | 17_42897174_42899046 | GJC1 | 10052 | 3.96066100727098 | -0.20891446 | 7 | only_DIvsD7 | UpDown | notDM | Gain | 5 |
| **17_42899606_42901031_10052** | 17_42899606_42901031 | GJC1 | 10052 | 3.85424529706612 | -0.21182215 | 8 | only_DIvsD7 | UpDown | notDM | Loss | 7 |
| **17_42904656_42907129_10052** | 17_42904656_42907129 | GJC1 | 10052 | 3.02160271602824 | 0.30172724 | 1 | only_DIvsD7 | UpDown | notDM | Loss | 6 |
| **17_42907772_42908752_10052** | 17_42907772_42908752 | GJC1 | 10052 | 0 | 0.67459222 | 2 | only_DIvsD7 | UpDown | notDM | Loss | 6 |
| **17_42974671_42976688_10052** | 17_42974671_42976688 | GJC1 | 10052 | 4.82278245907075 | -0.7130112 | 1 | only_DIvsD7 | UpDown | notDM | Loss | 7 |
| **17_42976883_42979892_10052** | 17_42976883_42979892 | GJC1 | 10052 | 4.8369946650524 | -0.7130112 | 1 | only_DIvsD7 | UpDown | notDM | Loss | 7 |
| **17_43025208_43026148_10052** | 17_43025208_43026148 | GJC1 | 10052 | 5.06830091598272 | -0.90177276 | 1 | only_DIvsD7 | UpDown | notDM | Gain | 2 |
| **17_43126667_43128820_10052** | 17_43126667_43128820 | GJC1 | 10052 | 5.33943156467393 | -0.78771663 | 1 | only_DIvsD7 | UpDown | notDM | Loss | 7 |
| **17_43129492_43130318_10052** | 17_43129492_43130318 | GJC1 | 10052 | 5.34501084997091 | -0.84326024 | 1 | only_DIvsD7 | UpDown | notDM | Gain | 2 |
| **17_43137106_43138348_10052** | 17_43137106_43138348 | GJC1 | 10052 | 5.35970081120763 | -0.78124319 | 1 | only_DIvsD7 | UpDown | notDM | Loss | 7 |
| **17_43223654_43224873_10052** | 17_43223654_43224873 | GJC1 | 10052 | 5.49896770228953 | -0.81058404 | 1 | only_DIvsD7 | UpDown | notDM | Gain | 2 |
| **17_43342989_43344111_124783** | 17_43342989_43344111 | SPATA32 | 124783 | 3.54777470538782 | 0.00133104 | 6 | only_DIvsD7 | UpDown | notDM | Loss | 3 |
| **17_43509417_43509984_201176** | 17_43509417_43509984 | ARHGAP27 | 201176 | 2.47567118832443 | 0.11859815 | 2 | notSpecific | UpDown | DM | Loss | 10 |
| **17_43970866_43971676_4137** | 17_43970866_43971676 | MAPT | 4137 | 2.3541084391474 | -0.03074811 | 2 | only_H3vsUI | UpDown | notDM | Gain | 4 |
| **17_43971882_43975586_4137** | 17_43971882_43975586 | MAPT | 4137 | 0 | -0.0409156 | 2 | only_H3vsUI | UpDown | notDM | Loss | 3 |
| **17_45897493_45900341_114881** | 17_45897493_45900341 | OSBPL7 | 114881 | 0 | 0.13027893 | 2 | only_H3vsUI | Down | notDM | Loss | 8 |
| **17_45906920_45908776_114881** | 17_45906920_45908776 | OSBPL7 | 114881 | 3.88772979728803 | 0.17038046 | 1 | only_H3vsUI | Down | notDM | Loss | 8 |
| **17_45908966_45910130_114881** | 17_45908966_45910130 | OSBPL7 | 114881 | 3.99215571104262 | 0.13889187 | 2 | only_H3vsUI | Down | notDM | Loss | 8 |
| **17_45939651_45940911_51226** | 17_45939651_45940911 | COPZ2 | 51226 | 5.24112044396528 | 0.71638669 | 7 | only_DIvsD7 | DownUp | notDM | Gain | 1 |
| **17_46025314_46027765_51226** | 17_46025314_46027765 | COPZ2 | 51226 | 4.94138718875374 | -0.59566535 | 7 | only_DIvsD7 | DownUp | notDM | Loss | 7 |
| **17_46099966_46102549_51226** | 17_46099966_46102549 | COPZ2 | 51226 | 4.10006022393115 | 0.30042297 | 1 | only_DIvsD7 | DownUp | notDM | Loss | 6 |
| **17_46102710_46104639_51226** | 17_46102710_46104639 | COPZ2 | 51226 | 4.0212306584797 | 0.31467589 | 2 | only_DIvsD7 | DownUp | notDM | Gain | 5 |
| **17_46114025_46115006_51226** | 17_46114025_46115006 | COPZ2 | 51226 | 2.12710479836481 | 0.05924529 | 1 | only_DIvsD7 | DownUp | notDM | Gain | 4 |
| **17_46122822_46125434_51226** | 17_46122822_46125434 | COPZ2 | 51226 | 3.88564387183576 | 0.69961367 | 1 | only_DIvsD7 | DownUp | notDM | Gain | 5 |
| **17_46126156_46131759_51226** | 17_46126156_46131759 | COPZ2 | 51226 | 4.04214218306496 | 0.69961367 | 1 | only_DIvsD7 | DownUp | notDM | Loss | 6 |
| **17_46176683_46178502_51226** | 17_46176683_46178502 | COPZ2 | 51226 | 4.78919983250084 | -0.52105518 | 1 | only_DIvsD7 | DownUp | notDM | Gain | 2 |

| | | | | | | | | | | | |
|---|---|---|---|---|---|---|---|---|---|---|---|
| **17_46178954_46179809_51226** | 17_46178954_46179809 | COPZ2 | 51226 | 4.80493638440671 | -0.52105518 | 1 | only_DIvsD7 | DownUp | notDM | Gain | 2 |
| **17_46204200_46205509_51226** | 17_46204200_46205509 | COPZ2 | 51226 | 4.94969731982622 | 0.98203722 | 6 | only_DIvsD7 | DownUp | notDM | Loss | 5 |
| **17_46871847_46874789_284076** | 17_46871847_46874789 | TTLL6 | 284076 | 2.1846914308176 | 0 | 7 | only_DIvsD7 | UpDown | notDM | Loss | 5 |
| **17_46876134_46879322_284076** | 17_46876134_46879322 | TTLL6 | 284076 | 3.64738297011462 | 0 | 6 | only_DIvsD7 | UpDown | notDM | Loss | 5 |
| **17_46881441_46883064_284076** | 17_46881441_46883064 | TTLL6 | 284076 | 4.05713336724909 | -0.25740119 | 4 | only_DIvsD7 | UpDown | notDM | Loss | 3 |
| **17_46885543_46887805_284076** | 17_46885543_46887805 | TTLL6 | 284076 | 3.82380015374988 | -0.30550943 | 3 | only_DIvsD7 | UpDown | notDM | Loss | 5 |
| **17_46893603_46895712_284076** | 17_46893603_46895712 | TTLL6 | 284076 | 0 | 0.40543691 | 2 | only_DIvsD7 | UpDown | notDM | Loss | 3 |
| **17_46908404_46910115_284076** | 17_46908404_46910115 | TTLL6 | 284076 | 4.14416930004167 | -0.40123706 | 1 | only_DIvsD7 | UpDown | notDM | Loss | 7 |
| **17_4710489_4711981_50488** | 17_4710489_4711981 | MINK1 | 50488 | 4.39190500686716 | 0.56201234 | 1 | only_H3vsUI | DownUp | notDM | Loss | 7 |
| **17_4736818_4739774_50488** | 17_4736818_4739774 | MINK1 | 50488 | 2.26717172840301 | -0.15520236 | 1 | only_H3vsUI | DownUp | notDM | Loss | 8 |
| **17_4742997_4744163_50488** | 17_4742997_4744163 | MINK1 | 50488 | 3.80373017097454 | -0.63128176 | 8 | only_H3vsUI | DownUp | notDM | Loss | 7 |
| **17_4750093_4756339_50488** | 17_4750093_4756339 | MINK1 | 50488 | 4.12904505988796 | 0.44340656 | 7 | only_H3vsUI | DownUp | notDM | Loss | 5 |
| **17_4760810_4763132_50488** | 17_4760810_4763132 | MINK1 | 50488 | 4.38340241049182 | 0.54741674 | 6 | only_H3vsUI | DownUp | notDM | Gain | 5 |
| **17_47645007_47646242_11248** | 17_47645007_47646242 | NXPH3 | 11248 | 3.85991848520072 | -0.10036213 | 2 | only_H3vsUI | UpDown | notDM | Loss | 3 |
| **17_47652327_47653037_11248** | 17_47652327_47653037 | NXPH3 | 11248 | 2.26481782300954 | -0.03880553 | 2 | only_H3vsUI | UpDown | notDM | Loss | 3 |
| **17_47653192_47654864_11248** | 17_47653192_47654864 | NXPH3 | 11248 | 0 | 0.06523705 | 2 | only_H3vsUI | UpDown | notDM | Loss | 3 |
| **17_48222791_48228314_84687** | 17_48222791_48228314 | PPP1R9B | 84687 | 0 | 0.03293028 | 1 | notSpecific | Down | DM | Loss | 10 |
| **17_48349527_48358617_162461** | 17_48349527_48358617 | TMEM92 | 162461 | 0 | 0.88755127 | 1 | notSpecific | UpDown | DM | Loss | 10 |
| **17_48845425_48846398_55018** | 17_48845425_48846398 | ANKRD40CL | 55018 | 2.70671778233676 | -0.02983894 | 6 | only_H3vsUI | UpDown | notDM | Loss | 3 |
| **17_4899921_4901059_50488** | 17_4899921_4901059 | MINK1 | 50488 | 5.2129542697013 | -0.71759451 | 1 | only_H3vsUI | DownUp | notDM | Loss | 7 |
| **17_4901525_4904154_50488** | 17_4901525_4904154 | MINK1 | 50488 | 5.21719958566189 | -0.7190946 | 1 | only_H3vsUI | DownUp | notDM | Loss | 7 |
| **17_4938208_4939577_7775** | 17_4938208_4939577 | ZNF232 | 7775 | 4.8784930713713 | -0.68177863 | 2 | only_H3vsUI | Down | notDM | Loss | 7 |
| **17_5014261_5016155_7775** | 17_5014261_5016155 | ZNF232 | 7775 | 0 | -0.38531564 | 2 | only_H3vsUI | Down | notDM | Loss | 8 |
| **17_5186093_5188084_7775** | 17_5186093_5188084 | ZNF232 | 7775 | 5.23280304771642 | -0.69200037 | 1 | only_H3vsUI | Down | notDM | Loss | 8 |
| **17_57838882_57842550_51651** | 17_57838882_57842550 | PTRH2 | 51651 | 4.7315645922794 | -0.81832902 | 7 | notSpecific | DownUp | DM | Gain | 9 |
| **17_57838882_57842550_81671** | 17_57838882_57842550 | VMP1 | 81671 | 4.73125022198505 | 0.74248989 | 7 | notSpecific | DownUp | DM | Gain | 9 |
| **17_57932737_57936263_6198** | 17_57932737_57936263 | RPS6KB1 | 6198 | 4.53482538625369 | 0.63963141 | 7 | notSpecific | Down | DM | Gain | 9 |
| **17_58216852_58217903_762** | 17_58216852_58217903 | CA4 | 762 | 3.9731278535997 | -0.05742702 | 4 | notSpecific | UpDown | DM | Loss | 10 |
| **17_61537502_61539129_117246** | 17_61537502_61539129 | FTSJ3 | 117246 | 5.56265204994689 | -0.81565578 | 6 | only_H3vsUI | Down | notDM | Gain | 2 |
| **17_61848670_61850735_117246** | 17_61848670_61850735 | FTSJ3 | 117246 | 4.72994193481259 | -0.57590303 | 1 | only_H3vsUI | Down | notDM | Gain | 2 |
| **17_61851973_61854642_117246** | 17_61851973_61854642 | FTSJ3 | 117246 | 4.69713340366363 | -0.60050737 | 1 | only_H3vsUI | Down | notDM | Gain | 2 |
| **17_61895477_61897497_117246** | 17_61895477_61897497 | FTSJ3 | 117246 | 3.84098383732038 | 0.60015661 | 8 | only_H3vsUI | Down | notDM | Loss | 5 |
| **17_61902748_61904321_117246** | 17_61902748_61904321 | FTSJ3 | 117246 | 2.04139268515822 | -0.24701758 | 1 | only_H3vsUI | Down | notDM | Loss | 8 |
| **17_61905145_61907220_117246** | 17_61905145_61907220 | FTSJ3 | 117246 | 2.8555191556678 | -0.25194241 | 2 | only_H3vsUI | Down | notDM | Gain | 4 |

| | | | | | | | | | | | |
|---|---|---|---|---|---|---|---|---|---|---|---|
| **17_61917246_61919815_117246** | 17_61917246_61919815 | FTSJ3 | 117246 | 4.10782026724805 | 0.16731962 | 1 | only_H3vsUI | Down | notDM | Loss | 8 |
| **17_61920565_61921816_117246** | 17_61920565_61921816 | FTSJ3 | 117246 | 4.20782279900151 | -0.45772257 | 1 | only_H3vsUI | Down | notDM | Gain | 2 |
| **17_62102457_62103929_117246** | 17_62102457_62103929 | FTSJ3 | 117246 | 5.29672879439152 | -0.75938436 | 4 | only_H3vsUI | Down | notDM | Gain | 2 |
| **17_62203562_62207146_117246** | 17_62203562_62207146 | FTSJ3 | 117246 | 5.4758657783713 | -0.78033578 | 1 | only_H3vsUI | Down | notDM | Gain | 2 |
| **17_62207792_62209070_117246** | 17_62207792_62209070 | FTSJ3 | 117246 | 5.48196404207699 | -0.78071678 | 2 | only_H3vsUI | Down | notDM | Gain | 2 |
| **17_62776096_62777450_109286553** | 17_62776096_62777450 | ARHGAP27P1-BPTFP1-KPNA2P3 | 109286553 | 2.82477646247555 | 0 | 2 | only_DIvsD7 | Up | notDM | Loss | 3 |
| **17_65373499_65374939_26207** | 17_65373499_65374939 | PITPNC1 | 26207 | 0 | 0.00968653 | 1 | notSpecific | Up | DM | Loss | 10 |
| **17_65373499_65374939_5718** | 17_65373499_65374939 | PSMD12 | 5718 | 4.03261876085072 | -0.51677504 | 1 | notSpecific | Down | DM | Loss | 10 |
| **17_6558759_6560881_51003** | 17_6558759_6560881 | MED31 | 51003 | 3.58058287681437 | 0.28091397 | 2 | notSpecific | DownUp | DM | Loss | 10 |
| **17_66244272_66246193_22901** | 17_66244272_66246193 | ARSG | 22901 | 3.96051834278071 | 0.03573787 | 1 | only_H3vsUI | UpDown | notDM | Loss | 6 |
| **17_67321394_67323163_23461** | 17_67321394_67323163 | ABCA5 | 23461 | 2.20682587603185 | 0.72034099 | 1 | only_H3vsUI | UpDown | notDM | Gain | 4 |
| **17_67323486_67324307_23461** | 17_67323486_67324307 | ABCA5 | 23461 | 2.21748394421391 | 0.2723456 | 1 | only_H3vsUI | UpDown | notDM | Loss | 3 |
| **17_67386541_67388217_23461** | 17_67386541_67388217 | ABCA5 | 23461 | 4.80085449150356 | 0.64852652 | 4 | only_H3vsUI | UpDown | notDM | Gain | 1 |
| **17_67468592_67471303_23461** | 17_67468592_67471303 | ABCA5 | 23461 | 5.16217892608745 | 0.76867008 | 7 | only_H3vsUI | UpDown | notDM | Gain | 5 |
| **17_6913646_6917949_239** | 17_6913646_6917949 | ALOX12 | 239 | 2.92427928606188 | 0 | 1 | only_H3vsUI | UpDown | notDM | Loss | 6 |
| **17_6918375_6919472_239** | 17_6918375_6919472 | ALOX12 | 239 | 3.74577721788976 | 0 | 1 | only_H3vsUI | UpDown | notDM | Loss | 3 |
| **17_7038818_7039847_1366** | 17_7038818_7039847 | CLDN7 | 1366 | 5.1026600561307 | -0.94248065 | 2 | only_DIvsD7 | UpDown | notDM | Loss | 3 |
| **17_7086975_7087902_1366** | 17_7086975_7087902 | CLDN7 | 1366 | 4.89548332091278 | -0.65394056 | 6 | only_DIvsD7 | UpDown | notDM | Gain | 1 |
| **17_7119306_7120633_1366** | 17_7119306_7120633 | CLDN7 | 1366 | 4.66162340922923 | 0.61247927 | 2 | only_DIvsD7 | UpDown | notDM | Loss | 3 |
| **17_7143184_7145714_1366** | 17_7143184_7145714 | CLDN7 | 1366 | 4.31804245491843 | 0.40379994 | 1 | only_DIvsD7 | UpDown | notDM | Loss | 6 |
| **17_7151992_7154806_1366** | 17_7151992_7154806 | CLDN7 | 1366 | 4.06844561835435 | -0.5382931 | 2 | only_DIvsD7 | UpDown | notDM | Loss | 6 |
| **17_7155568_7157537_1366** | 17_7155568_7157537 | CLDN7 | 1366 | 3.95308284391209 | -0.5382931 | 1 | only_DIvsD7 | UpDown | notDM | Loss | 6 |
| **17_7161849_7163055_1366** | 17_7161849_7163055 | CLDN7 | 1366 | 3.50650503240487 | -0.01328913 | 6 | only_DIvsD7 | UpDown | notDM | Gain | 1 |
| **17_7166269_7167289_1366** | 17_7166269_7167289 | CLDN7 | 1366 | 0 | 0.88564513 | 6 | only_DIvsD7 | UpDown | notDM | Loss | 3 |
| **17_7167402_7168852_1366** | 17_7167402_7168852 | CLDN7 | 1366 | 2.95036485437612 | 0.34913209 | 3 | only_DIvsD7 | UpDown | notDM | Gain | 1 |
| **17_7183618_7185902_1366** | 17_7183618_7185902 | CLDN7 | 1366 | 4.23319924158477 | 0.2554546 | 2 | only_DIvsD7 | UpDown | notDM | Gain | 1 |
| **17_7298072_7299110_6665** | 17_7298072_7299110 | SOX15 | 6665 | 5.28843033020024 | -0.86184466 | 2 | only_DIvsD7 | DownUp | notDM | Loss | 7 |
| **17_73073806_73076592_6613** | 17_73073806_73076592 | SUMO2 | 6613 | 5.01075352351442 | -0.64534406 | 7 | only_DIvsD7 | DownUp | notDM | Loss | 7 |
| **17_73081035_73083792_6613** | 17_73081035_73083792 | SUMO2 | 6613 | 4.9791247993763 | -0.64018978 | 1 | only_DIvsD7 | DownUp | notDM | Gain | 2 |
| **17_7308283_7309791_1366** | 17_7308283_7309791 | CLDN7 | 1366 | 5.1515935293836 | 0.95617624 | 1 | only_DIvsD7 | UpDown | notDM | Loss | 3 |
| **17_73118542_73120491_6613** | 17_73118542_73120491 | SUMO2 | 6613 | 4.76795690132047 | -0.66298747 | 7 | only_DIvsD7 | DownUp | notDM | Gain | 5 |
| **17_73120660_73123087_6613** | 17_73120660_73123087 | SUMO2 | 6613 | 4.74828108013984 | -0.67028487 | 7 | only_DIvsD7 | DownUp | notDM | Loss | 7 |
| **17_73154311_73156677_6613** | 17_73154311_73156677 | SUMO2 | 6613 | 4.35067434823328 | -0.80870461 | 3 | only_DIvsD7 | DownUp | notDM | Loss | 7 |
| **17_73175771_73178643_6613** | 17_73175771_73178643 | SUMO2 | 6613 | 2.65896484266443 | -0.00228325 | 1 | only_DIvsD7 | DownUp | notDM | Loss | 8 |

| | | | | | | | | | | | |
|---|---|---|---|---|---|---|---|---|---|---|---|
| **17_73507586_73508932_23558** | 17_73507586_73508932 | WBP2 | 23558 | 5.53460858417995 | 0.77727483 | 6 | notSpecific | Up | DM | Loss | 10 |
| **17_73507586_73508932_2584** | 17_73507586_73508932 | GALK1 | 2584 | 5.40204805277833 | 0.86711798 | 6 | notSpecific | Down | DM | Loss | 10 |
| **17_73507586_73508932_283989** | 17_73507586_73508932 | TSEN54 | 283989 | 3.56561172490206 | -0.30388994 | 6 | notSpecific | Down | DM | Loss | 10 |
| **17_73507586_73508932_2885** | 17_73507586_73508932 | GRB2 | 2885 | 5.07056609086762 | 0.64669015 | 6 | notSpecific | DownUp | DM | Loss | 10 |
| **17_73507586_73508932_30833** | 17_73507586_73508932 | NT5C | 30833 | 5.57945317857278 | -0.84810074 | 6 | notSpecific | Down | DM | Loss | 10 |
| **17_73507586_73508932_3993** | 17_73507586_73508932 | LLGL2 | 3993 | 4.65286493914142 | 0.67252524 | 6 | notSpecific | DownUp | DM | Loss | 10 |
| **17_73507586_73508932_57409** | 17_73507586_73508932 | MIF4GD | 57409 | 5.38081521469846 | -0.73392342 | 6 | notSpecific | UpDown | DM | Loss | 10 |
| **17_73507586_73508932_57513** | 17_73507586_73508932 | CASKIN2 | 57513 | 3.43071988786328 | 0.19967292 | 6 | notSpecific | Down | DM | Loss | 10 |
| **17_73507586_73508932_9772** | 17_73507586_73508932 | TMEM94 | 9772 | 4.73923205337659 | 0.86938047 | 6 | notSpecific | Down | DM | Loss | 10 |
| **17_73673446_73678848_6613** | 17_73673446_73678848 | SUMO2 | 6613 | 5.69403453849216 | -0.83724358 | 7 | only_DIvsD7 | DownUp | notDM | Loss | 7 |
| **17_7385732_7387558_6665** | 17_7385732_7387558 | SOX15 | 6665 | 5.02462110704954 | -0.63829146 | 1 | only_DIvsD7 | DownUp | notDM | Loss | 7 |
| **17_7388143_7391828_6665** | 17_7388143_7391828 | SOX15 | 6665 | 5.00673552071885 | -0.63829146 | 1 | only_DIvsD7 | DownUp | notDM | Loss | 7 |
| **17_73890501_73892659_85302** | 17_73890501_73892659 | FBF1 | 85302 | 4.64797922646009 | -0.49947878 | 1 | only_H3vsUI | Down | notDM | Gain | 4 |
| **17_73935861_73937066_85302** | 17_73935861_73937066 | FBF1 | 85302 | 1.73239375982297 | -0.36779377 | 2 | only_H3vsUI | Down | notDM | Loss | 8 |
| **17_73971868_73973570_85302** | 17_73971868_73973570 | FBF1 | 85302 | 4.54096730642925 | -0.41053446 | 2 | only_H3vsUI | Down | notDM | Loss | 2 |
| **17_74133966_74135642_8877** | 17_74133966_74135642 | SPHK1 | 8877 | 5.37493338566763 | 0.99965123 | 6 | only_DIvsD7 | DownUp | notDM | Loss | 5 |
| **17_74136662_74138335_8877** | 17_74136662_74138335 | SPHK1 | 8877 | 5.36997242943038 | 0.72789788 | 2 | only_DIvsD7 | DownUp | notDM | Loss | 6 |
| **17_74348758_74349557_8877** | 17_74348758_74349557 | SPHK1 | 8877 | 4.49323492107857 | 0.43767994 | 1 | only_DIvsD7 | DownUp | notDM | Loss | 7 |
| **17_74378586_74380508_8877** | 17_74378586_74380508 | SPHK1 | 8877 | 2.26245108973043 | 0.91233924 | 2 | only_DIvsD7 | DownUp | notDM | Loss | 6 |
| **17_74380630_74382583_8877** | 17_74380630_74382583 | SPHK1 | 8877 | 0 | 0.91271803 | 2 | only_DIvsD7 | DownUp | notDM | Loss | 6 |
| **17_74446277_74448733_8877** | 17_74446277_74448733 | SPHK1 | 8877 | 4.8665059085078 | 0.75868576 | 1 | only_DIvsD7 | DownUp | notDM | Loss | 6 |
| **17_74518847_74519277_8877** | 17_74518847_74519277 | SPHK1 | 8877 | 5.16467102352189 | -0.84007273 | 6 | only_DIvsD7 | DownUp | notDM | Gain | 2 |
| **17_74519405_74519980_8877** | 17_74519405_74519980 | SPHK1 | 8877 | 5.16632648657392 | -0.78815634 | 6 | only_DIvsD7 | DownUp | notDM | Gain | 2 |
| **17_74551946_74553697_8877** | 17_74551946_74553697 | SPHK1 | 8877 | 5.23365086789997 | 0.69758283 | 1 | only_DIvsD7 | DownUp | notDM | Gain | 5 |
| **17_74553938_74558284_8877** | 17_74553938_74558284 | SPHK1 | 8877 | 5.25815339977701 | 0.81129055 | 1 | only_DIvsD7 | DownUp | notDM | Loss | 6 |
| **17_74683842_74686279_8877** | 17_74683842_74686279 | SPHK1 | 8877 | 5.49290280309254 | 0.87120502 | 6 | only_DIvsD7 | DownUp | notDM | Gain | 1 |
| **17_74707176_74708246_8877** | 17_74707176_74708246 | SPHK1 | 8877 | 5.51386722349913 | 0.83544876 | 2 | only_DIvsD7 | DownUp | notDM | Loss | 6 |
| **17_74728347_74732445_8877** | 17_74728347_74732445 | SPHK1 | 8877 | 5.55097030131801 | 0.81475801 | 1 | only_DIvsD7 | DownUp | notDM | Loss | 6 |
| **17_74734142_74735842_8877** | 17_74734142_74735842 | SPHK1 | 8877 | 5.55799055161883 | 0.81475801 | 1 | only_DIvsD7 | DownUp | notDM | Gain | 5 |
| **17_7477152_7484021_6665** | 17_7477152_7484021 | SOX15 | 6665 | 3.97173959088778 | -0.25833955 | 1 | only_DIvsD7 | DownUp | notDM | Loss | 8 |
| **17_7485604_7487060_6665** | 17_7485604_7487060 | SOX15 | 6665 | 3.80147231352147 | -0.40594743 | 5 | only_DIvsD7 | DownUp | notDM | Gain | 1 |
| **17_7487432_7489038_6665** | 17_7487432_7489038 | SOX15 | 6665 | 3.6387886671574 | -0.40594743 | 1 | only_DIvsD7 | DownUp | notDM | Loss | 7 |
| **17_7493651_7495210_6665** | 17_7493651_7495210 | SOX15 | 6665 | 2.41995574848976 | 0.8114564 | 3 | only_DIvsD7 | DownUp | notDM | Loss | 5 |
| **17_76030659_76031648_147138** | 17_76030659_76031648 | TMC8 | 147138 | 4.9786916879365 | -0.73395862 | 3 | only_DIvsD7 | DownUp | notDM | Gain | 1 |

| | | | | | | | | | | | |
|---|---|---|---|---|---|---|---|---|---|---|---|
| **17_76036257_76038563_147138** | 17_76036257_76038563 | TMC8 | 147138 | 4.945945948135 | -0.70222345 | 1 | only_DIvsD7 | DownUp | notDM | Gain | 2 |
| **17_76123279_76124825_147138** | 17_76123279_76124825 | TMC8 | 147138 | 3.30856441356124 | 0.00625908 | 1 | only_DIvsD7 | DownUp | notDM | Loss | 3 |
| **17_76124901_76125930_147138** | 17_76124901_76125930 | TMC8 | 147138 | 2.96894968098134 | 0.30141591 | 4 | only_DIvsD7 | DownUp | notDM | Loss | 3 |
| **17_76135789_76136645_147138** | 17_76135789_76136645 | TMC8 | 147138 | 3.95090008936639 | 0.3820245 | 4 | only_DIvsD7 | DownUp | notDM | Loss | 3 |
| **17_76136811_76139106_147138** | 17_76136811_76139106 | TMC8 | 147138 | 3.9979540040694 | 0.25321208 | 4 | only_DIvsD7 | DownUp | notDM | Gain | 1 |
| **17_76139308_76141920_147138** | 17_76139308_76141920 | TMC8 | 147138 | 4.09516935143176 | 0.44762593 | 3 | only_DIvsD7 | DownUp | notDM | Loss | 5 |
| **17_76146206_76148793_147138** | 17_76146206_76148793 | TMC8 | 147138 | 4.28665852461569 | 0.50434457 | 3 | only_DIvsD7 | DownUp | notDM | Gain | 1 |
| **17_76167405_76169780_147138** | 17_76167405_76169780 | TMC8 | 147138 | 4.60795872706398 | 0.58071807 | 6 | only_DIvsD7 | DownUp | notDM | Loss | 5 |
| **17_76172903_76173642_147138** | 17_76172903_76173642 | TMC8 | 147138 | 4.66319190916089 | 0.74508764 | 6 | only_DIvsD7 | DownUp | notDM | Loss | 5 |
| **17_76180005_76183148_11322** | 17_76180005_76183148 | TMC6 | 11322 | 4.74151401648738 | 0.57967138 | 1 | notSpecific | Up | DM | Loss | 10 |
| **17_76180005_76183148_125061** | 17_76180005_76183148 | AFMID | 125061 | 2.39967372148104 | 0.52235032 | 1 | notSpecific | Up | DM | Loss | 10 |
| **17_76180005_76183148_7083** | 17_76180005_76183148 | TK1 | 7083 | 0 | -0.24923928 | 1 | notSpecific | Up | DM | Loss | 10 |
| **17_76180005_76183148_9144** | 17_76180005_76183148 | SYNGR2 | 9144 | 4.18571209987001 | -0.22865576 | 1 | notSpecific | Up | DM | Loss | 10 |
| **17_7619174_7621975_6665** | 17_7619174_7621975 | SOX15 | 6665 | 5.09963230676034 | 0.98838637 | 1 | only_DIvsD7 | DownUp | notDM | Loss | 6 |
| **17_76345668_76348980_147138** | 17_76345668_76348980 | TMC8 | 147138 | 5.34006915093022 | -0.74939184 | 7 | only_DIvsD7 | DownUp | notDM | Loss | 7 |
| **17_76349263_76350752_147138** | 17_76349263_76350752 | TMC8 | 147138 | 5.34714649932972 | -0.75387963 | 7 | only_DIvsD7 | DownUp | notDM | Loss | 7 |
| **17_7745598_7749245_1107** | 17_7745598_7749245 | CHD3 | 1107 | 4.58972625625424 | 0.48073859 | 1 | notSpecific | Down | DM | Loss | 10 |
| **17_7745598_7749245_124637** | 17_7745598_7749245 | CYB5D1 | 124637 | 4.08167128729092 | 0.2637475 | 1 | notSpecific | Up | DM | Loss | 10 |
| **17_7745598_7749245_23135** | 17_7745598_7749245 | KDM6B | 23135 | 3.37621185028267 | 0.00054199 | 1 | notSpecific | Up | DM | Loss | 10 |
| **17_7745598_7749245_84316** | 17_7745598_7749245 | NAA38 | 84316 | 4.06209289263017 | 0.75117682 | 1 | notSpecific | Up | DM | Loss | 10 |
| **17_7745598_7749245_92162** | 17_7745598_7749245 | TMEM88 | 92162 | 3.96094619573383 | 0.07851032 | 1 | notSpecific | Up | DM | Loss | 10 |
| **17_7745598_7749245_968** | 17_7745598_7749245 | CD68 | 968 | 5.41942370096004 | 0.77698839 | 1 | notSpecific | Up | DM | Loss | 10 |
| **17_78234866_78238482_57674** | 17_78234866_78238482 | RNF213 | 57674 | 2.30319605742049 | -0.27664421 | 1 | notSpecific | Down | DM | Loss | 10 |
| **17_79017021_79018598_124565** | 17_79017021_79018598 | SLC38A10 | 124565 | 5.39875571577337 | 0.89395168 | 6 | only_H3vsUI | DownUp | notDM | Loss | 5 |
| **17_7919070_7921013_6665** | 17_7919070_7921013 | SOX15 | 6665 | 5.62908528638585 | 0.82668866 | 2 | only_DIvsD7 | DownUp | notDM | Gain | 1 |
| **17_79211002_79212694_124565** | 17_79211002_79212694 | SLC38A10 | 124565 | 4.1264561134318 | -0.19210304 | 2 | only_H3vsUI | DownUp | notDM | Loss | 7 |
| **17_79213355_79215503_124565** | 17_79213355_79215503 | SLC38A10 | 124565 | 4.02411607282677 | -0.24116147 | 1 | only_H3vsUI | DownUp | notDM | Loss | 7 |
| **17_79267037_79269050_124565** | 17_79267037_79269050 | SLC38A10 | 124565 | 1.7481880270062 | -0.5887644 | 1 | only_H3vsUI | DownUp | notDM | Gain | 4 |
| **17_79269223_79270659_124565** | 17_79269223_79270659 | SLC38A10 | 124565 | 2.07918124604762 | -0.09312733 | 1 | only_H3vsUI | DownUp | notDM | Gain | 4 |
| **17_79281480_79284092_124565** | 17_79281480_79284092 | SLC38A10 | 124565 | 4.09394672389058 | -0.09153206 | 1 | only_H3vsUI | DownUp | notDM | Gain | 5 |
| **17_79301537_79305217_284186** | 17_79301537_79305217 | TMEM105 | 284186 | 0 | -0.00479055 | 5 | only_H3vsUI | UpDown | notDM | Gain | 4 |
| **17_79310316_79312003_284186** | 17_79310316_79312003 | TMEM105 | 284186 | 3.76671020726226 | 0.02131167 | 6 | only_H3vsUI | UpDown | notDM | Gain | 1 |
| **17_79312143_79314992_124565** | 17_79312143_79314992 | SLC38A10 | 124565 | 4.6342555319257 | 0.79392111 | 7 | only_H3vsUI | DownUp | notDM | Loss | 5 |
| **17_79312143_79314992_284186** | 17_79312143_79314992 | TMEM105 | 284186 | 3.884851982746 | 0.01781066 | 7 | only_H3vsUI | UpDown | notDM | Loss | 5 |

| | | | | | | | | | | | |
|---|---|---|---|---|---|---|---|---|---|---|---|
| **17_79361393_79362052_124565** | 17_79361393_79362052 | SLC38A10 | 124565 | 4.96533342801842 | 0.93221945 | 4 | only_H3vsUI | DownUp | notDM | Loss | 3 |
| **17_79466667_79467877_348262** | 17_79466667_79467877 | MCRIP1 | 348262 | 5.50959361458761 | -0.86378312 | 6 | only_H3vsUI | UpDown | notDM | Loss | 7 |
| **17_79541737_79546323_124565** | 17_79541737_79546323 | SLC38A10 | 124565 | 5.43564054362618 | 0.94888128 | 7 | only_H3vsUI | DownUp | notDM | Gain | 5 |
| **17_79541737_79546323_5986** | 17_79541737_79546323 | RFNG | 5986 | 5.66595321787074 | 0.90959299 | 7 | only_H3vsUI | UpDown | notDM | Gain | 5 |
| **17_79601501_79603782_5986** | 17_79601501_79603782 | RFNG | 5986 | 5.60838588146491 | 0.91266131 | 1 | only_H3vsUI | UpDown | notDM | Loss | 6 |
| **17_79604217_79605207_5986** | 17_79604217_79605207 | RFNG | 5986 | 5.60693248327225 | 0.89505867 | 1 | only_H3vsUI | UpDown | notDM | Gain | 1 |
| **17_79618262_79619298_348262** | 17_79618262_79619298 | MCRIP1 | 348262 | 5.23520007696928 | -0.69726118 | 6 | only_H3vsUI | UpDown | notDM | Loss | 7 |
| **17_79679453_79681475_348262** | 17_79679453_79681475 | MCRIP1 | 348262 | 5.04017891418938 | -0.64442993 | 1 | only_H3vsUI | UpDown | notDM | Loss | 7 |
| **17_79702222_79703469_348262** | 17_79702222_79703469 | MCRIP1 | 348262 | 4.94299464129126 | -0.65407106 | 4 | only_H3vsUI | UpDown | notDM | Loss | 7 |
| **17_79778941_79779753_348262** | 17_79778941_79779753 | MCRIP1 | 348262 | 4.05747591582625 | -0.77484936 | 6 | only_H3vsUI | UpDown | notDM | Loss | 2 |
| **17_79789419_79790877_348262** | 17_79789419_79790877 | MCRIP1 | 348262 | 2.46389298898591 | -0.92359767 | 1 | only_H3vsUI | UpDown | notDM | Loss | 8 |
| **17_79791257_79792696_348262** | 17_79791257_79792696 | MCRIP1 | 348262 | 1.96378782734556 | -0.81912676 | 1 | only_H3vsUI | UpDown | notDM | Loss | 8 |
| **17_79816094_79818107_348262** | 17_79816094_79818107 | MCRIP1 | 348262 | 4.39670485759754 | -0.52157442 | 1 | only_H3vsUI | UpDown | notDM | Loss | 7 |
| **17_79826722_79828820_348262** | 17_79826722_79828820 | MCRIP1 | 348262 | 4.55092511173554 | -0.50093824 | 1 | only_H3vsUI | UpDown | notDM | Loss | 7 |
| **17_79829306_79830797_348262** | 17_79829306_79830797 | MCRIP1 | 348262 | 4.58139207541131 | -0.67465184 | 1 | only_H3vsUI | UpDown | notDM | Gain | 2 |
| **17_79850012_79851370_348262** | 17_79850012_79851370 | MCRIP1 | 348262 | 4.76972432755914 | -0.87225541 | 1 | only_H3vsUI | UpDown | notDM | Loss | 7 |
| **17_79867498_79869034_348262** | 17_79867498_79869034 | MCRIP1 | 348262 | 4.88271233113363 | -0.68057794 | 1 | only_H3vsUI | UpDown | notDM | Loss | 7 |
| **17_79873777_79875782_5986** | 17_79873777_79875782 | RFNG | 5986 | 5.12690380964086 | -0.67827565 | 1 | only_H3vsUI | UpDown | notDM | Loss | 7 |
| **17_79925996_79926698_5986** | 17_79925996_79926698 | RFNG | 5986 | 4.91883209692923 | 0.63706205 | 6 | only_H3vsUI | UpDown | notDM | Gain | 1 |
| **17_79934035_79935368_5986** | 17_79934035_79935368 | RFNG | 5986 | 4.87129265513479 | 0.59506282 | 2 | only_H3vsUI | UpDown | notDM | Loss | 6 |
| **17_79935805_79937201_5986** | 17_79935805_79937201 | RFNG | 5986 | 4.8604518067655 | 0.59506282 | 1 | only_H3vsUI | UpDown | notDM | Loss | 6 |
| **17_79959956_79961563_5986** | 17_79959956_79961563 | RFNG | 5986 | 4.68265942417943 | 0.78690618 | 7 | only_H3vsUI | UpDown | notDM | Gain | 5 |
| **17_79961724_79962794_5986** | 17_79961724_79962794 | RFNG | 5986 | 4.67141353627164 | 0.80449905 | 7 | only_H3vsUI | UpDown | notDM | Loss | 1 |
| **17_79963514_79964901_5986** | 17_79963514_79964901 | RFNG | 5986 | 4.65079303965193 | 0.61915689 | 6 | only_H3vsUI | UpDown | notDM | Loss | 1 |
| **17_79988447_79989373_348262** | 17_79988447_79989373 | MCRIP1 | 348262 | 5.29508746205255 | -0.78370791 | 2 | only_H3vsUI | UpDown | notDM | Loss | 7 |
| **17_79988447_79989373_5986** | 17_79988447_79989373 | RFNG | 5986 | 4.30850038508503 | -0.48572709 | 2 | only_H3vsUI | UpDown | notDM | Loss | 7 |
| **17_79989890_79990610_5986** | 17_79989890_79990610 | RFNG | 5986 | 4.28126068705501 | -0.41133885 | 2 | only_H3vsUI | UpDown | notDM | Loss | 3 |
| **17_79994146_79996560_5986** | 17_79994146_79996560 | RFNG | 5986 | 4.11925588927794 | -0.44405604 | 2 | only_H3vsUI | UpDown | notDM | Loss | 8 |
| **17_80008225_80009379_5986** | 17_80008225_80009379 | RFNG | 5986 | 2.4345689040342 | 0.39827523 | 1 | only_H3vsUI | UpDown | notDM | Loss | 3 |
| **17_80010092_80012001_5986** | 17_80010092_80012001 | RFNG | 5986 | 2.57403126772772 | 0.37894567 | 1 | only_H3vsUI | UpDown | notDM | Loss | 6 |
| **17_80021678_80023462_5986** | 17_80021678_80023462 | RFNG | 5986 | 4.07776749038206 | 0.39201169 | 1 | only_H3vsUI | UpDown | notDM | Loss | 6 |
| **17_80023766_80024677_5986** | 17_80023766_80024677 | RFNG | 5986 | 4.14764541250174 | 0.44209052 | 1 | only_H3vsUI | UpDown | notDM | Loss | 3 |
| **17_80223416_80230706_5986** | 17_80223416_80230706 | RFNG | 5986 | 5.32994269402731 | 0.97043054 | 1 | only_H3vsUI | UpDown | notDM | Loss | 6 |
| **17_80231590_80232731_5986** | 17_80231590_80232731 | RFNG | 5986 | 5.34610445561436 | 0.76783269 | 4 | only_H3vsUI | UpDown | notDM | Loss | 3 |

| | | | | | | | | | | | |
|---|---|---|---|---|---|---|---|---|---|---|---|
| **17_8024956_8029850_1107** | 17_8024956_8029850 | CHD3 | 1107 | 5.37444405004593 | 0.73628912 | 2 | notSpecific | Down | DM | Loss | 10 |
| **17_8024956_8029850_5187** | 17_8024956_8029850 | PER1 | 5187 | 4.47463869413439 | 0.6517617 | 2 | notSpecific | Up | DM | Loss | 10 |
| **17_8024956_8029850_59344** | 17_8024956_8029850 | ALOXE3 | 59344 | 3.498448403174 | 0.19913055 | 2 | notSpecific | Up | DM | Loss | 10 |
| **17_8024956_8029850_84314** | 17_8024956_8029850 | TMEM107 | 84314 | 4.69709851146429 | 0.73065226 | 2 | notSpecific | Down | DM | Loss | 10 |
| **17_8024956_8029850_84667** | 17_8024956_8029850 | HES7 | 84667 | 0 | 0.45447847 | 2 | notSpecific | Up | DM | Loss | 10 |
| **17_80278829_80279735_5986** | 17_80278829_80279735 | RFNG | 5986 | 5.43004440218505 | 0.93965902 | 4 | only_H3vsUI | UpDown | notDM | Loss | 5 |
| **17_80287788_80289126_5986** | 17_80287788_80289126 | RFNG | 5986 | 5.44426345031715 | 0.94038486 | 6 | only_H3vsUI | UpDown | notDM | Loss | 5 |
| **17_8036922_8038331_5198** | 17_8036922_8038331 | PFAS | 5198 | 5.05794742951173 | 0.67733501 | 6 | only_H3vsUI | Down | notDM | Loss | 5 |
| **17_8061892_8062470_5198** | 17_8061892_8062470 | PFAS | 5198 | 4.95489346306892 | -0.66967924 | 6 | only_H3vsUI | Down | notDM | Loss | 7 |
| **17_8094750_8097704_5198** | 17_8094750_8097704 | PFAS | 5198 | 4.73965144370938 | 0.58383186 | 4 | only_H3vsUI | Down | notDM | Loss | 4 |
| **17_81038782_81039377_284207** | 17_81038782_81039377 | METRNL | 284207 | 3.08529057823006 | -0.00661568 | 2 | notSpecific | Up | DM | Loss | 10 |
| **17_8129234_8129923_5198** | 17_8129234_8129923 | PFAS | 5198 | 4.35568134610572 | -0.30753623 | 5 | only_H3vsUI | Down | notDM | Gain | 2 |
| **17_8130067_8131073_5198** | 17_8130067_8131073 | PFAS | 5198 | 4.33308437114365 | -0.41716588 | 1 | only_H3vsUI | Down | notDM | Loss | 7 |
| **17_8148953_8151292_5198** | 17_8148953_8151292 | PFAS | 5198 | 3.11826472608948 | -0.1251653 | 1 | only_H3vsUI | Down | notDM | Gain | 4 |
| **17_8151504_8155049_5198** | 17_8151504_8155049 | PFAS | 5198 | 0 | 0.50171507 | 1 | only_H3vsUI | Down | notDM | Loss | 8 |
| **17_8197461_8198713_22899** | 17_8197461_8198713 | ARHGEF15 | 22899 | 4.17163870853082 | 0.28197882 | 1 | only_H3vsUI | DownUp | notDM | Gain | 4 |
| **17_8197461_8198713_5198** | 17_8197461_8198713 | PFAS | 5198 | 4.65184958803156 | -0.57959356 | 1 | only_H3vsUI | Down | notDM | Gain | 2 |
| **17_8648655_8650165_146849** | 17_8648655_8650165 | CCDC42 | 146849 | 2.70156798505593 | 0 | 2 | only_DIvsD7 | UpDown | notDM | Gain | 4 |
| **17_9738694_9744268_9340** | 17_9738694_9744268 | GLP2R | 9340 | 3.20547503674089 | 0.77539892 | 7 | notSpecific | DownUp | DM | Gain | 9 |
| **18_29671293_29671799_51444** | 18_29671293_29671799 | RNF138 | 51444 | 1.47712125471966 | 0.15417454 | 5 | notSpecific | Up | DM | Gain | 9 |
| **18_2977908_2986972_9663** | 18_2977908_2986972 | LPIN2 | 9663 | 0 | 0.10493686 | 7 | only_H3vsUI | Down | notDM | Gain | 4 |
| **18_3011561_3012526_9663** | 18_3011561_3012526 | LPIN2 | 9663 | 2.89652621748956 | 0.18588092 | 2 | only_H3vsUI | Down | notDM | Gain | 4 |
| **18_3014679_3015735_9663** | 18_3014679_3015735 | LPIN2 | 9663 | 3.1360860973841 | 0.1662118 | 6 | only_H3vsUI | Down | notDM | Gain | 1 |
| **18_34409241_34410686_57536** | 18_34409241_34410686 | KIAA1328 | 57536 | 2.21218760440396 | 0.15187257 | 1 | only_DIvsD7 | UpDown | notDM | Gain | 4 |
| **18_3452677_3460289_7050** | 18_3452677_3460289 | TGIF1 | 7050 | 3.001733712809 | -0.21476593 | 5 | notSpecific | Down | DM | Gain | 9 |
| **18_47814717_47815167_30827** | 18_47814717_47815167 | CXXC1 | 30827 | 1.43136376415899 | 0.18156325 | 2 | notSpecific | UpDown | DM | Gain | 9 |
| **18_47814717_47815167_4152** | 18_47814717_47815167 | MBD1 | 4152 | 3.83129374437701 | 0.24049394 | 2 | notSpecific | Down | DM | Gain | 9 |
| **18_61090013_61091098_5055** | 18_61090013_61091098 | SERPINB2 | 5055 | 5.66642062777761 | 0.80852153 | 1 | only_H3vsUI | UpDown | notDM | Gain | 1 |
| **18_61603645_61604160_5055** | 18_61603645_61604160 | SERPINB2 | 5055 | 4.68760029736491 | 0.91106559 | 2 | only_H3vsUI | UpDown | notDM | Loss | 3 |
| **18_61615679_61616808_5055** | 18_61615679_61616808 | SERPINB2 | 5055 | 4.78348908746147 | 0.73362069 | 2 | only_H3vsUI | UpDown | notDM | Loss | 3 |
| **18_71815212_71815554_201456** | 18_71815212_71815554 | FBXO15 | 201456 | 2.17026171539496 | -0.10873051 | 5 | only_H3vsUI | DownUp | notDM | Loss | 3 |
| **18_71816058_71817943_201456** | 18_71816058_71817943 | FBXO15 | 201456 | 2.99738638439731 | -0.09610673 | 1 | only_H3vsUI | DownUp | notDM | Gain | 4 |
| **18_74766155_74767052_4155** | 18_74766155_74767052 | MBP | 4155 | 4.51207040139603 | -0.46763952 | 7 | only_H3vsUI | UpDown | notDM | Gain | 1 |
| **18_74767231_74768949_4155** | 18_74767231_74768949 | MBP | 4155 | 4.48596263419781 | -0.46499238 | 7 | only_H3vsUI | UpDown | notDM | Gain | 1 |

| | | | | | | | | | | | |
|---|---|---|---|---|---|---|---|---|---|---|---|
| **18_74776416_74782065_4155** | 18_74776416_74782065 | MBP | 4155 | 4.24306286480481 | 0.45418349 | 7 | only_H3vsUI | UpDown | notDM | Gain | 1 |
| **18_74782881_74784002_4155** | 18_74782881_74784002 | MBP | 4155 | 4.19212122213171 | 0.45342631 | 7 | only_H3vsUI | UpDown | notDM | Loss | 5 |
| **18_74813024_74815450_4155** | 18_74813024_74815450 | MBP | 4155 | 4.12907732425274 | 0.60780489 | 7 | only_H3vsUI | UpDown | notDM | Loss | 5 |
| **18_74825069_74827955_4155** | 18_74825069_74827955 | MBP | 4155 | 4.40664235535077 | 0.60705392 | 7 | only_H3vsUI | UpDown | notDM | Gain | 1 |
| **18_74838287_74840274_4155** | 18_74838287_74840274 | MBP | 4155 | 3.65292288756794 | 0.86300697 | 7 | only_H3vsUI | UpDown | notDM | Gain | 1 |
| **18_74840456_74841741_4155** | 18_74840456_74841741 | MBP | 4155 | 3.4814426285023 | 0.86622991 | 7 | only_H3vsUI | UpDown | notDM | Gain | 1 |
| **18_74842042_74843697_4155** | 18_74842042_74843697 | MBP | 4155 | 3.01241537476243 | 0.83075489 | 2 | only_H3vsUI | UpDown | notDM | Gain | 4 |
| **18_77437637_77438965_284241** | 18_77437637_77438965 | LOC284241 | 284241 | 2.8926510338773 | 0.00551759 | 1 | only_DIvsD7 | DownUp | notDM | Gain | 4 |
| **18_77440376_77441724_284241** | 18_77440376_77441724 | LOC284241 | 284241 | 2.80140371001736 | 0.00551759 | 1 | only_DIvsD7 | DownUp | notDM | Loss | 3 |
| **18_77746302_77748186_79863** | 18_77746302_77748186 | RBFA | 79863 | 4.66438809289896 | -0.75899566 | 2 | only_H3vsUI | DownUp | notDM | Loss | 7 |
| **19_10146438_10148855_388503** | 19_10146438_10148855 | C3P1 | 388503 | 3.56383691866455 | 0 | 7 | only_DIvsD7 | UpDown | notDM | Gain | 1 |
| **19_10399096_10403265_3386** | 19_10399096_10403265 | ICAM4 | 3386 | 3.16076856186113 | 0.4755256 | 2 | notSpecific | Down | DM | Loss | 10 |
| **19_10399096_10403265_7087** | 19_10399096_10403265 | ICAM5 | 7087 | 0 | 0.23379095 | 2 | notSpecific | Down | DM | Loss | 10 |
| **19_1093559_1094987_148223** | 19_1093559_1094987 | C19orf25 | 148223 | 5.58457092495149 | -0.99449423 | 1 | only_H3vsUI | DownUp | notDM | Loss | 7 |
| **19_11545234_11545896_1264** | 19_11545234_11545896 | CNN1 | 1264 | 5.01608437204493 | -0.72238381 | 2 | only_H3vsUI | UpDown | notDM | Gain | 2 |
| **19_11615276_11616000_1264** | 19_11615276_11616000 | CNN1 | 1264 | 4.52723021763753 | 0.82492934 | 2 | only_H3vsUI | UpDown | notDM | Loss | 3 |
| **19_11638568_11639664_1264** | 19_11638568_11639664 | CNN1 | 1264 | 4.00021709297223 | -0.40877401 | 1 | only_H3vsUI | UpDown | notDM | Loss | 3 |
| **19_11646115_11646990_1264** | 19_11646115_11646990 | CNN1 | 1264 | 3.42797271360821 | 0.50893208 | 1 | only_H3vsUI | UpDown | notDM | Loss | 3 |
| **19_11668496_11669780_1264** | 19_11668496_11669780 | CNN1 | 1264 | 4.27485032001666 | -0.58462541 | 1 | only_H3vsUI | UpDown | notDM | Gain | 4 |
| **19_11848187_11849334_1264** | 19_11848187_11849334 | CNN1 | 1264 | 5.29780645418088 | -0.82880082 | 2 | only_H3vsUI | UpDown | notDM | Gain | 2 |
| **19_1236972_1237683_1153** | 19_1236972_1237683 | CIRBP | 1153 | 4.50040115705547 | 0.3650948 | 2 | notSpecific | Down | DM | Loss | 10 |
| **19_1236972_1237683_255057** | 19_1236972_1237683 | CBARP | 255057 | 2.48855071650044 | 0.14596145 | 2 | notSpecific | Up | DM | Loss | 10 |
| **19_1236972_1237683_513** | 19_1236972_1237683 | ATP5F1D | 513 | 3.60927417240459 | -0.023452 | 2 | notSpecific | Up | DM | Loss | 10 |
| **19_1236972_1237683_90007** | 19_1236972_1237683 | MIDN | 90007 | 4.03622954408629 | -0.16982605 | 2 | notSpecific | Up | DM | Loss | 10 |
| **19_13042937_13044247_2193** | 19_13042937_13044247 | FARSA | 2193 | 2.45024910831936 | 0.71350436 | 2 | only_H3vsUI | DownUp | notDM | Gain | 4 |
| **19_13047938_13049109_2193** | 19_13047938_13049109 | FARSA | 2193 | 3.52917360326172 | -0.04728753 | 1 | only_H3vsUI | DownUp | notDM | Loss | 7 |
| **19_13049777_13052004_2193** | 19_13049777_13052004 | FARSA | 2193 | 3.71775369321072 | 0.2248764 | 1 | only_H3vsUI | DownUp | notDM | Loss | 7 |
| **19_13057140_13058663_2193** | 19_13057140_13058663 | FARSA | 2193 | 4.09981870961523 | -0.13850084 | 1 | only_H3vsUI | DownUp | notDM | Gain | 2 |
| **19_13136425_13138323_2193** | 19_13136425_13138323 | FARSA | 2193 | 4.96316898907749 | -0.64975668 | 6 | only_H3vsUI | DownUp | notDM | Loss | 7 |
| **19_1324486_1325939_148223** | 19_1324486_1325939 | C19orf25 | 148223 | 5.18551382421854 | -0.72593379 | 4 | only_H3vsUI | DownUp | notDM | Loss | 7 |
| **19_14017145_14018033_54862** | 19_14017145_14018033 | CC2D1A | 54862 | 2.28103336724773 | -0.0124277 | 1 | only_DIvsD7 | UpDown | notDM | Loss | 3 |
| **19_14168618_14169987_342979** | 19_14168618_14169987 | PALM3 | 342979 | 0 | 0.00441306 | 4 | only_H3vsUI | UpDown | notDM | Loss | 3 |
| **19_1477615_1478800_148223** | 19_1477615_1478800 | C19orf25 | 148223 | 2.60097289568675 | -0.56022563 | 1 | only_H3vsUI | DownUp | notDM | Gain | 4 |
| **19_1477615_1478800_54760** | 19_1477615_1478800 | PCSK4 | 54760 | 4.06228106997264 | -0.11241313 | 1 | only_H3vsUI | Up | notDM | Gain | 4 |

| | | | | | | | | | | | |
|---|---|---|---|---|---|---|---|---|---|---|---|
| **19_1489467_1490435_148223** | 19_1489467_1490435 | C19orf25 | 148223 | 4.01161272921942 | -0.62562668 | 2 | only_H3vsUI | DownUp | notDM | Loss | 7 |
| **19_1489467_1490435_54760** | 19_1489467_1490435 | PCSK4 | 54760 | 0 | -0.10785657 | 2 | only_H3vsUI | Up | notDM | Loss | 3 |
| **19_15541715_15543048_23581** | 19_15541715_15543048 | CASP14 | 23581 | 5.58141029351248 | 0.82284961 | 1 | only_DIvsD7 | UpDown | notDM | Loss | 3 |
| **19_1589729_1592178_148223** | 19_1589729_1592178 | C19orf25 | 148223 | 5.04349195746275 | -0.8233717 | 1 | only_H3vsUI | DownUp | notDM | Loss | 7 |
| **19_1593117_1593469_148223** | 19_1593117_1593469 | C19orf25 | 148223 | 5.05660378855205 | -0.8233717 | 1 | only_H3vsUI | DownUp | notDM | Loss | 7 |
| **19_17189464_17191465_84705** | 19_17189464_17191465 | GTPBP3 | 84705 | 5.40975057989907 | -0.81524529 | 5 | only_H3vsUI | DownUp | notDM | Loss | 7 |
| **19_17349959_17355582_84705** | 19_17349959_17355582 | GTPBP3 | 84705 | 4.96760881062458 | 0.6268136 | 1 | only_H3vsUI | DownUp | notDM | Loss | 6 |
| **19_17414399_17416293_84705** | 19_17414399_17416293 | GTPBP3 | 84705 | 4.50601769985113 | 0.6026597 | 3 | only_H3vsUI | DownUp | notDM | Loss | 5 |
| **19_17422114_17424138_84705** | 19_17422114_17424138 | GTPBP3 | 84705 | 4.38415620720308 | 0.4866083 | 5 | only_H3vsUI | DownUp | notDM | Loss | 5 |
| **19_17446994_17448171_84705** | 19_17446994_17448171 | GTPBP3 | 84705 | 2.26951294421792 | 0.31502912 | 1 | only_H3vsUI | DownUp | notDM | Gain | 4 |
| **19_17448398_17449673_84705** | 19_17448398_17449673 | GTPBP3 | 84705 | 1.64345267648619 | 0.31502912 | 1 | only_H3vsUI | DownUp | notDM | Loss | 3 |
| **19_17529498_17530557_84705** | 19_17529498_17530557 | GTPBP3 | 84705 | 4.90905298331726 | 0.57107667 | 1 | only_H3vsUI | DownUp | notDM | Gain | 1 |
| **19_17581722_17582276_84705** | 19_17581722_17582276 | GTPBP3 | 84705 | 5.12492787911053 | -0.76300536 | 2 | only_H3vsUI | DownUp | notDM | Gain | 2 |
| **19_17622789_17623882_84705** | 19_17622789_17623882 | GTPBP3 | 84705 | 5.24153900987033 | 0.74460811 | 1 | only_H3vsUI | DownUp | notDM | Loss | 7 |
| **19_17940468_17941568_3718** | 19_17940468_17941568 | JAK3 | 3718 | 4.23701562874516 | -0.64981472 | 4 | only_H3vsUI | DownUp | notDM | Loss | 3 |
| **19_17953167_17953881_3718** | 19_17953167_17953881 | JAK3 | 3718 | 3.69425411202528 | -0.67000089 | 2 | only_H3vsUI | DownUp | notDM | Loss | 3 |
| **19_17971140_17973862_3718** | 19_17971140_17973862 | JAK3 | 3718 | 4.08994041849863 | -0.22636602 | 2 | only_H3vsUI | DownUp | notDM | Loss | 8 |
| **19_18057956_18058542_3718** | 19_18057956_18058542 | JAK3 | 3718 | 4.99621386821002 | -0.75697501 | 2 | only_H3vsUI | DownUp | notDM | Gain | 2 |
| **19_18078063_18078381_3718** | 19_18078063_18078381 | JAK3 | 3718 | 5.07636368844452 | 0.70140653 | 4 | only_H3vsUI | DownUp | notDM | Loss | 3 |
| **19_18219322_18220747_3718** | 19_18219322_18220747 | JAK3 | 3718 | 5.41577938503628 | 0.81430163 | 4 | only_H3vsUI | DownUp | notDM | Loss | 3 |
| **19_1854062_1855390_83855** | 19_1854062_1855390 | KLF16 | 83855 | 3.91248776133232 | -0.11427358 | 5 | only_H3vsUI | Up | notDM | Loss | 1 |
| **19_1855600_1856747_83855** | 19_1855600_1856747 | KLF16 | 83855 | 3.83365699689287 | -0.16604366 | 7 | only_H3vsUI | Up | notDM | Loss | 1 |
| **19_1859606_1862585_148223** | 19_1859606_1862585 | C19orf25 | 148223 | 5.5802519248998 | -0.84797098 | 1 | only_H3vsUI | DownUp | notDM | Loss | 7 |
| **19_1859606_1862585_83855** | 19_1859606_1862585 | KLF16 | 83855 | 2.99122607569249 | 0.36628673 | 1 | only_H3vsUI | Up | notDM | Loss | 6 |
| **19_1873832_1875783_83855** | 19_1873832_1875783 | KLF16 | 83855 | 4.01157044359728 | -0.04171559 | 1 | only_H3vsUI | Up | notDM | Loss | 7 |
| **19_19301501_19302627_8625** | 19_19301501_19302627 | RFXANK | 8625 | 3.03901732199741 | 0.59951915 | 2 | only_H3vsUI | UpDown | notDM | Loss | 3 |
| **19_19303775_19304695_8625** | 19_19303775_19304695 | RFXANK | 8625 | 1.75587485567249 | 0.30668624 | 1 | only_H3vsUI | UpDown | notDM | Loss | 3 |
| **19_19312930_19314223_8625** | 19_19312930_19314223 | RFXANK | 8625 | 3.96435392929219 | -0.61411055 | 1 | only_H3vsUI | UpDown | notDM | Loss | 8 |
| **19_19431973_19433360_57130** | 19_19431973_19433360 | ATP13A1 | 57130 | 5.53291100242215 | -0.88150407 | 1 | only_H3vsUI | UpDown | notDM | Loss | 7 |
| **19_19495318_19496079_8625** | 19_19495318_19496079 | RFXANK | 8625 | 5.28227535447285 | 0.91242031 | 1 | only_H3vsUI | UpDown | notDM | Loss | 3 |
| **19_19772600_19774181_57130** | 19_19772600_19774181 | ATP13A1 | 57130 | 2.48000694295715 | 0.36213147 | 1 | only_H3vsUI | UpDown | notDM | Loss | 3 |
| **19_23868445_23869893_171392** | 19_23868445_23869893 | ZNF675 | 171392 | 2.09691001300806 | 0.23058206 | 1 | only_H3vsUI | Down | notDM | Loss | 3 |
| **19_2425871_2427321_4616** | 19_2425871_2427321 | GADD45B | 4616 | 4.68844651961336 | -0.71447449 | 1 | only_DIvsD7 | Up | notDM | Loss | 7 |
| **19_2477760_2479652_4616** | 19_2477760_2479652 | GADD45B | 4616 | 3.2145789535705 | 0.89530754 | 2 | only_DIvsD7 | Up | notDM | Loss | 6 |

| | | | | | | | | | | | |
|---|---|---|---|---|---|---|---|---|---|---|---|
| **19_2781328_2783036_4616** | 19_2781328_2783036 | GADD45B | 4616 | 5.48459449005123 | -0.93294004 | 1 | only_DIvsD7 | Up | notDM | Gain | 2 |
| **19_2783646_2784219_4616** | 19_2783646_2784219 | GADD45B | 4616 | 5.48788042717067 | -0.75916422 | 7 | only_DIvsD7 | Up | notDM | Gain | 2 |
| **19_30301292_30302449_898** | 19_30301292_30302449 | CCNE1 | 898 | 2.66558099101795 | -0.01985476 | 1 | only_H3vsUI | DownUp | notDM | Gain | 4 |
| **19_30303402_30304502_898** | 19_30303402_30304502 | CCNE1 | 898 | 2.70156798505593 | 0.14463803 | 2 | only_H3vsUI | DownUp | notDM | Loss | 8 |
| **19_33060675_33061577_91442** | 19_33060675_33061577 | FAAP24 | 91442 | 5.6037634235856 | -0.95366331 | 4 | only_H3vsUI | DownUp | notDM | Gain | 2 |
| **19_33460543_33462670_84902** | 19_33460543_33462670 | CEP89 | 84902 | 2.30102999566398 | 0.78825182 | 1 | only_H3vsUI | DownUp | notDM | Loss | 6 |
| **19_33460543_33462670_91442** | 19_33460543_33462670 | FAAP24 | 91442 | 2.68033551341456 | -0.13081877 | 1 | only_H3vsUI | DownUp | notDM | Loss | 8 |
| **19_33463181_33464283_84902** | 19_33463181_33464283 | CEP89 | 84902 | 2.49692964807321 | -0.42870856 | 1 | only_H3vsUI | DownUp | notDM | Gain | 4 |
| **19_33463181_33464283_91442** | 19_33463181_33464283 | FAAP24 | 91442 | 1.54406804435028 | -0.24036567 | 1 | only_H3vsUI | DownUp | notDM | Gain | 4 |
| **19_33467362_33469711_84902** | 19_33467362_33469711 | CEP89 | 84902 | 3.65272969606925 | -0.53094192 | 3 | only_H3vsUI | DownUp | notDM | Gain | 5 |
| **19_33467362_33469711_91442** | 19_33467362_33469711 | FAAP24 | 91442 | 3.62490060220449 | 0.05170667 | 3 | only_H3vsUI | DownUp | notDM | Gain | 5 |
| **19_33552313_33554682_84902** | 19_33552313_33554682 | CEP89 | 84902 | 4.95156092381038 | -0.59566059 | 1 | only_H3vsUI | DownUp | notDM | Gain | 2 |
| **19_33668219_33669753_84902** | 19_33668219_33669753 | CEP89 | 84902 | 5.3124989369668 | 0.84050348 | 2 | only_H3vsUI | DownUp | notDM | Gain | 5 |
| **19_33793016_33793349_1050** | 19_33793016_33793349 | CEBPA | 1050 | 2.08635983067475 | 0.32938027 | 4 | notSpecific | Down | DM | Loss | 10 |
| **19_33793016_33793349_1054** | 19_33793016_33793349 | CEBPG | 1054 | 4.85285191247802 | -0.80330457 | 4 | notSpecific | Down | DM | Loss | 10 |
| **19_35417408_35419007_90075** | 19_35417408_35419007 | ZNF30 | 90075 | 0 | 0.12535036 | 2 | only_H3vsUI | Down | notDM | Gain | 4 |
| **19_35464283_35465176_90075** | 19_35464283_35465176 | ZNF30 | 90075 | 4.66724743161008 | -0.47970418 | 1 | only_H3vsUI | Down | notDM | Gain | 2 |
| **19_35521550_35522796_5349** | 19_35521550_35522796 | FXYD3 | 5349 | 4.9261931965756 | -0.59807644 | 1 | only_H3vsUI | Up | notDM | Loss | 3 |
| **19_35614540_35616306_5349** | 19_35614540_35616306 | FXYD3 | 5349 | 3.8926510338773 | 0.38359202 | 2 | only_H3vsUI | Up | notDM | Gain | 4 |
| **19_35632405_35633876_5349** | 19_35632405_35633876 | FXYD3 | 5349 | 4.40951045226932 | -0.44080731 | 4 | only_H3vsUI | Up | notDM | Loss | 3 |
| **19_35999954_36001970_374897** | 19_35999954_36001970 | SBSN | 374897 | 4.23764425763919 | 0.41135518 | 1 | only_H3vsUI | Up | notDM | Loss | 6 |
| **19_36034972_36036336_374897** | 19_36034972_36036336 | SBSN | 374897 | 4.19761132316971 | 0.44315722 | 1 | only_H3vsUI | Up | notDM | Loss | 6 |
| **19_36036557_36038162_374897** | 19_36036557_36038162 | SBSN | 374897 | 4.23922437848713 | 0.44315722 | 1 | only_H3vsUI | Up | notDM | Loss | 6 |
| **19_36133288_36136952_374897** | 19_36133288_36136952 | SBSN | 374897 | 5.05720189858267 | 0.85139885 | 2 | only_H3vsUI | Up | notDM | Loss | 6 |
| **19_36230423_36231518_374897** | 19_36230423_36231518 | SBSN | 374897 | 5.32472064518049 | -0.82013845 | 1 | only_H3vsUI | Up | notDM | Loss | 7 |
| **19_36231618_36232329_374897** | 19_36231618_36232329 | SBSN | 374897 | 5.32717086970899 | -0.91010614 | 4 | only_H3vsUI | Up | notDM | Loss | 7 |
| **19_36249161_36250876_374897** | 19_36249161_36250876 | SBSN | 374897 | 5.3616353025534 | 0.80280464 | 1 | only_H3vsUI | Up | notDM | Gain | 5 |
| **19_36263559_36266315_115703** | 19_36263559_36266315 | ARHGAP33 | 115703 | 2.01283722470517 | 0.07482631 | 2 | only_DIvsD7 | DownUp | notDM | Loss | 6 |
| **19_36263559_36266315_374897** | 19_36263559_36266315 | SBSN | 374897 | 5.38801056601461 | -0.95593222 | 2 | only_H3vsUI | Up | notDM | Loss | 7 |
| **19_36704026_36705453_147929** | 19_36704026_36705453 | ZNF565 | 147929 | 2.05690485133647 | 0.10190575 | 1 | only_H3vsUI | DownUp | notDM | Loss | 6 |
| **19_36706283_36707808_147929** | 19_36706283_36707808 | ZNF565 | 147929 | 2.47567118832443 | 0.03750827 | 1 | only_H3vsUI | DownUp | notDM | Loss | 6 |
| **19_36868526_36869988_57677** | 19_36868526_36869988 | ZFP14 | 57677 | 1.95904139232109 | -0.1111033 | 1 | only_H3vsUI | Down | notDM | Gain | 4 |
| **19_39809908_39811327_9535** | 19_39809908_39811327 | GMFG | 9535 | 4.18752072083646 | 0.42078292 | 2 | only_H3vsUI | UpDown | notDM | Loss | 6 |
| **19_39830756_39832626_9535** | 19_39830756_39832626 | GMFG | 9535 | 3.60552052343747 | 0.7544968 | 1 | only_H3vsUI | UpDown | notDM | Loss | 6 |

| | | | | | | | | | | | |
|---|---|---|---|---|---|---|---|---|---|---|---|
| **19_39833413_39835440_9535** | 19_39833413_39835440 | GMFG | 9535 | 3.82536119595263 | 0.67857625 | 1 | only_H3vsUI | UpDown | notDM | Loss | 6 |
| **19_41305478_41307527_11100** | 19_41305478_41307527 | HNRNPUL1 | 11100 | 5.66547123102743 | -0.81927553 | 1 | notSpecific | Down | DM | Loss | 10 |
| **19_41305478_41307527_6626** | 19_41305478_41307527 | SNRPA | 6626 | 4.6844324061558 | -0.69298233 | 1 | notSpecific | Up | DM | Loss | 10 |
| **19_41305478_41307527_8425** | 19_41305478_41307527 | LTBP4 | 8425 | 5.26946157301525 | -0.70500133 | 1 | notSpecific | Up | DM | Loss | 10 |
| **19_41305478_41307527_92799** | 19_41305478_41307527 | SHKBP1 | 92799 | 5.34716602599351 | -0.72798221 | 1 | notSpecific | UpDown | DM | Loss | 10 |
| **19_42719449_42721844_116115** | 19_42719449_42721844 | ZNF526 | 116115 | 3.41580772763554 | -0.48579893 | 1 | notSpecific | Down | DM | Loss | 10 |
| **19_42719449_42721844_162989** | 19_42719449_42721844 | DEDD2 | 162989 | 0 | 0.34024341 | 1 | notSpecific | Up | DM | Loss | 10 |
| **19_42719449_42721844_2931** | 19_42719449_42721844 | GSK3A | 2931 | 4.3962342287858 | 0.35454782 | 1 | notSpecific | Up | DM | Loss | 10 |
| **19_43967699_43969058_7638** | 19_43967699_43969058 | ZNF221 | 7638 | 5.68693999045008 | -0.8399304 | 1 | only_DIvsD7 | Down | notDM | Loss | 7 |
| **19_44144131_44145049_7673** | 19_44144131_44145049 | ZNF222 | 7673 | 5.58484777272776 | 0.80651935 | 2 | only_DIvsD7 | UpDown | notDM | Loss | 3 |
| **19_44255312_44258668_7673** | 19_44255312_44258668 | ZNF222 | 7673 | 5.43269195900308 | -0.80383351 | 1 | only_DIvsD7 | UpDown | notDM | Loss | 7 |
| **19_44259279_44262888_7673** | 19_44259279_44262888 | ZNF222 | 7673 | 5.42587154800906 | -0.77153438 | 1 | only_DIvsD7 | UpDown | notDM | Gain | 5 |
| **19_44437955_44440288_7638** | 19_44437955_44440288 | ZNF221 | 7638 | 4.17926446433903 | 0.71638494 | 2 | only_DIvsD7 | Down | notDM | Gain | 4 |
| **19_44437955_44440288_7766** | 19_44437955_44440288 | ZNF223 | 7766 | 5.06399724300125 | 0.63307074 | 2 | only_H3vsUI | Down | notDM | Gain | 4 |
| **19_44455439_44455959_7638** | 19_44455439_44455959 | ZNF221 | 7638 | 1.64345267648619 | -0.11832128 | 2 | only_DIvsD7 | Down | notDM | Gain | 4 |
| **19_44455439_44455959_7673** | 19_44455439_44455959 | ZNF222 | 7673 | 4.8665649624474 | -0.77148864 | 2 | only_DIvsD7 | UpDown | notDM | Gain | 2 |
| **19_44455439_44455959_7773** | 19_44455439_44455959 | ZNF230 | 7773 | 4.70858234947879 | -0.57872163 | 2 | only_H3vsUI | Down | notDM | Gain | 4 |
| **19_44506072_44506948_7638** | 19_44506072_44506948 | ZNF221 | 7638 | 4.70495656049525 | -0.55926229 | 1 | only_DIvsD7 | Down | notDM | Gain | 4 |
| **19_44506072_44506948_7673** | 19_44506072_44506948 | ZNF222 | 7673 | 4.35308876483395 | -0.29803073 | 1 | only_DIvsD7 | UpDown | notDM | Gain | 4 |
| **19_44506072_44506948_7773** | 19_44506072_44506948 | ZNF230 | 7773 | 2.11394335230684 | 0.08300896 | 1 | only_H3vsUI | Down | notDM | Gain | 4 |
| **19_44507108_44508470_7638** | 19_44507108_44508470 | ZNF221 | 7638 | 4.71374247840908 | -0.55964756 | 1 | only_DIvsD7 | Down | notDM | Gain | 4 |
| **19_44507108_44508470_7673** | 19_44507108_44508470 | ZNF222 | 7673 | 4.32273600446995 | -0.29748754 | 1 | only_DIvsD7 | UpDown | notDM | Gain | 4 |
| **19_44507108_44508470_7773** | 19_44507108_44508470 | ZNF230 | 7773 | 1.51851393987789 | 0.08414773 | 1 | only_H3vsUI | Down | notDM | Gain | 4 |
| **19_44554937_44556042_7673** | 19_44554937_44556042 | ZNF222 | 7673 | 4.40560245520931 | -0.52628544 | 2 | only_DIvsD7 | UpDown | notDM | Gain | 4 |
| **19_44554937_44556042_7766** | 19_44554937_44556042 | ZNF223 | 7766 | 2.0899051114394 | -0.03748122 | 2 | only_H3vsUI | Down | notDM | Gain | 4 |
| **19_44554937_44556042_7773** | 19_44554937_44556042 | ZNF230 | 7773 | 4.67999084243487 | -0.54220192 | 2 | only_H3vsUI | Down | notDM | Gain | 4 |
| **19_44556205_44557252_7638** | 19_44556205_44557252 | ZNF221 | 7638 | 5.00350361474254 | -0.67641355 | 1 | only_DIvsD7 | Down | notDM | Loss | 7 |
| **19_44556205_44557252_7673** | 19_44556205_44557252 | ZNF222 | 7673 | 4.4265437915681 | -0.87217397 | 1 | only_DIvsD7 | UpDown | notDM | Loss | 3 |
| **19_44556205_44557252_7766** | 19_44556205_44557252 | ZNF223 | 7766 | 1.63346845557959 | -0.55307349 | 1 | only_H3vsUI | Down | notDM | Loss | 8 |
| **19_44556205_44557252_7773** | 19_44556205_44557252 | ZNF230 | 7773 | 4.69134676413482 | -0.62990879 | 1 | only_H3vsUI | Down | notDM | Loss | 7 |
| **19_44575639_44577170_7773** | 19_44575639_44577170 | ZNF230 | 7773 | 4.83609614624381 | -0.78024252 | 2 | only_H3vsUI | Down | notDM | Gain | 4 |
| **19_44616083_44617316_7773** | 19_44616083_44617316 | ZNF230 | 7773 | 5.03745837159532 | 0.77435447 | 2 | only_H3vsUI | Down | notDM | Loss | 4 |
| **19_44860898_44861978_7766** | 19_44860898_44861978 | ZNF223 | 7766 | 5.48392376265003 | -0.81705873 | 2 | only_H3vsUI | Down | notDM | Gain | 4 |
| **19_45197068_45200114_388551** | 19_45197068_45200114 | CEACAM16 | 388551 | 3.37639444203727 | 0 | 7 | only_DIvsD7 | Up | notDM | Loss | 5 |

| | | | | | | | | | | | |
|---|---|---|---|---|---|---|---|---|---|---|---|
| **19_45429594_45431116_342** | 19_45429594_45431116 | APOC1P1 | 342 | 0 | 0.01202084 | 2 | only_H3vsUI | Up | notDM | Gain | 4 |
| **19_45980636_45982377_6253** | 19_45980636_45982377 | RTN2 | 6253 | 4.14915750623186 | 0.46459148 | 1 | only_H3vsUI | UpDown | notDM | Loss | 6 |
| **19_45988719_45989915_6253** | 19_45988719_45989915 | RTN2 | 6253 | 3.81690383937566 | 0.15077962 | 1 | only_H3vsUI | UpDown | notDM | Gain | 4 |
| **19_45995376_45997836_6253** | 19_45995376_45997836 | RTN2 | 6253 | 0 | 0.61973387 | 2 | only_H3vsUI | UpDown | notDM | Loss | 3 |
| **19_45999198_46000213_6253** | 19_45999198_46000213 | RTN2 | 6253 | 1.99563519459755 | 0.02721689 | 2 | only_H3vsUI | UpDown | notDM | Loss | 3 |
| **19_46000483_46000879_6253** | 19_46000483_46000879 | RTN2 | 6253 | 2.23552844690755 | -0.01005678 | 4 | only_H3vsUI | UpDown | notDM | Loss | 3 |
| **19_46007873_46009848_6253** | 19_46007873_46009848 | RTN2 | 6253 | 4.05694294567288 | -0.4342924 | 5 | only_H3vsUI | UpDown | notDM | Loss | 6 |
| **19_46010236_46010656_6253** | 19_46010236_46010656 | RTN2 | 6253 | 4.13874466394889 | -0.43666623 | 1 | only_H3vsUI | UpDown | notDM | Loss | 3 |
| **19_46011110_46012604_6253** | 19_46011110_46012604 | RTN2 | 6253 | 4.16548174282216 | -0.48785945 | 1 | only_H3vsUI | UpDown | notDM | Loss | 8 |
| **19_46012726_46013970_6253** | 19_46012726_46013970 | RTN2 | 6253 | 4.21096025541681 | -0.46535782 | 5 | only_H3vsUI | UpDown | notDM | Loss | 6 |
| **19_46014812_46019896_6253** | 19_46014812_46019896 | RTN2 | 6253 | 4.263399331334 | 0.66390643 | 8 | only_H3vsUI | UpDown | notDM | Loss | 5 |
| **19_46184939_46185718_6253** | 19_46184939_46185718 | RTN2 | 6253 | 5.27523531755216 | -0.90698279 | 5 | only_H3vsUI | UpDown | notDM | Loss | 7 |
| **19_46226033_46227891_6253** | 19_46226033_46227891 | RTN2 | 6253 | 5.36089810792688 | -0.76502179 | 7 | only_H3vsUI | UpDown | notDM | Gain | 1 |
| **19_46281839_46287698_147912** | 19_46281839_46287698 | SIX5 | 147912 | 3.97053282976832 | -0.06727883 | 3 | notSpecific | Down | DM | Loss | 10 |
| **19_46281839_46287698_1760** | 19_46281839_46287698 | DMPK | 1760 | 0 | 0.58728866 | 3 | notSpecific | Up | DM | Loss | 10 |
| **19_46281839_46287698_1762** | 19_46281839_46287698 | DMWD | 1762 | 3.92059286208481 | -0.35401837 | 3 | notSpecific | DownUp | DM | Loss | 10 |
| **19_46281839_46287698_339344** | 19_46281839_46287698 | MYPOP | 339344 | 5.07248885930514 | -0.64665464 | 3 | notSpecific | DownUp | DM | Loss | 10 |
| **19_46363763_46366237_3171** | 19_46363763_46366237 | FOXA3 | 3171 | 3.1078880251828 | -0.01360944 | 1 | only_DIvsD7 | DownUp | notDM | Loss | 6 |
| **19_46366574_46368312_3171** | 19_46366574_46368312 | FOXA3 | 3171 | 0 | 0.04085562 | 4 | only_DIvsD7 | DownUp | notDM | Gain | 4 |
| **19_46497544_46498369_5536** | 19_46497544_46498369 | PPP5C | 5536 | 5.54653896208893 | -0.87714742 | 2 | notSpecific | Down | DM | Gain | 9 |
| **19_46497544_46498369_729440** | 19_46497544_46498369 | CCDC61 | 729440 | 2.57749179983723 | -0.35121304 | 2 | notSpecific | Down | DM | Gain | 9 |
| **19_46497544_46498369_8189** | 19_46497544_46498369 | SYMPK | 8189 | 5.11773849100372 | -0.8454939 | 2 | notSpecific | Down | DM | Gain | 9 |
| **19_4668839_4669909_55527** | 19_4668839_4669909 | FEM1A | 55527 | 5.0857185951654 | -0.71930838 | 1 | only_H3vsUI | UpDown | notDM | Loss | 7 |
| **19_47733890_47734358_26093** | 19_47733890_47734358 | CCDC9 | 26093 | 4.4048166180754 | 0.35205918 | 2 | notSpecific | Up | DM | Loss | 10 |
| **19_47733890_47734358_27113** | 19_47733890_47734358 | BBC3 | 27113 | 1.9731278535997 | 0.21700234 | 2 | notSpecific | Up | DM | Loss | 10 |
| **19_4792294_4792992_55527** | 19_4792294_4792992 | FEM1A | 55527 | 2.75434833571102 | 0.16611997 | 2 | only_H3vsUI | UpDown | notDM | Loss | 3 |
| **19_48015160_48018217_26093** | 19_48015160_48018217 | CCDC9 | 26093 | 5.40723109549703 | -0.79436112 | 1 | notSpecific | Up | DM | Loss | 10 |
| **19_48015160_48018217_29997** | 19_48015160_48018217 | NOP53 | 29997 | 5.36281598434959 | -0.80704618 | 1 | notSpecific | Up | DM | Loss | 10 |
| **19_48015160_48018217_8775** | 19_48015160_48018217 | NAPA | 8775 | 2.44870631990508 | -0.01142798 | 1 | notSpecific | Up | DM | Loss | 10 |
| **19_48015160_48018217_9704** | 19_48015160_48018217 | DHX34 | 9704 | 5.21108581767889 | -0.83763926 | 1 | notSpecific | Up | DM | Loss | 10 |
| **19_48064153_48065625_84215** | 19_48064153_48065625 | ZNF541 | 84215 | 3.70260284134043 | 0 | 6 | only_H3vsUI | UpDown | notDM | Loss | 3 |
| **19_48971344_48972123_4924** | 19_48971344_48972123 | NUCB1 | 4924 | 5.63498583240309 | 0.85877603 | 5 | only_H3vsUI | DownUp | notDM | Loss | 7 |
| **19_49013379_49014531_4924** | 19_49013379_49014531 | NUCB1 | 4924 | 5.59005788233685 | -0.88810522 | 4 | only_H3vsUI | DownUp | notDM | Gain | 2 |
| **19_49140968_49141484_4924** | 19_49140968_49141484 | NUCB1 | 4924 | 5.41853992195166 | 0.95521165 | 1 | only_H3vsUI | DownUp | notDM | Gain | 1 |

| | | | | | | | | | | | |
|---|---|---|---|---|---|---|---|---|---|---|---|
| **19_49224188_49225401_4924** | 19_49224188_49225401 | NUCB1 | 4924 | 5.25085895459929 | -0.81886763 | 5 | only_H3vsUI | DownUp | notDM | Loss | 7 |
| **19_49283445_49284893_4924** | 19_49283445_49284893 | NUCB1 | 4924 | 5.07440681164852 | 0.67839173 | 6 | only_H3vsUI | DownUp | notDM | Loss | 5 |
| **19_49374289_49375483_4924** | 19_49374289_49375483 | NUCB1 | 4924 | 4.44862903643092 | 0.53715721 | 5 | only_H3vsUI | DownUp | notDM | Gain | 1 |
| **19_49375708_49380266_4924** | 19_49375708_49380266 | NUCB1 | 4924 | 4.36757953442592 | 0.53715721 | 2 | only_H3vsUI | DownUp | notDM | Loss | 6 |
| **19_49402483_49403385_4924** | 19_49402483_49403385 | NUCB1 | 4924 | 2.38560627359831 | -0.35111409 | 2 | only_H3vsUI | DownUp | notDM | Gain | 2 |
| **19_49403519_49406136_4924** | 19_49403519_49406136 | NUCB1 | 4924 | 0 | -0.33176093 | 1 | only_H3vsUI | DownUp | notDM | Loss | 8 |
| **19_49553419_49554470_4909** | 19_49553419_49554470 | NTF4 | 4909 | 4.14188854970586 | -0.44188445 | 6 | only_H3vsUI | Down | notDM | Gain | 1 |
| **19_49558815_49561485_4909** | 19_49558815_49561485 | NTF4 | 4909 | 3.8356271662099 | 0.47445287 | 2 | only_H3vsUI | Down | notDM | Loss | 6 |
| **19_49566842_49568863_4909** | 19_49566842_49568863 | NTF4 | 4909 | 0 | 0.27984393 | 4 | only_H3vsUI | Down | notDM | Loss | 3 |
| **19_49566842_49568863_4924** | 19_49566842_49568863 | NUCB1 | 4924 | 5.21276539107034 | -0.74553072 | 4 | only_H3vsUI | DownUp | notDM | Loss | 7 |
| **19_49622701_49624216_23521** | 19_49622701_49624216 | RPL13A | 23521 | 5.56418772198079 | -0.77758606 | 2 | only_DIvsD7 | Up | notDM | Loss | 7 |
| **19_49648649_49650375_23521** | 19_49648649_49650375 | RPL13A | 23521 | 5.53210563694779 | 0.78232199 | 2 | only_DIvsD7 | Up | notDM | Gain | 1 |
| **19_49651545_49654288_4924** | 19_49651545_49654288 | NUCB1 | 4924 | 5.39431156323123 | 0.84996204 | 2 | only_H3vsUI | DownUp | notDM | Loss | 6 |
| **19_49673234_49674365_4924** | 19_49673234_49674365 | NUCB1 | 4924 | 5.43081008803194 | 0.77564339 | 6 | only_H3vsUI | DownUp | notDM | Loss | 5 |
| **19_49710640_49711790_23521** | 19_49710640_49711790 | RPL13A | 23521 | 5.44572248960316 | 0.76321457 | 4 | only_DIvsD7 | Up | notDM | Gain | 1 |
| **19_49864322_49866804_4924** | 19_49864322_49866804 | NUCB1 | 4924 | 5.66341538419171 | 0.97168151 | 1 | only_H3vsUI | DownUp | notDM | Gain | 5 |
| **19_49867105_49868189_23521** | 19_49867105_49868189 | RPL13A | 23521 | 5.08876314700252 | 0.8356793 | 3 | only_DIvsD7 | Up | notDM | Loss | 5 |
| **19_49925523_49927047_23521** | 19_49925523_49927047 | RPL13A | 23521 | 4.80494999481386 | 0.56327122 | 2 | only_DIvsD7 | Up | notDM | Gain | 1 |
| **19_49930870_49932025_23521** | 19_49930870_49932025 | RPL13A | 23521 | 4.7692812979729 | 0.73006766 | 2 | only_DIvsD7 | Up | notDM | Loss | 3 |
| **19_49943839_49947047_23521** | 19_49943839_49947047 | RPL13A | 23521 | 4.64166246223798 | 0.74621296 | 1 | only_DIvsD7 | Up | notDM | Loss | 6 |
| **19_49956819_49958300_23521** | 19_49956819_49958300 | RPL13A | 23521 | 4.51204368626781 | 0.49689984 | 1 | only_DIvsD7 | Up | notDM | Loss | 6 |
| **19_49976585_49978281_23521** | 19_49976585_49978281 | RPL13A | 23521 | 4.09985321988438 | 0.15669432 | 2 | only_DIvsD7 | Up | notDM | Gain | 7 |
| **19_49991307_49994616_23521** | 19_49991307_49994616 | RPL13A | 23521 | 2.69722934275972 | -0.54541486 | 1 | only_DIvsD7 | Up | notDM | Loss | 8 |
| **19_49999785_50002781_23521** | 19_49999785_50002781 | RPL13A | 23521 | 3.95308284391209 | -0.48251184 | 1 | only_DIvsD7 | Up | notDM | Gain | 4 |
| **19_50036205_50038361_23521** | 19_50036205_50038361 | RPL13A | 23521 | 4.65650067260142 | 0.54383121 | 1 | only_DIvsD7 | Up | notDM | Loss | 6 |
| **19_50081980_50083829_23521** | 19_50081980_50083829 | RPL13A | 23521 | 4.95985671834134 | -0.80399317 | 1 | only_DIvsD7 | Up | notDM | Loss | 7 |
| **19_50355046_50357341_23521** | 19_50355046_50357341 | RPL13A | 23521 | 5.5613196691401 | 0.87593884 | 1 | only_DIvsD7 | Up | notDM | Loss | 6 |
| **19_50430795_50433279_9476** | 19_50430795_50433279 | NAPSA | 9476 | 5.63914070902534 | -0.85355982 | 1 | only_H3vsUI | DownUp | notDM | Loss | 7 |
| **19_507911_508901_55658** | 19_507911_508901 | RNF126 | 55658 | 5.18845879829624 | -0.79645742 | 2 | only_H3vsUI | UpDown | notDM | Loss | 7 |
| **19_50817543_50818823_9476** | 19_50817543_50818823 | NAPSA | 9476 | 4.69991573583337 | 0.76883033 | 4 | only_H3vsUI | DownUp | notDM | Loss | 3 |
| **19_50829053_50830237_9476** | 19_50829053_50830237 | NAPSA | 9476 | 4.58765485099572 | -0.66015941 | 4 | only_H3vsUI | DownUp | notDM | Loss | 3 |
| **19_50831751_50832034_9476** | 19_50831751_50832034 | NAPSA | 9476 | 4.56700282652482 | 0.63858617 | 4 | only_H3vsUI | DownUp | notDM | Gain | 1 |
| **19_50860043_50861276_9476** | 19_50860043_50861276 | NAPSA | 9476 | 3.88400192476879 | 0.63635605 | 2 | only_H3vsUI | DownUp | notDM | Gain | 1 |
| **19_50871281_50873309_9476** | 19_50871281_50873309 | NAPSA | 9476 | 3.3714373174041 | 0.26739606 | 1 | only_H3vsUI | DownUp | notDM | Gain | 4 |

| | | | | | | | | | | | |
|---|---|---|---|---|---|---|---|---|---|---|---|
| **19_50878590_50879634_9476** | 19_50878590_50879634 | NAPSA | 9476 | 3.98502208210954 | -0.14868009 | 1 | only_H3vsUI | DownUp | notDM | Gain | 4 |
| **19_50886500_50887351_9476** | 19_50886500_50887351 | NAPSA | 9476 | 4.24479647874762 | -0.30451917 | 2 | only_H3vsUI | DownUp | notDM | Loss | 3 |
| **19_50887588_50889507_9476** | 19_50887588_50889507 | NAPSA | 9476 | 4.27088836469883 | -0.30561063 | 2 | only_H3vsUI | DownUp | notDM | Loss | 6 |
| **19_50935440_50936518_9476** | 19_50935440_50936518 | NAPSA | 9476 | 4.82289347754723 | 0.81434788 | 2 | only_H3vsUI | DownUp | notDM | Loss | 6 |
| **19_50962989_50964570_9476** | 19_50962989_50964570 | NAPSA | 9476 | 4.97340497441006 | 0.99779496 | 6 | only_H3vsUI | DownUp | notDM | Loss | 5 |
| **19_50979795_50982017_554235** | 19_50979795_50982017 | ASPDH | 554235 | 4.54562869699285 | -0.41866966 | 1 | only_DIvsD7 | DownUp | notDM | Loss | 7 |
| **19_51004075_51005224_126119** | 19_51004075_51005224 | JOSD2 | 126119 | 3.96435392929219 | 0.32673392 | 6 | notSpecific | Up | DM | Loss | 10 |
| **19_51004075_51005224_256236** | 19_51004075_51005224 | NAPSB | 256236 | 5.19327212206205 | 0.78535085 | 6 | notSpecific | UpDown | DM | Loss | 10 |
| **19_51004075_51005224_284361** | 19_51004075_51005224 | EMC10 | 284361 | 4.38633841635751 | 0.53473474 | 6 | notSpecific | Up | DM | Loss | 10 |
| **19_51004075_51005224_51231** | 19_51004075_51005224 | VRK3 | 51231 | 5.67710576517216 | 0.81479463 | 6 | notSpecific | DownUp | DM | Loss | 10 |
| **19_51004075_51005224_554235** | 19_51004075_51005224 | ASPDH | 554235 | 4.07623981977502 | -0.28768085 | 6 | only_DIvsD7 | DownUp | DM | Loss | 10 |
| **19_51004075_51005224_9476** | 19_51004075_51005224 | NAPSA | 9476 | 5.13080319614177 | 0.7844756 | 6 | only_H3vsUI | DownUp | DM | Loss | 10 |
| **19_51013348_51014446_554235** | 19_51013348_51014446 | ASPDH | 554235 | 3.43088094645289 | -0.37673918 | 1 | only_DIvsD7 | DownUp | notDM | Loss | 3 |
| **19_51014779_51015831_554235** | 19_51014779_51015831 | ASPDH | 554235 | 3.11793383503964 | -0.39748968 | 1 | only_DIvsD7 | DownUp | notDM | Loss | 3 |
| **19_51222089_51222646_9476** | 19_51222089_51222646 | NAPSA | 9476 | 5.54797150814724 | 0.99974415 | 2 | only_H3vsUI | DownUp | notDM | Loss | 3 |
| **19_52799125_52800233_147657** | 19_52799125_52800233 | ZNF480 | 147657 | 2.28555730900777 | 0.26928055 | 2 | only_H3vsUI | Down | notDM | Gain | 4 |
| **19_52800531_52801670_147657** | 19_52800531_52801670 | ZNF480 | 147657 | 2.00432137378264 | -0.30364052 | 1 | only_H3vsUI | Down | notDM | Loss | 8 |
| **19_53836136_53837938_91664** | 19_53836136_53837938 | ZNF845 | 91664 | 0 | 0.38093063 | 1 | only_H3vsUI | DownUp | notDM | Gain | 4 |
| **19_53935247_53936413_126017** | 19_53935247_53936413 | ZNF813 | 126017 | 4.53878730993371 | -0.74035519 | 1 | only_H3vsUI | Up | notDM | Loss | 3 |
| **19_53935247_53936413_55422** | 19_53935247_53936413 | ZNF331 | 55422 | 4.94428028325184 | -0.94925514 | 1 | only_H3vsUI | UpDown | notDM | Loss | 7 |
| **19_53935247_53936413_91664** | 19_53935247_53936413 | ZNF845 | 91664 | 4.99231929794479 | -0.93399385 | 1 | only_H3vsUI | DownUp | notDM | Loss | 7 |
| **19_54024358_54025586_55422** | 19_54024358_54025586 | ZNF331 | 55422 | 2.09691001300806 | -0.239838 | 1 | only_H3vsUI | UpDown | notDM | Loss | 8 |
| **19_54047761_54049234_126017** | 19_54047761_54049234 | ZNF813 | 126017 | 4.8851519392003 | -0.55737382 | 7 | only_H3vsUI | Up | notDM | Gain | 1 |
| **19_54047761_54049234_55422** | 19_54047761_54049234 | ZNF331 | 55422 | 3.80386663428498 | 0.16981283 | 7 | only_H3vsUI | UpDown | notDM | Gain | 1 |
| **19_54384003_54386491_55422** | 19_54384003_54386491 | ZNF331 | 55422 | 5.55597303584727 | 0.94151118 | 1 | only_H3vsUI | UpDown | notDM | Gain | 5 |
| **19_54384003_54386491_59284** | 19_54384003_54386491 | CACNG7 | 59284 | 4.41662386738418 | 0.55204632 | 1 | only_H3vsUI | UpDown | notDM | Gain | 4 |
| **19_54392063_54393223_59284** | 19_54392063_54393223 | CACNG7 | 59284 | 4.28706235255439 | 0.36707563 | 1 | only_H3vsUI | UpDown | notDM | Loss | 6 |
| **19_54397775_54398599_59284** | 19_54397775_54398599 | CACNG7 | 59284 | 4.14584875659054 | -0.16716319 | 6 | only_H3vsUI | UpDown | notDM | Loss | 3 |
| **19_54409516_54410612_55422** | 19_54409516_54410612 | ZNF331 | 55422 | 5.58562651938917 | 0.9538257 | 4 | only_H3vsUI | UpDown | notDM | Gain | 1 |
| **19_54409516_54410612_59284** | 19_54409516_54410612 | CACNG7 | 59284 | 3.29622628726116 | 0.55113328 | 4 | only_H3vsUI | UpDown | notDM | Gain | 1 |
| **19_54411249_54412490_55422** | 19_54411249_54412490 | ZNF331 | 55422 | 5.58757627918068 | 0.9421193 | 2 | only_H3vsUI | UpDown | notDM | Loss | 3 |
| **19_54411249_54412490_59284** | 19_54411249_54412490 | CACNG7 | 59284 | 2 | 0.55078107 | 2 | only_H3vsUI | UpDown | notDM | Loss | 6 |
| **19_54412739_54413400_55422** | 19_54412739_54413400 | ZNF331 | 55422 | 5.58924567505251 | 0.93748358 | 1 | only_H3vsUI | UpDown | notDM | Loss | 3 |
| **19_54412739_54413400_59284** | 19_54412739_54413400 | CACNG7 | 59284 | 2.18184358794477 | 0.55196589 | 1 | only_H3vsUI | UpDown | notDM | Loss | 3 |

| | | | | | | | | | | | |
|---|---|---|---|---|---|---|---|---|---|---|---|
| **19_55833832_55834827_51157** | 19_55833832_55834827 | ZNF580 | 51157 | 5.50183399878398 | 0.86105531 | 6 | only_H3vsUI | UpDown | notDM | Loss | 5 |
| **19_55894034_55895044_51157** | 19_55894034_55895044 | ZNF580 | 51157 | 5.41052248501935 | -0.74544313 | 1 | only_H3vsUI | UpDown | notDM | Loss | 7 |
| **19_56109485_56110364_51157** | 19_56109485_56110364 | ZNF580 | 51157 | 4.62354905691839 | 0.5945211 | 2 | only_H3vsUI | UpDown | notDM | Gain | 1 |
| **19_56111912_56113608_51157** | 19_56111912_56113608 | ZNF580 | 51157 | 4.5886637957802 | 0.71411299 | 1 | only_H3vsUI | UpDown | notDM | Loss | 6 |
| **19_56136067_56137159_51157** | 19_56136067_56137159 | ZNF580 | 51157 | 4.18281395125737 | 0.64460151 | 2 | only_H3vsUI | UpDown | notDM | Loss | 6 |
| **19_56145280_56148631_51157** | 19_56145280_56148631 | ZNF580 | 51157 | 3.57541879121436 | 0.56928982 | 1 | only_H3vsUI | UpDown | notDM | Loss | 6 |
| **19_56151412_56152310_51157** | 19_56151412_56152310 | ZNF580 | 51157 | 1.91907809237607 | 0.29207547 | 5 | only_H3vsUI | UpDown | notDM | Loss | 3 |
| **19_56152490_56155050_51157** | 19_56152490_56155050 | ZNF580 | 51157 | 2 | 0.59903581 | 1 | only_H3vsUI | UpDown | notDM | Loss | 6 |
| **19_56155267_56156934_51157** | 19_56155267_56156934 | ZNF580 | 51157 | 3.45893986189033 | 0.75561649 | 1 | only_H3vsUI | UpDown | notDM | Loss | 6 |
| **19_56158497_56160919_51157** | 19_56158497_56160919 | ZNF580 | 51157 | 3.78582791999587 | -0.44325841 | 1 | only_H3vsUI | UpDown | notDM | Loss | 8 |
| **19_56166533_56169606_51157** | 19_56166533_56169606 | ZNF580 | 51157 | 4.15054154137409 | -0.43413072 | 1 | only_H3vsUI | UpDown | notDM | Loss | 8 |
| **19_5620659_5622359_2528** | 19_5620659_5622359 | FUT6 | 2528 | 5.33722757577868 | 0.75680875 | 1 | only_H3vsUI | UpDown | notDM | Loss | 3 |
| **19_5620659_5622359_25873** | 19_5620659_5622359 | RPL36 | 25873 | 4.8319593103043 | -0.75349763 | 1 | only_H3vsUI | Up | notDM | Loss | 7 |
| **19_5623688_5625599_2528** | 19_5623688_5625599 | FUT6 | 2528 | 5.33070591061026 | 0.83647343 | 1 | only_H3vsUI | UpDown | notDM | Gain | 1 |
| **19_5623688_5625599_25873** | 19_5623688_5625599 | RPL36 | 25873 | 4.81072972230796 | -0.8148933 | 1 | only_H3vsUI | Up | notDM | Gain | 2 |
| **19_56651119_56652111_55311** | 19_56651119_56652111 | ZNF444 | 55311 | 2.64933485871214 | -0.01350817 | 1 | only_H3vsUI | UpDown | notDM | Loss | 3 |
| **19_56653077_56654680_55311** | 19_56653077_56654680 | ZNF444 | 55311 | 2.71850168886727 | -0.06191039 | 1 | only_H3vsUI | UpDown | notDM | Loss | 3 |
| **19_5679243_5680428_25873** | 19_5679243_5680428 | RPL36 | 25873 | 3.99646789026172 | 0.70804305 | 2 | only_H3vsUI | Up | notDM | Loss | 6 |
| **19_57752228_57753999_390980** | 19_57752228_57753999 | ZNF805 | 390980 | 2.24797326636181 | 0.12056919 | 1 | only_H3vsUI | Down | notDM | Gain | 4 |
| **19_57831168_57833601_147694** | 19_57831168_57833601 | ZNF548 | 147694 | 4.83006232109096 | 0.65217112 | 1 | only_DIvsD7 | Down | notDM | Gain | 5 |
| **19_57874997_57876218_147694** | 19_57874997_57876218 | ZNF548 | 147694 | 4.39795738010389 | 0.4595104 | 1 | only_DIvsD7 | Down | notDM | Loss | 7 |
| **19_57922554_57923921_147694** | 19_57922554_57923921 | ZNF548 | 147694 | 4.32915371079282 | 0.8091131 | 2 | only_DIvsD7 | Down | DM | Loss | 10 |
| **19_57922554_57923921_388567** | 19_57922554_57923921 | ZNF749 | 388567 | 4.35776338112395 | -0.43781223 | 2 | notSpecific | Down | DM | Loss | 10 |
| **19_57922554_57923921_7565** | 19_57922554_57923921 | ZNF17 | 7565 | 1.43136376415899 | 0.16166776 | 2 | notSpecific | Down | DM | Loss | 10 |
| **19_58069879_58071102_7694** | 19_58069879_58071102 | ZNF135 | 7694 | 5.69857635518541 | 0.88673243 | 1 | only_H3vsUI | UpDown | notDM | Loss | 3 |
| **19_58089535_58090199_79891** | 19_58089535_58090199 | ZNF671 | 79891 | 5.17259417518446 | 0.94763707 | 2 | only_DIvsD7 | Down | notDM | Gain | 4 |
| **19_58193536_58194320_284309** | 19_58193536_58194320 | ZNF776 | 284309 | 4.80512689130286 | -0.58994878 | 1 | only_DIvsD7 | Down | notDM | Gain | 2 |
| **19_58258377_58259358_284309** | 19_58258377_58259358 | ZNF776 | 284309 | 2.23044892137827 | 0.15383043 | 1 | only_DIvsD7 | Down | notDM | Gain | 4 |
| **19_58281103_58282278_284309** | 19_58281103_58282278 | ZNF776 | 284309 | 4.3597596164157 | -0.67205119 | 1 | only_DIvsD7 | Down | notDM | Gain | 2 |
| **19_58281103_58282278_79891** | 19_58281103_58282278 | ZNF671 | 79891 | 4.62443680502756 | 0.52470141 | 1 | only_DIvsD7 | Down | notDM | Gain | 4 |
| **19_5828445_5829059_2528** | 19_5828445_5829059 | FUT6 | 2528 | 4.02714566577434 | 0.26340388 | 2 | only_H3vsUI | UpDown | notDM | Loss | 3 |
| **19_58331296_58332852_79891** | 19_58331296_58332852 | ZNF671 | 79891 | 4.96523934138665 | -0.70426761 | 1 | only_DIvsD7 | Down | notDM | Loss | 7 |
| **19_58426811_58427859_284309** | 19_58426811_58427859 | ZNF776 | 284309 | 5.22698377039466 | -0.83501111 | 2 | only_DIvsD7 | Down | notDM | Loss | 7 |
| **19_58872483_58874121_7694** | 19_58872483_58874121 | ZNF135 | 7694 | 5.47979981275863 | -0.82787037 | 1 | only_H3vsUI | UpDown | notDM | Loss | 3 |

| | | | | | | | | | | | |
|---|---|---|---|---|---|---|---|---|---|---|---|
| **19_58896163_58898483_7694** | 19_58896163_58898483 | ZNF135 | 7694 | 5.51259902281766 | -0.76815634 | 1 | only_H3vsUI | UpDown | notDM | Loss | 6 |
| **19_58898780_58902694_7694** | 19_58898780_58902694 | ZNF135 | 7694 | 5.51607637896469 | -0.76815634 | 1 | only_H3vsUI | UpDown | notDM | Loss | 6 |
| **19_58911547_58912661_201514** | 19_58911547_58912661 | ZNF584 | 201514 | 3.8694077493822 | 0.04718933 | 1 | only_H3vsUI | Down | notDM | Loss | 8 |
| **19_58912920_58914391_201514** | 19_58912920_58914391 | ZNF584 | 201514 | 3.7538127835647 | 0.08954167 | 1 | only_H3vsUI | Down | notDM | Loss | 8 |
| **19_58917844_58919789_201514** | 19_58917844_58919789 | ZNF584 | 201514 | 2.43933269383026 | -0.06268054 | 1 | only_H3vsUI | Down | notDM | Loss | 8 |
| **19_58920148_58921876_201514** | 19_58920148_58921876 | ZNF584 | 201514 | 1.93951925261862 | 0.0550595 | 1 | only_H3vsUI | Down | notDM | Loss | 8 |
| **19_58920148_58921876_7694** | 19_58920148_58921876 | ZNF135 | 7694 | 5.54344966530768 | -0.83209157 | 1 | only_H3vsUI | UpDown | notDM | Loss | 3 |
| **19_58962005_58962924_25799** | 19_58962005_58962924 | ZNF324 | 25799 | 4.1914510144649 | 0.30143817 | 1 | only_H3vsUI | DownUp | notDM | Loss | 7 |
| **19_58963000_58964273_25799** | 19_58963000_58964273 | ZNF324 | 25799 | 4.15189056873509 | 0.19039653 | 1 | only_H3vsUI | DownUp | notDM | Loss | 7 |
| **19_58977493_58978336_25799** | 19_58977493_58978336 | ZNF324 | 25799 | 2.10720996964787 | 0.19066597 | 1 | only_H3vsUI | DownUp | notDM | Loss | 3 |
| **19_58978486_58980347_25799** | 19_58978486_58980347 | ZNF324 | 25799 | 1.46239799789896 | 0.24536315 | 1 | only_H3vsUI | DownUp | notDM | Loss | 8 |
| **19_58986025_58987491_25799** | 19_58986025_58987491 | ZNF324 | 25799 | 3.87898112339374 | -0.11696391 | 1 | only_H3vsUI | DownUp | notDM | Loss | 8 |
| **19_58987571_58989151_25799** | 19_58987571_58989151 | ZNF324 | 25799 | 3.95970902424643 | 0.10515415 | 1 | only_H3vsUI | DownUp | notDM | Gain | 4 |
| **19_5902540_5903420_2527** | 19_5902540_5903420 | FUT5 | 2527 | 2.57863920996807 | 0.26459722 | 1 | only_H3vsUI | DownUp | notDM | Loss | 3 |
| **19_5905067_5905940_2527** | 19_5905067_5905940 | FUT5 | 2527 | 3.10414555055401 | 0.24090509 | 1 | only_H3vsUI | DownUp | notDM | Loss | 3 |
| **19_59067483_59069689_7694** | 19_59067483_59069689 | ZNF135 | 7694 | 5.69621393076249 | -0.92468215 | 1 | only_H3vsUI | UpDown | notDM | Loss | 3 |
| **19_59070537_59072335_7694** | 19_59070537_59072335 | ZNF135 | 7694 | 5.69887531781776 | -0.92468215 | 1 | only_H3vsUI | UpDown | notDM | Loss | 3 |
| **19_6391657_6392994_2527** | 19_6391657_6392994 | FUT5 | 2527 | 5.68829610165034 | 0.87200346 | 1 | only_H3vsUI | DownUp | notDM | Loss | 6 |
| **19_660779_662643_55658** | 19_660779_662643 | RNF126 | 55658 | 2.73399928653839 | 0.14804583 | 1 | only_H3vsUI | UpDown | notDM | Loss | 6 |
| **19_663523_665351_55658** | 19_663523_665351 | RNF126 | 55658 | 2.5327543789925 | 0.14804583 | 1 | only_H3vsUI | UpDown | notDM | Loss | 6 |
| **19_677709_678487_55658** | 19_677709_678487 | RNF126 | 55658 | 4.16217593653016 | 0.41030122 | 3 | only_H3vsUI | UpDown | notDM | Gain | 1 |
| **19_7693152_7694172_126006** | 19_7693152_7694172 | PCP2 | 126006 | 3.64335396197686 | -0.0016269 | 1 | only_H3vsUI | Up | notDM | Loss | 3 |
| **19_7694720_7696028_126006** | 19_7694720_7696028 | PCP2 | 126006 | 3.40534636017571 | -0.0016269 | 1 | only_H3vsUI | Up | notDM | Loss | 3 |
| **19_782822_785022_55658** | 19_782822_785022 | RNF126 | 55658 | 5.07787640435928 | 0.69713347 | 5 | only_H3vsUI | UpDown | notDM | Loss | 5 |
| **19_7953521_7954305_80131** | 19_7953521_7954305 | LRRC8E | 80131 | 2.12385164096709 | 0.50493113 | 1 | only_DIvsD7 | Up | notDM | Loss | 3 |
| **19_7967449_7968358_80131** | 19_7967449_7968358 | LRRC8E | 80131 | 4.14801620823987 | 0.44522016 | 1 | only_DIvsD7 | Up | notDM | Loss | 3 |
| **19_7969251_7970224_80131** | 19_7969251_7970224 | LRRC8E | 80131 | 4.20038532423239 | 0.44522016 | 2 | only_DIvsD7 | Up | notDM | Gain | 1 |
| **19_859508_860339_55658** | 19_859508_860339 | RNF126 | 55658 | 5.29286941137632 | -0.74863095 | 5 | only_H3vsUI | UpDown | notDM | Gain | 2 |
| **19_9185278_9186125_10781** | 19_9185278_9186125 | ZNF266 | 10781 | 5.55643518158933 | -0.90757839 | 7 | only_H3vsUI | Down | notDM | Gain | 2 |
| **19_9545339_9545917_10781** | 19_9545339_9545917 | ZNF266 | 10781 | 2.50242711998443 | 0.83030528 | 2 | only_H3vsUI | Down | notDM | Gain | 4 |
| **2_100936615_100937913_164832** | 2_100936615_100937913 | LONRF2 | 164832 | 3.10822665637493 | 0 | 2 | only_DIvsD7 | Down | notDM | Loss | 7 |
| **2_100938448_100939952_164832** | 2_100938448_100939952 | LONRF2 | 164832 | 0 | 0 | 2 | only_DIvsD7 | Down | notDM | Loss | 8 |
| **2_101481355_101483464_6160** | 2_101481355_101483464 | RPL31 | 6160 | 5.13127211693336 | 0.83292321 | 8 | only_DIvsD7 | Up | notDM | Loss | 5 |
| **2_101507734_101510391_6160** | 2_101507734_101510391 | RPL31 | 6160 | 5.03488903532673 | 0.73495513 | 7 | only_DIvsD7 | Up | notDM | Gain | 5 |

| | | | | | | | | | | | |
|---|---|---|---|---|---|---|---|---|---|---|---|
| **2_101510474_101513229_6160** | 2_101510474_101513229 | RPL31 | 6160 | 5.02336359186869 | 0.85008801 | 7 | only_DIvsD7 | Up | notDM | Gain | 5 |
| **2_101619129_101623656_6160** | 2_101619129_101623656 | RPL31 | 6160 | 2.57518784492766 | 0.44445412 | 1 | only_DIvsD7 | Up | notDM | Loss | 6 |
| **2_101676496_101683366_6160** | 2_101676496_101683366 | RPL31 | 6160 | 4.76198043145343 | -0.60988182 | 7 | only_DIvsD7 | Up | notDM | Gain | 5 |
| **2_101721124_101724438_6160** | 2_101721124_101724438 | RPL31 | 6160 | 5.010176945666 | 0.82457594 | 8 | only_DIvsD7 | Up | notDM | Gain | 5 |
| **2_101975989_101976929_200407** | 2_101975989_101976929 | CREG2 | 200407 | 4.43136376415899 | 0.76210964 | 6 | only_H3vsUI | DownUp | notDM | Gain | 1 |
| **2_102002487_102003642_200407** | 2_102002487_102003642 | CREG2 | 200407 | 2.45788189673399 | 0.90340798 | 2 | only_H3vsUI | DownUp | notDM | Loss | 3 |
| **2_102004022_102005357_200407** | 2_102004022_102005357 | CREG2 | 200407 | 1.98227123303957 | 0.77464395 | 4 | only_H3vsUI | DownUp | notDM | Gain | 4 |
| **2_102011775_102014057_200407** | 2_102011775_102014057 | CREG2 | 200407 | 3.8948143290833 | 0.57459847 | 3 | only_H3vsUI | DownUp | notDM | Gain | 1 |
| **2_10410351_10412030_3241** | 2_10410351_10412030 | HPCAL1 | 3241 | 4.49116551654101 | -0.64673425 | 6 | only_DIvsD7 | Up | notDM | Loss | 7 |
| **2_10422590_10423580_3241** | 2_10422590_10423580 | HPCAL1 | 3241 | 4.28860689039097 | 0.53363062 | 7 | only_DIvsD7 | Up | notDM | Loss | 5 |
| **2_10423677_10425839_3241** | 2_10423677_10425839 | HPCAL1 | 3241 | 4.23494731565269 | 0.2728677 | 7 | only_DIvsD7 | Up | notDM | Gain | 5 |
| **2_10437625_10438927_3241** | 2_10437625_10438927 | HPCAL1 | 3241 | 3.61161711055434 | 0.63248412 | 3 | only_DIvsD7 | Up | notDM | Gain | 1 |
| **2_10441704_10442553_3241** | 2_10441704_10442553 | HPCAL1 | 3241 | 2.66558099101795 | 0.46464694 | 5 | only_DIvsD7 | Up | notDM | Loss | 6 |
| **2_10443259_10447588_3241** | 2_10443259_10447588 | HPCAL1 | 3241 | 2.39093510710338 | 0.49555073 | 1 | only_DIvsD7 | Up | notDM | Loss | 6 |
| **2_10522004_10531435_3241** | 2_10522004_10531435 | HPCAL1 | 3241 | 4.89757761188539 | 0.82559526 | 7 | only_DIvsD7 | Up | notDM | Loss | 5 |
| **2_10532891_10534215_3241** | 2_10532891_10534215 | HPCAL1 | 3241 | 4.95365339976592 | 0.80172275 | 8 | only_DIvsD7 | Up | DM | Loss | 10 |
| **2_10532891_10534215_4953** | 2_10532891_10534215 | ODC1 | 4953 | 4.73416752302826 | 0.62728027 | 8 | notSpecific | Up | DM | Loss | 10 |
| **2_105944434_105945900_79074** | 2_105944434_105945900 | C2orf49 | 79074 | 3.90794852161227 | 0.5707124 | 1 | only_DIvsD7 | UpDown | notDM | Loss | 6 |
| **2_105946394_105947741_79074** | 2_105946394_105947741 | C2orf49 | 79074 | 3.79581052466741 | 0.45163604 | 1 | only_DIvsD7 | UpDown | notDM | Gain | 4 |
| **2_105952498_105953690_79074** | 2_105952498_105953690 | C2orf49 | 79074 | 2.47712125471966 | 0.6922095 | 1 | only_DIvsD7 | UpDown | notDM | Loss | 6 |
| **2_105954343_105956383_79074** | 2_105954343_105956383 | C2orf49 | 79074 | 2.55144999797288 | 0.6922095 | 1 | only_DIvsD7 | UpDown | notDM | Loss | 6 |
| **2_106006705_106008807_79074** | 2_106006705_106008807 | C2orf49 | 79074 | 4.72195892576198 | -0.63673998 | 8 | only_DIvsD7 | UpDown | notDM | Loss | 7 |
| **2_106258957_106263234_79074** | 2_106258957_106263234 | C2orf49 | 79074 | 5.48425711975576 | 0.78322714 | 4 | only_DIvsD7 | UpDown | notDM | Loss | 6 |
| **2_106808915_106810408_80146** | 2_106808915_106810408 | UXS1 | 80146 | 2.54157924394658 | -0.3297556 | 1 | only_H3vsUI | DownUp | notDM | Loss | 8 |
| **2_106811125_106812057_80146** | 2_106811125_106812057 | UXS1 | 80146 | 2.52113808370404 | -0.33858671 | 2 | only_H3vsUI | DownUp | notDM | Gain | 4 |
| **2_108809604_108811323_3987** | 2_108809604_108811323 | LIMS1 | 3987 | 5.53095489297132 | -0.77625559 | 6 | only_H3vsUI | Down | notDM | Gain | 2 |
| **2_108887360_108888542_3987** | 2_108887360_108888542 | LIMS1 | 3987 | 5.41891583056647 | -0.82171619 | 4 | only_H3vsUI | Down | notDM | Gain | 2 |
| **2_108887360_108888542_6819** | 2_108887360_108888542 | SULT1C2 | 6819 | 4.22585181081816 | 0.19507318 | 4 | only_H3vsUI | UpDown | notDM | Gain | 1 |
| **2_108888639_108889289_6819** | 2_108888639_108889289 | SULT1C2 | 6819 | 4.20612396399185 | -0.14815085 | 4 | only_H3vsUI | UpDown | notDM | Gain | 1 |
| **2_108985818_108986979_3987** | 2_108985818_108986979 | LIMS1 | 3987 | 5.4540670052789 | 0.88904725 | 6 | only_H3vsUI | Down | notDM | Gain | 1 |
| **2_109000770_109005603_3987** | 2_109000770_109005603 | LIMS1 | 3987 | 5.45747464935084 | 0.84183137 | 7 | only_H3vsUI | Down | notDM | Gain | 5 |
| **2_109130808_109131781_3987** | 2_109130808_109131781 | LIMS1 | 3987 | 4.28176037219993 | -0.61499163 | 7 | only_H3vsUI | Down | notDM | Gain | 1 |
| **2_109148516_109150500_3987** | 2_109148516_109150500 | LIMS1 | 3987 | 2.6159500516564 | -0.48274283 | 5 | only_H3vsUI | Down | notDM | Gain | 4 |
| **2_109151289_109155469_3987** | 2_109151289_109155469 | LIMS1 | 3987 | 2.63748972951251 | -0.07119094 | 2 | only_H3vsUI | Down | notDM | Gain | 4 |

| | | | | | | | | | | | |
|---|---|---|---|---|---|---|---|---|---|---|---|
| **2_109186514_109193178_3987** | 2_109186514_109193178 | LIMS1 | 3987 | 4.06985302111362 | -0.57844148 | 8 | only_H3vsUI | Down | notDM | Gain | 5 |
| **2_109209104_109213204_3987** | 2_109209104_109213204 | LIMS1 | 3987 | 3.63738965012921 | 0.70164517 | 7 | only_H3vsUI | Down | notDM | Gain | 5 |
| **2_109236861_109239313_3987** | 2_109236861_109239313 | LIMS1 | 3987 | 0 | 0.63056641 | 6 | only_H3vsUI | Down | notDM | Gain | 4 |
| **2_109255787_109257976_3987** | 2_109255787_109257976 | LIMS1 | 3987 | 4.70775747841998 | 0.85103251 | 6 | only_H3vsUI | Down | notDM | Loss | 5 |
| **2_109336461_109339219_165055** | 2_109336461_109339219 | CCDC138 | 165055 | 4.80625461193395 | -0.52913267 | 2 | notSpecific | Down | DM | Loss | 10 |
| **2_109336461_109339219_5903** | 2_109336461_109339219 | RANBP2 | 5903 | 2.72098574415374 | -0.32899852 | 2 | notSpecific | Down | DM | Loss | 10 |
| **2_109350761_109352314_3987** | 2_109350761_109352314 | LIMS1 | 3987 | 4.76663588631027 | -0.54221266 | 8 | only_H3vsUI | Down | notDM | Gain | 2 |
| **2_109647719_109648794_3987** | 2_109647719_109648794 | LIMS1 | 3987 | 5.64620663141406 | 0.95614029 | 2 | only_H3vsUI | Down | notDM | Loss | 1 |
| **2_111878706_111880068_10018** | 2_111878706_111880068 | BCL2L11 | 10018 | 2.25527250510331 | -0.07475718 | 1 | notSpecific | Down | DM | Loss | 10 |
| **2_113400526_113402980_3553** | 2_113400526_113402980 | IL1B | 3553 | 5.28197369885562 | 0.79782262 | 1 | only_H3vsUI | UpDown | notDM | Gain | 1 |
| **2_113541443_113542647_3553** | 2_113541443_113542647 | IL1B | 3553 | 4.71388517690747 | 0.65525164 | 4 | only_H3vsUI | UpDown | notDM | Gain | 1 |
| **2_113560885_113562376_3553** | 2_113560885_113562376 | IL1B | 3553 | 4.50539420028499 | 0.51454407 | 3 | only_H3vsUI | UpDown | notDM | Gain | 1 |
| **2_113591445_113592639_3553** | 2_113591445_113592639 | IL1B | 3553 | 3.24427712080184 | 0.84655447 | 6 | only_H3vsUI | UpDown | notDM | Gain | 1 |
| **2_113626151_113628118_3553** | 2_113626151_113628118 | IL1B | 3553 | 4.50188049375506 | 0.68513676 | 6 | only_H3vsUI | UpDown | notDM | Gain | 1 |
| **2_113896062_113896541_3553** | 2_113896062_113896541 | IL1B | 3553 | 5.47953356292501 | 0.99638438 | 4 | only_H3vsUI | UpDown | notDM | Gain | 1 |
| **2_113961728_113962498_3553** | 2_113961728_113962498 | IL1B | 3553 | 5.56506467483735 | 0.80828113 | 7 | only_H3vsUI | UpDown | notDM | Gain | 1 |
| **2_114084328_114089460_3553** | 2_114084328_114089460 | IL1B | 3553 | 5.69014023857667 | 0.99583021 | 7 | only_H3vsUI | UpDown | notDM | Gain | 1 |
| **2_118943243_118944486_10050679** | 2_118943243_118944486 | THORLNC | 100506797 | 0 | 0.03117444 | 2 | only_H3vsUI | DownUp | notDM | Loss | 3 |
| **2_122353947_122354741_254128** | 2_122353947_122354741 | NIFK-AS1 | 254128 | 4.72375112163554 | 0.62043819 | 6 | only_H3vsUI | DownUp | notDM | Loss | 1 |
| **2_122390741_122393915_254128** | 2_122390741_122393915 | NIFK-AS1 | 254128 | 4.12424330894665 | -0.11736712 | 7 | only_H3vsUI | DownUp | notDM | Gain | 5 |
| **2_122404826_122406841_254128** | 2_122404826_122406841 | NIFK-AS1 | 254128 | 2.92220627743902 | 0.54736145 | 1 | only_H3vsUI | DownUp | notDM | Gain | 4 |
| **2_122407431_122410122_254128** | 2_122407431_122410122 | NIFK-AS1 | 254128 | 0 | 0.54736145 | 2 | only_H3vsUI | DownUp | notDM | Loss | 8 |
| **2_127974409_127977438_339761** | 2_127974409_127977438 | CYP27C1 | 339761 | 2.33645973384853 | 0.06327458 | 1 | only_H3vsUI | DownUp | notDM | Loss | 6 |
| **2_127977819_127978802_339761** | 2_127977819_127978802 | CYP27C1 | 339761 | 2.22271647114758 | 0.08056527 | 5 | only_H3vsUI | DownUp | notDM | Loss | 3 |
| **2_127982346_127985537_339761** | 2_127982346_127985537 | CYP27C1 | 339761 | 3.67154308526257 | 0.03078407 | 3 | only_H3vsUI | DownUp | notDM | Gain | 4 |
| **2_128457512_128458945_84826** | 2_128457512_128458945 | SFT2D3 | 84826 | 0 | 0.00057364 | 5 | only_H3vsUI | DownUp | notDM | Gain | 4 |
| **2_128459775_128461231_84826** | 2_128459775_128461231 | SFT2D3 | 84826 | 3.07188200730613 | 0.00057364 | 2 | only_H3vsUI | DownUp | notDM | Gain | 4 |
| **2_130937079_130938779_80097** | 2_130937079_130938779 | MZT2B | 80097 | 3.04453976039241 | -0.69964239 | 1 | only_H3vsUI | UpDown | notDM | Loss | 8 |
| **2_130940156_130941662_80097** | 2_130940156_130941662 | MZT2B | 80097 | 2.4345689040342 | -0.69964239 | 1 | only_H3vsUI | UpDown | notDM | Loss | 8 |
| **2_130997903_130998647_80097** | 2_130997903_130998647 | MZT2B | 80097 | 4.76398921847227 | 0.61969438 | 7 | only_H3vsUI | UpDown | notDM | Gain | 1 |
| **2_131148128_131150234_80097** | 2_131148128_131150234 | MZT2B | 80097 | 5.31977209142636 | 0.95212096 | 2 | only_H3vsUI | UpDown | notDM | Loss | 6 |
| **2_131721613_131722750_50649** | 2_131721613_131722750 | ARHGEF4 | 50649 | 5.10423438219647 | -0.74181422 | 4 | notSpecific | Up | DM | Loss | 10 |
| **2_132247572_132249752_150776** | 2_132247572_132249752 | SMPD4BP | 150776 | 2.80277372529198 | -0.00525632 | 1 | notSpecific | Down | DM | Loss | 10 |
| **2_132247572_132249752_653784** | 2_132247572_132249752 | MZT2A | 653784 | 2.75204844781944 | 0.77622283 | 1 | notSpecific | Up | DM | Loss | 10 |

| | | | | | | | | | | | |
|---|---|---|---|---|---|---|---|---|---|---|---|
| **2_134876004_134877412_4249** | 2_134876004_134877412 | MGAT5 | 4249 | 2.53402610605613 | 0.24605168 | 4 | only_H3vsUI | Down | notDM | Loss | 8 |
| **2_134877931_134879915_4249** | 2_134877931_134879915 | MGAT5 | 4249 | 2.25527250510331 | 0.2105197 | 1 | only_H3vsUI | Down | notDM | Loss | 8 |
| **2_135475394_135476219_81615** | 2_135475394_135476219 | TMEM163 | 81615 | 2.54654266347813 | -0.0006387 | 4 | only_H3vsUI | UpDown | notDM | Gain | 1 |
| **2_136286240_136288089_84083** | 2_136286240_136288089 | ZRANB3 | 84083 | 2.32837960343874 | -0.39299539 | 1 | only_DIvsD7 | DownUp | notDM | Loss | 7 |
| **2_136289479_136291944_84083** | 2_136289479_136291944 | ZRANB3 | 84083 | 3.07188200730613 | -0.39299539 | 2 | only_DIvsD7 | DownUp | notDM | Loss | 8 |
| **2_150441545_150443750_10192923** | 2_150441545_150443750 | MMADHC-DT | 101929231 | 0 | 0 | 2 | only_DIvsD7 | UpDown | notDM | Gain | 4 |
| **2_150444330_150445499_10192923** | 2_150444330_150445499 | MMADHC-DT | 101929231 | 2.79379038469082 | 0 | 1 | only_DIvsD7 | UpDown | notDM | Gain | 4 |
| **2_152230298_152231462_7130** | 2_152230298_152231462 | TNFAIP6 | 7130 | 4.20935413494016 | -0.19791013 | 3 | only_H3vsUI | UpDown | notDM | Gain | 1 |
| **2_162273159_162274444_10716** | 2_162273159_162274444 | TBR1 | 10716 | 0 | 0 | 4 | notSpecific | Up | DM | Loss | 10 |
| **2_162273159_162274444_57282** | 2_162273159_162274444 | SLC4A10 | 57282 | 3.80617997398389 | -0.0024228 | 4 | notSpecific | UpDown | DM | Loss | 10 |
| **2_165370173_165371706_22837** | 2_165370173_165371706 | COBLL1 | 22837 | 5.51451986129337 | 0.76735059 | 6 | only_H3vsUI | Down | notDM | Loss | 1 |
| **2_165548420_165555926_22837** | 2_165548420_165555926 | COBLL1 | 22837 | 5.15460349714295 | 0.93619898 | 4 | only_H3vsUI | Down | notDM | Gain | 5 |
| **2_165566260_165568004_22837** | 2_165566260_165568004 | COBLL1 | 22837 | 5.11621244913383 | 0.93293209 | 7 | only_H3vsUI | Down | notDM | Gain | 1 |
| **2_165613008_165615924_22837** | 2_165613008_165615924 | COBLL1 | 22837 | 4.91782572959952 | 0.93280847 | 6 | only_H3vsUI | Down | DM | Gain | 9 |
| **2_165696076_165697707_22837** | 2_165696076_165697707 | COBLL1 | 22837 | 2.98766626492627 | 0.72961672 | 2 | only_H3vsUI | Down | notDM | Loss | 3 |
| **2_165698478_165699329_22837** | 2_165698478_165699329 | COBLL1 | 22837 | 0 | 0.71449246 | 2 | only_H3vsUI | Down | notDM | Loss | 3 |
| **2_165753715_165755865_22837** | 2_165753715_165755865 | COBLL1 | 22837 | 4.74067053463103 | 0.68663069 | 3 | only_H3vsUI | Down | notDM | Gain | 1 |
| **2_165776296_165778082_22837** | 2_165776296_165778082 | COBLL1 | 22837 | 4.88994006628845 | 0.77209031 | 4 | only_H3vsUI | Down | notDM | Gain | 1 |
| **2_165793623_165795757_22837** | 2_165793623_165795757 | COBLL1 | 22837 | 4.97745380176821 | 0.93113376 | 4 | only_H3vsUI | Down | notDM | Gain | 1 |
| **2_171012936_171014699_140469** | 2_171012936_171014699 | MYO3B | 140469 | 4.30009525732372 | 0.29833607 | 3 | only_H3vsUI | UpDown | notDM | Loss | 3 |
| **2_176942974_176944830_344191** | 2_176942974_176944830 | EVX2 | 344191 | 3.58115289196629 | 0 | 4 | only_H3vsUI | UpDown | notDM | Loss | 6 |
| **2_176950141_176951608_344191** | 2_176950141_176951608 | EVX2 | 344191 | 3.17666993266815 | 0 | 4 | only_H3vsUI | UpDown | notDM | Loss | 3 |
| **2_176952943_176953771_344191** | 2_176952943_176953771 | EVX2 | 344191 | 3.63387226265833 | 0 | 4 | only_H3vsUI | UpDown | notDM | Loss | 3 |
| **2_179313547_179315383_494513** | 2_179313547_179315383 | PJVK | 494513 | 2.8926510338773 | -0.03110041 | 1 | only_H3vsUI | UpDown | notDM | Loss | 3 |
| **2_179316158_179317426_494513** | 2_179316158_179317426 | PJVK | 494513 | 0 | -0.03110041 | 5 | only_H3vsUI | UpDown | notDM | Loss | 3 |
| **2_182321375_182323019_3676** | 2_182321375_182323019 | ITGA4 | 3676 | 0 | 0.64232422 | 2 | only_H3vsUI | DownUp | notDM | Loss | 3 |
| **2_190412828_190414294_30061** | 2_190412828_190414294 | SLC40A1 | 30061 | 4.51140202968927 | 0.53322559 | 7 | notSpecific | Down | DM | Gain | 9 |
| **2_190412828_190414294_54529** | 2_190412828_190414294 | ASNSD1 | 54529 | 5.04881456187261 | -0.89173572 | 7 | notSpecific | DownUp | DM | Gain | 9 |
| **2_191134594_191136003_3628** | 2_191134594_191136003 | INPP1 | 3628 | 4.86118779279561 | -0.65450631 | 6 | notSpecific | Up | DM | Gain | 9 |
| **2_191134594_191136003_84281** | 2_191134594_191136003 | C2orf88 | 84281 | 4.94942416329465 | 0.77942489 | 6 | notSpecific | UpDown | DM | Gain | 9 |
| **2_200321744_200322807_23314** | 2_200321744_200322807 | SATB2 | 23314 | 1.11394335230684 | 0.0238221 | 2 | notSpecific | UpDown | DM | Loss | 10 |
| **2_200336477_200336797_79568** | 2_200336477_200336797 | MAIP1 | 79568 | 5.68461918249051 | -0.85954082 | 4 | only_H3vsUI | DownUp | notDM | Gain | 2 |
| **2_200774603_200775768_129450** | 2_200774603_200775768 | TYW5 | 129450 | 4.64793038377732 | -0.69303141 | 1 | only_H3vsUI | Down | notDM | Gain | 2 |
| **2_200776610_200777816_129450** | 2_200776610_200777816 | TYW5 | 129450 | 4.62744779121837 | -0.69303141 | 1 | only_H3vsUI | Down | notDM | Gain | 4 |

| | | | | | | | | | | | |
|---|---|---|---|---|---|---|---|---|---|---|---|
| **2_200817929_200819978_129450** | 2_200817929_200819978 | TYW5 | 129450 | 2.39093510710338 | -0.26152533 | 2 | only_H3vsUI | Down | notDM | Loss | 8 |
| **2_200817929_200819978_79568** | 2_200817929_200819978 | MAIP1 | 79568 | 2.75358305889291 | 0.34215789 | 2 | only_H3vsUI | DownUp | notDM | Loss | 8 |
| **2_200820812_200822828_129450** | 2_200820812_200822828 | TYW5 | 129450 | 2.77158748088126 | -0.23221833 | 1 | only_H3vsUI | Down | notDM | Gain | 4 |
| **2_200820812_200822828_79568** | 2_200820812_200822828 | MAIP1 | 79568 | 2.43136376415899 | 0.29008888 | 1 | only_H3vsUI | DownUp | notDM | Gain | 4 |
| **2_200841874_200844039_129450** | 2_200841874_200844039 | TYW5 | 129450 | 4.33551807589423 | 0.60799845 | 3 | only_H3vsUI | Down | notDM | Gain | 1 |
| **2_200841874_200844039_79568** | 2_200841874_200844039 | MAIP1 | 79568 | 4.33344727449675 | -0.33020602 | 3 | only_H3vsUI | DownUp | notDM | Gain | 5 |
| **2_201268814_201275788_129450** | 2_201268814_201275788 | TYW5 | 129450 | 5.65185249241883 | -0.90835947 | 7 | only_H3vsUI | Down | notDM | Gain | 5 |
| **2_20131611_20135035_130502** | 2_20131611_20135035 | TTC32 | 130502 | 4.48016510088712 | -0.41517498 | 7 | notSpecific | UpDown | DM | Gain | 9 |
| **2_20131611_20135035_57539** | 2_20131611_20135035 | WDR35 | 57539 | 4.73923997015154 | -0.76569707 | 7 | notSpecific | Down | DM | Gain | 9 |
| **2_202978069_202981415_729224** | 2_202978069_202981415 | KIAA2012-AS1 | 729224 | 2.56466606425209 | 0 | 6 | only_DIvsD7 | DownUp | notDM | Loss | 6 |
| **2_203239059_203240766_659** | 2_203239059_203240766 | BMPR2 | 659 | 2.95133751879592 | 0.04149637 | 5 | only_H3vsUI | Down | notDM | Gain | 4 |
| **2_203241798_203245495_659** | 2_203241798_203245495 | BMPR2 | 659 | 0 | 0.04764396 | 1 | only_H3vsUI | Down | notDM | Loss | 8 |
| **2_207138404_207139472_57683** | 2_207138404_207139472 | ZDBF2 | 57683 | 0 | 0.17868416 | 4 | only_DIvsD7 | Up | notDM | Loss | 3 |
| **2_207139914_207142375_57683** | 2_207139914_207142375 | ZDBF2 | 57683 | 2.72345567203519 | 0.13572074 | 2 | only_DIvsD7 | Up | notDM | Gain | 4 |
| **2_211033101_211035385_151050** | 2_211033101_211035385 | KANSL1L | 151050 | 0 | 0.01110703 | 4 | only_DIvsD7 | DownUp | notDM | Gain | 4 |
| **2_211036268_211037361_151050** | 2_211036268_211037361 | KANSL1L | 151050 | 2.21218760440396 | 0.00478124 | 2 | only_DIvsD7 | DownUp | notDM | Gain | 4 |
| **2_219860990_219861673_1412** | 2_219860990_219861673 | CRYBA2 | 1412 | 3.45469244923948 | 0.17672819 | 2 | only_DIvsD7 | DownUp | notDM | Gain | 1 |
| **2_219866698_219868436_1412** | 2_219866698_219868436 | CRYBA2 | 1412 | 3.93232153198929 | -0.10189811 | 2 | only_DIvsD7 | DownUp | notDM | Loss | 3 |
| **2_224466617_224467809_7857** | 2_224466617_224467809 | SCG2 | 7857 | 0 | 0 | 2 | only_H3vsUI | UpDown | notDM | Gain | 4 |
| **2_231191254_231192817_93349** | 2_231191254_231192817 | SP140L | 93349 | 0 | 0.77191834 | 4 | only_H3vsUI | DownUp | notDM | Gain | 4 |
| **2_231275793_231277829_93349** | 2_231275793_231277829 | SP140L | 93349 | 4.92374125499629 | 0.80544413 | 2 | only_H3vsUI | DownUp | notDM | Loss | 6 |
| **2_231279902_231283185_93349** | 2_231279902_231283185 | SP140L | 93349 | 4.94450734727201 | 0.75030392 | 4 | only_H3vsUI | DownUp | notDM | Loss | 6 |
| **2_231576527_231577413_93349** | 2_231576527_231577413 | SP140L | 93349 | 5.58504315490594 | 0.90874604 | 1 | only_H3vsUI | DownUp | notDM | Loss | 3 |
| **2_231578300_231580416_93349** | 2_231578300_231580416 | SP140L | 93349 | 5.58704048958193 | 0.90874604 | 1 | only_H3vsUI | DownUp | notDM | Gain | 5 |
| **2_231921972_231925169_80210** | 2_231921972_231925169 | ARMC9 | 80210 | 5.14042633016676 | -0.77742035 | 1 | only_H3vsUI | DownUp | notDM | Loss | 7 |
| **2_232062198_232063183_80210** | 2_232062198_232063183 | ARMC9 | 80210 | 2.20411998265592 | 0.10722588 | 2 | only_H3vsUI | DownUp | notDM | Loss | 8 |
| **2_232063530_232065990_80210** | 2_232063530_232065990 | ARMC9 | 80210 | 2.27875360095283 | 0.49145951 | 2 | only_H3vsUI | DownUp | notDM | Loss | 6 |
| **2_232229432_232231292_80210** | 2_232229432_232231292 | ARMC9 | 80210 | 3.60140806053468 | 0 | 7 | only_H3vsUI | DownUp | notDM | Loss | 5 |
| **2_232237654_232239170_80210** | 2_232237654_232239170 | ARMC9 | 80210 | 3.37493155397819 | 0 | 6 | only_H3vsUI | DownUp | notDM | Loss | 1 |
| **2_232326056_232328638_80210** | 2_232326056_232328638 | ARMC9 | 80210 | 5.41948652311005 | -0.78350678 | 1 | only_H3vsUI | DownUp | notDM | Loss | 7 |
| **2_232329558_232330716_80210** | 2_232329558_232330716 | ARMC9 | 80210 | 5.42523741641435 | -0.82784209 | 1 | only_H3vsUI | DownUp | notDM | Loss | 7 |
| **2_232789560_232790013_4880** | 2_232789560_232790013 | NPPC | 4880 | 3.04178731897175 | -0.07008005 | 4 | only_DIvsD7 | Up | notDM | Gain | 1 |
| **2_232790124_232791801_4880** | 2_232790124_232791801 | NPPC | 4880 | 0 | 0.01031918 | 2 | only_DIvsD7 | Up | notDM | Loss | 3 |
| **2_234475451_234476368_55230** | 2_234475451_234476368 | USP40 | 55230 | 1.39794000867204 | 0.76208223 | 1 | notSpecific | Down | DM | Gain | 9 |

| | | | | | | | | | | | |
|---|---|---|---|---|---|---|---|---|---|---|---|
| **2_234666381_234666818_10028692** | 2_234666381_234666818 | LOC100286922 | 100286922 | 3.37876117531637 | 0 | 7 | only_H3vsUI | UpDown | notDM | Gain | 1 |
| **2_239229683_239231036_26146** | 2_239229683_239231036 | TRAF3IP1 | 26146 | 2.69897000433602 | 0.32942106 | 1 | notSpecific | Down | DM | Loss | 10 |
| **2_239756914_239759389_117581** | 2_239756914_239759389 | TWIST2 | 117581 | 2.38560627359831 | 0.07432642 | 1 | only_H3vsUI | DownUp | notDM | Loss | 8 |
| **2_241526435_241528191_2859** | 2_241526435_241528191 | GPR35 | 2859 | 4.22162285774879 | 0.16338993 | 1 | only_H3vsUI | UpDown | notDM | Gain | 4 |
| **2_24234070_24236570_9540** | 2_24234070_24236570 | TP53I3 | 9540 | 4.84887851253801 | -0.58295671 | 6 | only_DIvsD7 | DownUp | notDM | Loss | 7 |
| **2_242822795_242823757_10192728** | 2_242822795_242823757 | LINC01237 | 101927289 | 0 | 0.15685437 | 2 | only_DIvsD7 | UpDown | notDM | Loss | 3 |
| **2_24297342_24299119_9540** | 2_24297342_24299119 | TP53I3 | 9540 | 3.90649665979935 | 0.68039265 | 1 | only_DIvsD7 | DownUp | notDM | Loss | 6 |
| **2_24305169_24307113_9540** | 2_24305169_24307113 | TP53I3 | 9540 | 1.83884909073726 | 0.1970467 | 1 | only_DIvsD7 | DownUp | notDM | Gain | 4 |
| **2_24307693_24308801_9540** | 2_24307693_24308801 | TP53I3 | 9540 | 1.60205999132796 | -0.22539667 | 5 | only_DIvsD7 | DownUp | notDM | Loss | 7 |
| **2_24345022_24346116_9540** | 2_24345022_24346116 | TP53I3 | 9540 | 4.57251147622018 | -0.50547574 | 1 | only_DIvsD7 | DownUp | notDM | Loss | 7 |
| **2_24346680_24348169_9540** | 2_24346680_24348169 | TP53I3 | 9540 | 4.59136516840857 | -0.50493988 | 2 | only_DIvsD7 | DownUp | notDM | Loss | 7 |
| **2_25013663_25015884_391356** | 2_25013663_25015884 | PTRHD1 | 391356 | 2.56584781867352 | 0.54260903 | 1 | only_H3vsUI | DownUp | notDM | Gain | 4 |
| **2_25016453_25019008_391356** | 2_25016453_25019008 | PTRHD1 | 391356 | 2.3096301674259 | 0.79600547 | 2 | only_H3vsUI | DownUp | notDM | Gain | 4 |
| **2_25028289_25030883_391356** | 2_25028289_25030883 | PTRHD1 | 391356 | 4.09233458952102 | -0.24393024 | 7 | only_H3vsUI | DownUp | notDM | Loss | 5 |
| **2_25031980_25033345_391356** | 2_25031980_25033345 | PTRHD1 | 391356 | 4.19675633105799 | 0.55069387 | 8 | only_H3vsUI | DownUp | notDM | Gain | 1 |
| **2_25219180_25221169_391356** | 2_25219180_25221169 | PTRHD1 | 391356 | 5.30734839548364 | 0.73215246 | 7 | only_H3vsUI | DownUp | notDM | Gain | 5 |
| **2_25300137_25302963_391356** | 2_25300137_25302963 | PTRHD1 | 391356 | 5.45314703520311 | 0.78749026 | 3 | only_H3vsUI | DownUp | notDM | Gain | 5 |
| **2_25887097_25888099_1838** | 2_25887097_25888099 | DTNB | 1838 | 3.92074926127571 | 0.52830247 | 6 | notSpecific | UpDown | DM | Gain | 9 |
| **2_26566828_26568607_165082** | 2_26566828_26568607 | ADGRF3 | 165082 | 1.88081359228079 | -0.19753294 | 1 | only_DIvsD7 | DownUp | notDM | Gain | 4 |
| **2_26569289_26572374_165082** | 2_26569289_26572374 | ADGRF3 | 165082 | 2.78461729263288 | -0.19753294 | 1 | only_DIvsD7 | DownUp | notDM | Gain | 4 |
| **2_26577407_26580267_165082** | 2_26577407_26580267 | ADGRF3 | 165082 | 3.94146173934733 | 0.47467731 | 7 | only_DIvsD7 | DownUp | notDM | Gain | 1 |
| **2_26634300_26635825_165082** | 2_26634300_26635825 | ADGRF3 | 165082 | 4.81711563868827 | 0.75277381 | 4 | only_DIvsD7 | DownUp | notDM | Gain | 1 |
| **2_27192207_27193810_22924** | 2_27192207_27193810 | MAPRE3 | 22924 | 0 | 0.81168528 | 1 | only_DIvsD7 | DownUp | notDM | Loss | 6 |
| **2_27192207_27193810_8884** | 2_27192207_27193810 | SLC5A6 | 8884 | 5.38167632116631 | -0.77656948 | 1 | only_H3vsUI | Down | notDM | Loss | 7 |
| **2_27193906_27194959_22924** | 2_27193906_27194959 | MAPRE3 | 22924 | 2.63144376901317 | 0.87367598 | 1 | only_DIvsD7 | DownUp | notDM | Loss | 6 |
| **2_27232561_27234672_22924** | 2_27232561_27234672 | MAPRE3 | 22924 | 3.47202469770028 | 0 | 6 | only_DIvsD7 | DownUp | notDM | Gain | 1 |
| **2_27237911_27240263_22924** | 2_27237911_27240263 | MAPRE3 | 22924 | 2.39093510710338 | 0 | 7 | only_DIvsD7 | DownUp | notDM | Loss | 5 |
| **2_27294757_27295866_22924** | 2_27294757_27295866 | MAPRE3 | 22924 | 5.00532210718478 | -0.77033323 | 2 | only_DIvsD7 | DownUp | notDM | Loss | 7 |
| **2_27294757_27295866_8884** | 2_27294757_27295866 | SLC5A6 | 8884 | 5.14384198349666 | 0.7273541 | 2 | only_H3vsUI | Down | notDM | Loss | 7 |
| **2_27309622_27310962_8884** | 2_27309622_27310962 | SLC5A6 | 8884 | 5.09270310416503 | 0.69628882 | 1 | only_H3vsUI | Down | notDM | Loss | 7 |
| **2_27355595_27357418_10113** | 2_27355595_27357418 | PREB | 10113 | 2.10037054511756 | -0.4937679 | 1 | notSpecific | UpDown | DM | Loss | 10 |
| **2_27355595_27357418_10669** | 2_27355595_27357418 | CGREF1 | 10669 | 4.33499628024323 | -0.5495284 | 1 | notSpecific | UpDown | DM | Loss | 10 |
| **2_27355595_27357418_339779** | 2_27355595_27357418 | PRR30 | 339779 | 3.68609977199592 | 0 | 1 | notSpecific | Up | DM | Loss | 10 |
| **2_27355595_27357418_84696** | 2_27355595_27357418 | ABHD1 | 84696 | 3.94885290619971 | -0.04632764 | 1 | notSpecific | Up | DM | Loss | 10 |

| | | | | | | | | | | | |
|---|---|---|---|---|---|---|---|---|---|---|---|
| **2_27355595_27357418_8890** | 2_27355595_27357418 | EIF2B4 | 8890 | 5.3725143310921 | -0.92517372 | 1 | notSpecific | DownUp | DM | Loss | 10 |
| **2_27357738_27358413_8884** | 2_27357738_27358413 | SLC5A6 | 8884 | 4.88277491081897 | 0.61357023 | 5 | only_H3vsUI | Down | notDM | Gain | 1 |
| **2_27392617_27393408_8884** | 2_27392617_27393408 | SLC5A6 | 8884 | 4.62037552786102 | 0.90789164 | 4 | only_H3vsUI | Down | notDM | Loss | 7 |
| **2_27433628_27434682_8884** | 2_27433628_27434682 | SLC5A6 | 8884 | 0 | -0.21837402 | 1 | only_H3vsUI | Down | notDM | Loss | 8 |
| **2_27435226_27437443_8884** | 2_27435226_27437443 | SLC5A6 | 8884 | 1.86923171973098 | 0.23592478 | 1 | only_H3vsUI | Down | notDM | Gain | 4 |
| **2_27440352_27442395_8884** | 2_27440352_27442395 | SLC5A6 | 8884 | 3.74803289413014 | 0.82497884 | 1 | only_H3vsUI | Down | notDM | Gain | 4 |
| **2_27485186_27488363_8884** | 2_27485186_27488363 | SLC5A6 | 8884 | 4.70352351311803 | -0.58976283 | 2 | only_H3vsUI | Down | notDM | Loss | 8 |
| **2_27514198_27517633_8884** | 2_27514198_27517633 | SLC5A6 | 8884 | 4.89787989802686 | 0.69644541 | 7 | only_H3vsUI | Down | notDM | Gain | 5 |
| **2_27514198_27517633_9553** | 2_27514198_27517633 | MRPL33 | 9553 | 5.67847467425172 | 0.90321141 | 7 | only_DIvsD7 | DownUp | notDM | Gain | 5 |
| **2_27525488_27526801_8884** | 2_27525488_27526801 | SLC5A6 | 8884 | 4.95586085646682 | 0.76059006 | 4 | only_H3vsUI | Down | notDM | Loss | 7 |
| **2_27525488_27526801_9553** | 2_27525488_27526801 | MRPL33 | 9553 | 5.67004536319664 | 0.93765799 | 4 | only_DIvsD7 | DownUp | notDM | Loss | 5 |
| **2_27601336_27603427_8884** | 2_27601336_27603427 | SLC5A6 | 8884 | 5.22187567503955 | -0.88238279 | 1 | only_H3vsUI | Down | notDM | Gain | 2 |
| **2_27629054_27632267_8884** | 2_27629054_27632267 | SLC5A6 | 8884 | 5.28868509032104 | -0.79021609 | 1 | only_H3vsUI | Down | notDM | Loss | 2 |
| **2_27718143_27718804_9553** | 2_27718143_27718804 | MRPL33 | 9553 | 5.44056434204506 | 0.84516626 | 2 | only_DIvsD7 | DownUp | notDM | Loss | 7 |
| **2_27850463_27851734_79635** | 2_27850463_27851734 | CCDC121 | 79635 | 2.16435285578444 | -0.33375135 | 2 | only_DIvsD7 | UpDown | notDM | Loss | 8 |
| **2_27852084_27853243_79635** | 2_27852084_27853243 | CCDC121 | 79635 | 2.31597034545692 | -0.33375135 | 2 | only_DIvsD7 | UpDown | notDM | Loss | 8 |
| **2_27927009_27931260_8884** | 2_27927009_27931260 | SLC5A6 | 8884 | 5.6923101062291 | -0.8459635 | 7 | only_H3vsUI | Down | notDM | Gain | 5 |
| **2_27939131_27941256_9553** | 2_27939131_27941256 | MRPL33 | 9553 | 4.72696344007602 | -0.53824569 | 7 | only_DIvsD7 | DownUp | notDM | Loss | 7 |
| **2_27945900_27948844_9553** | 2_27945900_27948844 | MRPL33 | 9553 | 4.66030565501788 | 0.71965197 | 7 | only_DIvsD7 | DownUp | notDM | Gain | 5 |
| **2_27988545_27989480_9553** | 2_27988545_27989480 | MRPL33 | 9553 | 3.70799574642293 | 0.12357812 | 4 | only_DIvsD7 | DownUp | notDM | Gain | 1 |
| **2_27993619_27994580_9553** | 2_27993619_27994580 | MRPL33 | 9553 | 0.698970004336019 | -0.48832944 | 1 | only_DIvsD7 | DownUp | notDM | Loss | 8 |
| **2_27994673_27998089_9553** | 2_27994673_27998089 | MRPL33 | 9553 | 1.95904139232109 | -0.27157102 | 1 | only_DIvsD7 | DownUp | notDM | Loss | 8 |
| **2_28129871_28130926_26160** | 2_28129871_28130926 | IFT172 | 26160 | 5.62039530450047 | 0.83821303 | 6 | notSpecific | Down | DM | Gain | 9 |
| **2_28129871_28130926_64080** | 2_28129871_28130926 | RBKS | 64080 | 4.22157071216646 | 0.157554 | 6 | notSpecific | Up | DM | Gain | 9 |
| **2_28129871_28130926_9553** | 2_28129871_28130926 | MRPL33 | 9553 | 5.13126248666778 | 0.70019458 | 6 | only_DIvsD7 | DownUp | DM | Gain | 9 |
| **2_28129871_28130926_9577** | 2_28129871_28130926 | BABAM2 | 9577 | 4.20989016868126 | -0.78946913 | 6 | notSpecific | UpDown | DM | Gain | 9 |
| **2_28142058_28147468_9553** | 2_28142058_28147468 | MRPL33 | 9553 | 5.1687213496997 | 0.74860854 | 7 | only_DIvsD7 | DownUp | notDM | Gain | 5 |
| **2_37788111_37790162_54497** | 2_37788111_37790162 | HEATR5B | 54497 | 5.67817955183399 | 0.81877207 | 7 | notSpecific | Down | DM | Gain | 9 |
| **2_38736321_38737386_101929596** | 2_38736321_38737386 | LINC02613 | 101929596 | 3.74012573696573 | 0.00177437 | 6 | notSpecific | UpDown | DM | Gain | 9 |
| **2_38736321_38737386_79833** | 2_38736321_38737386 | GEMIN6 | 79833 | 5.42808131417678 | -0.74698103 | 6 | notSpecific | Down | DM | Gain | 9 |
| **2_38736321_38737386_92906** | 2_38736321_38737386 | HNRNPLL | 92906 | 4.96718747453415 | -0.72029099 | 6 | notSpecific | Up | DM | Gain | 9 |
| **2_42273896_42274808_91461** | 2_42273896_42274808 | PKDCC | 91461 | 2.54777470538782 | 0.85267881 | 1 | only_DIvsD7 | DownUp | notDM | Loss | 6 |
| **2_42275806_42278287_91461** | 2_42275806_42278287 | PKDCC | 91461 | 2.81157500587059 | 0.79961907 | 1 | only_DIvsD7 | DownUp | notDM | Loss | 6 |
| **2_42339201_42340499_91461** | 2_42339201_42340499 | PKDCC | 91461 | 4.80647166760906 | 0.70143765 | 6 | only_DIvsD7 | DownUp | notDM | Loss | 5 |

| | | | | | | | | | | | |
|---|---|---|---|---|---|---|---|---|---|---|---|
| **2_42361756_42367123_91461** | 2_42361756_42367123 | PKDCC | 91461 | 4.93750786200587 | 0.61903697 | 7 | only_DIvsD7 | DownUp | notDM | Loss | 5 |
| **2_42396948_42401044_91461** | 2_42396948_42401044 | PKDCC | 91461 | 5.08561163047165 | 0.84061384 | 1 | only_DIvsD7 | DownUp | notDM | Loss | 6 |
| **2_42721249_42723344_91461** | 2_42721249_42723344 | PKDCC | 91461 | 5.64942346132777 | 0.87927735 | 1 | only_DIvsD7 | DownUp | notDM | Loss | 6 |
| **2_43267191_43270272_102723854** | 2_43267191_43270272 | LINC01819 | 102723854 | 2.70500795933334 | 0.42820999 | 3 | only_H3vsUI | UpDown | notDM | Gain | 1 |
| **2_43277820_43279969_102723854** | 2_43277820_43279969 | LINC01819 | 102723854 | 4.04696315412342 | 0.61226847 | 6 | only_H3vsUI | UpDown | notDM | Gain | 1 |
| **2_43336431_43338861_102723854** | 2_43336431_43338861 | LINC01819 | 102723854 | 4.84352553221136 | 0.99845224 | 6 | only_H3vsUI | UpDown | notDM | Gain | 1 |
| **2_43357013_43359631_102723854** | 2_43357013_43359631 | LINC01819 | 102723854 | 4.95582720234041 | 0.609818 | 3 | only_H3vsUI | UpDown | notDM | Loss | 5 |
| **2_43400165_43403241_102723854** | 2_43400165_43403241 | LINC01819 | 102723854 | 5.12543246582618 | -0.75026616 | 6 | only_H3vsUI | UpDown | notDM | Loss | 3 |
| **2_43638629_43640451_102723854** | 2_43638629_43640451 | LINC01819 | 102723854 | 5.57048573065805 | 0.82481306 | 6 | only_H3vsUI | UpDown | notDM | Gain | 1 |
| **2_61106998_61108273_400957** | 2_61106998_61108273 | LINC01185 | 400957 | 2.24797326636181 | 0.00236632 | 1 | only_H3vsUI | Up | notDM | Loss | 3 |
| **2_61108833_61115322_400957** | 2_61108833_61115322 | LINC01185 | 400957 | 2.58658730467176 | 0.00264077 | 1 | only_H3vsUI | Up | notDM | Gain | 4 |
| **2_64242555_64245993_51542** | 2_64242555_64245993 | VPS54 | 51542 | 2.75966784468963 | 0.27729356 | 1 | only_H3vsUI | Down | notDM | Loss | 8 |
| **2_64246680_64248156_51542** | 2_64246680_64248156 | VPS54 | 51542 | 2.06069784035361 | 0.72552222 | 5 | only_H3vsUI | Down | notDM | Gain | 4 |
| **2_65085512_65088051_10097** | 2_65085512_65088051 | ACTR2 | 10097 | 5.56447313311966 | 0.77903814 | 1 | only_H3vsUI | DownUp | notDM | Gain | 5 |
| **2_65169888_65171974_10097** | 2_65169888_65171974 | ACTR2 | 10097 | 5.45170816338798 | 0.75053119 | 6 | only_H3vsUI | DownUp | notDM | Gain | 5 |
| **2_65291182_65293544_10097** | 2_65291182_65293544 | ACTR2 | 10097 | 5.20797886614084 | 0.8965841 | 4 | only_H3vsUI | DownUp | notDM | Loss | 5 |
| **2_65452897_65454626_10097** | 2_65452897_65454626 | ACTR2 | 10097 | 2.41830129131975 | 0.18210097 | 2 | only_H3vsUI | DownUp | notDM | Loss | 8 |
| **2_65455383_65460017_10097** | 2_65455383_65460017 | ACTR2 | 10097 | 2.69722934275972 | 0.26226105 | 1 | only_H3vsUI | DownUp | notDM | Gain | 4 |
| **2_65638174_65640953_10097** | 2_65638174_65640953 | ACTR2 | 10097 | 5.26313640177264 | -0.84807952 | 6 | only_H3vsUI | DownUp | notDM | Loss | 7 |
| **2_68349407_68350371_116143** | 2_68349407_68350371 | WDR92 | 116143 | 4.53447033220334 | -0.50980918 | 6 | only_H3vsUI | Down | notDM | Gain | 2 |
| **2_68382793_68384514_116143** | 2_68382793_68384514 | WDR92 | 116143 | 2.00432137378264 | -0.3578401 | 1 | only_H3vsUI | Down | notDM | Gain | 4 |
| **2_68385178_68387339_116143** | 2_68385178_68387339 | WDR92 | 116143 | 2.68841982200271 | -0.35698955 | 1 | only_H3vsUI | Down | notDM | Gain | 4 |
| **2_68417685_68423412_116143** | 2_68417685_68423412 | WDR92 | 116143 | 4.51947359087236 | -0.41807772 | 8 | only_H3vsUI | Down | notDM | Gain | 5 |
| **2_68443619_68446886_116143** | 2_68443619_68446886 | WDR92 | 116143 | 4.77096977045032 | -0.88126814 | 8 | only_H3vsUI | Down | notDM | Gain | 5 |
| **2_68869953_68871650_10887** | 2_68869953_68871650 | PROKR1 | 10887 | 0 | 0.00100681 | 4 | only_H3vsUI | Up | notDM | Loss | 3 |
| **2_69662523_69664316_27247** | 2_69662523_69664316 | NFU1 | 27247 | 2.36921585741014 | -0.17105422 | 1 | only_H3vsUI | DownUp | notDM | Loss | 8 |
| **2_69664674_69665531_27247** | 2_69664674_69665531 | NFU1 | 27247 | 0 | 0.12367123 | 1 | only_H3vsUI | DownUp | notDM | Gain | 4 |
| **2_69710078_69712764_27247** | 2_69710078_69712764 | NFU1 | 27247 | 4.65830718884789 | 0.58596594 | 6 | only_H3vsUI | DownUp | notDM | Gain | 5 |
| **2_69792604_69797034_27247** | 2_69792604_69797034 | NFU1 | 27247 | 5.10734227359317 | -0.68446396 | 3 | only_H3vsUI | DownUp | notDM | Loss | 7 |
| **2_69901588_69903993_22848** | 2_69901588_69903993 | AAK1 | 22848 | 2.03742649794062 | 0.01758425 | 6 | notSpecific | Down | DM | Gain | 9 |
| **2_70120294_70121071_27247** | 2_70120294_70121071 | NFU1 | 27247 | 5.65872381854976 | -0.83823333 | 4 | only_H3vsUI | DownUp | notDM | Gain | 2 |
| **2_70121276_70123042_27247** | 2_70121276_70123042 | NFU1 | 27247 | 5.65965858800315 | -0.83823333 | 1 | only_H3vsUI | DownUp | notDM | Loss | 7 |
| **2_71088698_71089134_25806** | 2_71088698_71089134 | VAX2 | 25806 | 4.58644101505516 | 0.45984403 | 6 | only_DIvsD7 | UpDown | notDM | Loss | 3 |
| **2_71098173_71099808_25806** | 2_71098173_71099808 | VAX2 | 25806 | 4.44580651557958 | 0.61418565 | 4 | only_DIvsD7 | UpDown | notDM | Gain | 1 |

| | | | | | | | | | | | |
|---|---|---|---|---|---|---|---|---|---|---|---|
| **2_71113879_71115432_25806** | 2_71113879_71115432 | VAX2 | 25806 | 4.08951654422482 | 0.14151648 | 2 | only_DIvsD7 | UpDown | notDM | Loss | 6 |
| **2_71126809_71130057_25806** | 2_71126809_71130057 | VAX2 | 25806 | 0 | 0.72727386 | 1 | only_DIvsD7 | UpDown | notDM | Loss | 6 |
| **2_72371755_72375625_56603** | 2_72371755_72375625 | CYP26B1 | 56603 | 0 | -0.00119978 | 2 | notSpecific | DownUp | DM | Loss | 10 |
| **2_73144629_73146285_2016** | 2_73144629_73146285 | EMX1 | 2016 | 1.43136376415899 | 0.02900515 | 4 | notSpecific | Up | DM | Loss | 10 |
| **2_73144629_73146285_6697** | 2_73144629_73146285 | SPR | 6697 | 4.47885496752866 | -0.59928357 | 4 | notSpecific | Down | DM | Loss | 10 |
| **2_73495884_73496495_10574** | 2_73495884_73496495 | CCT7 | 10574 | 4.53683610026608 | 0.76911906 | 2 | notSpecific | DownUp | DM | Loss | 10 |
| **2_73495884_73496495_150726** | 2_73495884_73496495 | FBXO41 | 150726 | 3.19005141775921 | 0.00228891 | 2 | notSpecific | DownUp | DM | Loss | 10 |
| **2_73495884_73496495_94097** | 2_73495884_73496495 | SFXN5 | 94097 | 5.29467560702956 | 0.80564811 | 2 | notSpecific | Down | DM | Loss | 10 |
| **2_74197823_74198563_100874048** | 2_74197823_74198563 | DGUOK-AS1 | 100874048 | 4.00026049854739 | 0.73618308 | 6 | only_H3vsUI | UpDown | DM | Loss | 10 |
| **2_74206317_74209823_100874048** | 2_74206317_74209823 | DGUOK-AS1 | 100874048 | 0 | 0.5349108 | 2 | only_H3vsUI | UpDown | notDM | Gain | 4 |
| **2_74210384_74211847_100874048** | 2_74210384_74211847 | DGUOK-AS1 | 100874048 | 3.27829620809127 | -0.19394076 | 2 | only_H3vsUI | UpDown | notDM | Loss | 3 |
| **2_74212725_74215405_100874048** | 2_74212725_74215405 | DGUOK-AS1 | 100874048 | 3.62726341656822 | -0.18959489 | 1 | only_H3vsUI | UpDown | notDM | Loss | 6 |
| **2_74226380_74228488_100874048** | 2_74226380_74228488 | DGUOK-AS1 | 100874048 | 4.25270743300728 | 0.52247054 | 6 | only_H3vsUI | UpDown | notDM | Loss | 5 |
| **2_88924459_88926451_101928403** | 2_88924459_88926451 | EIF2AK3-DT | 101928403 | 2.78318869107526 | 0 | 1 | only_H3vsUI | Up | notDM | Loss | 6 |
| **2_88927396_88928954_101928403** | 2_88927396_88928954 | EIF2AK3-DT | 101928403 | 2.5327543789925 | 0 | 1 | only_H3vsUI | Up | notDM | Gain | 4 |
| **2_95824227_95825271_84874** | 2_95824227_95825271 | ZNF514 | 84874 | 1.91381385238372 | 0.03544079 | 2 | only_H3vsUI | Down | notDM | Loss | 8 |
| **2_95829891_95831173_84874** | 2_95829891_95831173 | ZNF514 | 84874 | 0 | 0.06095318 | 1 | only_H3vsUI | Down | notDM | Loss | 8 |
| **2_95831599_95832914_84874** | 2_95831599_95832914 | ZNF514 | 84874 | 2.64640372622307 | 0.06095318 | 1 | only_H3vsUI | Down | notDM | Gain | 4 |
| **2_98702194_98704456_200403** | 2_98702194_98704456 | VWA3B | 200403 | 0 | 0 | 2 | only_H3vsUI | UpDown | notDM | Loss | 3 |
| **2_99223001_99224677_493753** | 2_99223001_99224677 | COA5 | 493753 | 2.4456042032736 | -0.16592546 | 1 | only_H3vsUI | Down | notDM | Loss | 8 |
| **2_99225243_99227315_493753** | 2_99225243_99227315 | COA5 | 493753 | 2.46239799789896 | -0.16592546 | 1 | only_H3vsUI | Down | notDM | Loss | 8 |
| **2_99458947_99460261_493753** | 2_99458947_99460261 | COA5 | 493753 | 5.36916203125429 | -0.81343963 | 6 | only_H3vsUI | Down | notDM | Gain | 2 |
| **2_99527711_99530809_493753** | 2_99527711_99530809 | COA5 | 493753 | 5.48107411020809 | -0.86819857 | 6 | only_H3vsUI | Down | notDM | Loss | 2 |
| **2_99533165_99534106_493753** | 2_99533165_99534106 | COA5 | 493753 | 5.48882840696583 | -0.92686513 | 7 | only_H3vsUI | Down | notDM | Gain | 2 |
| **2_9982388_9983483_3241** | 2_9982388_9983483 | HPCAL1 | 3241 | 5.66231670441265 | 0.80021702 | 1 | only_DIvsD7 | Up | notDM | Gain | 1 |
| **20_18037270_18040489_58495** | 20_18037270_18040489 | OVOL2 | 58495 | 0 | 0.18553918 | 4 | only_H3vsUI | DownUp | notDM | Loss | 3 |
| **20_22774156_22777639_22918** | 20_22774156_22777639 | CD93 | 22918 | 5.46138295966264 | 0.79946936 | 7 | only_DIvsD7 | DownUp | notDM | Gain | 5 |
| **20_22989733_22992025_7056** | 20_22989733_22992025 | THBD | 7056 | 4.58333482403455 | -0.55521361 | 7 | notSpecific | Down | DM | Gain | 9 |
| **20_22992577_22995306_22918** | 20_22992577_22995306 | CD93 | 22918 | 4.85534952358276 | 0.7510301 | 6 | only_DIvsD7 | DownUp | notDM | Gain | 5 |
| **20_23026288_23032067_22918** | 20_23026288_23032067 | CD93 | 22918 | 4.54276320266193 | 0.42577093 | 1 | only_DIvsD7 | DownUp | notDM | Gain | 5 |
| **20_23340957_23342430_22918** | 20_23340957_23342430 | CD93 | 22918 | 5.43772203158879 | 0.98899474 | 1 | only_DIvsD7 | DownUp | notDM | Gain | 1 |
| **20_23343010_23344867_22918** | 20_23343010_23344867 | CD93 | 22918 | 5.44096415214913 | 0.87161664 | 1 | only_DIvsD7 | DownUp | notDM | Loss | 6 |
| **20_2633689_2637755_10528** | 20_2633689_2637755 | NOP56 | 10528 | 0 | -0.30351662 | 1 | notSpecific | Up | DM | Loss | 10 |
| **20_2633689_2637755_3420** | 20_2633689_2637755 | IDH3B | 3420 | 3.85003325768977 | -0.46237333 | 1 | notSpecific | DownUp | DM | Loss | 10 |

| | | | | | | | | | | | |
|---|---|---|---|---|---|---|---|---|---|---|---|
| **20_30448172_30450148_128853** | 20_30448172_30450148 | DUSP15 | 128853 | 3.89960165914612 | 0.07703403 | 2 | only_H3vsUI | Up | notDM | Loss | 3 |
| **20_30457332_30459041_128853** | 20_30457332_30459041 | DUSP15 | 128853 | 0 | -0.05741228 | 2 | only_H3vsUI | Up | notDM | Loss | 3 |
| **20_30466580_30467509_128853** | 20_30466580_30467509 | DUSP15 | 128853 | 3.91418443342325 | -0.09203265 | 1 | only_H3vsUI | Up | notDM | Loss | 3 |
| **20_33813222_33814373_55245** | 20_33813222_33814373 | UQCC1 | 55245 | 5.26825493885189 | -0.76026851 | 2 | only_H3vsUI | DownUp | notDM | Loss | 7 |
| **20_33891835_33896879_55245** | 20_33891835_33896879 | UQCC1 | 55245 | 0 | 0.22600103 | 7 | only_H3vsUI | DownUp | notDM | Gain | 4 |
| **20_33898715_33901122_55245** | 20_33898715_33901122 | UQCC1 | 55245 | 3.5864747785714 | 0.12720696 | 7 | only_H3vsUI | DownUp | notDM | Gain | 5 |
| **20_33905049_33909311_55245** | 20_33905049_33909311 | UQCC1 | 55245 | 4.008302024212 | 0.19939618 | 7 | only_H3vsUI | DownUp | notDM | Loss | 5 |
| **20_33998092_33999685_55245** | 20_33998092_33999685 | UQCC1 | 55245 | 2.17609125905568 | -0.37101135 | 2 | only_H3vsUI | DownUp | notDM | Loss | 8 |
| **20_33999866_34000513_55245** | 20_33999866_34000513 | UQCC1 | 55245 | 1.53147891704226 | -0.37101135 | 3 | only_H3vsUI | DownUp | notDM | Gain | 4 |
| **20_3745995_3748412_1059** | 20_3745995_3748412 | CENPB | 1059 | 4.23902404660009 | -0.52689835 | 1 | notSpecific | Up | DM | Loss | 10 |
| **20_3745995_3748412_54976** | 20_3745995_3748412 | C20orf27 | 54976 | 0 | -0.81519494 | 1 | notSpecific | UpDown | DM | Loss | 10 |
| **20_3745995_3748412_57506** | 20_3745995_3748412 | MAVS | 57506 | 4.89806116876976 | -0.66414515 | 1 | notSpecific | Down | DM | Loss | 10 |
| **20_3745995_3748412_994** | 20_3745995_3748412 | CDC25B | 994 | 4.45525610211795 | 0.63435504 | 1 | notSpecific | DownUp | DM | Loss | 10 |
| **20_3777188_3781302_55317** | 20_3777188_3781302 | AP5S1 | 55317 | 4.29835083771916 | 0.27301007 | 1 | notSpecific | Up | DM | Loss | 10 |
| **20_3777188_3781302_994** | 20_3777188_3781302 | CDC25B | 994 | 2.40140054078154 | 0.58166184 | 1 | notSpecific | DownUp | DM | Loss | 10 |
| **20_39958201_39959247_64900** | 20_39958201_39959247 | LPIN3 | 64900 | 4.0134271270707 | 0.04260169 | 3 | only_H3vsUI | Down | notDM | Loss | 2 |
| **20_39968299_39969327_64900** | 20_39968299_39969327 | LPIN3 | 64900 | 2.36921585741014 | 0.03579929 | 4 | only_H3vsUI | Down | notDM | Gain | 4 |
| **20_39969455_39970987_64900** | 20_39969455_39970987 | LPIN3 | 64900 | 0 | 0.19874771 | 1 | only_H3vsUI | Down | notDM | Gain | 4 |
| **20_42834803_42839028_51526** | 20_42834803_42839028 | OSER1 | 51526 | 2.60638136511061 | -0.03220875 | 1 | only_DIvsD7 | Up | notDM | Gain | 4 |
| **20_42839733_42840944_51526** | 20_42839733_42840944 | OSER1 | 51526 | 2.48287358360875 | 0.01509027 | 1 | only_DIvsD7 | Up | notDM | Gain | 4 |
| **20_42983117_42986141_51526** | 20_42983117_42986141 | OSER1 | 51526 | 5.15742049985729 | 0.78509488 | 1 | only_DIvsD7 | Up | notDM | Loss | 6 |
| **20_43012650_43013678_51526** | 20_43012650_43013678 | OSER1 | 51526 | 5.23860054144153 | 0.79269899 | 7 | only_DIvsD7 | Up | notDM | Loss | 5 |
| **20_43028837_43030996_51526** | 20_43028837_43030996 | OSER1 | 51526 | 5.27739831829176 | 0.79440202 | 1 | only_DIvsD7 | Up | notDM | Loss | 6 |
| **20_43940075_43940974_11317** | 20_43940075_43940974 | RBPJL | 11317 | 3.23095955574857 | 0 | 4 | only_DIvsD7 | DownUp | notDM | Loss | 3 |
| **20_48779958_48782660_101927512** | 20_48779958_48782660 | LINC01275 | 101927512 | 0 | 0 | 5 | only_DIvsD7 | DownUp | notDM | Gain | 4 |
| **20_48782824_48784289_101927512** | 20_48782824_48784289 | LINC01275 | 101927512 | 3.45101845215546 | 0 | 5 | only_DIvsD7 | DownUp | notDM | Gain | 1 |
| **20_48935717_48939200_101927586** | 20_48935717_48939200 | LINC01271 | 101927586 | 0 | 0.0020427 | 1 | only_H3vsUI | DownUp | notDM | Loss | 3 |
| **20_52722363_52725204_1591** | 20_52722363_52725204 | CYP24A1 | 1591 | 4.81497303397042 | 0.946296 | 7 | notSpecific | Down | DM | Gain | 9 |
| **20_52722363_52725204_8537** | 20_52722363_52725204 | BCAS1 | 8537 | 4.54754088621962 | 0.40619487 | 7 | notSpecific | Up | DM | Gain | 9 |
| **20_52737328_52740047_1591** | 20_52737328_52740047 | CYP24A1 | 1591 | 4.70299888335918 | 0.72186035 | 7 | notSpecific | Down | DM | Gain | 9 |
| **20_5493296_5495694_101929207** | 20_5493296_5495694 | LINC01729 | 101929207 | 3.42324587393681 | 0 | 3 | only_DIvsD7 | Up | notDM | Gain | 1 |
| **20_5493296_5495694_149837** | 20_5493296_5495694 | LINC00654 | 149837 | 3.90520202866232 | 0.23331294 | 3 | only_H3vsUI | UpDown | notDM | Gain | 1 |
| **20_55839363_55841276_655** | 20_55839363_55841276 | BMP7 | 655 | 2.61278385671974 | 0.17095998 | 4 | only_H3vsUI | Up | notDM | Loss | 3 |
| **20_5700068_5702072_149840** | 20_5700068_5702072 | SHLD1 | 149840 | 4.46191851180896 | 0.38893046 | 6 | only_DIvsD7 | Up | notDM | Loss | 5 |

| | | | | | | | | | | | |
|---|---|---|---|---|---|---|---|---|---|---|---|
| **20_5714770_5719741_149840** | 20_5714770_5719741 | SHLD1 | 149840 | 4.05534009954418 | -0.09435911 | 7 | only_DIvsD7 | Up | notDM | Gain | 5 |
| **20_5731283_5732636_149840** | 20_5731283_5732636 | SHLD1 | 149840 | 2.26951294421792 | 0.49537436 | 1 | only_DIvsD7 | Up | notDM | Gain | 4 |
| **20_5759998_5761975_149840** | 20_5759998_5761975 | SHLD1 | 149840 | 4.46181355362262 | 0.85435178 | 3 | only_DIvsD7 | Up | notDM | Loss | 5 |
| **20_5766464_5769109_149840** | 20_5766464_5769109 | SHLD1 | 149840 | 4.54933437732304 | 0.72625459 | 3 | only_DIvsD7 | Up | notDM | Gain | 5 |
| **20_6102104_6104823_149840** | 20_6102104_6104823 | SHLD1 | 149840 | 5.56938210377325 | -0.78754017 | 2 | only_DIvsD7 | Up | notDM | Loss | 6 |
| **20_61147752_61148735_128826** | 20_61147752_61148735 | MIR1-1HG | 128826 | 1.9731278535997 | 0 | 4 | only_H3vsUI | UpDown | notDM | Loss | 3 |
| **20_61491221_61492272_63910** | 20_61491221_61492272 | SLC17A9 | 63910 | 4.96275278524049 | -0.6271513 | 2 | only_H3vsUI | DownUp | notDM | Loss | 3 |
| **20_61548029_61548584_63910** | 20_61548029_61548584 | SLC17A9 | 63910 | 4.54984894430493 | 0.74124383 | 8 | only_H3vsUI | DownUp | notDM | Gain | 1 |
| **20_61553519_61557387_63910** | 20_61553519_61557387 | SLC17A9 | 63910 | 4.43155674104815 | 0.3662522 | 1 | only_H3vsUI | DownUp | notDM | Loss | 6 |
| **20_61557924_61559197_63910** | 20_61557924_61559197 | SLC17A9 | 63910 | 4.40143500722987 | 0.49545604 | 1 | only_H3vsUI | DownUp | notDM | Loss | 3 |
| **20_61566824_61568984_63910** | 20_61566824_61568984 | SLC17A9 | 63910 | 4.17808443287921 | 0.58408517 | 1 | only_H3vsUI | DownUp | notDM | Loss | 6 |
| **20_61569858_61571518_63910** | 20_61569858_61571518 | SLC17A9 | 63910 | 4.10994958023059 | 0.76273861 | 1 | only_H3vsUI | DownUp | notDM | Gain | 1 |
| **20_61885347_61885790_63910** | 20_61885347_61885790 | SLC17A9 | 63910 | 5.47849579073694 | -0.78909412 | 4 | only_H3vsUI | DownUp | notDM | Loss | 7 |
| **20_62152578_62153205_54923** | 20_62152578_62153205 | LIME1 | 54923 | 5.33516492894019 | -0.90265696 | 2 | only_H3vsUI | DownUp | notDM | Loss | 7 |
| **20_62167713_62169381_54923** | 20_62167713_62169381 | LIME1 | 54923 | 5.30141634585253 | -0.79756582 | 5 | only_H3vsUI | DownUp | notDM | Gain | 2 |
| **20_62313077_62315965_54923** | 20_62313077_62315965 | LIME1 | 54923 | 4.72911617191763 | -0.59897029 | 7 | only_H3vsUI | DownUp | notDM | Loss | 7 |
| **20_62329184_62331824_54923** | 20_62329184_62331824 | LIME1 | 54923 | 4.57674435432742 | -0.72026686 | 7 | only_H3vsUI | DownUp | notDM | Loss | 7 |
| **20_62337379_62338948_54923** | 20_62337379_62338948 | LIME1 | 54923 | 4.4635047861021 | 0.3603382 | 1 | only_H3vsUI | DownUp | notDM | Gain | 4 |
| **20_62339754_62340896_54923** | 20_62339754_62340896 | LIME1 | 54923 | 4.43338575741794 | 0.35720272 | 1 | only_H3vsUI | DownUp | notDM | Gain | 4 |
| **20_62371723_62373609_54923** | 20_62371723_62373609 | LIME1 | 54923 | 3.57170883180869 | 0.22777239 | 5 | only_H3vsUI | DownUp | notDM | Loss | 6 |
| **20_6748505_6749562_650** | 20_6748505_6749562 | BMP2 | 650 | 2.29225607135648 | 0.13292718 | 4 | notSpecific | Up | DM | Loss | 10 |
| **20_7999196_7999979_56255** | 20_7999196_7999979 | TMX4 | 56255 | 2.69722934275972 | -0.37547955 | 2 | only_H3vsUI | DownUp | notDM | Loss | 8 |
| **20_8000504_8001325_56255** | 20_8000504_8001325 | TMX4 | 56255 | 1.47712125471966 | -0.37547955 | 2 | only_H3vsUI | DownUp | notDM | Gain | 4 |
| **20_8112160_8112757_56255** | 20_8112160_8112757 | TMX4 | 56255 | 5.04809205181237 | 0.70768328 | 4 | only_H3vsUI | DownUp | notDM | Loss | 3 |
| **21_30363490_30364856_10069** | 21_30363490_30364856 | RWDD2B | 10069 | 4.42842638644059 | 0.4555004 | 1 | only_H3vsUI | DownUp | notDM | Gain | 4 |
| **21_30365436_30366214_10069** | 21_30365436_30366214 | RWDD2B | 10069 | 4.40585839931764 | 0.4555004 | 5 | only_H3vsUI | DownUp | notDM | Gain | 1 |
| **21_30389904_30391351_10069** | 21_30389904_30391351 | RWDD2B | 10069 | 2.5092025223311 | 0.57660317 | 1 | only_H3vsUI | DownUp | notDM | Loss | 6 |
| **21_30391941_30392982_10069** | 21_30391941_30392982 | RWDD2B | 10069 | 2.43136376415899 | 0.57647549 | 2 | only_H3vsUI | DownUp | notDM | Loss | 3 |
| **21_30396331_30396833_10069** | 21_30396331_30396833 | RWDD2B | 10069 | 3.66595602945396 | 0.63389444 | 2 | only_H3vsUI | DownUp | notDM | Gain | 4 |
| **21_30397450_30398744_10069** | 21_30397450_30398744 | RWDD2B | 10069 | 3.7598943740255 | 0.63389444 | 2 | only_H3vsUI | DownUp | notDM | Loss | 6 |
| **21_30449430_30451297_10069** | 21_30449430_30451297 | RWDD2B | 10069 | 4.76161966554771 | 0.72009647 | 5 | only_H3vsUI | DownUp | notDM | Loss | 7 |
| **21_30671737_30675948_571** | 21_30671737_30675948 | BACH1 | 571 | 2.73798732633343 | -0.61923529 | 1 | only_H3vsUI | Up | notDM | Gain | 4 |
| **21_40683519_40684390_103091865** | 21_40683519_40684390 | BRWD1-AS2 | 103091865 | 3.29666519026153 | -0.00703885 | 2 | only_H3vsUI | DownUp | notDM | Gain | 4 |
| **21_40683519_40684390_3150** | 21_40683519_40684390 | HMGN1 | 3150 | 4.55791604022211 | -0.6674084 | 2 | only_DIvsD7 | Down | notDM | Gain | 2 |

| | | | | | | | | | | | |
|---|---|---|---|---|---|---|---|---|---|---|---|
| **21_40686344_40687080_103091865** | 21_40686344_40687080 | BRWD1-AS2 | 103091865 | 0 | -0.00486987 | 5 | only_H3vsUI | DownUp | notDM | Gain | 4 |
| **21_40686344_40687080_3150** | 21_40686344_40687080 | HMGN1 | 3150 | 4.52431821476093 | -0.88327849 | 5 | only_DIvsD7 | Down | notDM | Gain | 2 |
| **21_40717630_40720358_3150** | 21_40717630_40720358 | HMGN1 | 3150 | 2.22010808804005 | -0.43887335 | 2 | only_DIvsD7 | Down | notDM | Loss | 8 |
| **21_40721495_40722849_3150** | 21_40721495_40722849 | HMGN1 | 3150 | 2.98855895687862 | -0.43887335 | 5 | only_DIvsD7 | Down | notDM | Loss | 7 |
| **21_40752424_40753974_3150** | 21_40752424_40753974 | HMGN1 | 3150 | 4.49810369763808 | -0.80084761 | 1 | only_DIvsD7 | Down | notDM | Loss | 7 |
| **21_40757903_40759511_3150** | 21_40757903_40759511 | HMGN1 | 3150 | 4.5726625331511 | -0.85703634 | 5 | only_DIvsD7 | Down | notDM | Gain | 2 |
| **21_40759754_40761129_3150** | 21_40759754_40761129 | HMGN1 | 3150 | 4.58857420665492 | -0.57069416 | 1 | only_DIvsD7 | Down | notDM | Gain | 2 |
| **21_40816399_40817497_3150** | 21_40816399_40817497 | HMGN1 | 3150 | 4.98171896614152 | -0.63217428 | 2 | only_DIvsD7 | Down | notDM | Loss | 7 |
| **21_40817888_40818661_3150** | 21_40817888_40818661 | HMGN1 | 3150 | 4.98841178905203 | -0.7037495 | 1 | only_DIvsD7 | Down | notDM | Loss | 7 |
| **21_41003694_41006895_3150** | 21_41003694_41006895 | HMGN1 | 3150 | 5.45205184184959 | 0.78379452 | 7 | only_DIvsD7 | Down | notDM | Gain | 5 |
| **21_44297569_44299158_10785** | 21_44297569_44299158 | WDR4 | 10785 | 2.68484536164441 | 0.26913907 | 1 | only_DIvsD7 | UpDown | notDM | Loss | 6 |
| **21_44312684_44313171_10785** | 21_44312684_44313171 | WDR4 | 10785 | 4.11614265385379 | -0.16809308 | 1 | only_DIvsD7 | UpDown | notDM | Gain | 2 |
| **21_45553677_45555114_23308** | 21_45553677_45555114 | ICOSLG | 23308 | 5.02422287699035 | 0.63778147 | 2 | only_H3vsUI | Up | notDM | Loss | 6 |
| **21_45659568_45660352_23308** | 21_45659568_45660352 | ICOSLG | 23308 | 2.69722934275972 | 0.70614255 | 2 | only_H3vsUI | Up | notDM | Loss | 3 |
| **21_46358752_46359677_85395** | 21_46358752_46359677 | FAM207A | 85395 | 2.39619934709574 | 0.26603861 | 1 | only_H3vsUI | UpDown | notDM | Gain | 4 |
| **21_46360329_46361941_85395** | 21_46360329_46361941 | FAM207A | 85395 | 2.60852603357719 | 0.53611593 | 1 | only_H3vsUI | UpDown | notDM | Gain | 4 |
| **21_46432012_46433553_85395** | 21_46432012_46433553 | FAM207A | 85395 | 4.8578690011444 | -0.57477305 | 6 | only_H3vsUI | UpDown | notDM | Loss | 3 |
| **21_46826005_46828087_85395** | 21_46826005_46828087 | FAM207A | 85395 | 5.66844835368292 | -0.83251882 | 1 | only_H3vsUI | UpDown | notDM | Loss | 7 |
| **22_18110056_18111259_55670** | 22_18110056_18111259 | PEX26 | 55670 | 5.6527451545521 | -0.9043432 | 1 | only_H3vsUI | DownUp | notDM | Gain | 2 |
| **22_18111759_18112787_55670** | 22_18111759_18112787 | PEX26 | 55670 | 5.65126638095444 | -0.9043432 | 2 | only_H3vsUI | DownUp | notDM | Loss | 7 |
| **22_18119943_18121332_55670** | 22_18119943_18121332 | PEX26 | 55670 | 5.6429025508754 | -0.86223585 | 1 | only_H3vsUI | DownUp | notDM | Loss | 7 |
| **22_18121944_18123031_55670** | 22_18121944_18123031 | PEX26 | 55670 | 5.64122020207759 | -0.86223585 | 1 | only_H3vsUI | DownUp | notDM | Gain | 2 |
| **22_18518272_18519701_55670** | 22_18518272_18519701 | PEX26 | 55670 | 4.61356699877128 | -0.45032831 | 6 | only_H3vsUI | DownUp | notDM | Gain | 2 |
| **22_18542970_18544098_55670** | 22_18542970_18544098 | PEX26 | 55670 | 4.22211792875822 | -0.20513463 | 7 | only_H3vsUI | DownUp | notDM | Gain | 1 |
| **22_18559118_18560594_55670** | 22_18559118_18560594 | PEX26 | 55670 | 1.98227123303957 | 0.11690727 | 1 | only_H3vsUI | DownUp | notDM | Loss | 8 |
| **22_18573996_18576005_55670** | 22_18573996_18576005 | PEX26 | 55670 | 4.12136284056339 | 0.30922317 | 7 | only_H3vsUI | DownUp | notDM | Loss | 5 |
| **22_18632905_18634327_11274** | 22_18632905_18634327 | USP18 | 11274 | 2.38201704257487 | 0.50284122 | 2 | only_DIvsD7 | UpDown | notDM | Gain | 4 |
| **22_20103314_20104354_27037** | 22_20103314_20104354 | TRMT2A | 27037 | 2.61172330800734 | -0.39014682 | 1 | only_H3vsUI | UpDown | notDM | Loss | 3 |
| **22_20105616_20107738_27037** | 22_20105616_20107738 | TRMT2A | 27037 | 2.90308998699194 | -0.16992006 | 1 | only_H3vsUI | UpDown | notDM | Loss | 6 |
| **22_20193400_20195375_27037** | 22_20193400_20195375 | TRMT2A | 27037 | 4.94760034919971 | 0.79428623 | 1 | only_H3vsUI | UpDown | notDM | Gain | 1 |
| **22_21304638_21306024_150209** | 22_21304638_21306024 | AIFM3 | 150209 | 4.12801133257597 | 0.34292535 | 8 | only_H3vsUI | UpDown | notDM | Loss | 3 |
| **22_21310153_21311381_150209** | 22_21310153_21311381 | AIFM3 | 150209 | 3.90692734730896 | 0.33305238 | 1 | only_H3vsUI | UpDown | notDM | Loss | 3 |
| **22_21335402_21336193_150209** | 22_21335402_21336193 | AIFM3 | 150209 | 3.73423960443546 | -0.00137456 | 5 | only_H3vsUI | UpDown | notDM | Loss | 3 |
| **22_23907957_23908480_23523** | 22_23907957_23908480 | CABIN1 | 23523 | 5.69837809206785 | -0.81848874 | 4 | only_DIvsD7 | Down | notDM | Loss | 7 |

| | | | | | | | | | | | |
|---|---|---|---|---|---|---|---|---|---|---|---|
| **22_24408138_24409520_23523** | 22_24408138_24409520 | CABIN1 | 23523 | 2.43775056282039 | -0.62387591 | 1 | only_DIvsD7 | Down | notDM | Gain | 4 |
| **22_24541286_24543098_23523** | 22_24541286_24543098 | CABIN1 | 23523 | 3.93996829055134 | 0.25919684 | 7 | only_DIvsD7 | Down | notDM | Loss | 5 |
| **22_24553133_24553727_23523** | 22_24553133_24553727 | CABIN1 | 23523 | 2.80208925788173 | 0.23153065 | 3 | only_DIvsD7 | Down | notDM | Gain | 1 |
| **22_24670924_24671264_23523** | 22_24670924_24671264 | CABIN1 | 23523 | 5.42016706472756 | 0.74467748 | 8 | only_DIvsD7 | Down | notDM | Gain | 2 |
| **22_24684575_24686122_23523** | 22_24684575_24686122 | CABIN1 | 23523 | 5.44213313589275 | 0.75183327 | 6 | only_DIvsD7 | Down | notDM | Gain | 2 |
| **22_24949273_24950891_388886** | 22_24949273_24950891 | LRRC75B | 388886 | 4.58138068870999 | 0.54027817 | 1 | only_H3vsUI | UpDown | notDM | Loss | 6 |
| **22_24952132_24954178_388886** | 22_24952132_24954178 | LRRC75B | 388886 | 4.54224016628972 | 0.47442968 | 1 | only_H3vsUI | UpDown | notDM | Loss | 6 |
| **22_24973983_24976065_388886** | 22_24973983_24976065 | LRRC75B | 388886 | 4.11280601726688 | 0.75614749 | 6 | only_H3vsUI | UpDown | notDM | Gain | 1 |
| **22_24995506_25000928_388886** | 22_24995506_25000928 | LRRC75B | 388886 | 3.81144094367416 | -0.39679617 | 5 | only_H3vsUI | UpDown | notDM | Loss | 6 |
| **22_25002232_25005608_388886** | 22_25002232_25005608 | LRRC75B | 388886 | 4.12070551565827 | -0.6193619 | 5 | only_H3vsUI | UpDown | notDM | Loss | 6 |
| **22_25796139_25800548_1416** | 22_25796139_25800548 | CRYBB2P1 | 1416 | 4.63873877978937 | -0.7209961 | 1 | only_H3vsUI | Down | notDM | Loss | 2 |
| **22_25844300_25845773_1416** | 22_25844300_25845773 | CRYBB2P1 | 1416 | 2.36172783601759 | -0.2754982 | 1 | only_H3vsUI | Down | notDM | Loss | 8 |
| **22_28990339_28991936_150274** | 22_28990339_28991936 | HSCB | 150274 | 5.16658994742734 | 0.69796428 | 6 | notSpecific | Up | DM | Gain | 9 |
| **22_29074313_29075309_11200** | 22_29074313_29075309 | CHEK2 | 11200 | 4.51621796516111 | 0.5045875 | 1 | only_H3vsUI | DownUp | notDM | Gain | 4 |
| **22_29076562_29077568_11200** | 22_29076562_29077568 | CHEK2 | 11200 | 4.48525281696033 | -0.43751871 | 6 | only_H3vsUI | DownUp | notDM | Gain | 2 |
| **22_29105522_29109335_11200** | 22_29105522_29109335 | CHEK2 | 11200 | 0 | 0.41232802 | 7 | only_H3vsUI | DownUp | notDM | Loss | 4 |
| **22_29136680_29137716_11200** | 22_29136680_29137716 | CHEK2 | 11200 | 1.92427928606188 | -0.40313657 | 1 | only_H3vsUI | DownUp | notDM | Loss | 8 |
| **22_29138587_29139672_11200** | 22_29138587_29139672 | CHEK2 | 11200 | 2.87909587950007 | -0.2938382 | 2 | only_H3vsUI | DownUp | notDM | Loss | 8 |
| **22_29179244_29180891_11200** | 22_29179244_29180891 | CHEK2 | 11200 | 4.61714717916289 | -0.61623382 | 3 | only_H3vsUI | DownUp | notDM | Loss | 7 |
| **22_29193719_29195852_11200** | 22_29193719_29195852 | CHEK2 | 11200 | 4.74780009086437 | -0.62403119 | 1 | only_H3vsUI | DownUp | notDM | Gain | 2 |
| **22_29372194_29375993_11200** | 22_29372194_29375993 | CHEK2 | 11200 | 5.36995204893401 | -0.79243381 | 3 | only_H3vsUI | DownUp | notDM | Gain | 5 |
| **22_29661973_29663660_10634** | 22_29661973_29663660 | GAS2L1 | 10634 | 4.59530879544407 | 0.41246864 | 1 | only_DIvsD7 | DownUp | notDM | Loss | 6 |
| **22_29664698_29668534_10634** | 22_29664698_29668534 | GAS2L1 | 10634 | 4.53793237451101 | 0.41246864 | 1 | only_DIvsD7 | DownUp | notDM | Loss | 6 |
| **22_29670754_29672476_10634** | 22_29670754_29672476 | GAS2L1 | 10634 | 4.48533805630935 | -0.34390853 | 8 | only_DIvsD7 | DownUp | notDM | Gain | 2 |
| **22_30013377_30015146_10634** | 22_30013377_30015146 | GAS2L1 | 10634 | 5.49182516087736 | 0.76263135 | 7 | only_DIvsD7 | DownUp | notDM | Loss | 5 |
| **22_30781886_30783038_200312** | 22_30781886_30783038 | RNF215 | 200312 | 1.88649072517248 | 0.17147648 | 2 | only_H3vsUI | UpDown | notDM | Loss | 3 |
| **22_30792478_30792937_200312** | 22_30792478_30792937 | RNF215 | 200312 | 3.95918454273119 | 0.40141283 | 2 | only_H3vsUI | UpDown | notDM | Loss | 3 |
| **22_30793104_30794946_200312** | 22_30793104_30794946 | RNF215 | 200312 | 3.99965242536608 | 0.25386269 | 1 | only_H3vsUI | UpDown | DM | Loss | 10 |
| **22_30793104_30794946_23541** | 22_30793104_30794946 | SEC14L2 | 23541 | 1.81954393554187 | 0.64734681 | 1 | notSpecific | DownUp | DM | Loss | 10 |
| **22_30793104_30794946_51537** | 22_30793104_30794946 | MTFP1 | 51537 | 4.42844258027999 | -0.60855866 | 1 | notSpecific | Down | DM | Loss | 10 |
| **22_30815959_30819907_200312** | 22_30815959_30819907 | RNF215 | 200312 | 0 | 0.07950466 | 5 | only_H3vsUI | UpDown | notDM | Loss | 4 |
| **22_30821972_30822544_200312** | 22_30821972_30822544 | RNF215 | 200312 | 3.62469453127208 | 0.07907932 | 5 | only_H3vsUI | UpDown | notDM | Loss | 3 |
| **22_31226394_31227007_6525** | 22_31226394_31227007 | SMTN | 6525 | 5.36753854725421 | -0.83291357 | 7 | only_H3vsUI | UpDown | notDM | Gain | 2 |
| **22_31227161_31229738_6525** | 22_31227161_31229738 | SMTN | 6525 | 5.36239575799458 | -0.83526798 | 7 | only_H3vsUI | UpDown | notDM | Gain | 1 |

| | | | | | | | | | | | |
|---|---|---|---|---|---|---|---|---|---|---|---|
| **22_31450917_31451691_6525** | 22_31450917_31451691 | SMTN | 6525 | 4.40851131457522 | 0.57863561 | 7 | only_H3vsUI | UpDown | notDM | Gain | 1 |
| **22_31451833_31453228_6525** | 22_31451833_31453228 | SMTN | 6525 | 4.38163844672616 | 0.60116733 | 7 | only_H3vsUI | UpDown | notDM | Gain | 1 |
| **22_31477364_31480656_6525** | 22_31477364_31480656 | SMTN | 6525 | 1.77815125038364 | 0.13102719 | 1 | only_H3vsUI | UpDown | notDM | Loss | 6 |
| **22_31502431_31503399_6525** | 22_31502431_31503399 | SMTN | 6525 | 4.40015792339045 | 0.88453703 | 1 | only_H3vsUI | UpDown | notDM | Loss | 3 |
| **22_31503871_31504483_6525** | 22_31503871_31504483 | SMTN | 6525 | 4.42435886259033 | 0.33947448 | 2 | only_H3vsUI | UpDown | notDM | Loss | 3 |
| **22_31556404_31557763_6525** | 22_31556404_31557763 | SMTN | 6525 | 4.89818197391154 | 0.66576567 | 1 | only_H3vsUI | UpDown | notDM | Loss | 6 |
| **22_31639147_31641210_6525** | 22_31639147_31641210 | SMTN | 6525 | 5.20909392012959 | 0.79312506 | 7 | only_H3vsUI | UpDown | notDM | Gain | 1 |
| **22_36236743_36237796_23543** | 22_36236743_36237796 | RBFOX2 | 23543 | 2.45484486000851 | 0.93118762 | 6 | notSpecific | UpDown | DM | Gain | 9 |
| **22_37913701_37915337_23464** | 22_37913701_37915337 | GCAT | 23464 | 5.46026020929046 | -0.79162534 | 1 | only_H3vsUI | Up | notDM | Loss | 7 |
| **22_38140272_38141848_23464** | 22_38140272_38141848 | GCAT | 23464 | 4.79284675972606 | -0.65357465 | 1 | only_H3vsUI | Up | notDM | Gain | 2 |
| **22_38198031_38199894_23464** | 22_38198031_38199894 | GCAT | 23464 | 3.61573968861916 | 0.56720224 | 2 | only_H3vsUI | Up | notDM | Loss | 6 |
| **22_38201333_38203766_23464** | 22_38201333_38203766 | GCAT | 23464 | 2.40823996531185 | 0.70064251 | 2 | only_H3vsUI | Up | notDM | Loss | 6 |
| **22_38245647_38247232_23464** | 22_38245647_38247232 | GCAT | 23464 | 4.62052122934205 | -0.69060756 | 1 | only_H3vsUI | Up | notDM | Gain | 2 |
| **22_38302828_38304011_23464** | 22_38302828_38304011 | GCAT | 23464 | 4.99522702924928 | -0.8120531 | 2 | only_H3vsUI | Up | notDM | Gain | 2 |
| **22_38613629_38614738_23464** | 22_38613629_38614738 | GCAT | 23464 | 5.612486104065 | -0.85037481 | 7 | only_H3vsUI | Up | notDM | Loss | 7 |
| **22_38667509_38668752_23464** | 22_38667509_38668752 | GCAT | 23464 | 5.66608627952098 | -0.84877496 | 2 | only_H3vsUI | Up | notDM | Loss | 7 |
| **22_38669185_38669684_23464** | 22_38669185_38669684 | GCAT | 23464 | 5.66760796368321 | -0.86322142 | 4 | only_H3vsUI | Up | notDM | Loss | 7 |
| **22_38788034_38788912_25777** | 22_38788034_38788912 | SUN2 | 25777 | 5.60340097458495 | -0.82484168 | 8 | only_H3vsUI | Up | notDM | Gain | 2 |
| **22_39150007_39151511_25777** | 22_39150007_39151511 | SUN2 | 25777 | 2.69460519893357 | 0.3733425 | 1 | only_H3vsUI | Up | notDM | Loss | 3 |
| **22_39152303_39153732_25777** | 22_39152303_39153732 | SUN2 | 25777 | 2.47712125471966 | 0.3733425 | 1 | only_H3vsUI | Up | notDM | Gain | 4 |
| **22_39188700_39189867_25777** | 22_39188700_39189867 | SUN2 | 25777 | 2.45024910831936 | 0.41106873 | 2 | only_H3vsUI | Up | notDM | Loss | 3 |
| **22_39190369_39190863_25777** | 22_39190369_39190863 | SUN2 | 25777 | 2.34830486304816 | 0.32768252 | 4 | only_H3vsUI | Up | notDM | Gain | 2 |
| **22_39319059_39322117_25777** | 22_39319059_39322117 | SUN2 | 25777 | 5.11029671520898 | 0.83023047 | 3 | only_H3vsUI | Up | notDM | Gain | 5 |
| **22_39638759_39639950_25777** | 22_39638759_39639950 | SUN2 | 25777 | 5.65187185450433 | 0.95413885 | 2 | only_H3vsUI | Up | notDM | Loss | 3 |
| **22_41040623_41042345_57591** | 22_41040623_41042345 | MRTFA | 57591 | 3.89867034296553 | -0.23214277 | 7 | notSpecific | Down | DM | Gain | 9 |
| **22_41592706_41593384_83746** | 22_41592706_41593384 | L3MBTL2 | 83746 | 3.89888965592659 | 0.26675662 | 2 | only_DIvsD7 | Down | notDM | Gain | 2 |
| **22_41593669_41594985_83746** | 22_41593669_41594985 | L3MBTL2 | 83746 | 3.80085449150356 | 0.11736785 | 4 | only_DIvsD7 | Down | notDM | Loss | 7 |
| **22_41601459_41602574_83746** | 22_41601459_41602574 | L3MBTL2 | 83746 | 2.19033169817029 | -0.41935302 | 2 | only_DIvsD7 | Down | notDM | Loss | 8 |
| **22_41808493_41809529_83746** | 22_41808493_41809529 | L3MBTL2 | 83746 | 5.31636669428693 | 0.88076385 | 5 | only_DIvsD7 | Down | notDM | Loss | 7 |
| **22_41810003_41811740_83746** | 22_41810003_41811740 | L3MBTL2 | 83746 | 5.31952036810943 | 0.81751118 | 1 | only_DIvsD7 | Down | notDM | Loss | 7 |
| **22_41838951_41840396_84844** | 22_41838951_41840396 | PHF5A | 84844 | 4.38621350371676 | -0.40707744 | 1 | only_DIvsD7 | DownUp | notDM | Loss | 7 |
| **22_41840595_41842381_84844** | 22_41840595_41842381 | PHF5A | 84844 | 4.34925809551279 | -0.40707744 | 2 | only_DIvsD7 | DownUp | notDM | Loss | 7 |
| **22_41843117_41844284_84844** | 22_41843117_41844284 | PHF5A | 84844 | 4.31060835645852 | -0.40707744 | 1 | only_DIvsD7 | DownUp | notDM | Loss | 7 |
| **22_41862854_41864343_84844** | 22_41862854_41864343 | PHF5A | 84844 | 2.58771096501891 | -0.35046309 | 1 | only_DIvsD7 | DownUp | notDM | Loss | 7 |

| | | | | | | | | | | | |
|---|---|---|---|---|---|---|---|---|---|---|---|
| **22_41865417_41867056_84844** | 22_41865417_41867056 | PHF5A | 84844 | 2.83884909073726 | -0.35046309 | 1 | only_DIvsD7 | DownUp | notDM | Gain | 2 |
| **22_41908366_41909732_84844** | 22_41908366_41909732 | PHF5A | 84844 | 4.63987479004972 | 0.64131082 | 7 | only_DIvsD7 | DownUp | notDM | Loss | 5 |
| **22_42976242_42977734_91695** | 22_42976242_42977734 | RRP7BP | 91695 | 2.43136376415899 | -0.36153849 | 1 | only_H3vsUI | DownUp | notDM | Loss | 8 |
| **22_43008157_43010394_91695** | 22_43008157_43010394 | RRP7BP | 91695 | 4.47917206550173 | -0.44696635 | 1 | only_H3vsUI | DownUp | notDM | Gain | 2 |
| **22_43083636_43085266_53947** | 22_43083636_43085266 | A4GALT | 53947 | 3.67135800344349 | -0.01289725 | 3 | only_DIvsD7 | UpDown | notDM | Loss | 3 |
| **22_43083636_43085266_91695** | 22_43083636_43085266 | RRP7BP | 91695 | 5.02363924167391 | 0.66054152 | 3 | only_H3vsUI | DownUp | notDM | Loss | 1 |
| **22_43115148_43116493_53947** | 22_43115148_43116493 | A4GALT | 53947 | 2.90955602924118 | 0.05289503 | 2 | only_DIvsD7 | UpDown | notDM | Gain | 4 |
| **22_50311044_50311817_79087** | 22_50311044_50311817 | ALG12 | 79087 | 2.46239799789896 | -0.4106198 | 1 | only_H3vsUI | Down | notDM | Loss | 8 |
| **22_50363499_50364475_79087** | 22_50363499_50364475 | ALG12 | 79087 | 4.71092087039542 | -0.57300361 | 7 | only_H3vsUI | Down | notDM | Loss | 2 |
| **22_50730794_50731968_410** | 22_50730794_50731968 | ARSA | 410 | 5.52454670334956 | 0.82531139 | 5 | only_DIvsD7 | UpDown | notDM | Loss | 5 |
| **22_50732143_50733474_410** | 22_50732143_50733474 | ARSA | 410 | 5.5225876704565 | 0.82551459 | 7 | only_DIvsD7 | UpDown | notDM | Loss | 5 |
| **22_50740145_50743149_410** | 22_50740145_50743149 | ARSA | 410 | 5.50980984085687 | 0.86281428 | 7 | only_DIvsD7 | UpDown | notDM | Loss | 5 |
| **3_10066372_10067998_132014** | 3_10066372_10067998 | IL17RE | 132014 | 5.08586830149103 | -0.6398374 | 2 | only_DIvsD7 | DownUp | notDM | Loss | 7 |
| **3_10068143_10069529_132014** | 3_10068143_10069529 | IL17RE | 132014 | 5.09213440768669 | -0.6398374 | 1 | only_DIvsD7 | DownUp | notDM | Loss | 7 |
| **3_105106441_105108441_214** | 3_105106441_105108441 | ALCAM | 214 | 4.31576049066573 | 0.31816633 | 7 | notSpecific | Down | DM | Gain | 9 |
| **3_105235551_105238452_214** | 3_105235551_105238452 | ALCAM | 214 | 3.52943035436699 | 0 | 6 | notSpecific | Down | DM | Gain | 9 |
| **3_112738727_112739603_25871** | 3_112738727_112739603 | NEPRO | 25871 | 2.37106786227174 | -0.19217933 | 4 | notSpecific | Down | DM | Gain | 9 |
| **3_113413853_113414991_205717** | 3_113413853_113414991 | USF3 | 205717 | 2.70156798505593 | -0.40711078 | 1 | only_DIvsD7 | UpDown | notDM | Loss | 7 |
| **3_113415873_113418455_205717** | 3_113415873_113418455 | USF3 | 205717 | 2.58206336291171 | -0.40711078 | 1 | only_DIvsD7 | UpDown | notDM | Loss | 8 |
| **3_114326505_114328119_26137** | 3_114326505_114328119 | ZBTB20 | 26137 | 5.69110006643884 | 0.94692717 | 6 | only_H3vsUI | DownUp | notDM | Loss | 5 |
| **3_114340470_114345543_26137** | 3_114340470_114345543 | ZBTB20 | 26137 | 0 | 0.7100394 | 7 | only_H3vsUI | DownUp | notDM | Loss | 4 |
| **3_114367705_114370913_26137** | 3_114367705_114370913 | ZBTB20 | 26137 | 4.37867038520798 | 0.50939891 | 6 | only_H3vsUI | DownUp | notDM | Gain | 5 |
| **3_114424776_114427860_26137** | 3_114424776_114427860 | ZBTB20 | 26137 | 3.69766516264767 | 0.00203977 | 6 | only_H3vsUI | DownUp | notDM | Gain | 4 |
| **3_114432818_114434801_26137** | 3_114432818_114434801 | ZBTB20 | 26137 | 0 | 0.00301381 | 6 | only_H3vsUI | DownUp | notDM | Gain | 4 |
| **3_114436207_114441762_26137** | 3_114436207_114441762 | ZBTB20 | 26137 | 3.52698506856 | 0.00411388 | 6 | only_H3vsUI | DownUp | notDM | Gain | 4 |
| **3_114475522_114478875_26137** | 3_114475522_114478875 | ZBTB20 | 26137 | 0 | 0.00236665 | 1 | only_H3vsUI | DownUp | notDM | Gain | 4 |
| **3_114617493_114622239_26137** | 3_114617493_114622239 | ZBTB20 | 26137 | 0 | -0.00376575 | 5 | only_H3vsUI | DownUp | notDM | Gain | 4 |
| **3_114817910_114820425_26137** | 3_114817910_114820425 | ZBTB20 | 26137 | 0 | 0.41890606 | 4 | only_H3vsUI | DownUp | notDM | Loss | 3 |
| **3_114863915_114865654_2596** | 3_114863915_114865654 | GAP43 | 2596 | 5.67807931033222 | -0.86517891 | 2 | only_DIvsD7 | DownUp | notDM | Loss | 7 |
| **3_114863915_114865654_26137** | 3_114863915_114865654 | ZBTB20 | 26137 | 2.66745295288995 | 0.44857663 | 2 | only_H3vsUI | DownUp | notDM | Loss | 6 |
| **3_114867045_114868282_2596** | 3_114867045_114868282 | GAP43 | 2596 | 5.67567754433848 | -0.8669449 | 1 | only_DIvsD7 | DownUp | notDM | Loss | 7 |
| **3_114867045_114868282_26137** | 3_114867045_114868282 | ZBTB20 | 26137 | 2.96801571399364 | 0.44773487 | 1 | only_H3vsUI | DownUp | notDM | Loss | 6 |
| **3_114936515_114937643_26137** | 3_114936515_114937643 | ZBTB20 | 26137 | 4.84756649014259 | 0.8041084 | 3 | only_H3vsUI | DownUp | notDM | Gain | 1 |
| **3_115230776_115232493_2596** | 3_115230776_115232493 | GAP43 | 2596 | 5.04012348211277 | 0.7215002 | 2 | only_DIvsD7 | DownUp | notDM | Loss | 6 |

| | | | | | | | | | | | |
|---|---|---|---|---|---|---|---|---|---|---|---|
| **3_115342213_115343628_2596** | 3_115342213_115343628 | GAP43 | 2596 | 1.64345267648619 | 0.99860492 | 4 | only_DIvsD7 | DownUp | notDM | Loss | 3 |
| **3_118903992_118907186_8702** | 3_118903992_118907186 | B4GALT4 | 8702 | 4.71850999269768 | 0.57668856 | 7 | only_DIvsD7 | UpDown | notDM | Gain | 5 |
| **3_118912521_118915417_8702** | 3_118912521_118915417 | B4GALT4 | 8702 | 4.64649194864631 | -0.55379329 | 7 | only_DIvsD7 | UpDown | notDM | Gain | 5 |
| **3_118922428_118928643_8702** | 3_118922428_118928643 | B4GALT4 | 8702 | 4.49252292842261 | 0.7073445 | 5 | only_DIvsD7 | UpDown | notDM | Gain | 5 |
| **3_118930333_118932525_8702** | 3_118930333_118932525 | B4GALT4 | 8702 | 4.43504764133996 | -0.4551366 | 7 | only_DIvsD7 | UpDown | notDM | Gain | 5 |
| **3_118945974_118947759_8702** | 3_118945974_118947759 | B4GALT4 | 8702 | 4.08044609461105 | 0.92085367 | 8 | only_DIvsD7 | UpDown | notDM | Loss | 5 |
| **3_118953685_118959459_8702** | 3_118953685_118959459 | B4GALT4 | 8702 | 1.44715803134222 | 0.54854608 | 1 | only_DIvsD7 | UpDown | notDM | Gain | 4 |
| **3_118959845_118960451_8702** | 3_118959845_118960451 | B4GALT4 | 8702 | 1.73239375982297 | -0.45290398 | 7 | only_DIvsD7 | UpDown | notDM | Loss | 7 |
| **3_119040495_119042965_8702** | 3_119040495_119042965 | B4GALT4 | 8702 | 4.90689506055697 | -0.70359483 | 1 | only_DIvsD7 | UpDown | notDM | Loss | 7 |
| **3_119297659_119299677_141** | 3_119297659_119299677 | ADPRH | 141 | 0 | 0.07385637 | 2 | only_DIvsD7 | UpDown | notDM | Gain | 4 |
| **3_120459651_120461006_285282** | 3_120459651_120461006 | RABL3 | 285282 | 2.55870857053317 | 0.08226889 | 1 | only_H3vsUI | DownUp | notDM | Loss | 8 |
| **3_120461694_120463886_285282** | 3_120461694_120463886 | RABL3 | 285282 | 2.4996870826184 | 0.08220125 | 1 | only_H3vsUI | DownUp | notDM | Loss | 8 |
| **3_120472192_120473095_285282** | 3_120472192_120473095 | RABL3 | 285282 | 4.03450813677917 | 0.22426619 | 8 | only_H3vsUI | DownUp | notDM | Gain | 1 |
| **3_12196709_12198451_5468** | 3_12196709_12198451 | PPARG | 5468 | 5.12096856501938 | 0.67556415 | 7 | only_H3vsUI | UpDown | notDM | Loss | 1 |
| **3_12198747_12201381_5468** | 3_12198747_12201381 | PPARG | 5468 | 5.11122889823416 | 0.73909658 | 1 | only_H3vsUI | UpDown | notDM | Loss | 6 |
| **3_12262274_12265140_5468** | 3_12262274_12265140 | PPARG | 5468 | 4.8157835577913 | 0.54022203 | 6 | only_H3vsUI | UpDown | notDM | Gain | 5 |
| **3_12308077_12309908_5468** | 3_12308077_12309908 | PPARG | 5468 | 4.28987868279042 | 0.50285573 | 6 | only_H3vsUI | UpDown | notDM | Gain | 1 |
| **3_12328165_12329123_5468** | 3_12328165_12329123 | PPARG | 5468 | 2.44404479591808 | 0.95367548 | 5 | only_H3vsUI | UpDown | notDM | Loss | 3 |
| **3_12329903_12333543_5468** | 3_12329903_12333543 | PPARG | 5468 | 2.70329137811866 | 0.9616465 | 1 | only_H3vsUI | UpDown | notDM | Gain | 4 |
| **3_12333921_12336493_5468** | 3_12333921_12336493 | PPARG | 5468 | 3.65934563574618 | -0.17924868 | 8 | only_H3vsUI | UpDown | notDM | Loss | 5 |
| **3_12340344_12343903_5468** | 3_12340344_12343903 | PPARG | 5468 | 3.99016119289848 | 0.20840505 | 7 | only_H3vsUI | UpDown | notDM | Gain | 5 |
| **3_12345362_12348327_5468** | 3_12345362_12348327 | PPARG | 5468 | 4.20314171911199 | 0.83012476 | 7 | only_H3vsUI | UpDown | notDM | Gain | 5 |
| **3_12348682_12351953_5468** | 3_12348682_12351953 | PPARG | 5468 | 4.28519712281097 | 0.63458415 | 7 | only_H3vsUI | UpDown | notDM | Gain | 5 |
| **3_12352265_12355126_5468** | 3_12352265_12355126 | PPARG | 5468 | 4.3592091917643 | 0.83887355 | 7 | only_H3vsUI | UpDown | notDM | Gain | 5 |
| **3_12356456_12360048_5468** | 3_12356456_12360048 | PPARG | 5468 | 4.43229569244401 | 0.82584018 | 5 | only_H3vsUI | UpDown | notDM | Gain | 5 |
| **3_12386010_12387384_5468** | 3_12386010_12387384 | PPARG | 5468 | 3.74725639744214 | 0.11437425 | 6 | only_H3vsUI | UpDown | notDM | Gain | 1 |
| **3_124231492_124233287_7372** | 3_124231492_124233287 | UMPS | 7372 | 5.33447988845822 | 0.71905395 | 4 | only_H3vsUI | Down | notDM | Gain | 1 |
| **3_124448323_124449234_7372** | 3_124448323_124449234 | UMPS | 7372 | 1.81954393554187 | -0.43656912 | 2 | only_H3vsUI | Down | notDM | Gain | 4 |
| **3_124449432_124452030_7372** | 3_124449432_124452030 | UMPS | 7372 | 2.13033376849501 | -0.47220498 | 1 | only_H3vsUI | Down | notDM | Loss | 8 |
| **3_12445736_12448649_5468** | 3_12445736_12448649 | PPARG | 5468 | 5.06572159344525 | 0.72356143 | 4 | only_H3vsUI | UpDown | notDM | Gain | 5 |
| **3_12511446_12513560_5468** | 3_12511446_12513560 | PPARG | 5468 | 5.25738574729304 | 0.80986199 | 6 | only_H3vsUI | UpDown | notDM | Gain | 1 |
| **3_12721257_12722101_5468** | 3_12721257_12722101 | PPARG | 5468 | 5.5918311836677 | 0.81617036 | 6 | only_H3vsUI | UpDown | notDM | Gain | 1 |
| **3_128967007_128968302_22820** | 3_128967007_128968302 | COPG1 | 22820 | 2.2405492482826 | -0.47746883 | 1 | only_H3vsUI | DownUp | notDM | Loss | 8 |
| **3_128968569_128969754_22820** | 3_128968569_128969754 | COPG1 | 22820 | 1.98227123303957 | -0.47746883 | 2 | only_H3vsUI | DownUp | notDM | Loss | 8 |

| | | | | | | | | | | | |
|---|---|---|---|---|---|---|---|---|---|---|---|
| **3_129115310_129117939_22820** | 3_129115310_129117939 | COPG1 | 22820 | 5.16683550293114 | -0.68340172 | 1 | only_H3vsUI | DownUp | notDM | Loss | 7 |
| **3_129131316_129134342_22820** | 3_129131316_129134342 | COPG1 | 22820 | 5.21183842985717 | -0.728995 | 7 | only_H3vsUI | DownUp | notDM | Loss | 7 |
| **3_129146281_129147493_90288** | 3_129146281_129147493 | EFCAB12 | 90288 | 0 | 0.04616403 | 2 | only_H3vsUI | Up | notDM | Gain | 4 |
| **3_129360686_129361935_22820** | 3_129360686_129361935 | COPG1 | 22820 | 5.59355077302051 | -0.83903518 | 7 | only_H3vsUI | DownUp | notDM | Gain | 2 |
| **3_131068036_131070449_131870** | 3_131068036_131070449 | NUDT16 | 131870 | 4.47973362534305 | 0.7471934 | 7 | only_H3vsUI | Down | notDM | Gain | 1 |
| **3_131071064_131073495_131870** | 3_131071064_131073495 | NUDT16 | 131870 | 4.4335298259155 | 0.85841485 | 6 | only_H3vsUI | Down | notDM | Gain | 1 |
| **3_131100791_131101891_131870** | 3_131100791_131101891 | NUDT16 | 131870 | 2.2148438480477 | -0.53680595 | 1 | only_H3vsUI | Down | notDM | Loss | 8 |
| **3_132377059_132379159_79876** | 3_132377059_132379159 | UBA5 | 79876 | 1.68124123737559 | 0.49790417 | 1 | only_H3vsUI | Down | notDM | Gain | 4 |
| **3_132441915_132442754_79876** | 3_132441915_132442754 | UBA5 | 79876 | 4.79734372616236 | -0.52641063 | 1 | only_H3vsUI | Down | notDM | Loss | 7 |
| **3_13577741_13580586_2199** | 3_13577741_13580586 | FBLN2 | 2199 | 4.00199317382353 | -0.09666051 | 3 | only_DIvsD7 | DownUp | notDM | Loss | 5 |
| **3_148801873_148804300_10087394** | 3_148801873_148804300 | HLTF-AS1 | 100873945 | 0 | -0.00306271 | 1 | only_H3vsUI | UpDown | notDM | Loss | 3 |
| **3_151593006_151594478_56670** | 3_151593006_151594478 | SUCNR1 | 56670 | 3.1978316933289 | 0.01714431 | 4 | notSpecific | Up | DM | Gain | 9 |
| **3_152685422_152689115_5912** | 3_152685422_152689115 | RAP2B | 5912 | 5.28084005181591 | 0.76972305 | 7 | only_H3vsUI | Up | notDM | Gain | 5 |
| **3_152760242_152762037_5912** | 3_152760242_152762037 | RAP2B | 5912 | 5.07185624330995 | 0.71257631 | 3 | only_H3vsUI | Up | notDM | Gain | 1 |
| **3_152861548_152863633_5912** | 3_152861548_152863633 | RAP2B | 5912 | 4.21476439666804 | 0.78351283 | 6 | only_H3vsUI | Up | notDM | Gain | 5 |
| **3_152875737_152879007_5912** | 3_152875737_152879007 | RAP2B | 5912 | 3.00987563371216 | 0.45776767 | 1 | only_H3vsUI | Up | notDM | Gain | 4 |
| **3_152880802_152885079_5912** | 3_152880802_152885079 | RAP2B | 5912 | 2.88930170250631 | 0.48411422 | 1 | only_H3vsUI | Up | notDM | Gain | 4 |
| **3_153837793_153839055_26084** | 3_153837793_153839055 | ARHGEF26 | 26084 | 1.97772360528885 | 0.05059917 | 2 | only_DIvsD7 | Down | notDM | Gain | 4 |
| **3_153839317_153844921_26084** | 3_153839317_153844921 | ARHGEF26 | 26084 | 1.95424250943932 | 0.02161783 | 2 | only_DIvsD7 | Down | notDM | Loss | 8 |
| **3_154145698_154147013_344758** | 3_154145698_154147013 | GPR149 | 344758 | 2.69196510276736 | 0 | 4 | only_H3vsUI | UpDown | notDM | Loss | 3 |
| **3_157052455_157055277_152078** | 3_157052455_157055277 | SLC66A1L | 152078 | 5.31356563941553 | 0.93300575 | 3 | only_H3vsUI | DownUp | notDM | Gain | 1 |
| **3_157087009_157089042_152078** | 3_157087009_157089042 | SLC66A1L | 152078 | 5.2357606818522 | 0.85236024 | 6 | only_H3vsUI | DownUp | notDM | Loss | 5 |
| **3_157154247_157156721_152078** | 3_157154247_157156721 | SLC66A1L | 152078 | 5.01875457411434 | 0.90393642 | 2 | only_H3vsUI | DownUp | notDM | Loss | 6 |
| **3_157216549_157218863_152078** | 3_157216549_157218863 | SLC66A1L | 152078 | 4.62604252200325 | 0.66554068 | 4 | only_H3vsUI | DownUp | notDM | Gain | 1 |
| **3_157260523_157261607_152078** | 3_157260523_157261607 | SLC66A1L | 152078 | 0 | 0.84585891 | 2 | only_H3vsUI | DownUp | notDM | Gain | 4 |
| **3_16926726_16928822_23228** | 3_16926726_16928822 | PLCL2 | 23228 | 2.44090908206522 | 0.2340811 | 1 | only_DIvsD7 | UpDown | notDM | Loss | 6 |
| **3_16946533_16949452_23180** | 3_16946533_16949452 | RFTN1 | 23180 | 5.59253426448648 | 0.83095755 | 6 | notSpecific | Down | DM | Gain | 9 |
| **3_16946533_16949452_23228** | 3_16946533_16949452 | PLCL2 | 23228 | 4.30282858826026 | 0.23245711 | 6 | only_DIvsD7 | UpDown | DM | Gain | 9 |
| **3_169755962_169757014_80012** | 3_169755962_169757014 | PHC3 | 80012 | 5.15388800251949 | 0.83178376 | 2 | only_H3vsUI | Down | notDM | Loss | 7 |
| **3_169896494_169899294_80012** | 3_169896494_169899294 | PHC3 | 80012 | 2.34830486304816 | 0.11040456 | 1 | only_H3vsUI | Down | notDM | Loss | 8 |
| **3_169899632_169900315_80012** | 3_169899632_169900315 | PHC3 | 80012 | 1.98677173426624 | 0.28126637 | 1 | only_H3vsUI | Down | notDM | Gain | 4 |
| **3_169950495_169951784_80012** | 3_169950495_169951784 | PHC3 | 80012 | 4.70722941932729 | 0.51781717 | 8 | only_H3vsUI | Down | notDM | Gain | 1 |
| **3_170206270_170207613_80012** | 3_170206270_170207613 | PHC3 | 80012 | 5.48679306607356 | 0.83967499 | 6 | only_H3vsUI | Down | notDM | Gain | 1 |
| **3_17055614_17057185_23228** | 3_17055614_17057185 | PLCL2 | 23228 | 3.55990662503611 | 0.0169875 | 6 | only_DIvsD7 | UpDown | notDM | Gain | 1 |

| | | | | | | | | | | | |
|---|---|---|---|---|---|---|---|---|---|---|---|
| **3_183164434_183165044_10050568** | 3_183164434_183165044 | LINC00888 | 100505687 | 2.59769518592551 | 0.7721289 | 2 | only_H3vsUI | DownUp | notDM | Gain | 1 |
| **3_183165712_183166904_10050568** | 3_183165712_183166904 | LINC00888 | 100505687 | 2.43933269383026 | 0.77527616 | 1 | only_H3vsUI | DownUp | notDM | Gain | 4 |
| **3_183978004_183979076_94032** | 3_183978004_183979076 | CAMK2N2 | 94032 | 2.24551266781415 | 0.16826917 | 4 | only_DIvsD7 | DownUp | notDM | Gain | 4 |
| **3_183979576_183980428_94032** | 3_183979576_183980428 | CAMK2N2 | 94032 | 2.51454775266029 | 0.16886133 | 2 | only_DIvsD7 | DownUp | notDM | Loss | 3 |
| **3_184078391_184079309_1181** | 3_184078391_184079309 | CLCN2 | 1181 | 2.11727129565576 | -0.0369949 | 1 | only_DIvsD7 | UpDown | notDM | Loss | 3 |
| **3_184081276_184082768_1181** | 3_184081276_184082768 | CLCN2 | 1181 | 3.27600198996205 | 0.21949616 | 1 | only_DIvsD7 | UpDown | notDM | Gain | 4 |
| **3_184530450_184531440_23355** | 3_184530450_184531440 | VPS8 | 23355 | 2.71683772329952 | -0.69259553 | 2 | notSpecific | DownUp | DM | Loss | 10 |
| **3_184972001_184972586_1962** | 3_184972001_184972586 | EHHADH | 1962 | 2.06818586174616 | 0.2220346 | 2 | only_H3vsUI | DownUp | notDM | Gain | 2 |
| **3_184999575_185000615_1962** | 3_184999575_185000615 | EHHADH | 1962 | 0 | 0.00176296 | 2 | only_H3vsUI | DownUp | notDM | Gain | 4 |
| **3_185000885_185002319_1962** | 3_185000885_185002319 | EHHADH | 1962 | 3.04493154614916 | 0.00176269 | 1 | only_H3vsUI | DownUp | notDM | Loss | 8 |
| **3_185079741_185080715_1962** | 3_185079741_185080715 | EHHADH | 1962 | 5.03304559384366 | -0.84903286 | 4 | only_H3vsUI | DownUp | notDM | Gain | 2 |
| **3_188664297_188665459_10087404** | 3_188664297_188665459 | TPRG1-AS1 | 100874043 | 0 | 0.04821061 | 5 | only_H3vsUI | UpDown | notDM | Loss | 3 |
| **3_188665912_188667485_10087404** | 3_188665912_188667485 | TPRG1-AS1 | 100874043 | 2.68663626926229 | 0.04824822 | 7 | only_H3vsUI | UpDown | notDM | Gain | 1 |
| **3_189835597_189836838_10192915** | 3_189835597_189836838 | P3H2-AS1 | 101929152 | 3.28239550474253 | 0.02152355 | 2 | only_H3vsUI | UpDown | notDM | Loss | 3 |
| **3_189837015_189838166_10192915** | 3_189837015_189838166 | P3H2-AS1 | 101929152 | 2.76937732607614 | 0.04515065 | 2 | only_H3vsUI | UpDown | notDM | Loss | 3 |
| **3_189842909_189843826_10192915** | 3_189842909_189843826 | P3H2-AS1 | 101929152 | 3.61888448499545 | 0.03557034 | 6 | only_H3vsUI | UpDown | notDM | Gain | 1 |
| **3_196358423_196359327_10192976** | 3_196358423_196359327 | LINC01063 | 101929769 | 2.12057393120585 | 0.03469253 | 1 | only_DIvsD7 | UpDown | notDM | Gain | 4 |
| **3_196359739_196360108_10192976** | 3_196359739_196360108 | LINC01063 | 101929769 | 2.45178643552429 | 0.03708526 | 4 | only_DIvsD7 | UpDown | notDM | Loss | 3 |
| **3_196437966_196438953_5062** | 3_196437966_196438953 | PAK2 | 5062 | 4.4436697034964 | 0.48632959 | 1 | only_DIvsD7 | UpDown | notDM | Loss | 7 |
| **3_196439707_196440596_5062** | 3_196439707_196440596 | PAK2 | 5062 | 4.41718926845901 | 0.48688739 | 1 | only_DIvsD7 | UpDown | notDM | Loss | 7 |
| **3_196467390_196469441_5062** | 3_196467390_196469441 | PAK2 | 5062 | 2.82216807936802 | 0.56559854 | 1 | only_DIvsD7 | UpDown | notDM | Gain | 4 |
| **3_196516324_196518290_5062** | 3_196516324_196518290 | PAK2 | 5062 | 4.69546416426286 | 0.74326375 | 8 | only_DIvsD7 | UpDown | notDM | Gain | 1 |
| **3_196518694_196521301_5062** | 3_196518694_196521301 | PAK2 | 5062 | 4.71573600322506 | 0.83488769 | 6 | only_DIvsD7 | UpDown | notDM | Loss | 5 |
| **3_196667298_196668801_5062** | 3_196667298_196668801 | PAK2 | 5062 | 5.30227030508398 | 0.87035238 | 2 | only_DIvsD7 | UpDown | notDM | Loss | 6 |
| **3_196669917_196671173_5062** | 3_196669917_196671173 | PAK2 | 5062 | 5.30790446770194 | 0.87035238 | 1 | only_DIvsD7 | UpDown | notDM | Loss | 6 |
| **3_197022546_197024498_10050708** | 3_197022546_197024498 | DLG1-AS1 | 100507086 | 2.79657433321043 | 0.01014373 | 1 | only_H3vsUI | UpDown | notDM | Loss | 3 |
| **3_197025506_197026934_10050708** | 3_197025506_197026934 | DLG1-AS1 | 100507086 | 2.5854607295085 | 0.0081619 | 2 | only_H3vsUI | UpDown | notDM | Loss | 3 |
| **3_23495965_23497540_7324** | 3_23495965_23497540 | UBE2E1 | 7324 | 5.54388808507078 | 0.77495063 | 6 | only_DIvsD7 | DownUp | notDM | Gain | 1 |
| **3_23743140_23744495_7324** | 3_23743140_23744495 | UBE2E1 | 7324 | 5.01241537476243 | -0.70258309 | 6 | only_DIvsD7 | DownUp | notDM | Loss | 7 |
| **3_23779740_23785778_7324** | 3_23779740_23785778 | UBE2E1 | 7324 | 4.7897005496258 | -0.52608852 | 3 | only_DIvsD7 | DownUp | notDM | Gain | 5 |
| **3_23788918_23791061_7324** | 3_23788918_23791061 | UBE2E1 | 7324 | 4.75139459247808 | -0.6395509 | 3 | only_DIvsD7 | DownUp | notDM | Gain | 5 |
| **3_23846251_23847271_7324** | 3_23846251_23847271 | UBE2E1 | 7324 | 2.09342168516224 | -0.25167801 | 1 | only_DIvsD7 | DownUp | notDM | Gain | 4 |
| **3_23848231_23852189_7324** | 3_23848231_23852189 | UBE2E1 | 7324 | 2.89042101880091 | -0.69181199 | 1 | only_DIvsD7 | DownUp | notDM | Loss | 8 |
| **3_23852438_23854365_7324** | 3_23852438_23854365 | UBE2E1 | 7324 | 0 | 0.13333991 | 2 | only_DIvsD7 | DownUp | notDM | Gain | 4 |

| | | | | | | | | | | | |
|---|---|---|---|---|---|---|---|---|---|---|---|
| **3_23906422_23908797_7324** | 3_23906422_23908797 | UBE2E1 | 7324 | 0 | 0.07657395 | 8 | only_DIvsD7 | DownUp | notDM | Gain | 4 |
| **3_23987698_23990085_7324** | 3_23987698_23990085 | UBE2E1 | 7324 | 5.14688429008769 | -0.74310872 | 1 | only_DIvsD7 | DownUp | notDM | Loss | 7 |
| **3_27490921_27492690_9497** | 3_27490921_27492690 | SLC4A7 | 9497 | 4.5211119205978 | -0.48604694 | 8 | notSpecific | Down | DM | Gain | 9 |
| **3_33259226_33261276_25827** | 3_33259226_33261276 | FBXL2 | 25827 | 4.76104783925584 | 0.71769121 | 2 | notSpecific | DownUp | DM | Loss | 10 |
| **3_33259226_33261276_26032** | 3_33259226_33261276 | SUSD5 | 26032 | 0 | 0.81493582 | 2 | notSpecific | DownUp | DM | Loss | 10 |
| **3_33259226_33261276_2720** | 3_33259226_33261276 | GLB1 | 2720 | 5.08110956914827 | 0.86172139 | 2 | notSpecific | Down | DM | Loss | 10 |
| **3_37902120_37902948_101928153** | 3_37902120_37902948 | ITGA9-AS1 | 101928153 | 2.4345689040342 | 0.58425262 | 1 | only_DIvsD7 | UpDown | notDM | Gain | 4 |
| **3_37903780_37906556_101928153** | 3_37903780_37906556 | ITGA9-AS1 | 101928153 | 2.73479982958885 | 0.43625165 | 1 | only_DIvsD7 | UpDown | notDM | Gain | 4 |
| **3_37975514_37978939_101928153** | 3_37975514_37978939 | ITGA9-AS1 | 101928153 | 4.85906020136226 | 0.85272399 | 7 | only_DIvsD7 | UpDown | notDM | Gain | 1 |
| **3_43147681_43148258_84892** | 3_43147681_43148258 | POMGNT2 | 84892 | 2.06069784035361 | 0.17411435 | 4 | only_DIvsD7 | UpDown | notDM | Loss | 3 |
| **3_44517502_44518821_285346** | 3_44517502_44518821 | ZNF852 | 285346 | 4.52254855598626 | -0.38831865 | 2 | only_DIvsD7 | Down | notDM | Loss | 7 |
| **3_44550951_44551824_285346** | 3_44550951_44551824 | ZNF852 | 285346 | 2.48429983934679 | -0.44685535 | 2 | only_DIvsD7 | Down | notDM | Loss | 8 |
| **3_44552432_44553303_285346** | 3_44552432_44553303 | ZNF852 | 285346 | 2.48572142648158 | 0.4389571 | 1 | only_DIvsD7 | Down | notDM | Gain | 4 |
| **3_4533251_4534834_3708** | 3_4533251_4534834 | ITPR1 | 3708 | 2.29885307640971 | 0.17491556 | 2 | only_DIvsD7 | UpDown | notDM | Loss | 6 |
| **3_4535184_4536366_3708** | 3_4535184_4536366 | ITPR1 | 3708 | 2.18184358794477 | 0.56867608 | 1 | only_DIvsD7 | UpDown | notDM | Gain | 4 |
| **3_48468665_48470129_5364** | 3_48468665_48470129 | PLXNB1 | 5364 | 3.12450422483428 | 0.59556494 | 4 | only_H3vsUI | DownUp | notDM | Loss | 3 |
| **3_48471238_48472174_5364** | 3_48471238_48472174 | PLXNB1 | 5364 | 0 | 0.24438805 | 2 | only_H3vsUI | DownUp | notDM | Loss | 3 |
| **3_48480279_48481458_5364** | 3_48480279_48481458 | PLXNB1 | 5364 | 3.94551782207784 | -0.54577737 | 1 | only_H3vsUI | DownUp | notDM | Gain | 2 |
| **3_48482079_48483775_5364** | 3_48482079_48483775 | PLXNB1 | 5364 | 4.02616540883925 | -0.55764176 | 2 | only_H3vsUI | DownUp | notDM | Loss | 7 |
| **3_48486970_48487953_5364** | 3_48486970_48487953 | PLXNB1 | 5364 | 4.19066779607065 | -0.36075065 | 1 | only_H3vsUI | DownUp | notDM | Loss | 7 |
| **3_48488442_48489878_5364** | 3_48488442_48489878 | PLXNB1 | 5364 | 4.23003998115794 | -0.36075065 | 2 | only_H3vsUI | DownUp | notDM | Loss | 7 |
| **3_5028959_5029891_55207** | 3_5028959_5029891 | ARL8B | 55207 | 5.12739962959279 | 0.75700357 | 7 | notSpecific | DownUp | DM | Gain | 9 |
| **3_50382631_50383779_51364** | 3_50382631_50383779 | ZMYND10 | 51364 | 0 | 0.00368542 | 4 | only_H3vsUI | UpDown | notDM | Loss | 3 |
| **3_50387011_50388182_51364** | 3_50387011_50388182 | ZMYND10 | 51364 | 3.5854607295085 | -0.01844016 | 1 | only_H3vsUI | UpDown | notDM | Loss | 3 |
| **3_50388461_50389527_51364** | 3_50388461_50389527 | ZMYND10 | 51364 | 3.72427586960079 | -0.03133743 | 1 | only_H3vsUI | UpDown | notDM | Loss | 3 |
| **3_50607045_50608197_51409** | 3_50607045_50608197 | HEMK1 | 51409 | 2.0899051114394 | -0.09117743 | 1 | only_H3vsUI | Down | notDM | Loss | 8 |
| **3_50649551_50650149_51409** | 3_50649551_50650149 | HEMK1 | 51409 | 4.62970514499581 | 0.61319885 | 2 | only_H3vsUI | Down | notDM | Gain | 2 |
| **3_50654679_50655778_51409** | 3_50654679_50655778 | HEMK1 | 51409 | 4.67903703746036 | -0.5025817 | 2 | only_H3vsUI | Down | notDM | Gain | 4 |
| **3_50708440_50709417_51409** | 3_50708440_50709417 | HEMK1 | 51409 | 5.00654305316151 | 0.74921853 | 6 | only_H3vsUI | Down | notDM | Gain | 1 |
| **3_52008191_52009938_6159** | 3_52008191_52009938 | RPL29 | 6159 | 4.2997687119198 | 0.43995694 | 5 | only_H3vsUI | DownUp | notDM | Loss | 6 |
| **3_52015965_52018732_6159** | 3_52015965_52018732 | RPL29 | 6159 | 4.0502637226458 | 0.43828733 | 5 | only_H3vsUI | DownUp | notDM | Loss | 5 |
| **3_52028110_52029533_6159** | 3_52028110_52029533 | RPL29 | 6159 | 2.54032947479087 | 0.53873227 | 1 | only_H3vsUI | DownUp | notDM | Loss | 6 |
| **3_52089259_52090080_211** | 3_52089259_52090080 | ALAS1 | 211 | 5.15258490905204 | 0.77446187 | 2 | only_DIvsD7 | DownUp | notDM | Loss | 7 |
| **3_52090711_52091299_11188** | 3_52090711_52091299 | NISCH | 11188 | 5.60012961151235 | 0.7965189 | 1 | only_H3vsUI | Down | notDM | Loss | 2 |

| | | | | | | | | | | | |
|---|---|---|---|---|---|---|---|---|---|---|---|
| **3_52230828_52232034_211** | 3_52230828_52232034 | ALAS1 | 211 | 1.83884909073726 | 0.34084139 | 4 | only_DIvsD7 | DownUp | notDM | Loss | 7 |
| **3_52232347_52233563_11188** | 3_52232347_52233563 | NISCH | 11188 | 5.40836039747228 | -0.77515005 | 1 | only_H3vsUI | Down | notDM | Loss | 7 |
| **3_52232347_52233563_211** | 3_52232347_52233563 | ALAS1 | 211 | 2.37839790094814 | 0.17363355 | 1 | only_DIvsD7 | DownUp | notDM | Loss | 7 |
| **3_52309919_52312046_11188** | 3_52309919_52312046 | NISCH | 11188 | 5.24941361622943 | -0.79619577 | 1 | only_H3vsUI | Down | notDM | Gain | 2 |
| **3_52312903_52314108_11188** | 3_52312903_52314108 | NISCH | 11188 | 5.24434145595964 | -0.79619577 | 1 | only_H3vsUI | Down | notDM | Gain | 2 |
| **3_52442705_52443791_11188** | 3_52442705_52443791 | NISCH | 11188 | 4.66127303044306 | -0.44489829 | 2 | only_H3vsUI | Down | notDM | Loss | 7 |
| **3_52444737_52446291_11188** | 3_52444737_52446291 | NISCH | 11188 | 4.63691896785955 | -0.55457769 | 1 | only_H3vsUI | Down | notDM | Loss | 7 |
| **3_52483664_52485252_11188** | 3_52483664_52485252 | NISCH | 11188 | 3.64167237322469 | -0.09466348 | 7 | only_H3vsUI | Down | notDM | Loss | 7 |
| **3_52488598_52489438_11188** | 3_52488598_52489438 | NISCH | 11188 | 2.29225607135648 | -0.68974992 | 1 | only_H3vsUI | Down | notDM | Loss | 8 |
| **3_52490060_52491314_11188** | 3_52490060_52491314 | NISCH | 11188 | 2.63245729218472 | -0.68974992 | 2 | only_H3vsUI | Down | notDM | Loss | 8 |
| **3_52569784_52570344_11188** | 3_52569784_52570344 | NISCH | 11188 | 4.90391978194984 | -0.83748362 | 4 | only_H3vsUI | Down | notDM | Loss | 7 |
| **3_52570766_52572651_11188** | 3_52570766_52572651 | NISCH | 11188 | 4.90979129748957 | 0.6601521 | 1 | only_H3vsUI | Down | notDM | Loss | 7 |
| **3_52639940_52642774_211** | 3_52639940_52642774 | ALAS1 | 211 | 5.61048129912281 | -0.79954794 | 3 | only_DIvsD7 | DownUp | notDM | Loss | 7 |
| **3_52803321_52804530_11188** | 3_52803321_52804530 | NISCH | 11188 | 5.49650067421562 | -0.82628333 | 1 | only_H3vsUI | Down | notDM | Loss | 7 |
| **3_53511921_53513583_55540** | 3_53511921_53513583 | IL17RB | 55540 | 5.5647015637125 | 0.83130001 | 7 | only_DIvsD7 | DownUp | notDM | Loss | 5 |
| **3_53880566_53880881_55540** | 3_53880566_53880881 | IL17RB | 55540 | 0 | 0.78166186 | 2 | only_DIvsD7 | DownUp | notDM | Gain | 1 |
| **3_53914524_53915764_55540** | 3_53914524_53915764 | IL17RB | 55540 | 4.53044303990296 | 0.69646532 | 2 | only_DIvsD7 | DownUp | notDM | Loss | 3 |
| **3_57198428_57199209_378** | 3_57198428_57199209 | ARF4 | 378 | 5.5843617596248 | 0.84894318 | 2 | only_DIvsD7 | DownUp | notDM | Loss | 3 |
| **3_57204418_57204697_378** | 3_57204418_57204697 | ARF4 | 378 | 5.57811063049989 | 0.83289769 | 4 | only_DIvsD7 | DownUp | notDM | Gain | 2 |
| **3_57262306_57264159_378** | 3_57262306_57264159 | ARF4 | 378 | 5.50389550010988 | -0.80915603 | 1 | only_DIvsD7 | DownUp | notDM | Loss | 7 |
| **3_57540693_57541768_378** | 3_57540693_57541768 | ARF4 | 378 | 4.61771308986343 | -0.85982801 | 1 | only_DIvsD7 | DownUp | notDM | Gain | 2 |
| **3_57542130_57544656_378** | 3_57542130_57544656 | ARF4 | 378 | 4.58636222330787 | -0.85982801 | 1 | only_DIvsD7 | DownUp | notDM | Loss | 7 |
| **3_57579670_57582273_378** | 3_57579670_57582273 | ARF4 | 378 | 2.98362628712453 | -0.67746043 | 2 | only_DIvsD7 | DownUp | notDM | Loss | 7 |
| **3_57583601_57585367_378** | 3_57583601_57585367 | ARF4 | 378 | 2.56584781867352 | -0.67746043 | 1 | only_DIvsD7 | DownUp | notDM | Gain | 2 |
| **3_57676241_57678307_378** | 3_57676241_57678307 | ARF4 | 378 | 4.96937397202059 | 0.70666524 | 1 | only_DIvsD7 | DownUp | notDM | Loss | 6 |
| **3_57737032_57739834_378** | 3_57737032_57739834 | ARF4 | 378 | 5.18746995617695 | 0.77375471 | 6 | only_DIvsD7 | DownUp | notDM | Gain | 5 |
| **3_57773461_57775722_378** | 3_57773461_57775722 | ARF4 | 378 | 5.27950268398316 | -0.74538304 | 8 | only_DIvsD7 | DownUp | notDM | Loss | 7 |
| **3_57781529_57782740_378** | 3_57781529_57782740 | ARF4 | 378 | 5.29731395376431 | -0.73571684 | 8 | only_DIvsD7 | DownUp | notDM | Gain | 2 |
| **3_57896793_57897882_378** | 3_57896793_57897882 | ARF4 | 378 | 5.49646467649652 | 0.96131707 | 6 | only_DIvsD7 | DownUp | notDM | Gain | 2 |
| **3_57955876_57956863_378** | 3_57955876_57956863 | ARF4 | 378 | 5.57149570743157 | 0.83775894 | 7 | only_DIvsD7 | DownUp | notDM | Gain | 2 |
| **3_58010595_58012814_378** | 3_58010595_58012814 | ARF4 | 378 | 5.63098183186692 | 0.94344232 | 8 | only_DIvsD7 | DownUp | notDM | Loss | 5 |
| **3_58046581_58049615_378** | 3_58046581_58049615 | ARF4 | 378 | 5.66607878417358 | 0.81409178 | 7 | only_DIvsD7 | DownUp | notDM | Gain | 5 |
| **3_58146924_58153215_105377105** | 3_58146924_58153215 | FLNB-AS1 | 105377105 | 0 | 0 | 7 | only_H3vsUI | DownUp | notDM | Loss | 4 |
| **3_72708361_72711201_55164** | 3_72708361_72711201 | SHQ1 | 55164 | 5.2705181290992 | 0.70306944 | 6 | only_DIvsD7 | UpDown | notDM | Loss | 5 |

| | | | | | | | | | | | |
|---|---|---|---|---|---|---|---|---|---|---|---|
| **3_72895365_72898918_55164** | 3_72895365_72898918 | SHQ1 | 55164 | 0 | -0.61206009 | 1 | only_DIvsD7 | UpDown | notDM | Loss | 8 |
| **3_72937645_72938828_55164** | 3_72937645_72938828 | SHQ1 | 55164 | 4.60223367438643 | -0.61110326 | 4 | only_DIvsD7 | UpDown | notDM | Gain | 2 |
| **3_81789244_81795468_2632** | 3_81789244_81795468 | GBE1 | 2632 | 0 | 0.00750062 | 7 | notSpecific | Down | DM | Gain | 9 |
| **3_93560126_93560606_200894** | 3_93560126_93560606 | ARL13B | 200894 | 5.14130237047762 | 0.96078127 | 1 | notSpecific | Up | DM | Gain | 9 |
| **3_9691273_9692892_64419** | 3_9691273_9692892 | MTMR14 | 64419 | 2.19865708695442 | 0.12926166 | 1 | notSpecific | Up | DM | Loss | 10 |
| **3_9745878_9747060_132014** | 3_9745878_9747060 | IL17RE | 132014 | 5.29546373657116 | -0.80479365 | 2 | only_DIvsD7 | DownUp | notDM | Loss | 7 |
| **3_9771463_9772788_132014** | 3_9771463_9772788 | IL17RE | 132014 | 5.23430463975834 | -0.78934707 | 1 | only_DIvsD7 | DownUp | notDM | Loss | 7 |
| **3_9850157_9852909_132014** | 3_9850157_9852909 | IL17RE | 132014 | 4.96092243718015 | -0.74860575 | 1 | only_DIvsD7 | DownUp | notDM | Gain | 2 |
| **3_9905791_9906902_132014** | 3_9905791_9906902 | IL17RE | 132014 | 4.57289482588338 | 0.56877709 | 7 | only_DIvsD7 | DownUp | notDM | Loss | 5 |
| **3_9919938_9921773_132014** | 3_9919938_9921773 | IL17RE | 132014 | 4.35278046757133 | 0.71826536 | 6 | only_DIvsD7 | DownUp | notDM | Loss | 5 |
| **3_9932659_9934164_132014** | 3_9932659_9934164 | IL17RE | 132014 | 4.00603795499732 | -0.22740199 | 1 | only_DIvsD7 | DownUp | notDM | Gain | 4 |
| **3_9938227_9941651_132014** | 3_9938227_9941651 | IL17RE | 132014 | 3.46478751964594 | 0.64882576 | 7 | only_DIvsD7 | DownUp | notDM | Loss | 5 |
| **3_9944657_9945067_132014** | 3_9944657_9945067 | IL17RE | 132014 | 1.96848294855394 | 0.41441069 | 6 | only_DIvsD7 | DownUp | notDM | Gain | 1 |
| **3_9958093_9959646_132014** | 3_9958093_9959646 | IL17RE | 132014 | 4.13299570069227 | 0.58000098 | 2 | only_DIvsD7 | DownUp | notDM | Loss | 6 |
| **3_9975553_9977096_132014** | 3_9975553_9977096 | IL17RE | 132014 | 4.49196368489569 | -0.37002055 | 2 | only_DIvsD7 | DownUp | notDM | Loss | 7 |
| **3_9987023_9988155_132014** | 3_9987023_9988155 | IL17RE | 132014 | 4.62852175275528 | 0.688333 | 1 | only_DIvsD7 | DownUp | notDM | Gain | 1 |
| **4_106628335_106629587_57117** | 4_106628335_106629587 | INTS12 | 57117 | 2.36172783601759 | 0.64168522 | 1 | only_H3vsUI | Down | notDM | Gain | 4 |
| **4_106630328_106632124_57117** | 4_106630328_106632124 | INTS12 | 57117 | 2.69196510276736 | 0.64025449 | 1 | only_H3vsUI | Down | notDM | Gain | 4 |
| **4_108664845_108665282_166929** | 4_108664845_108665282 | SGMS2 | 166929 | 4.90546126381727 | -0.66864046 | 7 | notSpecific | Up | DM | Gain | 9 |
| **4_120024261_120025916_54532** | 4_120024261_120025916 | USP53 | 54532 | 5.03335538893766 | 0.73611647 | 7 | notSpecific | Up | DM | Gain | 9 |
| **4_120548659_120550005_8654** | 4_120548659_120550005 | PDE5A | 8654 | 0 | 0.20894689 | 2 | only_DIvsD7 | UpDown | notDM | Loss | 3 |
| **4_123702938_123706265_11162** | 4_123702938_123706265 | NUDT6 | 11162 | 5.13829322254694 | 0.85038461 | 7 | only_H3vsUI | DownUp | notDM | Gain | 5 |
| **4_123718992_123720130_11162** | 4_123718992_123720130 | NUDT6 | 11162 | 5.09213089490095 | 0.79240026 | 7 | only_H3vsUI | DownUp | notDM | Loss | 1 |
| **4_123746787_123747569_11162** | 4_123746787_123747569 | NUDT6 | 11162 | 4.98477476818465 | 0.9052756 | 3 | only_H3vsUI | DownUp | notDM | Gain | 1 |
| **4_123793474_123797972_11162** | 4_123793474_123797972 | NUDT6 | 11162 | 4.6607706435277 | 0.76199023 | 7 | only_H3vsUI | DownUp | notDM | Gain | 5 |
| **4_123809379_123812160_11162** | 4_123809379_123812160 | NUDT6 | 11162 | 4.49971456874112 | 0.6181508 | 7 | only_H3vsUI | DownUp | notDM | Loss | 5 |
| **4_123818193_123820076_11162** | 4_123818193_123820076 | NUDT6 | 11162 | 4.37449172494382 | 0.37705307 | 7 | only_H3vsUI | DownUp | notDM | Gain | 1 |
| **4_123840990_123843475_11162** | 4_123840990_123843475 | NUDT6 | 11162 | 2.81224469680037 | 0.39194681 | 1 | only_H3vsUI | DownUp | notDM | Gain | 4 |
| **4_123844412_123846500_11162** | 4_123844412_123846500 | NUDT6 | 11162 | 2.46389298898591 | 0.39194681 | 2 | only_H3vsUI | DownUp | notDM | Loss | 6 |
| **4_128800335_128801870_10733** | 4_128800335_128801870 | PLK4 | 10733 | 2.25285303097989 | 0.29462282 | 1 | notSpecific | Down | DM | Gain | 9 |
| **4_128800335_128801870_22824** | 4_128800335_128801870 | HSPA4L | 22824 | 4.98614446471052 | 0.65109653 | 1 | notSpecific | UpDown | DM | Gain | 9 |
| **4_128800335_128801870_80167** | 4_128800335_128801870 | ABHD18 | 80167 | 4.92819094803876 | 0.71177812 | 1 | notSpecific | DownUp | DM | Gain | 9 |
| **4_140999751_141000634_10012985** | 4_140999751_141000634 | SCOC-AS1 | 100129858 | 5.46817150896512 | -0.89876285 | 7 | notSpecific | Up | DM | Gain | 9 |
| **4_141176979_141179230_60592** | 4_141176979_141179230 | SCOC | 60592 | 0 | 0.23422636 | 3 | only_H3vsUI | DownUp | notDM | Gain | 4 |

| | | | | | | | | | | | |
|---|---|---|---|---|---|---|---|---|---|---|---|
| **4_141264371_141264958_60592** | 4_141264371_141264958 | SCOC | 60592 | 0 | 0.01368488 | 4 | only_H3vsUI | DownUp | notDM | Gain | 1 |
| **4_141293091_141294329_60592** | 4_141293091_141294329 | SCOC | 60592 | 2.55870857053317 | 0.17518699 | 2 | only_H3vsUI | DownUp | notDM | Loss | 3 |
| **4_141295158_141296501_60592** | 4_141295158_141296501 | SCOC | 60592 | 2.67209785793572 | 0.18052726 | 1 | only_H3vsUI | DownUp | notDM | Loss | 3 |
| **4_144435394_144438276_10012805** | 4_144435394_144438276 | SMARCA5-AS1 | 100128055 | 0 | -0.04350486 | 1 | only_H3vsUI | Up | notDM | Gain | 4 |
| **4_144621561_144622850_166752** | 4_144621561_144622850 | FREM3 | 166752 | 0 | 0 | 4 | only_H3vsUI | UpDown | notDM | Loss | 3 |
| **4_146538891_146540207_166785** | 4_146538891_146540207 | MMAA | 166785 | 0 | 0 | 1 | only_H3vsUI | DownUp | notDM | Gain | 4 |
| **4_146540747_146542080_166785** | 4_146540747_146542080 | MMAA | 166785 | 3.12515582958053 | 0 | 1 | only_H3vsUI | DownUp | notDM | Gain | 4 |
| **4_153272124_153274763_201799** | 4_153272124_153274763 | TMEM154 | 201799 | 5.51372353808501 | 0.80017892 | 4 | only_DIvsD7 | DownUp | notDM | Loss | 3 |
| **4_153699123_153700759_23321** | 4_153699123_153700759 | TRIM2 | 23321 | 5.62822540016721 | -0.94342545 | 1 | only_DIvsD7 | UpDown | notDM | Loss | 7 |
| **4_153701490_153702960_23321** | 4_153701490_153702960 | TRIM2 | 23321 | 5.62596957010223 | -0.94342545 | 2 | only_DIvsD7 | UpDown | notDM | Gain | 2 |
| **4_154057497_154058665_23321** | 4_154057497_154058665 | TRIM2 | 23321 | 4.82577652796543 | -0.7469341 | 7 | only_DIvsD7 | UpDown | notDM | Gain | 1 |
| **4_154064148_154065797_23321** | 4_154064148_154065797 | TRIM2 | 23321 | 3.92808845966497 | 0.02161968 | 3 | only_DIvsD7 | UpDown | notDM | Loss | 3 |
| **4_154073551_154075947_23321** | 4_154073551_154075947 | TRIM2 | 23321 | 0 | 0.04572625 | 2 | only_DIvsD7 | UpDown | notDM | Loss | 3 |
| **4_154154010_154156367_23321** | 4_154154010_154156367 | TRIM2 | 23321 | 4.34566773563535 | 0.46892027 | 3 | only_DIvsD7 | UpDown | notDM | Gain | 1 |
| **4_165674895_165676187_10050598** | 4_165674895_165676187 | SMIM31 | 100505989 | 0 | 0 | 3 | only_H3vsUI | Up | notDM | Gain | 4 |
| **4_170578719_170583341_54969** | 4_170578719_170583341 | HPF1 | 54969 | 4.98120227797528 | -0.89297903 | 5 | only_H3vsUI | Down | notDM | Gain | 4 |
| **4_170677181_170678890_54969** | 4_170677181_170678890 | HPF1 | 54969 | 2.33243845991561 | 0.33501147 | 1 | only_H3vsUI | Down | notDM | Loss | 8 |
| **4_170712138_170712992_54969** | 4_170712138_170712992 | HPF1 | 54969 | 4.51898745744096 | -0.40005039 | 4 | only_H3vsUI | Down | notDM | Gain | 4 |
| **4_178228821_178230841_55247** | 4_178228821_178230841 | NEIL3 | 55247 | 2.17609125905568 | -0.16276924 | 7 | notSpecific | Down | DM | Gain | 9 |
| **4_184557632_184560401_201965** | 4_184557632_184560401 | RWDD4 | 201965 | 4.30055200879289 | -0.3237672 | 5 | only_H3vsUI | UpDown | notDM | Gain | 1 |
| **4_184580752_184582026_201965** | 4_184580752_184582026 | RWDD4 | 201965 | 2.57518784492766 | 0.35621535 | 1 | only_H3vsUI | UpDown | notDM | Gain | 4 |
| **4_186123883_186124848_441054** | 4_186123883_186124848 | C4orf47 | 441054 | 5.34743930714142 | -0.75354931 | 1 | only_H3vsUI | Up | notDM | Gain | 2 |
| **4_186125430_186126573_441054** | 4_186125430_186126573 | C4orf47 | 441054 | 5.34406003909306 | -0.75354931 | 1 | only_H3vsUI | Up | notDM | Gain | 2 |
| **4_186315881_186317419_441054** | 4_186315881_186317419 | C4orf47 | 441054 | 4.4769040531738 | -0.58580508 | 1 | only_H3vsUI | Up | notDM | Loss | 3 |
| **4_186317903_186318760_441054** | 4_186317903_186318760 | C4orf47 | 441054 | 4.45703366505438 | -0.58580508 | 2 | only_H3vsUI | Up | notDM | Loss | 3 |
| **4_186345492_186346784_441054** | 4_186345492_186346784 | C4orf47 | 441054 | 2.79239168949825 | -0.94465361 | 2 | only_H3vsUI | Up | notDM | Loss | 3 |
| **4_186391443_186392201_441054** | 4_186391443_186392201 | C4orf47 | 441054 | 4.64386703258041 | -0.60481108 | 2 | only_H3vsUI | Up | notDM | Loss | 3 |
| **4_2413507_2415192_402160** | 4_2413507_2415192 | CFAP99 | 402160 | 3.74115159885178 | 0.00417617 | 5 | only_H3vsUI | UpDown | notDM | Loss | 6 |
| **4_2418011_2419036_402160** | 4_2418011_2419036 | CFAP99 | 402160 | 3.22167499707077 | 0.00589734 | 2 | only_H3vsUI | UpDown | notDM | Loss | 3 |
| **4_2420701_2421624_402160** | 4_2420701_2421624 | CFAP99 | 402160 | 0.301029995663981 | 0.00729498 | 2 | only_H3vsUI | UpDown | notDM | Loss | 3 |
| **4_24552785_24554480_1665** | 4_24552785_24554480 | DHX15 | 1665 | 4.50075775447104 | -0.57295156 | 8 | notSpecific | DownUp | DM | Gain | 9 |
| **4_38671629_38677792_81793** | 4_38671629_38677792 | TLR10 | 81793 | 5.02852245281812 | 0.66826171 | 7 | only_H3vsUI | UpDown | notDM | Loss | 5 |
| **4_39184324_39185744_81793** | 4_39184324_39185744 | TLR10 | 81793 | 5.60175153273005 | -0.95294583 | 1 | only_H3vsUI | UpDown | notDM | Loss | 3 |
| **4_4387543_4389259_27065** | 4_4387543_4389259 | NSG1 | 27065 | 0 | -0.36289516 | 4 | notSpecific | Up | DM | Loss | 10 |

| | | | | | | | | | | | |
|---|---|---|---|---|---|---|---|---|---|---|---|
| **4_4387543_4389259_53407** | 4_4387543_4389259 | STX18 | 53407 | 5.18892005077642 | 0.89109571 | 4 | notSpecific | UpDown | DM | Loss | 10 |
| **4_5713388_5714766_132884** | 4_5713388_5714766 | EVC2 | 132884 | 3.49080095201086 | 0.26837661 | 1 | only_H3vsUI | Down | notDM | Loss | 3 |
| **4_5722915_5725602_132884** | 4_5722915_5725602 | EVC2 | 132884 | 4.10116258221484 | -0.16971944 | 7 | only_H3vsUI | Down | notDM | Loss | 4 |
| **4_57841212_57843155_5431** | 4_57841212_57843155 | POLR2B | 5431 | 3.27323283404305 | -0.47495001 | 1 | notSpecific | Down | DM | Loss | 10 |
| **4_57841212_57843155_5978** | 4_57841212_57843155 | REST | 5978 | 4.81860853776137 | -0.8326305 | 1 | notSpecific | Down | DM | Loss | 10 |
| **4_57841212_57843155_84273** | 4_57841212_57843155 | NOA1 | 84273 | 3.26363606858811 | 0.01637619 | 1 | notSpecific | Up | DM | Loss | 10 |
| **4_6274549_6275760_7466** | 4_6274549_6275760 | WFS1 | 7466 | 3.46731206298055 | 0.67906651 | 6 | only_H3vsUI | DownUp | notDM | Loss | 3 |
| **4_6577073_6577951_23324** | 4_6577073_6577951 | MAN2B2 | 23324 | 2.20139712432045 | 0.32112802 | 1 | only_DIvsD7 | DownUp | notDM | Loss | 3 |
| **4_6710122_6711514_23324** | 4_6710122_6711514 | MAN2B2 | 23324 | 5.12453030780368 | -0.7132174 | 1 | only_DIvsD7 | DownUp | notDM | Gain | 2 |
| **4_74210235_74212266_213** | 4_74210235_74212266 | ALB | 213 | 4.76110806696504 | 0.9832167 | 3 | only_H3vsUI | DownUp | notDM | Gain | 1 |
| **4_74558288_74558989_213** | 4_74558288_74558989 | ALB | 213 | 5.43620400645052 | 0.9175517 | 6 | only_H3vsUI | DownUp | notDM | Gain | 1 |
| **4_74569167_74572213_2920** | 4_74569167_74572213 | CXCL2 | 2920 | 5.59406090127042 | 0.84226785 | 7 | only_H3vsUI | Up | DM | Gain | 9 |
| **4_74569167_74572213_3576** | 4_74569167_74572213 | CXCL8 | 3576 | 4.53206609481055 | 0.83330817 | 7 | notSpecific | Up | DM | Gain | 9 |
| **4_74869675_74872996_2920** | 4_74869675_74872996 | CXCL2 | 2920 | 4.96339584134703 | 0.77262033 | 7 | only_H3vsUI | Up | notDM | Gain | 5 |
| **4_74869675_74872996_2921** | 4_74869675_74872996 | CXCL3 | 2921 | 4.49654220633805 | 0.91165734 | 7 | only_H3vsUI | Up | notDM | Gain | 5 |
| **4_74888603_74889670_2921** | 4_74888603_74889670 | CXCL3 | 2921 | 4.16725824304363 | 0.15604912 | 7 | only_H3vsUI | Up | DM | Gain | 9 |
| **4_74888603_74889670_6374** | 4_74888603_74889670 | CXCL5 | 6374 | 4.38217919696025 | -0.64386986 | 7 | notSpecific | Up | DM | Gain | 9 |
| **4_74889739_74890560_2921** | 4_74889739_74890560 | CXCL3 | 2921 | 4.14013077837113 | 0.3097002 | 7 | only_H3vsUI | Up | notDM | Loss | 1 |
| **4_74895853_74896385_2921** | 4_74895853_74896385 | CXCL3 | 2921 | 3.9106244048892 | 0.31388677 | 3 | only_H3vsUI | Up | notDM | Gain | 1 |
| **4_74901760_74904108_2920** | 4_74901760_74904108 | CXCL2 | 2920 | 4.78393929281101 | 0.64786248 | 1 | only_H3vsUI | Up | notDM | Loss | 6 |
| **4_74901760_74904108_2921** | 4_74901760_74904108 | CXCL3 | 2921 | 2.47567118832443 | 0.79560044 | 1 | only_H3vsUI | Up | notDM | Loss | 6 |
| **4_74904597_74906168_2920** | 4_74904597_74906168 | CXCL2 | 2920 | 4.76897090813836 | 0.64866498 | 1 | only_H3vsUI | Up | notDM | Loss | 6 |
| **4_74904597_74906168_2921** | 4_74904597_74906168 | CXCL3 | 2921 | 2.28555730900777 | 0.7975263 | 1 | only_H3vsUI | Up | notDM | Loss | 6 |
| **4_74961972_74964618_2920** | 4_74961972_74964618 | CXCL2 | 2920 | 2.46982201597816 | 0.79986713 | 1 | only_H3vsUI | Up | notDM | Loss | 6 |
| **4_74961972_74964618_2921** | 4_74961972_74964618 | CXCL3 | 2921 | 4.76047525905971 | 0.71913938 | 1 | only_H3vsUI | Up | notDM | Loss | 6 |
| **4_74969743_74970775_2920** | 4_74969743_74970775 | CXCL2 | 2920 | 3.68421679513888 | 0.34592206 | 3 | only_H3vsUI | Up | notDM | Gain | 1 |
| **4_74969743_74970775_2921** | 4_74969743_74970775 | CXCL3 | 2921 | 4.81543163080156 | 0.59324685 | 3 | only_H3vsUI | Up | notDM | Gain | 1 |
| **4_74973416_74975445_2920** | 4_74973416_74975445 | CXCL2 | 2920 | 3.929725378378 | 0.84941765 | 7 | only_H3vsUI | Up | notDM | Gain | 5 |
| **4_74973416_74975445_2921** | 4_74973416_74975445 | CXCL3 | 2921 | 4.83892461364535 | 0.78100013 | 7 | only_H3vsUI | Up | notDM | Gain | 5 |
| **4_74975519_74977805_2920** | 4_74975519_74977805 | CXCL2 | 2920 | 4.02567444941034 | 0.8270661 | 7 | only_H3vsUI | Up | notDM | Gain | 5 |
| **4_74975519_74977805_2921** | 4_74975519_74977805 | CXCL3 | 2921 | 4.85219931944628 | 0.70879015 | 7 | only_H3vsUI | Up | notDM | Gain | 5 |
| **4_74982006_74984033_2920** | 4_74982006_74984033 | CXCL2 | 2920 | 4.23289450915601 | 0.91725443 | 7 | only_H3vsUI | Up | notDM | Loss | 5 |
| **4_74982006_74984033_2921** | 4_74982006_74984033 | CXCL3 | 2921 | 4.89009112037533 | 0.78632613 | 7 | only_H3vsUI | Up | notDM | Loss | 5 |
| **4_75241301_75242734_2921** | 4_75241301_75242734 | CXCL3 | 2921 | 5.52754741574834 | 0.92084786 | 7 | only_H3vsUI | Up | notDM | Loss | 5 |

| | | | | | | | | | | | |
|---|---|---|---|---|---|---|---|---|---|---|---|
| **4_76438181_76439291_25898** | 4_76438181_76439291 | RCHY1 | 25898 | 2.33645973384853 | -0.2199299 | 1 | only_H3vsUI | Down | notDM | Gain | 4 |
| **4_76439954_76441618_25898** | 4_76439954_76441618 | RCHY1 | 25898 | 2.65224634100332 | -0.2199299 | 2 | only_H3vsUI | Down | notDM | Loss | 8 |
| **4_77490420_77493472_55752** | 4_77490420_77493472 | SEPTIN11 | 55752 | 5.57678463417896 | -0.82987733 | 4 | only_H3vsUI | DownUp | notDM | Loss | 7 |
| **4_77751346_77752237_55752** | 4_77751346_77752237 | SEPTIN11 | 55752 | 5.07454583622544 | -0.80395111 | 3 | only_H3vsUI | DownUp | notDM | Gain | 2 |
| **4_77752355_77753614_55752** | 4_77752355_77753614 | SEPTIN11 | 55752 | 5.06947939307111 | -0.80250264 | 3 | only_H3vsUI | DownUp | notDM | Gain | 2 |
| **4_77818784_77819773_345079** | 4_77818784_77819773 | SOWAHB | 345079 | 0 | 0 | 4 | only_H3vsUI | UpDown | notDM | Loss | 3 |
| **4_77869283_77870609_55752** | 4_77869283_77870609 | SEPTIN11 | 55752 | 2.39445168082622 | 0.71747846 | 1 | only_H3vsUI | DownUp | notDM | Loss | 3 |
| **4_77871433_77873459_55752** | 4_77871433_77873459 | SEPTIN11 | 55752 | 2.76267856372744 | 0.16319848 | 2 | only_H3vsUI | DownUp | notDM | Loss | 8 |
| **4_78738529_78739868_246175** | 4_78738529_78739868 | CNOT6L | 246175 | 2.81624129999178 | 0.56692805 | 1 | only_H3vsUI | Down | notDM | Loss | 8 |
| **4_78741603_78742494_246175** | 4_78741603_78742494 | CNOT6L | 246175 | 3.03462845662532 | 0.46109065 | 2 | only_H3vsUI | Down | notDM | Gain | 4 |
| **4_78841490_78844320_246175** | 4_78841490_78844320 | CNOT6L | 246175 | 5.00409771904421 | 0.64000157 | 6 | only_H3vsUI | Down | notDM | Gain | 1 |
| **4_83351953_83353889_3184** | 4_83351953_83353889 | HNRNPD | 3184 | 4.75681861048214 | -0.7552476 | 1 | notSpecific | DownUp | DM | Loss | 10 |
| **4_83351953_83353889_58478** | 4_83351953_83353889 | ENOPH1 | 58478 | 2.38021124171161 | -0.8800294 | 1 | notSpecific | DownUp | DM | Loss | 10 |
| **4_83351953_83353889_9987** | 4_83351953_83353889 | HNRNPDL | 9987 | 3.13893394025692 | -0.39987703 | 1 | notSpecific | Up | DM | Loss | 10 |
| **4_84203511_84205505_27235** | 4_84203511_84205505 | COQ2 | 27235 | 2.6170003411209 | -0.01248954 | 1 | only_DIvsD7 | Down | notDM | Loss | 8 |
| **4_84206067_84207103_27235** | 4_84206067_84207103 | COQ2 | 27235 | 2.17897694729317 | -0.01248954 | 1 | only_DIvsD7 | Down | notDM | Loss | 8 |
| **4_90031490_90032176_166815** | 4_90031490_90032176 | TIGD2 | 166815 | 3.25358028956218 | 0.04162443 | 2 | only_H3vsUI | DownUp | notDM | Gain | 2 |
| **4_90032872_90034477_166815** | 4_90032872_90034477 | TIGD2 | 166815 | 0 | 0.03818567 | 1 | only_H3vsUI | DownUp | notDM | Loss | 8 |
| **4_90040300_90041585_166815** | 4_90040300_90041585 | TIGD2 | 166815 | 3.80167805903589 | 0.00860174 | 7 | only_H3vsUI | DownUp | notDM | Gain | 1 |
| **4_90043581_90044797_166815** | 4_90043581_90044797 | TIGD2 | 166815 | 3.9829492885745 | 0.04977543 | 7 | only_H3vsUI | DownUp | notDM | Gain | 1 |
| **4_90044879_90046086_166815** | 4_90044879_90046086 | TIGD2 | 166815 | 4.0379441552031 | 0.05004504 | 7 | only_H3vsUI | DownUp | notDM | Gain | 1 |
| **4_96023073_96024062_658** | 4_96023073_96024062 | BMPR1B | 658 | 3.15986784709257 | 0 | 6 | notSpecific | Down | DM | Gain | 9 |
| **5_110844793_110847818_10050567** | 5_110844793_110847818 | STARD4-AS1 | 100505678 | 2.02938377768521 | -0.0974439 | 2 | only_H3vsUI | UpDown | notDM | Gain | 4 |
| **5_110848472_110849933_10050567** | 5_110848472_110849933 | STARD4-AS1 | 100505678 | 2.74036268949424 | -0.09297996 | 2 | only_H3vsUI | UpDown | notDM | Gain | 4 |
| **5_115174890_115176792_9140** | 5_115174890_115176792 | ATG12 | 9140 | 2.66745295288995 | 0.26161141 | 1 | only_DIvsD7 | Up | notDM | Gain | 4 |
| **5_115178151_115179721_9140** | 5_115178151_115179721 | ATG12 | 9140 | 2.95279244304409 | 0.28461272 | 2 | only_DIvsD7 | Up | notDM | Loss | 6 |
| **5_115185982_115187410_9140** | 5_115185982_115187410 | ATG12 | 9140 | 3.94091473758029 | -0.36380086 | 8 | only_DIvsD7 | Up | notDM | Loss | 7 |
| **5_122423240_122426797_28966** | 5_122423240_122426797 | SNX24 | 28966 | 5.38367356901615 | -0.78584127 | 4 | notSpecific | Down | DM | Loss | 10 |
| **5_122423240_122426797_5480** | 5_122423240_122426797 | PPIC | 5480 | 4.70591500387879 | -0.90549219 | 4 | notSpecific | Down | DM | Loss | 10 |
| **5_122423240_122426797_93166** | 5_122423240_122426797 | PRDM6 | 93166 | 0 | 0 | 4 | notSpecific | DownUp | DM | Loss | 10 |
| **5_126852437_126853157_133619** | 5_126852437_126853157 | PRRC1 | 133619 | 2.16136800223497 | -0.1925699 | 2 | notSpecific | Down | DM | Gain | 9 |
| **5_127040633_127041690_84466** | 5_127040633_127041690 | MEGF10 | 84466 | 5.61711781552598 | 0.99981828 | 6 | notSpecific | Up | DM | Gain | 9 |
| **5_129240157_129242600_337876** | 5_129240157_129242600 | CHSY3 | 337876 | 0 | 0.2217118 | 4 | notSpecific | Up | DM | Loss | 10 |
| **5_130497955_130500072_90624** | 5_130497955_130500072 | LYRM7 | 90624 | 3.81730087839332 | 0.08209276 | 1 | only_H3vsUI | DownUp | notDM | Gain | 5 |

| | | | | | | | | | | | |
|---|---|---|---|---|---|---|---|---|---|---|---|
| **5_130501047_130502127_90624** | 5_130501047_130502127 | LYRM7 | 90624 | 3.65427282709771 | 0.08209276 | 5 | only_H3vsUI | DownUp | notDM | Loss | 7 |
| **5_130505425_130506323_90624** | 5_130505425_130506323 | LYRM7 | 90624 | 2.55630250076729 | 0.38713139 | 1 | only_H3vsUI | DownUp | notDM | Loss | 6 |
| **5_130506718_130507929_90624** | 5_130506718_130507929 | LYRM7 | 90624 | 1.91907809237607 | 0.19993004 | 1 | only_H3vsUI | DownUp | notDM | Loss | 8 |
| **5_131698142_131702128_553103** | 5_131698142_131702128 | MIR3936HG | 553103 | 3.50691072555152 | 0.38328115 | 7 | only_DIvsD7 | UpDown | notDM | Gain | 5 |
| **5_131705556_131709571_553103** | 5_131705556_131709571 | MIR3936HG | 553103 | 2.3384564936046 | 0.23733295 | 2 | only_DIvsD7 | UpDown | notDM | Loss | 6 |
| **5_132201133_132202104_2661** | 5_132201133_132202104 | GDF9 | 2661 | 2.13033376849501 | -0.00579172 | 1 | only_H3vsUI | Up | notDM | Loss | 3 |
| **5_132202532_132204236_2661** | 5_132202532_132204236 | GDF9 | 2661 | 0 | 0 | 1 | only_H3vsUI | Up | notDM | Loss | 3 |
| **5_137468089_137469552_10902** | 5_137468089_137469552 | BRD8 | 10902 | 4.65440759055263 | -0.57477384 | 2 | only_H3vsUI | Down | notDM | Gain | 4 |
| **5_137512703_137514609_10902** | 5_137512703_137514609 | BRD8 | 10902 | 0 | -0.23149873 | 1 | only_H3vsUI | Down | notDM | Gain | 4 |
| **5_137514757_137517034_10902** | 5_137514757_137517034 | BRD8 | 10902 | 2.6730209071289 | -0.23149873 | 1 | only_H3vsUI | Down | notDM | Loss | 8 |
| **5_137547422_137548845_10902** | 5_137547422_137548845 | BRD8 | 10902 | 4.51519804322513 | 0.36245861 | 1 | only_H3vsUI | Down | notDM | Gain | 1 |
| **5_137801319_137802710_1958** | 5_137801319_137802710 | EGR1 | 1958 | 2.15228834438306 | -0.11804094 | 1 | notSpecific | Up | DM | Loss | 10 |
| **5_137801319_137802710_51307** | 5_137801319_137802710 | FAM53C | 51307 | 5.10505009993651 | -0.69176823 | 1 | notSpecific | Up | DM | Loss | 10 |
| **5_137801319_137802710_51308** | 5_137801319_137802710 | REEP2 | 51308 | 4.42512647050877 | -0.65661511 | 1 | notSpecific | DownUp | DM | Loss | 10 |
| **5_138606391_138614868_340061** | 5_138606391_138614868 | STING1 | 340061 | 5.39272156844364 | 0.86889048 | 1 | only_H3vsUI | DownUp | notDM | Loss | 6 |
| **5_138628694_138629203_340061** | 5_138628694_138629203 | STING1 | 340061 | 5.36685983379705 | 0.73805797 | 3 | only_H3vsUI | DownUp | notDM | Gain | 1 |
| **5_138629978_138634880_340061** | 5_138629978_138634880 | STING1 | 340061 | 5.35613489542224 | 0.74307877 | 1 | only_H3vsUI | DownUp | notDM | Loss | 6 |
| **5_138726478_138728382_202051** | 5_138726478_138728382 | SPATA24 | 202051 | 4.05568406491941 | -0.08291723 | 2 | only_H3vsUI | UpDown | notDM | Loss | 6 |
| **5_138738566_138739657_202051** | 5_138738566_138739657 | SPATA24 | 202051 | 2.07918124604762 | -0.0961343 | 1 | only_H3vsUI | UpDown | notDM | Loss | 3 |
| **5_138739843_138740571_202051** | 5_138739843_138740571 | SPATA24 | 202051 | 1.83250891270624 | -0.0961343 | 3 | only_H3vsUI | UpDown | notDM | Loss | 4 |
| **5_139174732_139175278_340061** | 5_139174732_139175278 | STING1 | 340061 | 5.49534032385629 | 0.82482965 | 4 | only_H3vsUI | DownUp | notDM | Loss | 3 |
| **5_139283404_139284587_9542** | 5_139283404_139284587 | NRG2 | 9542 | 0 | 0 | 4 | notSpecific | Up | DM | Loss | 10 |
| **5_139492691_139493401_492311** | 5_139492691_139493401 | IGIP | 492311 | 4.08353845123014 | 0.06324837 | 5 | only_H3vsUI | Down | notDM | Loss | 2 |
| **5_139494045_139496345_492311** | 5_139494045_139496345 | IGIP | 492311 | 3.96270073170434 | 0.05928564 | 1 | only_H3vsUI | Down | notDM | Loss | 8 |
| **5_139502344_139503720_492311** | 5_139502344_139503720 | IGIP | 492311 | 3.25575478664304 | 0.04073167 | 6 | only_H3vsUI | Down | notDM | Gain | 1 |
| **5_140012307_140014667_929** | 5_140012307_140014667 | CD14 | 929 | 0 | -0.00229081 | 3 | only_DIvsD7 | Up | notDM | Gain | 4 |
| **5_140017837_140018861_929** | 5_140017837_140018861 | CD14 | 929 | 3.65829765030819 | 0.00013573 | 1 | only_DIvsD7 | Up | notDM | Loss | 3 |
| **5_140019307_140020277_929** | 5_140019307_140020277 | CD14 | 929 | 3.77988496319264 | -0.00159968 | 1 | only_DIvsD7 | Up | notDM | Gain | 4 |
| **5_140043196_140044336_373863** | 5_140043196_140044336 | DND1 | 373863 | 3.9462557071994 | -0.05546452 | 1 | only_H3vsUI | DownUp | notDM | Loss | 7 |
| **5_140044678_140046733_373863** | 5_140044678_140046733 | DND1 | 373863 | 3.80881842509212 | -0.08241474 | 1 | only_H3vsUI | DownUp | notDM | Loss | 8 |
| **5_141392719_141394027_80762** | 5_141392719_141394027 | NDFIP1 | 80762 | 4.97463602224574 | 0.70047004 | 1 | only_DIvsD7 | UpDown | notDM | Loss | 6 |
| **5_141452221_141454025_80762** | 5_141452221_141454025 | NDFIP1 | 80762 | 4.53566115265944 | -0.70088648 | 6 | only_DIvsD7 | UpDown | notDM | Loss | 7 |
| **5_141488766_141491326_80762** | 5_141488766_141491326 | NDFIP1 | 80762 | 2.61804809671209 | 0.92978347 | 2 | only_DIvsD7 | UpDown | notDM | Gain | 4 |
| **5_141594963_141596793_80762** | 5_141594963_141596793 | NDFIP1 | 80762 | 5.027708313254 | 0.76547661 | 6 | only_DIvsD7 | UpDown | notDM | Loss | 5 |

| | | | | | | | | | | | |
|---|---|---|---|---|---|---|---|---|---|---|---|
| **5_146833809_146834867_1809** | 5_146833809_146834867 | DPYSL3 | 1809 | 2.46089784275655 | 0.98694326 | 2 | only_H3vsUI | DownUp | notDM | Gain | 4 |
| **5_146887533_146890530_1809** | 5_146887533_146890530 | DPYSL3 | 1809 | 0 | 0.57367232 | 1 | only_H3vsUI | DownUp | notDM | Loss | 8 |
| **5_149641475_149644796_815** | 5_149641475_149644796 | CAMK2A | 815 | 2.21748394421391 | -0.00106193 | 3 | only_H3vsUI | UpDown | notDM | Loss | 6 |
| **5_153823864_153825322_386627** | 5_153823864_153825322 | SAP30L-AS1 | 386627 | 1.94939000664491 | -0.0246915 | 1 | only_DIvsD7 | UpDown | notDM | Gain | 4 |
| **5_153825981_153827717_386627** | 5_153825981_153827717 | SAP30L-AS1 | 386627 | 2.78103693862113 | 0.35060413 | 1 | only_DIvsD7 | UpDown | notDM | Loss | 3 |
| **5_156119381_156121501_26762** | 5_156119381_156121501 | HAVCR1 | 26762 | 5.56185239744695 | 0.99940624 | 6 | only_H3vsUI | DownUp | notDM | Loss | 5 |
| **5_156472193_156475640_26762** | 5_156472193_156475640 | HAVCR1 | 26762 | 4.02081688702891 | 0.99971167 | 7 | only_H3vsUI | DownUp | notDM | Gain | 5 |
| **5_156475801_156488549_26762** | 5_156475801_156488549 | HAVCR1 | 26762 | 0 | 0.83064618 | 5 | only_H3vsUI | DownUp | notDM | Gain | 4 |
| **5_156498653_156501077_26762** | 5_156498653_156501077 | HAVCR1 | 26762 | 4.09777773453928 | 0.42587225 | 6 | only_H3vsUI | DownUp | notDM | Gain | 5 |
| **5_156568289_156569681_9443** | 5_156568289_156569681 | MED7 | 9443 | 2.02530586526477 | -0.18842282 | 2 | only_H3vsUI | Down | notDM | Loss | 8 |
| **5_156570084_156571240_9443** | 5_156570084_156571240 | MED7 | 9443 | 2.47712125471966 | -0.18842282 | 1 | only_H3vsUI | Down | notDM | Loss | 8 |
| **5_159300357_159302216_10192776** | 5_159300357_159302216 | LINC01847 | 101927766 | 3.29402509409532 | 0 | 7 | only_H3vsUI | UpDown | notDM | Gain | 1 |
| **5_159337245_159343289_147** | 5_159337245_159343289 | ADRA1B | 147 | 2.70070371714502 | 0.00165193 | 7 | only_H3vsUI | DownUp | notDM | Loss | 4 |
| **5_159343661_159347241_147** | 5_159343661_159347241 | ADRA1B | 147 | 0 | 0.00330521 | 2 | only_H3vsUI | DownUp | notDM | Loss | 8 |
| **5_169415541_169417410_10013189** | 5_169415541_169417410 | INSYN2B | 100131897 | 3.90800220117255 | 0.57104398 | 3 | notSpecific | Down | DM | Gain | 9 |
| **5_169415541_169417410_1794** | 5_169415541_169417410 | DOCK2 | 1794 | 3.73487980279263 | 0.01151941 | 3 | notSpecific | DownUp | DM | Gain | 9 |
| **5_170844903_170846474_8817** | 5_170844903_170846474 | FGF18 | 8817 | 2.2718416065365 | 0.0086067 | 2 | only_DIvsD7 | Up | notDM | Loss | 3 |
| **5_172192637_172197155_1843** | 5_172192637_172197155 | DUSP1 | 1843 | 3.01870049866624 | 0.30082566 | 1 | notSpecific | Up | DM | Loss | 10 |
| **5_172378115_172379362_10026816** | 5_172378115_172379362 | LOC100268168 | 100268168 | 3.80753502806885 | 0.03255955 | 7 | notSpecific | Down | DM | Loss | 10 |
| **5_172378115_172379362_51121** | 5_172378115_172379362 | RPL26L1 | 51121 | 3.81524559191656 | 0.66657786 | 7 | notSpecific | Up | DM | Loss | 10 |
| **5_175083896_175084963_3274** | 5_175083896_175084963 | HRH2 | 3274 | 1.85125834871908 | 0.10846802 | 2 | only_DIvsD7 | DownUp | notDM | Loss | 3 |
| **5_175841619_175843352_1212** | 5_175841619_175843352 | CLTB | 1212 | 0 | 0.43369068 | 1 | only_H3vsUI | UpDown | notDM | Gain | 4 |
| **5_175863342_175864355_1212** | 5_175863342_175864355 | CLTB | 1212 | 4.29609452982012 | 0.44335912 | 4 | only_H3vsUI | UpDown | notDM | Loss | 3 |
| **5_175865670_175866526_1212** | 5_175865670_175866526 | CLTB | 1212 | 4.35044185653506 | -0.28380049 | 6 | only_H3vsUI | UpDown | notDM | Loss | 3 |
| **5_175914962_175919771_1212** | 5_175914962_175919771 | CLTB | 1212 | 4.85553126971034 | 0.88287223 | 7 | only_H3vsUI | UpDown | notDM | Loss | 5 |
| **5_175959819_175963869_54825** | 5_175959819_175963869 | CDHR2 | 54825 | 4.09774305894488 | 0.30819827 | 1 | only_H3vsUI | Up | notDM | Loss | 6 |
| **5_175964512_175964901_54825** | 5_175964512_175964901 | CDHR2 | 54825 | 4.06039561731991 | 0.30819827 | 1 | only_H3vsUI | Up | notDM | Gain | 1 |
| **5_175965058_175965857_54825** | 5_175965058_175965857 | CDHR2 | 54825 | 4.02267576195373 | 0.30819827 | 4 | only_H3vsUI | Up | notDM | Gain | 1 |
| **5_175968223_175970080_54825** | 5_175968223_175970080 | CDHR2 | 54825 | 0 | 0.01450648 | 1 | only_H3vsUI | Up | notDM | Loss | 3 |
| **5_175970553_175972081_54825** | 5_175970553_175972081 | CDHR2 | 54825 | 3.01828430842653 | 0.01323185 | 5 | only_H3vsUI | Up | notDM | Loss | 3 |
| **5_175972144_175973309_54825** | 5_175972144_175973309 | CDHR2 | 54825 | 3.48911436937892 | 0.08672419 | 6 | only_H3vsUI | Up | notDM | Loss | 3 |
| **5_176074524_176076084_1212** | 5_176074524_176076084 | CLTB | 1212 | 5.36410803303791 | -0.79970876 | 1 | only_H3vsUI | UpDown | notDM | Loss | 7 |
| **5_176880731_176883436_1627** | 5_176880731_176883436 | DBN1 | 1627 | 4.2382217936778 | -0.36010709 | 2 | notSpecific | Down | DM | Loss | 10 |
| **5_176880731_176883436_2870** | 5_176880731_176883436 | GRK6 | 2870 | 4.43223148569021 | -0.79264409 | 2 | notSpecific | Up | DM | Loss | 10 |

| | | | | | | | | | | | |
|---|---|---|---|---|---|---|---|---|---|---|---|
| **5_176880731_176883436_80758** | 5_176880731_176883436 | PRR7 | 80758 | 0 | -0.00548241 | 2 | notSpecific | DownUp | DM | Loss | 10 |
| **5_176880731_176883436_83463** | 5_176880731_176883436 | MXD3 | 83463 | 5.15182934013187 | 0.67579377 | 2 | notSpecific | Up | DM | Loss | 10 |
| **5_177539762_177540360_64777** | 5_177539762_177540360 | RMND5B | 64777 | 4.24644933826704 | 0.34229438 | 2 | only_DIvsD7 | Down | notDM | Gain | 2 |
| **5_177540538_177541528_23138** | 5_177540538_177541528 | N4BP3 | 23138 | 1.98227123303957 | 0.02465396 | 2 | notSpecific | Up | DM | Loss | 10 |
| **5_177540538_177541528_55651** | 5_177540538_177541528 | NHP2 | 55651 | 4.59594791765295 | 0.82822219 | 2 | notSpecific | Up | DM | Loss | 10 |
| **5_177540538_177541528_64777** | 5_177540538_177541528 | RMND5B | 64777 | 4.21669359916975 | 0.33454173 | 2 | only_DIvsD7 | Down | DM | Loss | 10 |
| **5_177556520_177557871_64777** | 5_177556520_177557871 | RMND5B | 64777 | 2.22010808804005 | -0.20013056 | 1 | only_DIvsD7 | Down | notDM | Loss | 8 |
| **5_177558174_177559716_64777** | 5_177558174_177559716 | RMND5B | 64777 | 2.14612803567824 | -0.20013056 | 1 | only_DIvsD7 | Down | notDM | Loss | 8 |
| **5_177578953_177580568_64777** | 5_177578953_177580568 | RMND5B | 64777 | 4.32054091992458 | -0.80640874 | 2 | only_DIvsD7 | Down | notDM | Gain | 2 |
| **5_178051200_178053575_64777** | 5_178051200_178053575 | RMND5B | 64777 | 5.69299312769382 | -0.8578126 | 1 | only_DIvsD7 | Down | notDM | Loss | 7 |
| **5_178979148_178979998_3187** | 5_178979148_178979998 | HNRNPH1 | 3187 | 4.85493728389172 | 0.69380741 | 8 | only_DIvsD7 | DownUp | notDM | Loss | 5 |
| **5_179043971_179049749_3187** | 5_179043971_179049749 | HNRNPH1 | 3187 | 1.68124123737559 | 0.2613381 | 2 | only_DIvsD7 | DownUp | notDM | Loss | 6 |
| **5_179050939_179052240_3187** | 5_179050939_179052240 | HNRNPH1 | 3187 | 0 | 0.085371 | 1 | only_DIvsD7 | DownUp | notDM | Loss | 6 |
| **5_179050939_179052240_9794** | 5_179050939_179052240 | MAML1 | 9794 | 5.03192123489974 | -0.62779242 | 1 | only_DIvsD7 | UpDown | notDM | Loss | 7 |
| **5_179060412_179062407_3187** | 5_179060412_179062407 | HNRNPH1 | 3187 | 0 | 0.00596957 | 1 | only_DIvsD7 | DownUp | notDM | Gain | 4 |
| **5_179084223_179086281_3187** | 5_179084223_179086281 | HNRNPH1 | 3187 | 4.59307551611353 | -0.68835531 | 1 | only_DIvsD7 | DownUp | notDM | Loss | 7 |
| **5_179084223_179086281_9794** | 5_179084223_179086281 | MAML1 | 9794 | 4.8667066591488 | -0.57763628 | 1 | only_DIvsD7 | UpDown | notDM | Loss | 7 |
| **5_179103867_179105531_3187** | 5_179103867_179105531 | HNRNPH1 | 3187 | 4.71823588242831 | 0.71090092 | 1 | only_DIvsD7 | DownUp | notDM | Gain | 5 |
| **5_179103867_179105531_9794** | 5_179103867_179105531 | MAML1 | 9794 | 4.73508766422784 | 0.53948884 | 1 | only_DIvsD7 | UpDown | notDM | Gain | 1 |
| **5_179105715_179106588_3187** | 5_179105715_179106588 | HNRNPH1 | 3187 | 4.73332568810799 | 0.61195575 | 2 | only_DIvsD7 | DownUp | notDM | Gain | 1 |
| **5_179157433_179159562_9794** | 5_179157433_179159562 | MAML1 | 9794 | 2.48429983934679 | 0.16835431 | 4 | only_DIvsD7 | UpDown | notDM | Loss | 6 |
| **5_179160432_179162069_9794** | 5_179160432_179162069 | MAML1 | 9794 | 2.75434833571102 | 0.16835431 | 1 | only_DIvsD7 | UpDown | notDM | Loss | 6 |
| **5_179230701_179232969_3187** | 5_179230701_179232969 | HNRNPH1 | 3187 | 5.25310043556491 | 0.74076774 | 1 | only_DIvsD7 | DownUp | notDM | Loss | 6 |
| **5_179230701_179232969_4056** | 5_179230701_179232969 | LTC4S | 4056 | 3.98748750613309 | 0.08383918 | 1 | only_H3vsUI | UpDown | notDM | Loss | 6 |
| **5_179234224_179236206_9794** | 5_179234224_179236206 | MAML1 | 9794 | 4.87133938109986 | 0.59196308 | 5 | only_DIvsD7 | UpDown | notDM | Gain | 1 |
| **5_179244517_179246096_3187** | 5_179244517_179246096 | HNRNPH1 | 3187 | 5.28746580534323 | 0.80516561 | 7 | only_DIvsD7 | DownUp | notDM | Gain | 5 |
| **5_179246376_179247800_3187** | 5_179246376_179247800 | HNRNPH1 | 3187 | 5.29161079786272 | 0.80987625 | 7 | only_DIvsD7 | DownUp | notDM | Gain | 5 |
| **5_179248619_179256536_11282** | 5_179248619_179256536 | MGAT4B | 11282 | 4.16693309487115 | 0.30325895 | 1 | notSpecific | Down | DM | Loss | 10 |
| **5_179248619_179256536_3187** | 5_179248619_179256536 | HNRNPH1 | 3187 | 5.29655989398707 | 0.79463807 | 1 | only_DIvsD7 | DownUp | DM | Loss | 10 |
| **5_179248619_179256536_51149** | 5_179248619_179256536 | MRNIP | 51149 | 4.46660073629688 | -0.40305965 | 1 | notSpecific | UpDown | DM | Loss | 10 |
| **5_179248619_179256536_80230** | 5_179248619_179256536 | RUFY1 | 80230 | 5.41810067387471 | -0.74132778 | 1 | notSpecific | Down | DM | Loss | 10 |
| **5_179248619_179256536_8878** | 5_179248619_179256536 | SQSTM1 | 8878 | 2.85491302230786 | -0.04027554 | 1 | notSpecific | Up | DM | Loss | 10 |
| **5_216634_218168_116349** | 5_216634_218168 | EXOC3-AS1 | 116349 | 5.35235813059147 | 0.75214327 | 1 | only_H3vsUI | UpDown | notDM | Loss | 6 |
| **5_216634_218168_6389** | 5_216634_218168 | SDHA | 6389 | 2.36921585741014 | 0.6968355 | 1 | only_H3vsUI | DownUp | notDM | Loss | 6 |

| | | | | | | | | | | | |
|---|---|---|---|---|---|---|---|---|---|---|---|
| **5_218829_220930_6389** | 5_218829_220930 | SDHA | 6389 | 2.63346845557959 | 0.59451974 | 1 | only_H3vsUI | DownUp | notDM | Loss | 6 |
| **5_270019_271415_6389** | 5_270019_271415 | SDHA | 6389 | 4.71281800020785 | 0.68106139 | 1 | only_H3vsUI | DownUp | notDM | Loss | 6 |
| **5_272198_274565_6389** | 5_272198_274565 | SDHA | 6389 | 4.73051580512519 | 0.6267395 | 2 | only_H3vsUI | DownUp | notDM | Loss | 6 |
| **5_31530243_31532047_29102** | 5_31530243_31532047 | DROSHA | 29102 | 2.13033376849501 | 0.05073278 | 1 | only_H3vsUI | Down | notDM | Loss | 8 |
| **5_31532652_31534807_29102** | 5_31532652_31534807 | DROSHA | 29102 | 2.54530711646582 | 0.03228577 | 1 | only_H3vsUI | Down | notDM | Loss | 8 |
| **5_31638304_31639295_23037** | 5_31638304_31639295 | PDZD2 | 23037 | 2.34830486304816 | 0 | 2 | only_H3vsUI | UpDown | notDM | Loss | 3 |
| **5_32059243_32060552_23037** | 5_32059243_32060552 | PDZD2 | 23037 | 0 | 0 | 6 | only_H3vsUI | UpDown | notDM | Gain | 4 |
| **5_441889_443112_116349** | 5_441889_443112 | EXOC3-AS1 | 116349 | 2.16731733474818 | 0.78720504 | 2 | only_H3vsUI | UpDown | notDM | Loss | 6 |
| **5_441889_443112_6389** | 5_441889_443112 | SDHA | 6389 | 5.34919590741286 | 0.72388193 | 2 | only_H3vsUI | DownUp | notDM | Loss | 3 |
| **5_443564_445028_116349** | 5_443564_445028 | EXOC3-AS1 | 116349 | 2.48855071650044 | 0.78914827 | 1 | only_H3vsUI | UpDown | notDM | Loss | 6 |
| **5_443564_445028_6389** | 5_443564_445028 | SDHA | 6389 | 5.35243915856107 | 0.7260834 | 1 | only_H3vsUI | DownUp | notDM | Loss | 6 |
| **5_5135202_5137382_101929176** | 5_5135202_5137382 | CTD-2297D10.2 | 101929176 | 3.44498111208794 | 0.1792432 | 6 | only_H3vsUI | UpDown | notDM | Loss | 3 |
| **5_5139577_5141948_101929176** | 5_5139577_5141948 | CTD-2297D10.2 | 101929176 | 0 | 0.60786374 | 4 | only_H3vsUI | UpDown | notDM | Loss | 3 |
| **5_5177446_5180357_101929176** | 5_5177446_5180357 | CTD-2297D10.2 | 101929176 | 4.5714875530558 | 0.61012036 | 6 | only_H3vsUI | UpDown | notDM | Loss | 5 |
| **5_52314662_52315843_3673** | 5_52314662_52315843 | ITGA2 | 3673 | 4.46993977478634 | 0.8578659 | 6 | notSpecific | Down | DM | Gain | 9 |
| **5_52314662_52315843_4338** | 5_52314662_52315843 | MOCS2 | 4338 | 4.95292314746539 | 0.77831711 | 6 | notSpecific | Down | DM | Gain | 9 |
| **5_52403877_52405400_257396** | 5_52403877_52405400 | LOC257396 | 257396 | 2.43616264704076 | -0.15565913 | 2 | only_H3vsUI | Down | notDM | Gain | 4 |
| **5_52405743_52407062_257396** | 5_52405743_52407062 | LOC257396 | 257396 | 1.86332286012046 | -0.16528709 | 1 | only_H3vsUI | Down | notDM | Loss | 8 |
| **5_53812522_53813403_11082** | 5_53812522_53813403 | ESM1 | 11082 | 5.67032843541193 | 0.99939563 | 4 | only_H3vsUI | UpDown | notDM | Gain | 1 |
| **5_54450290_54451305_11082** | 5_54450290_54451305 | ESM1 | 11082 | 5.22737501511662 | 0.99992864 | 6 | only_H3vsUI | UpDown | notDM | Loss | 3 |
| **5_55759795_55762654_102467147** | 5_55759795_55762654 | LINC01948 | 102467147 | 4.17443779645205 | 0.19477626 | 3 | only_DIvsD7 | UpDown | notDM | Loss | 5 |
| **5_55770931_55773744_102467147** | 5_55770931_55773744 | LINC01948 | 102467147 | 3.585799009013 | 0.19439614 | 6 | only_DIvsD7 | UpDown | notDM | Loss | 5 |
| **5_55776964_55778319_102467147** | 5_55776964_55778319 | LINC01948 | 102467147 | 0 | 0.14258284 | 2 | only_DIvsD7 | UpDown | notDM | Loss | 3 |
| **5_58845008_58846528_5144** | 5_58845008_58846528 | PDE4D | 5144 | 5.33832698282188 | 0.81396874 | 7 | notSpecific | Down | DM | Gain | 9 |
| **5_60239130_60240503_1161** | 5_60239130_60240503 | ERCC8 | 1161 | 2.58319877396862 | -0.83780209 | 1 | only_H3vsUI | Down | notDM | Loss | 8 |
| **5_60241128_60243065_1161** | 5_60241128_60243065 | ERCC8 | 1161 | 2.38916608436453 | -0.725104 | 2 | only_H3vsUI | Down | notDM | Loss | 8 |
| **5_60594969_60604004_1161** | 5_60594969_60604004 | ERCC8 | 1161 | 5.54910875578114 | -0.92564239 | 5 | only_H3vsUI | Down | notDM | Loss | 7 |
| **5_60606715_60611315_1161** | 5_60606715_60611315 | ERCC8 | 1161 | 5.56326388384855 | -0.80150828 | 6 | only_H3vsUI | Down | notDM | Loss | 2 |
| **5_60623177_60626648_1161** | 5_60623177_60626648 | ERCC8 | 1161 | 5.5823804412714 | -0.81672416 | 1 | only_H3vsUI | Down | notDM | Loss | 2 |
| **5_60629342_60631927_1161** | 5_60629342_60631927 | ERCC8 | 1161 | 5.58932841761832 | -0.78414931 | 1 | only_H3vsUI | Down | notDM | Loss | 2 |
| **5_63801666_63803370_401190** | 5_63801666_63803370 | RGS7BP | 401190 | 0 | 0.08918417 | 4 | only_H3vsUI | Up | notDM | Loss | 3 |
| **5_6631120_6632827_6715** | 5_6631120_6632827 | SRD5A1 | 6715 | 2.85369821177617 | -0.19644221 | 1 | only_DIvsD7 | Down | notDM | Gain | 4 |
| **5_68461875_68462783_92259** | 5_68461875_68462783 | MRPS36 | 92259 | 4.7069907304057 | 0.72122609 | 5 | only_H3vsUI | UpDown | notDM | Gain | 1 |
| **5_68463057_68466762_92259** | 5_68463057_68466762 | MRPS36 | 92259 | 4.6716633461617 | 0.74785198 | 1 | only_H3vsUI | UpDown | notDM | Loss | 6 |

| | | | | | | | | | | | |
|---|---|---|---|---|---|---|---|---|---|---|---|
| **5_68483778_68485250_92259** | 5_68483778_68485250 | MRPS36 | 92259 | 4.45347123372294 | 0.35480516 | 1 | only_H3vsUI | UpDown | notDM | Loss | 6 |
| **5_68485471_68486914_92259** | 5_68485471_68486914 | MRPS36 | 92259 | 4.42725884027041 | 0.35480516 | 1 | only_H3vsUI | UpDown | notDM | Gain | 4 |
| **5_68512281_68513457_92259** | 5_68512281_68513457 | MRPS36 | 92259 | 2.41161970596323 | 0.88794474 | 2 | only_H3vsUI | UpDown | notDM | Gain | 4 |
| **5_68514018_68515495_92259** | 5_68514018_68515495 | MRPS36 | 92259 | 2.48572142648158 | 0.88794474 | 2 | only_H3vsUI | UpDown | notDM | Gain | 4 |
| **5_68529494_68530534_92259** | 5_68529494_68530534 | MRPS36 | 92259 | 4.19816203904608 | 0.47637237 | 1 | only_H3vsUI | UpDown | notDM | Gain | 4 |
| **5_68530829_68536389_92259** | 5_68530829_68536389 | MRPS36 | 92259 | 4.2348208798074 | 0.70079998 | 1 | only_H3vsUI | UpDown | notDM | Gain | 4 |
| **5_692388_693353_11076** | 5_692388_693353 | TPPP | 11076 | 2.19865708695442 | 0.02419227 | 2 | only_DIvsD7 | UpDown | notDM | Loss | 3 |
| **5_692388_693353_6389** | 5_692388_693353 | SDHA | 6389 | 5.67573894186432 | -0.91571473 | 2 | only_H3vsUI | DownUp | notDM | Loss | 7 |
| **5_693621_694614_11076** | 5_693621_694614 | TPPP | 11076 | 2.05307844348342 | 0.01154731 | 2 | only_DIvsD7 | UpDown | notDM | Loss | 3 |
| **5_72416622_72417818_134285** | 5_72416622_72417818 | TMEM171 | 134285 | 2.35793484700045 | 0.68641029 | 1 | only_H3vsUI | Down | notDM | Loss | 8 |
| **5_72899966_72904037_134285** | 5_72899966_72904037 | TMEM171 | 134285 | 5.6844611462054 | 0.80850272 | 7 | only_H3vsUI | Down | notDM | Gain | 5 |
| **5_74905582_74908070_728780** | 5_74905582_74908070 | ANKDD1B | 728780 | 0 | 0.00039552 | 1 | only_DIvsD7 | DownUp | notDM | Gain | 4 |
| **5_76031660_76032545_2149** | 5_76031660_76032545 | F2R | 2149 | 4.29653356592899 | 0.7753206 | 6 | notSpecific | UpDown | DM | Gain | 9 |
| **5_76031660_76032545_2151** | 5_76031660_76032545 | F2RL2 | 2151 | 5.05077790256809 | 0.78073538 | 6 | notSpecific | UpDown | DM | Gain | 9 |
| **5_77716152_77717086_9522** | 5_77716152_77717086 | SCAMP1 | 9522 | 3.57135939275384 | -0.00171206 | 6 | notSpecific | DownUp | DM | Gain | 9 |
| **5_79949084_79950215_4437** | 5_79949084_79950215 | MSH3 | 4437 | 2.87909587950007 | 0.75316797 | 2 | only_H3vsUI | DownUp | notDM | Loss | 6 |
| **5_79950992_79952390_4437** | 5_79950992_79952390 | MSH3 | 4437 | 1.36172783601759 | 0.74468546 | 1 | only_H3vsUI | DownUp | notDM | Loss | 6 |
| **5_80256644_80258031_4437** | 5_80256644_80258031 | MSH3 | 4437 | 5.48597540028492 | -0.86025623 | 2 | only_H3vsUI | DownUp | notDM | Loss | 7 |
| **5_80595032_80597265_100131067** | 5_80595032_80597265 | CKMT2-AS1 | 100131067 | 1.60205999132796 | -0.56542692 | 1 | only_H3vsUI | UpDown | notDM | Loss | 8 |
| **5_80597698_80599989_100131067** | 5_80597698_80599989 | CKMT2-AS1 | 100131067 | 2.59769518592551 | -0.56542692 | 2 | only_H3vsUI | UpDown | notDM | Gain | 4 |
| **5_80746492_80747737_23635** | 5_80746492_80747737 | SSBP2 | 23635 | 5.47596594372756 | -0.84289667 | 6 | only_H3vsUI | DownUp | notDM | Gain | 2 |
| **5_81045168_81045872_23635** | 5_81045168_81045872 | SSBP2 | 23635 | 2.98677173426624 | 0.68776973 | 2 | only_H3vsUI | DownUp | notDM | Loss | 3 |
| **5_81047335_81048491_23635** | 5_81047335_81048491 | SSBP2 | 23635 | 2.67209785793572 | 0.9371174 | 5 | only_H3vsUI | DownUp | notDM | Loss | 6 |
| **5_81050462_81052589_23635** | 5_81050462_81052589 | SSBP2 | 23635 | 3.55606115900953 | 0.75721805 | 6 | only_H3vsUI | DownUp | notDM | Gain | 1 |
| **5_81064117_81065771_23635** | 5_81064117_81065771 | SSBP2 | 23635 | 4.23172438332852 | -0.16902911 | 3 | only_H3vsUI | DownUp | notDM | Gain | 1 |
| **5_81146122_81148031_23635** | 5_81146122_81148031 | SSBP2 | 23635 | 4.99676552012874 | 0.84975718 | 3 | only_H3vsUI | DownUp | notDM | Gain | 1 |
| **5_90672389_90675819_57561** | 5_90672389_90675819 | ARRDC3 | 57561 | 3.52608069180203 | 0.45142088 | 1 | notSpecific | Down | DM | Gain | 9 |
| **5_93951910_93954182_84250** | 5_93951910_93954182 | SLF1 | 84250 | 2.32014628611105 | -0.20830264 | 1 | only_H3vsUI | Down | notDM | Gain | 4 |
| **5_93954715_93956945_84250** | 5_93954715_93956945 | SLF1 | 84250 | 2.51188336097887 | -0.350626 | 2 | only_H3vsUI | Down | notDM | Loss | 8 |
| **5_95589942_95591854_831** | 5_95589942_95591854 | CAST | 831 | 5.6088190300108 | 0.83651086 | 7 | notSpecific | Down | DM | Gain | 9 |
| **6_105401735_105405089_389421** | 6_105401735_105405089 | LIN28B | 389421 | 0 | 0.97479424 | 5 | only_DIvsD7 | Down | notDM | Gain | 4 |
| **6_105406831_105408421_389421** | 6_105406831_105408421 | LIN28B | 389421 | 3.28103336724773 | 0.97219663 | 2 | only_DIvsD7 | Down | notDM | Gain | 4 |
| **6_107436904_107438110_57673** | 6_107436904_107438110 | BEND3 | 57673 | 2.63648789635337 | 0.01837766 | 1 | only_H3vsUI | Down | notDM | Loss | 8 |
| **6_108438829_108439575_10028736** | 6_108438829_108439575 | OSTM1-AS1 | 100287366 | 3.71138537909845 | 0 | 2 | notSpecific | UpDown | DM | Loss | 10 |

| | | | | | | | | | | | |
|---|---|---|---|---|---|---|---|---|---|---|---|
| **6_108438829_108439575_8724** | 6_108438829_108439575 | SNX3 | 8724 | 5.1541012518713 | 0.7545072 | 2 | notSpecific | DownUp | DM | Loss | 10 |
| **6_117811044_117820555_57120** | 6_117811044_117820555 | GOPC | 57120 | 5.01337238415717 | 0.90030198 | 7 | only_H3vsUI | Down | notDM | Gain | 5 |
| **6_117824890_117827775_57120** | 6_117824890_117827775 | GOPC | 57120 | 4.98185030634092 | 0.61896803 | 7 | only_H3vsUI | Down | notDM | Loss | 5 |
| **6_117869387_117872382_57120** | 6_117869387_117872382 | GOPC | 57120 | 4.71011736511182 | 0.64309716 | 7 | only_H3vsUI | Down | notDM | Gain | 5 |
| **6_117919842_117923312_57120** | 6_117919842_117923312 | GOPC | 57120 | 2.57978359661681 | 0.76643688 | 1 | only_H3vsUI | Down | notDM | Gain | 4 |
| **6_117923880_117925132_57120** | 6_117923880_117925132 | GOPC | 57120 | 2.28103336724773 | 0.85524264 | 2 | only_H3vsUI | Down | notDM | Loss | 8 |
| **6_117997291_117998695_57120** | 6_117997291_117998695 | GOPC | 57120 | 4.86688961565764 | 0.88191137 | 2 | only_H3vsUI | Down | notDM | Gain | 4 |
| **6_126068053_126070562_135112** | 6_126068053_126070562 | NCOA7 | 135112 | 4.61857102812013 | 0.60063788 | 2 | only_H3vsUI | Up | notDM | Loss | 6 |
| **6_126070837_126071758_135112** | 6_126070837_126071758 | NCOA7 | 135112 | 4.60588658966101 | 0.58113759 | 2 | only_H3vsUI | Up | notDM | Loss | 3 |
| **6_126101780_126104110_135112** | 6_126101780_126104110 | NCOA7 | 135112 | 0 | 0.40217426 | 1 | only_H3vsUI | Up | notDM | Gain | 8 |
| **6_126104563_126106044_135112** | 6_126104563_126106044 | NCOA7 | 135112 | 3.35372393758895 | 0.6099631 | 7 | only_H3vsUI | Up | notDM | Gain | 1 |
| **6_126110361_126111767_135112** | 6_126110361_126111767 | NCOA7 | 135112 | 2.50514997831991 | 0.03573822 | 5 | only_H3vsUI | Up | notDM | Gain | 4 |
| **6_126112502_126116298_135112** | 6_126112502_126116298 | NCOA7 | 135112 | 2.59439255037543 | 0.29636571 | 1 | only_H3vsUI | Up | notDM | Loss | 6 |
| **6_126132816_126134691_135112** | 6_126132816_126134691 | NCOA7 | 135112 | 3.22375545365724 | 0.01796233 | 7 | only_H3vsUI | Up | notDM | Gain | 4 |
| **6_126135747_126136440_135112** | 6_126135747_126136440 | NCOA7 | 135112 | 0 | 0.03665174 | 8 | only_H3vsUI | Up | notDM | Gain | 1 |
| **6_126192161_126195255_135112** | 6_126192161_126195255 | NCOA7 | 135112 | 4.95344557213989 | 0.63800322 | 7 | only_H3vsUI | Up | notDM | Loss | 5 |
| **6_126219328_126224214_135112** | 6_126219328_126224214 | NCOA7 | 135112 | 0 | 0.30718268 | 7 | only_H3vsUI | Up | notDM | Gain | 4 |
| **6_126238459_126246973_135112** | 6_126238459_126246973 | NCOA7 | 135112 | 0 | 0.5423059 | 5 | only_H3vsUI | Up | notDM | Loss | 8 |
| **6_126261390_126262058_135112** | 6_126261390_126262058 | NCOA7 | 135112 | 4.32099741679422 | 0.45290235 | 7 | only_H3vsUI | Up | notDM | Loss | 1 |
| **6_126317382_126317876_135112** | 6_126317382_126317876 | NCOA7 | 135112 | 4.88611266806809 | 0.60452511 | 6 | only_H3vsUI | Up | notDM | Gain | 2 |
| **6_132128000_132128917_5167** | 6_132128000_132128917 | ENPP1 | 5167 | 2.38021124171161 | 0 | 2 | only_H3vsUI | UpDown | notDM | Loss | 3 |
| **6_132654494_132656570_8876** | 6_132654494_132656570 | VNN1,00 | 8876 | 5.57820240410245 | 0.99733539 | 3 | only_H3vsUI | UpDown | notDM | Gain | 1 |
| **6_132831791_132833738_8417** | 6_132831791_132833738 | STX7 | 8417 | 2.66651798055488 | 0.27732807 | 2 | only_DIvsD7 | UpDown | notDM | Gain | 4 |
| **6_133118547_133119657_116843** | 6_133118547_133119657 | SLC18B1 | 116843 | 0.698970004336019 | -0.88074445 | 2 | only_H3vsUI | Down | notDM | Loss | 8 |
| **6_133133512_133134930_116843** | 6_133133512_133134930 | SLC18B1 | 116843 | 4.14154383422065 | -0.70962922 | 2 | only_H3vsUI | Down | notDM | Gain | 4 |
| **6_133136122_133139845_116843** | 6_133136122_133139845 | SLC18B1 | 116843 | 4.2165089781866 | -0.71558596 | 1 | only_H3vsUI | Down | notDM | Loss | 8 |
| **6_133355258_133356703_8876** | 6_133355258_133356703 | VNN1,00 | 8876 | 5.5052476835869 | 0.99781659 | 6 | only_H3vsUI | UpDown | notDM | Loss | 3 |
| **6_139457080_139458066_51696** | 6_139457080_139458066 | HECA | 51696 | 2.92064500140679 | 0.18298854 | 2 | only_H3vsUI | Up | notDM | Loss | 3 |
| **6_143770313_143771621_8504** | 6_143770313_143771621 | PEX3 | 8504 | 2.51054501020661 | 0.24101106 | 2 | only_DIvsD7 | UpDown | notDM | Gain | 4 |
| **6_143772121_143773484_8504** | 6_143772121_143773484 | PEX3 | 8504 | 2.25285303097989 | 0.23933214 | 2 | only_DIvsD7 | UpDown | notDM | Gain | 4 |
| **6_145077580_145079408_7402** | 6_145077580_145079408 | UTRN | 7402 | 3.23854788768133 | -0.00831173 | 6 | notSpecific | Down | DM | Gain | 9 |
| **6_150208333_150210366_10065273** | 6_150208333_150210366 | RAET1E-AS1 | 100652739 | 2.9415114326344 | 0.0037337 | 7 | only_H3vsUI | Up | notDM | Gain | 1 |
| **6_150258853_150260655_10537805** | 6_150258853_150260655 | LOC105378052 | 105378052 | 3.50406288267869 | 0 | 4 | only_H3vsUI | Down | notDM | Loss | 3 |
| **6_153438505_153440895_26271** | 6_153438505_153440895 | FBXO5 | 26271 | 5.12825060081807 | -0.82209648 | 7 | notSpecific | DownUp | DM | Gain | 9 |

| | | | | | | | | | | | |
|---|---|---|---|---|---|---|---|---|---|---|---|
| **6_153438505_153440895_26575** | 6_153438505_153440895 | RGS17 | 26575 | 4.06032002868829 | -0.16215518 | 7 | notSpecific | Up | DM | Gain | 9 |
| **6_159270343_159272964_7430** | 6_159270343_159272964 | EZR | 7430 | 4.47659978839897 | 0.62941004 | 7 | notSpecific | DownUp | DM | Gain | 9 |
| **6_16238978_16239615_2766** | 6_16238978_16239615 | GMPR | 2766 | 2.22788670461367 | 0.34412614 | 4 | only_H3vsUI | UpDown | notDM | Loss | 3 |
| **6_170100381_170101661_387263** | 6_170100381_170101661 | C6orf120 | 387263 | 2.75815462196739 | -0.54817666 | 1 | only_H3vsUI | DownUp | notDM | Gain | 4 |
| **6_170102623_170104838_387263** | 6_170102623_170104838 | C6orf120 | 387263 | 2.59328606702046 | -0.54817666 | 1 | only_H3vsUI | DownUp | notDM | Loss | 8 |
| **6_170121352_170123596_387263** | 6_170121352_170123596 | C6orf120 | 387263 | 4.28151060149266 | -0.64832278 | 1 | only_H3vsUI | DownUp | notDM | Gain | 2 |
| **6_170125052_170125964_387263** | 6_170125052_170125964 | C6orf120 | 387263 | 4.35787769922987 | 0.57698075 | 1 | only_H3vsUI | DownUp | notDM | Loss | 7 |
| **6_170149825_170151365_387263** | 6_170149825_170151365 | C6orf120 | 387263 | 4.67755220626809 | -0.61373345 | 1 | only_H3vsUI | DownUp | notDM | Loss | 7 |
| **6_170152135_170153500_387263** | 6_170152135_170153500 | C6orf120 | 387263 | 4.69813535741307 | -0.61373345 | 1 | only_H3vsUI | DownUp | notDM | Gain | 2 |
| **6_170199838_170201527_387263** | 6_170199838_170201527 | C6orf120 | 387263 | 4.98937416552269 | 0.60303943 | 3 | only_H3vsUI | DownUp | notDM | Loss | 5 |
| **6_170604346_170605388_28514** | 6_170604346_170605388 | DLL1 | 28514 | 3.68006342748195 | 0.01775839 | 4 | notSpecific | UpDown | DM | Loss | 10 |
| **6_170604346_170605388_84498** | 6_170604346_170605388 | FAM120B | 84498 | 0 | -0.00202071 | 4 | notSpecific | Down | DM | Loss | 10 |
| **6_18386834_18389891_255488** | 6_18386834_18389891 | RNF144B | 255488 | 0 | 0.89017652 | 2 | only_H3vsUI | Down | notDM | Gain | 4 |
| **6_26027576_26031250_10475** | 6_26027576_26031250 | TRIM38 | 10475 | 4.80988279032947 | 0.78233016 | 5 | notSpecific | DownUp | DM | Gain | 9 |
| **6_26027576_26031250_3006** | 6_26027576_26031250 | H1-2 | 3006 | 4.40568778667278 | 0.32579178 | 5 | notSpecific | Down | DM | Gain | 9 |
| **6_28104826_28106156_9753** | 6_28104826_28106156 | ZSCAN12 | 9753 | 5.41722748961061 | 0.92529612 | 1 | only_H3vsUI | Down | notDM | Loss | 7 |
| **6_28218541_28220006_387032** | 6_28218541_28220006 | ZKSCAN4 | 387032 | 0 | 0.18695225 | 2 | only_DIvsD7 | Down | notDM | Loss | 3 |
| **6_28220165_28220693_387032** | 6_28220165_28220693 | ZKSCAN4 | 387032 | 2.21748394421391 | 0.17560821 | 2 | only_DIvsD7 | Down | notDM | Gain | 2 |
| **6_28233914_28234887_387032** | 6_28233914_28234887 | ZKSCAN4 | 387032 | 4.14345199902163 | -0.16917179 | 2 | only_DIvsD7 | Down | notDM | Gain | 4 |
| **6_28235131_28236533_387032** | 6_28235131_28236533 | ZKSCAN4 | 387032 | 4.17986763127041 | -0.16917179 | 2 | only_DIvsD7 | Down | notDM | Loss | 3 |
| **6_28367620_28368555_9753** | 6_28367620_28368555 | ZSCAN12 | 9753 | 2.05690485133647 | 0.44665697 | 2 | only_H3vsUI | Down | notDM | Gain | 4 |
| **6_28651354_28652145_9753** | 6_28651354_28652145 | ZSCAN12 | 9753 | 5.45308124842391 | -0.86247246 | 4 | only_H3vsUI | Down | notDM | Gain | 2 |
| **6_28697131_28698584_5987** | 6_28697131_28698584 | TRIM27 | 5987 | 5.28596665797566 | 0.7140209 | 2 | only_H3vsUI | Down | notDM | Gain | 1 |
| **6_28723859_28726145_5987** | 6_28723859_28726145 | TRIM27 | 5987 | 5.21911540253278 | 0.69726909 | 1 | only_H3vsUI | Down | notDM | Gain | 4 |
| **6_28777220_28780336_5987** | 6_28777220_28780336 | TRIM27 | 5987 | 5.04700602796563 | 0.62705835 | 4 | only_H3vsUI | Down | notDM | Gain | 4 |
| **6_28784781_28786547_5987** | 6_28784781_28786547 | TRIM27 | 5987 | 5.02209829746114 | 0.63184312 | 1 | only_H3vsUI | Down | notDM | Loss | 7 |
| **6_28804688_28805734_5987** | 6_28804688_28805734 | TRIM27 | 5987 | 4.93466001913304 | 0.70067002 | 2 | only_H3vsUI | Down | notDM | Gain | 1 |
| **6_28830753_28831420_5987** | 6_28830753_28831420 | TRIM27 | 5987 | 4.78064848842674 | 0.55814694 | 4 | only_H3vsUI | Down | notDM | Gain | 1 |
| **6_28844841_28846202_5987** | 6_28844841_28846202 | TRIM27 | 5987 | 4.65863137460951 | 0.51513271 | 4 | only_H3vsUI | Down | notDM | Gain | 1 |
| **6_28864221_28866037_414760** | 6_28864221_28866037 | HCG14 | 414760 | 0 | 0.00284313 | 1 | notSpecific | Up | DM | Gain | 10 |
| **6_28889260_28891013_5987** | 6_28889260_28891013 | TRIM27 | 5987 | 2.8767949762007 | 0.4021631 | 1 | only_H3vsUI | Down | notDM | Gain | 4 |
| **6_28891884_28892996_5987** | 6_28891884_28892996 | TRIM27 | 5987 | 2.08278537031645 | 0.04635795 | 1 | only_H3vsUI | Down | notDM | Gain | 4 |
| **6_28907505_28908728_5987** | 6_28907505_28908728 | TRIM27 | 5987 | 4.197032318918 | -0.33675983 | 2 | only_H3vsUI | Down | notDM | Loss | 7 |
| **6_28908938_28911397_5987** | 6_28908938_28911397 | TRIM27 | 5987 | 4.2348714585627 | -0.32476876 | 2 | only_H3vsUI | Down | notDM | Gain | 4 |

| | | | | | | | | | | | |
|---|---|---|---|---|---|---|---|---|---|---|---|
| **6_30639175_30640486_9656** | 6_30639175_30640486 | MDC1 | 9656 | 4.6490718653987 | -0.54168429 | 1 | only_H3vsUI | DownUp | notDM | Loss | 7 |
| **6_30655260_30656521_9656** | 6_30655260_30656521 | MDC1 | 9656 | 4.45455523380111 | 0.33966696 | 5 | only_H3vsUI | DownUp | notDM | Gain | 1 |
| **6_30658923_30659674_9656** | 6_30658923_30659674 | MDC1 | 9656 | 4.40360089751828 | 0.29112778 | 2 | only_H3vsUI | DownUp | notDM | Gain | 1 |
| **6_30677088_30678199_9656** | 6_30677088_30678199 | MDC1 | 9656 | 3.83632411570675 | 0.62222968 | 7 | only_H3vsUI | DownUp | notDM | Gain | 1 |
| **6_30683969_30685037_9656** | 6_30683969_30685037 | MDC1 | 9656 | 1.34242268082221 | 0.55448802 | 1 | only_H3vsUI | DownUp | notDM | Loss | 6 |
| **6_30686894_30687892_9656** | 6_30686894_30687892 | MDC1 | 9656 | 3.26434550705009 | 0.55448802 | 5 | only_H3vsUI | DownUp | notDM | Gain | 4 |
| **6_30689581_30690552_9656** | 6_30689581_30690552 | MDC1 | 9656 | 3.65561858354122 | 0.55448802 | 1 | only_H3vsUI | DownUp | notDM | Loss | 7 |
| **6_30714793_30720106_9656** | 6_30714793_30720106 | MDC1 | 9656 | 4.47329715284889 | -0.35450492 | 7 | only_H3vsUI | DownUp | notDM | Gain | 5 |
| **6_30726613_30727899_9656** | 6_30726613_30727899 | MDC1 | 9656 | 4.6186441882963 | -0.90002696 | 7 | only_H3vsUI | DownUp | notDM | Gain | 2 |
| **6_30730751_30733242_9656** | 6_30730751_30733242 | MDC1 | 9656 | 4.65986868166268 | -0.57895942 | 7 | only_H3vsUI | DownUp | notDM | Gain | 5 |
| **6_31081591_31083893_29113** | 6_31081591_31083893 | C6orf15 | 29113 | 3.09933527768596 | 0.03634069 | 5 | only_H3vsUI | UpDown | notDM | Gain | 1 |
| **6_31548798_31550028_629** | 6_31548798_31550028 | CFB | 629 | 5.56722051947358 | -0.84814513 | 2 | only_H3vsUI | DownUp | notDM | Loss | 7 |
| **6_31587200_31587773_629** | 6_31587200_31587773 | CFB | 629 | 5.52037871297756 | 0.82085753 | 1 | only_H3vsUI | DownUp | notDM | Loss | 3 |
| **6_31618606_31619919_55937** | 6_31618606_31619919 | APOM | 55937 | 2.44090908206522 | 0.08904236 | 1 | only_H3vsUI | Up | notDM | Loss | 3 |
| **6_31620580_31621813_55937** | 6_31620580_31621813 | APOM | 55937 | 2.58883172559421 | 0.08904236 | 1 | only_H3vsUI | Up | notDM | Loss | 3 |
| **6_31626737_31628530_55937** | 6_31626737_31628530 | APOM | 55937 | 3.81590965088677 | 0.17137774 | 1 | only_H3vsUI | Up | notDM | Loss | 3 |
| **6_31631379_31633267_55937** | 6_31631379_31633267 | APOM | 55937 | 4.04871363808097 | 0.24200129 | 1 | only_H3vsUI | Up | notDM | Loss | 6 |
| **6_31634090_31635384_55937** | 6_31634090_31635384 | APOM | 55937 | 4.14295230734343 | 0.24200129 | 2 | only_H3vsUI | Up | notDM | Loss | 3 |
| **6_31829585_31830588_629** | 6_31829585_31830588 | CFB | 629 | 4.81216438839285 | 0.54547622 | 2 | only_H3vsUI | DownUp | notDM | Loss | 3 |
| **6_31870239_31871080_629** | 6_31870239_31871080 | CFB | 629 | 4.38731862468566 | 0.35443807 | 5 | only_H3vsUI | DownUp | notDM | Gain | 4 |
| **6_31925153_31926542_629** | 6_31925153_31926542 | CFB | 629 | 3.77546480934574 | 0.43369693 | 1 | only_H3vsUI | DownUp | notDM | Loss | 6 |
| **6_31927020_31927875_629** | 6_31927020_31927875 | CFB | 629 | 3.89376176205794 | 0.43369693 | 2 | only_H3vsUI | DownUp | notDM | Loss | 3 |
| **6_31940398_31941518_629** | 6_31940398_31941518 | CFB | 629 | 4.32649971472504 | 0.23789191 | 1 | only_H3vsUI | DownUp | notDM | Loss | 3 |
| **6_32094633_32095840_629** | 6_32094633_32095840 | CFB | 629 | 5.2441360449822 | 0.71540336 | 1 | only_H3vsUI | DownUp | notDM | Loss | 3 |
| **6_32096280_32097876_629** | 6_32096280_32097876 | CFB | 629 | 5.24819403794086 | 0.79113385 | 1 | only_H3vsUI | DownUp | notDM | Loss | 6 |
| **6_32156133_32157668_4855** | 6_32156133_32157668 | NOTCH4 | 4855 | 4.5337339384132 | 0.45672535 | 1 | only_H3vsUI | UpDown | notDM | Loss | 3 |
| **6_32162525_32163340_4855** | 6_32162525_32163340 | NOTCH4 | 4855 | 3.30449052777349 | -0.06852752 | 1 | only_H3vsUI | UpDown | notDM | Gain | 1 |
| **6_32163810_32165275_4855** | 6_32163810_32165275 | NOTCH4 | 4855 | 1.90848501887865 | -0.06852752 | 1 | only_H3vsUI | UpDown | notDM | Loss | 3 |
| **6_32340715_32342685_4855** | 6_32340715_32342685 | NOTCH4 | 4855 | 5.17281594010092 | 0.99962758 | 6 | only_H3vsUI | UpDown | notDM | Gain | 1 |
| **6_32805197_32806043_3112** | 6_32805197_32806043 | HLA-DOB | 3112 | 4.30907630182233 | 0.75058629 | 2 | only_DIvsD7 | DownUp | notDM | Loss | 3 |
| **6_32810881_32812581_3112** | 6_32810881_32812581 | HLA-DOB | 3112 | 4.41594107974587 | 0.46107298 | 2 | only_DIvsD7 | DownUp | notDM | Gain | 3 |
| **6_32819864_32821410_3112** | 6_32819864_32821410 | HLA-DOB | 3112 | 4.5445764915681 | 0.71767497 | 1 | only_DIvsD7 | DownUp | notDM | Loss | 3 |
| **6_32822043_32823029_3112** | 6_32822043_32823029 | HLA-DOB | 3112 | 4.57077636879475 | 0.63299938 | 1 | only_DIvsD7 | DownUp | notDM | Loss | 3 |
| **6_32887610_32890692_3112** | 6_32887610_32890692 | HLA-DOB | 3112 | 5.01193819067795 | 0.81258455 | 7 | only_DIvsD7 | DownUp | notDM | Gain | 1 |

| | | | | | | | | | | | |
|---|---|---|---|---|---|---|---|---|---|---|---|
| **6_33240005_33244891_6222** | 6_33240005_33244891 | RPS18 | 6222 | 2.34242268082221 | -0.08502948 | 1 | only_DIvsD7 | DownUp | notDM | Loss | 8 |
| **6_33255774_33256766_6222** | 6_33255774_33256766 | RPS18 | 6222 | 4.20205216891206 | 0.34189347 | 1 | only_DIvsD7 | DownUp | notDM | Loss | 7 |
| **6_33257393_33258561_6222** | 6_33257393_33258561 | RPS18 | 6222 | 4.24410386337651 | 0.34189347 | 2 | only_DIvsD7 | DownUp | notDM | Loss | 7 |
| **6_33392248_33393451_6222** | 6_33392248_33393451 | RPS18 | 6222 | 5.18316446108945 | 0.72506355 | 5 | only_DIvsD7 | DownUp | notDM | Loss | 5 |
| **6_33394940_33395607_6222** | 6_33394940_33395607 | RPS18 | 6222 | 5.19076577564961 | 0.72024071 | 1 | only_DIvsD7 | DownUp | notDM | Loss | 3 |
| **6_33398441_33399724_6222** | 6_33398441_33399724 | RPS18 | 6222 | 5.20045650075171 | 0.806047 | 4 | only_DIvsD7 | DownUp | notDM | Loss | 5 |
| **6_34192040_34192908_221491** | 6_34192040_34192908 | SMIM29 | 221491 | 4.37821614974988 | 0.84737 | 1 | only_DIvsD7 | DownUp | notDM | Loss | 3 |
| **6_34203476_34204539_221491** | 6_34203476_34204539 | SMIM29 | 221491 | 4.08969320878484 | 0.43452981 | 5 | only_DIvsD7 | DownUp | notDM | Gain | 1 |
| **6_34206060_34208523_221491** | 6_34206060_34208523 | SMIM29 | 221491 | 3.91613798310718 | -0.05033199 | 5 | only_DIvsD7 | DownUp | notDM | Loss | 6 |
| **6_34209636_34212335_221491** | 6_34209636_34212335 | SMIM29 | 221491 | 3.64659975172037 | 0.3927467 | 7 | only_DIvsD7 | DownUp | notDM | Loss | 5 |
| **6_34215403_34216742_221491** | 6_34215403_34216742 | SMIM29 | 221491 | 1.7481880270062 | 0.73222205 | 5 | only_DIvsD7 | DownUp | notDM | Gain | 4 |
| **6_4125218_4127406_404220** | 6_4125218_4127406 | C6orf201 | 404220 | 3.4848690327204 | 0 | 3 | only_H3vsUI | UpDown | notDM | Gain | 1 |
| **6_41861957_41862629_25862** | 6_41861957_41862629 | USP49 | 25862 | 0 | 0.41070206 | 2 | only_H3vsUI | Down | notDM | Loss | 3 |
| **6_41863411_41864569_25862** | 6_41863411_41864569 | USP49 | 25862 | 2.49692964807321 | 0.10323182 | 2 | only_H3vsUI | Down | notDM | Loss | 3 |
| **6_41885044_41886377_25862** | 6_41885044_41886377 | USP49 | 25862 | 4.35894321963233 | -0.27494447 | 7 | only_H3vsUI | Down | notDM | Gain | 1 |
| **6_41886913_41888601_25862** | 6_41886913_41888601 | USP49 | 25862 | 4.39308360208946 | -0.41891093 | 1 | only_H3vsUI | Down | notDM | Loss | 8 |
| **6_41889621_41891753_25862** | 6_41889621_41891753 | USP49 | 25862 | 4.43822580760453 | -0.41891093 | 2 | only_H3vsUI | Down | notDM | Gain | 4 |
| **6_47755507_47755962_221391** | 6_47755507_47755962 | OPN5 | 221391 | 2.8481891169914 | 0 | 4 | only_DIvsD7 | UpDown | notDM | Gain | 1 |
| **6_5004491_5005103_10799** | 6_5004491_5005103 | RPP40 | 10799 | 2.34830486304816 | 0.50490431 | 5 | only_H3vsUI | DownUp | notDM | Gain | 4 |
| **6_52589589_52591081_2938** | 6_52589589_52591081 | GSTA1 | 2938 | 4.88973857907739 | 0.86587056 | 7 | only_H3vsUI | UpDown | notDM | Gain | 1 |
| **6_55766281_55768552_653** | 6_55766281_55768552 | BMP5 | 653 | 4.4136517520637 | 0.99832626 | 7 | notSpecific | DownUp | DM | Gain | 9 |
| **6_56816954_56819314_221336** | 6_56816954_56819314 | BEND6 | 221336 | 2.78604121024255 | -0.01466413 | 1 | only_H3vsUI | DownUp | notDM | Loss | 6 |
| **6_56820263_56821928_221336** | 6_56820263_56821928 | BEND6 | 221336 | 2.24303804868629 | -0.00814429 | 2 | only_H3vsUI | DownUp | notDM | Loss | 3 |
| **6_7388476_7389760_83732** | 6_7388476_7389760 | RIOK1 | 83732 | 0 | 0.13518422 | 1 | only_H3vsUI | DownUp | notDM | Loss | 8 |
| **6_7390275_7392147_83732** | 6_7390275_7392147 | RIOK1 | 83732 | 2.73878055848437 | 0.13518422 | 1 | only_H3vsUI | DownUp | notDM | Loss | 8 |
| **6_73972996_73973740_80759** | 6_73972996_73973740 | KHDC1 | 80759 | 2.01283722470517 | 0.01567433 | 2 | notSpecific | Down | DM | Loss | 10 |
| **6_74753880_74754973_26503** | 6_74753880_74754973 | SLC17A5 | 26503 | 5.59127168271144 | 0.7842717 | 6 | notSpecific | DownUp | DM | Gain | 9 |
| **6_8102990_8103945_9521** | 6_8102990_8103945 | EEF1E1 | 9521 | 2.44404479591808 | 0.27760698 | 4 | notSpecific | Down | DM | Gain | 9 |
| **6_8433148_8435412_51000** | 6_8433148_8435412 | SLC35B3 | 51000 | 2.58319877396862 | 0.26706947 | 1 | only_DIvsD7 | DownUp | notDM | Gain | 4 |
| **6_8435908_8436586_51000** | 6_8435908_8436586 | SLC35B3 | 51000 | 2.06445798922692 | 0.26706947 | 4 | only_DIvsD7 | DownUp | notDM | Loss | 3 |
| **6_88118421_88119016_57038** | 6_88118421_88119016 | RARS2 | 57038 | 5.25697257272491 | 0.85252194 | 1 | only_H3vsUI | DownUp | notDM | Gain | 1 |
| **6_88287853_88289303_57038** | 6_88287853_88289303 | RARS2 | 57038 | 4.01782603803042 | 0.35834446 | 7 | only_H3vsUI | DownUp | notDM | Gain | 5 |
| **6_88297613_88299402_57038** | 6_88297613_88299402 | RARS2 | 57038 | 2.52374646681156 | 0.01095251 | 1 | only_H3vsUI | DownUp | notDM | Loss | 8 |
| **6_88300068_88302048_57038** | 6_88300068_88302048 | RARS2 | 57038 | 2.52504480703685 | 0.01095251 | 1 | only_H3vsUI | DownUp | notDM | Gain | 4 |

| | | | | | | | | | | | |
|---|---|---|---|---|---|---|---|---|---|---|---|
| **6_88406611_88410603_57038** | 6_88406611_88410603 | RARS2 | 57038 | 5.02894520300236 | -0.63823687 | 2 | only_H3vsUI | DownUp | notDM | Loss | 7 |
| **6_88412185_88413315_57038** | 6_88412185_88413315 | RARS2 | 57038 | 5.05102124916611 | -0.63823687 | 5 | only_H3vsUI | DownUp | notDM | Gain | 2 |
| **6_88433263_88433706_57038** | 6_88433263_88433706 | RARS2 | 57038 | 5.12562438037366 | -0.75768427 | 7 | only_H3vsUI | DownUp | notDM | Gain | 2 |
| **6_88439411_88442666_57038** | 6_88439411_88442666 | RARS2 | 57038 | 5.14512800789656 | -0.88309854 | 7 | only_H3vsUI | DownUp | notDM | Gain | 5 |
| **7_100075646_100076750_221914** | 7_100075646_100076750 | GPC2 | 221914 | 5.47806987520593 | -0.81406889 | 2 | only_DIvsD7 | DownUp | notDM | Gain | 2 |
| **7_100271965_100272714_221914** | 7_100271965_100272714 | GPC2 | 221914 | 5.69633629013329 | -0.84004981 | 2 | only_DIvsD7 | DownUp | notDM | Loss | 7 |
| **7_101655167_101657496_79706** | 7_101655167_101657496 | PRKRIP1 | 79706 | 5.57900001720429 | -0.84372529 | 7 | only_H3vsUI | Down | notDM | Gain | 2 |
| **7_101795038_101797392_79706** | 7_101795038_101797392 | PRKRIP1 | 79706 | 5.37928919740077 | -0.78078806 | 7 | only_H3vsUI | Down | notDM | Gain | 2 |
| **7_101915932_101918180_79706** | 7_101915932_101918180 | PRKRIP1 | 79706 | 5.07446169508754 | -0.63375008 | 4 | only_H3vsUI | Down | notDM | Loss | 2 |
| **7_102035786_102036688_79706** | 7_102035786_102036688 | PRKRIP1 | 79706 | 2.0899051114394 | 0.13840708 | 2 | only_H3vsUI | Down | notDM | Loss | 8 |
| **7_102036997_102038394_79706** | 7_102036997_102038394 | PRKRIP1 | 79706 | 2.27646180417324 | 0.13840708 | 1 | only_H3vsUI | Down | notDM | Loss | 8 |
| **7_102071727_102072922_79706** | 7_102071727_102072922 | PRKRIP1 | 79706 | 4.82872432713879 | 0.64014837 | 7 | only_H3vsUI | Down | notDM | Gain | 1 |
| **7_102074202_102075790_79706** | 7_102074202_102075790 | PRKRIP1 | 79706 | 4.57280192369961 | 0.65603416 | 1 | only_H3vsUI | Down | notDM | Loss | 7 |
| **7_102080035_102081447_79706** | 7_102080035_102081447 | PRKRIP1 | 79706 | 4.63575509607829 | 0.829419 | 3 | only_H3vsUI | Down | notDM | Gain | 1 |
| **7_107382526_107384061_10192797** | 7_107382526_107384061 | LOC101927974 | 101927974 | 0 | 0.00226154 | 2 | only_DIvsD7 | DownUp | notDM | Gain | 4 |
| **7_107384941_107386869_10192797** | 7_107384941_107386869 | LOC101927974 | 101927974 | 3.13735411137073 | 0.00059713 | 1 | only_DIvsD7 | DownUp | notDM | Loss | 6 |
| **7_1125954_1128474_2852** | 7_1125954_1128474 | GPER1 | 2852 | 0 | 0.04202439 | 2 | only_DIvsD7 | DownUp | notDM | Loss | 6 |
| **7_123173846_123174651_154865** | 7_123173846_123174651 | IQUB | 154865 | 0 | 0.17964263 | 2 | only_H3vsUI | UpDown | notDM | Gain | 4 |
| **7_123175338_123176097_154865** | 7_123175338_123176097 | IQUB | 154865 | 2.85003325768977 | 0.15570519 | 2 | only_H3vsUI | UpDown | notDM | Loss | 3 |
| **7_123177502_123179294_154865** | 7_123177502_123179294 | IQUB | 154865 | 3.45818443557026 | 0.18859898 | 6 | only_H3vsUI | UpDown | notDM | Loss | 3 |
| **7_127223317_127225587_381** | 7_127223317_127225587 | ARF5 | 381 | 3.4491697321652 | -0.48677081 | 2 | only_DIvsD7 | DownUp | notDM | Loss | 8 |
| **7_127225767_127226489_381** | 7_127225767_127226489 | ARF5 | 381 | 3.28126068705501 | -0.48677081 | 2 | only_DIvsD7 | DownUp | notDM | Gain | 2 |
| **7_127228684_127230086_381** | 7_127228684_127230086 | ARF5 | 381 | 2.35602585719312 | -0.80874344 | 1 | only_DIvsD7 | DownUp | notDM | Loss | 7 |
| **7_128043816_128045896_55798** | 7_128043816_128045896 | METTL2B | 55798 | 4.85072585874763 | 0.71355345 | 1 | only_DIvsD7 | Down | notDM | Loss | 7 |
| **7_128050128_128051711_55798** | 7_128050128_128051711 | METTL2B | 55798 | 4.64526503021562 | -0.50380088 | 2 | only_DIvsD7 | Down | notDM | Gain | 2 |
| **7_128094488_128095833_55798** | 7_128094488_128095833 | METTL2B | 55798 | 1.79239168949825 | 0.3163868 | 5 | only_DIvsD7 | Down | notDM | Gain | 4 |
| **7_128096100_128100481_55798** | 7_128096100_128100481 | METTL2B | 55798 | 2.31806333496276 | 0.23750913 | 1 | only_DIvsD7 | Down | notDM | Loss | 8 |
| **7_128115829_128116721_55798** | 7_128115829_128116721 | METTL2B | 55798 | 1.94448267215017 | 0.21728358 | 3 | only_DIvsD7 | Down | notDM | Gain | 1 |
| **7_128116978_128118752_55798** | 7_128116978_128118752 | METTL2B | 55798 | 2.23552844690755 | 0.21728358 | 1 | only_DIvsD7 | Down | notDM | Loss | 8 |
| **7_128169706_128170714_55798** | 7_128169706_128170714 | METTL2B | 55798 | 4.72345567203519 | 0.67945076 | 4 | only_DIvsD7 | Down | notDM | Loss | 7 |
| **7_128171941_128174414_55798** | 7_128171941_128174414 | METTL2B | 55798 | 4.8810935077074 | 0.60127036 | 1 | only_DIvsD7 | Down | notDM | Loss | 4 |
| **7_128578086_128579026_55798** | 7_128578086_128579026 | METTL2B | 55798 | 5.68322180209473 | 0.81520045 | 2 | only_DIvsD7 | Down | notDM | Loss | 3 |
| **7_129007076_129008338_4899** | 7_129007076_129008338 | NRF1 | 4899 | 5.38604035107763 | 0.83858238 | 4 | only_H3vsUI | Up | notDM | Loss | 5 |
| **7_129208991_129210655_4899** | 7_129208991_129210655 | NRF1 | 4899 | 4.61173392632452 | -0.68620973 | 7 | only_H3vsUI | Up | notDM | Gain | 1 |

| | | | | | | | | | | | |
|---|---|---|---|---|---|---|---|---|---|---|---|
| **7_129249637_129251098_4899** | 7_129249637_129251098 | NRF1 | 4899 | 2.66086547800387 | -0.48497288 | 1 | only_H3vsUI | Up | notDM | Loss | 8 |
| **7_129252248_129253597_4899** | 7_129252248_129253597 | NRF1 | 4899 | 2.82607480270083 | -0.49581153 | 1 | only_H3vsUI | Up | notDM | Loss | 8 |
| **7_129304010_129305895_4899** | 7_129304010_129305895 | NRF1 | 4899 | 3.83384804953115 | 0.00436548 | 6 | only_H3vsUI | Up | notDM | Gain | 1 |
| **7_134671531_134673817_340351** | 7_134671531_134673817 | AGBL3 | 340351 | 2.35602585719312 | 0.49425399 | 2 | only_H3vsUI | UpDown | notDM | Gain | 4 |
| **7_134675609_134677987_340351** | 7_134675609_134677987 | AGBL3 | 340351 | 3.63397315578967 | 0.55270713 | 3 | only_H3vsUI | UpDown | notDM | Gain | 1 |
| **7_134976165_134977416_340351** | 7_134976165_134977416 | AGBL3 | 340351 | 5.48413748254072 | -0.7709363 | 7 | only_H3vsUI | UpDown | notDM | Gain | 1 |
| **7_138718749_138720023_92092** | 7_138718749_138720023 | ZC3HAV1L | 92092 | 2.8767949762007 | 0.85117318 | 1 | only_DIvsD7 | Down | notDM | Loss | 8 |
| **7_138720845_138722043_92092** | 7_138720845_138722043 | ZC3HAV1L | 92092 | 1.85733249643127 | 0.85139731 | 1 | only_DIvsD7 | Down | notDM | Gain | 4 |
| **7_140096050_140098001_401409** | 7_140096050_140098001 | RAB19 | 401409 | 3.76663588631027 | -0.10118843 | 2 | only_H3vsUI | UpDown | notDM | Gain | 4 |
| **7_140098345_140099014_401409** | 7_140098345_140099014 | RAB19 | 401409 | 3.68394713075151 | -0.10775851 | 4 | only_H3vsUI | UpDown | notDM | Gain | 1 |
| **7_140395661_140396444_10013471** | 7_140395661_140396444 | NDUFB2-AS1 | 100134713 | 2.63748972951251 | 5.416e-05 | 5 | only_H3vsUI | DownUp | notDM | Loss | 3 |
| **7_140397058_140398762_10013471** | 7_140397058_140398762 | NDUFB2-AS1 | 100134713 | 2.26245108973043 | 5.416e-05 | 1 | only_H3vsUI | DownUp | notDM | Loss | 8 |
| **7_150076629_150078433_285971** | 7_150076629_150078433 | ZNF775 | 285971 | 4.03148925570975 | 0.82742494 | 1 | notSpecific | Down | DM | Loss | 10 |
| **7_150076629_150078433_29803** | 7_150076629_150078433 | REPIN1 | 29803 | 4.02938377768521 | 0.88912216 | 1 | notSpecific | Up | DM | Loss | 10 |
| **7_150076629_150078433_65999** | 7_150076629_150078433 | LRRC61 | 65999 | 4.75013024750556 | 0.68365087 | 1 | notSpecific | Up | DM | Loss | 10 |
| **7_150735239_150737610_9311** | 7_150735239_150737610 | ASIC3 | 9311 | 3.91200937558698 | 0.01613182 | 7 | only_H3vsUI | UpDown | notDM | Loss | 5 |
| **7_150753513_150754992_9311** | 7_150753513_150754992 | ASIC3 | 9311 | 3.71725431276255 | -0.01298073 | 1 | only_H3vsUI | UpDown | notDM | Loss | 3 |
| **7_150755211_150756578_9311** | 7_150755211_150756578 | ASIC3 | 9311 | 3.83966655688243 | -0.01102105 | 1 | only_H3vsUI | UpDown | notDM | Loss | 6 |
| **7_150863786_150864696_2636** | 7_150863786_150864696 | GBX1 | 2636 | 0 | 0 | 4 | only_H3vsUI | DownUp | notDM | Loss | 3 |
| **7_150864909_150865981_2636** | 7_150864909_150865981 | GBX1 | 2636 | 2.44090908206522 | 0 | 2 | only_H3vsUI | DownUp | notDM | Loss | 3 |
| **7_155742910_155745979_389602** | 7_155742910_155745979 | LOC389602 | 389602 | 3.97071870372019 | 0.03361363 | 7 | only_DIvsD7 | UpDown | notDM | Gain | 1 |
| **7_156433533_156434762_64434** | 7_156433533_156434762 | NOM1 | 64434 | 5.48806538924847 | 0.9398379 | 1 | only_DIvsD7 | Down | notDM | Loss | 7 |
| **7_156724045_156725759_64434** | 7_156724045_156725759 | NOM1 | 64434 | 4.22164892819223 | 0.27109852 | 6 | only_DIvsD7 | Down | notDM | Gain | 1 |
| **7_156742779_156745323_64434** | 7_156742779_156745323 | NOM1 | 64434 | 2.56110138364906 | -0.21866141 | 1 | only_DIvsD7 | Down | notDM | Loss | 8 |
| **7_156793343_156794070_64434** | 7_156793343_156794070 | NOM1 | 64434 | 4.70695662127693 | 0.47153618 | 1 | only_DIvsD7 | Down | notDM | Gain | 2 |
| **7_156800285_156802482_64434** | 7_156800285_156802482 | NOM1 | 64434 | 4.76245348236355 | -0.79706196 | 1 | only_DIvsD7 | Down | notDM | Loss | 7 |
| **7_156930373_156931284_64434** | 7_156930373_156931284 | NOM1 | 64434 | 5.27406081518904 | -0.69920635 | 3 | only_DIvsD7 | Down | notDM | Gain | 2 |
| **7_2287802_2289530_55501** | 7_2287802_2289530 | CHST12 | 55501 | 5.18665691357812 | 0.71465671 | 7 | only_H3vsUI | UpDown | notDM | Loss | 1 |
| **7_23338372_23338828_29896** | 7_23338372_23338828 | TRA2A | 29896 | 5.36704453397708 | -0.8516685 | 3 | only_DIvsD7 | DownUp | notDM | Gain | 2 |
| **7_23339150_23341345_29896** | 7_23339150_23341345 | TRA2A | 29896 | 5.36232410935087 | -0.80084095 | 1 | only_DIvsD7 | DownUp | notDM | Gain | 2 |
| **7_23502260_23510092_29896** | 7_23502260_23510092 | TRA2A | 29896 | 4.78890336165243 | 0.52068447 | 1 | only_DIvsD7 | DownUp | notDM | Loss | 6 |
| **7_23514395_23517319_29896** | 7_23514395_23517319 | TRA2A | 29896 | 4.73482382309632 | 0.76599757 | 2 | only_DIvsD7 | DownUp | notDM | Gain | 5 |
| **7_23567585_23571118_29896** | 7_23567585_23571118 | TRA2A | 29896 | 2.70243053644553 | -0.37106535 | 1 | only_DIvsD7 | DownUp | notDM | Loss | 8 |
| **7_23571744_23572650_29896** | 7_23571744_23572650 | TRA2A | 29896 | 2.09691001300806 | -0.37106535 | 1 | only_DIvsD7 | DownUp | notDM | Loss | 7 |

| | | | | | | | | | | | |
|---|---|---|---|---|---|---|---|---|---|---|---|
| **7_2395045_2397530_55501** | 7_2395045_2397530 | CHST12 | 55501 | 4.65985917735731 | -0.47049481 | 1 | only_H3vsUI | UpDown | notDM | Loss | 7 |
| **7_24323657_24325326_4852** | 7_24323657_24325326 | NPY | 4852 | 0 | -0.01365784 | 4 | only_DIvsD7 | DownUp | notDM | Loss | 3 |
| **7_2439338_2440370_55501** | 7_2439338_2440370 | CHST12 | 55501 | 3.45545396877863 | 0.92430348 | 3 | only_H3vsUI | UpDown | notDM | Gain | 1 |
| **7_2442172_2442938_55501** | 7_2442172_2442938 | CHST12 | 55501 | 2.45636603312904 | 0.27671153 | 1 | only_H3vsUI | UpDown | notDM | Loss | 3 |
| **7_2443451_2444813_55501** | 7_2443451_2444813 | CHST12 | 55501 | 2.36172783601759 | 0.17691068 | 1 | only_H3vsUI | UpDown | notDM | Loss | 3 |
| **7_2491268_2491984_55501** | 7_2491268_2491984 | CHST12 | 55501 | 4.6816662759979 | -0.48703979 | 4 | only_H3vsUI | UpDown | notDM | Loss | 2 |
| **7_26997595_26999321_3204** | 7_26997595_26999321 | HOXA7 | 3204 | 5.2944177235619 | 0.9859578 | 6 | notSpecific | UpDown | DM | Gain | 9 |
| **7_27173865_27176921_3202** | 7_27173865_27176921 | HOXA5 | 3202 | 3.80393484986384 | 0.16195625 | 4 | only_H3vsUI | DownUp | notDM | Gain | 4 |
| **7_27188458_27191346_3202** | 7_27188458_27191346 | HOXA5 | 3202 | 3.71374247840908 | 0.15533317 | 1 | only_H3vsUI | DownUp | notDM | Loss | 6 |
| **7_28448898_28451090_9586** | 7_28448898_28451090 | CREB5 | 9586 | 0 | 0.93405073 | 4 | notSpecific | Up | DM | Loss | 10 |
| **7_38971978_38972541_27072** | 7_38971978_38972541 | VPS41 | 27072 | 0 | 0 | 4 | notSpecific | UpDown | DM | Gain | 9 |
| **7_45806410_45808275_641977** | 7_45806410_45808275 | SEPTIN7P2 | 641977 | 2.4456042032736 | -0.16689438 | 1 | only_DIvsD7 | UpDown | notDM | Loss | 6 |
| **7_45808740_45809905_641977** | 7_45808740_45809905 | SEPTIN7P2 | 641977 | 2.19865708695442 | -0.16073277 | 1 | only_DIvsD7 | UpDown | notDM | Loss | 3 |
| **7_45908793_45909722_641977** | 7_45908793_45909722 | SEPTIN7P2 | 641977 | 5.00104972299307 | -0.61238419 | 6 | only_DIvsD7 | UpDown | notDM | Gain | 1 |
| **7_47631527_47633420_64759** | 7_47631527_47633420 | TNS3 | 64759 | 3.99064958831885 | 0.769636 | 7 | notSpecific | Down | DM | Gain | 9 |
| **7_47813708_47816461_64759** | 7_47813708_47816461 | TNS3 | 64759 | 5.28333063306365 | 0.71711767 | 7 | notSpecific | Down | DM | Gain | 9 |
| **7_47813708_47816461_84847** | 7_47813708_47816461 | LINC00525 | 84847 | 4.03193334027561 | -0.07505988 | 7 | notSpecific | UpDown | DM | Gain | 9 |
| **7_5534207_5535737_60** | 7_5534207_5535737 | ACTB | 60 | 4.53780650664498 | 0.60194512 | 3 | only_H3vsUI | Down | notDM | Loss | 5 |
| **7_5547987_5551390_60** | 7_5547987_5551390 | ACTB | 60 | 4.27521918678215 | 0.29524514 | 8 | only_H3vsUI | Down | notDM | Loss | 5 |
| **7_5551410_5553149_60** | 7_5551410_5553149 | ACTB | 60 | 4.23258956275479 | -0.1851215 | 1 | only_H3vsUI | Down | notDM | Loss | 8 |
| **7_5553524_5555361_60** | 7_5553524_5555361 | ACTB | 60 | 4.17236937676384 | -0.1851215 | 5 | only_H3vsUI | Down | notDM | Gain | 4 |
| **7_5565583_5569061_60** | 7_5565583_5569061 | ACTB | 60 | 3.06892761168207 | -0.15621015 | 1 | only_H3vsUI | Down | notDM | Loss | 8 |
| **7_5570384_5572067_60** | 7_5570384_5572067 | ACTB | 60 | 2.17897694729317 | -0.24107724 | 5 | only_H3vsUI | Down | notDM | Loss | 4 |
| **7_5594971_5595656_60** | 7_5594971_5595656 | ACTB | 60 | 3.83033199345196 | 0.08631142 | 5 | only_H3vsUI | Down | notDM | Loss | 2 |
| **7_5596323_5598628_60** | 7_5596323_5598628 | ACTB | 60 | 3.57909732655264 | 0.04014872 | 1 | only_H3vsUI | Down | notDM | Loss | 4 |
| **7_5600927_5603861_60** | 7_5600927_5603861 | ACTB | 60 | 0 | -0.00557989 | 5 | only_H3vsUI | Down | notDM | Gain | 4 |
| **7_5608646_5611894_60** | 7_5608646_5611894 | ACTB | 60 | 4.58451214271295 | 0.58650563 | 2 | only_H3vsUI | Down | notDM | Loss | 4 |
| **7_5633777_5639595_60** | 7_5633777_5639595 | ACTB | 60 | 4.80307454954133 | 0.63972851 | 1 | only_H3vsUI | Down | notDM | Loss | 4 |
| **7_5727229_5734825_60** | 7_5727229_5734825 | ACTB | 60 | 5.19589688619353 | 0.68965602 | 7 | only_H3vsUI | Down | notDM | Loss | 5 |
| **7_6018905_6019962_7965** | 7_6018905_6019962 | AIMP2 | 7965 | 4.46112319663961 | 0.38778863 | 7 | only_H3vsUI | DownUp | notDM | Gain | 2 |
| **7_6034773_6036087_7965** | 7_6034773_6036087 | AIMP2 | 7965 | 4.10927473823029 | 0.46588659 | 3 | only_H3vsUI | DownUp | notDM | Loss | 5 |
| **7_6046512_6048638_7965** | 7_6046512_6048638 | AIMP2 | 7965 | 2.37839790094814 | -0.29072726 | 1 | only_H3vsUI | DownUp | notDM | Loss | 8 |
| **7_6049647_6051091_60** | 7_6049647_6051091 | ACTB | 60 | 5.68071071228831 | -0.81329745 | 1 | only_H3vsUI | Down | notDM | Loss | 7 |
| **7_6049647_6051091_7965** | 7_6049647_6051091 | AIMP2 | 7965 | 2.84633711212981 | 0.68800368 | 1 | only_H3vsUI | DownUp | notDM | Loss | 8 |

| | | | | | | | | | | | |
|---|---|---|---|---|---|---|---|---|---|---|---|
| **7_64022412_64023159_340252** | 7_64022412_64023159 | ZNF680 | 340252 | 2.41664050733828 | 0.20535469 | 2 | only_DIvsD7 | Down | notDM | Gain | 4 |
| **7_65234391_65235463_2990** | 7_65234391_65235463 | GUSB | 2990 | 5.32579470436874 | 0.86605594 | 2 | only_H3vsUI | UpDown | notDM | Loss | 3 |
| **7_65445353_65447099_2990** | 7_65445353_65447099 | GUSB | 2990 | 2 | 0.31728409 | 1 | only_H3vsUI | UpDown | notDM | Loss | 6 |
| **7_65447351_65447741_2990** | 7_65447351_65447741 | GUSB | 2990 | 2.09691001300806 | 0.26093357 | 3 | only_H3vsUI | UpDown | notDM | Gain | 2 |
| **7_7216106_7218810_54468** | 7_7216106_7218810 | MIOS | 54468 | 5.58869178859222 | -0.854085 | 3 | only_DIvsD7 | UpDown | notDM | Gain | 5 |
| **7_73868660_73870451_9569** | 7_73868660_73870451 | GTF2IRD1 | 9569 | 2.34830486304816 | 0.67387368 | 1 | notSpecific | UpDown | DM | Loss | 10 |
| **7_7477591_7479593_340267** | 7_7477591_7479593 | COL28A1 | 340267 | 2.72181061521255 | 0 | 7 | notSpecific | Down | DM | Gain | 9 |
| **7_7558712_7560195_54468** | 7_7558712_7560195 | MIOS | 54468 | 4.66649926056169 | 0.9230659 | 6 | only_DIvsD7 | UpDown | notDM | Gain | 1 |
| **7_7607018_7609130_54468** | 7_7607018_7609130 | MIOS | 54468 | 2.63144376901317 | 0.69722906 | 1 | only_DIvsD7 | UpDown | notDM | Loss | 6 |
| **7_76234814_76236679_22932** | 7_76234814_76236679 | POMZP3 | 22932 | 3.68160299873087 | 0 | 7 | only_H3vsUI | DownUp | notDM | Gain | 1 |
| **7_76254744_76256091_22932** | 7_76254744_76256091 | POMZP3 | 22932 | 0 | 0 | 2 | only_H3vsUI | DownUp | notDM | Loss | 8 |
| **7_76256670_76257785_22932** | 7_76256670_76257785 | POMZP3 | 22932 | 2.05690485133647 | 0.00375152 | 1 | only_H3vsUI | DownUp | notDM | Gain | 4 |
| **7_7770688_7772538_54468** | 7_7770688_7772538 | MIOS | 54468 | 5.21533876757166 | -0.78313072 | 6 | only_DIvsD7 | UpDown | notDM | Loss | 7 |
| **7_8009406_8010762_54468** | 7_8009406_8010762 | MIOS | 54468 | 5.60500751179512 | 0.79338712 | 4 | only_DIvsD7 | UpDown | notDM | Gain | 1 |
| **7_90892261_90893535_8321** | 7_90892261_90893535 | FZD1 | 8321 | 2.39619934709574 | 0.07985797 | 1 | only_DIvsD7 | UpDown | notDM | Gain | 4 |
| **7_90894523_90895231_8321** | 7_90894523_90895231 | FZD1 | 8321 | 2.87040390527903 | 0.01133284 | 2 | only_DIvsD7 | UpDown | notDM | Loss | 3 |
| **7_91760467_91763140_5189** | 7_91760467_91763140 | PEX1 | 5189 | 5.59616808671899 | -0.8496619 | 1 | only_H3vsUI | Down | notDM | Loss | 2 |
| **7_91760467_91763140_613126** | 7_91760467_91763140 | CYP51A1-AS1 | 613126 | 2.89153745767256 | 0.00806859 | 1 | only_H3vsUI | UpDown | notDM | Loss | 6 |
| **7_91764044_91765223_5189** | 7_91764044_91765223 | PEX1 | 5189 | 5.59386953508819 | -0.89466051 | 1 | only_H3vsUI | Down | notDM | Gain | 2 |
| **7_91764044_91765223_613126** | 7_91764044_91765223 | CYP51A1-AS1 | 613126 | 2.10720996964787 | 0.00744816 | 1 | only_H3vsUI | UpDown | notDM | Gain | 4 |
| **7_91872775_91875180_5189** | 7_91872775_91875180 | PEX1 | 5189 | 5.45113527318776 | -0.82428041 | 1 | only_H3vsUI | Down | notDM | Gain | 2 |
| **7_91875802_91878105_5189** | 7_91875802_91878105 | PEX1 | 5189 | 5.44661637665445 | -0.82428041 | 2 | only_H3vsUI | Down | notDM | Loss | 2 |
| **7_92053054_92055779_5189** | 7_92053054_92055779 | PEX1 | 5189 | 5.00888534997626 | 0.82593947 | 2 | only_H3vsUI | Down | notDM | Gain | 4 |
| **7_92155835_92157642_5189** | 7_92155835_92157642 | PEX1 | 5189 | 2.03342375548695 | -0.9500895 | 2 | only_H3vsUI | Down | notDM | Gain | 4 |
| **7_92158278_92159444_5189** | 7_92158278_92159444 | PEX1 | 5189 | 2.72509452108147 | -0.9500895 | 2 | only_H3vsUI | Down | notDM | Loss | 8 |
| **7_92214339_92219234_5189** | 7_92214339_92219234 | PEX1 | 5189 | 4.75275504247068 | -0.89234605 | 1 | only_H3vsUI | Down | notDM | Gain | 2 |
| **7_92219759_92220574_5189** | 7_92219759_92220574 | PEX1 | 5189 | 4.79247573836125 | -0.76071217 | 2 | only_H3vsUI | Down | notDM | Loss | 7 |
| **7_92250532_92251796_5189** | 7_92250532_92251796 | PEX1 | 5189 | 4.96744968725126 | -0.75013003 | 7 | only_H3vsUI | Down | notDM | Gain | 2 |
| **7_92260971_92266057_5189** | 7_92260971_92266057 | PEX1 | 5189 | 5.01375543972348 | -0.8749029 | 6 | only_H3vsUI | Down | notDM | Gain | 5 |
| **7_92368691_92372595_5189** | 7_92368691_92372595 | PEX1 | 5189 | 5.32396948586744 | -0.73668413 | 8 | only_H3vsUI | Down | notDM | Gain | 5 |
| **7_92436830_92441735_5189** | 7_92436830_92441735 | PEX1 | 5189 | 5.44572404578751 | -0.93259391 | 7 | only_H3vsUI | Down | notDM | Gain | 5 |
| **7_92462899_92465647_5189** | 7_92462899_92465647 | PEX1 | 5189 | 5.48450768138973 | -0.75878745 | 1 | only_H3vsUI | Down | notDM | Loss | 2 |
| **7_92466282_92467256_5189** | 7_92466282_92467256 | PEX1 | 5189 | 5.4892959915396 | -0.84640543 | 2 | only_H3vsUI | Down | notDM | Loss | 7 |
| **7_96177706_96178705_7979** | 7_96177706_96178705 | SEM1 | 7979 | 5.20539114621576 | -0.70164554 | 6 | only_DIvsD7 | DownUp | notDM | Loss | 3 |

| | | | | | | | | | | | |
|---|---|---|---|---|---|---|---|---|---|---|---|
| **7_96337346_96338733_7979** | 7_96337346_96338733 | SEM1 | 7979 | 2.6730209071289 | -0.2082313 | 1 | only_DIvsD7 | DownUp | notDM | Loss | 6 |
| **7_96339317_96340117_7979** | 7_96339317_96340117 | SEM1 | 7979 | 2.06445798922692 | -0.2082313 | 1 | only_DIvsD7 | DownUp | notDM | Gain | 4 |
| **7_99514509_99516833_221914** | 7_99514509_99516833 | GPC2 | 221914 | 5.41188895239585 | -0.88498609 | 1 | only_DIvsD7 | DownUp | notDM | Gain | 2 |
| **7_99517290_99518590_221914** | 7_99517290_99518590 | GPC2 | 221914 | 5.40891971465971 | -0.88498796 | 2 | only_DIvsD7 | DownUp | notDM | Gain | 2 |
| **7_99646284_99647374_221914** | 7_99646284_99647374 | GPC2 | 221914 | 5.10591533724466 | -0.73744577 | 2 | only_DIvsD7 | DownUp | notDM | Gain | 2 |
| **7_99647614_99648745_221914** | 7_99647614_99648745 | GPC2 | 221914 | 5.10121762674755 | -0.67823342 | 1 | only_DIvsD7 | DownUp | notDM | Loss | 7 |
| **7_99715661_99716773_221914** | 7_99715661_99716773 | GPC2 | 221914 | 4.76507220110279 | -0.79169996 | 1 | only_DIvsD7 | DownUp | notDM | Loss | 7 |
| **7_99717210_99718620_221914** | 7_99717210_99718620 | GPC2 | 221914 | 4.75105573910668 | -0.6749991 | 1 | only_DIvsD7 | DownUp | notDM | Gain | 2 |
| **7_99717210_99718620_255374** | 7_99717210_99718620 | MBLAC1 | 255374 | 3.7626035495668 | -0.00281407 | 1 | only_DIvsD7 | DownUp | notDM | Gain | 4 |
| **7_99722255_99724640_221914** | 7_99722255_99724640 | GPC2 | 221914 | 4.70200810031504 | -0.78364853 | 1 | only_DIvsD7 | DownUp | notDM | Loss | 7 |
| **7_99722255_99724640_255374** | 7_99722255_99724640 | MBLAC1 | 255374 | 0 | -0.02140741 | 1 | only_DIvsD7 | DownUp | notDM | Loss | 8 |
| **7_99724869_99726291_221914** | 7_99724869_99726291 | GPC2 | 221914 | 4.68752896121463 | -0.78364853 | 2 | only_DIvsD7 | DownUp | notDM | Loss | 7 |
| **7_99724869_99726291_255374** | 7_99724869_99726291 | MBLAC1 | 255374 | 2.66558099101795 | -0.02140741 | 2 | only_DIvsD7 | DownUp | notDM | Loss | 7 |
| **7_99745643_99746218_221914** | 7_99745643_99746218 | GPC2 | 221914 | 4.45901533230183 | -0.33911925 | 5 | only_DIvsD7 | DownUp | notDM | Loss | 3 |
| **7_99746609_99748388_221914** | 7_99746609_99748388 | GPC2 | 221914 | 4.42493061444988 | -0.73461663 | 1 | only_DIvsD7 | DownUp | notDM | Gain | 2 |
| **7_99754359_99756169_221914** | 7_99754359_99756169 | GPC2 | 221914 | 4.27466576908139 | -0.65506192 | 1 | only_DIvsD7 | DownUp | notDM | Loss | 3 |
| **7_99756348_99757294_221914** | 7_99756348_99757294 | GPC2 | 221914 | 4.24789965088881 | -0.76502915 | 1 | only_DIvsD7 | DownUp | notDM | Gain | 2 |
| **7_99760355_99761636_221914** | 7_99760355_99761636 | GPC2 | 221914 | 4.12570892597265 | -0.33716131 | 6 | only_DIvsD7 | DownUp | notDM | Gain | 1 |
| **7_99774090_99775065_221914** | 7_99774090_99775065 | GPC2 | 221914 | 0 | 0.33810549 | 1 | only_DIvsD7 | DownUp | notDM | Gain | 4 |
| **7_99869931_99871149_221914** | 7_99869931_99871149 | GPC2 | 221914 | 4.97746295039542 | -0.84753973 | 1 | only_DIvsD7 | DownUp | notDM | Gain | 2 |
| **8_100023418_100025143_10537566** | 8_100023418_100025143 | VPS13B-DT | 105375666 | 2.11394335230684 | 0.41881649 | 1 | only_H3vsUI | DownUp | notDM | Gain | 4 |
| **8_100025762_100027022_10537566** | 8_100025762_100027022 | VPS13B-DT | 105375666 | 2.69196510276736 | 0.41881649 | 2 | only_H3vsUI | DownUp | notDM | Gain | 4 |
| **8_100030985_100032090_10537566** | 8_100030985_100032090 | VPS13B-DT | 105375666 | 3.7570162347313 | -0.70719401 | 6 | only_H3vsUI | DownUp | notDM | Loss | 7 |
| **8_10533806_10535519_203076** | 8_10533806_10535519 | C8orf74 | 203076 | 3.21801004298436 | 0.00473115 | 6 | only_DIvsD7 | UpDown | notDM | Gain | 1 |
| **8_10535666_10539827_203076** | 8_10535666_10539827 | C8orf74 | 203076 | 3.54555450723407 | -0.0011363 | 6 | only_DIvsD7 | UpDown | notDM | Loss | 5 |
| **8_11313593_11314240_252969** | 8_11313593_11314240 | NEIL2 | 252969 | 5.49541250291948 | 0.81797715 | 3 | only_H3vsUI | DownUp | notDM | Loss | 5 |
| **8_11314699_11317572_252969** | 8_11314699_11317572 | NEIL2 | 252969 | 5.49076307584048 | 0.78845744 | 7 | only_H3vsUI | DownUp | notDM | Loss | 5 |
| **8_11593244_11594615_252969** | 8_11593244_11594615 | NEIL2 | 252969 | 4.51364369230904 | -0.55413612 | 2 | only_H3vsUI | DownUp | notDM | Gain | 2 |
| **8_11619230_11620701_252969** | 8_11619230_11620701 | NEIL2 | 252969 | 3.80915553174718 | -0.13199009 | 6 | only_H3vsUI | DownUp | notDM | Gain | 1 |
| **8_11627268_11628997_252969** | 8_11627268_11628997 | NEIL2 | 252969 | 1.34242268082221 | -0.25911023 | 1 | only_H3vsUI | DownUp | notDM | Gain | 4 |
| **8_11657509_11660001_252969** | 8_11657509_11660001 | NEIL2 | 252969 | 4.48091197784664 | -0.39915119 | 1 | only_H3vsUI | DownUp | notDM | Loss | 7 |
| **8_11660348_11665868_2222** | 8_11660348_11665868 | FDFT1 | 2222 | 1.86332286012046 | -0.22999265 | 2 | notSpecific | Down | DM | Loss | 10 |
| **8_11660348_11665868_252969** | 8_11660348_11665868 | NEIL2 | 252969 | 4.51985423434138 | -0.39032412 | 2 | only_H3vsUI | DownUp | DM | Loss | 10 |
| **8_11710062_11712754_252969** | 8_11710062_11712754 | NEIL2 | 252969 | 4.9181247383681 | -0.77526342 | 7 | only_H3vsUI | DownUp | notDM | Gain | 5 |

| | | | | | | | | | | | |
|---|---|---|---|---|---|---|---|---|---|---|---|
| **8_11725797_11726947_252969** | 8_11725797_11726947 | NEIL2 | 252969 | 4.99366103543716 | -0.64654945 | 1 | only_H3vsUI | DownUp | notDM | Gain | 2 |
| **8_11740781_11742145_252969** | 8_11740781_11742145 | NEIL2 | 252969 | 5.05513741463258 | -0.63114772 | 6 | only_H3vsUI | DownUp | notDM | Gain | 2 |
| **8_11758832_11760881_252969** | 8_11758832_11760881 | NEIL2 | 252969 | 5.11921628615176 | -0.78890073 | 2 | only_H3vsUI | DownUp | notDM | Gain | 2 |
| **8_117876358_117886234_644660** | 8_117876358_117886234 | RAD21-AS1 | 644660 | 2.63346845557959 | 0 | 2 | only_DIvsD7 | DownUp | notDM | Gain | 4 |
| **8_117887251_117887839_644660** | 8_117887251_117887839 | RAD21-AS1 | 644660 | 2.77085201164214 | 0 | 2 | only_DIvsD7 | DownUp | notDM | Gain | 4 |
| **8_124552063_124553475_114907** | 8_124552063_124553475 | FBXO32 | 114907 | 0 | 0.99697348 | 2 | notSpecific | Up | DM | Loss | 10 |
| **8_128745110_128748153_10027068** | 8_128745110_128748153 | CASC11 | 100270680 | 0 | 0.01843688 | 1 | only_DIvsD7 | Up | notDM | Loss | 3 |
| **8_128750243_128758282_10027068** | 8_128750243_128758282 | CASC11 | 100270680 | 3.60552052343747 | 0.01843688 | 1 | only_DIvsD7 | Up | notDM | Gain | 4 |
| **8_133686452_133687770_23639** | 8_133686452_133687770 | LRRC6 | 23639 | 1.75587485567249 | 0.12865739 | 2 | only_H3vsUI | DownUp | notDM | Loss | 3 |
| **8_133687890_133688637_23639** | 8_133687890_133688637 | LRRC6 | 23639 | 1.73239375982297 | 0.02250522 | 2 | only_H3vsUI | DownUp | notDM | Loss | 3 |
| **8_142126112_142128268_22898** | 8_142126112_142128268 | DENND3 | 22898 | 4.01994668167884 | 0.89352188 | 6 | notSpecific | UpDown | DM | Loss | 10 |
| **8_144348398_144349491_10050731** | 8_144348398_144349491 | MINCR | 100507316 | 4.15670055258202 | 0.16617087 | 1 | notSpecific | Down | DM | Loss | 10 |
| **8_144348398_144349491_2738** | 8_144348398_144349491 | GLI4 | 2738 | 2.05307844348342 | -0.03222497 | 1 | only_H3vsUI | UpDown | DM | Loss | 10 |
| **8_144348398_144349491_286128** | 8_144348398_144349491 | ZFP41 | 286128 | 4.28553480612003 | -0.27532489 | 1 | notSpecific | Down | DM | Loss | 10 |
| **8_144348398_144349491_79943** | 8_144348398_144349491 | ZNF696 | 79943 | 4.38145804692216 | -0.66002062 | 1 | notSpecific | UpDown | DM | Loss | 10 |
| **8_144349731_144351858_2738** | 8_144349731_144351858 | GLI4 | 2738 | 2.11058971029925 | -0.03222497 | 1 | only_H3vsUI | UpDown | notDM | Loss | 3 |
| **8_144360998_144363759_2738** | 8_144360998_144363759 | GLI4 | 2738 | 3.62644302533129 | -0.00050505 | 1 | only_H3vsUI | UpDown | notDM | Loss | 6 |
| **8_145548826_145550254_4796** | 8_145548826_145550254 | TONSL | 4796 | 5.07763675754486 | -0.71325714 | 1 | only_H3vsUI | Down | notDM | Loss | 7 |
| **8_145550655_145551703_4796** | 8_145550655_145551703 | TONSL | 4796 | 5.07234182151732 | -0.79173836 | 1 | only_H3vsUI | Down | notDM | Gain | 2 |
| **8_145560741_145561667_4796** | 8_145560741_145561667 | TONSL | 4796 | 5.03395021938242 | 0.69414059 | 2 | only_H3vsUI | Down | notDM | Loss | 7 |
| **8_145632725_145634533_4796** | 8_145632725_145634533 | TONSL | 4796 | 4.5477131862317 | -0.61864091 | 2 | only_H3vsUI | Down | notDM | Loss | 8 |
| **8_145634873_145636166_4796** | 8_145634873_145636166 | TONSL | 4796 | 4.52713991565525 | -0.40299878 | 1 | only_H3vsUI | Down | notDM | Loss | 7 |
| **8_145638539_145639770_4796** | 8_145638539_145639770 | TONSL | 4796 | 4.47796008011449 | -0.63551099 | 2 | only_H3vsUI | Down | notDM | Loss | 7 |
| **8_145651864_145653884_4796** | 8_145651864_145653884 | TONSL | 4796 | 4.20259728569243 | -0.27562499 | 1 | only_H3vsUI | Down | notDM | Loss | 8 |
| **8_145654045_145656031_4796** | 8_145654045_145656031 | TONSL | 4796 | 4.1397846642937 | -0.12692082 | 1 | only_H3vsUI | Down | notDM | Loss | 8 |
| **8_145667807_145669765_4796** | 8_145667807_145669765 | TONSL | 4796 | 1.79934054945358 | -0.01195445 | 1 | only_H3vsUI | Down | notDM | Loss | 8 |
| **8_145670058_145671039_4796** | 8_145670058_145671039 | TONSL | 4796 | 2.36735592102602 | -0.04466356 | 1 | only_H3vsUI | Down | notDM | Loss | 8 |
| **8_145686339_145687699_4796** | 8_145686339_145687699 | TONSL | 4796 | 4.21785228025989 | -0.65495794 | 2 | only_H3vsUI | Down | notDM | Loss | 7 |
| **8_145691475_145693418_4796** | 8_145691475_145693418 | TONSL | 4796 | 4.33605927786635 | 0.27093624 | 2 | only_H3vsUI | Down | notDM | Loss | 7 |
| **8_145701941_145703191_4796** | 8_145701941_145703191 | TONSL | 4796 | 4.50712694037055 | 0.44160792 | 2 | only_H3vsUI | Down | notDM | Loss | 7 |
| **8_145703460_145705866_4796** | 8_145703460_145705866 | TONSL | 4796 | 4.52717861880396 | -0.55892633 | 1 | only_H3vsUI | Down | notDM | Loss | 8 |
| **8_145741385_145743051_4796** | 8_145741385_145743051 | TONSL | 4796 | 4.85485236241783 | 0.67841152 | 1 | only_H3vsUI | Down | notDM | Loss | 7 |
| **8_145746252_145748146_4796** | 8_145746252_145748146 | TONSL | 4796 | 4.88324681249592 | 0.64618225 | 1 | only_H3vsUI | Down | notDM | Gain | 4 |
| **8_145748316_145750049_4796** | 8_145748316_145750049 | TONSL | 4796 | 4.89481986216669 | 0.64498757 | 5 | only_H3vsUI | Down | notDM | Loss | 4 |

| | | | | | | | | | | | |
|---|---|---|---|---|---|---|---|---|---|---|---|
| **8_145751275_145752480_4796** | 8_145751275_145752480 | TONSL | 4796 | 4.91105102032813 | 0.69297764 | 1 | only_H3vsUI | Down | notDM | Loss | 7 |
| **8_145908852_145910781_29894** | 8_145908852_145910781 | CPSF1 | 29894 | 5.4379106199747 | -0.78978477 | 1 | notSpecific | UpDown | DM | Loss | 10 |
| **8_145908852_145910781_7553** | 8_145908852_145910781 | ZNF7 | 7553 | 5.15286294552391 | 0.69514036 | 1 | notSpecific | Down | DM | Loss | 10 |
| **8_145908852_145910781_7564** | 8_145908852_145910781 | ZNF16 | 7564 | 5.42394182602921 | -0.78583411 | 1 | notSpecific | Down | DM | Loss | 10 |
| **8_145908852_145910781_80728** | 8_145908852_145910781 | ARHGAP39 | 80728 | 2.6170003411209 | 0.03667402 | 1 | notSpecific | Down | DM | Loss | 10 |
| **8_146052980_146054752_4796** | 8_146052980_146054752 | TONSL | 4796 | 5.58340850002881 | -0.7908327 | 1 | only_H3vsUI | Down | notDM | Loss | 7 |
| **8_146226908_146228223_286101** | 8_146226908_146228223 | ZNF252P | 286101 | 1.5910646070265 | -0.18328918 | 1 | only_H3vsUI | Down | notDM | Loss | 8 |
| **8_146228314_146229537_286101** | 8_146228314_146229537 | ZNF252P | 286101 | 1.74036268949424 | -0.25734298 | 2 | only_H3vsUI | Down | notDM | Loss | 8 |
| **8_25900725_25904880_64641** | 8_25900725_25904880 | EBF2 | 64641 | 0 | 0.0011955 | 4 | notSpecific | Up | DM | Loss | 10 |
| **8_26122775_26124657_5520** | 8_26122775_26124657 | PPP2R2A | 5520 | 4.38737202701981 | 0.6496331 | 6 | only_H3vsUI | UpDown | notDM | Gain | 1 |
| **8_26149669_26153823_5520** | 8_26149669_26153823 | PPP2R2A | 5520 | 2.50514997831991 | -0.35942655 | 1 | only_H3vsUI | UpDown | notDM | Loss | 8 |
| **8_26161420_26164564_5520** | 8_26161420_26164564 | PPP2R2A | 5520 | 4.08174324992272 | -0.25999267 | 8 | only_H3vsUI | UpDown | notDM | Gain | 4 |
| **8_26164693_26166565_5520** | 8_26164693_26166565 | PPP2R2A | 5520 | 4.17799796309657 | 0.24357875 | 8 | only_H3vsUI | UpDown | notDM | Loss | 5 |
| **8_28257233_28259274_7976** | 8_28257233_28259274 | FZD3 | 7976 | 4.96593510026899 | -0.72521404 | 1 | only_H3vsUI | Down | notDM | Loss | 7 |
| **8_28346481_28348920_7976** | 8_28346481_28348920 | FZD3 | 7976 | 3.44901531634779 | 0.81971337 | 2 | only_H3vsUI | Down | notDM | Gain | 4 |
| **8_28350752_28351646_7976** | 8_28350752_28351646 | FZD3 | 7976 | 1.93449845124357 | 0.53455235 | 2 | only_H3vsUI | Down | notDM | Loss | 8 |
| **8_28352158_28353801_7976** | 8_28352158_28353801 | FZD3 | 7976 | 2.58771096501891 | 0.59933745 | 2 | only_H3vsUI | Down | notDM | Loss | 8 |
| **8_32083093_32087357_3084** | 8_32083093_32087357 | NRG1 | 3084 | 5.50301261298037 | 0.99780783 | 7 | notSpecific | UpDown | DM | Gain | 9 |
| **8_32125946_32130162_3084** | 8_32125946_32130162 | NRG1 | 3084 | 5.44038635540571 | 0.99160483 | 7 | notSpecific | UpDown | DM | Gain | 9 |
| **8_40028089_40032229_56892** | 8_40028089_40032229 | TCIM | 56892 | 4.23304690209818 | 0.94902297 | 7 | notSpecific | Down | DM | Gain | 9 |
| **8_40066120_40074084_56892** | 8_40066120_40074084 | TCIM | 56892 | 4.74141162476655 | 0.74959432 | 7 | notSpecific | Down | DM | Gain | 9 |
| **8_42194999_42195930_5423** | 8_42194999_42195930 | POLB | 5423 | 1.64345267648619 | 0.28815495 | 2 | only_DIvsD7 | UpDown | notDM | Loss | 3 |
| **8_42196194_42197768_5423** | 8_42196194_42197768 | POLB | 5423 | 2.34830486304816 | 0.28815495 | 1 | only_DIvsD7 | UpDown | notDM | Loss | 6 |
| **8_42248261_42249183_5423** | 8_42248261_42249183 | POLB | 5423 | 4.71842694724808 | 0.87307996 | 1 | only_DIvsD7 | UpDown | notDM | Gain | 1 |
| **8_42353573_42356309_5423** | 8_42353573_42356309 | POLB | 5423 | 5.19756172446986 | -0.79033838 | 6 | only_DIvsD7 | UpDown | notDM | Loss | 7 |
| **8_42358219_42362088_5423** | 8_42358219_42362088 | POLB | 5423 | 5.21007494692755 | -0.81447921 | 2 | only_DIvsD7 | UpDown | notDM | Gain | 5 |
| **8_42575649_42578077_5423** | 8_42575649_42578077 | POLB | 5423 | 5.57937082075941 | -0.84584727 | 7 | only_DIvsD7 | UpDown | notDM | Loss | 7 |
| **8_48870561_48872457_7336** | 8_48870561_48872457 | UBE2V2 | 7336 | 4.68591184126796 | -0.71636717 | 2 | only_H3vsUI | DownUp | notDM | Loss | 7 |
| **8_48873616_48876053_7336** | 8_48873616_48876053 | UBE2V2 | 7336 | 4.65246875114999 | -0.62678152 | 1 | only_H3vsUI | DownUp | notDM | Gain | 2 |
| **8_48904950_48906395_7336** | 8_48904950_48906395 | UBE2V2 | 7336 | 4.16334030561825 | 0.53583671 | 4 | only_H3vsUI | DownUp | notDM | Gain | 1 |
| **8_48920081_48920797_7336** | 8_48920081_48920797 | UBE2V2 | 7336 | 2.2148438480477 | -0.46399982 | 2 | only_H3vsUI | DownUp | notDM | Loss | 8 |
| **8_48921377_48923092_7336** | 8_48921377_48923092 | UBE2V2 | 7336 | 2.56584781867352 | -0.46364113 | 1 | only_H3vsUI | DownUp | notDM | Loss | 8 |
| **8_54426236_54427701_100507516** | 8_54426236_54427701 | LOC100507516 | 100507516 | 3.94403828005442 | 0.03363718 | 7 | only_H3vsUI | UpDown | notDM | Gain | 1 |
| **8_54435524_54437666_100507516** | 8_54435524_54437666 | LOC100507516 | 100507516 | 0 | 0.05557334 | 4 | only_H3vsUI | UpDown | notDM | Loss | 3 |

| | | | | | | | | | | | |
|---|---|---|---|---|---|---|---|---|---|---|---|
| **8_54624318_54625012_8601** | 8_54624318_54625012 | RGS20 | 8601 | 5.22639624165003 | 0.70385359 | 2 | only_H3vsUI | Down | notDM | Gain | 1 |
| **8_54791504_54793435_8601** | 8_54791504_54793435 | RGS20 | 8601 | 0 | 0.72088579 | 1 | only_H3vsUI | Down | notDM | Loss | 8 |
| **8_54793581_54800049_8601** | 8_54793581_54800049 | RGS20 | 8601 | 2.17897694729317 | 0.65668447 | 1 | only_H3vsUI | Down | notDM | Loss | 8 |
| **8_54856573_54857564_8601** | 8_54856573_54857564 | RGS20 | 8601 | 2.50785587169583 | 0.09697344 | 7 | only_H3vsUI | Down | notDM | Gain | 1 |
| **8_54857881_54859655_8601** | 8_54857881_54859655 | RGS20 | 8601 | 0 | 0.1087709 | 7 | only_H3vsUI | Down | notDM | Gain | 4 |
| **8_67339882_67340796_100505676** | 8_67339882_67340796 | RRS1-AS1 | 100505676 | 2.62013605497376 | 0.01689134 | 2 | only_H3vsUI | UpDown | notDM | Loss | 3 |
| **8_67341619_67345688_100505676** | 8_67341619_67345688 | RRS1-AS1 | 100505676 | 2.61172330800734 | 0.03952046 | 1 | only_H3vsUI | UpDown | notDM | Loss | 6 |
| **8_67350636_67352784_100505676** | 8_67350636_67352784 | RRS1-AS1 | 100505676 | 3.97432743542362 | 0.03965478 | 2 | only_H3vsUI | UpDown | notDM | Loss | 3 |
| **8_67375073_67376938_254778** | 8_67375073_67376938 | VXN | 254778 | 3.40414924920969 | 0.0049489 | 3 | only_H3vsUI | UpDown | notDM | Loss | 3 |
| **8_71125605_71129063_10499** | 8_71125605_71129063 | NCOA2 | 10499 | 5.27179051006231 | 0.76074003 | 6 | notSpecific | UpDown | DM | Gain | 9 |
| **8_72740390_72741612_9242** | 8_72740390_72741612 | MSC | 9242 | 4.17874679652896 | 0.79333347 | 3 | only_H3vsUI | UpDown | notDM | Gain | 1 |
| **8_72743807_72745258_9242** | 8_72743807_72745258 | MSC | 9242 | 4.05865374157237 | 0.26036603 | 7 | only_H3vsUI | UpDown | notDM | Gain | 1 |
| **8_74202402_74205564_6129** | 8_74202402_74205564 | RPL7 | 6129 | 3.04532297878666 | -0.53518493 | 1 | only_H3vsUI | DownUp | notDM | Gain | 4 |
| **8_74207457_74209754_6129** | 8_74207457_74209754 | RPL7 | 6129 | 2.89542254603941 | -0.53518493 | 2 | only_H3vsUI | DownUp | notDM | Loss | 8 |
| **8_74218295_74221439_6129** | 8_74218295_74221439 | RPL7 | 6129 | 4.06535560128996 | 0.84661772 | 3 | only_H3vsUI | DownUp | notDM | Gain | 1 |
| **8_74267695_74268774_6129** | 8_74267695_74268774 | RPL7 | 6129 | 4.79119222243299 | 0.63547914 | 3 | only_H3vsUI | DownUp | notDM | Gain | 1 |
| **8_74331201_74336270_6129** | 8_74331201_74336270 | RPL7 | 6129 | 5.09807236536218 | 0.71644183 | 3 | only_H3vsUI | DownUp | notDM | Gain | 5 |
| **8_74750439_74751430_6921** | 8_74750439_74751430 | ELOC | 6921 | 5.1229167390064 | -0.6571722 | 8 | only_H3vsUI | Up | notDM | Gain | 2 |
| **8_74872739_74873409_6921** | 8_74872739_74873409 | ELOC | 6921 | 4.03080204877227 | 0.13472052 | 8 | only_H3vsUI | Up | notDM | Gain | 1 |
| **8_74881566_74884055_6921** | 8_74881566_74884055 | ELOC | 6921 | 2.45788189673399 | -0.31120322 | 1 | only_H3vsUI | Up | notDM | Gain | 4 |
| **8_74884776_74886017_6921** | 8_74884776_74886017 | ELOC | 6921 | 2.64048143697042 | -0.34909072 | 1 | only_H3vsUI | Up | notDM | Loss | 8 |
| **8_74887142_74888286_6921** | 8_74887142_74888286 | ELOC | 6921 | 3.44762309776029 | -0.62981989 | 2 | only_H3vsUI | Up | notDM | Loss | 7 |
| **8_74888837_74890814_6921** | 8_74888837_74890814 | ELOC | 6921 | 3.65301945109961 | -0.75625574 | 1 | only_H3vsUI | Up | notDM | Gain | 4 |
| **8_76320282_76321371_3174** | 8_76320282_76321371 | HNF4G | 3174 | 2.13033376849501 | 0.08806839 | 2 | only_H3vsUI | UpDown | notDM | Loss | 3 |
| **8_80978956_80980200_7163** | 8_80978956_80980200 | TPD52 | 7163 | 3.27415784926368 | 0.00402038 | 8 | notSpecific | Down | DM | Gain | 9 |
| **8_86376210_86376681_760** | 8_86376210_86376681 | CA2 | 760 | 2.11727129565576 | 0.14790125 | 4 | notSpecific | Down | DM | Loss | 10 |
| **8_89338706_89341329_4325** | 8_89338706_89341329 | MMP16 | 4325 | 0 | 0.23776764 | 4 | only_H3vsUI | UpDown | DM | Loss | 3 |
| **8_91654492_91657819_100127983** | 8_91654492_91657819 | C8orf88 | 100127983 | 5.5310533554833 | 0.92963975 | 1 | only_DIvsD7 | DownUp | notDM | Gain | 5 |
| **8_96111565_96114543_79666** | 8_96111565_96114543 | PLEKHF2 | 79666 | 4.49817266063654 | 0.51672328 | 3 | only_DIvsD7 | DownUp | notDM | Gain | 5 |
| **8_96123831_96126104_79666** | 8_96123831_96126104 | PLEKHF2 | 79666 | 4.29948550716121 | 0.67119032 | 3 | only_DIvsD7 | DownUp | notDM | Gain | 5 |
| **8_96126385_96127733_79666** | 8_96126385_96127733 | PLEKHF2 | 79666 | 4.26245108973043 | 0.74552012 | 6 | only_DIvsD7 | DownUp | notDM | Gain | 1 |
| **8_96145083_96145812_79666** | 8_96145083_96145812 | PLEKHF2 | 79666 | 2.34439227368511 | 0.26198312 | 2 | only_DIvsD7 | DownUp | notDM | Loss | 3 |
| **8_96146652_96148493_79666** | 8_96146652_96148493 | PLEKHF2 | 79666 | 2.79379038469082 | 0.52692633 | 2 | only_DIvsD7 | DownUp | notDM | Gain | 4 |
| **8_96149563_96150453_79666** | 8_96149563_96150453 | PLEKHF2 | 79666 | 3.54814363743485 | 0.4884595 | 6 | only_DIvsD7 | DownUp | notDM | Gain | 1 |

| | | | | | | | | | | | |
|---|---|---|---|---|---|---|---|---|---|---|---|
| **8_96186638_96187892_79666** | 8_96186638_96187892 | PLEKHF2 | 79666 | 4.60861160041461 | 0.61728239 | 4 | only_DIvsD7 | DownUp | notDM | Gain | 1 |
| **8_97505186_97508108_6383** | 8_97505186_97508108 | SDC2 | 6383 | 0 | 0.65574962 | 2 | only_H3vsUI | UpDown | notDM | Loss | 3 |
| **8_99410703_99412075_6788** | 8_99410703_99412075 | STK3 | 6788 | 5.62924135386647 | 0.9555655 | 3 | only_H3vsUI | UpDown | notDM | Loss | 5 |
| **8_99835461_99837567_6788** | 8_99835461_99837567 | STK3 | 6788 | 2.52244423350632 | -0.42333109 | 2 | only_H3vsUI | UpDown | notDM | Loss | 3 |
| **8_99838387_99839550_6788** | 8_99838387_99839550 | STK3 | 6788 | 2.68124123737559 | -0.05236866 | 3 | only_H3vsUI | UpDown | notDM | Gain | 1 |
| **8_99894960_99896544_6788** | 8_99894960_99896544 | STK3 | 6788 | 4.7563546002898 | 0.95617702 | 6 | only_H3vsUI | UpDown | notDM | Loss | 5 |
| **8_99952760_99954401_6788** | 8_99952760_99954401 | STK3 | 6788 | 2.60097289568675 | 0 | 2 | only_H3vsUI | UpDown | notDM | Loss | 3 |
| **8_99954990_99956530_6788** | 8_99954990_99956530 | STK3 | 6788 | 0 | 0 | 2 | only_H3vsUI | UpDown | notDM | Loss | 3 |
| **8_99956766_99958903_6788** | 8_99956766_99958903 | STK3 | 6788 | 3.23375736296551 | 0 | 1 | only_H3vsUI | UpDown | notDM | Loss | 6 |
| **8_99978092_99979432_6788** | 8_99978092_99979432 | STK3 | 6788 | 5.14670154597199 | 0.96496254 | 3 | only_H3vsUI | UpDown | notDM | Gain | 1 |
| **9_100638591_100640855_51531** | 9_100638591_100640855 | TRMO | 51531 | 4.64343293537925 | 0.69733998 | 7 | only_H3vsUI | UpDown | notDM | Gain | 1 |
| **9_100683268_100684701_51531** | 9_100683268_100684701 | TRMO | 51531 | 1.73239375982297 | 0.64893961 | 5 | only_H3vsUI | UpDown | notDM | Gain | 4 |
| **9_100684887_100687276_51531** | 9_100684887_100687276 | TRMO | 51531 | 2.13033376849501 | 0.64893961 | 2 | only_H3vsUI | UpDown | notDM | Gain | 4 |
| **9_100743817_100745366_51531** | 9_100743817_100745366 | TRMO | 51531 | 4.77119783731714 | 0.69436598 | 1 | only_H3vsUI | UpDown | notDM | Gain | 1 |
| **9_100747461_100748745_51531** | 9_100747461_100748745 | TRMO | 51531 | 4.79720519743536 | 0.69426531 | 1 | only_H3vsUI | UpDown | notDM | Gain | 1 |
| **9_102580604_102581823_8013** | 9_102580604_102581823 | NR4A3 | 8013 | 3.36791473879375 | 0.63648843 | 2 | only_H3vsUI | Up | notDM | Gain | 4 |
| **9_102582532_102584043_8013** | 9_102582532_102584043 | NR4A3 | 8013 | 1.97772360528885 | 0.47786814 | 2 | only_H3vsUI | Up | notDM | Loss | 6 |
| **9_102584244_102587059_27130** | 9_102584244_102587059 | INVS | 27130 | 5.43851070504812 | 0.92620284 | 2 | notSpecific | Down | DM | Loss | 10 |
| **9_102584244_102587059_8013** | 9_102584244_102587059 | NR4A3 | 8013 | 2.03742649794062 | 0.52464256 | 2 | only_H3vsUI | Up | DM | Loss | 10 |
| **9_102668226_102668777_55014** | 9_102668226_102668777 | STX17 | 55014 | 2.33243845991561 | -0.09217475 | 4 | only_H3vsUI | Down | notDM | Loss | 8 |
| **9_102669338_102671061_55014** | 9_102669338_102671061 | STX17 | 55014 | 2.54282542695918 | 0.02623129 | 2 | only_H3vsUI | Down | notDM | Gain | 4 |
| **9_102669338_102671061_8013** | 9_102669338_102671061 | NR4A3 | 8013 | 4.9304548865567 | -0.75850752 | 2 | only_H3vsUI | Up | notDM | Gain | 2 |
| **9_103111906_103113598_54881** | 9_103111906_103113598 | TEX10 | 54881 | 3.19534605834842 | -0.26598978 | 8 | only_DIvsD7 | Up | notDM | Loss | 7 |
| **9_103113736_103114875_54881** | 9_103113736_103114875 | TEX10 | 54881 | 2.46389298898591 | -0.37033067 | 1 | only_DIvsD7 | Up | notDM | Loss | 7 |
| **9_103115342_103116413_54881** | 9_103115342_103116413 | TEX10 | 54881 | 2.25285303097989 | -0.37033067 | 2 | only_DIvsD7 | Up | notDM | Loss | 7 |
| **9_103141400_103143179_54881** | 9_103141400_103143179 | TEX10 | 54881 | 4.41798623068638 | -0.67442744 | 7 | only_DIvsD7 | Up | notDM | Gain | 5 |
| **9_103141400_103143179_55014** | 9_103141400_103143179 | STX17 | 55014 | 5.67431724356984 | -0.8443336 | 7 | only_H3vsUI | Down | notDM | Gain | 2 |
| **9_103189942_103193563_54881** | 9_103189942_103193563 | TEX10 | 54881 | 4.87345429974895 | -0.55952612 | 1 | only_DIvsD7 | Up | notDM | Loss | 7 |
| **9_103234729_103235246_54881** | 9_103234729_103235246 | TEX10 | 54881 | 5.07760770045816 | 0.66545858 | 4 | only_DIvsD7 | Up | notDM | Loss | 3 |
| **9_104127029_104132496_570** | 9_104127029_104132496 | BAAT | 570 | 3.10105935490812 | 0.00195999 | 7 | only_DIvsD7 | Up | notDM | Gain | 5 |
| **9_104144946_104146770_570** | 9_104144946_104146770 | BAAT | 570 | 0 | 0.2320528 | 3 | only_DIvsD7 | Up | notDM | Gain | 4 |
| **9_104149908_104150934_570** | 9_104149908_104150934 | BAAT | 570 | 3.61373614126187 | 0.23227758 | 7 | only_DIvsD7 | Up | notDM | Gain | 1 |
| **9_104159385_104160648_570** | 9_104159385_104160648 | BAAT | 570 | 4.13309161025471 | -0.21564902 | 2 | only_DIvsD7 | Up | notDM | Gain | 7 |
| **9_104161270_104162845_570** | 9_104161270_104162845 | BAAT | 570 | 4.18951838612638 | -0.18057266 | 1 | only_DIvsD7 | Up | notDM | Loss | 7 |

| | | | | | | | | | | | |
|---|---|---|---|---|---|---|---|---|---|---|---|
| **9_112954975_112956763_7295** | 9_112954975_112956763 | TXN | 7295 | 4.79349703212913 | -0.57838047 | 6 | notSpecific | Up | DM | Gain | 9 |
| **9_114118073_114120174_22949** | 9_114118073_114120174 | PTGR1 | 22949 | 5.38345988868923 | 0.91995531 | 3 | only_H3vsUI | DownUp | notDM | Loss | 5 |
| **9_114243580_114245512_22949** | 9_114243580_114245512 | PTGR1 | 22949 | 5.06619170203592 | -0.91985195 | 1 | only_H3vsUI | DownUp | notDM | Loss | 7 |
| **9_114246705_114248107_22949** | 9_114246705_114248107 | PTGR1 | 22949 | 5.0553056880186 | -0.79874909 | 1 | only_H3vsUI | DownUp | notDM | Gain | 2 |
| **9_114286607_114288791_22949** | 9_114286607_114288791 | PTGR1 | 22949 | 4.86513952252993 | 0.86550406 | 2 | only_H3vsUI | DownUp | notDM | Loss | 6 |
| **9_114357532_114361144_22949** | 9_114357532_114361144 | PTGR1 | 22949 | 2.92012332629072 | 0.59201036 | 1 | only_H3vsUI | DownUp | notDM | Gain | 4 |
| **9_114361858_114365797_22949** | 9_114361858_114365797 | PTGR1 | 22949 | 0 | 0.64455255 | 5 | only_H3vsUI | DownUp | notDM | Gain | 4 |
| **9_114366378_114368316_22949** | 9_114366378_114368316 | PTGR1 | 22949 | 3.63184946215982 | 0.66447435 | 7 | only_H3vsUI | DownUp | notDM | Gain | 5 |
| **9_114392193_114393434_22949** | 9_114392193_114393434 | PTGR1 | 22949 | 4.48441373771668 | -0.81663106 | 2 | only_H3vsUI | DownUp | notDM | Gain | 2 |
| **9_114393818_114396562_22949** | 9_114393818_114396562 | PTGR1 | 22949 | 4.50695127403032 | -0.89373121 | 1 | only_H3vsUI | DownUp | notDM | Loss | 7 |
| **9_114424257_114425866_22949** | 9_114424257_114425866 | PTGR1 | 22949 | 4.79638003663138 | -0.57371483 | 1 | only_H3vsUI | DownUp | notDM | Loss | 7 |
| **9_114489890_114491984_22949** | 9_114489890_114491984 | PTGR1 | 22949 | 5.10692826547611 | 0.78223457 | 6 | only_H3vsUI | DownUp | notDM | Gain | 5 |
| **9_114567755_114569005_22949** | 9_114567755_114569005 | PTGR1 | 22949 | 5.31340738382506 | 0.78583611 | 6 | only_H3vsUI | DownUp | notDM | Loss | 5 |
| **9_115249818_115251770_158405** | 9_115249818_115251770 | KIAA1958 | 158405 | 2.72015930340596 | -0.01337377 | 2 | only_H3vsUI | UpDown | notDM | Gain | 4 |
| **9_115981442_115983486_1317** | 9_115981442_115983486 | SLC31A1 | 1317 | 2.58206336291171 | 0.13002717 | 1 | only_DIvsD7 | DownUp | notDM | Loss | 8 |
| **9_115983908_115986131_1317** | 9_115983908_115986131 | SLC31A1 | 1317 | 1.63346845557959 | 0.13002717 | 1 | only_DIvsD7 | DownUp | notDM | Loss | 8 |
| **9_116101176_116102421_114987** | 9_116101176_116102421 | WDR31 | 114987 | 2.11058971029925 | 0.13384736 | 1 | only_DIvsD7 | UpDown | notDM | Gain | 4 |
| **9_116102685_116103692_114987** | 9_116102685_116103692 | WDR31 | 114987 | 2.03742649794062 | 0.13219841 | 1 | only_DIvsD7 | UpDown | notDM | Gain | 4 |
| **9_116111886_116112743_114987** | 9_116111886_116112743 | WDR31 | 114987 | 3.97099736557249 | -0.06014578 | 4 | only_DIvsD7 | UpDown | notDM | Loss | 3 |
| **9_116111886_116112743_1317** | 9_116111886_116112743 | SLC31A1 | 1317 | 5.10723711220479 | -0.68618358 | 4 | only_DIvsD7 | DownUp | notDM | Loss | 7 |
| **9_116225004_116228722_1317** | 9_116225004_116228722 | SLC31A1 | 1317 | 5.38224404176581 | 0.78558729 | 2 | only_DIvsD7 | DownUp | notDM | Loss | 6 |
| **9_116280840_116284230_1317** | 9_116280840_116284230 | SLC31A1 | 1317 | 5.47275644931721 | -0.77961374 | 7 | only_DIvsD7 | DownUp | notDM | Gain | 5 |
| **9_116637528_116638123_114991** | 9_116637528_116638123 | ZNF618 | 114991 | 2.70586371228392 | 0.31399813 | 4 | only_H3vsUI | Down | notDM | Gain | 4 |
| **9_116639076_116641312_114991** | 9_116639076_116641312 | ZNF618 | 114991 | 2.65127801399814 | 0.44435011 | 2 | only_H3vsUI | Down | notDM | Gain | 4 |
| **9_116717141_116718418_114991** | 9_116717141_116718418 | ZNF618 | 114991 | 4.89494157216981 | 0.81438786 | 6 | only_H3vsUI | Down | notDM | Loss | 5 |
| **9_116718544_116721529_114991** | 9_116718544_116721529 | ZNF618 | 114991 | 4.9026337382134 | 0.6658765 | 8 | only_H3vsUI | Down | notDM | Gain | 1 |
| **9_116734891_116736877_114991** | 9_116734891_116736877 | ZNF618 | 114991 | 4.98345939217558 | 0.81212002 | 6 | only_H3vsUI | Down | notDM | Loss | 5 |
| **9_122130554_122132357_1620** | 9_122130554_122132357 | BRINP1 | 1620 | 0 | 0.30399393 | 4 | notSpecific | Down | DM | Loss | 10 |
| **9_123448517_123463110_1955** | 9_123448517_123463110 | MEGF9 | 1955 | 4.13478252938186 | 0.64335574 | 7 | only_DIvsD7 | UpDown | notDM | Gain | 5 |
| **9_123464205_123466172_1955** | 9_123464205_123466172 | MEGF9 | 1955 | 4.02436250435328 | -0.17425845 | 7 | only_DIvsD7 | UpDown | notDM | Gain | 5 |
| **9_123466779_123475919_1955** | 9_123466779_123475919 | MEGF9 | 1955 | 2.91907809237607 | 0.06492338 | 1 | only_DIvsD7 | UpDown | notDM | Gain | 4 |
| **9_123477017_123478854_1955** | 9_123477017_123478854 | MEGF9 | 1955 | 2.43296929087441 | 0.02641457 | 1 | only_DIvsD7 | UpDown | notDM | Gain | 4 |
| **9_123833362_123835766_51552** | 9_123833362_123835766 | RAB14 | 51552 | 5.10850413706097 | 0.80024233 | 4 | only_DIvsD7 | UpDown | notDM | Gain | 5 |
| **9_123837392_123838641_51552** | 9_123837392_123838641 | RAB14 | 51552 | 5.09866794873812 | 0.64693511 | 2 | only_DIvsD7 | UpDown | notDM | Gain | 5 |

| | | | | | | | | | | | |
|---|---|---|---|---|---|---|---|---|---|---|---|
| **9_123944216_123946956_51552** | 9_123944216_123946956 | RAB14 | 51552 | 4.23532640248339 | -0.49365089 | 8 | only_DIvsD7 | UpDown | notDM | Gain | 5 |
| **9_123958272_123963549_51552** | 9_123958272_123963549 | RAB14 | 51552 | 2.77742682238931 | 0.27436086 | 1 | only_DIvsD7 | UpDown | notDM | Gain | 4 |
| **9_123964439_123965121_51552** | 9_123964439_123965121 | RAB14 | 51552 | 2.46834733041216 | 0.35391481 | 1 | only_DIvsD7 | UpDown | notDM | Loss | 7 |
| **9_124029498_124030283_51552** | 9_124029498_124030283 | RAB14 | 51552 | 4.81526552846654 | 0.60749378 | 2 | only_DIvsD7 | UpDown | notDM | Loss | 7 |
| **9_124030491_124033139_51552** | 9_124030491_124033139 | RAB14 | 51552 | 4.82181474437764 | 0.83919034 | 1 | only_DIvsD7 | UpDown | notDM | Loss | 6 |
| **9_124119075_124119574_51552** | 9_124119075_124119574 | RAB14 | 51552 | 5.19013552087703 | 0.77004404 | 6 | only_DIvsD7 | UpDown | notDM | Gain | 1 |
| **9_127631720_127634242_11224** | 9_127631720_127634242 | RPL35 | 11224 | 3.87546641586639 | -0.21247568 | 1 | notSpecific | Up | DM | Loss | 10 |
| **9_127631720_127634242_2649** | 9_127631720_127634242 | NR6A1 | 2649 | 4.99196109502514 | -0.807311 | 1 | notSpecific | DownUp | DM | Loss | 10 |
| **9_127631720_127634242_81873** | 9_127631720_127634242 | ARPC5L | 81873 | 2.37657695705651 | 0.1036514 | 1 | notSpecific | UpDown | DM | Loss | 10 |
| **9_129089553_129090818_89853** | 9_129089553_129090818 | MVB12B | 89853 | 2.6170003411209 | 0.04950876 | 1 | only_DIvsD7 | UpDown | notDM | Gain | 4 |
| **9_129181794_129183979_89853** | 9_129181794_129183979 | MVB12B | 89853 | 2.19033169817029 | 0 | 6 | only_DIvsD7 | UpDown | notDM | Loss | 3 |
| **9_129236379_129238095_89853** | 9_129236379_129238095 | MVB12B | 89853 | 3.69879625179043 | 0 | 3 | only_DIvsD7 | UpDown | notDM | Gain | 1 |
| **9_130073076_130074663_27433** | 9_130073076_130074663 | TOR2A | 27433 | 5.62624898138046 | 0.80847297 | 3 | only_DIvsD7 | UpDown | notDM | Gain | 1 |
| **9_130324363_130330961_27433** | 9_130324363_130330961 | TOR2A | 27433 | 5.22165935593142 | 0.73133882 | 1 | only_DIvsD7 | UpDown | notDM | Loss | 6 |
| **9_130373563_130374333_27433** | 9_130373563_130374333 | TOR2A | 27433 | 5.09082216339466 | 0.91099692 | 2 | only_DIvsD7 | UpDown | notDM | Gain | 1 |
| **9_130399432_130400389_27433** | 9_130399432_130400389 | TOR2A | 27433 | 4.9877020078572 | 0.69227273 | 6 | only_DIvsD7 | UpDown | notDM | Gain | 1 |
| **9_130476439_130477142_27433** | 9_130476439_130477142 | TOR2A | 27433 | 4.30990683556868 | 0.63567527 | 2 | only_DIvsD7 | UpDown | notDM | Loss | 3 |
| **9_130477330_130477834_27433** | 9_130477330_130477834 | TOR2A | 27433 | 4.29530314463715 | 0.43033205 | 1 | only_DIvsD7 | UpDown | notDM | Gain | 2 |
| **9_130478100_130479013_27433** | 9_130478100_130479013 | TOR2A | 27433 | 4.26913919679218 | 0.69878733 | 1 | only_DIvsD7 | UpDown | notDM | Loss | 3 |
| **9_130483216_130483726_27433** | 9_130483216_130483726 | TOR2A | 27433 | 4.14138705486734 | 0.83539619 | 3 | only_DIvsD7 | UpDown | notDM | Loss | 5 |
| **9_130483874_130484990_27433** | 9_130483874_130484990 | TOR2A | 27433 | 4.09981870961523 | 0.8379308 | 6 | only_DIvsD7 | UpDown | notDM | Loss | 5 |
| **9_130496250_130497554_27433** | 9_130496250_130497554 | TOR2A | 27433 | 0 | 0.86651092 | 1 | only_DIvsD7 | UpDown | notDM | Loss | 6 |
| **9_130497772_130498432_27433** | 9_130497772_130498432 | TOR2A | 27433 | 2.23044892137827 | 0.91052612 | 4 | only_DIvsD7 | UpDown | notDM | Gain | 1 |
| **9_130502027_130503101_27433** | 9_130502027_130503101 | TOR2A | 27433 | 3.64894518216567 | 0.5325849 | 6 | only_DIvsD7 | UpDown | notDM | Gain | 1 |
| **9_130503266_130505537_27433** | 9_130503266_130505537 | TOR2A | 27433 | 3.75694023604672 | 0.60157154 | 2 | only_DIvsD7 | UpDown | notDM | Loss | 6 |
| **9_130516565_130517229_27433** | 9_130516565_130517229 | TOR2A | 27433 | 4.27905064819903 | 0.36803181 | 4 | only_DIvsD7 | UpDown | notDM | Loss | 3 |
| **9_130533959_130534515_27433** | 9_130533959_130534515 | TOR2A | 27433 | 4.56095818603873 | 0.40900725 | 2 | only_DIvsD7 | UpDown | notDM | Gain | 2 |
| **9_130562427_130565013_27433** | 9_130562427_130565013 | TOR2A | 27433 | 4.81179611805981 | 0.64428108 | 1 | only_DIvsD7 | UpDown | notDM | Loss | 6 |
| **9_130565492_130567192_27433** | 9_130565492_130567192 | TOR2A | 27433 | 4.83185698191296 | 0.64428108 | 1 | only_DIvsD7 | UpDown | notDM | Gain | 5 |
| **9_130639944_130640850_27433** | 9_130639944_130640850 | TOR2A | 27433 | 5.15348559004906 | 0.77322592 | 1 | only_DIvsD7 | UpDown | notDM | Loss | 3 |
| **9_130660413_130661622_27433** | 9_130660413_130661622 | TOR2A | 27433 | 5.21181709712235 | 0.75325233 | 1 | only_DIvsD7 | UpDown | notDM | Loss | 3 |
| **9_130691850_130695594_27433** | 9_130691850_130695594 | TOR2A | 27433 | 5.28846833022699 | 0.76429515 | 5 | only_DIvsD7 | UpDown | notDM | Gain | 5 |
| **9_130714066_130719848_27433** | 9_130714066_130719848 | TOR2A | 27433 | 5.33548598348924 | 0.77585906 | 8 | only_DIvsD7 | UpDown | notDM | Loss | 5 |
| **9_130733944_130742730_138429** | 9_130733944_130742730 | PIP5KL1 | 138429 | 4.64623703495538 | -0.81275402 | 1 | notSpecific | Up | DM | Loss | 10 |

| | | | | | | | | | | | |
|---|---|---|---|---|---|---|---|---|---|---|---|
| **9_130733944_130742730_203245** | 9_130733944_130742730 | NAIF1 | 203245 | 4.93887482084512 | 0.66616581 | 1 | notSpecific | Down | DM | Loss | 10 |
| **9_130733944_130742730_2356** | 9_130733944_130742730 | FPGS | 2356 | 5.22735185912501 | -0.74938421 | 1 | notSpecific | DownUp | DM | Loss | 10 |
| **9_130733944_130742730_30815** | 9_130733944_130742730 | ST6GALNAC6 | 30815 | 4.85812797401624 | -0.87363583 | 1 | notSpecific | Up | DM | Loss | 10 |
| **9_130733944_130742730_399665** | 9_130733944_130742730 | FAM102A | 399665 | 1.79934054945358 | -0.04830663 | 1 | notSpecific | Down | DM | Loss | 10 |
| **9_131987191_131988829_10050611** | 9_131987191_131988829 | LINC01503 | 100506119 | 5.03083036700478 | 0.63809565 | 3 | only_H3vsUI | UpDown | notDM | Loss | 5 |
| **9_132096818_132098479_10050611** | 9_132096818_132098479 | LINC01503 | 100506119 | 2.80277372529198 | 0.36330635 | 7 | only_H3vsUI | UpDown | notDM | Gain | 1 |
| **9_132106049_132106820_10050611** | 9_132106049_132106820 | LINC01503 | 100506119 | 3.99414111112612 | 0.58241575 | 7 | only_H3vsUI | UpDown | notDM | Loss | 5 |
| **9_132227040_132228952_10050611** | 9_132227040_132228952 | LINC01503 | 100506119 | 5.11679695953272 | 0.94344205 | 7 | only_H3vsUI | UpDown | notDM | Loss | 5 |
| **9_132381918_132382543_375759** | 9_132381918_132382543 | C9orf50 | 375759 | 2.71011736511182 | 0 | 2 | only_DIvsD7 | UpDown | notDM | Gain | 1 |
| **9_132716764_132717938_23048** | 9_132716764_132717938 | FNBP1 | 23048 | 4.46861314362982 | 0.61679335 | 7 | only_H3vsUI | Down | notDM | Gain | 1 |
| **9_132785248_132788617_23048** | 9_132785248_132788617 | FNBP1 | 23048 | 4.22595507289601 | 0.53477733 | 6 | only_H3vsUI | Down | notDM | Gain | 1 |
| **9_132798344_132799077_23048** | 9_132798344_132799077 | FNBP1 | 23048 | 3.80597635071756 | 0.73875332 | 6 | only_H3vsUI | Down | notDM | Loss | 1 |
| **9_132803324_132805043_23048** | 9_132803324_132805043 | FNBP1 | 23048 | 2.60097289568675 | 0.61005362 | 1 | only_H3vsUI | Down | notDM | Loss | 8 |
| **9_132805641_132806757_23048** | 9_132805641_132806757 | FNBP1 | 23048 | 2.30535136944662 | 0.73352921 | 1 | only_H3vsUI | Down | notDM | Loss | 8 |
| **9_132814107_132816034_23048** | 9_132814107_132816034 | FNBP1 | 23048 | 3.93631263366219 | 0.13324396 | 1 | only_H3vsUI | Down | notDM | Gain | 4 |
| **9_132816525_132817949_23048** | 9_132816525_132817949 | FNBP1 | 23048 | 4.04351946024576 | 0.13324396 | 2 | only_H3vsUI | Down | notDM | Loss | 8 |
| **9_135904057_135905690_6837** | 9_135904057_135905690 | MED22 | 6837 | 5.49022263002341 | -0.86562212 | 1 | only_DIvsD7 | UpDown | notDM | Loss | 7 |
| **9_135906523_135908368_6837** | 9_135906523_135908368 | MED22 | 6837 | 5.48644464810068 | -0.86562212 | 1 | only_DIvsD7 | UpDown | notDM | Loss | 7 |
| **9_136089431_136090415_29989** | 9_136089431_136090415 | OBP2B | 29989 | 3.68151258663896 | 0 | 6 | only_H3vsUI | UpDown | notDM | Gain | 1 |
| **9_136200818_136202870_6837** | 9_136200818_136202870 | MED22 | 6837 | 4.07947067923521 | 0.30225821 | 1 | only_DIvsD7 | UpDown | notDM | Loss | 6 |
| **9_136203125_136203729_6837** | 9_136203125_136203729 | MED22 | 6837 | 4.04929556833272 | -0.27295125 | 4 | only_DIvsD7 | UpDown | notDM | Gain | 2 |
| **9_136212913_136214620_6837** | 9_136212913_136214620 | MED22 | 6837 | 2.41161970596323 | 0.14339231 | 1 | only_DIvsD7 | UpDown | notDM | Loss | 6 |
| **9_136215431_136217960_6837** | 9_136215431_136217960 | MED22 | 6837 | 2.74507479158206 | 0.36490999 | 1 | only_DIvsD7 | UpDown | notDM | Gain | 4 |
| **9_136221914_136223060_6837** | 9_136221914_136223060 | MED22 | 6837 | 3.84751096520325 | -0.64688163 | 2 | only_DIvsD7 | UpDown | notDM | Gain | 2 |
| **9_136240936_136242492_169436** | 9_136240936_136242492 | STKLD1 | 169436 | 2.8733206018154 | -0.01460239 | 1 | only_DIvsD7 | DownUp | notDM | Loss | 6 |
| **9_136243266_136245687_169436** | 9_136243266_136245687 | STKLD1 | 169436 | 0 | 0.01213115 | 1 | only_DIvsD7 | DownUp | notDM | Loss | 3 |
| **9_136243266_136245687_6837** | 9_136243266_136245687 | MED22 | 6837 | 4.45318068970209 | -0.46412683 | 1 | only_DIvsD7 | UpDown | notDM | Loss | 7 |
| **9_136281177_136282941_6837** | 9_136281177_136282941 | MED22 | 6837 | 4.82086454972976 | -0.80819001 | 1 | only_DIvsD7 | UpDown | notDM | Loss | 7 |
| **9_136283248_136284450_6837** | 9_136283248_136284450 | MED22 | 6837 | 4.83424262551524 | -0.63772493 | 2 | only_DIvsD7 | UpDown | notDM | Gain | 2 |
| **9_136293108_136293644_6837** | 9_136293108_136293644 | MED22 | 6837 | 4.89278447134719 | 0.93090984 | 4 | only_DIvsD7 | UpDown | notDM | Loss | 3 |
| **9_136373995_136375787_6837** | 9_136373995_136375787 | MED22 | 6837 | 5.20172477011638 | 0.82656965 | 4 | only_DIvsD7 | UpDown | notDM | Gain | 1 |
| **9_136995753_136996701_11091** | 9_136995753_136996701 | WDR5 | 11091 | 3.65417654187796 | 0.82584987 | 6 | notSpecific | Down | DM | Loss | 10 |
| **9_139872190_139872836_401562** | 9_139872190_139872836 | LCNL1 | 401562 | 3.66370092538965 | 0 | 2 | only_H3vsUI | UpDown | notDM | Gain | 1 |
| **9_140062037_140062953_10050599** | 9_140062037_140062953 | TMEM210 | 100505993 | 3.48869169831694 | 0 | 4 | notSpecific | Down | DM | Loss | 10 |

| | | | | | | | | | | | |
|---|---|---|---|---|---|---|---|---|---|---|---|
| **9_140062037_140062953_10383** | 9_140062037_140062953 | TUBB4B | 10383 | 4.86161206317733 | 0.77481403 | 4 | notSpecific | Up | DM | Loss | 10 |
| **9_140062037_140062953_116225** | 9_140062037_140062953 | ZMYND19 | 116225 | 5.6253021595013 | 0.84949904 | 4 | notSpecific | Down | DM | Loss | 10 |
| **9_140062037_140062953_27158** | 9_140062037_140062953 | NDOR1 | 27158 | 4.57015750766483 | 0.89453959 | 4 | notSpecific | Down | DM | Loss | 10 |
| **9_140062037_140062953_2902** | 9_140062037_140062953 | GRIN1 | 2902 | 0 | 0 | 4 | notSpecific | Up | DM | Loss | 10 |
| **9_140062037_140062953_29882** | 9_140062037_140062953 | ANAPC2 | 29882 | 4.3018326981844 | 0.57568025 | 4 | notSpecific | DownUp | DM | Loss | 10 |
| **9_140062037_140062953_375791** | 9_140062037_140062953 | CYSRT1 | 375791 | 4.75352177845229 | 0.67045113 | 4 | notSpecific | UpDown | DM | Loss | 10 |
| **9_140062037_140062953_389816** | 9_140062037_140062953 | LRRC26 | 389816 | 3.19061179781361 | -0.00560108 | 4 | notSpecific | Down | DM | Loss | 10 |
| **9_140062037_140062953_7186** | 9_140062037_140062953 | TRAF2 | 7186 | 5.44881294851972 | 0.86179165 | 4 | notSpecific | UpDown | DM | Loss | 10 |
| **9_140062037_140062953_85026** | 9_140062037_140062953 | ARRDC1-AS1 | 85026 | 5.65351448470837 | 0.85455644 | 4 | notSpecific | Up | DM | Loss | 10 |
| **9_140062037_140062953_8636** | 9_140062037_140062953 | SSNA1 | 8636 | 4.30548034865321 | 0.61635034 | 4 | notSpecific | UpDown | DM | Loss | 10 |
| **9_140062037_140062953_94107** | 9_140062037_140062953 | TMEM203 | 94107 | 4.56981851188756 | 0.65167403 | 4 | notSpecific | Down | DM | Loss | 10 |
| **9_140117105_140117846_10383** | 9_140117105_140117846 | TUBB4B | 10383 | 4.25090769970086 | 0.2104281 | 4 | notSpecific | Up | DM | Loss | 10 |
| **9_140117105_140117846_27158** | 9_140117105_140117846 | NDOR1 | 27158 | 4.23014225231864 | 0.26760526 | 4 | notSpecific | Down | DM | Loss | 10 |
| **9_140117105_140117846_375791** | 9_140117105_140117846 | CYSRT1 | 375791 | 3.25503116334555 | 0.05530989 | 4 | notSpecific | UpDown | DM | Loss | 10 |
| **9_140117105_140117846_441478** | 9_140117105_140117846 | NRARP | 441478 | 4.89684575828148 | -0.76296616 | 4 | notSpecific | Up | DM | Loss | 10 |
| **9_140117105_140117846_643596** | 9_140117105_140117846 | RNF224 | 643596 | 3.62044838471171 | 0.00315988 | 4 | notSpecific | DownUp | DM | Loss | 10 |
| **9_140500202_140502123_116225** | 9_140500202_140502123 | ZMYND19 | 116225 | 4.18361144921843 | -0.18134197 | 2 | notSpecific | Down | DM | Loss | 10 |
| **9_140500202_140502123_79813** | 9_140500202_140502123 | EHMT1 | 79813 | 4.0547279320822 | -0.74801063 | 2 | notSpecific | Down | DM | Loss | 10 |
| **9_140500202_140502123_85026** | 9_140500202_140502123 | ARRDC1-AS1 | 85026 | 4.04700213051851 | -0.09684085 | 2 | notSpecific | Up | DM | Loss | 10 |
| **9_140500202_140502123_92714** | 9_140500202_140502123 | ARRDC1 | 92714 | 1.82607480270083 | -0.23544681 | 2 | notSpecific | Down | DM | Loss | 10 |
| **9_140500202_140502123_92715** | 9_140500202_140502123 | DPH7 | 92715 | 4.42917067879397 | -0.60964751 | 2 | notSpecific | Down | DM | Loss | 10 |
| **9_20682771_20684072_101929548** | 9_20682771_20684072 | FOCAD-AS1 | 101929548 | 0 | 0 | 4 | only_H3vsUI | DownUp | notDM | Loss | 3 |
| **9_20684383_20686706_101929548** | 9_20684383_20686706 | FOCAD-AS1 | 101929548 | 0 | 0.00763402 | 1 | only_H3vsUI | DownUp | notDM | Gain | 4 |
| **9_21560022_21573837_1030** | 9_21560022_21573837 | CDKN2B | 1030 | 5.63893231076784 | 0.87828932 | 5 | only_H3vsUI | DownUp | notDM | Gain | 5 |
| **9_21574491_21591722_1030** | 9_21574491_21591722 | CDKN2B | 1030 | 5.6207178488354 | 0.82977424 | 5 | only_H3vsUI | DownUp | notDM | Gain | 5 |
| **9_21591922_21594488_1030** | 9_21591922_21594488 | CDKN2B | 1030 | 5.61783141867403 | 0.78964923 | 7 | only_H3vsUI | DownUp | DM | Gain | 9 |
| **9_21591922_21594488_554202** | 9_21591922_21594488 | MIR31HG | 554202 | 4.50861051042941 | 0.84178618 | 7 | notSpecific | Down | DM | Gain | 9 |
| **9_21614500_21617242_1030** | 9_21614500_21617242 | CDKN2B | 1030 | 5.5933292727406 | 0.94786233 | 7 | only_H3vsUI | DownUp | notDM | Loss | 5 |
| **9_21674632_21691186_1030** | 9_21674632_21691186 | CDKN2B | 1030 | 5.50255684267078 | 0.78718367 | 7 | only_H3vsUI | DownUp | DM | Gain | 9 |
| **9_21674632_21691186_554202** | 9_21674632_21691186 | MIR31HG | 554202 | 5.06056942126129 | 0.81666453 | 7 | notSpecific | Down | DM | Gain | 9 |
| **9_22143688_22144631_1030** | 9_22143688_22144631 | CDKN2B | 1030 | 5.12843158109325 | 0.89653369 | 8 | only_H3vsUI | DownUp | notDM | Gain | 1 |
| **9_22147165_22149116_1030** | 9_22147165_22149116 | CDKN2B | 1030 | 5.13952332277918 | 0.8504525 | 7 | only_H3vsUI | DownUp | notDM | Gain | 1 |
| **9_22156926_22160309_1030** | 9_22156926_22160309 | CDKN2B | 1030 | 5.16922756850498 | 0.84766499 | 7 | only_H3vsUI | DownUp | notDM | Gain | 1 |
| **9_22168803_22171011_1030** | 9_22168803_22171011 | CDKN2B | 1030 | 5.20282875329443 | 0.81171772 | 7 | only_H3vsUI | DownUp | notDM | Gain | 1 |

| | | | | | | | | | | | |
|---|---|---|---|---|---|---|---|---|---|---|---|
| **9_22208394_22213026_1030** | 9_22208394_22213026 | CDKN2B | 1030 | 5.29910615923645 | 0.79454466 | 7 | only_H3vsUI | DownUp | notDM | Gain | 1 |
| **9_22237251_22239109_1030** | 9_22237251_22239109 | CDKN2B | 1030 | 5.35788341434534 | 0.88030627 | 7 | only_H3vsUI | DownUp | notDM | Gain | 1 |
| **9_26723057_26724403_79886** | 9_26723057_26724403 | CAAP1 | 79886 | 5.22634208716363 | 0.70750257 | 4 | only_H3vsUI | Down | notDM | Gain | 1 |
| **9_26890204_26892335_79886** | 9_26890204_26892335 | CAAP1 | 79886 | 2.66931688056611 | -0.66342465 | 1 | only_H3vsUI | Down | notDM | Loss | 8 |
| **9_26892861_26893513_79886** | 9_26892861_26893513 | CAAP1 | 79886 | 1.78532983501077 | -0.66350111 | 1 | only_H3vsUI | Down | notDM | Gain | 4 |
| **9_26944839_26946966_79886** | 9_26944839_26946966 | CAAP1 | 79886 | 4.71996072339248 | -0.7090282 | 1 | only_H3vsUI | Down | notDM | Loss | 2 |
| **9_26947591_26948649_79886** | 9_26947591_26948649 | CAAP1 | 79886 | 4.74215931617876 | -0.7090282 | 2 | only_H3vsUI | Down | notDM | Gain | 2 |
| **9_26954938_26956317_101929602** | 9_26954938_26956317 | IFT74-AS1 | 101929602 | 0 | -0.00135542 | 1 | only_H3vsUI | UpDown | notDM | Gain | 4 |
| **9_26954938_26956317_79886** | 9_26954938_26956317 | CAAP1 | 79886 | 4.79336425989455 | -0.80136211 | 1 | only_H3vsUI | Down | notDM | Gain | 2 |
| **9_26956476_26957955_101929602** | 9_26956476_26957955 | IFT74-AS1 | 101929602 | 2.26717172840301 | -0.00135542 | 1 | only_H3vsUI | UpDown | notDM | Gain | 4 |
| **9_26956476_26957955_79886** | 9_26956476_26957955 | CAAP1 | 79886 | 4.80398259439456 | -0.80136211 | 1 | only_H3vsUI | Down | notDM | Gain | 2 |
| **9_35514034_35524666_203260** | 9_35514034_35524666 | CCDC107 | 203260 | 5.12592346783465 | 0.70602338 | 7 | only_H3vsUI | DownUp | notDM | Loss | 5 |
| **9_35514034_35524666_7169** | 9_35514034_35524666 | TPM2 | 7169 | 5.20648855991244 | 0.73117047 | 7 | only_DIvsD7 | DownUp | notDM | Loss | 5 |
| **9_35524914_35530468_7169** | 9_35524914_35530468 | TPM2 | 7169 | 5.20257004609748 | 0.72084013 | 7 | only_DIvsD7 | DownUp | notDM | Loss | 5 |
| **9_35603068_35604027_7169** | 9_35603068_35604027 | TPM2 | 7169 | 4.9112322050662 | -0.87174432 | 1 | only_DIvsD7 | DownUp | notDM | Gain | 2 |
| **9_35604291_35605176_7169** | 9_35604291_35605176 | TPM2 | 7169 | 4.9281089592747 | -0.86714782 | 1 | only_DIvsD7 | DownUp | notDM | Loss | 7 |
| **9_35605779_35607365_203260** | 9_35605779_35607365 | CCDC107 | 203260 | 4.70768086450618 | 0.50706201 | 1 | only_H3vsUI | DownUp | notDM | Loss | 7 |
| **9_35646123_35647699_203260** | 9_35646123_35647699 | CCDC107 | 203260 | 4.02853058651133 | 0.55326766 | 4 | only_H3vsUI | DownUp | notDM | Loss | 6 |
| **9_35646123_35647699_7169** | 9_35646123_35647699 | TPM2 | 7169 | 4.57797408137696 | -0.63269272 | 4 | only_DIvsD7 | DownUp | notDM | Loss | 7 |
| **9_35651631_35653502_203260** | 9_35651631_35653502 | CCDC107 | 203260 | 3.68214507637383 | -0.26186106 | 6 | only_H3vsUI | DownUp | notDM | Gain | 1 |
| **9_35656536_35657820_203260** | 9_35656536_35657820 | CCDC107 | 203260 | 2.74663419893758 | 0.49630509 | 2 | only_H3vsUI | DownUp | notDM | Loss | 6 |
| **9_35656536_35657820_7169** | 9_35656536_35657820 | TPM2 | 7169 | 4.44280889284628 | -0.57193772 | 2 | only_DIvsD7 | DownUp | notDM | Loss | 7 |
| **9_35658460_35660435_203260** | 9_35658460_35660435 | CCDC107 | 203260 | 1.92941892571429 | 0.49630509 | 2 | only_H3vsUI | DownUp | notDM | Loss | 6 |
| **9_35658460_35660435_7169** | 9_35658460_35660435 | TPM2 | 7169 | 4.3997775244888 | -0.57193772 | 2 | only_DIvsD7 | DownUp | notDM | Loss | 7 |
| **9_35663291_35665209_203260** | 9_35663291_35665209 | CCDC107 | 203260 | 3.69827457667437 | -0.29828364 | 1 | only_H3vsUI | DownUp | notDM | Loss | 7 |
| **9_35663291_35665209_7169** | 9_35663291_35665209 | TPM2 | 7169 | 4.30818010103067 | -0.80064212 | 1 | only_DIvsD7 | DownUp | notDM | Loss | 7 |
| **9_35668254_35670929_203260** | 9_35668254_35670929 | CCDC107 | 203260 | 3.99804126436343 | -0.05093426 | 7 | only_H3vsUI | DownUp | notDM | Loss | 5 |
| **9_35672699_35674614_203260** | 9_35672699_35674614 | CCDC107 | 203260 | 4.15606431233987 | -0.34418426 | 6 | only_H3vsUI | DownUp | notDM | Loss | 7 |
| **9_35672699_35674614_7169** | 9_35672699_35674614 | TPM2 | 7169 | 4.03850094307667 | 0.17643073 | 6 | only_DIvsD7 | DownUp | notDM | Loss | 7 |
| **9_35687165_35691824_7169** | 9_35687165_35691824 | TPM2 | 7169 | 0 | 0.43377902 | 1 | only_DIvsD7 | DownUp | notDM | Loss | 8 |
| **9_35815596_35816041_7169** | 9_35815596_35816041 | TPM2 | 7169 | 5.09933873268003 | -0.84556892 | 2 | only_DIvsD7 | DownUp | notDM | Loss | 7 |
| **9_35923299_35925186_646962** | 9_35923299_35925186 | HRCT1 | 646962 | 4.23330077156347 | 0.32740584 | 7 | only_H3vsUI | UpDown | notDM | Gain | 1 |
| **9_35945179_35945878_646962** | 9_35945179_35945878 | HRCT1 | 646962 | 4.59097551184065 | 0.73709505 | 4 | only_H3vsUI | UpDown | notDM | Gain | 1 |
| **9_36136755_36137938_152007** | 9_36136755_36137938 | GLIPR2 | 152007 | 1.17609125905568 | 0.71253111 | 2 | only_H3vsUI | DownUp | notDM | Loss | 3 |

| | | | | | | | | | | | |
|---|---|---|---|---|---|---|---|---|---|---|---|
| **9_36148164_36150393_152007** | 9_36148164_36150393 | GLIPR2 | 152007 | 4.05793222738061 | 0.34525205 | 3 | only_H3vsUI | DownUp | notDM | Loss | 5 |
| **9_36258576_36260230_152007** | 9_36258576_36260230 | GLIPR2 | 152007 | 5.08581840511621 | -0.91093284 | 1 | only_H3vsUI | DownUp | notDM | Loss | 7 |
| **9_36936068_36937384_5079** | 9_36936068_36937384 | PAX5 | 5079 | 4.81484001677711 | 0.54933318 | 6 | only_H3vsUI | Up | notDM | Loss | 3 |
| **9_37025061_37027680_5079** | 9_37025061_37027680 | PAX5 | 5079 | 4.35007348995169 | 0.54940747 | 4 | only_H3vsUI | Up | notDM | Loss | 3 |
| **9_37027864_37028237_5079** | 9_37027864_37028237 | PAX5 | 5079 | 4.40129712502158 | 0.54937356 | 4 | only_H3vsUI | Up | notDM | Gain | 1 |
| **9_37034182_37037383_5079** | 9_37034182_37037383 | PAX5 | 5079 | 0 | 0.17157154 | 2 | only_H3vsUI | Up | notDM | Loss | 3 |
| **9_37462282_37464986_92014** | 9_37462282_37464986 | SLC25A51 | 92014 | 5.64282545788198 | 0.81926826 | 1 | only_DIvsD7 | UpDown | notDM | Loss | 6 |
| **9_37465960_37467473_92014** | 9_37465960_37467473 | SLC25A51 | 92014 | 5.64036017535922 | 0.81926826 | 5 | only_DIvsD7 | UpDown | notDM | Gain | 5 |
| **9_37486006_37487963_92014** | 9_37486006_37487963 | SLC25A51 | 92014 | 5.61949820496703 | 0.89553528 | 1 | only_DIvsD7 | UpDown | notDM | Loss | 6 |
| **9_37574925_37577326_51010** | 9_37574925_37577326 | EXOSC3 | 51010 | 5.3175117649345 | -0.7921918 | 1 | only_H3vsUI | Down | notDM | Loss | 2 |
| **9_37753336_37753750_51010** | 9_37753336_37753750 | EXOSC3 | 51010 | 4.49571080831393 | 0.35278628 | 4 | only_H3vsUI | Down | notDM | Loss | 2 |
| **9_37753336_37753750_92014** | 9_37753336_37753750 | SLC25A51 | 92014 | 5.17712654027441 | 0.77733881 | 4 | only_DIvsD7 | UpDown | notDM | Loss | 3 |
| **9_37753954_37755392_51010** | 9_37753954_37755392 | EXOSC3 | 51010 | 4.47221507176179 | -0.38543688 | 2 | only_H3vsUI | Down | notDM | Loss | 7 |
| **9_37782947_37784756_51010** | 9_37782947_37784756 | EXOSC3 | 51010 | 2.47567118832443 | 0.18190486 | 2 | only_H3vsUI | Down | notDM | Gain | 4 |
| **9_37799121_37800651_51010** | 9_37799121_37800651 | EXOSC3 | 51010 | 1.7481880270062 | 0.33148046 | 1 | only_H3vsUI | Down | notDM | Gain | 4 |
| **9_37801300_37803375_51010** | 9_37801300_37803375 | EXOSC3 | 51010 | 2.77524625974024 | 0.33148046 | 2 | only_H3vsUI | Down | notDM | Loss | 8 |
| **9_37901464_37903924_92014** | 9_37901464_37903924 | SLC25A51 | 92014 | 2.26481782300954 | 0.53037078 | 2 | only_DIvsD7 | UpDown | notDM | Loss | 6 |
| **9_37904389_37905376_92014** | 9_37904389_37905376 | SLC25A51 | 92014 | 1.61278385671974 | 0.47805967 | 1 | only_DIvsD7 | UpDown | notDM | Loss | 3 |
| **9_37925872_37927136_92014** | 9_37925872_37927136 | SLC25A51 | 92014 | 4.33733943938842 | 0.81367999 | 6 | only_DIvsD7 | UpDown | notDM | Loss | 5 |
| **9_37942943_37946283_92014** | 9_37942943_37946283 | SLC25A51 | 92014 | 4.58899959049944 | 0.74109455 | 7 | only_DIvsD7 | UpDown | DM | Loss | 10 |
| **9_37949484_37955529_51010** | 9_37949484_37955529 | EXOSC3 | 51010 | 5.21598633918365 | -0.70107797 | 7 | only_H3vsUI | Down | notDM | Loss | 2 |
| **9_37963361_37970443_6461** | 9_37963361_37970443 | SHB | 6461 | 4.99461625962303 | 0.66487444 | 7 | only_H3vsUI | UpDown | notDM | Loss | 5 |
| **9_38015337_38022113_6461** | 9_38015337_38022113 | SHB | 6461 | 4.67300246535807 | 0.50087251 | 7 | only_H3vsUI | UpDown | notDM | Loss | 5 |
| **9_38022386_38029999_92014** | 9_38022386_38029999 | SLC25A51 | 92014 | 5.07291498758764 | 0.75700825 | 7 | only_DIvsD7 | UpDown | notDM | Loss | 5 |
| **9_38043792_38049785_6461** | 9_38043792_38049785 | SHB | 6461 | 4.28838338439977 | 0.57750768 | 7 | only_H3vsUI | UpDown | notDM | Loss | 5 |
| **9_38056882_38059175_6461** | 9_38056882_38059175 | SHB | 6461 | 4.00156065264257 | 0.54772261 | 8 | only_H3vsUI | UpDown | notDM | Loss | 5 |
| **9_38059317_38061260_6461** | 9_38059317_38061260 | SHB | 6461 | 3.90042175345774 | 0.54042877 | 8 | only_H3vsUI | UpDown | notDM | Loss | 5 |
| **9_38061342_38068442_6461** | 9_38061342_38068442 | SHB | 6461 | 2.88592633980143 | 0.57342397 | 1 | only_H3vsUI | UpDown | notDM | Loss | 6 |
| **9_38069499_38071070_6461** | 9_38069499_38071070 | SHB | 6461 | 2.46389298898591 | 0.43272433 | 1 | only_H3vsUI | UpDown | notDM | Gain | 4 |
| **9_38118980_38120586_92014** | 9_38118980_38120586 | SLC25A51 | 92014 | 5.3316944724639 | 0.88509678 | 6 | only_DIvsD7 | UpDown | notDM | Loss | 5 |
| **9_38436817_38438880_6461** | 9_38436817_38438880 | SHB | 6461 | 5.56538613547384 | 0.84405999 | 7 | only_H3vsUI | UpDown | notDM | Loss | 5 |
| **9_38471922_38473762_6461** | 9_38471922_38473762 | SHB | 6461 | 5.6049967277377 | 0.84371748 | 6 | only_H3vsUI | UpDown | notDM | Gain | 1 |
| **9_4679809_4682015_403313** | 9_4679809_4682015 | PLPP6 | 403313 | 4.24336054767377 | -0.42028255 | 1 | notSpecific | Down | DM | Loss | 10 |
| **9_4679809_4682015_50808** | 9_4679809_4682015 | AK3 | 50808 | 4.77241706497391 | 0.61553949 | 1 | notSpecific | DownUp | DM | Loss | 10 |

| | | | | | | | | | | | |
|---|---|---|---|---|---|---|---|---|---|---|---|
| **9_4679809_4682015_55664** | 9_4679809_4682015 | CDC37L1 | 55664 | 2.38916608436453 | -0.68906299 | 1 | notSpecific | Down | DM | Loss | 10 |
| **9_74523338_74525236_138241** | 9_74523338_74525236 | C9orf85 | 138241 | 3.1300119496719 | -0.2194837 | 2 | only_H3vsUI | DownUp | notDM | Gain | 4 |
| **9_74526629_74529099_138241** | 9_74526629_74529099 | C9orf85 | 138241 | 2.23044892137827 | -0.1615685 | 2 | only_H3vsUI | DownUp | notDM | Gain | 4 |
| **9_74763860_74765291_138241** | 9_74763860_74765291 | C9orf85 | 138241 | 5.37525747306131 | 0.87196386 | 2 | only_H3vsUI | DownUp | notDM | Loss | 6 |
| **9_74920362_74922303_100507540** | 9_74920362_74922303 | LINC01504 | 100507540 | 1.27875360095283 | 0.033569 | 1 | only_H3vsUI | DownUp | notDM | Gain | 4 |
| **9_75591264_75598545_216** | 9_75591264_75598545 | ALDH1A1 | 216 | 4.3672999996814 | 0.41434969 | 7 | notSpecific | Down | DM | Gain | 9 |
| **9_75601023_75609702_216** | 9_75601023_75609702 | ALDH1A1 | 216 | 4.51925029983953 | 0.64519944 | 7 | notSpecific | Down | DM | Gain | 9 |
| **9_75610061_75612653_216** | 9_75610061_75612653 | ALDH1A1 | 216 | 4.62422019672419 | 0.80742876 | 7 | notSpecific | Down | DM | Gain | 9 |
| **9_75697202_75702009_216** | 9_75697202_75702009 | ALDH1A1 | 216 | 3.26623169668989 | -0.00582203 | 7 | notSpecific | Down | DM | Gain | 9 |
| **9_77701310_77702915_26578** | 9_77701310_77702915 | OSTF1 | 26578 | 2.73639650227664 | -0.1775307 | 2 | only_DIvsD7 | DownUp | notDM | Gain | 2 |
| **9_77703601_77705039_26578** | 9_77703601_77705039 | OSTF1 | 26578 | 2.15836249209525 | -0.35015173 | 1 | only_DIvsD7 | DownUp | notDM | Gain | 2 |
| **9_84297802_84302830_101927502** | 9_84297802_84302830 | LOC101927502 | 101927502 | 3.25503116334555 | 0 | 1 | only_H3vsUI | UpDown | notDM | Loss | 6 |
| **9_84304951_84306378_101927502** | 9_84304951_84306378 | LOC101927502 | 101927502 | 2.51188336097887 | 0 | 1 | only_H3vsUI | UpDown | notDM | Loss | 3 |
| **9_95638639_95640264_83744** | 9_95638639_95640264 | ZNF484 | 83744 | 1.43136376415899 | -0.30714765 | 1 | notSpecific | Down | DM | Gain | 9 |
| **9_95856839_95857748_101927993** | 9_95856839_95857748 | LOC101927993 | 101927993 | 1.80617997398389 | -0.00164119 | 5 | only_DIvsD7 | UpDown | notDM | Gain | 4 |
| **9_95858833_95862091_101927993** | 9_95858833_95862091 | LOC101927993 | 101927993 | 3.01029995663981 | -0.00076537 | 1 | only_DIvsD7 | UpDown | notDM | Loss | 6 |
| **9_95893772_95896393_203328** | 9_95893772_95896393 | SUSD3 | 203328 | 4.86204188633984 | 0.6358082 | 1 | only_H3vsUI | Down | notDM | Loss | 6 |
| **9_96091279_96093057_203328** | 9_96091279_96093057 | SUSD3 | 203328 | 5.43183319180575 | -0.76313944 | 6 | only_H3vsUI | Down | notDM | Gain | 1 |
| **9_98636025_98637568_100128782** | 9_98636025_98637568 | LINC00476 | 100128782 | 2.29003461136252 | -0.3464515 | 1 | only_H3vsUI | UpDown | notDM | Gain | 4 |
| **9_98638416_98640488_100128782** | 9_98638416_98640488 | LINC00476 | 100128782 | 2.81690383937566 | -0.3464515 | 1 | only_H3vsUI | UpDown | notDM | Gain | 4 |
| **X_10124318_10126915_1183** | X_10124318_10126915 | CLCN4 | 1183 | 0 | 0.29322621 | 2 | only_H3vsUI | Down | notDM | Loss | 8 |
| **X_101854737_101856014_80823** | X_101854737_101856014 | BHLHB9 | 80823 | 5.16139495760766 | -0.74079126 | 2 | only_H3vsUI | Down | notDM | Loss | 7 |
| **X_101967946_101968457_80823** | X_101967946_101968457 | BHLHB9 | 80823 | 3.85491302230786 | 0.8943359 | 4 | only_H3vsUI | Down | notDM | Loss | 2 |
| **X_101974614_101976677_80823** | X_101974614_101976677 | BHLHB9 | 80823 | 0 | 0.68439958 | 2 | only_H3vsUI | Down | notDM | Loss | 8 |
| **X_102024322_102025597_80823** | X_102024322_102025597 | BHLHB9 | 80823 | 4.36739319792248 | -0.60962011 | 1 | only_H3vsUI | Down | notDM | Loss | 7 |
| **X_102206075_102206884_80823** | X_102206075_102206884 | BHLHB9 | 80823 | 5.31187036345981 | -0.7780641 | 4 | only_H3vsUI | Down | notDM | Loss | 7 |
| **X_102469379_102470375_80823** | X_102469379_102470375 | BHLHB9 | 80823 | 5.69352030219052 | 0.82042465 | 2 | only_H3vsUI | Down | notDM | Loss | 2 |
| **X_106142005_106143412_9075** | X_106142005_106143412 | CLDN2 | 9075 | 0 | 0.04164191 | 7 | only_H3vsUI | DownUp | notDM | Gain | 4 |
| **X_106160221_106162596_9075** | X_106160221_106162596 | CLDN2 | 9075 | 0 | 0 | 7 | only_H3vsUI | DownUp | notDM | Gain | 4 |
| **X_111922691_111923809_340596** | X_111922691_111923809 | LHFPL1 | 340596 | 0 | 0.38060507 | 2 | only_H3vsUI | UpDown | notDM | Loss | 3 |
| **X_118532272_118533156_292** | X_118532272_118533156 | SLC25A5 | 292 | 4.84015629900937 | 0.59849671 | 5 | only_DIvsD7 | DownUp | notDM | Gain | 2 |
| **X_118533723_118535362_292** | X_118533723_118535362 | SLC25A5 | 292 | 4.82608776652172 | 0.68377235 | 1 | only_DIvsD7 | DownUp | notDM | Loss | 6 |
| **X_118603246_118604454_292** | X_118603246_118604454 | SLC25A5 | 292 | 2.94694327069783 | 0.5091505 | 1 | only_DIvsD7 | DownUp | notDM | Loss | 7 |
| **X_119003222_119005423_292** | X_119003222_119005423 | SLC25A5 | 292 | 5.6029699699969 | 0.80097006 | 1 | only_DIvsD7 | DownUp | notDM | Gain | 7 |

| | | | | | | | | | | | |
|---|---|---|---|---|---|---|---|---|---|---|---|
| **X_119383625_119384434_10009** | X_119383625_119384434 | ZBTB33 | 10009 | 2.2405492482826 | 0.26917841 | 2 | notSpecific | Down | DM | Gain | 9 |
| **X_119383625_119384434_158801** | X_119383625_119384434 | NKAPP1 | 158801 | 3.65369479531508 | 0.20141855 | 2 | notSpecific | Down | DM | Gain | 9 |
| **X_119383625_119384434_3920** | X_119383625_119384434 | LAMP2 | 3920 | 5.34002151329213 | 0.77623226 | 2 | notSpecific | Up | DM | Gain | 9 |
| **X_119738182_119740297_28985** | X_119738182_119740297 | MCTS1 | 28985 | 2.54282542695918 | 0.26989425 | 1 | only_H3vsUI | DownUp | notDM | Loss | 6 |
| **X_129040656_129041714_10813** | X_129040656_129041714 | UTP14A | 10813 | 2.74896286125616 | 0.25747412 | 2 | only_DIvsD7 | DownUp | notDM | Loss | 7 |
| **X_129091421_129091904_10813** | X_129091421_129091904 | UTP14A | 10813 | 4.7097447109838 | 0.68603807 | 1 | only_DIvsD7 | DownUp | notDM | Loss | 3 |
| **X_129091421_129091904_63035** | X_129091421_129091904 | BCORL1 | 63035 | 4.39239794321637 | 0.42198229 | 1 | only_DIvsD7 | DownUp | notDM | Loss | 3 |
| **X_129095222_129096060_10813** | X_129095222_129096060 | UTP14A | 10813 | 4.74081254343687 | 0.70940391 | 4 | only_DIvsD7 | DownUp | notDM | Loss | 3 |
| **X_129095222_129096060_63035** | X_129095222_129096060 | BCORL1 | 63035 | 4.31232548231513 | 0.41582556 | 4 | only_DIvsD7 | DownUp | notDM | Loss | 3 |
| **X_129114936_129115827_63035** | X_129114936_129115827 | BCORL1 | 63035 | 2.88081359228079 | 0.52488874 | 2 | only_DIvsD7 | DownUp | notDM | Loss | 3 |
| **X_129116654_129117963_10813** | X_129116654_129117963 | UTP14A | 10813 | 4.88359898309093 | 0.72820163 | 1 | only_DIvsD7 | DownUp | notDM | Loss | 7 |
| **X_129116654_129117963_63035** | X_129116654_129117963 | BCORL1 | 63035 | 1.84509804001426 | 0.53308984 | 1 | only_DIvsD7 | DownUp | notDM | Loss | 3 |
| **X_129118436_129119183_63035** | X_129118436_129119183 | BCORL1 | 63035 | 3.26764098234592 | 0.52316395 | 1 | only_DIvsD7 | DownUp | notDM | Loss | 3 |
| **X_133683870_133684837_10050675** | X_133683870_133684837 | LINC00629 | 100506757 | 0 | 0.00162213 | 3 | notSpecific | Down | DM | Loss | 10 |
| **X_133683870_133684837_8933** | X_133683870_133684837 | RTL8C | 8933 | 5.68266483512833 | 0.82909309 | 3 | notSpecific | Down | DM | Loss | 10 |
| **X_149531846_149533234_10046** | X_149531846_149533234 | MAMLD1 | 10046 | 2.47275644931721 | 0.80026891 | 2 | notSpecific | Up | DM | Loss | 10 |
| **X_149531846_149533234_8776** | X_149531846_149533234 | MTMR1 | 8776 | 5.51668341862012 | 0.84457434 | 2 | notSpecific | Down | DM | Loss | 10 |
| **X_1509293_1510421_8623** | X_1509293_1510421 | ASMTL | 8623 | 4.78777917079731 | -0.57177441 | 1 | only_DIvsD7 | DownUp | notDM | Loss | 7 |
| **X_152734789_152735807_633** | X_152734789_152735807 | BGN | 633 | 4.39077618980628 | -0.3257352 | 1 | only_DIvsD7 | UpDown | notDM | Loss | 3 |
| **X_152862893_152864127_139728** | X_152862893_152864127 | PNCK | 139728 | 4.87883189510874 | -0.6351626 | 1 | only_H3vsUI | UpDown | notDM | Loss | 3 |
| **X_152888062_152888920_139728** | X_152888062_152888920 | PNCK | 139728 | 4.70638489477426 | 0.74743197 | 6 | only_H3vsUI | UpDown | notDM | Gain | 1 |
| **X_152987363_152989172_139728** | X_152987363_152989172 | PNCK | 139728 | 4.6771413877498 | 0.56571699 | 1 | only_H3vsUI | UpDown | notDM | Loss | 3 |
| **X_153094746_153095981_57595** | X_153094746_153095981 | PDZD4 | 57595 | 0 | 0.99912716 | 4 | notSpecific | DownUp | DM | Loss | 10 |
| **X_153094746_153095981_6748** | X_153094746_153095981 | SSR4 | 6748 | 4.53975337105938 | -0.37599999 | 4 | notSpecific | Up | DM | Loss | 10 |
| **X_153150072_153151450_393** | X_153150072_153151450 | ARHGAP4 | 393 | 4.60492770342845 | -0.44882315 | 2 | only_H3vsUI | DownUp | notDM | Loss | 3 |
| **X_153150072_153151450_5973** | X_153150072_153151450 | RENBP | 5973 | 4.76856410951357 | -0.50136483 | 2 | only_H3vsUI | UpDown | notDM | Loss | 3 |
| **X_153192383_153193616_393** | X_153192383_153193616 | ARHGAP4 | 393 | 1.85125834871908 | 0.13803989 | 1 | only_H3vsUI | DownUp | notDM | Loss | 3 |
| **X_153194061_153195317_393** | X_153194061_153195317 | ARHGAP4 | 393 | 2.27875360095283 | -0.00568791 | 1 | only_H3vsUI | DownUp | notDM | Loss | 3 |
| **X_153194061_153195317_5973** | X_153194061_153195317 | RENBP | 5973 | 4.17093610860908 | -0.13373794 | 1 | only_H3vsUI | UpDown | notDM | Loss | 3 |
| **X_153198815_153200104_2010** | X_153198815_153200104 | EMD | 2010 | 5.61015958449904 | -0.80456942 | 2 | only_DIvsD7 | Up | notDM | Loss | 7 |
| **X_153198815_153200104_393** | X_153198815_153200104 | ARHGAP4 | 393 | 3.8131137540079 | 0.1169292 | 2 | only_H3vsUI | DownUp | notDM | Loss | 3 |
| **X_153198815_153200104_5973** | X_153198815_153200104 | RENBP | 5973 | 4.001733712809 | -0.24447076 | 2 | only_H3vsUI | UpDown | notDM | Loss | 3 |
| **X_153234104_153235541_2010** | X_153234104_153235541 | EMD | 2010 | 5.57065149993293 | -0.8876941 | 1 | only_DIvsD7 | Up | notDM | Loss | 7 |
| **X_153234104_153235541_5973** | X_153234104_153235541 | RENBP | 5973 | 4.37961367587827 | -0.36425139 | 1 | only_H3vsUI | UpDown | notDM | Loss | 3 |

| | | | | | | | | | | | |
|---|---|---|---|---|---|---|---|---|---|---|---|
| **X_153283696_153284648_2010** | X_153283696_153284648 | EMD | 2010 | 5.50918369801371 | -0.78030285 | 1 | only_DIvsD7 | Up | notDM | Loss | 7 |
| **X_153311102_153313716_5973** | X_153311102_153313716 | RENBP | 5973 | 5.00415364352774 | 0.64662811 | 7 | only_H3vsUI | UpDown | notDM | Loss | 5 |
| **X_153595685_153597789_2010** | X_153595685_153597789 | EMD | 2010 | 3.99321572047414 | 0.59418518 | 5 | only_DIvsD7 | Up | notDM | Gain | 5 |
| **X_153595685_153597789_5973** | X_153595685_153597789 | RENBP | 5973 | 5.58607394883772 | 0.8076469 | 5 | only_H3vsUI | UpDown | notDM | Gain | 1 |
| **X_153599978_153602864_2010** | X_153599978_153602864 | EMD | 2010 | 3.67154308526257 | -0.50818889 | 5 | only_DIvsD7 | Up | notDM | Loss | 7 |
| **X_153603202_153603653_2010** | X_153603202_153603653 | EMD | 2010 | 3.59162103821332 | -0.68375223 | 5 | only_DIvsD7 | Up | notDM | Loss | 7 |
| **X_153608033_153609322_2010** | X_153608033_153609322 | EMD | 2010 | 2.67942789661212 | -0.67793009 | 1 | only_DIvsD7 | Up | notDM | Loss | 7 |
| **X_153627362_153629042_2010** | X_153627362_153629042 | EMD | 2010 | 4.29514909657892 | -0.7994684 | 1 | only_DIvsD7 | Up | notDM | Loss | 7 |
| **X_153636396_153637628_2010** | X_153636396_153637628 | EMD | 2010 | 4.45886437836152 | -0.60524754 | 1 | only_DIvsD7 | Up | notDM | Loss | 7 |
| **X_153638943_153639756_2010** | X_153638943_153639756 | EMD | 2010 | 4.49571080831393 | -0.90711631 | 1 | only_DIvsD7 | Up | notDM | Gain | 2 |
| **X_153640010_153641732_2010** | X_153640010_153641732 | EMD | 2010 | 4.51128161346714 | -0.411482 | 1 | only_DIvsD7 | Up | notDM | Loss | 7 |
| **X_153666112_153666990_393** | X_153666112_153666990 | ARHGAP4 | 393 | 5.67615017308473 | 0.84912408 | 2 | only_H3vsUI | DownUp | notDM | Loss | 3 |
| **X_153672835_153673948_2010** | X_153672835_153673948 | EMD | 2010 | 4.8142742387479 | -0.67703593 | 1 | only_DIvsD7 | Up | notDM | Loss | 7 |
| **X_153717665_153718760_2010** | X_153717665_153718760 | EMD | 2010 | 5.04152690089309 | -0.87980275 | 1 | only_DIvsD7 | Up | notDM | Loss | 7 |
| **X_153743423_153744369_2010** | X_153743423_153744369 | EMD | 2010 | 5.13287418483258 | -0.81371122 | 1 | only_DIvsD7 | Up | notDM | Loss | 7 |
| **X_153770469_153774846_2010** | X_153770469_153774846 | EMD | 2010 | 5.21175575967213 | -0.7747464 | 5 | only_DIvsD7 | Up | notDM | Loss | 7 |
| **X_153776116_153777554_2010** | X_153776116_153777554 | EMD | 2010 | 5.22656124225659 | -0.80552481 | 1 | only_DIvsD7 | Up | notDM | Loss | 7 |
| **X_153991756_153993286_2010** | X_153991756_153993286 | EMD | 2010 | 5.58455849091067 | -0.80293663 | 1 | only_DIvsD7 | Up | notDM | Loss | 7 |
| **X_15509129_15511140_8905** | X_15509129_15511140 | AP1S2 | 8905 | 5.5584685625238 | -0.84699031 | 1 | only_H3vsUI | UpDown | notDM | Gain | 2 |
| **X_15691839_15693007_8905** | X_15691839_15693007 | AP1S2 | 8905 | 5.25511082095307 | -0.91807248 | 1 | only_H3vsUI | UpDown | notDM | Gain | 2 |
| **X_15694172_15695782_8905** | X_15694172_15695782 | AP1S2 | 8905 | 5.24836076873472 | -0.91807248 | 2 | only_H3vsUI | UpDown | notDM | Loss | 7 |
| **X_15871179_15872351_8905** | X_15871179_15872351 | AP1S2 | 8905 | 2.76789761601809 | -0.10151684 | 2 | only_H3vsUI | UpDown | notDM | Gain | 4 |
| **X_15873438_15874510_8905** | X_15873438_15874510 | AP1S2 | 8905 | 2.69983772586725 | -0.93747753 | 2 | only_H3vsUI | UpDown | notDM | Gain | 4 |
| **X_16729361_16730165_56474** | X_16729361_16730165 | CTPS2 | 56474 | 2.78604121024255 | 0.1556609 | 2 | only_H3vsUI | UpDown | notDM | Loss | 3 |
| **X_16736782_16737537_56474** | X_16736782_16737537 | CTPS2 | 56474 | 3.77880220401324 | -0.26328645 | 1 | only_H3vsUI | UpDown | notDM | Loss | 7 |
| **X_16738065_16739834_56474** | X_16738065_16739834 | CTPS2 | 56474 | 3.86284665998294 | -0.22050916 | 2 | only_H3vsUI | UpDown | notDM | Loss | 8 |
| **X_16743212_16745261_56474** | X_16743212_16745261 | CTPS2 | 56474 | 4.08475496317935 | 0.38132861 | 8 | only_H3vsUI | UpDown | notDM | Gain | 1 |
| **X_16805339_16806795_56474** | X_16805339_16806795 | CTPS2 | 56474 | 4.87254084653009 | 0.58661437 | 2 | only_H3vsUI | UpDown | notDM | Loss | 6 |
| **X_16825508_16826853_56474** | X_16825508_16826853 | CTPS2 | 56474 | 4.97651045945345 | -0.62615365 | 8 | only_H3vsUI | UpDown | notDM | Loss | 7 |
| **X_16900015_16901762_56474** | X_16900015_16901762 | CTPS2 | 56474 | 5.22777876002687 | -0.71404331 | 1 | only_H3vsUI | UpDown | notDM | Loss | 7 |
| **X_18692107_18693017_6247** | X_18692107_18693017 | RS1 | 6247 | 3.27415784926368 | 0 | 1 | only_DIvsD7 | UpDown | notDM | Loss | 3 |
| **X_21943591_21946162_6611** | X_21943591_21946162 | SMS | 6611 | 4.10284516424542 | -0.38963574 | 3 | notSpecific | DownUp | DM | Gain | 9 |
| **X_23684859_23685430_6303** | X_23684859_23685430 | SAT1 | 6303 | 5.06455157709107 | 0.6647505 | 3 | only_DIvsD7 | Up | notDM | Loss | 5 |
| **X_23685870_23687912_6303** | X_23685870_23687912 | SAT1 | 6303 | 5.05516036484301 | 0.6647505 | 2 | only_DIvsD7 | Up | notDM | Loss | 7 |

| | | | | | | | | | | | |
|---|---|---|---|---|---|---|---|---|---|---|---|
| **X_23802096_23804903_6303** | X_23802096_23804903 | SAT1 | 6303 | 0 | 0.29622938 | 1 | only_DIvsD7 | Up | notDM | Gain | 8 |
| **X_23806120_23807089_6303** | X_23806120_23807089 | SAT1 | 6303 | 3.66913084737333 | 0.67808296 | 7 | only_DIvsD7 | Up | notDM | Gain | 1 |
| **X_23820395_23823397_6303** | X_23820395_23823397 | SAT1 | 6303 | 4.27744875926448 | 0.60402089 | 7 | only_DIvsD7 | Up | notDM | Gain | 5 |
| **X_23862113_23864389_6303** | X_23862113_23864389 | SAT1 | 6303 | 4.78391072221534 | 0.65281712 | 7 | only_DIvsD7 | Up | notDM | Loss | 5 |
| **X_23914348_23920735_6303** | X_23914348_23920735 | SAT1 | 6303 | 5.05321678075253 | 0.6872546 | 7 | only_DIvsD7 | Up | notDM | Gain | 5 |
| **X_24073359_24076727_1968** | X_24073359_24076727 | EIF2S3 | 1968 | 2.43616264704076 | 0.25682754 | 1 | only_H3vsUI | DownUp | notDM | Gain | 4 |
| **X_24135156_24136222_1968** | X_24135156_24136222 | EIF2S3 | 1968 | 4.7928817453854 | 0.73429188 | 3 | only_H3vsUI | DownUp | notDM | Gain | 1 |
| **X_24511599_24513934_1968** | X_24511599_24513934 | EIF2S3 | 1968 | 5.64200723162516 | -0.84989193 | 6 | only_H3vsUI | DownUp | notDM | Loss | 7 |
| **X_30319594_30326279_4113** | X_30319594_30326279 | MAGEB2 | 4113 | 4.93408921368824 | -0.64015233 | 1 | only_H3vsUI | DownUp | notDM | Loss | 6 |
| **X_38660905_38663092_58526** | X_38660905_38663092 | MID1IP1 | 58526 | 2.34635297445064 | -0.1658731 | 1 | only_DIvsD7 | UpDown | notDM | Gain | 4 |
| **X_38665421_38667553_58526** | X_38665421_38667553 | MID1IP1 | 58526 | 3.67559505638675 | -0.1658731 | 2 | only_DIvsD7 | UpDown | notDM | Gain | 5 |
| **X_45046117_45047163_79742** | X_45046117_45047163 | DIPK2B | 79742 | 4.11340850681818 | 0.78973628 | 2 | only_H3vsUI | UpDown | notDM | Loss | 3 |
| **X_46616147_46618142_84679** | X_46616147_46618142 | SLC9A7 | 84679 | 2.54282542695918 | 0.72712703 | 1 | only_H3vsUI | Down | notDM | Loss | 8 |
| **X_46618983_46620058_84679** | X_46618983_46620058 | SLC9A7 | 84679 | 2.69460519893357 | 0.41987633 | 4 | only_H3vsUI | Down | notDM | Loss | 4 |
| **X_46628122_46629193_84679** | X_46628122_46629193 | SLC9A7 | 84679 | 3.98380664197842 | 0.03198568 | 6 | only_H3vsUI | Down | notDM | Gain | 1 |
| **X_46629388_46630479_84679** | X_46629388_46630479 | SLC9A7 | 84679 | 4.03742649794062 | -0.20319339 | 3 | only_H3vsUI | Down | notDM | Gain | 1 |
| **X_46772616_46774369_84679** | X_46772616_46774369 | SLC9A7 | 84679 | 5.18788154294209 | -0.74465059 | 1 | only_H3vsUI | Down | notDM | Loss | 7 |
| **X_48896753_48897421_90060** | X_48896753_48897421 | CCDC120 | 90060 | 4.28110157568415 | 0.31261353 | 7 | only_DIvsD7 | UpDown | notDM | Loss | 5 |
| **X_48901513_48902267_90060** | X_48901513_48902267 | CCDC120 | 90060 | 4.1540281496032 | 0.22773404 | 5 | only_DIvsD7 | UpDown | notDM | Loss | 3 |
| **X_48936477_48937613_90060** | X_48936477_48937613 | CCDC120 | 90060 | 4.30007349526714 | 0.21844564 | 1 | only_DIvsD7 | UpDown | notDM | Loss | 3 |
| **X_54467215_54468851_90121** | X_54467215_54468851 | TSR2 | 90121 | 2.58319877396862 | -0.51991725 | 1 | only_DIvsD7 | DownUp | notDM | Loss | 8 |
| **X_55025436_55026341_5207** | X_55025436_55026341 | PFKFB1 | 5207 | 2.6730209071289 | 0 | 1 | only_H3vsUI | UpDown | notDM | Loss | 3 |
| **X_55026930_55028786_5207** | X_55026930_55028786 | PFKFB1 | 5207 | 3.29336255471145 | 0 | 1 | only_H3vsUI | UpDown | notDM | Loss | 3 |
| **X_66766532_66769692_367** | X_66766532_66769692 | AR | 367 | 3.42488163663107 | 0.43282154 | 2 | only_DIvsD7 | UpDown | notDM | Gain | 4 |
| **X_68048955_68050408_1947** | X_68048955_68050408 | EFNB1 | 1947 | 2.06818586174616 | 0.58119952 | 2 | only_H3vsUI | Down | DM | Loss | 10 |
| **X_69508830_69509686_54857** | X_69508830_69509686 | GDPD2 | 54857 | 5.12484969711265 | -0.96834449 | 1 | only_H3vsUI | DownUp | notDM | Loss | 3 |
| **X_70286415_70287746_29934** | X_70286415_70287746 | SNX12 | 29934 | 2.63144376901317 | -0.09420304 | 1 | only_DIvsD7 | DownUp | notDM | Gain | 2 |
| **X_70288645_70289385_29934** | X_70288645_70289385 | SNX12 | 29934 | 2.67577834167409 | -0.15802459 | 4 | only_DIvsD7 | DownUp | notDM | Loss | 7 |
| **X_70314769_70315557_29934** | X_70314769_70315557 | SNX12 | 29934 | 4.42484898168298 | -0.58209106 | 1 | only_DIvsD7 | DownUp | notDM | Loss | 7 |
| **X_70315998_70317052_29934** | X_70315998_70317052 | SNX12 | 29934 | 4.44321603416583 | -0.63301687 | 2 | only_DIvsD7 | DownUp | notDM | Loss | 7 |
| **X_70401035_70401769_29934** | X_70401035_70401769 | SNX12 | 29934 | 5.0525554379874 | -0.62490744 | 1 | only_DIvsD7 | DownUp | notDM | Loss | 7 |
| **X_80375327_80377169_6451** | X_80375327_80377169 | SH3BGRL | 6451 | 4.90557463055713 | 0.57104475 | 2 | only_DIvsD7 | DownUp | notDM | Loss | 6 |
| **X_80456430_80457404_6451** | X_80456430_80457404 | SH3BGRL | 6451 | 1.5910646070265 | 0.11401018 | 1 | only_DIvsD7 | DownUp | notDM | Loss | 3 |
| **X_80457651_80459759_6451** | X_80457651_80459759 | SH3BGRL | 6451 | 1.41497334797082 | -0.14838843 | 1 | only_DIvsD7 | DownUp | notDM | Loss | 8 |
